# Cell type-specific genetic regulation of expression in the granule cell layer of the human dentate gyrus

**DOI:** 10.1101/612200

**Authors:** AE Jaffe, DJ Hoeppner, T Saito, L Blanpain, J Ukaigwe, EE Burke, R Tao, K Tajinda, A Deep-Soboslay, JH Shin, JE Kleinman, DR Weinberger, M Matsumoto, TM Hyde

## Abstract

Laser capture microdissection followed by RNA-seq (LCM-seq) was used to profile the transcriptional landscape of the granule cell layer of the dentate gyrus (DG-GCL) in human hippocampus, and contrasted to homogenate tissue. We identified widespread cell type-specific aging and genetic effects in the DG-GCL that were either missing or directionally discordant in corresponding bulk hippocampus RNA-seq data from largely the same subjects. Of the ∼9 million eQTLs in the DG-GCL, 15% were not in bulk hippocampus, including 15 schizophrenia genome-wide association study (GWAS) risk variants. We then created custom transcriptome-wide association study (TWAS) genetic weights from the DG-GCL which identified many novel schizophrenia-associated genetic signals not found in TWAS from bulk hippocampus, including GRM3 and CACNA1C. These results highlight the biological resolution of cell type-specific expression profiling using targeted sampling strategies like LCM, and complement homogenate and single nuclei approaches in human brain.

## Introduction

Extensive effort has been spent over the past ten years to more fully characterize the human brain transcriptome within and across cell types and to better understand changes in RNA expression associated with brain development and aging, developmental or psychiatric brain disorders, and local genetic variation. Large consortia have primarily focused on the molecular profiling of RNA extracted from homogenate/bulk tissue from different brain regions across tens or hundreds of individuals ^1–6^, though single cell expression approaches are increasingly coming into focus.

We have previously identified extensive gene expression associations in human brain with schizophrenia and its genetic risk ^7^, development/aging, and local genetic variation in the dorsolateral prefrontal cortex (DLPFC) ^8^ and more recently, the hippocampal formation (HIPPO) ^9^. While eQTLs in these two brain regions were highly overlapping, consistent with previous work across many tissues in the body ^6^, there were distinct region-specific expression profiles associated with brain development and subsequently dysregulated in schizophrenia. We further identified stronger effects of schizophrenia diagnosis in the DLPFC compared to HIPPO, with an order of magnitude more genes differentially expressed. While these differences in signatures across brain regions likely related to the unique cell types underlying each region, particularly for changes across development ^9^, the specific cell types in which these signals act within and across brain regions is largely unknown.

To better understand expression within and across individual cell types, there has been a dramatic shift to RNA-seq that profiles tens or hundreds of thousands of cells or nuclei from few individuals. While these single cell (scRNA-seq) or nucleus (ncRNA-seq) approaches have cataloged dozens of transcriptionally disjoint cell classes in the human brain ^10,11^, the limited number of subjects and high cost have largely prevented the use of these approaches for association with genetic variation and with human traits. Furthermore, rarer cell populations have lower probabilities of ascertainment and subsequent characterization in these analyses. An alternative strategy for cell type-specific expression involves isolating specific cell populations using nuclear antibodies followed by flow cytometry or laser capture microdissection (LCM) of cell bodies for cells clearly defined by morphology. While previous research utilized LCM for expression profiling of precise anatomical regions in the human and primate brains ^12,13^ or layer-specific analysis of human cortex using microarray technologies ^14^, there have been few efforts to transcriptionally profile individual cell populations using this technique.

We therefore sought to evaluate LCM followed by RNA-seq (LCM-seq) as a tool for cell type-specific expression analysis in human brain tissue. Here we profiled the granule cell layer of the dentate gyrus subfield (DG-GCL) in the hippocampal formation, which plays a critical neuromodulatory role controlling information flow from the entorhinal cortex to CA3, CA1 and downstream targets including the prefrontal cortex ^15^. This layer primarily contains cell bodies of granule neurons, the primary excitatory neuronal cell type in the dentate gyrus. Single nuclei sequencing studies of human hippocampus estimated these cells constitute ∼5-10% of the hippocampus ^16^. The dentate gyrus plays a role in pattern separation and the downstream target CA3 plays a role in pattern completion in both rodent and human systems ^17–20^. Deficits in their activity previously have been associated with bipolar disorder and schizophrenia. However, many of the previous findings linking this important cell type to these debilitating disorders have been based on animal models ^21^, induced pluripotent stem cell (iPSC)-based approaches ^22^, or low-resolution functional imaging ^23–25^. We additionally selected this cell population to permit direct comparisons to bulk HIPPO RNA-seq data from a largely overlapping set of individuals ^9^.

We isolated and characterized human DG-GCL neurons to define their role in psychiatric disorders and their genetic risk, explicitly contrasting these data against bulk tissue to understand the benefits of cell type enrichment strategies. The LCM-based enrichment strategy is the only available high throughput approach that isolates the cell body and the nucleus, in contrast to droplet and sorting techniques which are nucleus restricted. This strategy of deeply sequencing target cell populations revealed new expression associations to aging and genetic polymorphisms, including those linked to schizophrenia. These data further provide a benchmark for comparing emerging technologies in cell type-specific RNA-seq and subsequent association with age, genotype, and diagnosis of psychiatric disorders.

## Results

### LCM-seq produces cell type-specific expression data

We performed laser capture microdissection (LCM) to extract the DG-GCL in postmortem human hippocampal tissue from 263 human subjects, including 75 donors with schizophrenia, 66 with bipolar disorder, 29 with major depression, and 93 neurotypical controls, all with genome-wide genotype data (Table 1, see Methods, Figure S1, Table S1). Furthermore, 112 subjects also had bulk hippocampal formation (HIPPO) RNA-seq data available (from 333 total age-matched samples). We first demonstrated that the LCM-seq procedure generates high quality RNA-seq data, as the LCM-seq data and bulk hippocampus RNA-seq data generated with the same RiboZero Gold library types had similar chrM (Figure S2A) and genome mapping rates (Figure S2B) albeit with slightly lower exonic mapping rates (Figure S2C) and correspondingly lower RNA integrity numbers (RINs) due to the laser methodology (Figure S2D).

**Table 1:** TWAS analysis in the DG-GCL integrated with the PGC2+CLOZUK schizophrenia GWAS. We further tabulated the subset of features that were heritable/had weights in the bulk hippocampus. GWAS In: best GWAS SNP from TWAS was an index SNP or proxy to GWAS loci. HIPPO P: p-value < 0.05, FDR: FDR < 0.5. *unique genes (can be overlaps across features)

| Type | Considered |  | InHippo | TWAS FDR < 5% |  |  |  |  | TWAS Bonf < 5% |  |  |  |  |
| --- | --- | --- | --- | --- | --- | --- | --- | --- | --- | --- | --- | --- | --- |
|  |  |  |  | Overall | GWAS |  | HIPPO |  | Overall | GWAS |  | HIPPO |  |
|  | #Features | #Genes | #Features |  | In | Out | P | FDR |  | In | Out | P | FDR |
| Gene | 3528 | 3528 | 1675 | 231 | 73 | 158 | 100 | 75 | 53 | 33 | 20 | 23 | 20 |
| Exon | 22806 | 5430 | 6852 | 1666 | 517 | 1149 | 543 | 315 | 212 | 183 | 29 | 63 | 61 |
| Junction | 9697 | 3790 | 2948 | 734 | 235 | 499 | 279 | 188 | 125 | 93 | 32 | 49 | 48 |
| Transcript | 5648 | 4073 | 2190 | 461 | 137 | 324 | 213 | 146 | 81 | 57 | 24 | 36 | 36 |
| Total | 41679 | 8546* | 13662 | 3092 | 962 | 2130 | 1135 | 724 | 471 | 366 | 105 | 171 | 165 |
|  |  | *Unique |  |  |  |  |  |  |  |  |  |  |  |

We then used the paired DG-GCL and hippocampus RNA-seq data on 129 subjects to confirm that the LCM-seq sample was enriched for neurons compared to bulk tissue (see Methods). We identified 1,899 genes differentially expressed in DG-GCL compared with bulk tissue at a conservative Bonferroni-adjusted p-value < 1% (p<4.65 × 10-7, Table S2). As expected, the top enriched genes (more highly expressed in DG-GCL) were *KCNK1* (3.9x up), *CAMK1* (4.5x up), and *GABRD* (6.3x up), which are relatively specific to neurons, and top depleted genes (more highly expressed in bulk tissue) were *MOBP* (53.7x down) and *MBP* (11.4x down) which are highly enriched in non-neuronal cells (Figure 1). Other significant and expected differentially expressed genes included enrichment of the dentate gyrus-associated gene *PROX1* (5.0x up) and relative depletion of the astrocyte gene *GFAP* (3.8x down). As a set, genes most enriched for preferential DG-GCL expression related to neuronal processes and localization (Table S3), demonstrating that the LCM-seq data are highly enriched for neuronal cells and produce high-quality data when compared to RNA-seq data derived from homogenate brain tissue.

**Figure 1.**
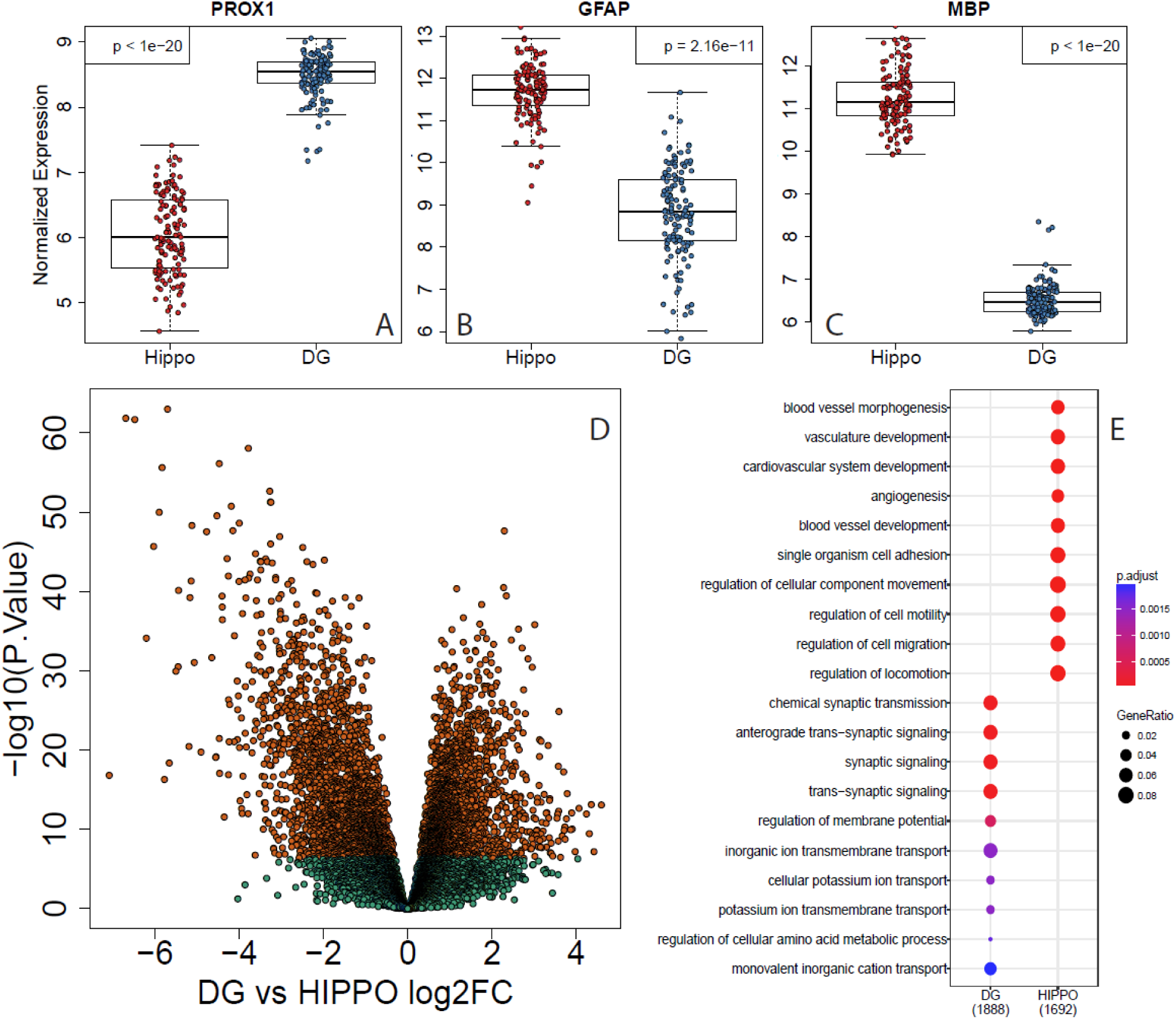
LCM-seq confirms expected strong cell-type-specific enrichment in DG. (A) DG-specific PROX1 (B) Astrocyte-specific GFAP (C) and Oligodendrocyte-specific MBP expression (log2 scale) (D) Volcano plot of differential expression between DG and HIPPO reveals many significant hits (E) Gene-set enrichment analysis demonstrates significant association of neuronal functions with the DG and non-neuronal functions with the homogenate hippocampus.

### Cell-type specific signatures of aging in the hippocampus

Because the hippocampal formation undergoes considerable cellular alterations with advancing age, we sought to identify genes with unique patterns of expression associated with postnatal aging, ranging from 16-84 years, in the DG-GCL against a background of aging associations in the bulk hippocampus. Here we expanded our comparisons to the full cohort of DG-GCL samples (N=263) and age- and library-matched bulk hippocampus samples (N=333, see Methods) on 21,460 expressed genes. Using linear modeling that adjusted for observed and latent confounders (see Methods), we first identified genes associated with age within each dataset (Figure 2A, Table S4). In the DG-GCL, we identified 1,709 genes whose expression was significantly associated with age (at FDR < 5%) with 833 genes increasing in expression and 876 genes decreasing in expression, with a median 3.7% change in expression per decade of life. In the hippocampus we identified 1,428 genes significantly associated with age (at FDR < 5%) with 733 genes increasing in expression and 695 genes decreasing in expression, with a median 3.0% change in expression per decade of life (IQR: 2.1%-4.4%). While the overlap between datasets was statistically significant (372 genes, odds ratio, OR: 4.93, p < 2.2e-16), there were over one thousand unique age-associated genes in each dataset (DG: 1,337 and Hippo: 1,056 genes). We then assessed the age-related directional changes of expression within the 2,765 genes significant in either the DG-GCL or hippocampus. While most genes showed directionally consistent age-related changes across dataset (N=2,353, 85.1%), including 369 of the 372 genes significant in both, there were 412 genes that showed opposite directionality, i.e. increased in expression across age in the DG-GCL and decreased in the hippocampus, or visa versa. These data indicate that ageing of dentate gyrus neurons has cell specific patterns not represented in bulk hippocampal formation tissue.

**Figure 2.**
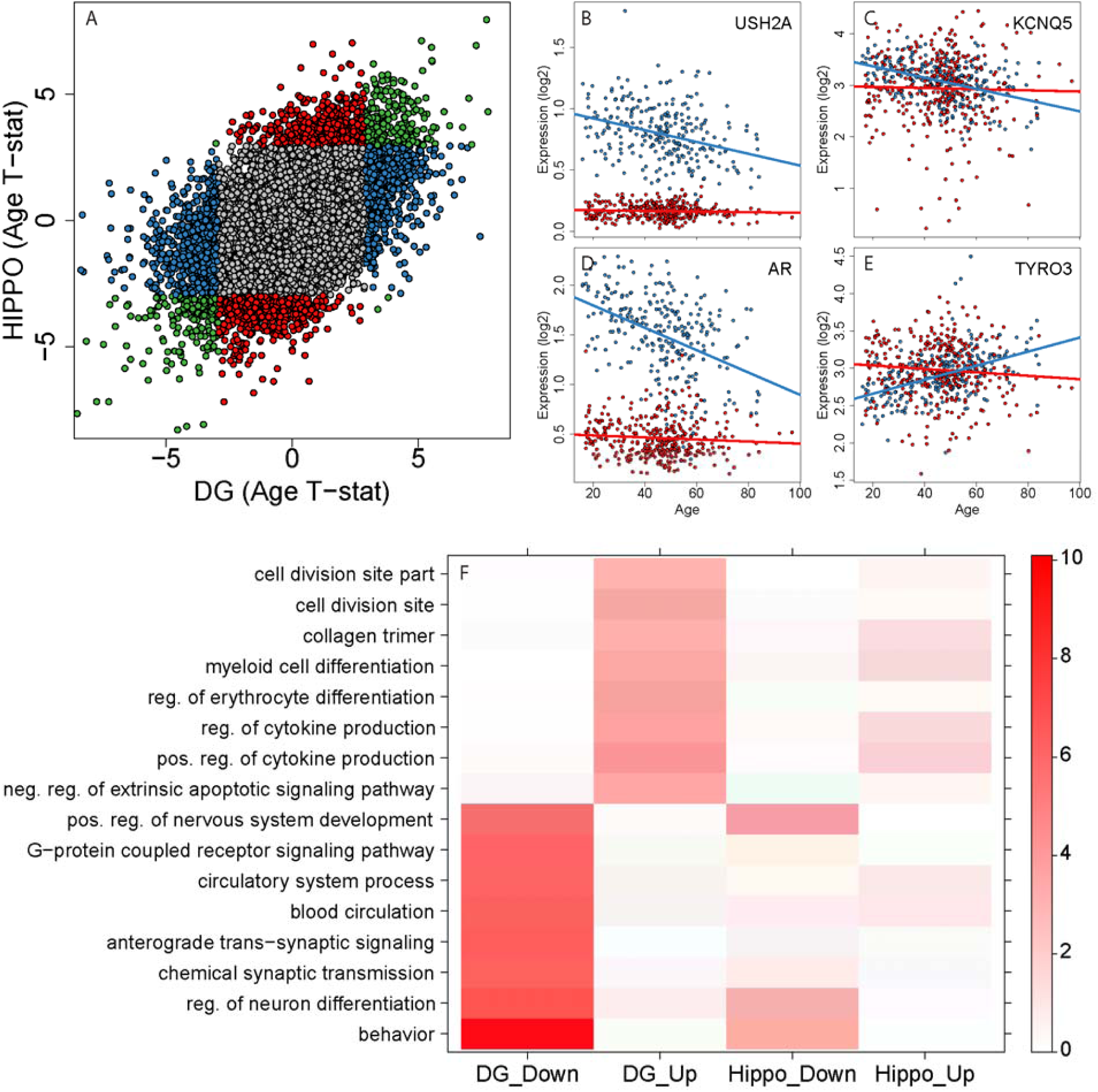
Cell-type specific changes in gene expression associated with aging. (A) DG vs Homogenate hippocampus T-statistic (Age) reveals both shared (gray) and differential expression (red = Hippo, Blue = DG) by age (B-E) Example genes with stable expression in Hippocampus (red) but age-dependent expression in DG (blue) (F) Heat map of association with age-dependent gene set enrichment terms. Temperature scale is p-value (−log10).

There are several individual gene expression patterns within these 2765 genes showing age-related changes in DG-DGL that are missed in sequencing bulk hippocampus. For example, USH2A shows early high expression in the DG-GCL that decreases across the lifespan, compared to low stable expression in the bulk hippocampus (Figure 2B). Another gene, KCNQ5, a voltage gated potassium channel subunit, shows similar expression levels in both datasets, but only changes expression levels across the lifespan in the DG-GCL (decreases, Figure 2C). The androgen receptor gene AR also shows higher expression in the DG-GCL that significantly decreases in expression across the lifespan (Figure 2D). A further pattern of age-related changes in the DG-GCL is shown by TYRO3, a tyrosine-protein kinase, which involves similar expression levels in both datasets but increasing expression across the lifespan only in the DG-GCL (Figure 2E). Additional examples are shown in Data S1.

We next performed gene set enrichment analyses to assess whether the age-related genes in the DG-GCL and hippocampus converged on the same biological functions (see Methods). We found more enrichment for predefined gene sets in the DG-GCL (N=245 sets) than in bulk hippocampus (N=130 sets, Figure 2B) with largely shared pathways for genes increasing in expression across age but no overlapping gene sets for those genes decreasing in expression across age. The most divergent gene sets were enriched for many processes related to the structure and function of neurons (Figure 2F, Table S5), including decreasing expression of genes in the DG-GCL - but not bulk tissue-that associated with “behavior” (GO:0007610, 56/630 genes; DG-GCL q=8.52e-7, HIPPO q=0.186), “anterograde trans-synaptic signaling” (GO:0098916, 55/630 genes; DG-GCL q=4.03e-4, HIPPO q=0.997) “G-protein coupled receptor signaling pathway” (GO:0007186, 53/630 genes; DG-GCL q=4.56e-04, HIPPO q=0.997) and “regulation of neuron projection development” (GO:0010975, 42/630 genes; DG-GCL q=0.0042, HIPPO q=0.26). Conversely, many of the gene sets associated with increasing expression over age, regardless of dataset (and cell specificity), related to immune cells and their processes, at least as represented in peripheral inflammatory cells (i.e. neutrophils and leukocytes), with some specific classes of cell types showing preferential enrichment in DG-GCL (cytokines, myeloid cells, erythrocytes) and bulk hippocampus (T cells).

Lastly, we analyzed the effects of age within individual subjects across brain regions using linear mixed effects modeling with an interaction between age and region/dataset. We found 406 genes with significantly different expression trajectories with aging contrasting the DG-GCL with hippocampus. The majority of these genes (N=247, 60.8%) were differentially expressed by age in either the DG-GCL or hippocampus (as in Figure 2B-E, Table S4). The remaining significant genes in this interaction model were those that did not show significant changes with aging in either dataset, but had significantly different age associations when contrasting DG-GCL with hippocampus in single statistical model (Data S2). These results highlight the value of generating cell type-specific expression data to better characterize aging-related expression phenotypes that are missed in homogenate tissue.

### Cell type specific eQTLs

We next assessed the potential for cell type-specific genetic regulation of expression using expression quantitative trait loci (eQTL) analysis. We first calculated local (cis) eQTLs in the DG-GCL dataset (N=263) at different expression features: genes, exons, junctions and transcripts (see Methods). We identified widespread cell type-specific genetic regulation of transcription. There were 8,988,986 significant SNP-feature pairs in the DG-GCL, which were driven by exon-level signals (4,734,526 pairs, Data S1). The majority of expressed genes showed association with a neighboring SNP at at least one feature type at genome-wide FDR

<1% significance (N=17,683, Figure 3A). The exon, junction, and transcript eQTL features were annotated back to 71,346 specific Gencode transcripts, and half had support from at least two feature types (Figure 3B). Lastly, we identified 5,853 exon-exon splice junctions associated with nearby genetic variation that were only partially annotated (1,622 exon skipping and 4,231 alternative exonic boundary events) to 3,086 unique genes. Of these, 162 showed association only to unannotated sequence, and not to any other annotated feature in the gene, potentially suggesting novel DG-GCL specific transcripts for a small number of genes.

**Figure 3.**
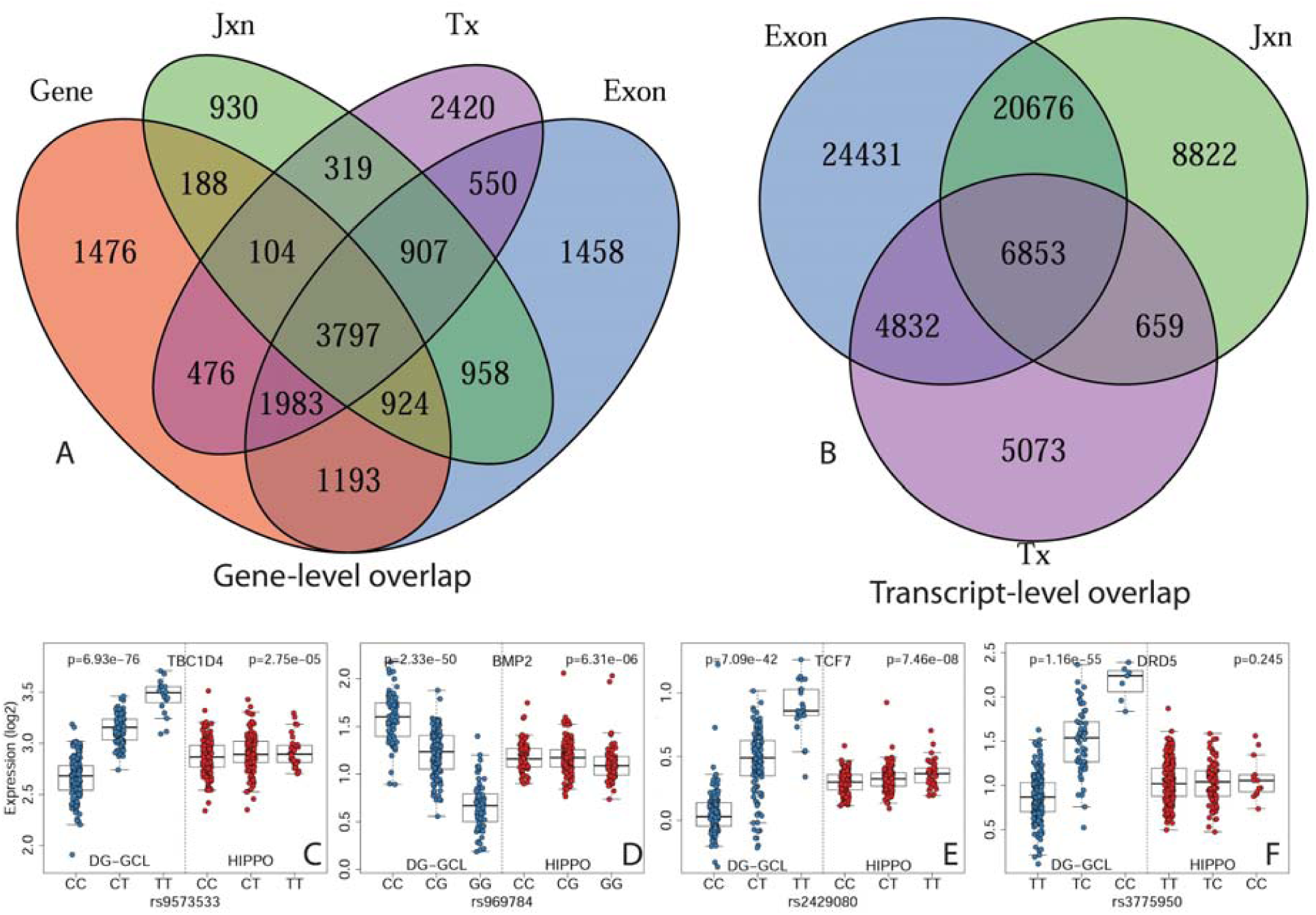
DG-specific eQTLs. (A) Venn diagram showing cis-eQTL feature overlap (B) GENCODE-mapped feature overlap in DG (C-F) Example SNPS with strong cis-eQTL gene signals in DG, but not homogenate hippocampus.

We then assessed cell type specificity of these eQTLs - and the extent that would have been missed in homogenate sequencing of hippocampus tissue. We calculated corresponding eQTL statistics for all ∼9M eQTL SNP-feature pairs in the 333 homogenate hippocampus samples, as well as the region-by-genotype interaction effects in the joint set of 596 samples. Here 3,420,833 SNP-feature pairs significant in DG-GCL (38.5%) across 13,767 genes showed different eQTL effects in the bulk hippocampus (interaction model, at FDR < 5%), including 1,501,540 pairs across 12,531 genes that were not even marginally significant in bulk tissue (hippocampus model, at p > 0.05). There were additionally 13,732 SNP-feature pairs that were genome-wide significant in both DG-GCL and hippocampus (each at FDR < 1%) in 159 genes with opposite eQTL directionality (Figure 3C-4F, Data S3), suggesting different mechanisms of genetic regulation across different cell types. Log fold changes per allele copy (i.e. the effect sizes) were also larger in DG-GCL than hippocampus when constraining to genome-wide eQTL significance in both datasets (4,781,015 at FDR < 1%) on average (85.6% pairs), even among the subset of these eQTLs more statistically significant in hippocampus (70.7% pairs). These results underscore that genetic regulation of gene expression can differ remarkably across specific cell populations and that bulk tissue eQTL analyses do not capture the complexity of genetic regulation in brain ^6^.

### Cell type specific eQTLs for schizophrenia

Hippocampus is one of the brain regions prominently implicated in the pathogenesis of schizophrenia ^26,27^, so we then performed focused eQTL analyses around schizophrenia risk variants from the latest genome-wide association study (GWAS) ^28^. We profiled 6,277 proxy SNPs (LD R^2^ > 0.8) from 156 loci with lead “index” SNPs present and common in our sample (from 179 loci/index SNPs identified in the GWAS, 87.5%, see Methods), and performed cis eQTL analyses within the DG-GGL (N=263), HIPPO (N=333), and joint interaction (N=596) datasets. We identified 100/156 loci (64.1%) with significant eQTLs in either the DG-GCL or HIPPO (FDR < 1%), with 60 loci with significant eQTLs in both datasets (Table S6). Combining eQTL data from the DLPFC ^9^ identified an additional 11 loci with FDR < 1% significant signal (N=111/156), and reducing the significance (at FDR < 5%, controlling only within these GWAS-centric analyses) found eQTL evidence for 136/156 (87.2%) of tested loci across all three datasets. Combining data across multiple brain regions and cell types even in a relatively limited number of brain specimens has therefore identified a transcript feature associated with almost every current schizophrenia risk locus.

Importantly, there were many loci with significant eQTLs only in DG-GCL without corresponding marginal significance in bulk hippocampus, including seven loci where the index SNPs themselves associated with expression levels of gene features: *PSD3* (Figure 4A, DGp=1.51e-6), *MARS* (Figure 4B, DGp=1.63e-6), *NLGN4X* (Figure 4C, DGp=1.79e-5), GRM3 (Figure 4D. DGp=2.65e-5), *SEMA6D* (Figure 4E, DGp=5.85e-5), *MMP16* (Figure 4F, DGp=1.47e-4), and *THEMIS* (Figure 4G, DGp=2.75e-4). Integrating nearby proxies further identified association to SATB2 (Figure 4H, DGp=2.81e-6), CACNA1C (Figure 4I, DGp=4.27e-6), KCTD18 (Figure 4J, DGp=6.34e-6), PRKD1 (Figure 4K, DGp=8.07e-5), HDAC2-AS2 (Figure 4L, DGp=1.16e-4), IGSF9B (Figure 4M, DGp=1.18e-4), TMPRSS5 (Figure 4N, DGp=1.38e-4) and TNKS (Figure 4O, DGp=1.53e-4). Two eQTL associations of particular interest involved genetic regulation of *GRM3* and *CACNA1C*, as ion channels (e.g. *CACNA1C*) and G protein coupled receptors (e.g. GRM3) are classic therapeutic drug targets and eQTLs for these genes in bulk tissue have been elusive. We explored the tissue specificity of these eQTLs using the BrainSeq DLPFC ^8,9^, CommonMind Consortium DLPFC ^3^ and GTEx ^5^ bulk tissue projects. The GWAS index SNP that associated with GRM3 expression (rs12704290) was not significantly associated with any expression features in the BrainSeq or CommonMind Consortium DLPFC, or GTEx datasets across any brain region or body tissue. The top proxy SNP that associated with *CACNA1C* expression levels (rs7297582) did show association in GTEx, exclusively in the cerebellum (p=6.6e-7) with no significant association in any other tissue or in BrainSeq.

**Figure 4.**
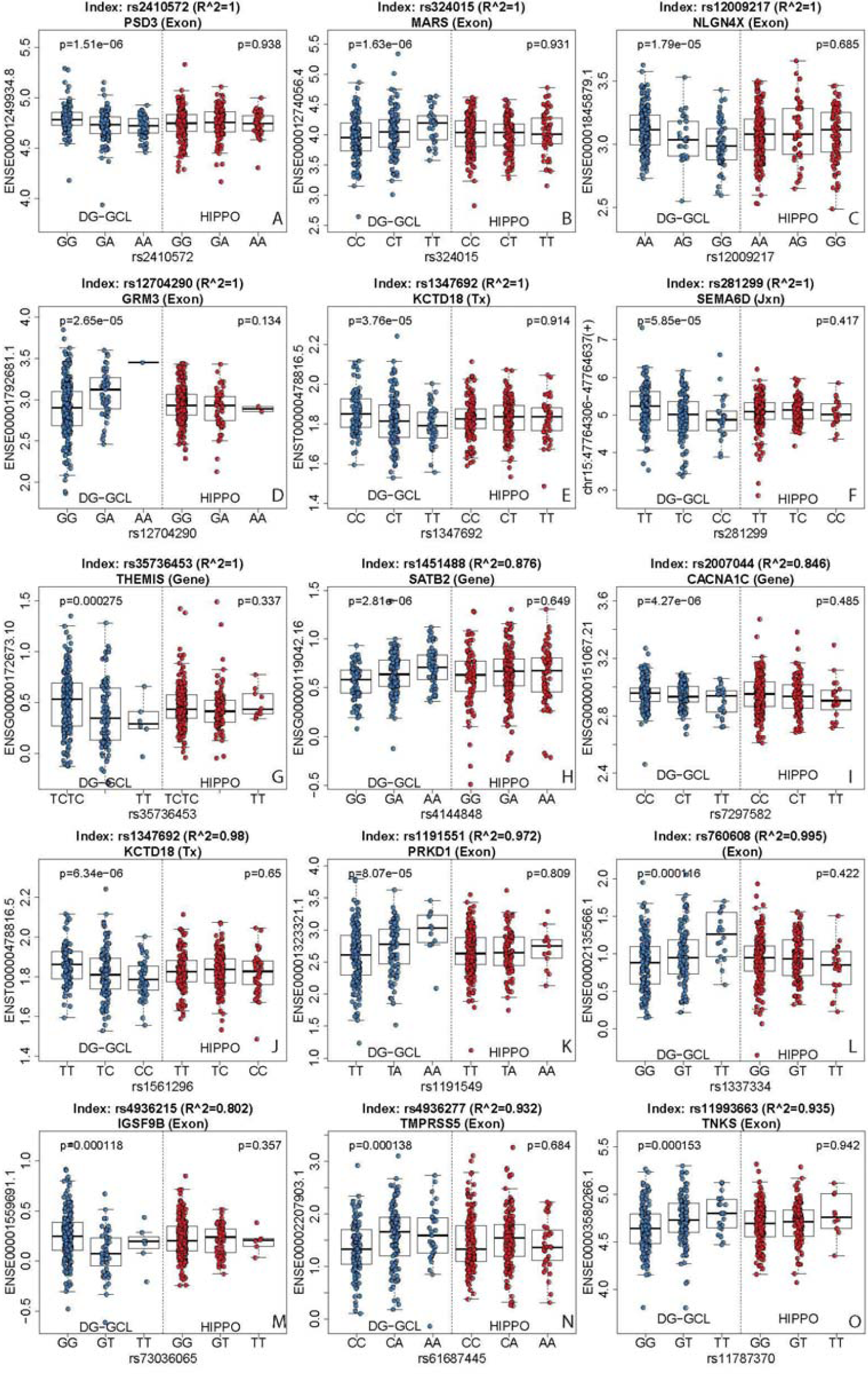
Schizophrenia risk eQTLs reveal feature associations in DG not seen in homogenate hippocampus. X=Schizophrenia risk SNP genotype. Y=log2(FPKM) from DG or Homogenate hippocampus. (A-G) Example eQTLs with PGC2 index SNPs (R∧2=1). (H-O) Example eQTLs with less stringent LD R∧2 >0.7.

### Cell-type specific TWAS

In light of the unique DG-GCL eQTLs, we performed a transcriptome-wide association study (TWAS) ^29^ after constructing SNP weights across the four feature summarizations (gene, exon, junction and transcript) using the DG-GCL dataset and applying these weights to the summary statistics from the entire collection of schizophrenia GWAS summary statistics from above (see Methods). We identified 3092 features (231 genes, 1666 exons, 734 junctions and 461 transcripts, Table S7) significantly associated with schizophrenia risk with TWAS FDR < 5% that were annotated to 1069 unique genes, of which 471 features (53 genes, 212 exons, 125 junctions and 81 transcripts) in 170 genes were significant following more conservative Bonferroni adjustment (at < 5%). As the TWAS approach combines GWAS and eQTL information, it’s possible for GWAS signal that does not reach GWAS genome-wide significance (p < 5e-8) to still achieve TWAS transcriptome-wide significance. We therefore annotated the strongest GWAS variant for each significant TWAS feature back to the clumped GWAS risk loci, and found that 77.7% of the TWAS Bonferroni-significant features (N=366) mapped back to the published GWAS risk loci. We confirmed these TWAS-significant features included GRM3 (TWAS gene p=7.28e-06) and *CACNA1C* (TWAS gene p=1.65e-10), which were both associated with decreased expression with increased schizophrenia genetic risk. While a small fraction of Bonferroni-significant TWAS features were outside of GWAS risk loci, a much larger fraction of FDR-significant TWAS features identified potentially novel genes and corresponding GWAS loci implicated in schizophrenia (N=2130 features in 799 genes, 68.9%).

We compared these TWAS weights and corresponding schizophrenia associations to those from the bulk hippocampus. Only half of the genes, and a third of the features overall, in the DG dataset had heritable expression in hippocampus and were considered for integration with GWAS data. Among TWAS-significant features in the DG-GCL, there were fewer than half that showed even marginally significant association in the hippocampus, largely in line with the schizophrenia risk variant-focused eQTL analysis above. Profiling the DG-GCL transcriptome, therefore, identified unique schizophrenia-associated signal that was missed in homogenate tissue, and these genetic associations to expression highlight that the potential pathogenic role of these genes in schizophrenia is mediated in a cell-specific context.

### Unique signatures of illness in the DG-GCL

We lastly explored unique illness state-associated gene expression differences in the DG-GCL compared to the bulk hippocampus. Our previous work suggested that the hippocampus had fewer genes differentially expressed in samples from patients with schizophrenia compared to genes differentially expressed between patients and controls in the DLPFC (both using ribosomal-depletion sequencing methods) ^9^. In the DG-GCL, we similarly found relatively small numbers of genes associated with diagnosis, particularly compared to aging and eQTL effects. We identified 26 genes in SCZD, 20 genes in BPD, and 7 genes in MDD (all at FDR< 10%, Table S8). While only a single gene RPL12P20 was significant in more than one disorder (SCZD and BPD), we found a significant correlation between these two disorders across the entire transcriptome (□, Pearson correlation = 0.53, Figure S3), with less correlation between both SCZD and MDD (□ = 0.39) and BPD and MDD (□= 0.35), as previously reported ^30^. We compared the effects of SCZD in the DG-GCL to bulk hippocampus, which was the only diagnostic group shared between the two datasets. There were only two significant genes shared across datasets: GMIP and ZNF766 (at FDR < 10%) with larger log-fold changes in the DG-GCL (−0.45 versus −0.17 and 0.14 versus 0.08, respectively). There was also less global correlation for SCZD effects between datasets (□= 0.197) than shared diagnosis effects within the DG-GCL (Figure S4). These results highlight the cell type-specificity of schizophrenia-associated differential expression analysis.

Antipsychotic and antidepressant drugs have been associated with changes in DG gene expression and in putative neurogenesis ^31^. We, therefore, compared two different treatment effects on differential expression specificity in the DG-GCL. First, we tested for differences in expression between patients with MDD with (N=18) and without (N=8) selective serotonin reuptake inhibitors (SSRIs) compared to unaffected controls negative for SSRIs (N=93, with 63 explicitly testing negative for SSRIs), and found 31 genes significantly different between patients with MDD on SSRIs and unaffected controls. These genes were largely non-overlapping with the above 7 genes associated with MDD overall, ignoring SSRI status (only two genes, RAD18 and DCAF16, were shared). None of these genes have been associated with adult neurogenesis, an effect of SSRIs that has been prominently hypothesized ^32^. We performed an analogous analysis within schizophrenia patients stratifying by antipsychotics status at time of death (49 with antipsychotics and 25 without) compared to 94 controls (of which 55 explicitly tested negative for antipsychotics). We found 110 genes differentially expressed between patients on antipsychotics and unaffected controls (at FDR<10%), compared to 0 genes different between patients not on antipsychotics and unaffected controls. Here there was more overlap among the 27 genes with overall effects above (23/27), and only two genes (SRR and GRN) were previously associated with adult neurogenesis. Both analyses are in line with previous observations that gene expression differences between patients and controls largely reflect treatment effects ^8^.

## Discussion

We performed LCM followed by RNA-seq to generate the transcriptional landscape of the granule cell layer of the dentate gyrus in human hippocampus. This approach identified widespread cell type-specific aging and genetic effects in the DG-GCL that were either missing or directionally discordant in corresponding bulk hippocampus RNA-seq data from largely the same subjects. We identified 1337 genes with expression that only associated with age across the lifespan in the DG-GCL and these genes were enriched for diverse neuronal processes. We further identified ∼9 million SNP-feature eQTL pairs in the DG-GCL, of which 15% were not even marginally significant (p> 0.05) in bulk hippocampus. Using these eQTL maps, we identified novel schizophrenia-associated genes and their features that were completely missed in bulk brain tissue, including associations to GRM3 and CACNA1C. We lastly found a small number of genes differentially expressed in the DG-GCL in patients with schizophrenia, bipolar disorder or major depression compared to neurotypical individuals that were largely missed in bulk tissue. These results together highlight the importance, and biological resolution, of exploring cell type-specific gene expression levels using targeted sampling strategies like LCM.

Cellular neuroscience has rapidly evolved via the development of innovative tools that enable the characterization of single cell transcriptional profiles after dissociation or sorting of fresh tissue ^33–35^ or single nuclei from frozen tissue ^10,11,16,35,36^. However, while these approaches characterize the expression levels of many cell classes in the brain, there are typically few individuals profiled and few genes characterized per cell, making disease-or trait-related associations within individual cell types difficult. The other extreme in large-scale neurogenomics has involved profiling homogenate tissue, mixing and potentially diluting cell type-specific associations across individuals ^2,3,37^. Although some disease-relevant targets can be readily assigned to a single cell class from homogenate tissue (e.g. C4) ^38^, the cellular specificity of most others require further interrogation ^3,39,40^.

Our data illustrate a balance between these two extremes - deeply profiling gene expression from a relatively homogeneous cell population, that permits cell type-specific inference of age, genotype, and neuropsychiatric illness. As proof of principle, we used the granule cell layer of the well-characterized hippocampal dentate gyrus as a practical intermediate between homogenate tissue and sorted nuclei. The anatomically-distinct granule neuron layer enables direct comparison between deep sequencing of homogenate tissue and a highly-enriched single cell type. Using the unique tissue resource of contralateral cerebral hemispheres, we directly compared transcript diversity between homogenate and laser-captured granule neuron cell bodies.

Our most striking finding involved the extensive cell type-specific genetic regulation of expression, with highly significant eQTLs in the DG-GCL without any corresponding signal in homogenate hippocampal tissue. While larger consortia like GTEx ^6^, psychENCODE ^41^ and our BrainSeq ^8,9^ have saturated the landscape of homogenate tissue eQTLs in human brain, here we demonstrate much of genetic regulation within individual cell populations is clearly incomplete. Only one other study - to the best of our knowledge - has profiled a selective cell population from human brain tissue to identify cell type eQTLs, namely dopamine neurons from the substantia nigra in 84 subjects, and only a small number of eQTLs (only 3461 SNPs to 151 expressed sequences) were reported ^42^. This report further did not assess the cellular or regional specificity of these associations, and many of these reported SNPs show association in both the DG-GCL and bulk hippocampus. For example, the top disease-related SNP (rs17649553) showed strong association to 11 nearby genes in our DG-GCL (including 8 genes at p < 1e-20) of which 9 were also FDR-significant in bulk hippocampus (including 5 genes at p < 1e-20). Other studies have profiled human brain tissue with LCM^14,43–45^ but all other have focused on comparisons of illness state, which showed the least amount of signal in the DG-GCL compared to age and genotype.

In addition to the extensive genome-wide cell type-specific eQTL associations, we found many novel schizophrenia-associated eQTLs that were not identified in homogenate hippocampus. Two of the most prominent and longest-implicated genes with unique DG-GCL eQTL associations were *GRM3* ^46^ and *CACNA1C* ^47^, which finally provides molecular evidence implicating these genes, rather than merely variants proximal to these genes, with schizophrenia genetic risk. GRM3 and CACNA1c have been especially alluring as schizophrenia gene targets given the druggability of this G-protein coupled receptor and ion channel, respectively. These cell type specific associations are heuristic in terms of generating models for therapeutic discovery, as cell and animal models targeting *GRM3* and CACNA1C might focus on dentate gyrus granule cell experimental models. In this regard, it should be noted that the risk associated alleles for both of these genes was associated with relatively reduced expression. Prior work had been inconsistent in suggesting the directionality of risk association with expression of CACNA1C in neocortical samples ^48,49^, and the widely held assumption that CACNA1C antagonism is a therapeutic translation of the risk association may have to be re-examined based on the current data.

Overall, we demonstrate that the LCM-based enrichment strategy detects signals unique to the granule cell layer that were completely missed in homogenate tissue and generates TWAS evidence of novel schizophrenia risk associated loci that also were dependent on gene expression data in DG-GCL in contrast to bulk hippocampal tissue. This strategy of deeply sequencing target cell populations provides a powerful balance between unbiased single cell and homogenate tissue sequencing that can provide cell type-specific associations to common molecular and clinical traits.

## Methods

### Human Postmortem Brain Tissue Collection

Postmortem human brain tissues were collected at the Clinical Brain Disorders Branch (CBDB) at National Institute of Mental Health (NIMH) through the Northern Virginia and District of Columbia Medical Examiners’ Office, according to NIH Institutional Review Board guidelines (Protocol #90-M-0142) and the Lieber Institute for Brain Development (LIBD) according to with a protocol approved by the Institutional Review Board of the State of Maryland Department of Health and Mental Hygiene (#12-24) and the Western Institutional Review Board (#20111080). Details of the donation process, specimens handling, clinical characterization, neuropathological screening, and toxicological analyses are described previously ^50,51^. Each subject was diagnosed retrospectively by two board-certified psychiatrists, according to the criteria in the DSM-IV. Brain specimens from the CBDB were transferred from the NIMH to the LIBD under a Material Transfer Agreement.

Briefly, all patients met DSM-IV criteria for a lifetime axis I diagnosis of schizophrenia or schizoaffective disorder (N=75), bipolar disorder (N=66), or major depression (N=29), and neurotypical control subjects (N=93) were defined as those individuals with no history of significant psychological problems or psychological care, psychiatric admissions, or drug detoxification and with no known history of psychiatric symptoms or substance abuse, as determined by both telephone screening and medical examiner documentation, as well as negative toxicology results. Additional selection criteria include high-integrity RNA in each sample from prior studies of other brain areas, age match to control samples, and a broad age range. Due to the relatively large sample set, we further attempted to use gender and ethnic diversity as inclusion rather than exclusion criteria. A majority of these cases were Caucasians (N=169). A total of 263 hippocampal samples (Table 1) were used for the laser capture microdissection (LCM) described below.

### Brain tissue processing

After removal from the calverium, brains were wrapped in plastic and cooled on wet ice. A detailed macroscopic inspection was performed of the brain, meninges, attached blood vessels, and when possible, the pituitary and pineal glands. Brains were removed from the skull, wrapped in plastic, and transported on wet ice. The brains then were hemisected, cut into 1.5 cm coronal slabs, flash-rapidly frozen in a pre-chilled dry-ice/isopentane slurry bath (−40°C), and stored at −80°C. The time from when the tissue was stored at −80°C until the RNA was extracted was considered the freezer time (mean ± SD: 43.8 ± 2.8 months). A block of lateral superior cerebellar cortex hemisphere was cut transversely to the folia. pH was measured by inserting a probe into the right parietal neocortex and again into the right cerebellar hemisphere. For the purpose of this study, the slab containing the hippocampus at the level of the midbody was identified by visual inspection. An approximately 2×2×1 cm block was then taken from the medial temporal lobe, encompassing the hippocampal formation, entorhinal cortex, and adjacent white matter. This block was kept frozen at all times, and then was mounted for sectioning for laser capture microdissection.

### Dentate Gyrus Laser Capture Microdissection (LCM)

The dentate gyrus granule cell layer was isolated from the neighboring polymorphic and molecular layers using laser capture microdissection (LCM) (Figure S1). The mid-body of the dentate gyrus (DG) was exposed by gross block dissection followed by 30 micron cryosection onto 20 Glass slides coated with pre-charged PEN membrane (Zeiss Microscopy, Jena, Germany). To enhance signal in the densely-packed granule cell layer, sections were briefly stained with the nucleic acid intercalating agent Acridine Orange (Molecular Probes A3568) for 1 minute in EtOH prior to LCM. Emitted green light from blue excitation was used to distinguish the granule cell layer from the adjacent polymorphic layer and sub-granule zone. The limits of the granule cell layer were defined manually for each section and entered into PALM Robo software for LCM using laser pressure capture (Zeiss, Jena, Germany). LCM caps (MMI, Switzerland) were stored on dry ice immediately after harvesting fragments until RNA lysis and RNA extraction with RNeasy Micro kit (Qiagen, Venlo, the Netherlands). Over 100ng total RNA was isolated from the pooled fragments of each donor. This relatively high output quantity from LCM enables more accurate steady-state mRNA quantitation and avoids the amplification bias and computational confounds associated with RNA pre-amplification ^52^. The quantity and integrity of RNAs was determined using nanodrop (Wilmington DE) and BioAnalyzer (Agilent, Santa Clara CA).

### RNA sequencing data generation and processing

RNA was made into RNA-seq libraries using the Illumina RiboZero Gold library preparation kit and sequenced on an Illumina HiSeq 2000 sequencer. Raw sequencing reads were quality checked with FastQC ^53^, and where needed leading bases were trimmed from the reads using Trimmomatic ^54^ as appropriate. Quality checked reads were mapped to the hg38/GRCh38 human reference genome with splice-aware aligner HISAT2 version 2.0.4 ^55^, with an average overall alignment rate of 92.5% (SD=3.8%). Feature-level quantification based on GENCODE release 25 (GRCh38.p7) annotation was run on aligned reads using featureCounts (subread version 1.5.0-p3) ^56^ with a mean 27.2% (SD=4.0%) of mapped reads assigned to genes. Exon-exon junction counts were extracted from the BAM files using regtools ^57^ v. 0.1.0 and the bed_to_juncs program from TopHat2 ^58^ to retain the number of supporting reads (in addition to returning the coordinates of the spliced sequence, rather than the maximum fragment range) as described in ^8^. Annotated transcripts were quantified with Salmon version 0.7.2 ^59^. For an additional QC check of sample labeling, variant calling on 740 common missense SNVs was performed on each sample using bcftools version 1.2 and verified against the genotype data described below. We generated strand-specific base-pair coverage BigWig files for each sample using bam2wig.py version 2.6.4 from RSeQC ^60^ and wigToBigWig version 4 from UCSC tools ^61^ to calculate quality surrogate variables (qSVs) for hippocampus-susceptible degradation regions ^9^.

We retained 21460 expressed genes with RPKM > 0.5 when using the number of reads assigned to genes as the denominator (not the number mapped to the genome). For secondary analyses, we retained 358280 expressed exons with RPKM > 0.5 (using assigned genes as the denominator), 241957 exon-exon splice junctions with reads per 10 million spliced (RP10M) > and not completely novel, and 95,027 transcripts with TPM > 0.05.

### Integration with hippocampus RNA-seq data

We further integrated existing RNA-seq data from the bulk hippocampus on the same 21460 genes, which has been described previously ^9^, on 333 samples over the age of 17 (200 controls and 133 patients with schizophrenia) that were sequenced with the RiboZero HMR kit, resulting in a joint dataset of 596 samples across 484 unique donors. The bulk hippocampus dissections here consisted of the entire hippocampal formation including all of the CA subfields being dissected from the medial temporal lobe under visual guidance using a handheld dental drill.

The anterior half of the hippocampal formation was included in this dissection, beginning just posterior to the amygdala. Sensitivity analyses related to library preparation kits were performed with an additional 17 hippocampus RNA-seq samples that were sequenced with the same RiboZero HMR kit which were excluded from subsequent analysis (Figure S2).

### DNA genotyping and imputation

Cerebellar DNA was extracted and genotyped for all 484 unique donors across the 596 samples as described previously ^8,9^ which briefly involved phasing and imputation to the 1000 Genomes Phase 3 reference panel with SHAPEIT2 ^62^ and IMPUTE2 ^63^. We retained 6,521,503 SNPs that were well-imputed (missingness < 10%) and common (MAF > 5%, HWE > 1×10-6) across the 263 DG-GCL samples for eQTL analysis. The same sets of SNPs were extracted from the 333 bulk hippocampus samples. We used LD-independent SNPs to calculate the top 5 multidimensional scaling (MDS) components as a measure of quantitative ancestry.

### Cell type-specific expression analysis

We performed differential expression analysis among the 112 donors with both hippocampus and DG-GCL samples (N=224) using linear mixed effects (LME) modeling using the limma voom approach ^64^. We adjusted for the mitochondrial chromosome (chrM) mapping rate, ribosomal RNA (rRNA) assignment rate, overall genome mapping rate, and exonic assignment rates to account for differences in library preparations and other technical factors, and used subject as a random intercept in the LME modeling with the ‘duplicateCorrelation()’ argument.

### Differential expression across age and diagnosis

We next modeled age and diagnosis effects, adjusting for exonic assignment rate, sex, chrM mapping rate, five MDS ancestry components, and quality surrogate variables (qSVs) using different sample subsets of the combined 596 RNA-seq samples. We first calculated qSVs ^65^ from the 488 significant degradation-susceptible regions of the hippocampus by extracting library size-normalized read coverage across all 596 samples, and performing PCA to retain the top 9 PCs as the qSVs. We fixed the qSVs for the different models below for comparability, and ran four differential expression (DE) analyses using limma voom ^64^

1. DG-GCL only (N=263) - we assessed the significance of the age main effects and diagnosis main effects separately using the ‘topTable()’ function
2. Hippocampus only (N=333) - we assessed the significance of the age main effects and diagnosis main effects separately using the ‘topTable()’ function
3. Age interaction model (N=596) - we further adjusted for brain region and the interaction between brain region and age, and treated donor as a random intercept using LME modeling. We assessed the significance of the interaction term.
4. Schizophrenia interaction model (N=596) - we further adjusted for brain region and the interaction between brain region and schizophrenia diagnosis, and treated donor as a random intercept using LME modeling

We performed two sensitivity analyses for drug effects within the DG-GCL - one for SSRIs among MDDs and another for antipsychotics among patients with schizophrenia. Here we recoded the diagnosis effect to be Control < Case (No Treatment) < Case (Treatment) and extracted the main effects of Case (Treatment) versus Control and Case (No Treatment) versus Control, adjusting for the same model as #1 above. Controls without toxicology data were still included as detailed chart review suggested these individuals were neurotypical, but patients without definitive toxicology data were excluded.

### Gene set testing

We used the clusterProfiler Bioconductor package ^66^ to perform gene set enrichment analyses using sets of significant genes as the inputs, along with the Entrez IDs corresponding to the 21640 expressed genes as the gene universe. Gene set results were corrected using Benjamini-Hochberg for multiple testing. We used the MANGO database ^32^ to test for enrichment of adult neurogenesis genes, which used mouse Entrez gene IDs as identifiers. We mapped these IDs to human ENSEMBL orthologs, resulting in 257/259 matches in Gencode V25, of which 172/259 were expressed in the DG and HIPPO datasets, which formed the gene set for enrichment analyses.

### Expression quantitative trait loci (eQTL) mapping

We performed eQTL mapping with the 263 DG-GCL samples, using the MatrixEQTL package ^67^ and adjusting for diagnosis, sex, the 5 MDS components, and then overall expression principal components estimated by the ‘num.sv’ function in the sva package ^68^ (gene: 22, exon: 28, junction: 46, transcript: 35) using 500Mb as the window size for cis-eQTL detection. We retained those eQTLs with empirical FDR < 0.01 significance for testing in other datasets (N=8,988,986 SNP-feature pairs), and tested their effects in the hippocampus dataset (N=333) using analogous statistical models with dataset-specific PCs (gene: 24, exon: 28, junction: 32, transcript: 26). We also computed the interaction between genotype and dataset using the full set of 596 samples for all ∼9M SNP-feature pairs using LinearCROSS argument in MatrixEQTL, again adjusting for an analogous statistical model with dataset-specific PCs (the top 25 feature-specific PCs). We merged DG-GCL-significant eQTLs across the three datasets to create a single database. We also performed more focused eQTL analyses within the latest schizophrenia risk variants ^28^ and their highly correlated proxies identified using rAggr ^69^ (rather than all common SNPs, R∧2> 0.8) using the same statistical models as above. Corrected p-values were re-calculated in these analyses as they were relative to those SNPs and features proximal to the GWAS risk loci.

### Transcriptome-wide association study (TWAS) analysis

We constructed TWAS weights following this guide: https://github.com/LieberInstitute/brainseq_phase2/tree/master/twas which was adapted from the README of the published TWAS approach ^29,70^, adapted for hg38 coordinates (rather than hg19). First, three sets of SNPs were homogenized in terms of coordinates, names, and reference alleles: 1) the LD reference set from: gusevlab.org/projects/fusion/, 2) the SNPs used to calculate the TWAS weights, i.e. the same set of SNPs used in the DG-GCL eQTL analyses, and 3) the GWAS summary statistic SNPs from PGC2 and the Walters Group Data Repository ^28^. Feature weights were then computed for the DG-GCL for genes, exons, exon-exon junctions, and transcripts by running the weight computing TWAS-FUSION R script based off of Gusev’s work. After the functional weight was computed for each feature, two more TWAS-FUSION scripts were run (adapted from FUSION.assoc_test.R and FUSION.post_process.R) which applied the weights to the GWAS summary statistic SNPs and calculated the functional-GWAS association statistics. We reclassified variants as proxies if they have GWAS P-values < 5e-8 even if they were not in strong LD (R∧2 < 0.8) with the index SNP. Weights from the hippocampus were obtained from a larger sequencing study described in Collado-Torres et al 2018 for comparability ^9^. We computed TWAS FDR-and Bonferroni-adjusted p-values within each feature type separately.

## Data Availability

Raw sequencing reads will be made available through SRA at accession

**[TBD]** and processed data will be made available through GEO at accession **[TBD]**

## Code Availability

https://github.com/lieberinstitute/dg_hippo_paper

## Supplementary Figures

**Figure S1.**
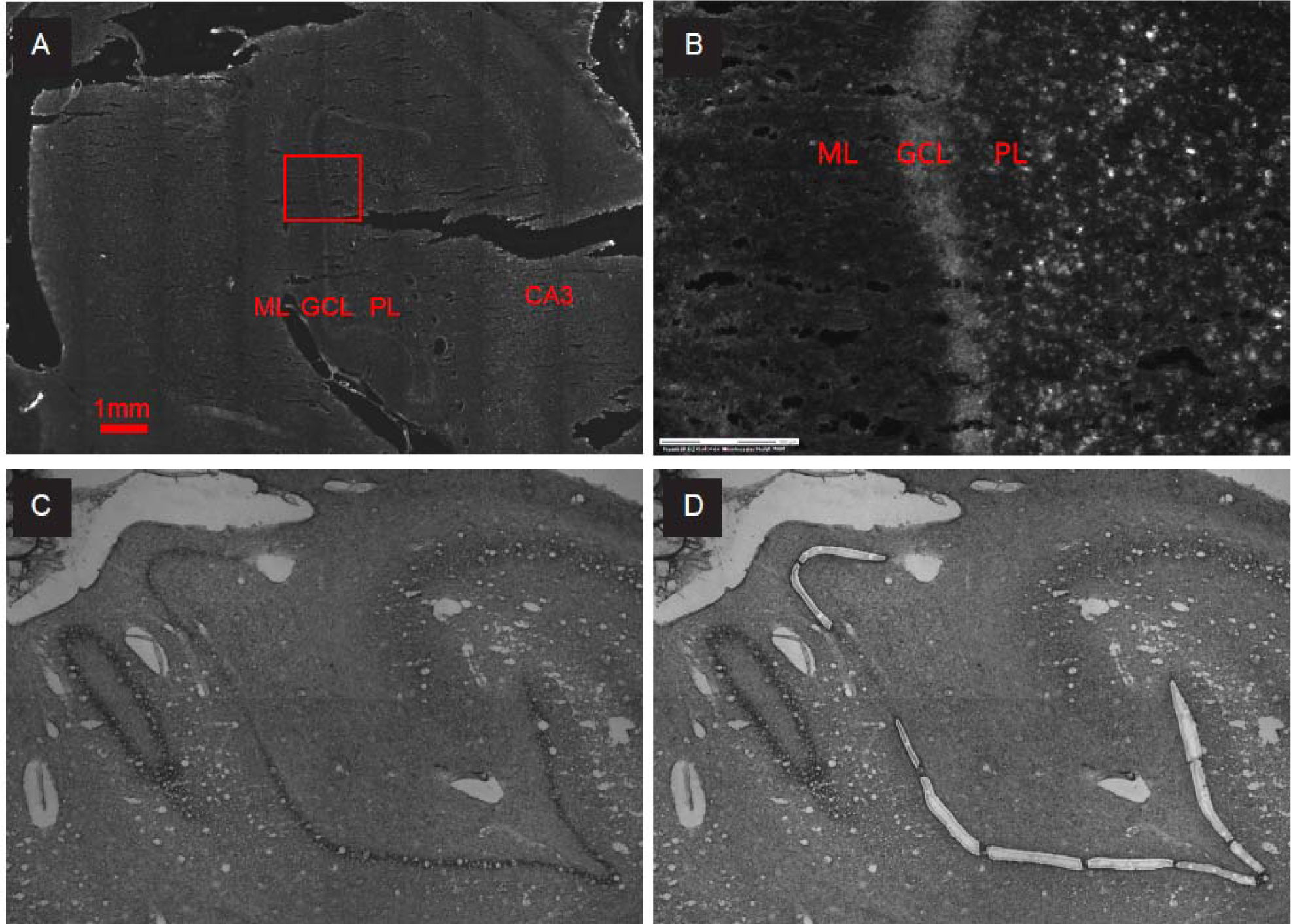
LCM of granule cell layer from human postmortem dentate gyrus. Micrographs of 30 um frozen hippocampal sections (A) original block at the mid-body. (B) Magnified from box in A showing cytoarchitectural distinction between DG laminae with phase contrast. Scale bar = 0.3 mm (C) Cresyl violet stain before LCM (D) post-LCM. The difference indicates the captured granule cell layer.

**Figure S2.**
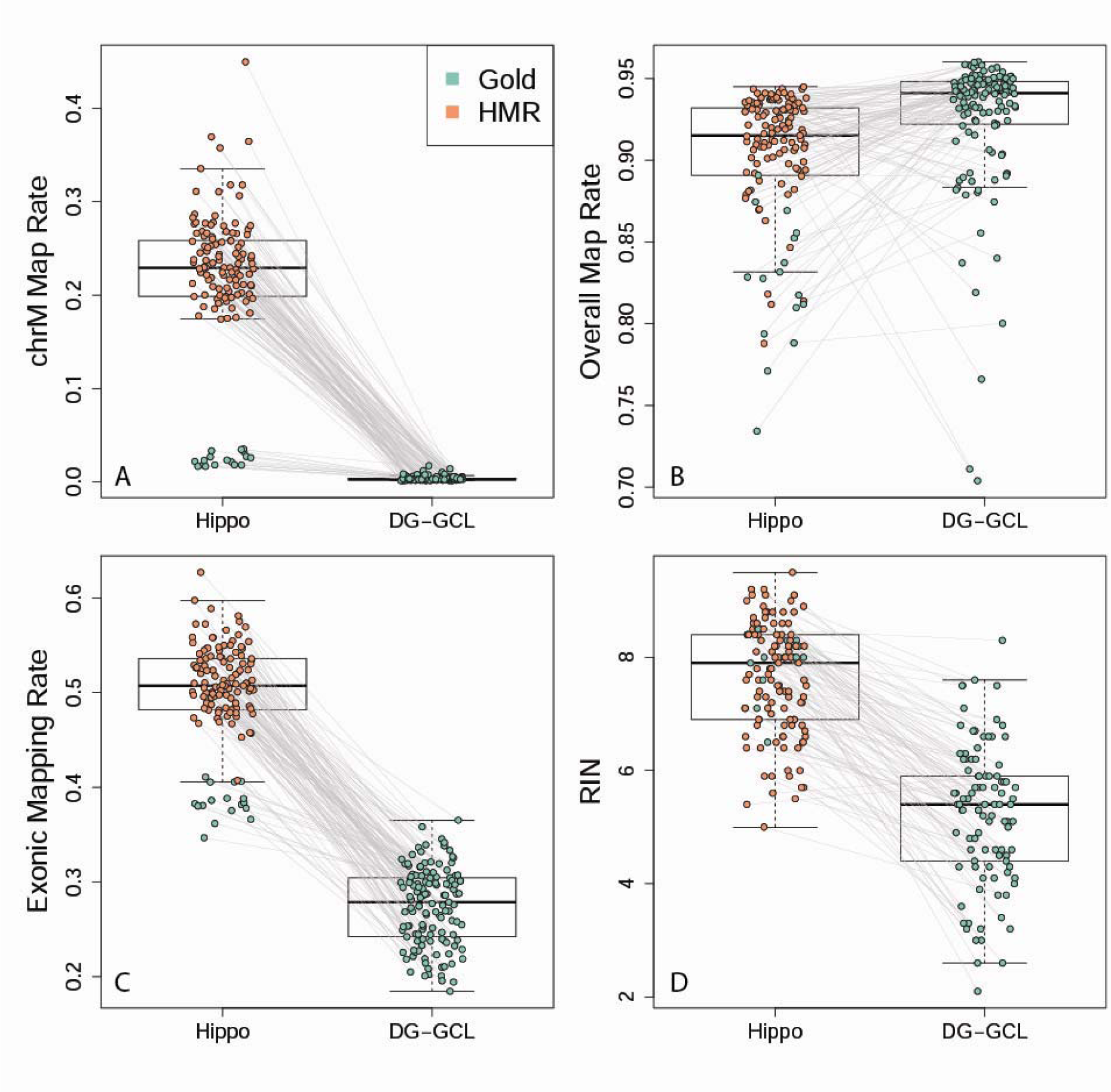
Quality control metrics across the DG-GCL and bulk hippocampus.

**Figure S3:**
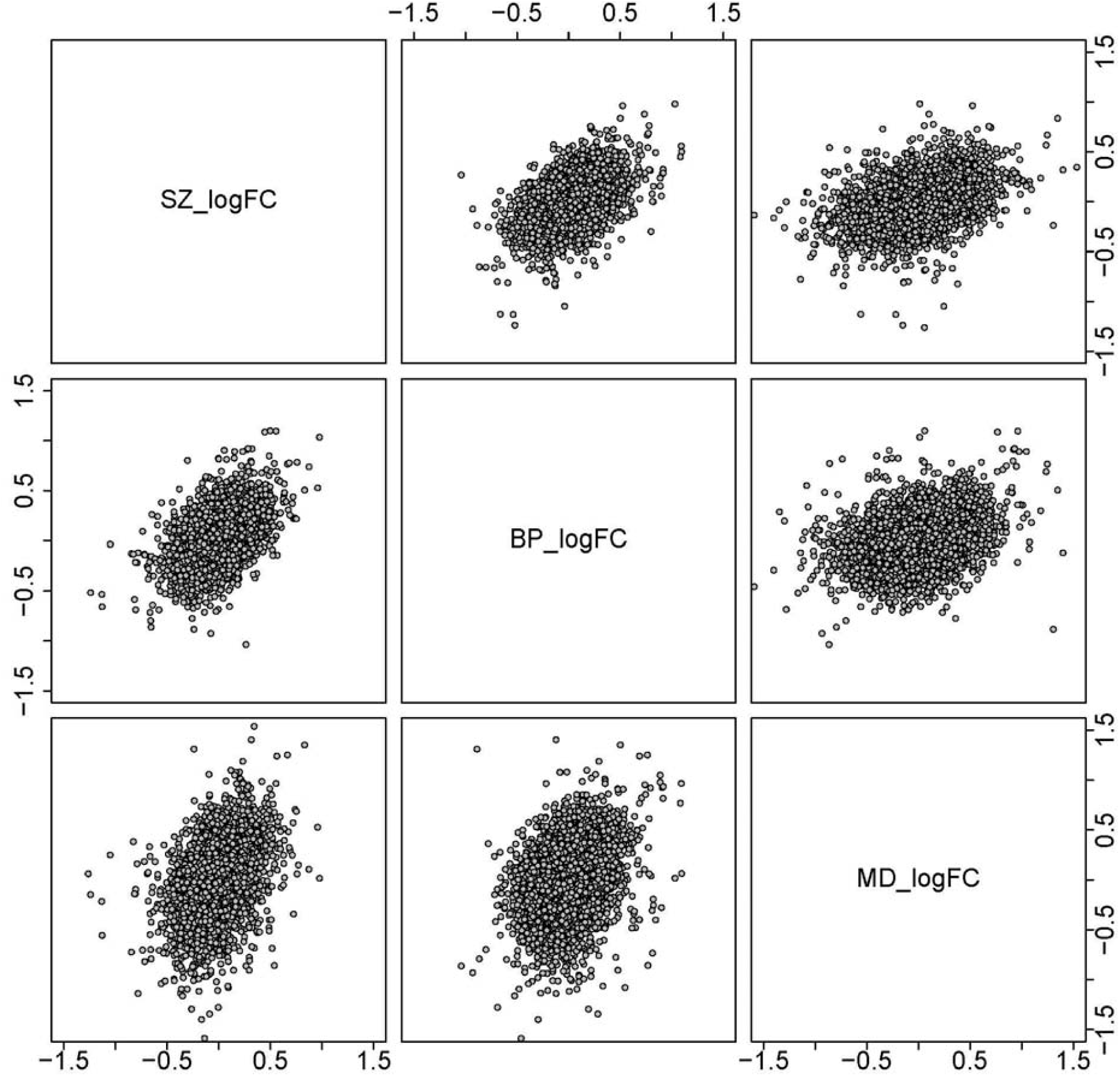
Comparisons of effect sizes across diagnosis

**Figure S4.**
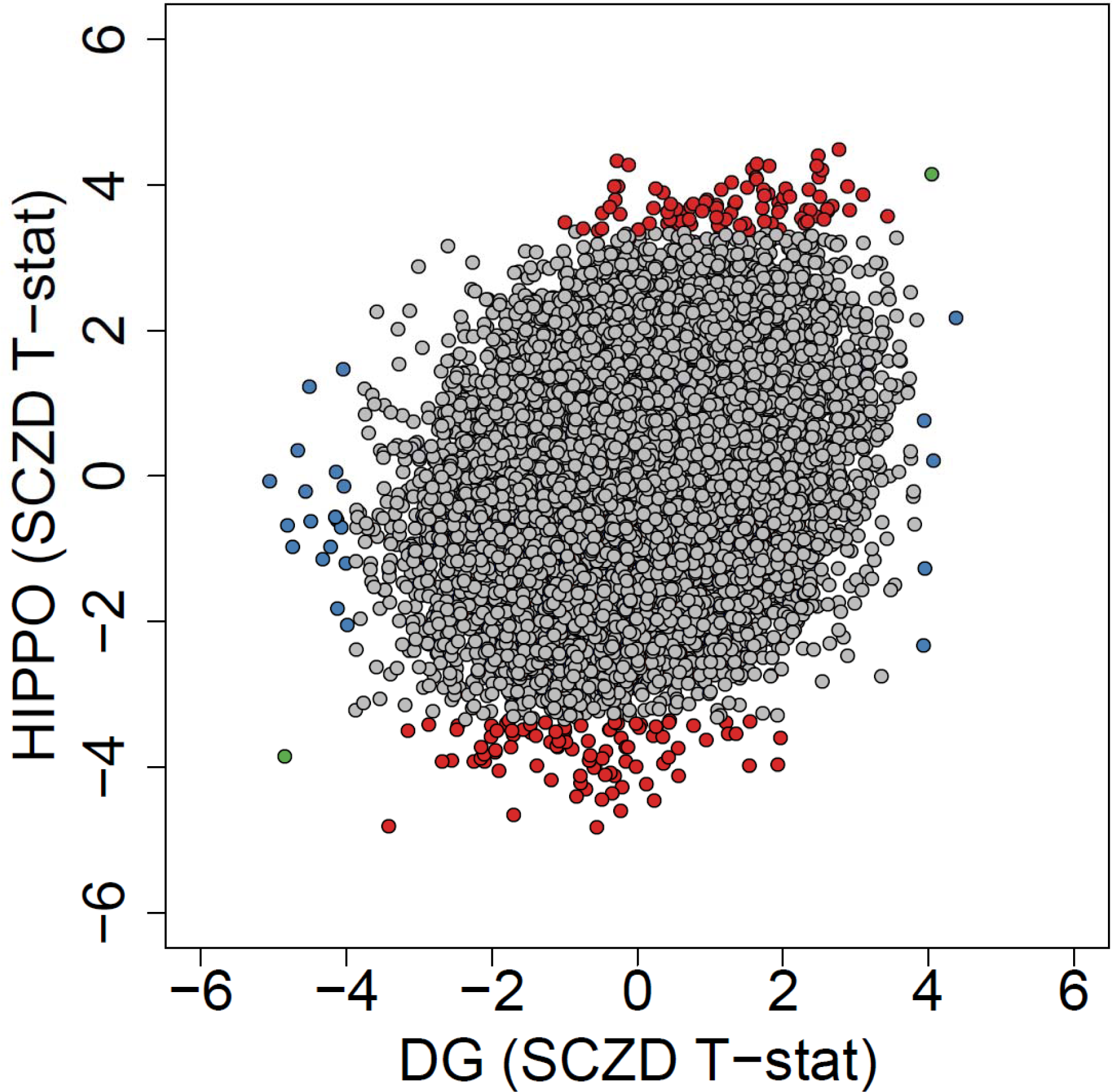
Differential expression by diagnosis within the DG-GCL versus hippocampus reveals cell type-specific illness associations.

## Supplementary Table Legends

Table S1: Demographic data for DG-GCL subjects. * p <0.05 using logistic regression for binary traits and linear regression for quantitative traits.

Table S2: DG-GCL versus hippocampus differential expression statistics via the limma voom approach. Positive log2 fold changes and t-statistics correspond to higher expression in the DG-GCL than hippocampus

Table S3: Gene ontology enrichments for genes significantly differentially expressed between the DG-GCL and hippocampus

Table S4: Age-associated differential expression statistics via limma voom in the DG-GCL, with corresponding aging statistics calculated in the hippocampus and age-interaction statistics calculated using both datasets.

Table S5: Gene ontology enrichments for genes significantly associated with age, stratified by dataset and expression association directionality across the lifespan

Table S6: eQTLs to schizophrenia risk loci or highly correlated proxies from the PGC2+CLOZUK GWAS.

Table S7: Significant TWAS signals at FDR < 5% using the PGC2+CLOZUK schizophrenia GWAS.

Table S8: Differential expression analysis for each diagnostic group using the limma voom approach

Data S1: genome-wide eQTL associations at FDR < 1% in the DG-GCL with corresponding statistics in homogenate hippocampus. Available at: https://s3.us-east-2.amazonaws.com/lcmseq-dggcl/DataS1_DGGCL_eQTLs_plusHippo.csv.gz

