## Supplementary material for "Cell type-specific genetic regulation of expression in the granule cell layer of the human dentate gyrus": suppData2_ageIntExamples.pdf

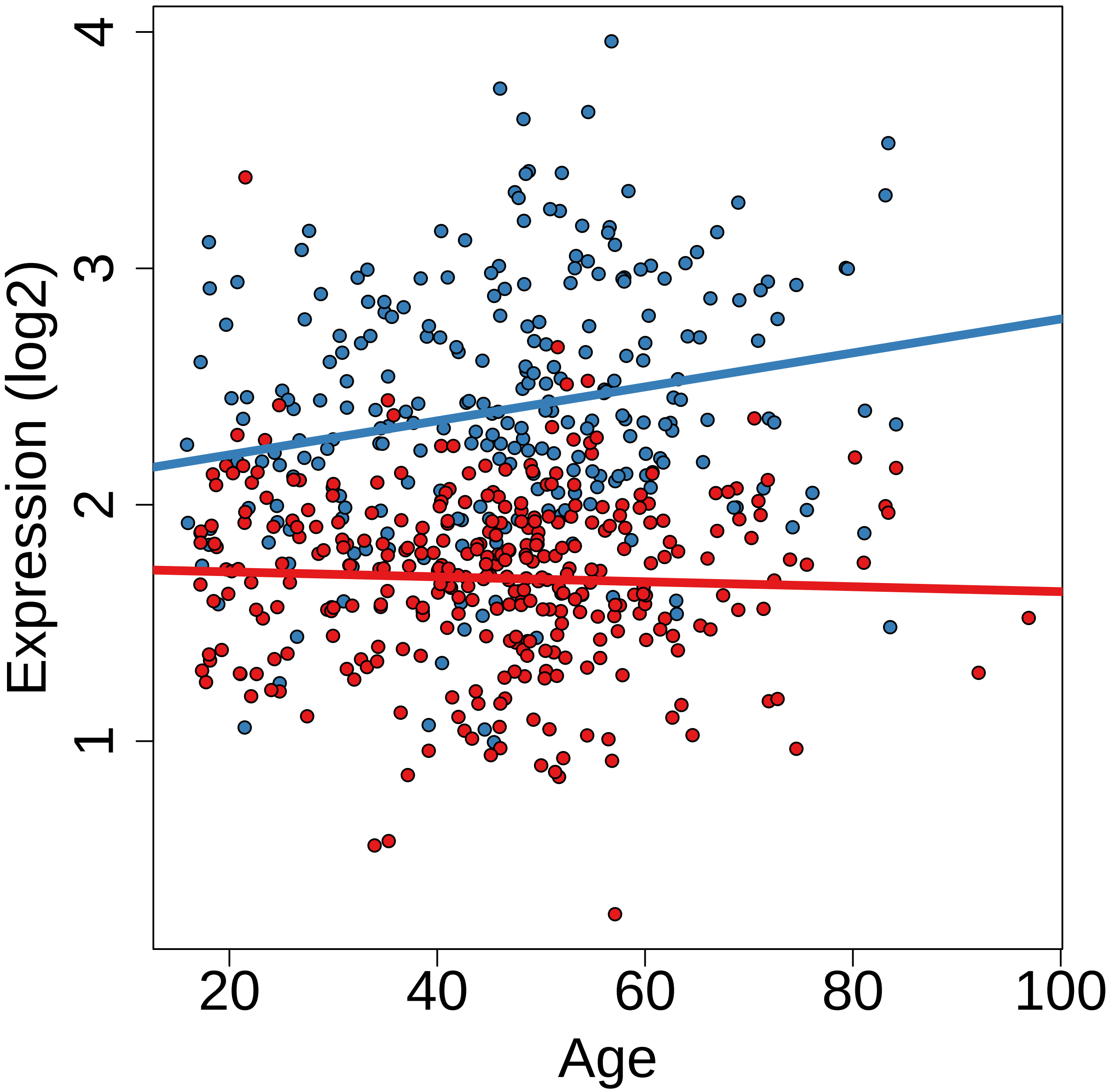

Expression (log2)
1
2
3
4
20
40
60
80
100
Age

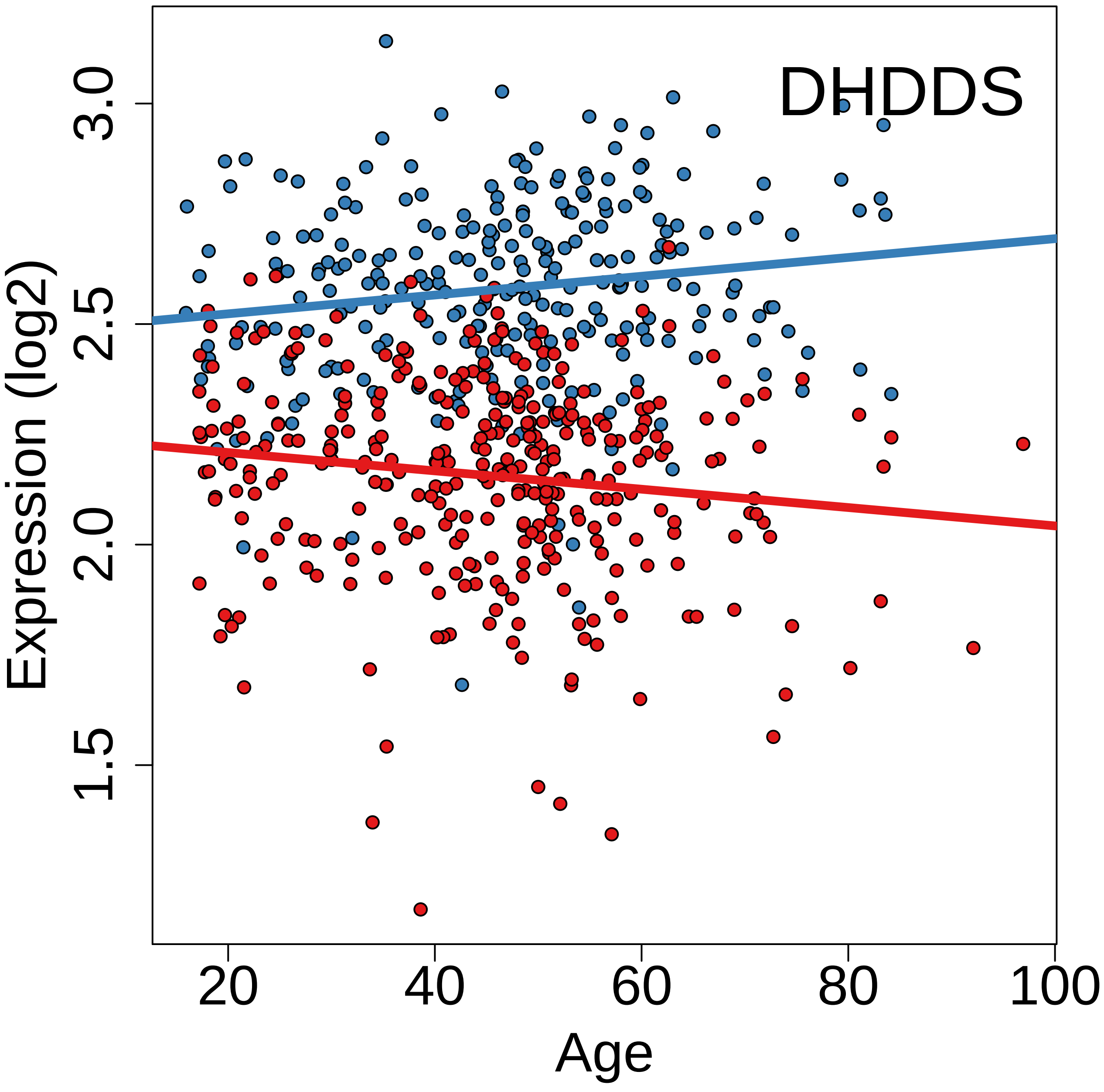
DHDDS
Expression (log2)
1.5
2.0
2.5
3.0
20
40
60
80
100
Age

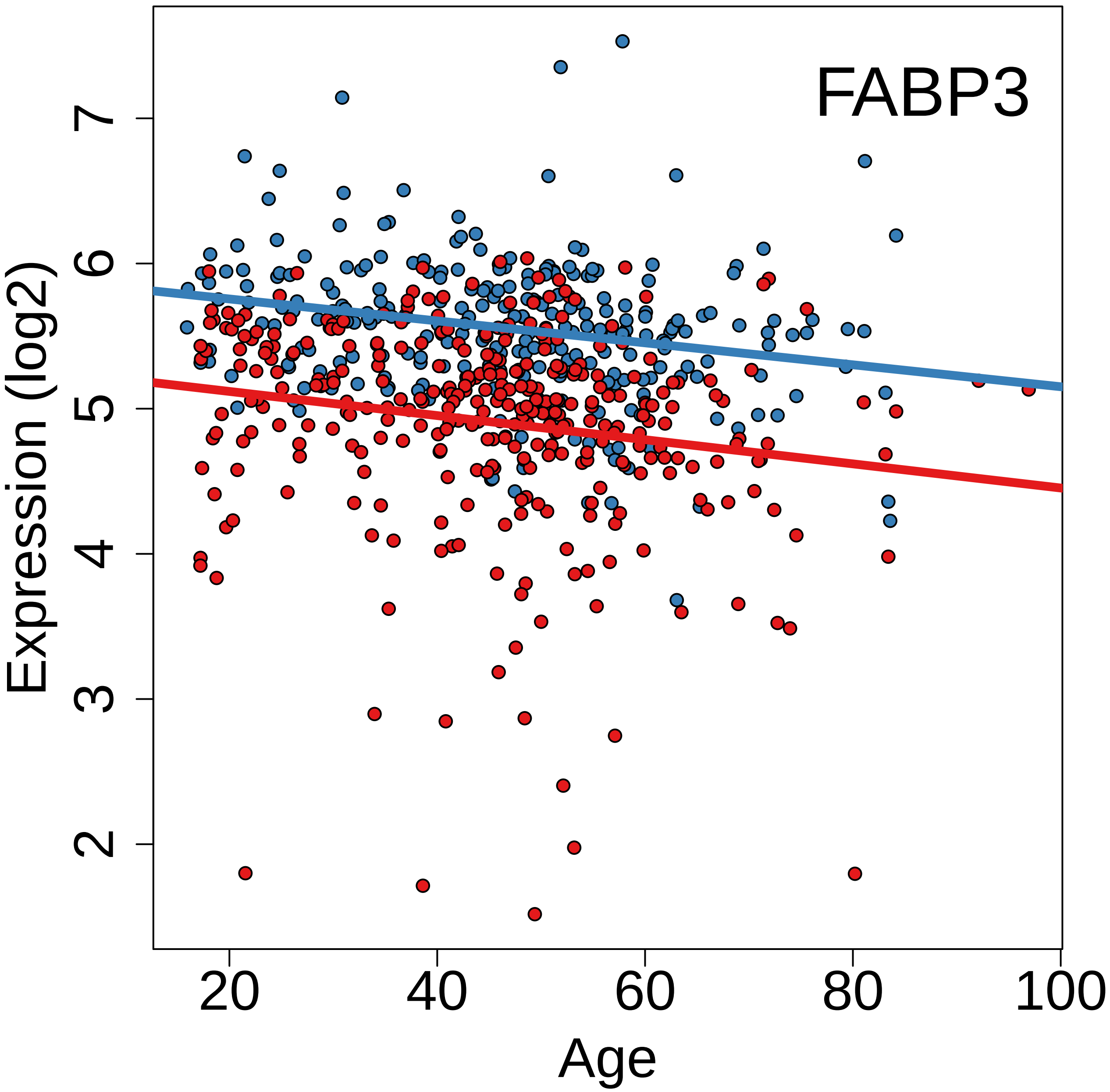
FABP3
Expression (log2)
Age
2
3
4
5
6
7
20
40
60
80
100

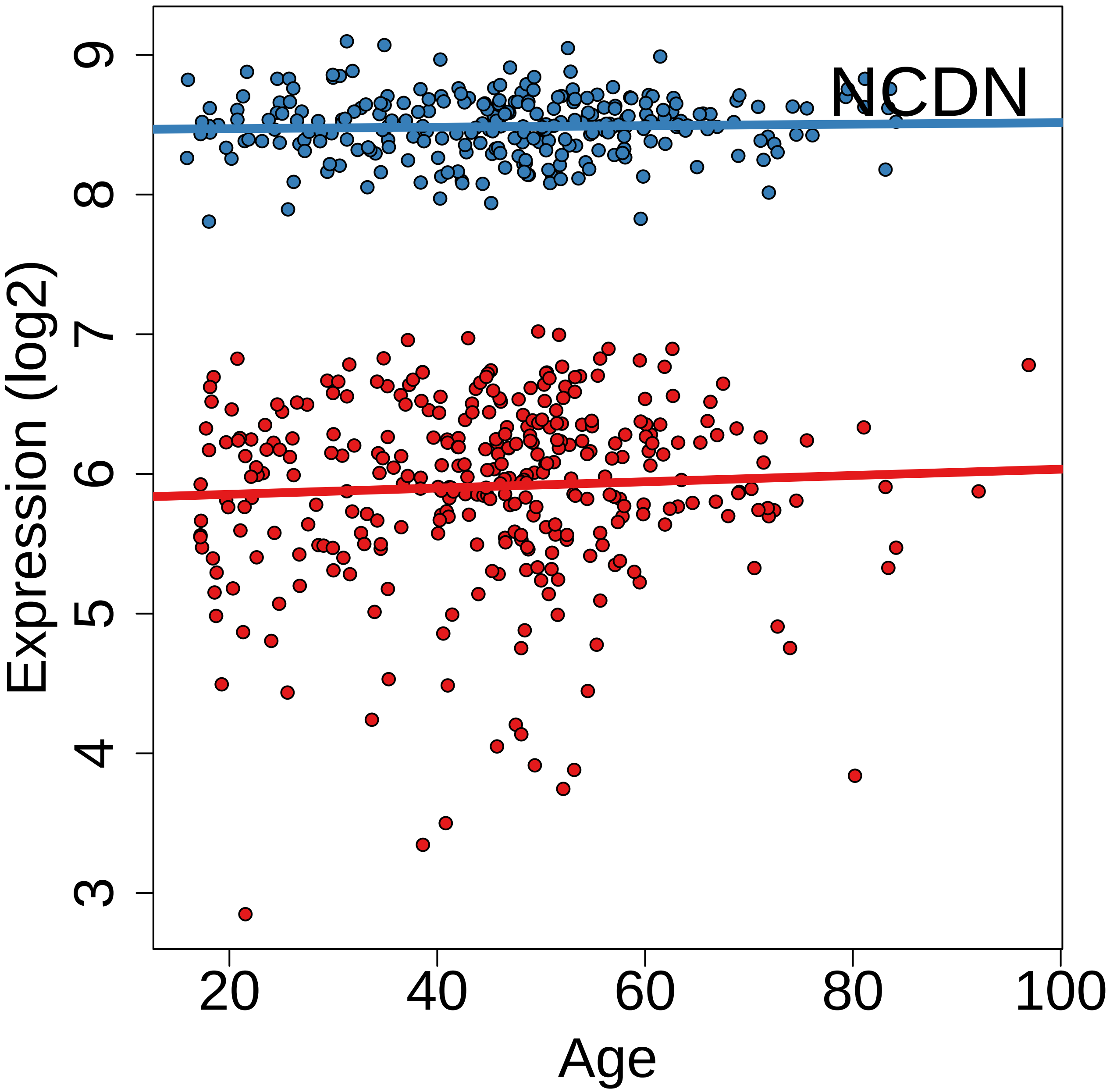

NCDN
Expression (log2)
Age
3
4
5
6
7
8
9
20
40
60
80
100

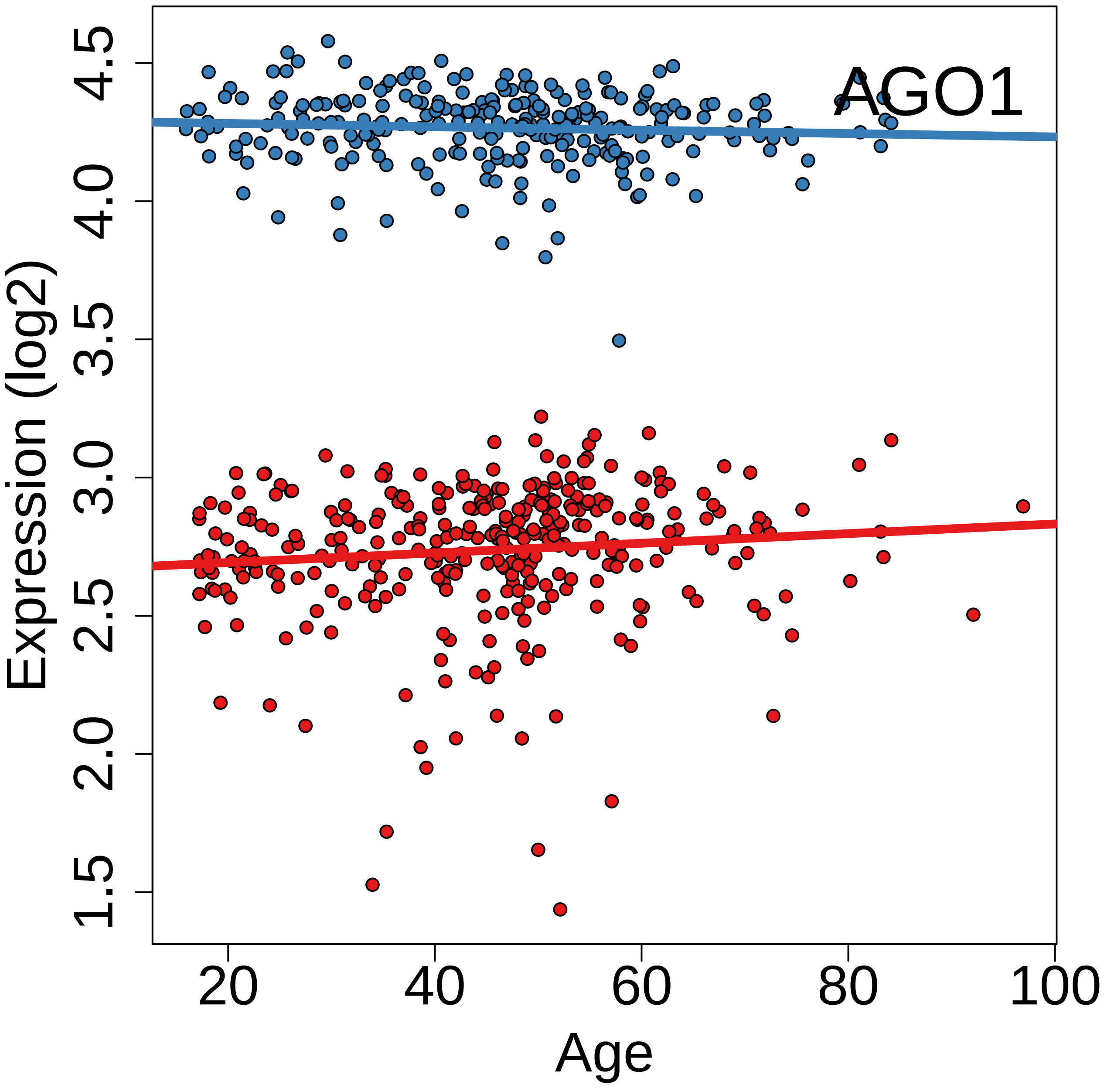
AGO1
Expression (log2)
1.5
2.0
2.5
3.0
3.5
4.0
4.5
20
40
60
80
100
Age

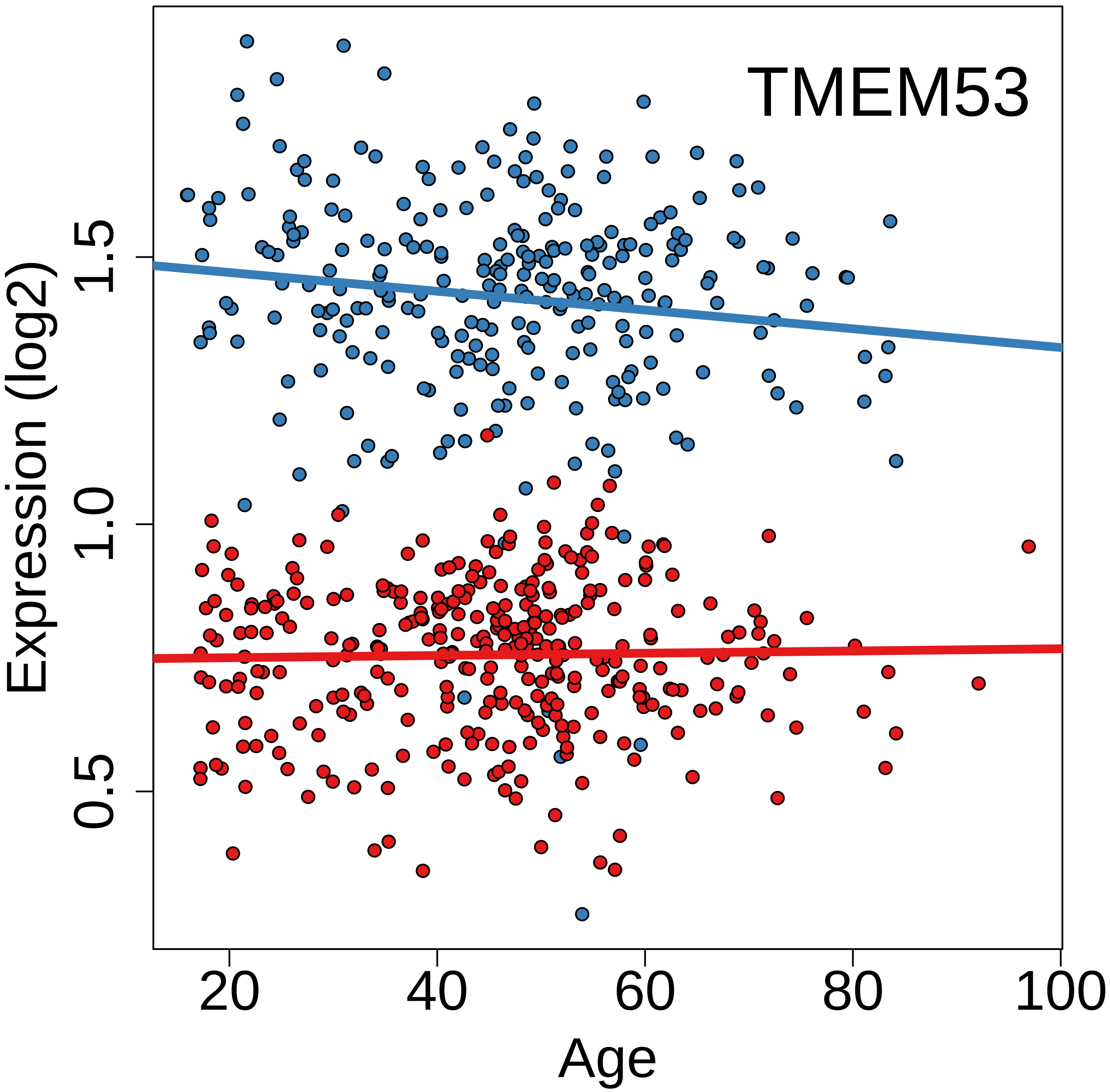
TMEM53
Expression (log2)
0.5
1.0
1.5
20
40
60
80
100
Age

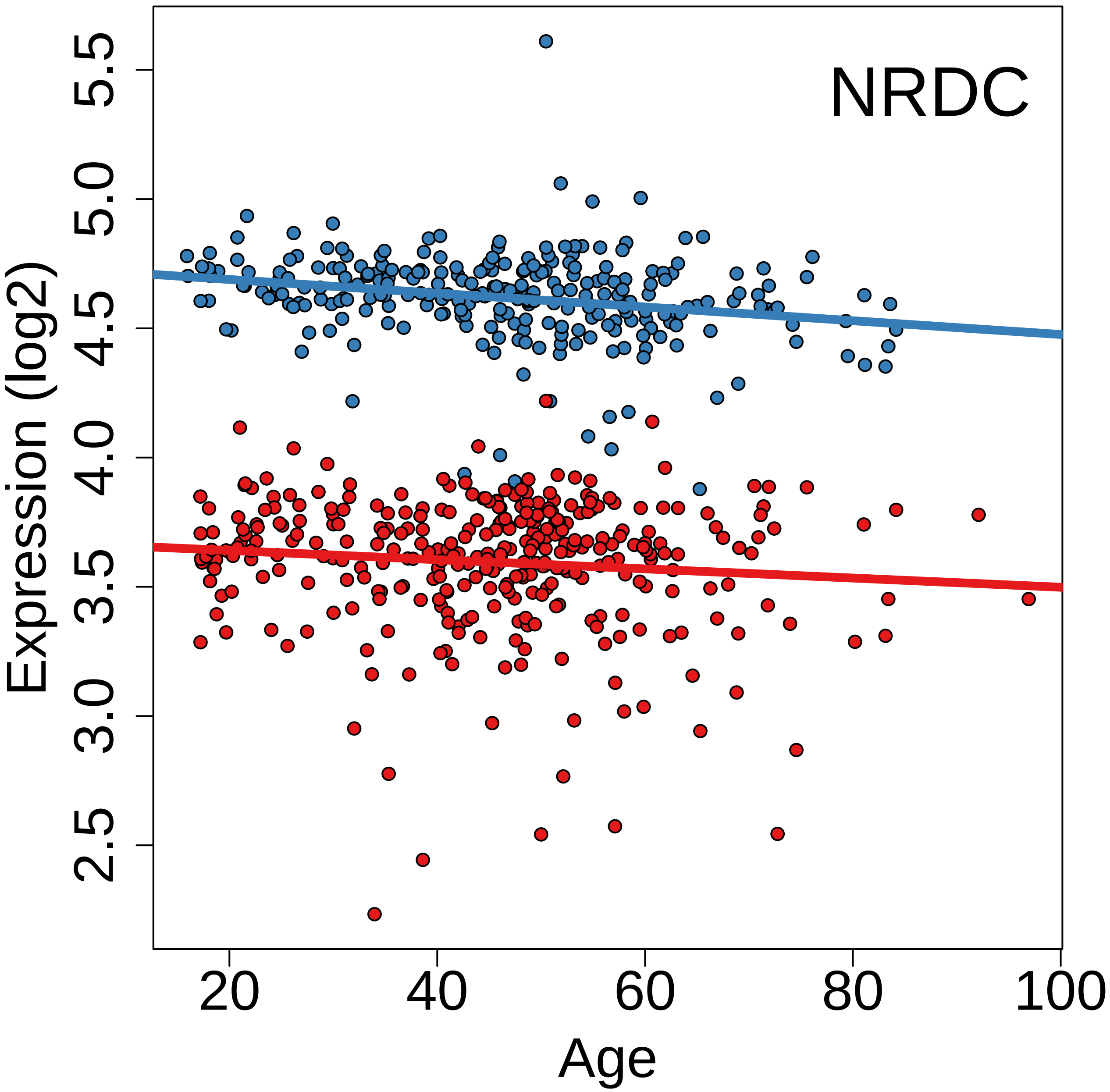
NRDC
Expression (log2)
5.5
5.0
4.5
4.0
3.5
3.0
2.5
20
40
60
80
100
Age

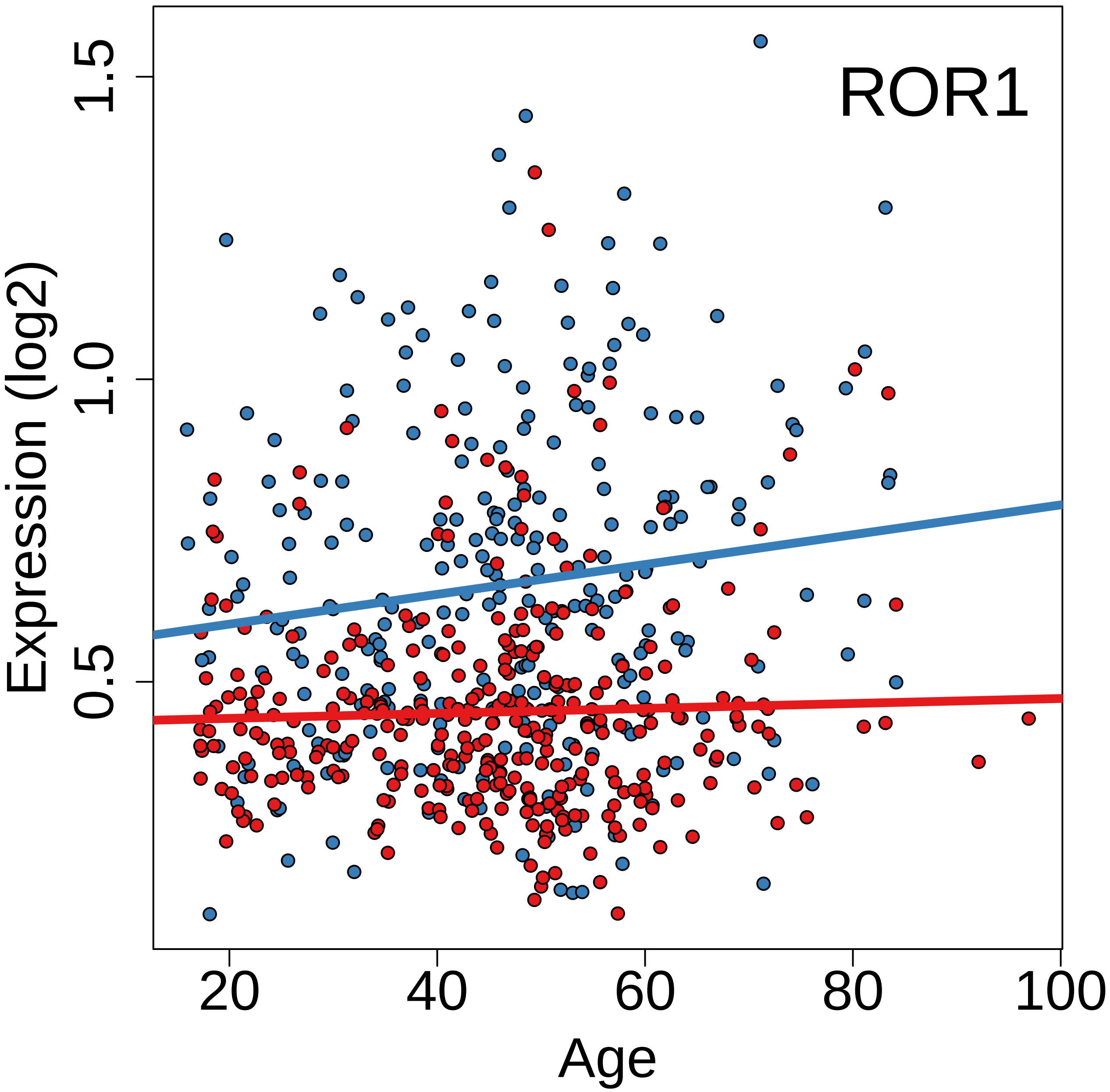
ROR1
Expression (log2)
0.5
1.0
1.5
20
40
60
80
100
Age

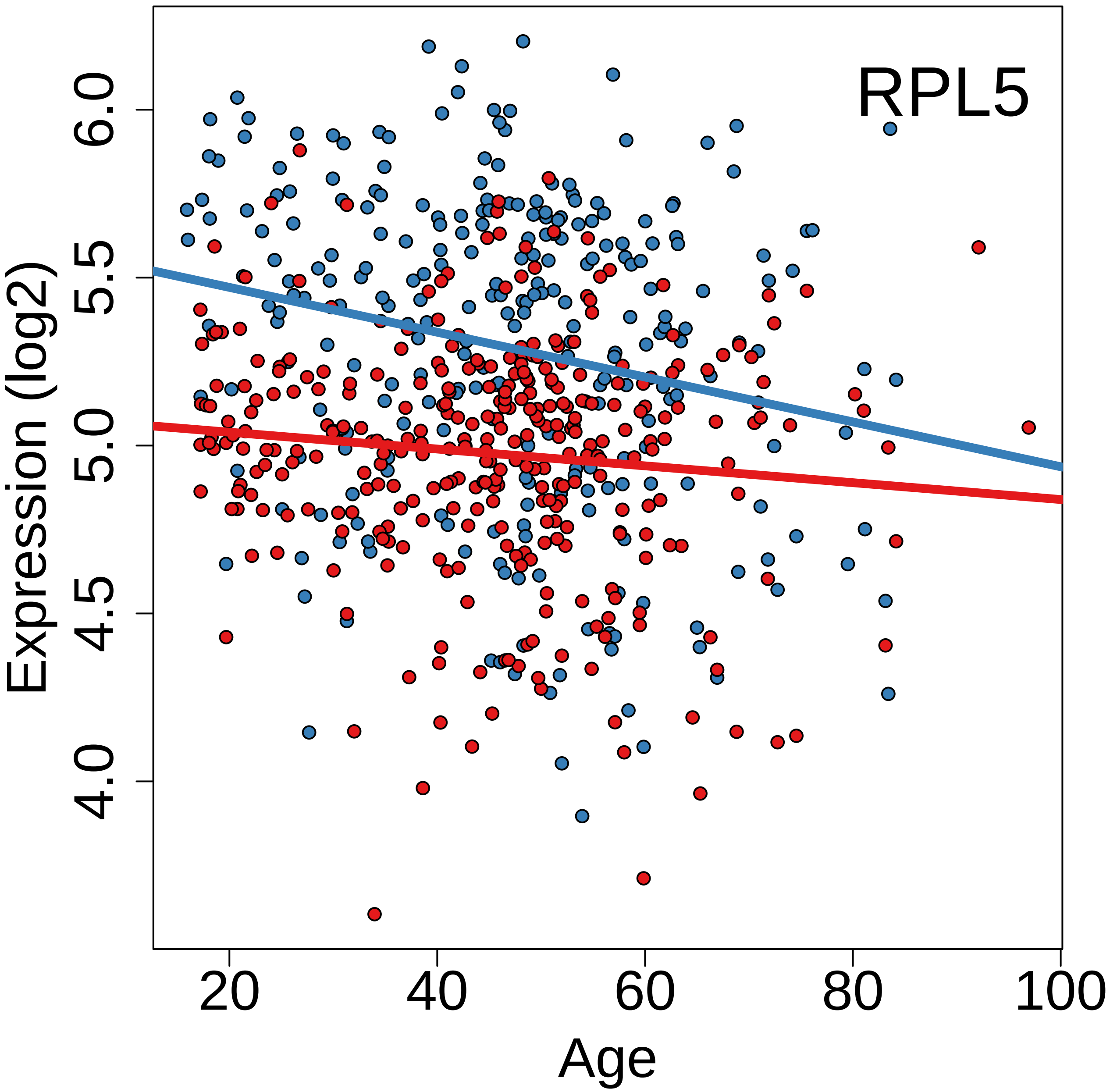
RPL5
Expression (log2)
4.0
4.5
5.0
5.5
6.0
20
40
60
80
100
Age

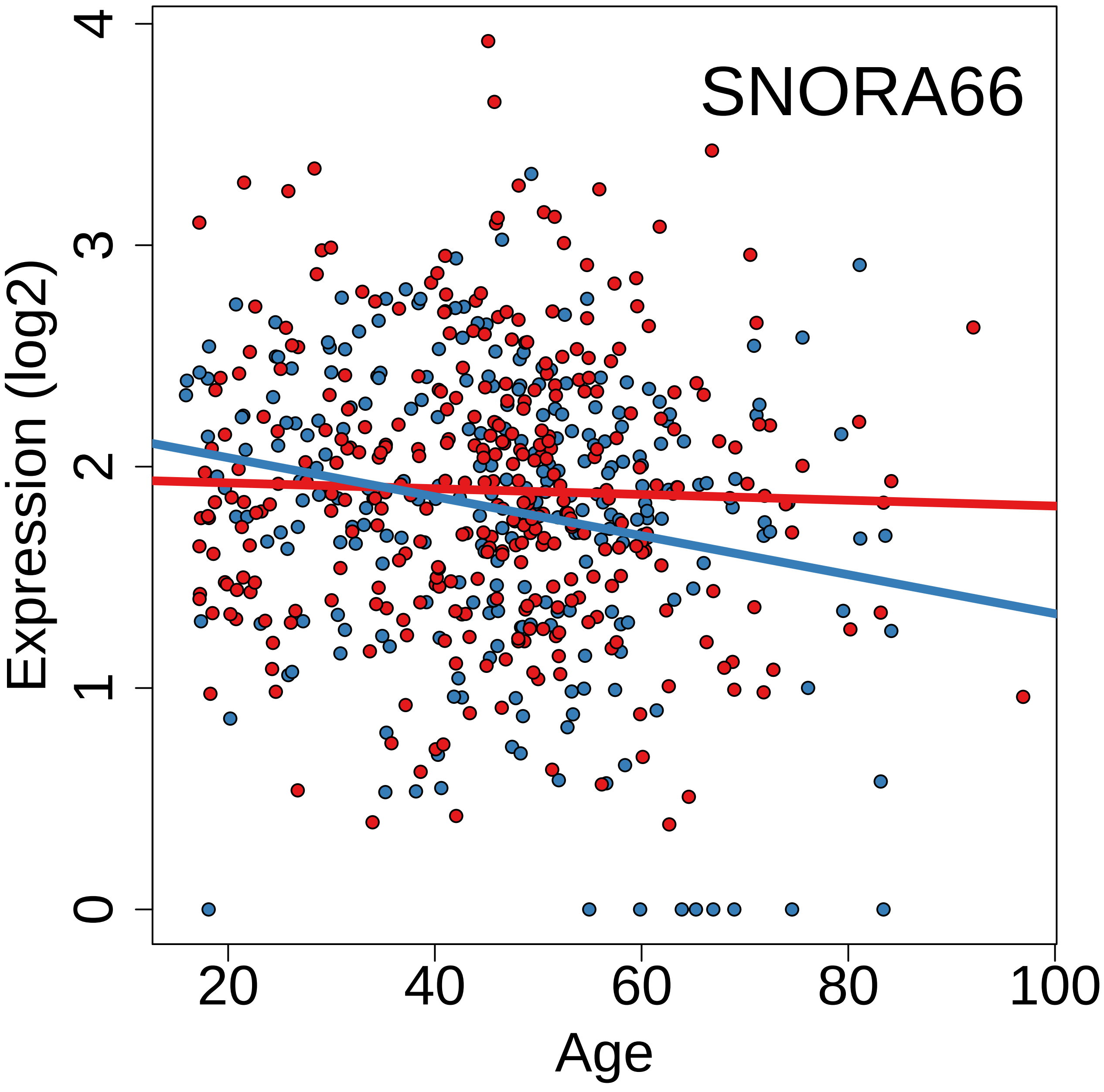
SNORA66
Expression (log2)
0
1
2
3
4
20
40
60
80
100
Age

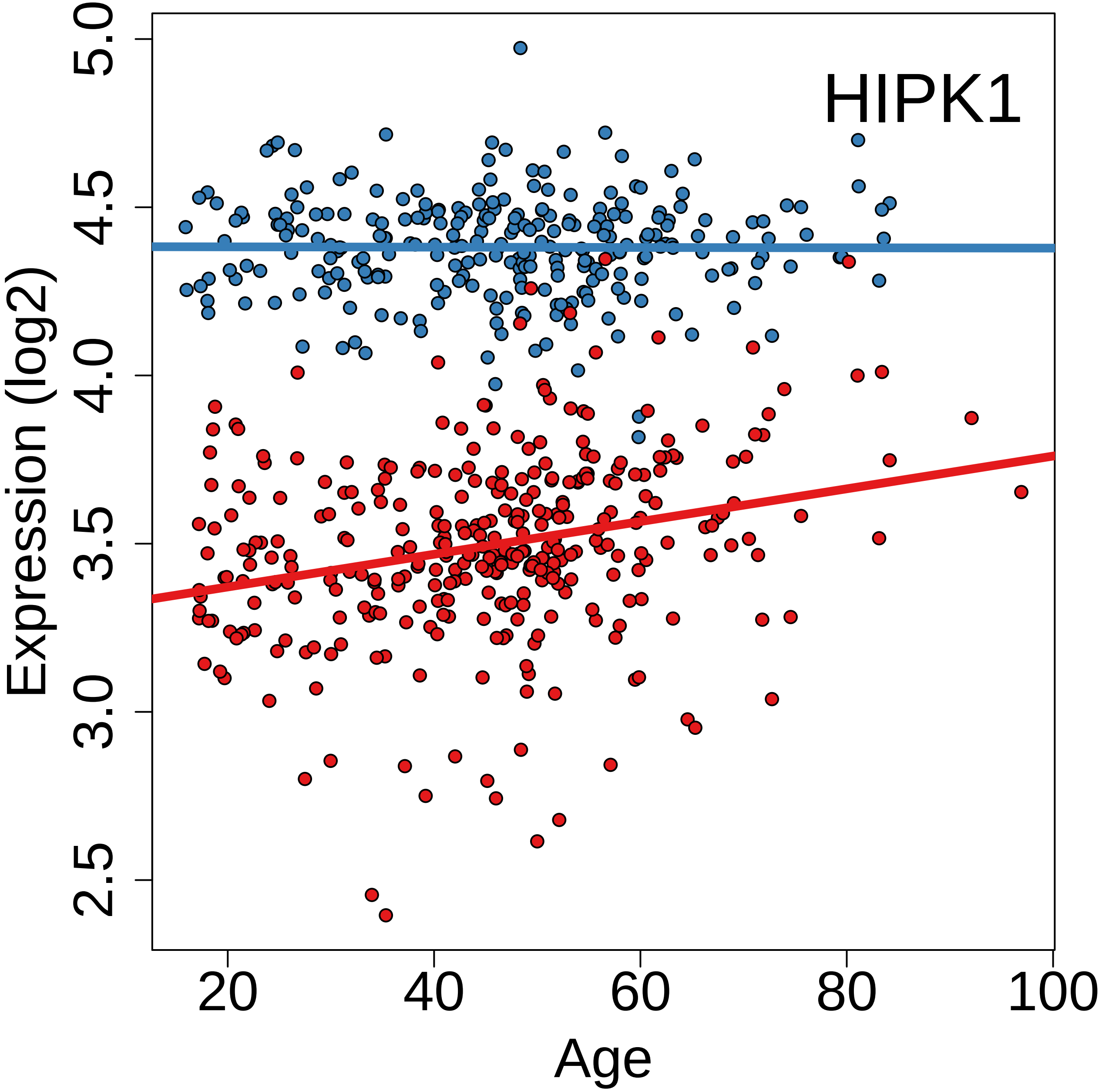
HIPK1
Expression (log2)
5.0
4.5
4.0
3.5
3.0
2.5
20
40
60
80
100
Age

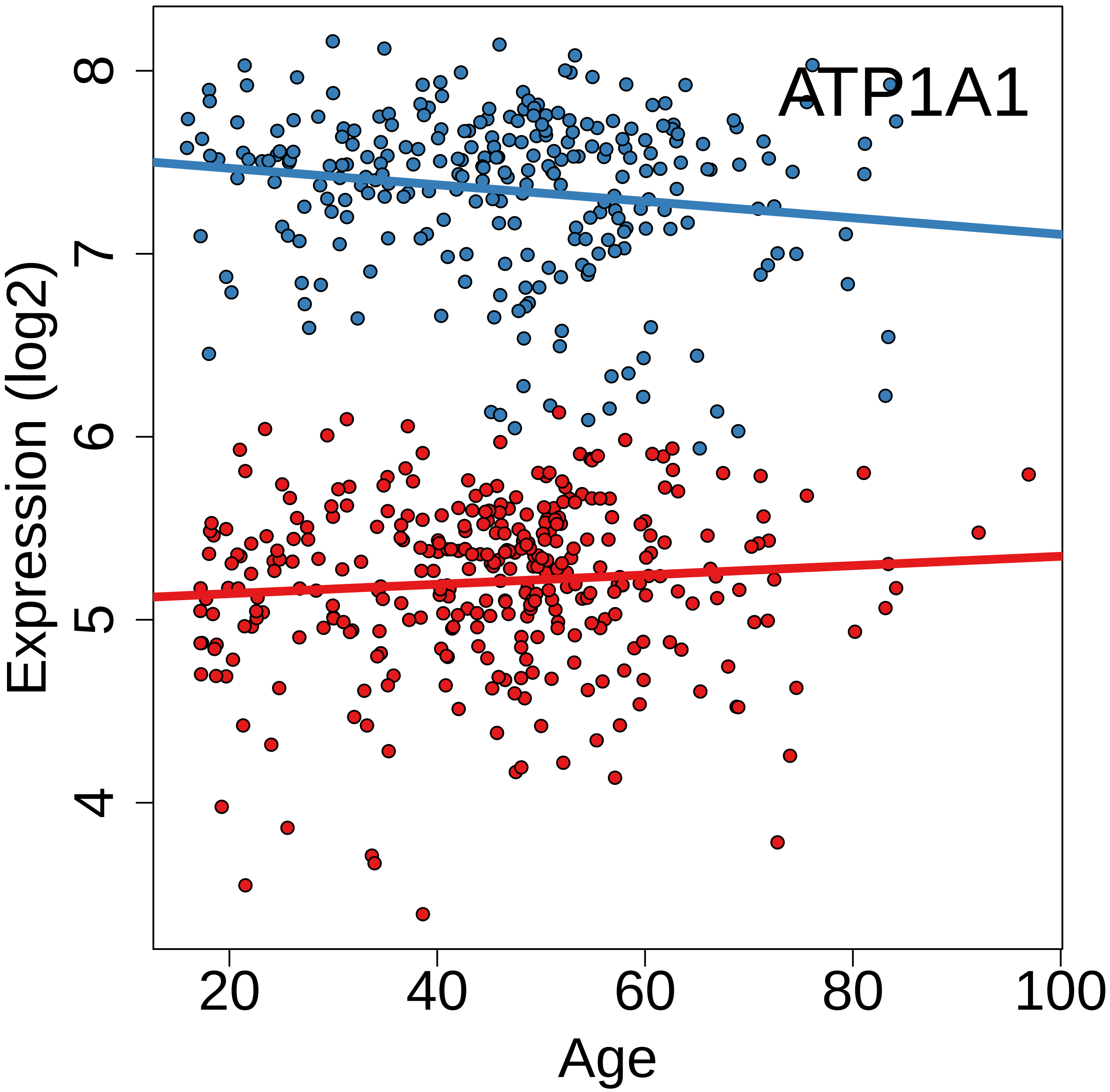
ATP1A1
Expression (log2)
8
7
6
5
4
20
40
60
80
100
Age

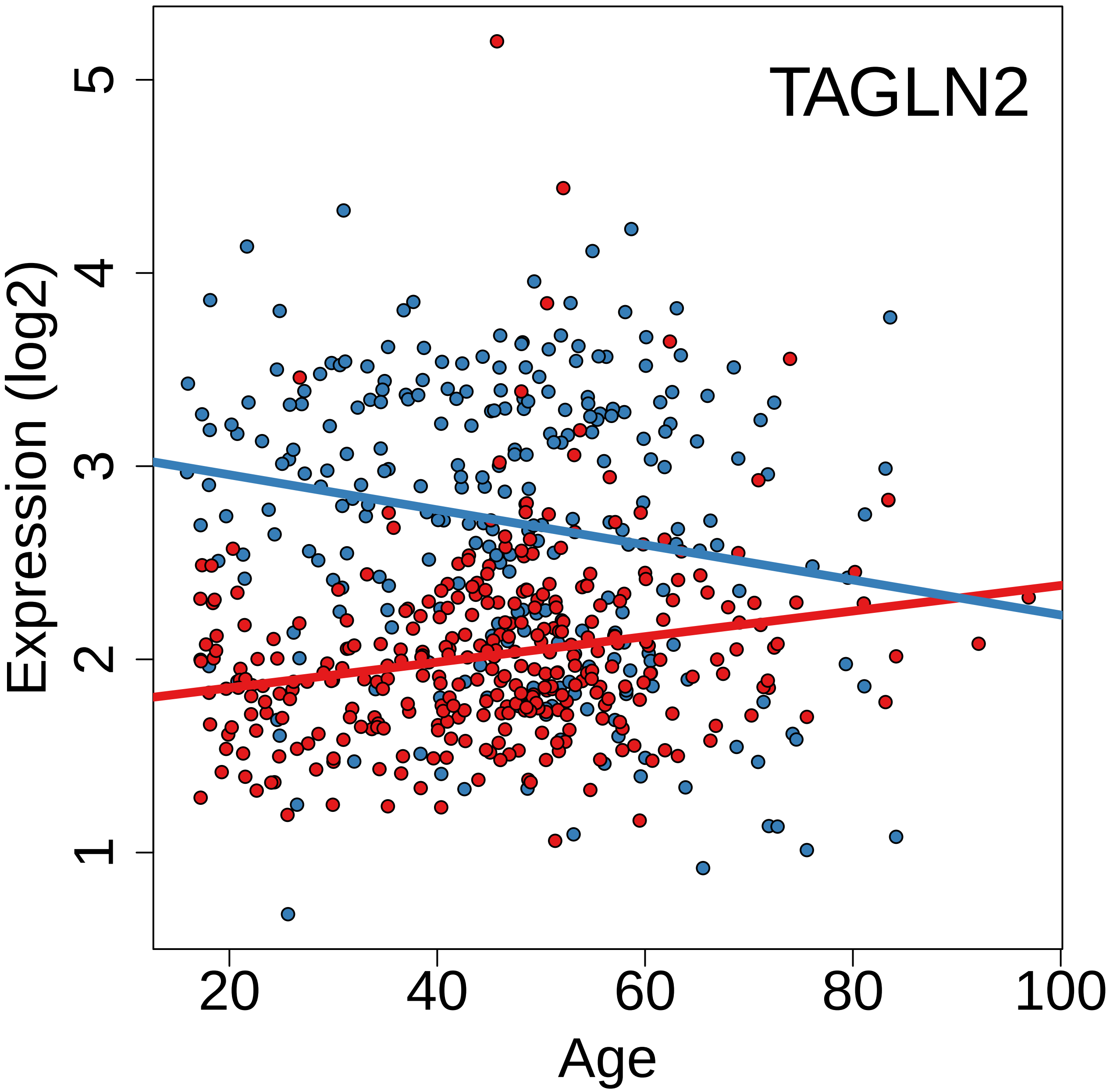
TAGLN2
Expression (log2)
1
2
3
4
5
20
40
60
80
100
Age

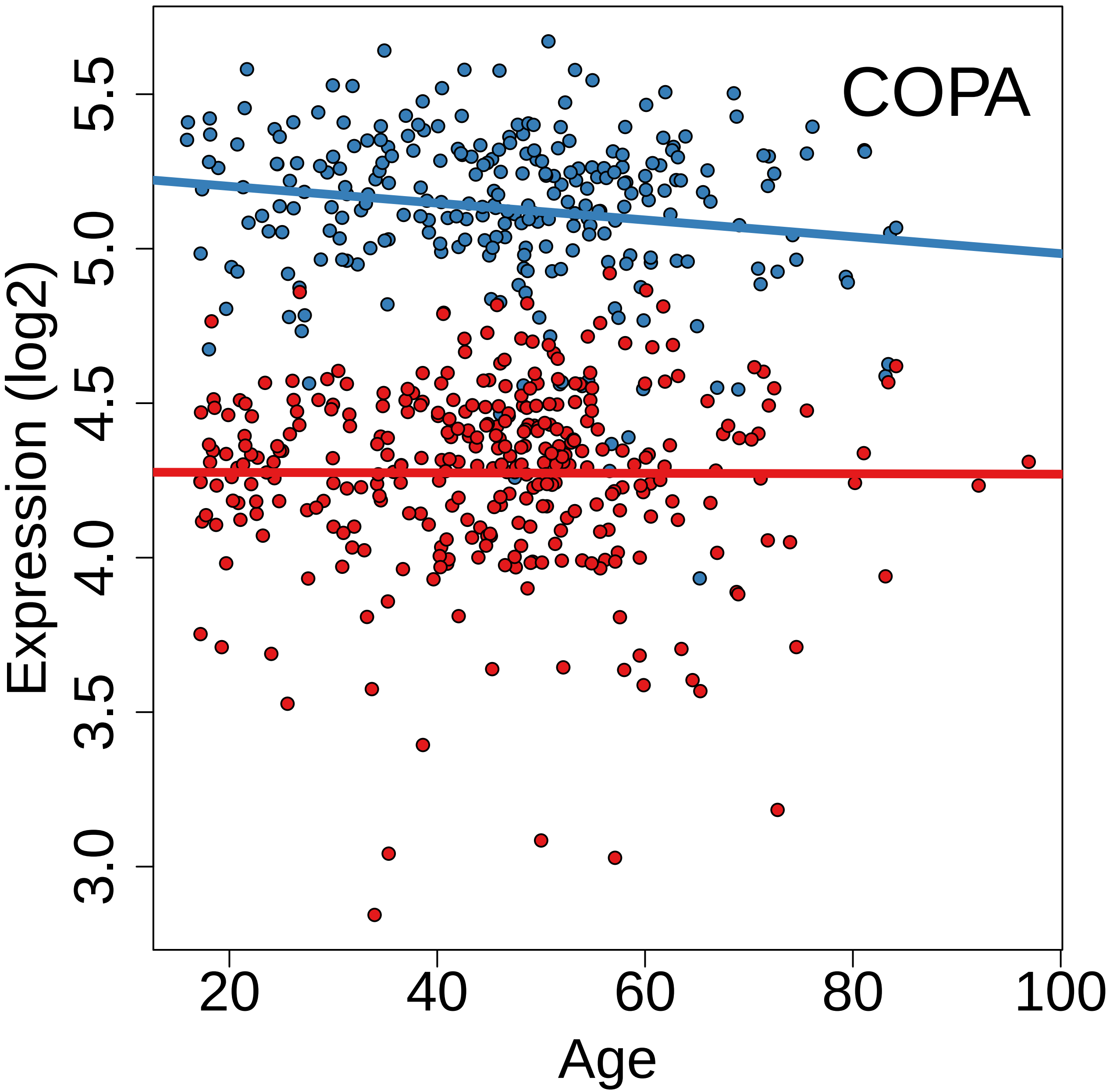

COPA
Expression (log2)
5.5
5.0
4.5
4.0
3.5
3.0
20
40
60
80
100
Age

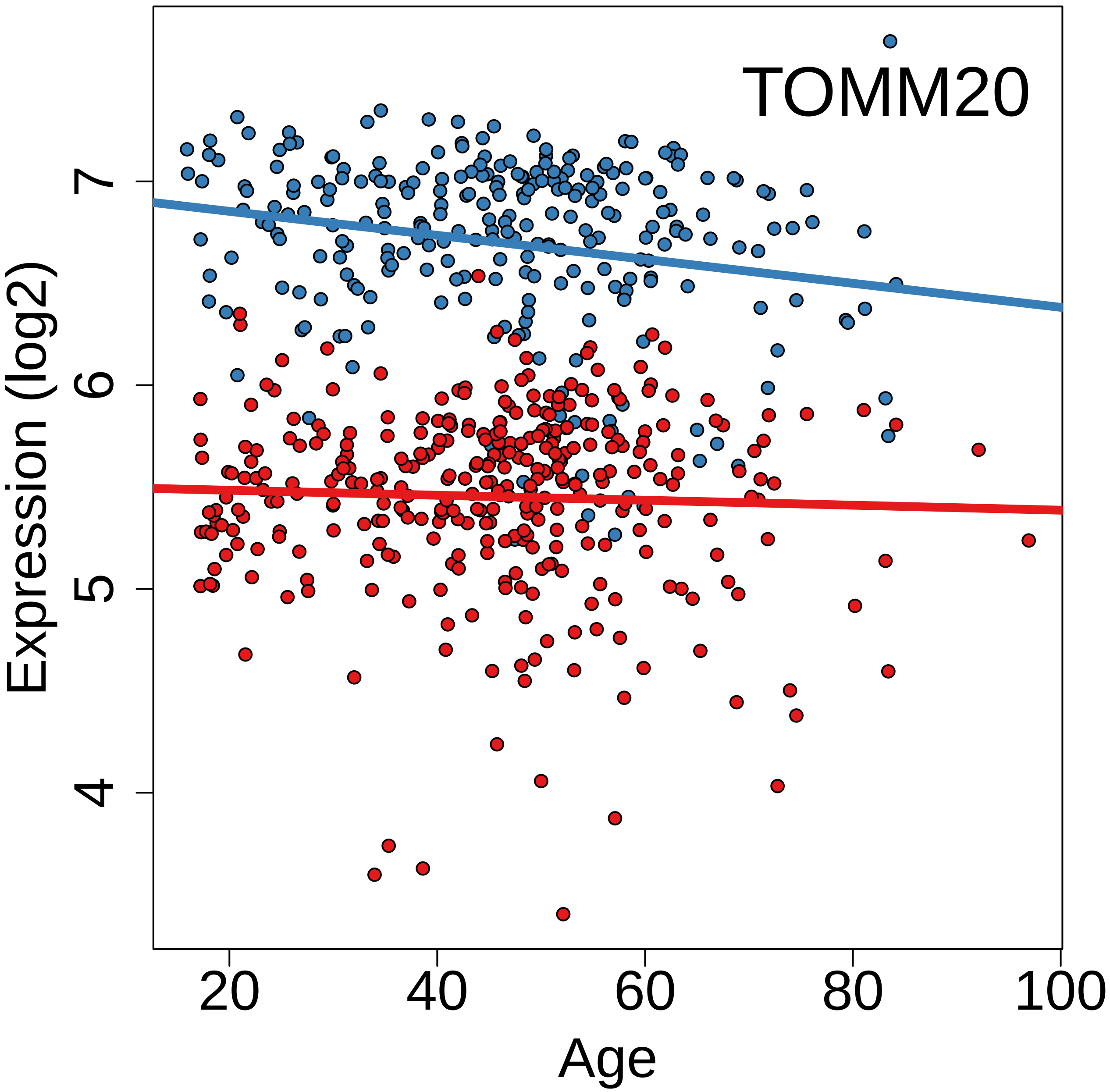
TOMM20
Expression (log2)
4
5
6
7
20
40
60
80
100
Age

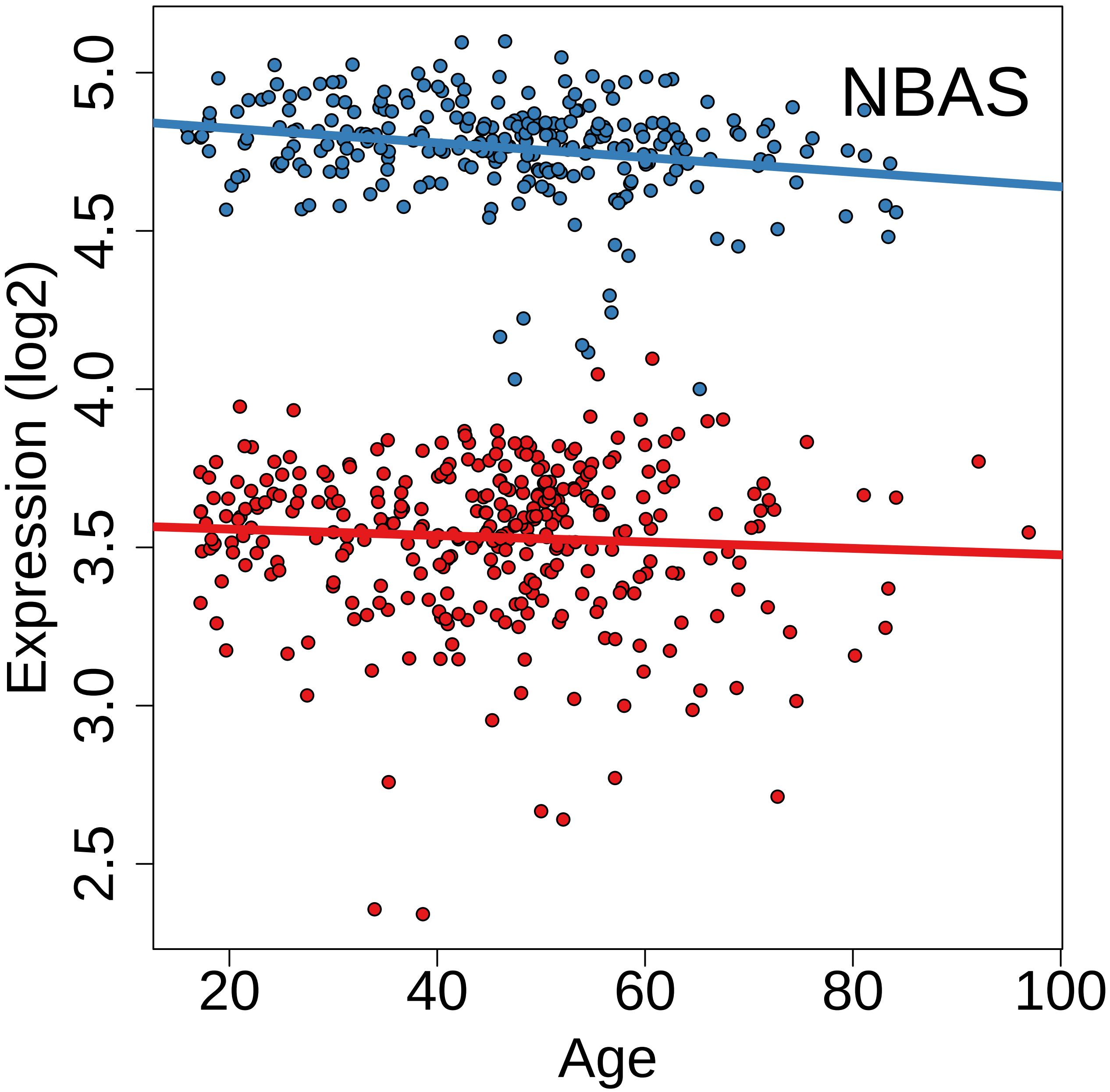
NBAS
Expression (log2)
5.0
4.5
4.0
3.5
3.0
2.5
20
40
60
80
100
Age

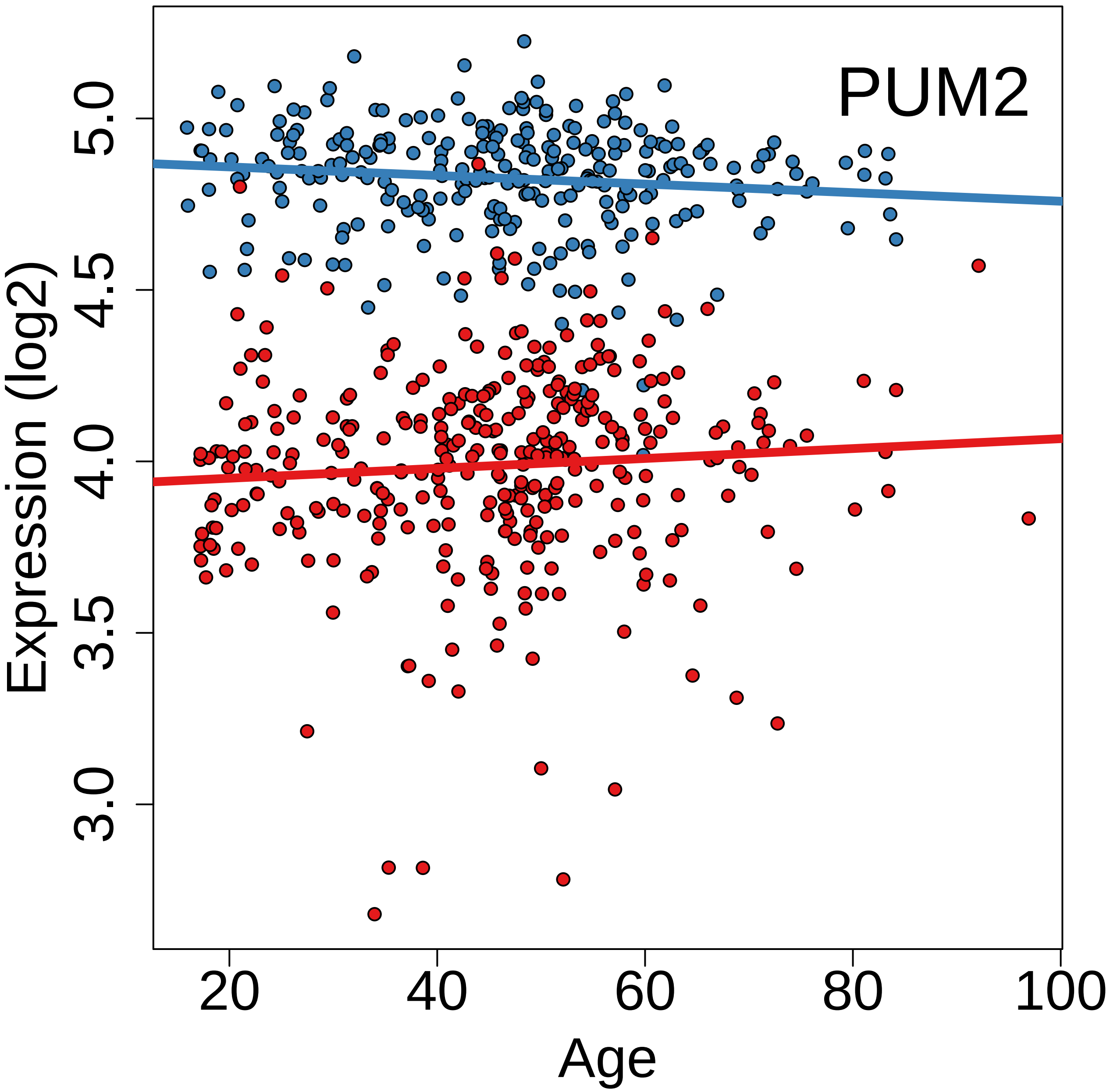

PUM2
Expression (log2)
3.0
3.5
4.0
4.5
5.0
20
40
60
80
100
Age

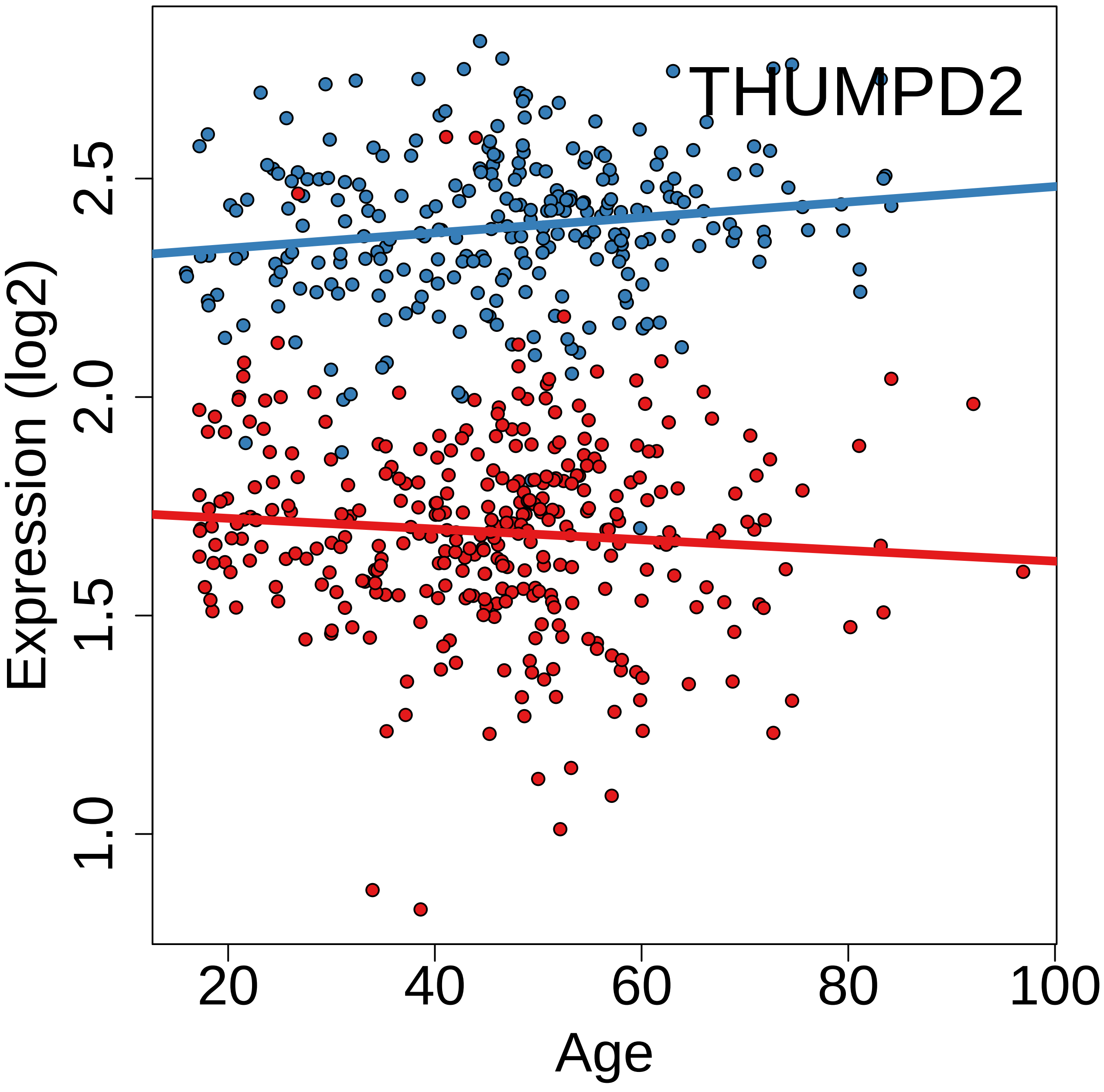
THUMPD2
Expression (log2)
1.0
1.5
2.0
2.5
20
40
60
80
100
Age

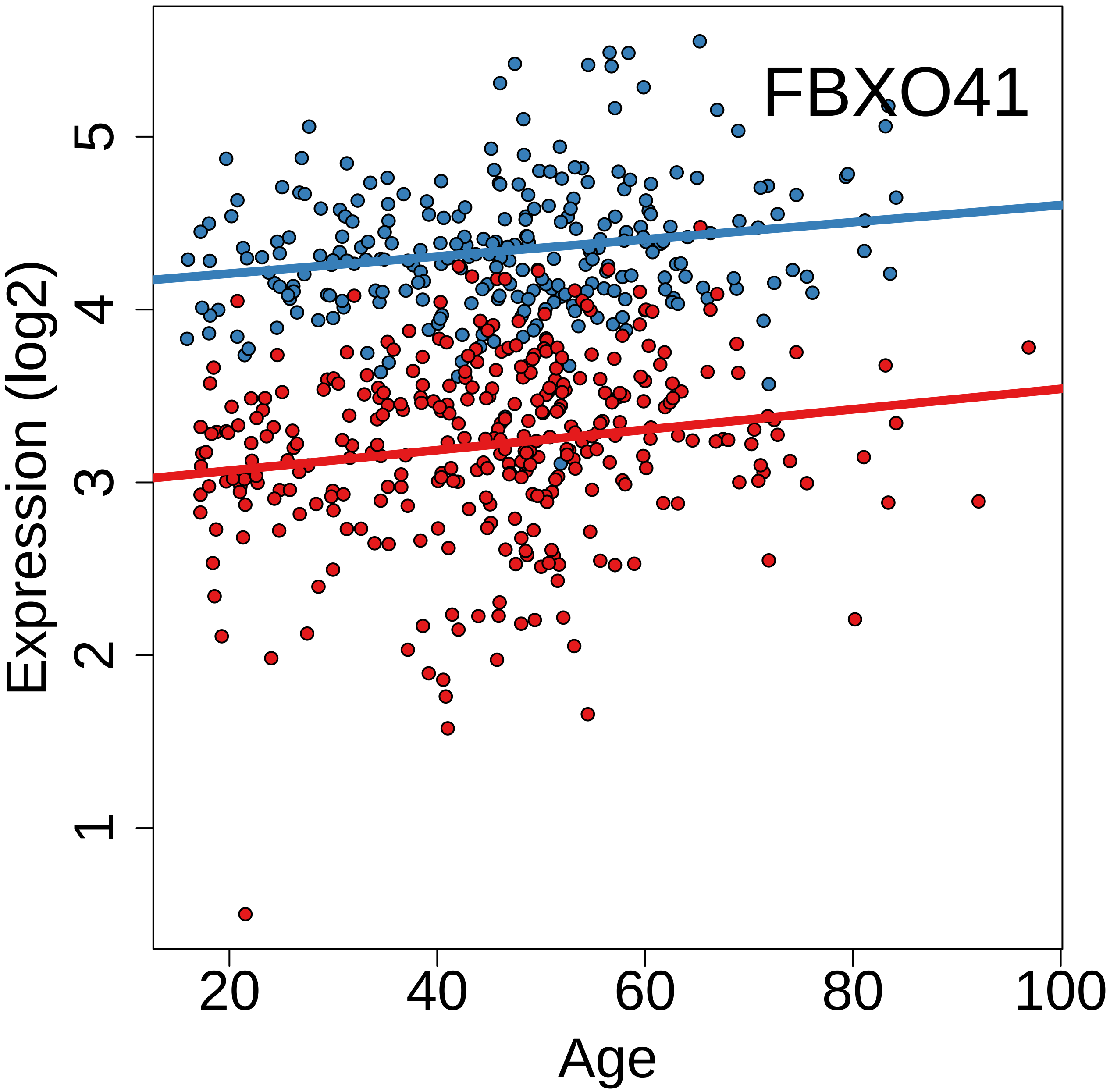
FBXO41
Expression (log2)
1
2
3
4
5
20
40
60
80
100
Age

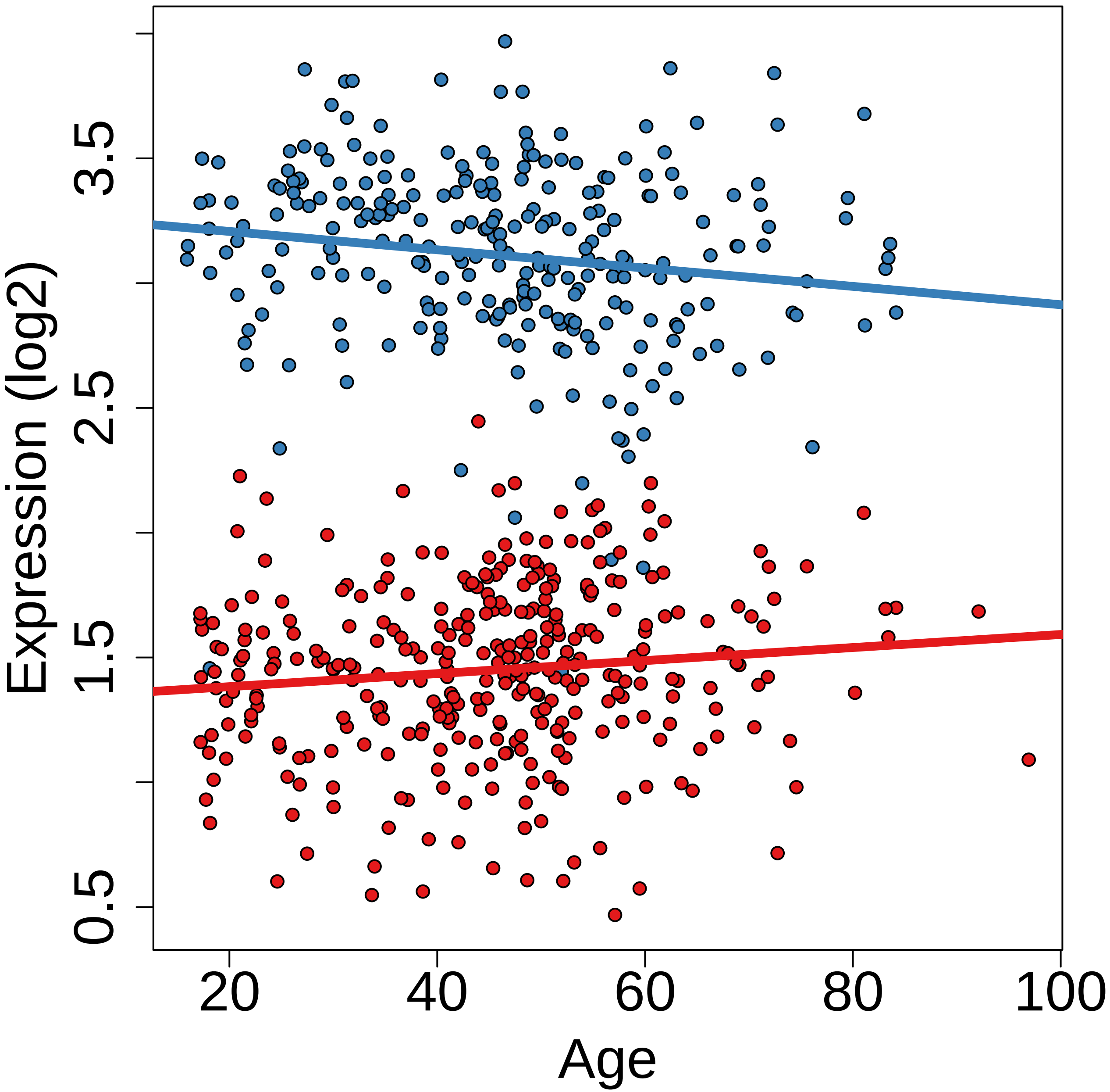
Expression (log2)
0.5
1.5
2.5
3.5
20
40
60
80
100
Age

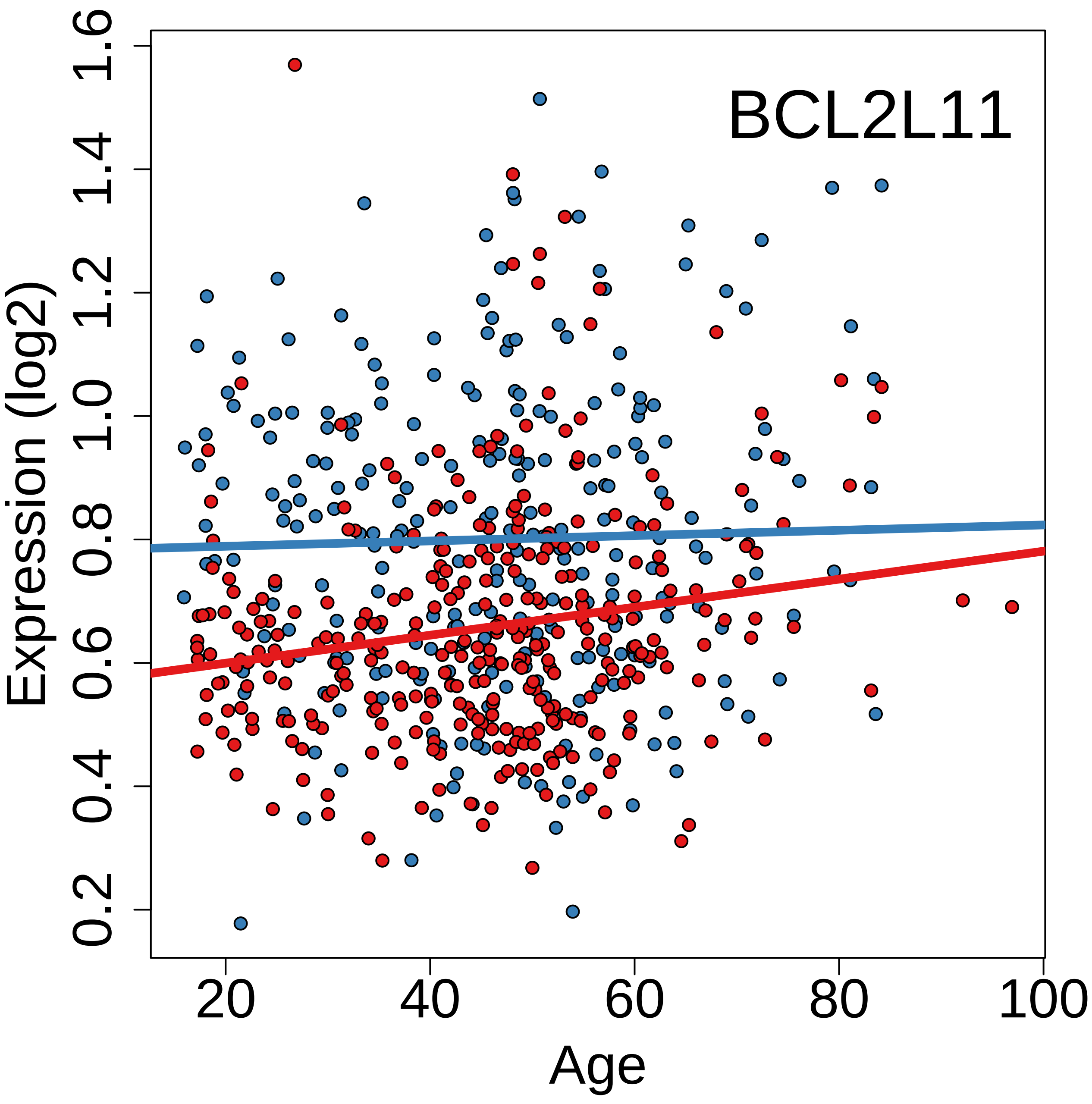

BCL2L11
Expression (log2)
0.2
0.4
0.6
0.8
1.0
1.2
1.4
1.6
20
40
60
80
100
Age

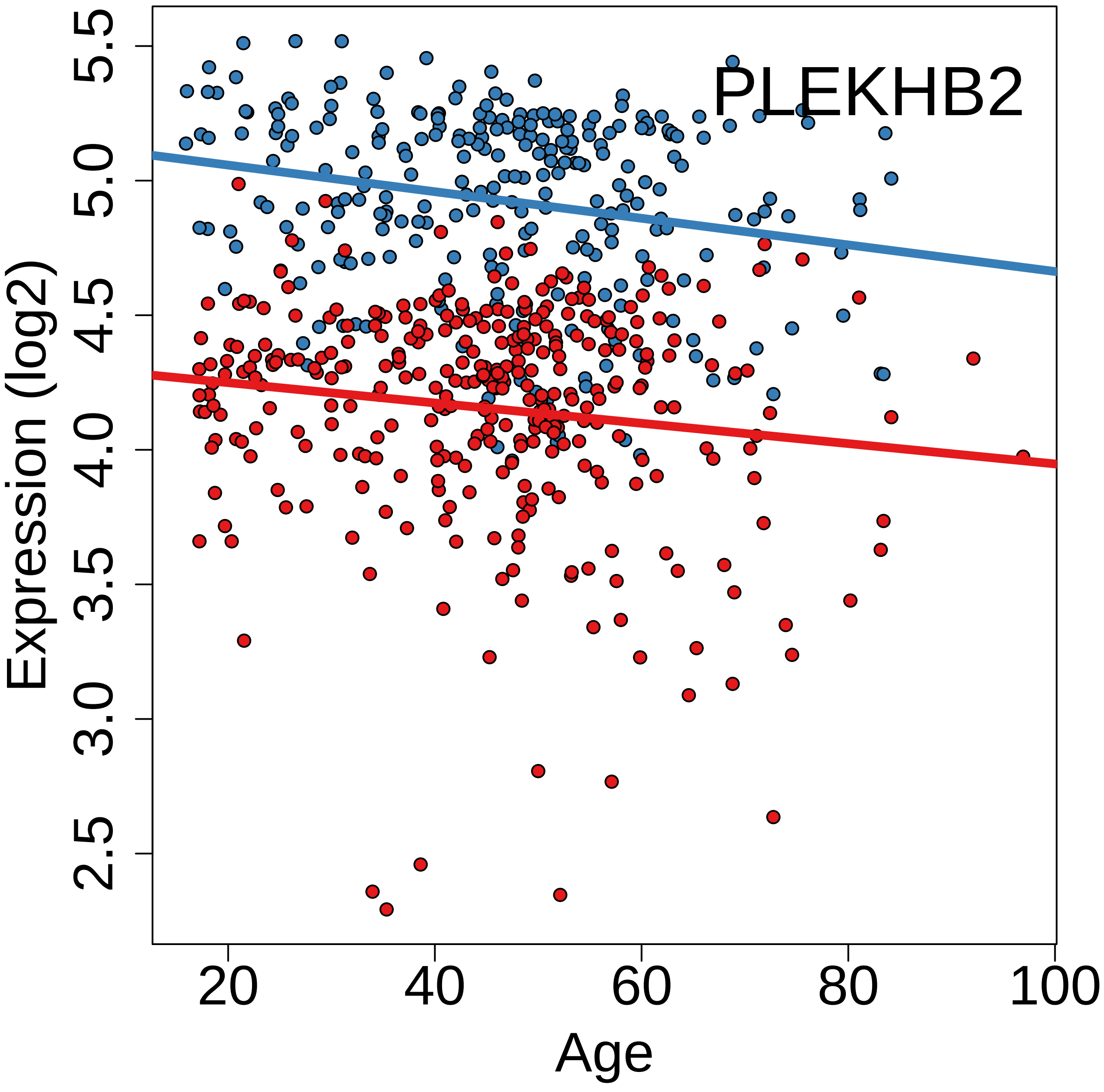
PLEKHB2
Expression (log2)
5.5
5.0
4.5
4.0
3.5
3.0
2.5
Age
20
40
60
80
100

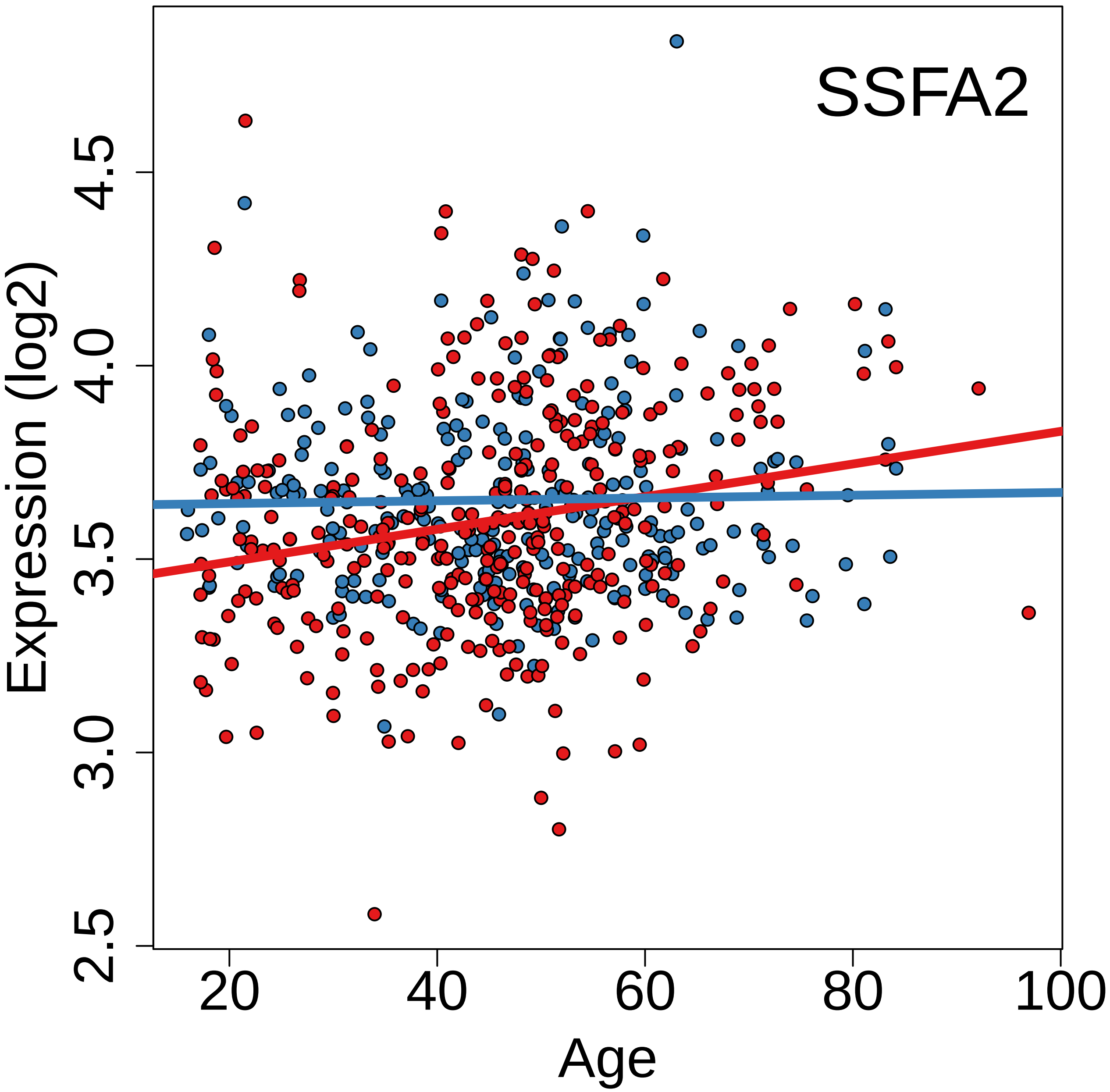
SSFA2
Expression (log2)
2.5
3.0
3.5
4.0
4.5
20
40
60
80
100
Age

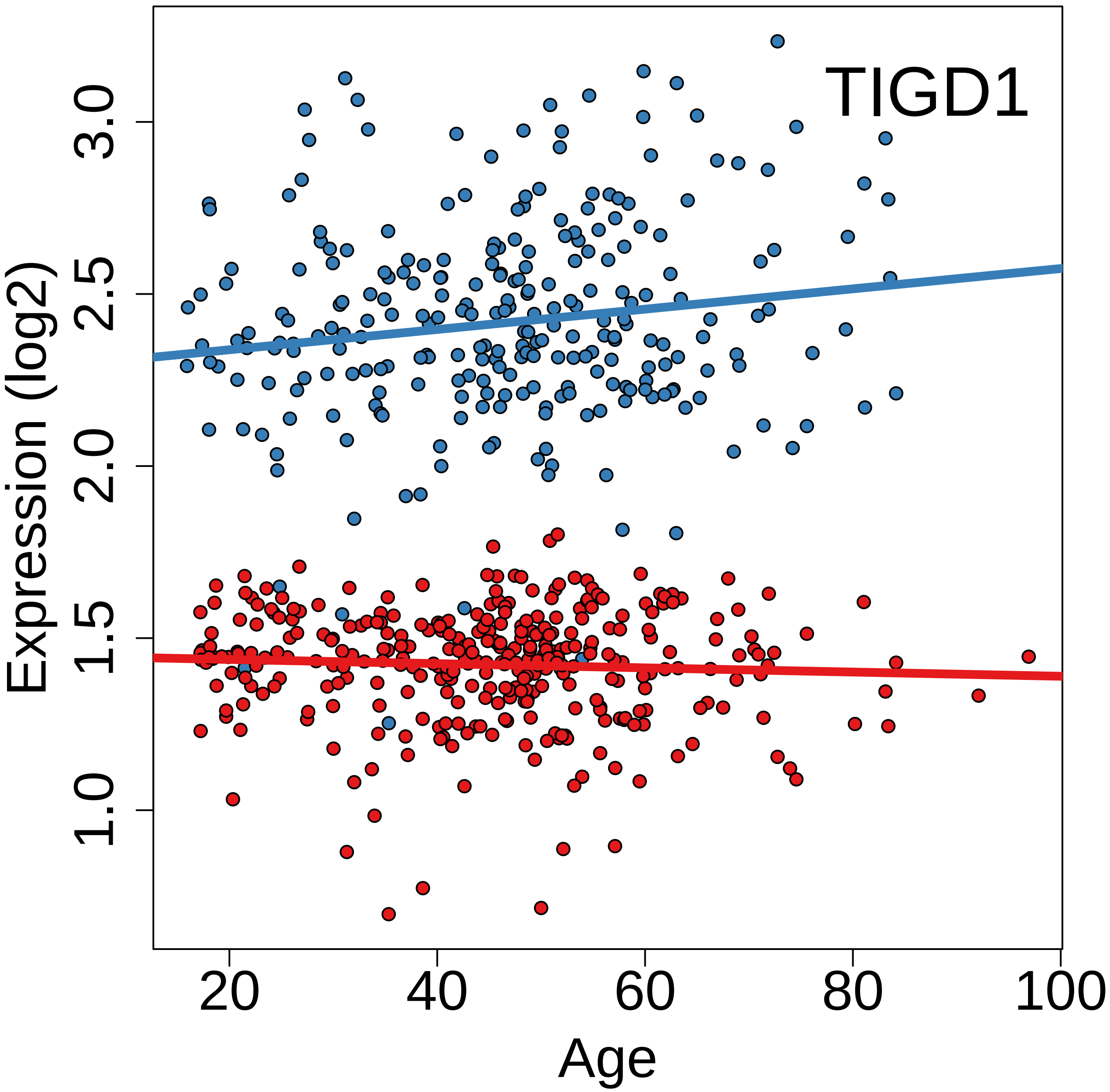
TIGD1
Expression (log2)
1.0
1.5
2.0
2.5
3.0
20
40
60
80
100
Age

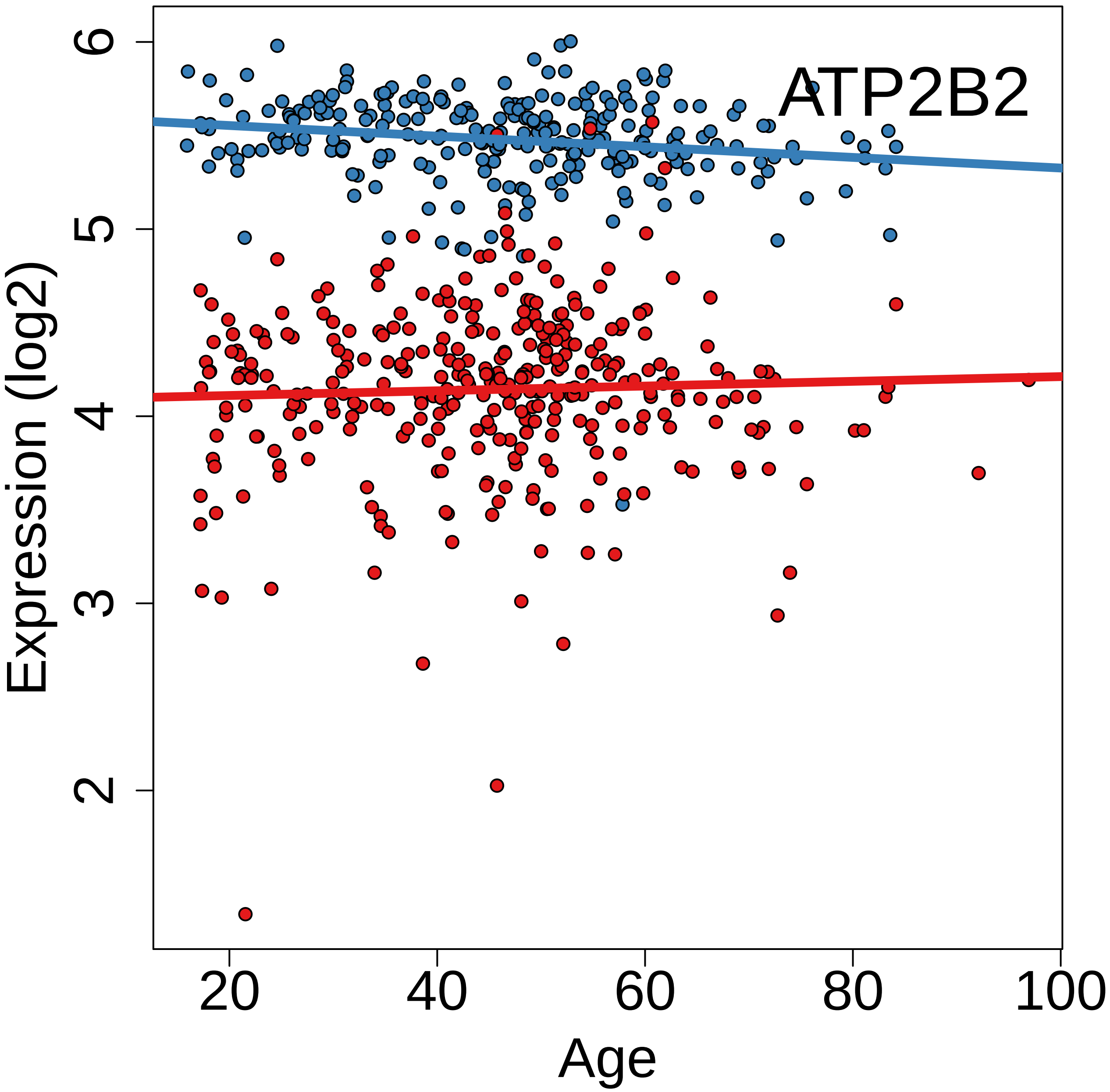
ATP2B2
Expression (log2)
Age
2
3
4
5
6
20
40
60
80
100

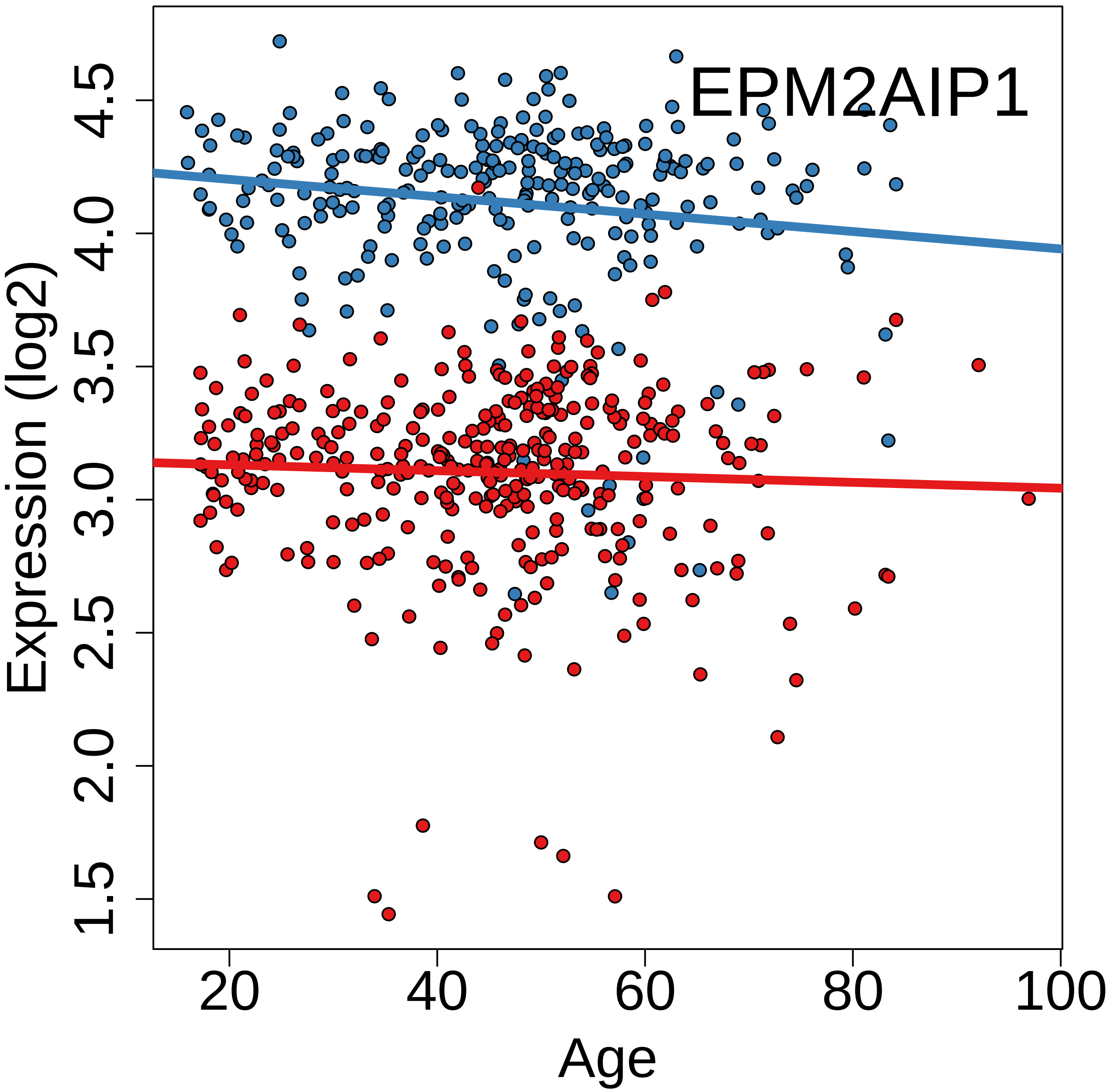
EPM2AIP1
Expression (log2)
1.5
2.0
2.5
3.0
3.5
4.0
4.5
20
40
60
80
100
Age

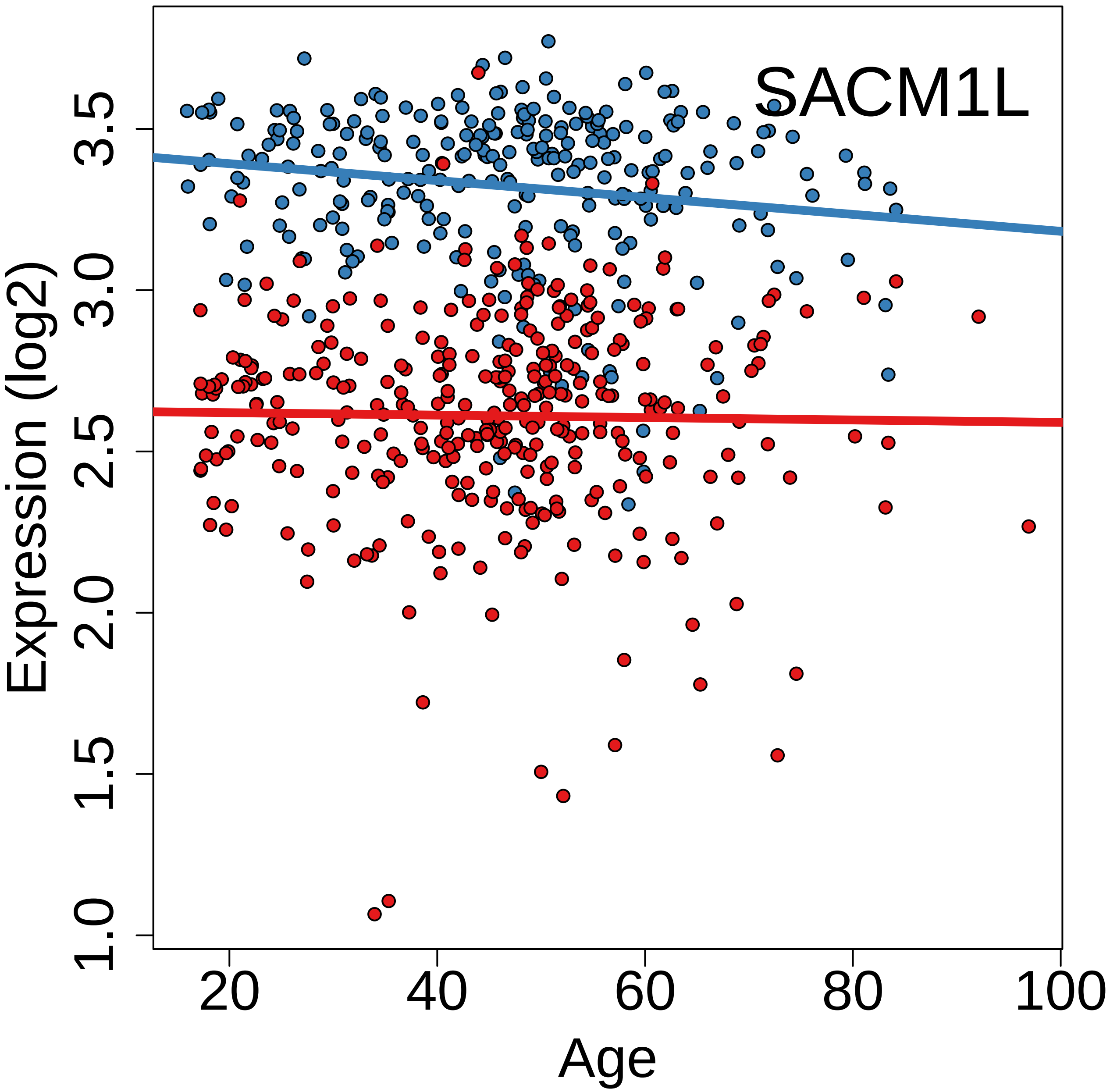
SACM1L
Expression (log2)
Age
1.0
1.5
2.0
2.5
3.0
3.5
20
40
60
80
100

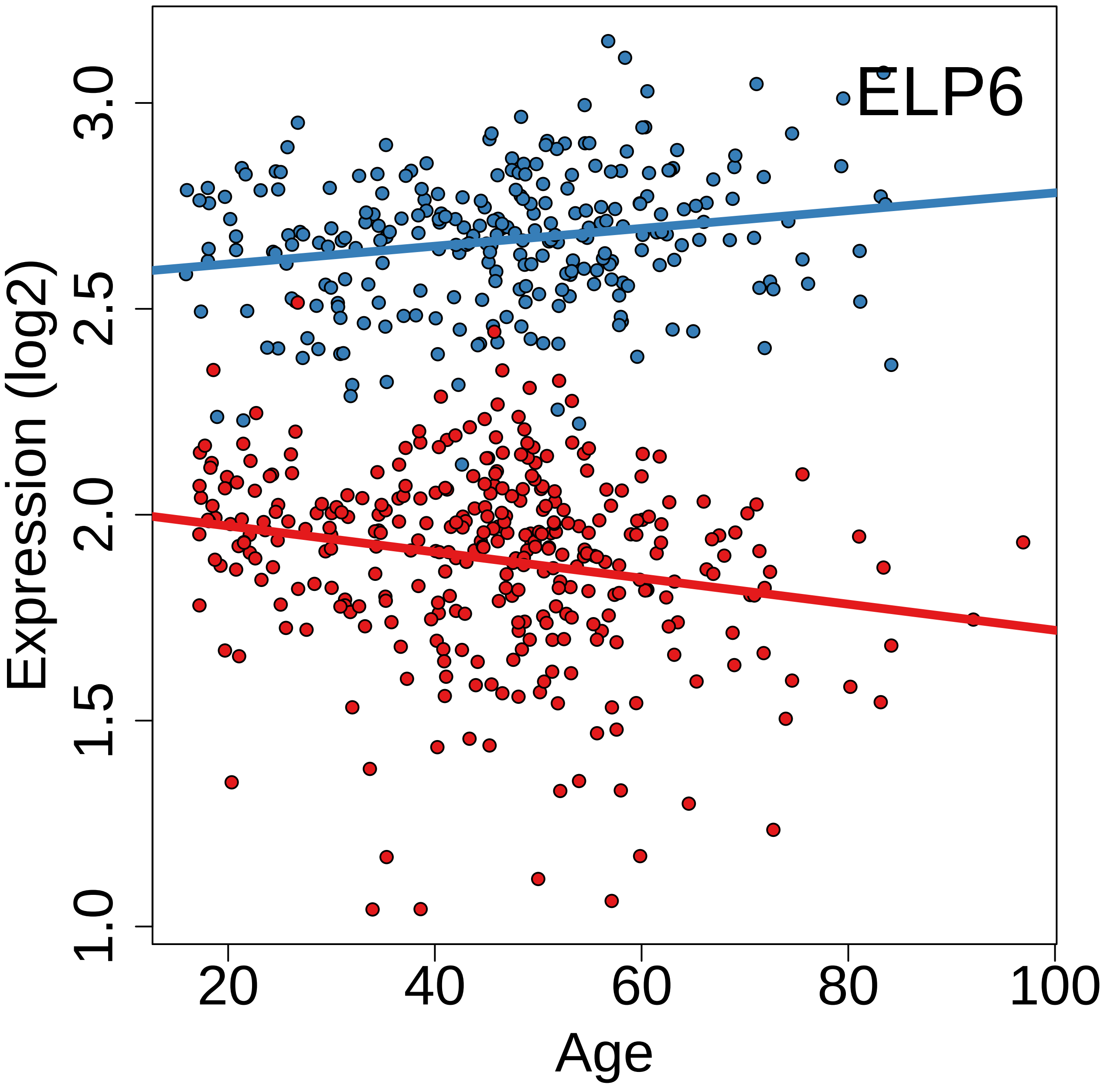
ELP6
Expression (log2)
1.0
1.5
2.0
2.5
3.0
20
40
60
80
100
Age

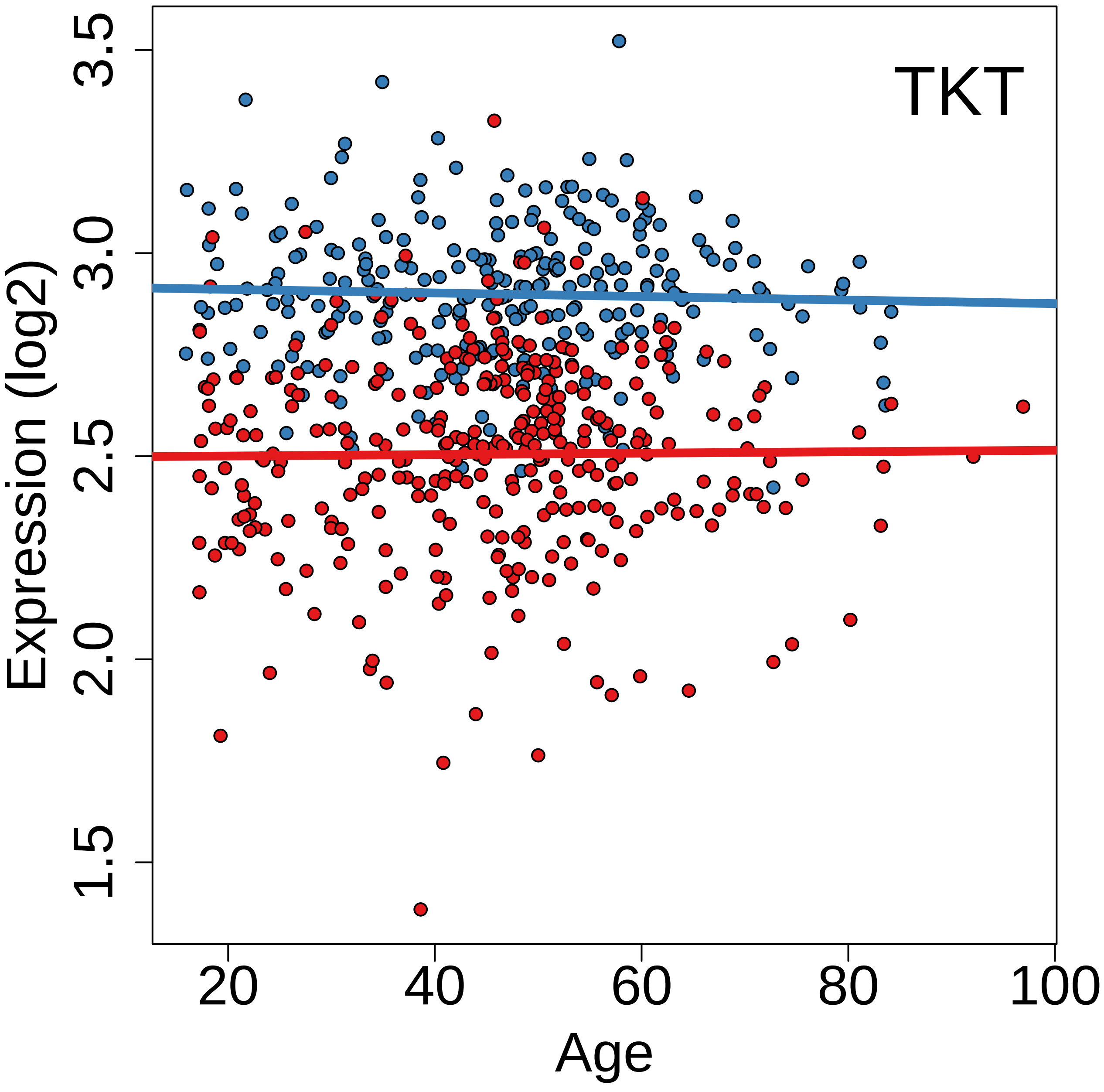
TKT
Expression (log2)
3.5
3.0
2.5
2.0
1.5
20
40
60
80
100
Age

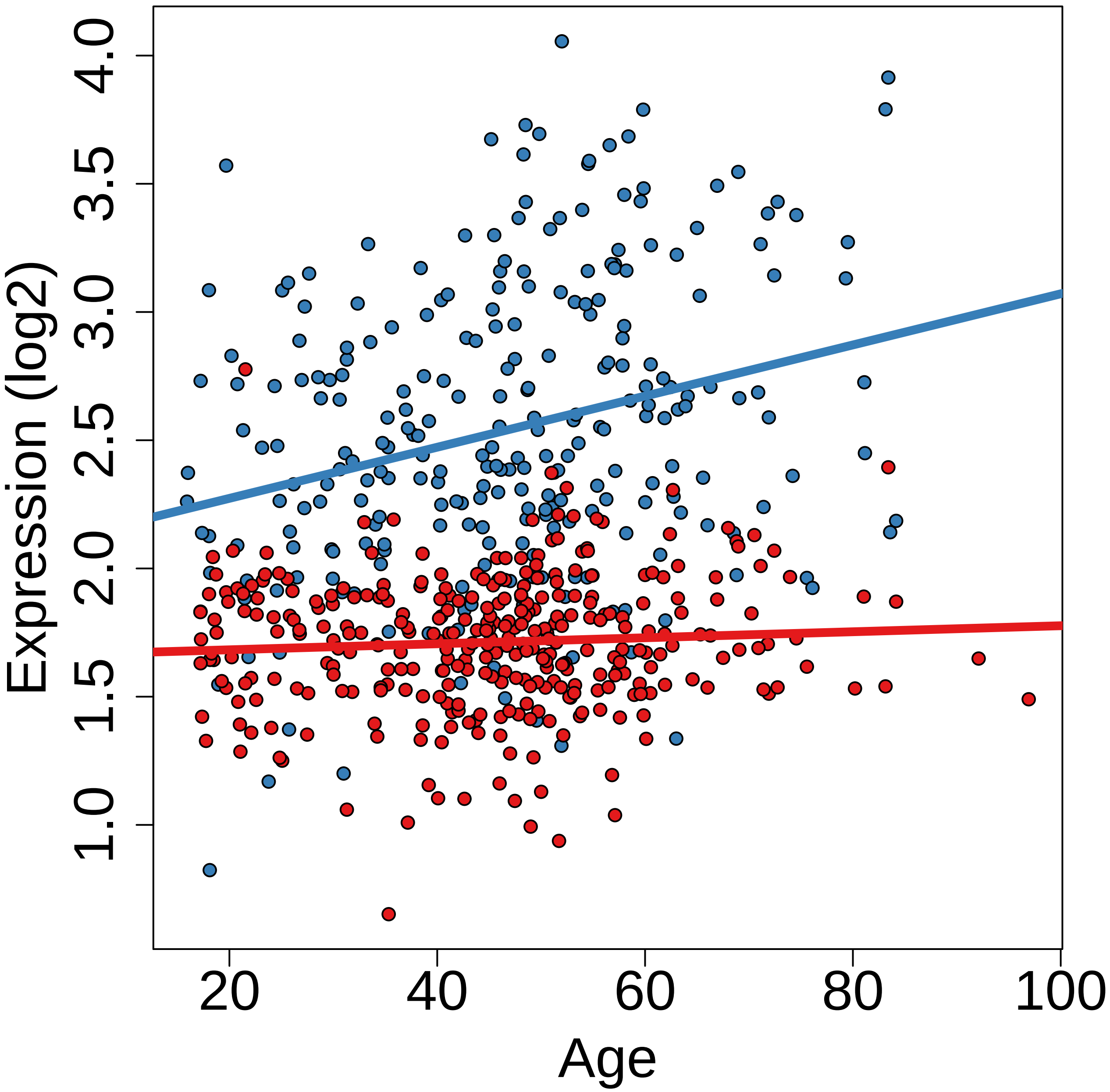
Expression (log2)
1.0
1.5
2.0
2.5
3.0
3.5
4.0
20
40
60
80
100
Age

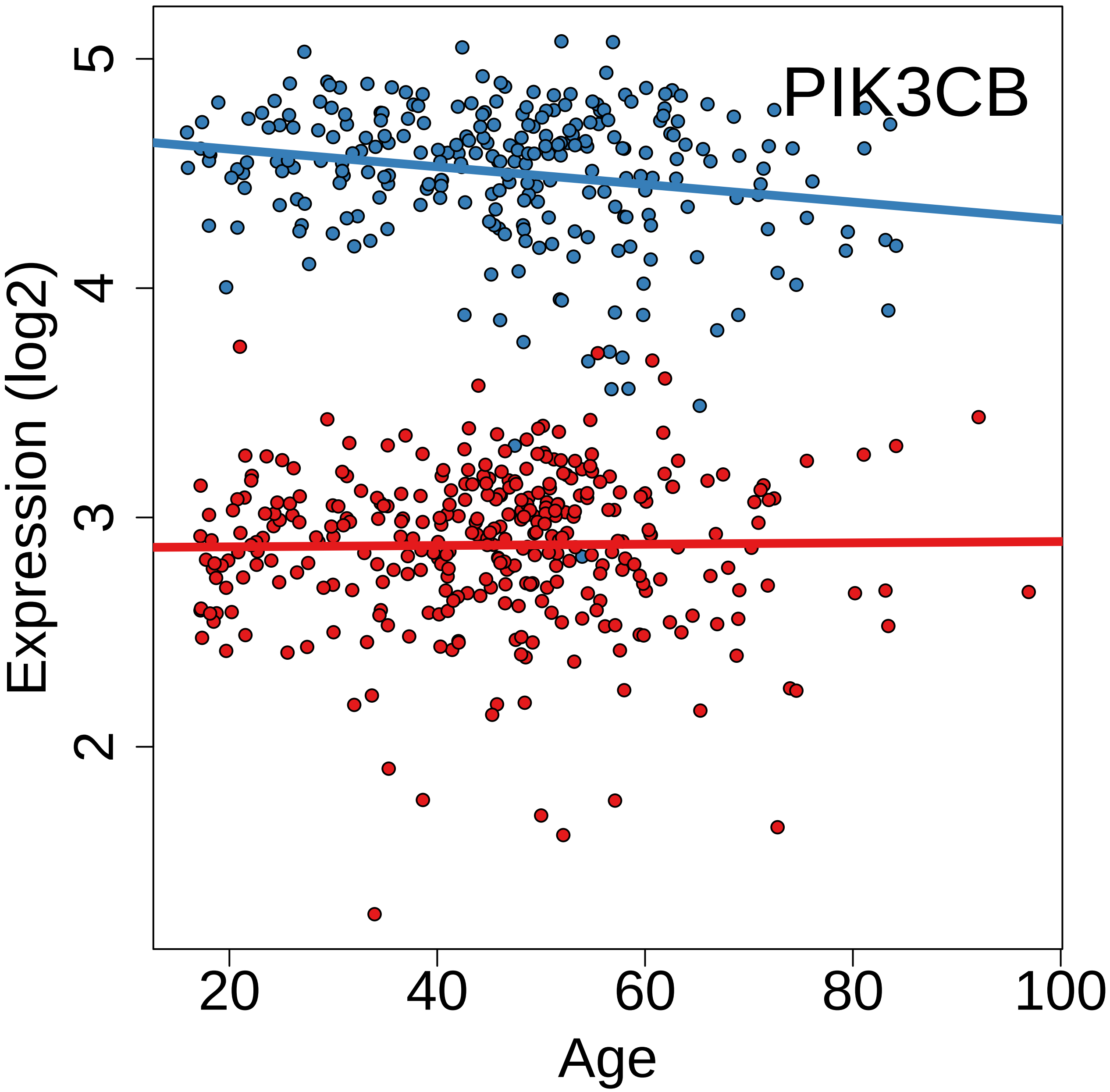

PIK3CB
Expression (log2)
Age
2
3
4
5
20
40
60
80
100

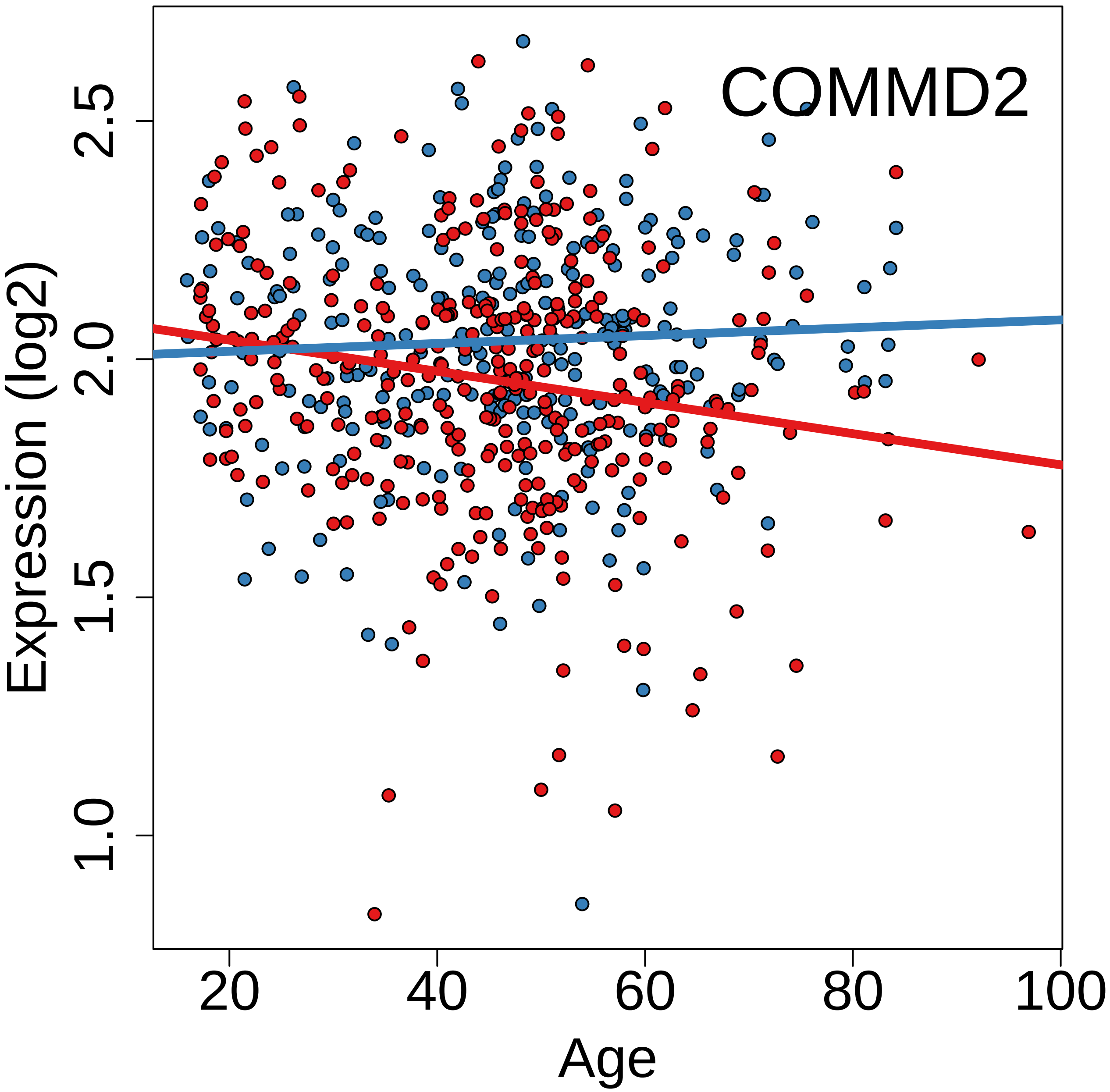
COMMD2
Expression (log2)
2.5
2.0
1.5
1.0
20
40
60
80
100
Age

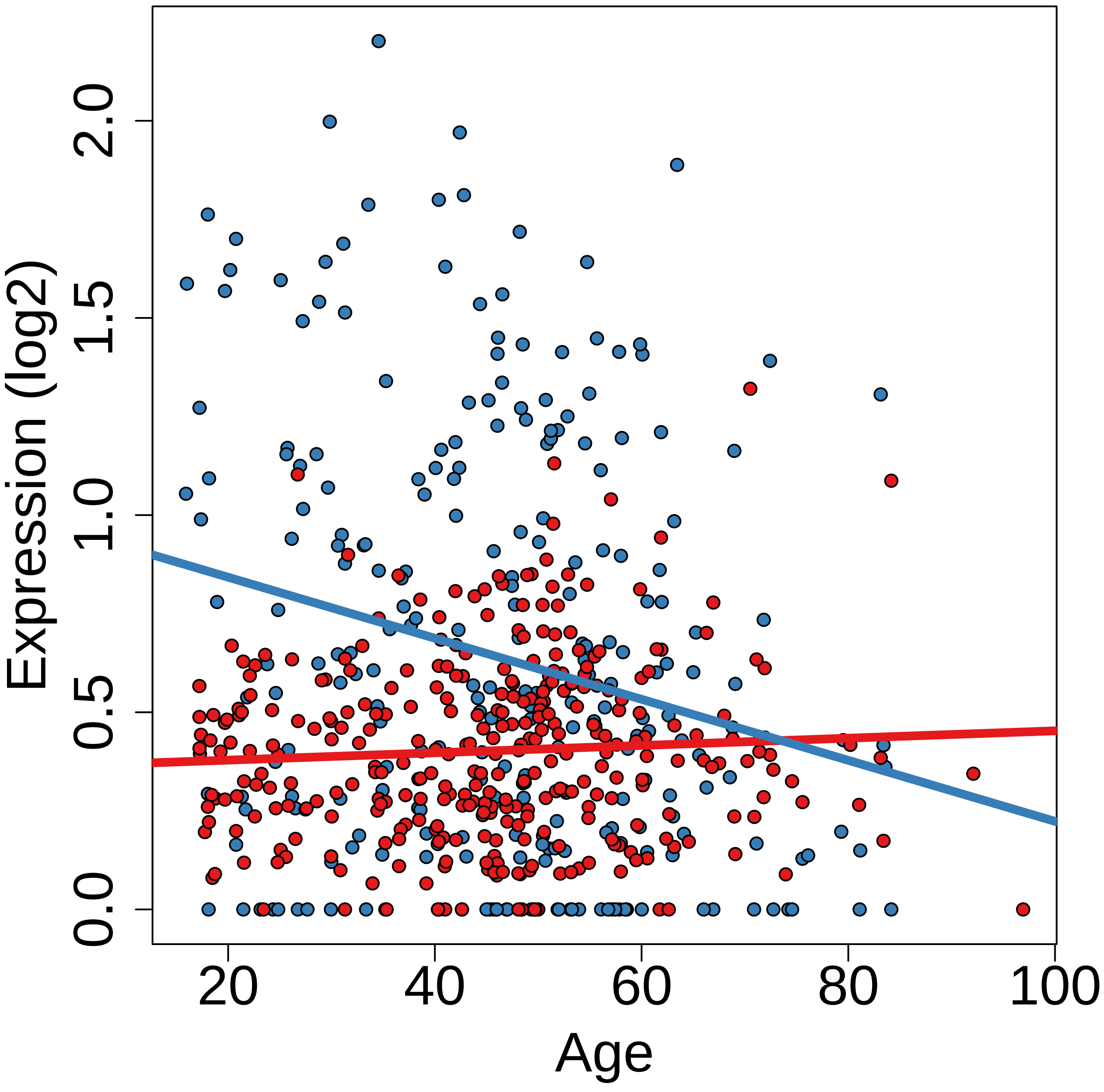

Expression (log2)
0.0
0.5
1.0
1.5
2.0
20
40
60
80
100
Age

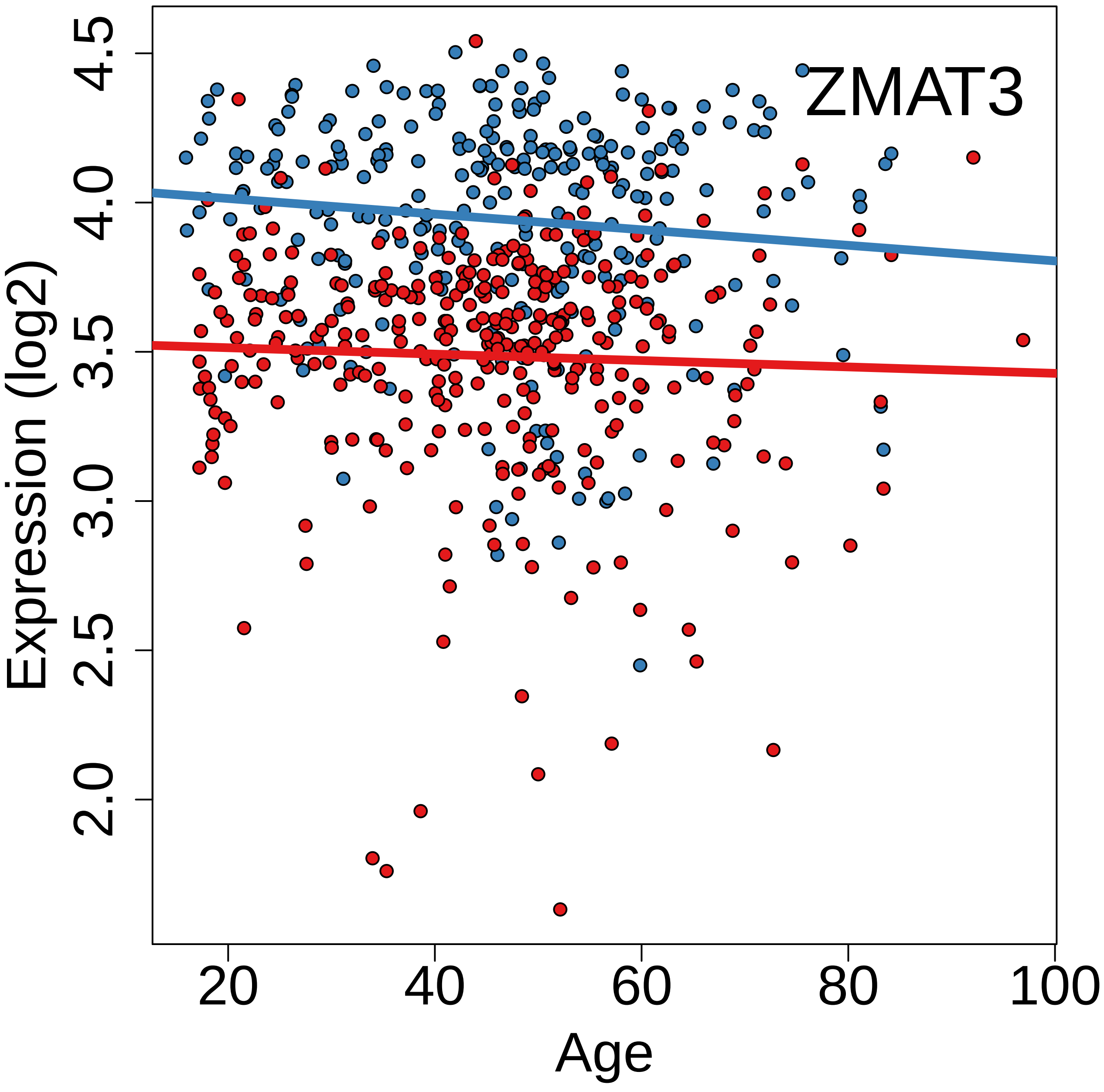
ZMAT3
Expression (log2)
Age
2.0
2.5
3.0
3.5
4.0
4.5
20
40
60
80
100

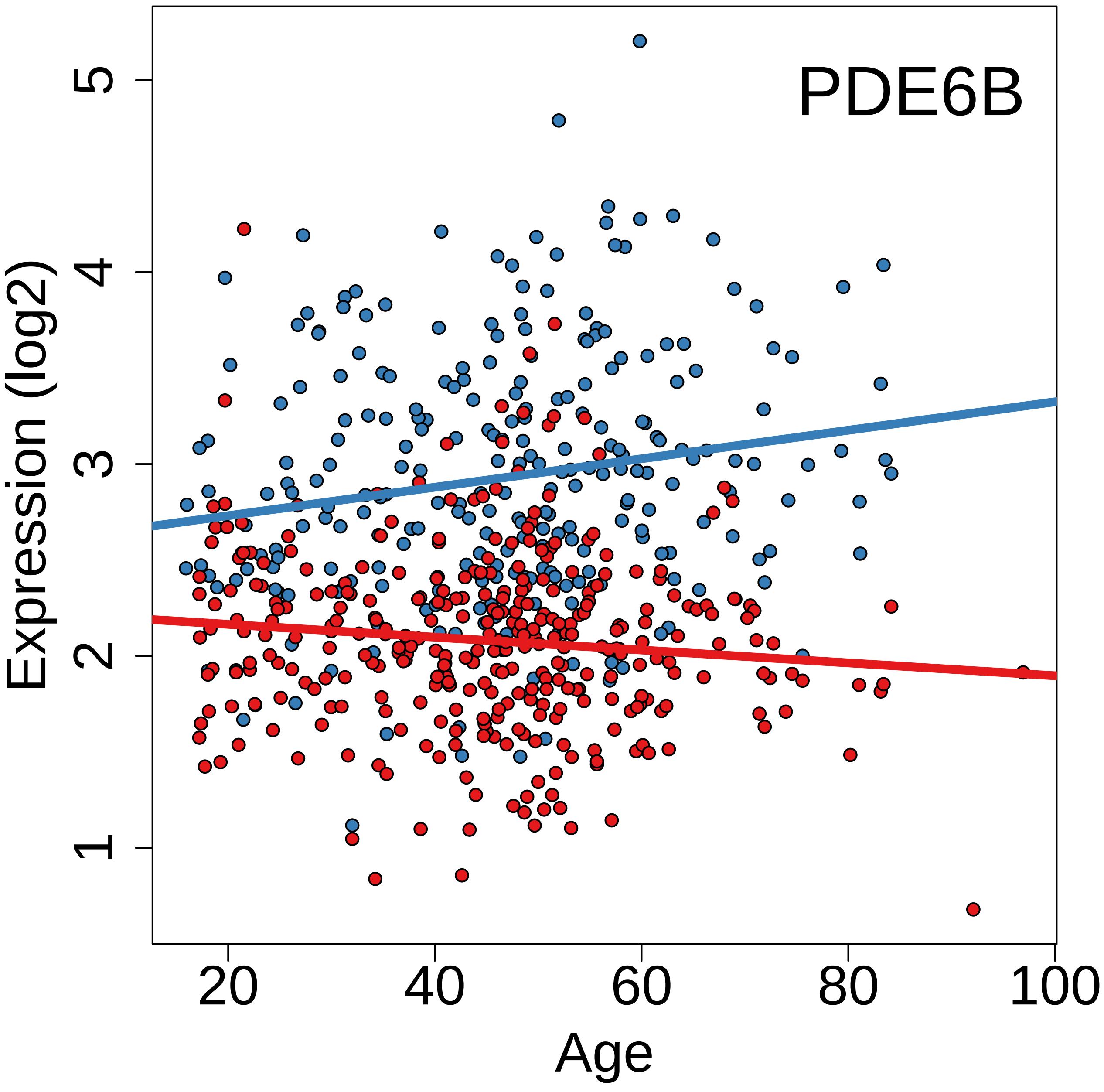
PDE6B
Expression (log2)
1
2
3
4
5
20
40
60
80
100
Age

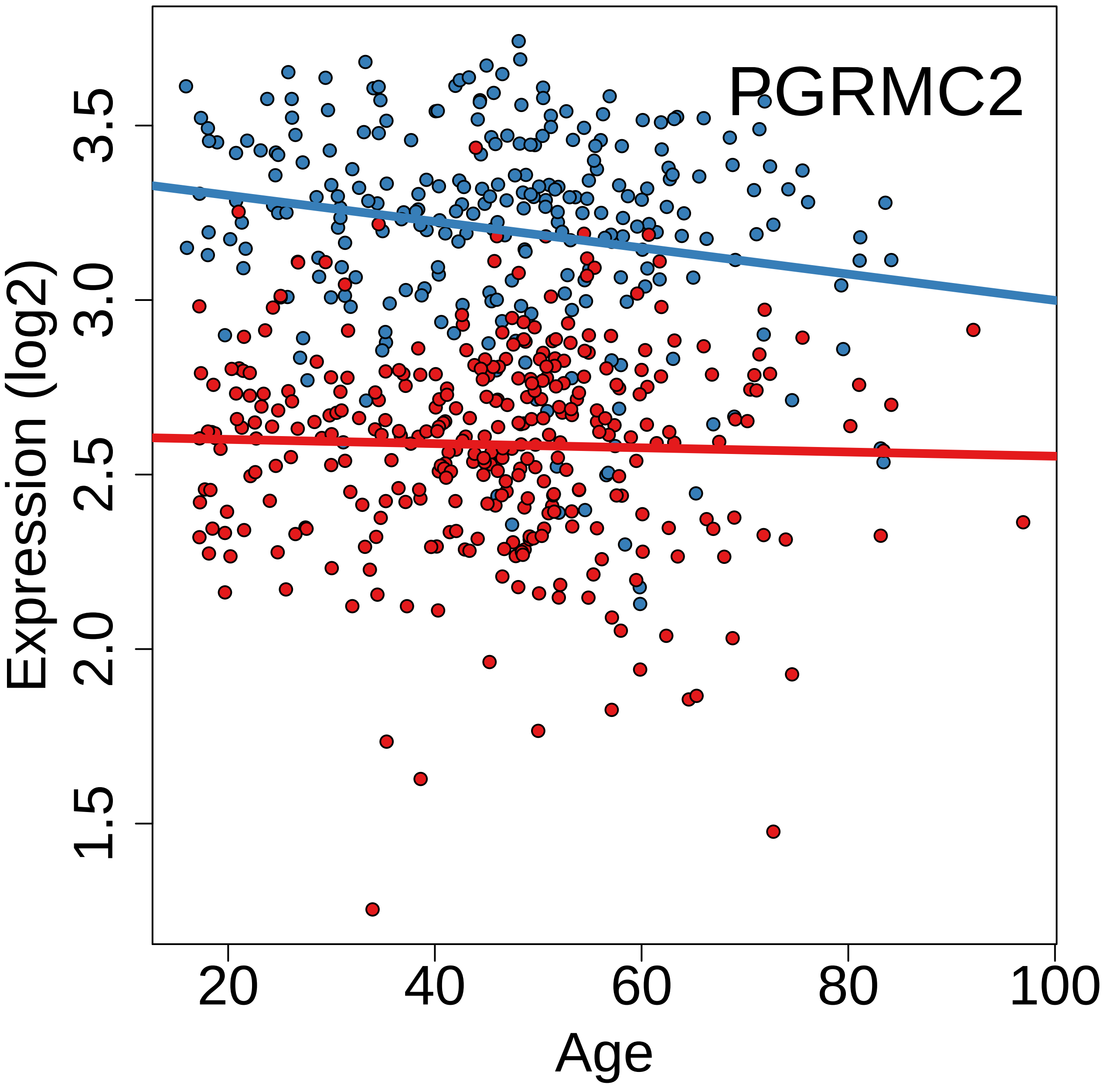

PGRMC2
Expression (log2)
1.5
2.0
2.5
3.0
3.5
20
40
60
80
100
Age

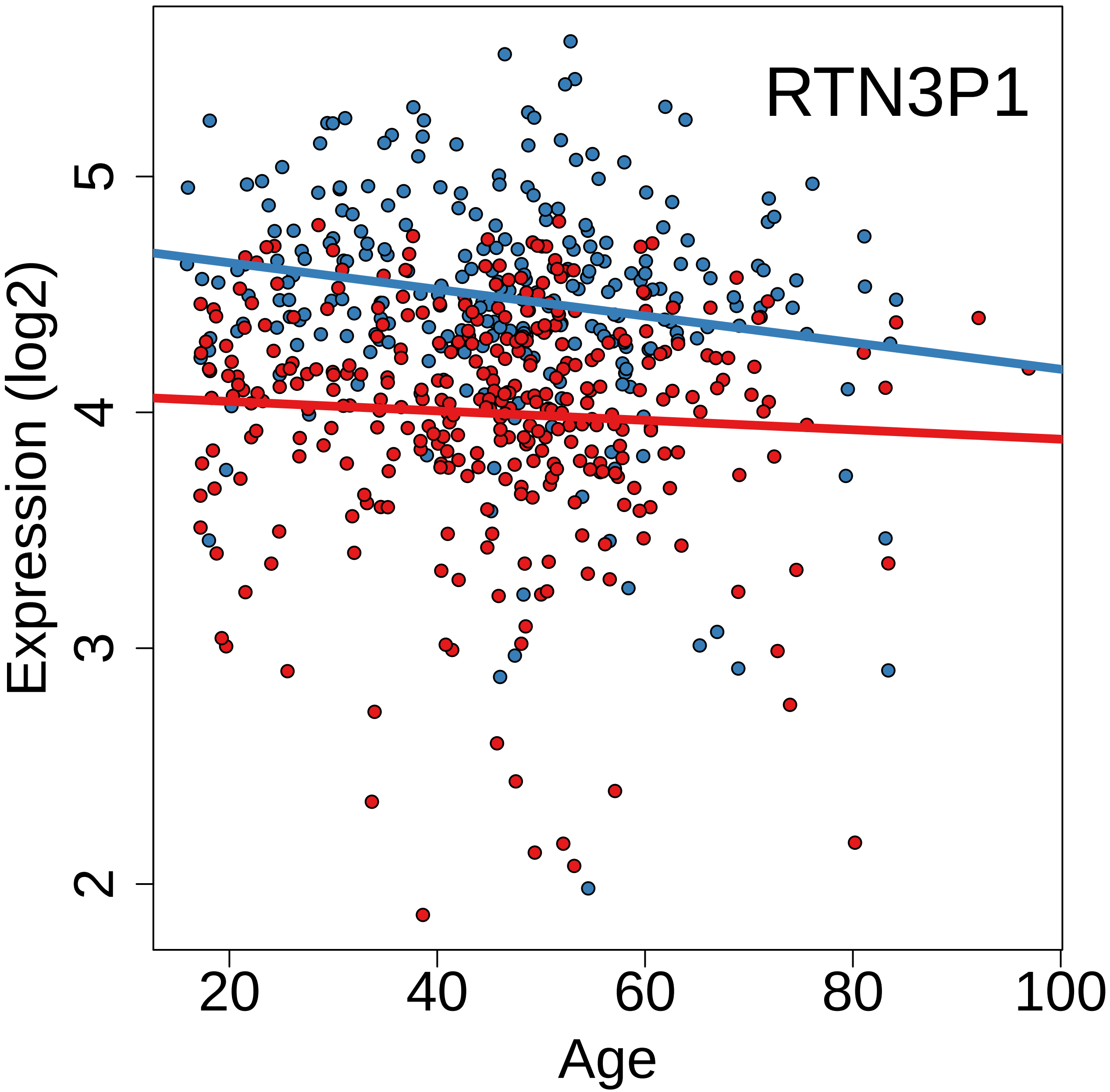

RTN3P1
Expression (log2)
2
3
4
5
20
40
60
80
100
Age

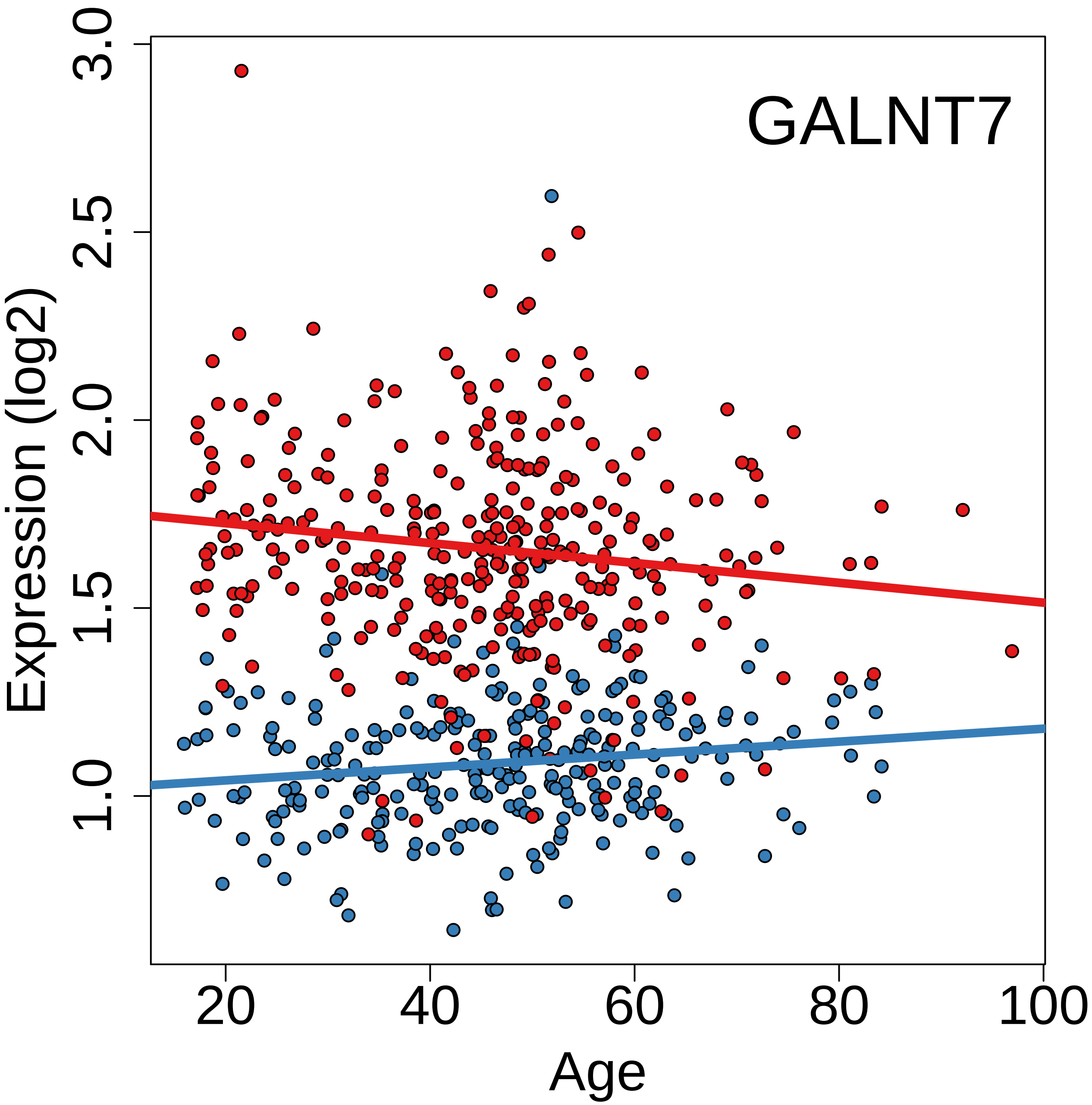

GALNT7
Expression (log2)
Age
3.0
2.5
2.0
1.5
1.0
20
40
60
80
100

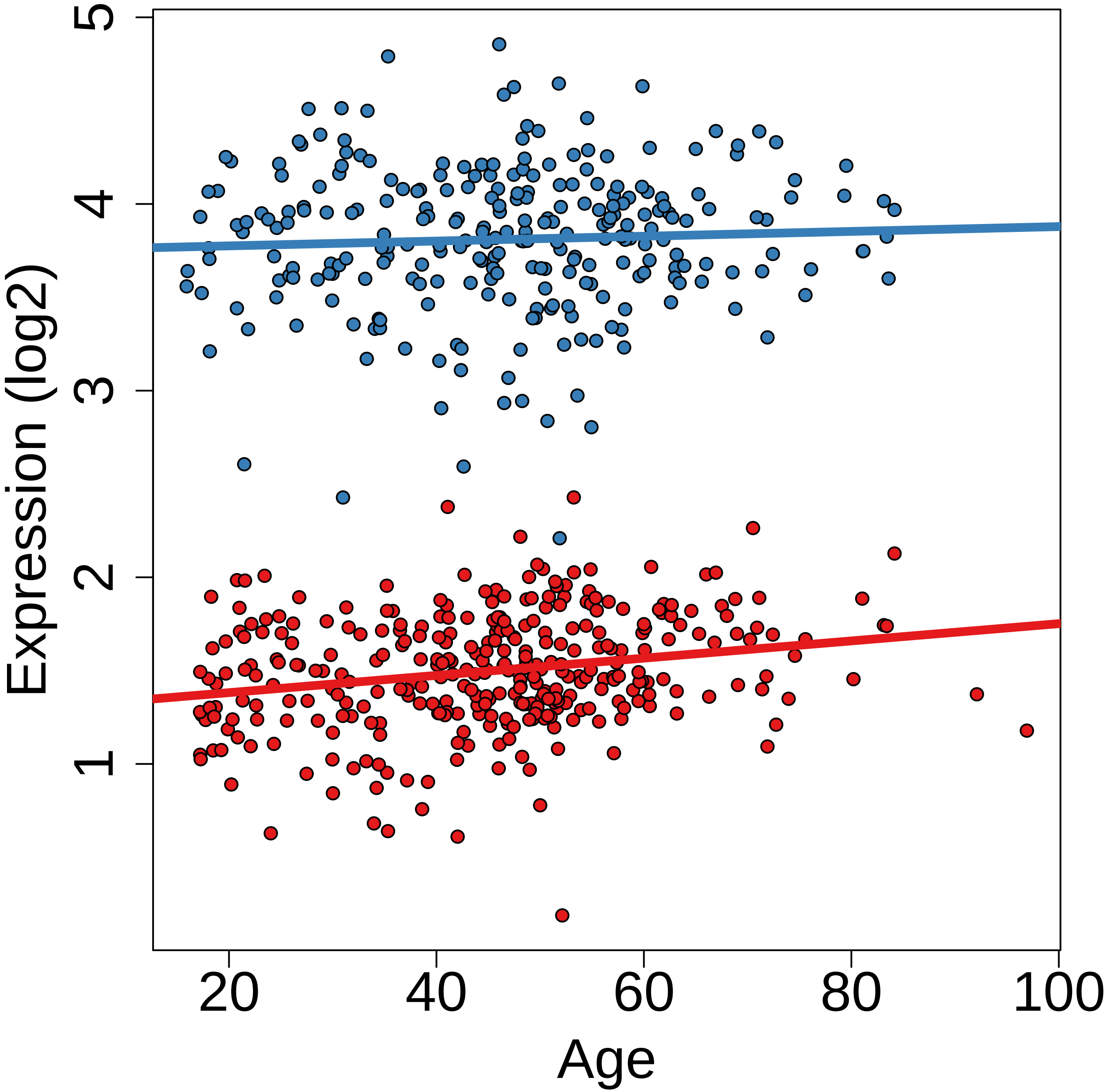

Expression (log2)
1
2
3
4
5
20
40
60
80
100
Age

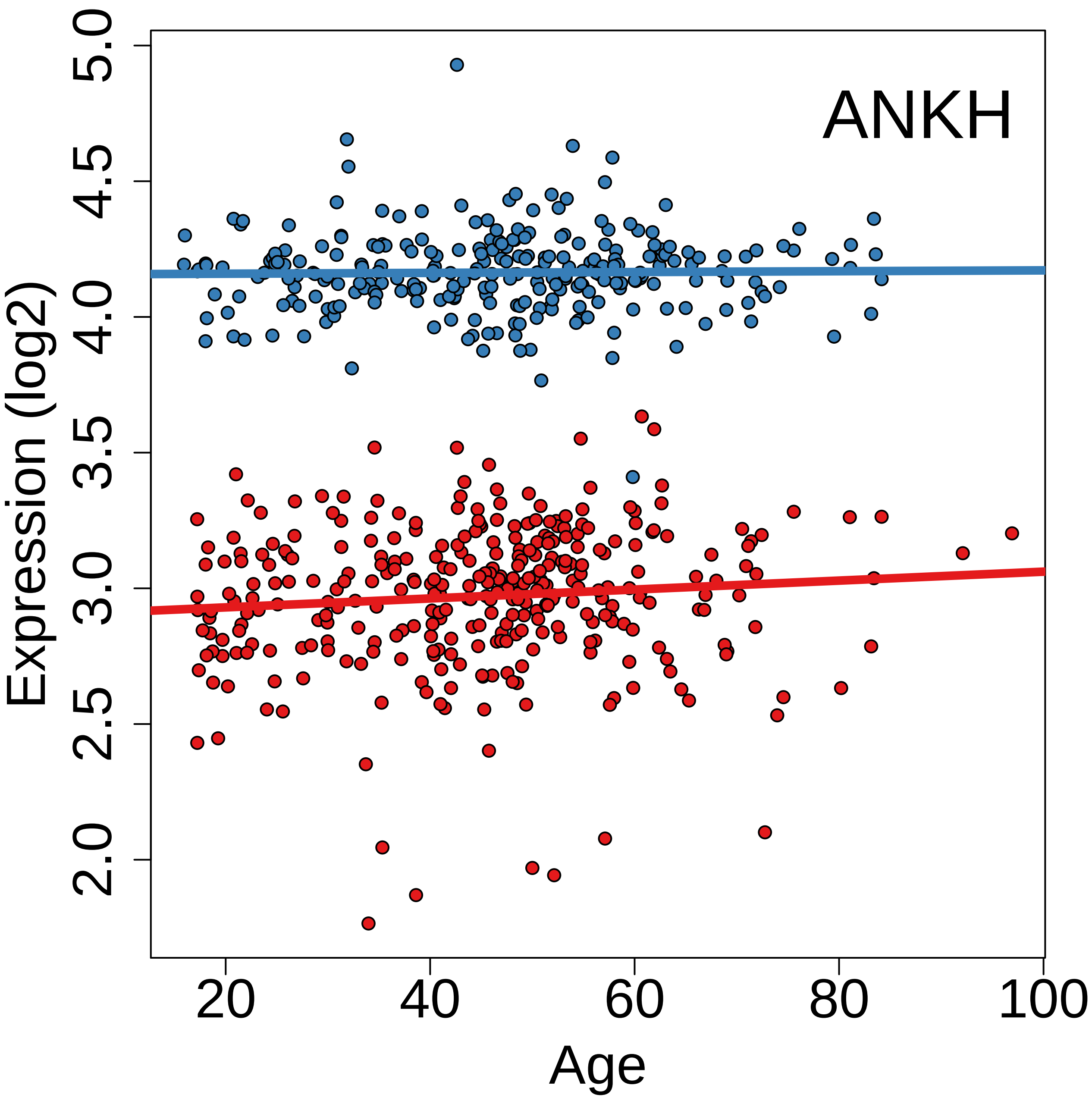

ANKH
Expression (log2)
Age
2.0
2.5
3.0
3.5
4.0
4.5
5.0
20
40
60
80
100

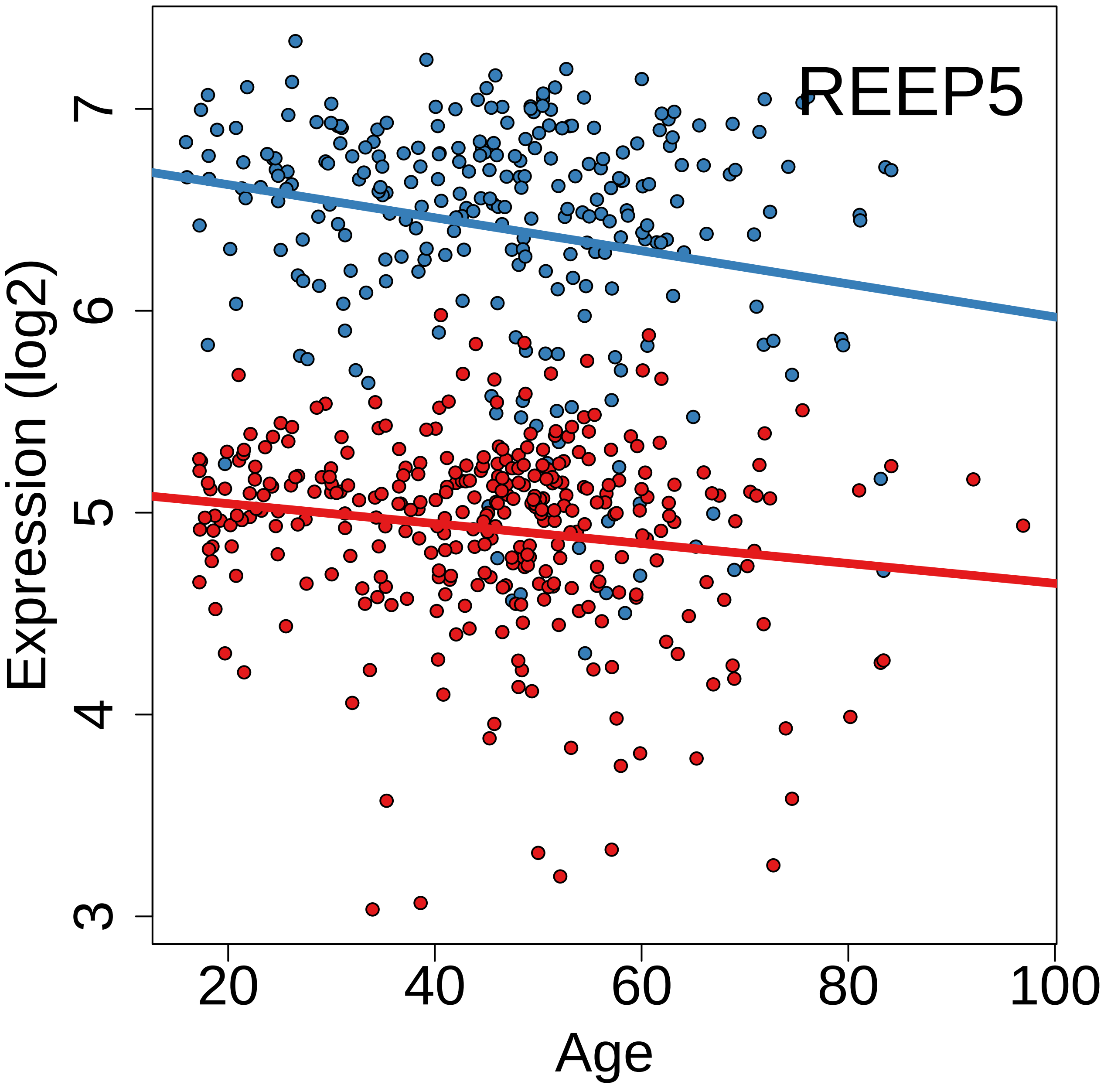
REEP5
Expression (log2)
Age
3
4
5
6
7
20
40
60
80
100

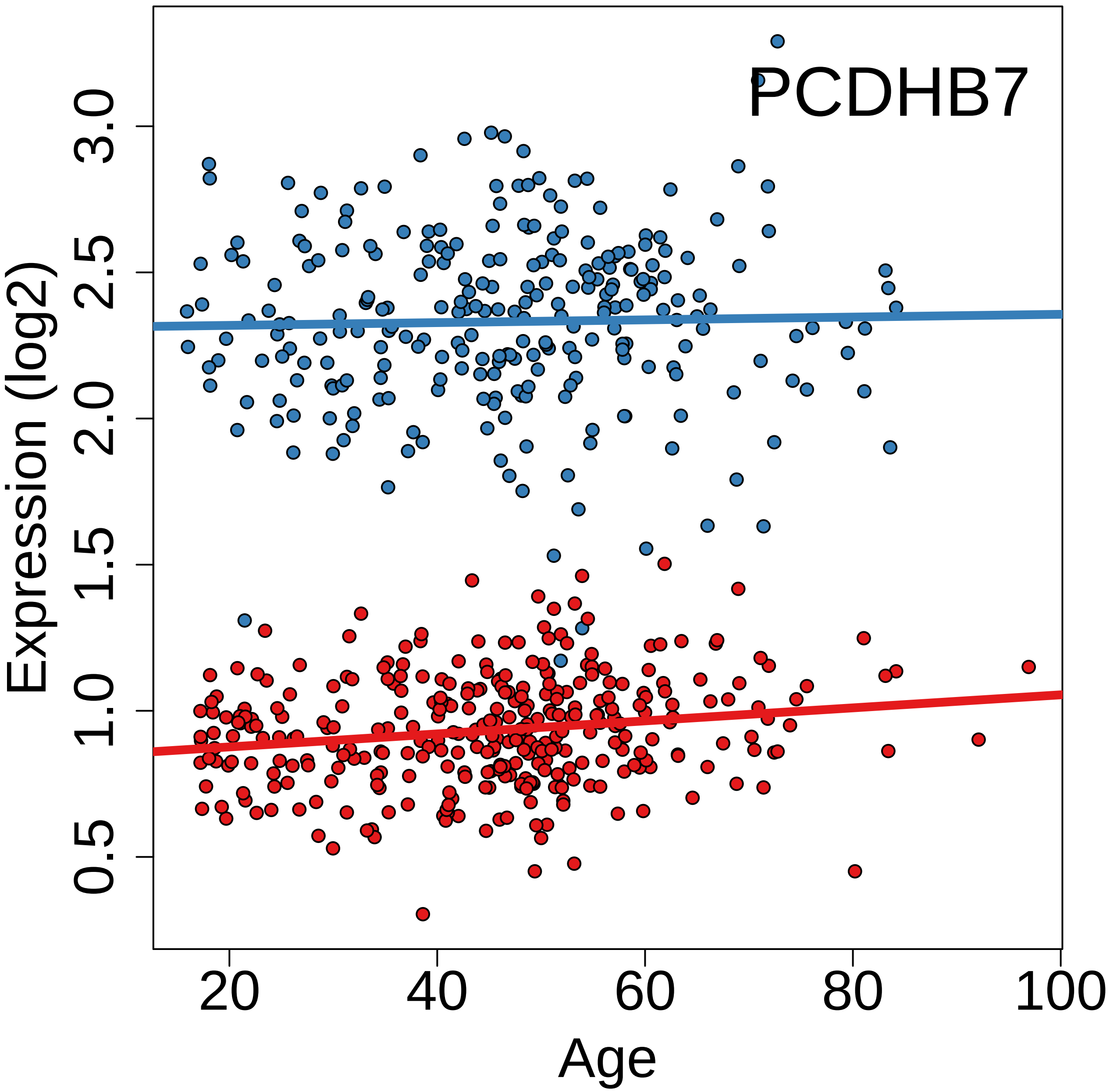
PCDHB7
Expression (log2)
0.5
1.0
1.5
2.0
2.5
3.0
20
40
60
80
100
Age

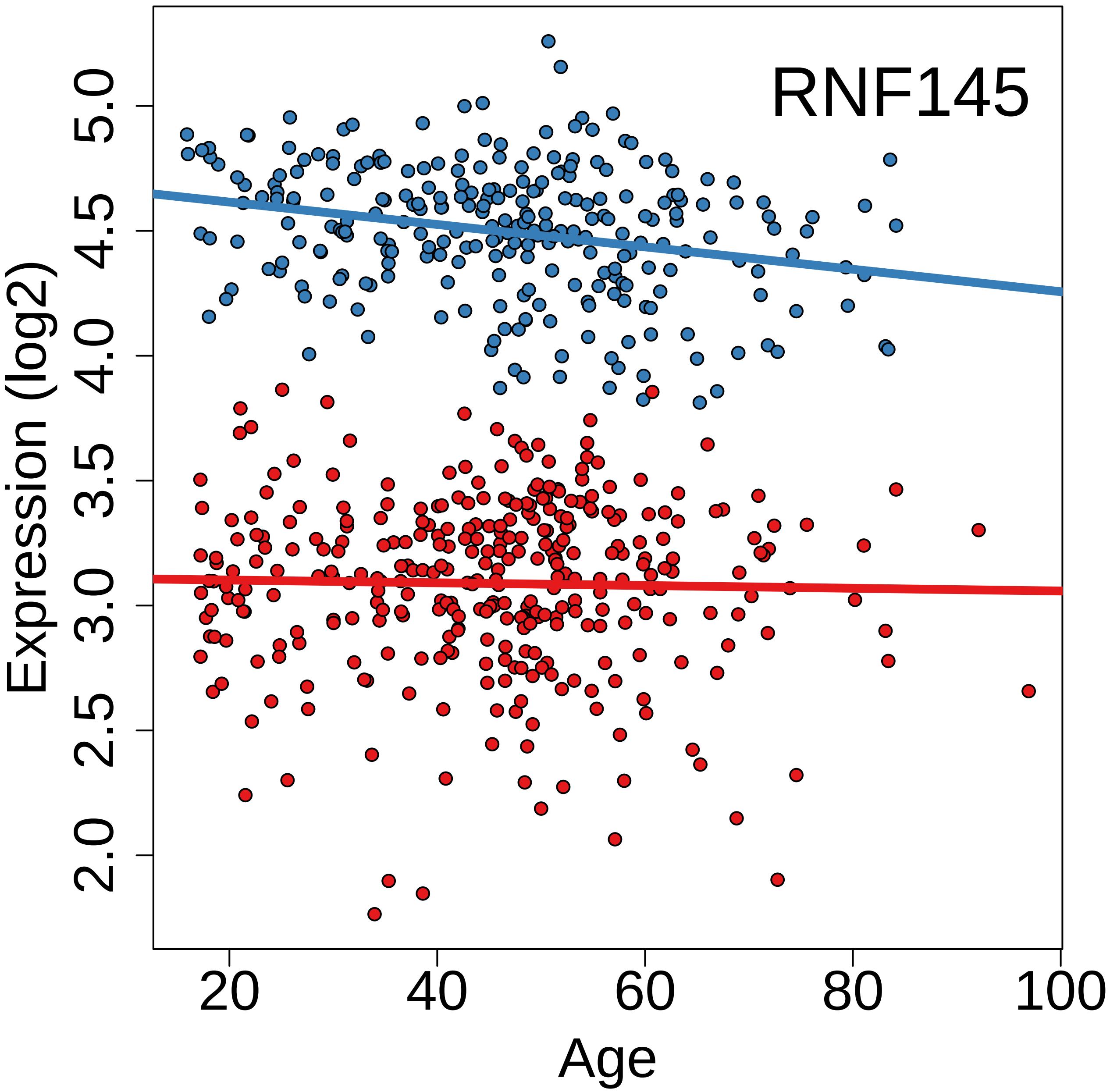

RNF145
Expression (log2)
2.0
2.5
3.0
3.5
4.0
4.5
5.0
20
40
60
80
100
Age

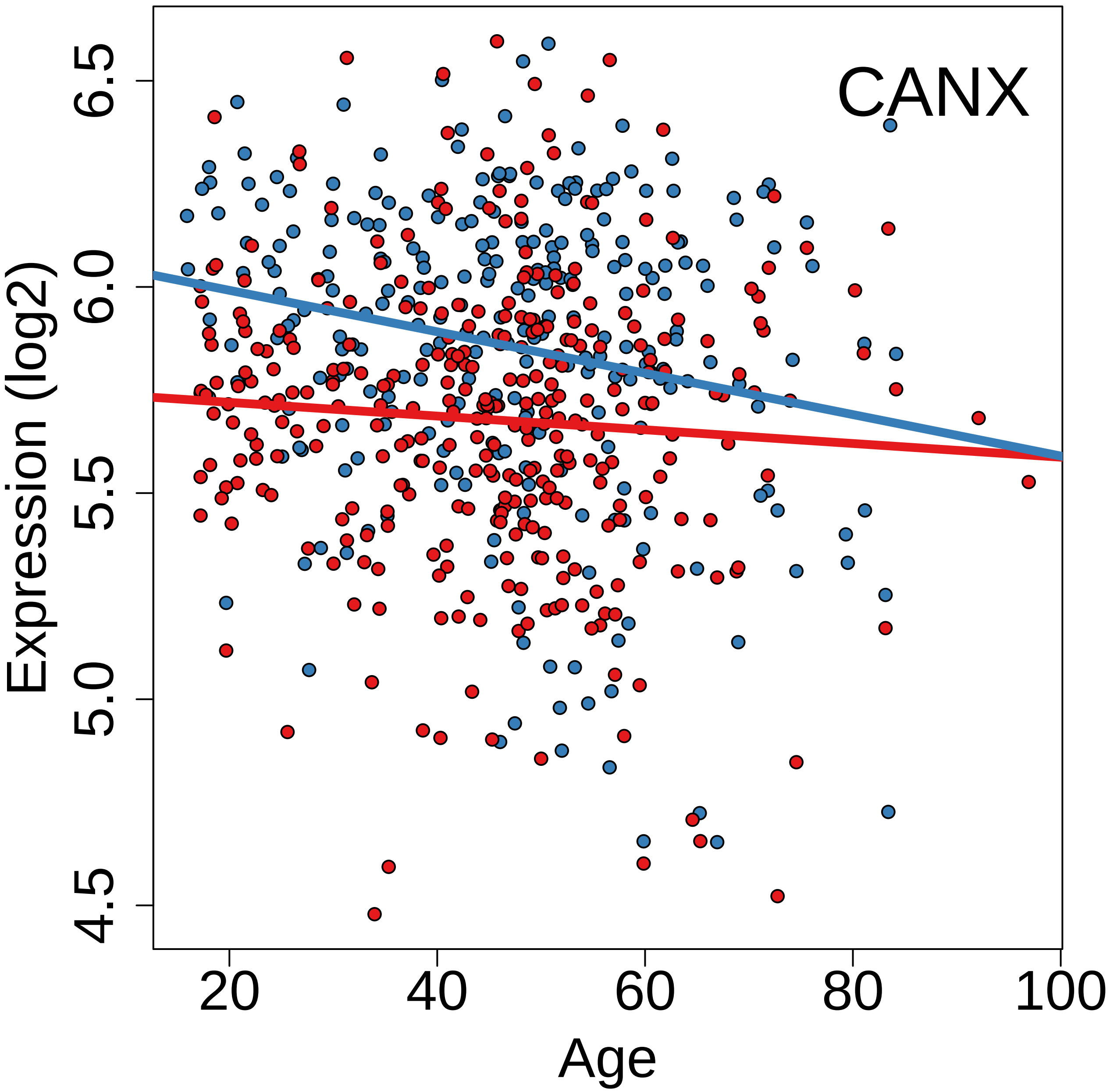
CANX
Expression (log2)
Age
6.5
6.0
5.5
5.0
4.5
20
40
60
80
100

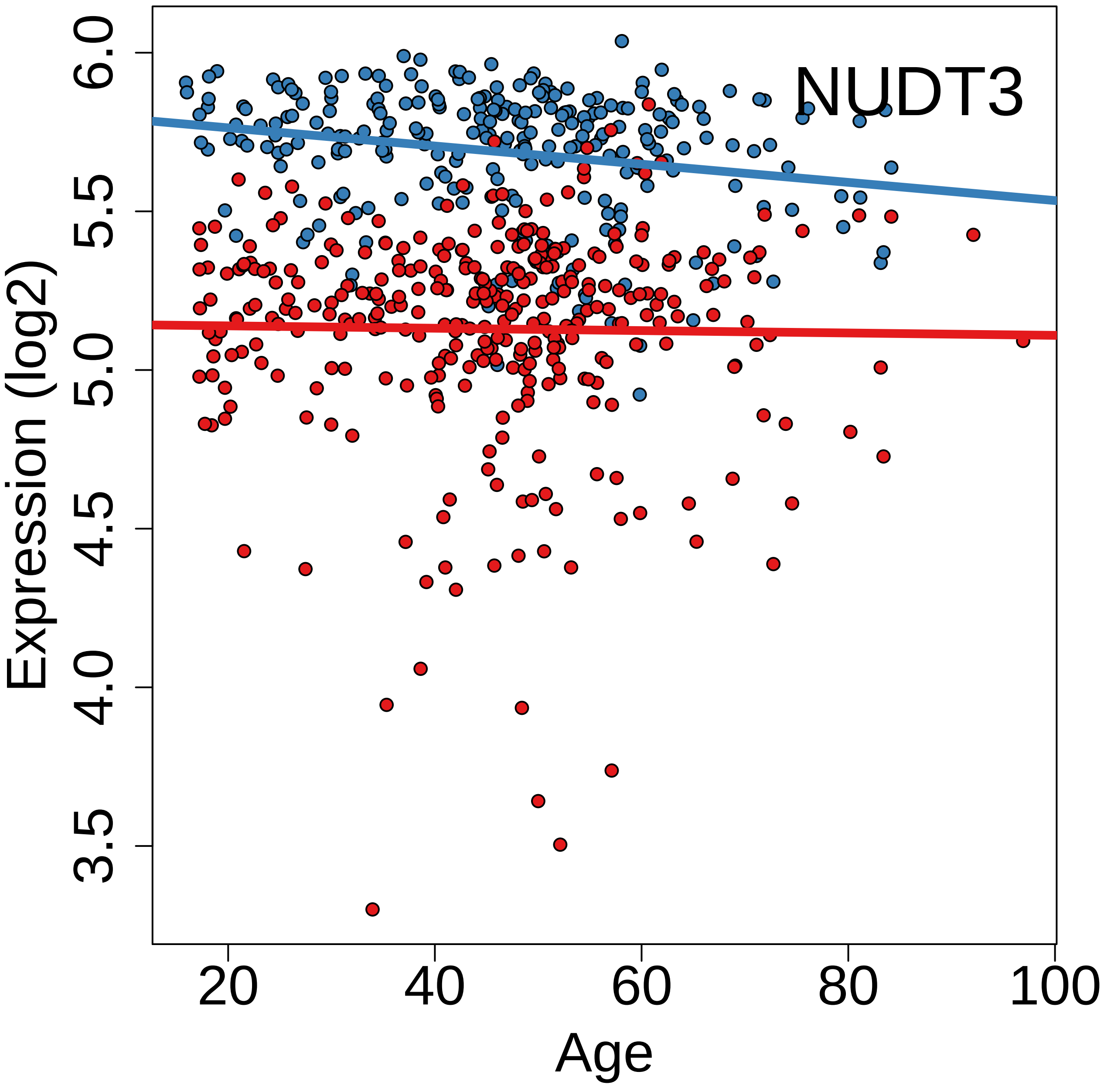
NUDT3
Expression (log2)
Age
6.0
5.5
5.0
4.5
4.0
3.5
20
40
60
80
100

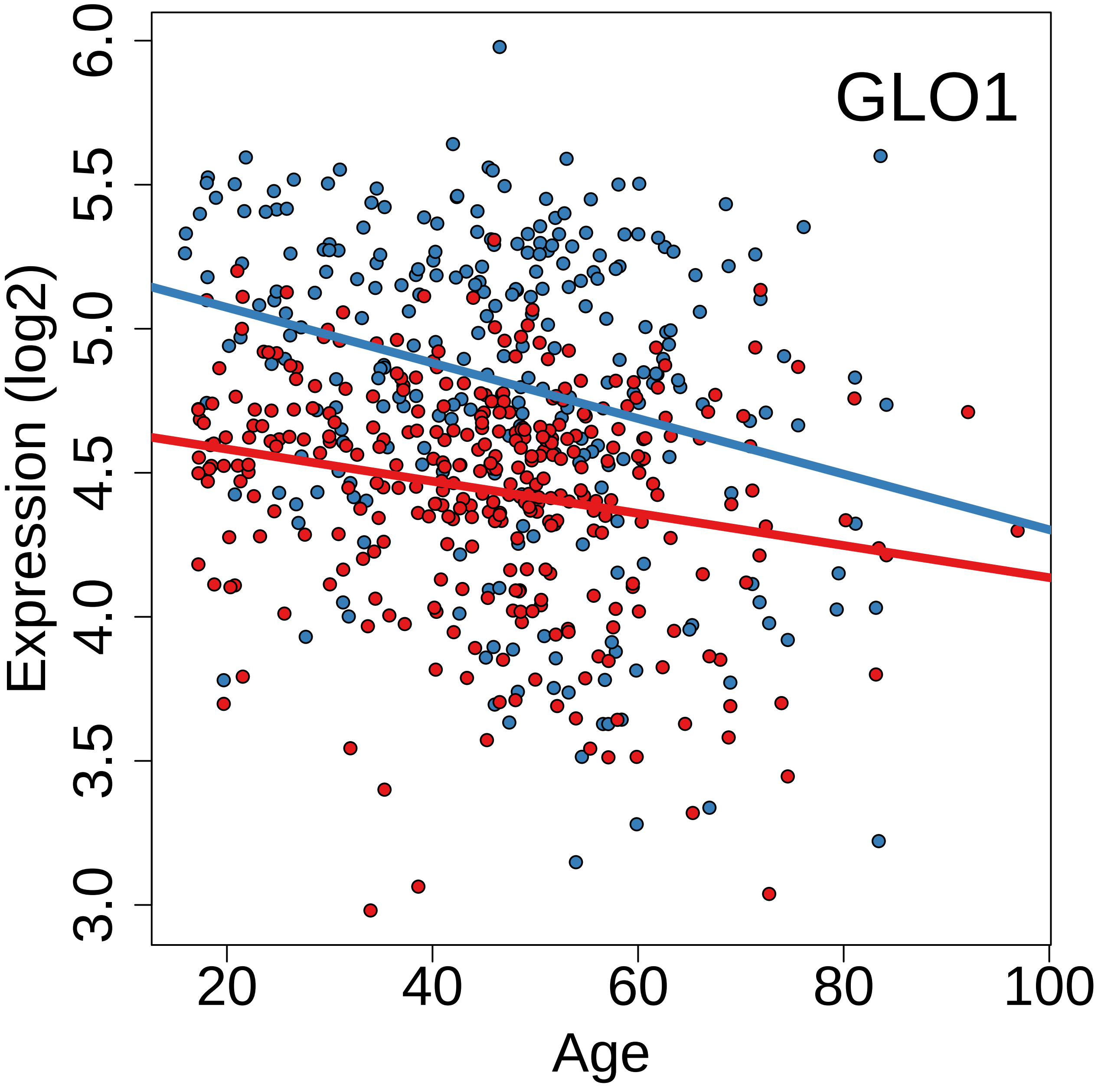

GLO1
Expression (log2)
Age
3.0
3.5
4.0
4.5
5.0
5.5
6.0
20
40
60
80
100

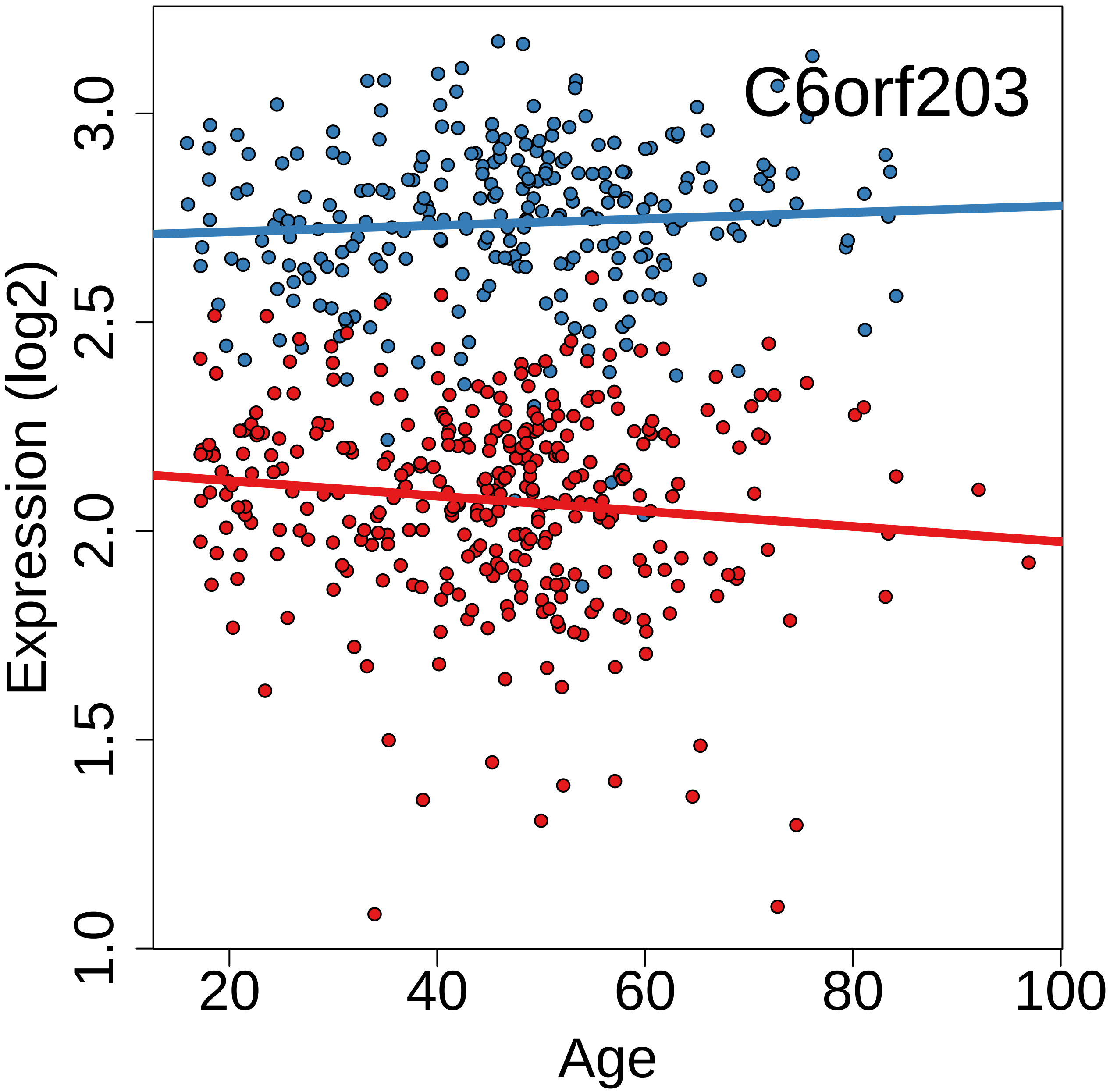
C6orf203
Expression (log2)
1.0
1.5
2.0
2.5
3.0
20
40
60
80
100
Age

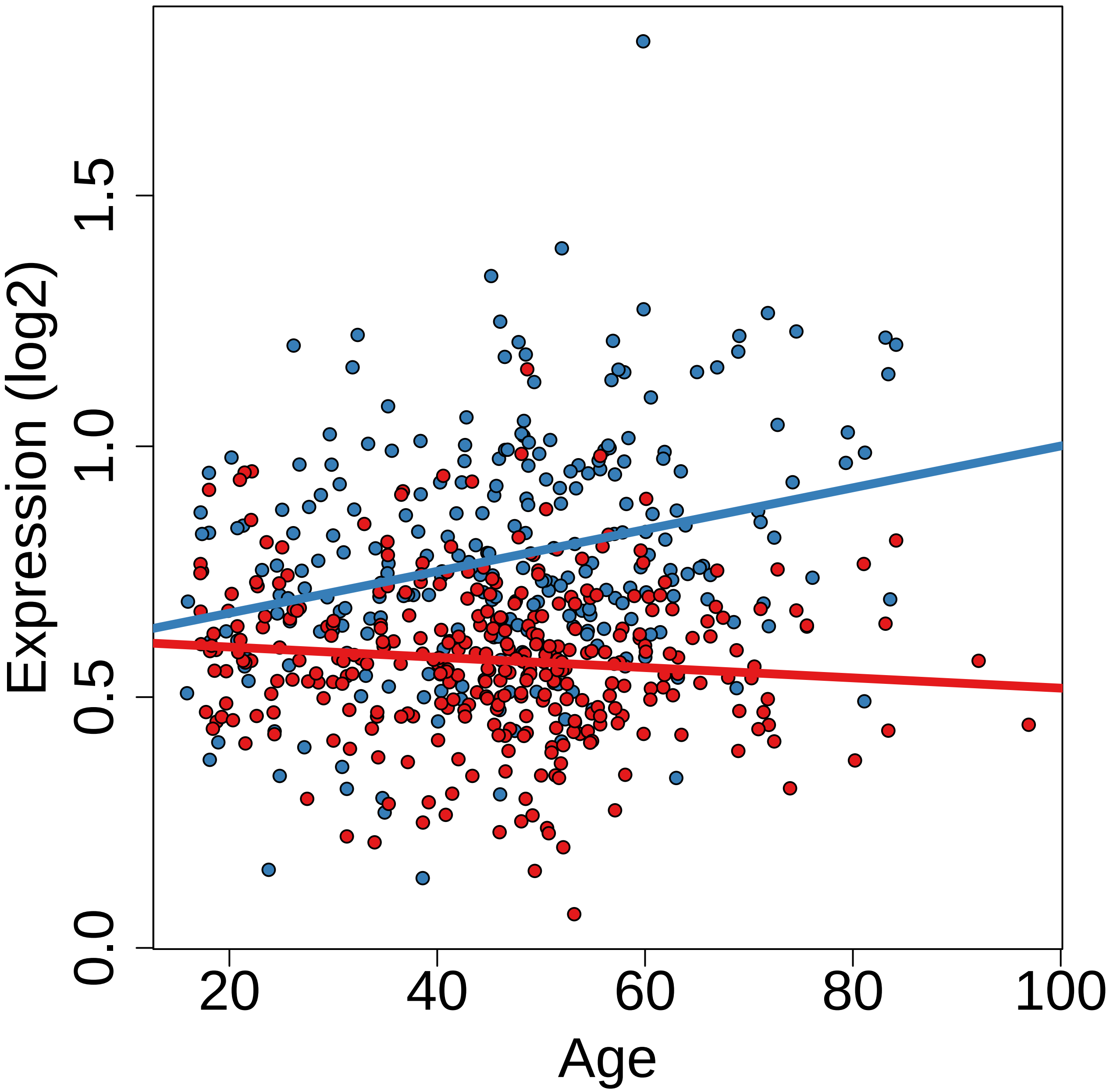

Expression (log2)
0.0
0.5
1.0
1.5
20
40
60
80
100
Age

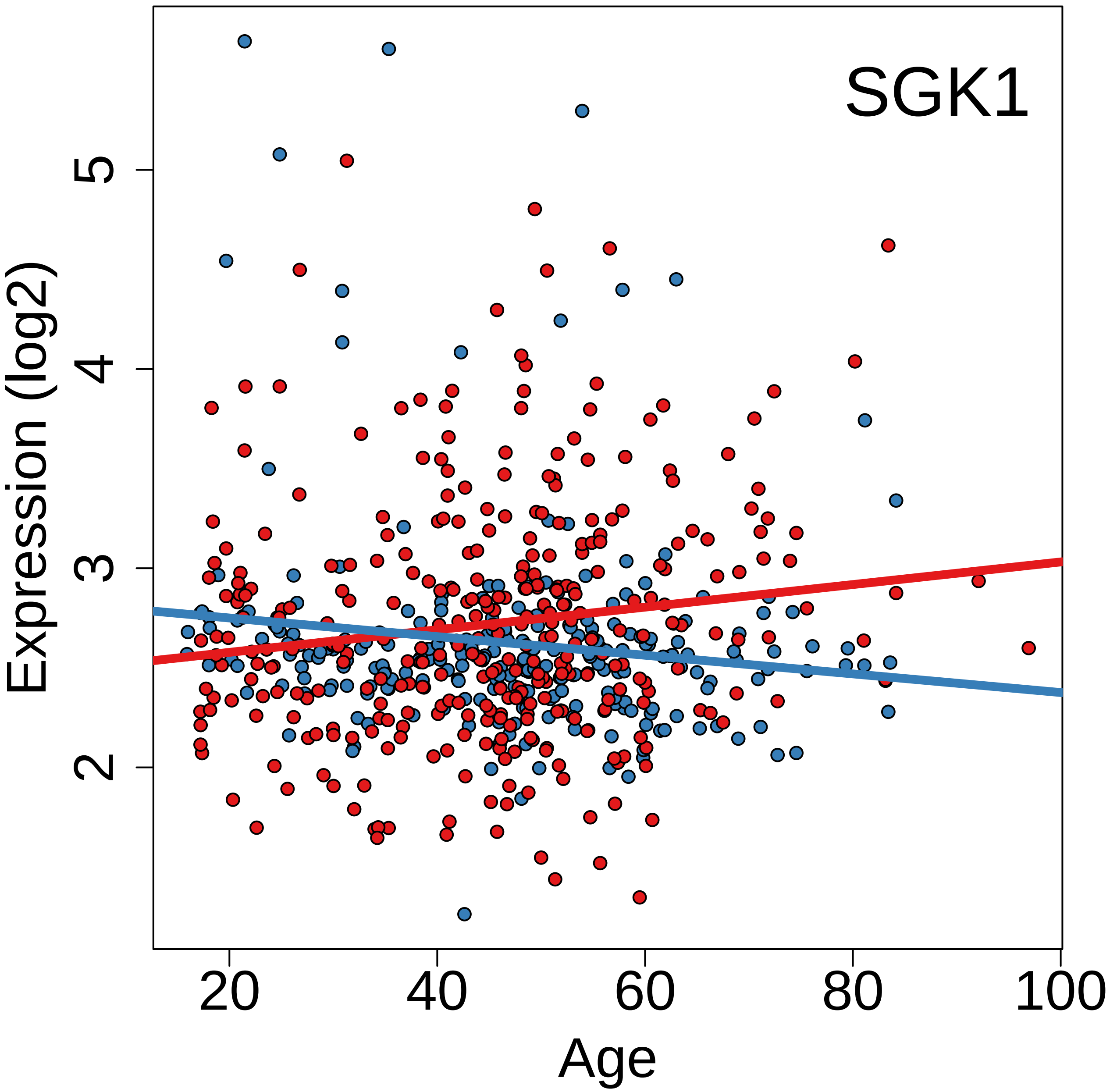
SGK1
Expression (log2)
Age
2
3
4
5
20
40
60
80
100

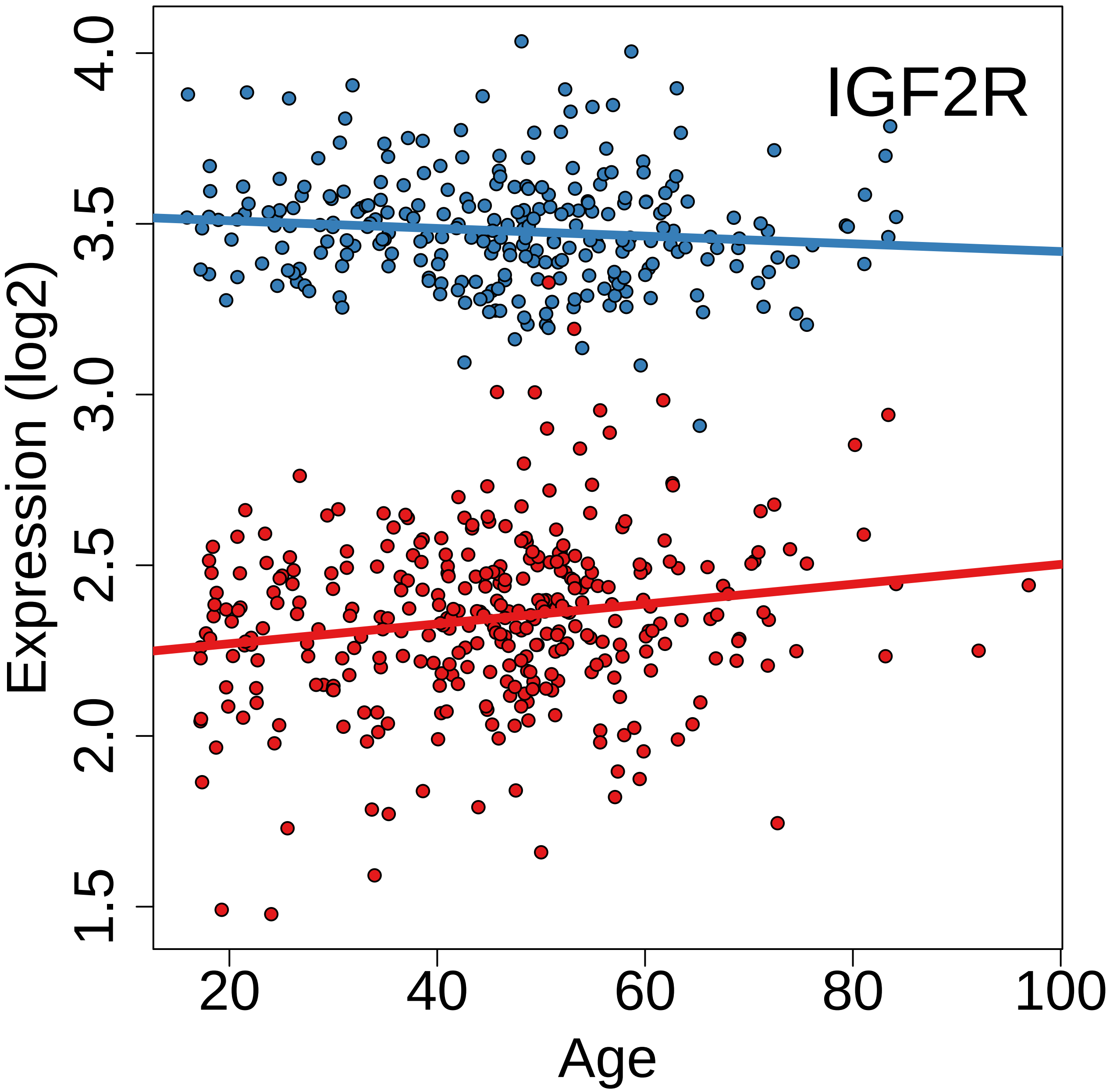
IGF2R
Expression (log2)
1.5
2.0
2.5
3.0
3.5
4.0
20
40
60
80
100
Age

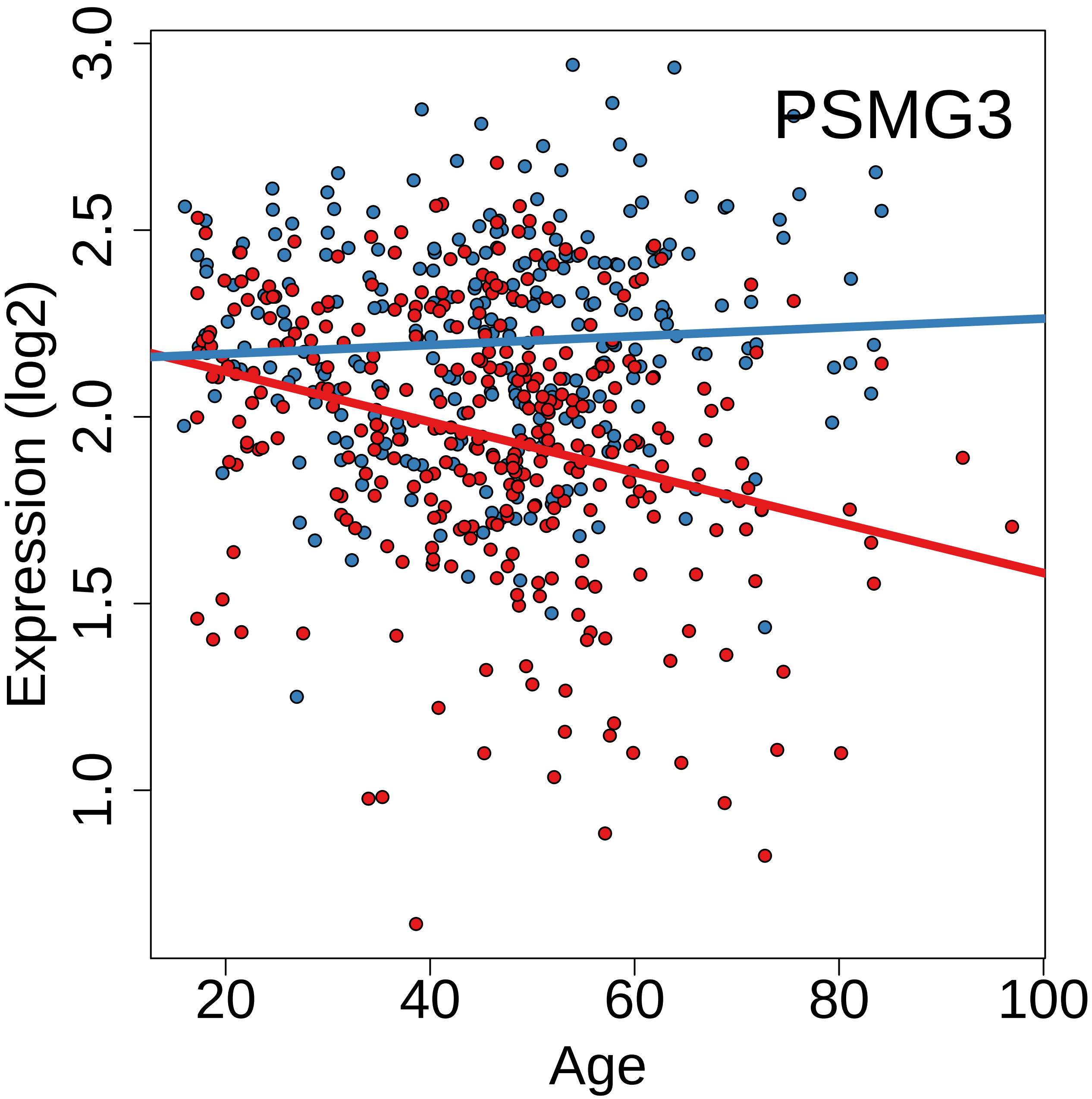
PSMG3
Expression (log2)
3.0
2.5
2.0
1.5
1.0
20
40
60
80
100
Age

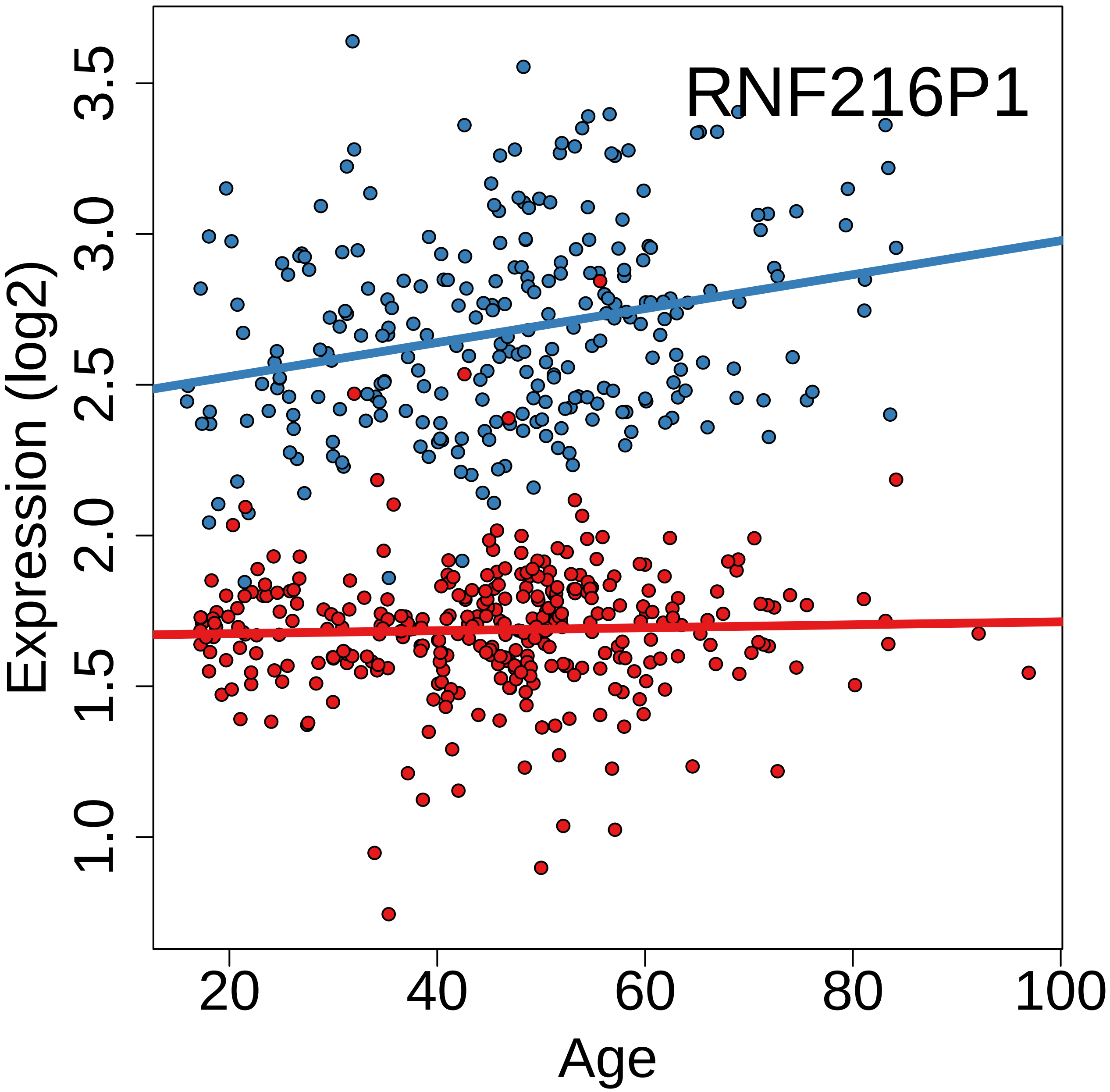
RNF216P1
Expression (log2)
3.5
3.0
2.5
2.0
1.5
1.0
20
40
60
80
100
Age

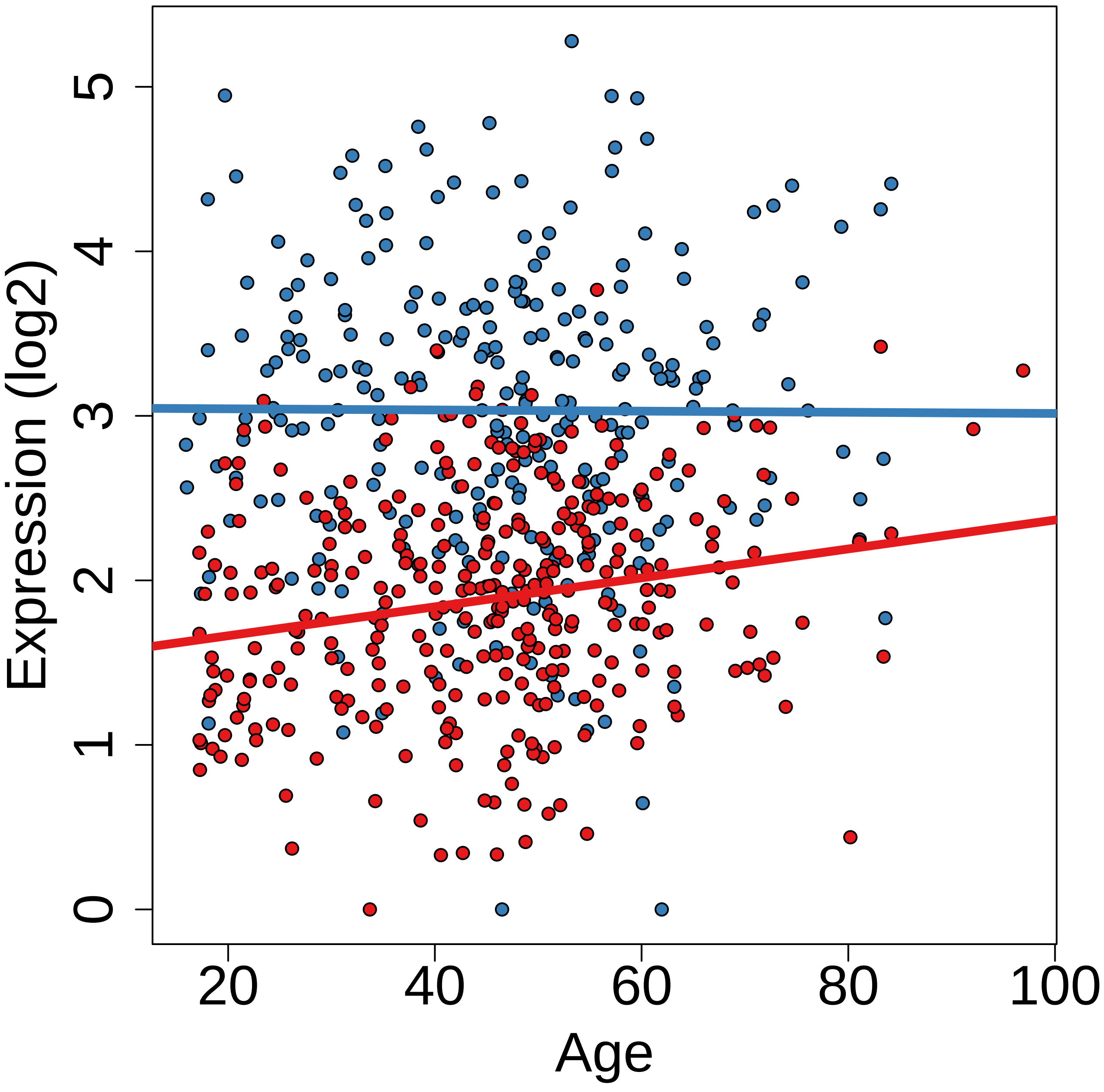

Expression (log2)
0
1
2
3
4
5
20
40
60
80
100
Age

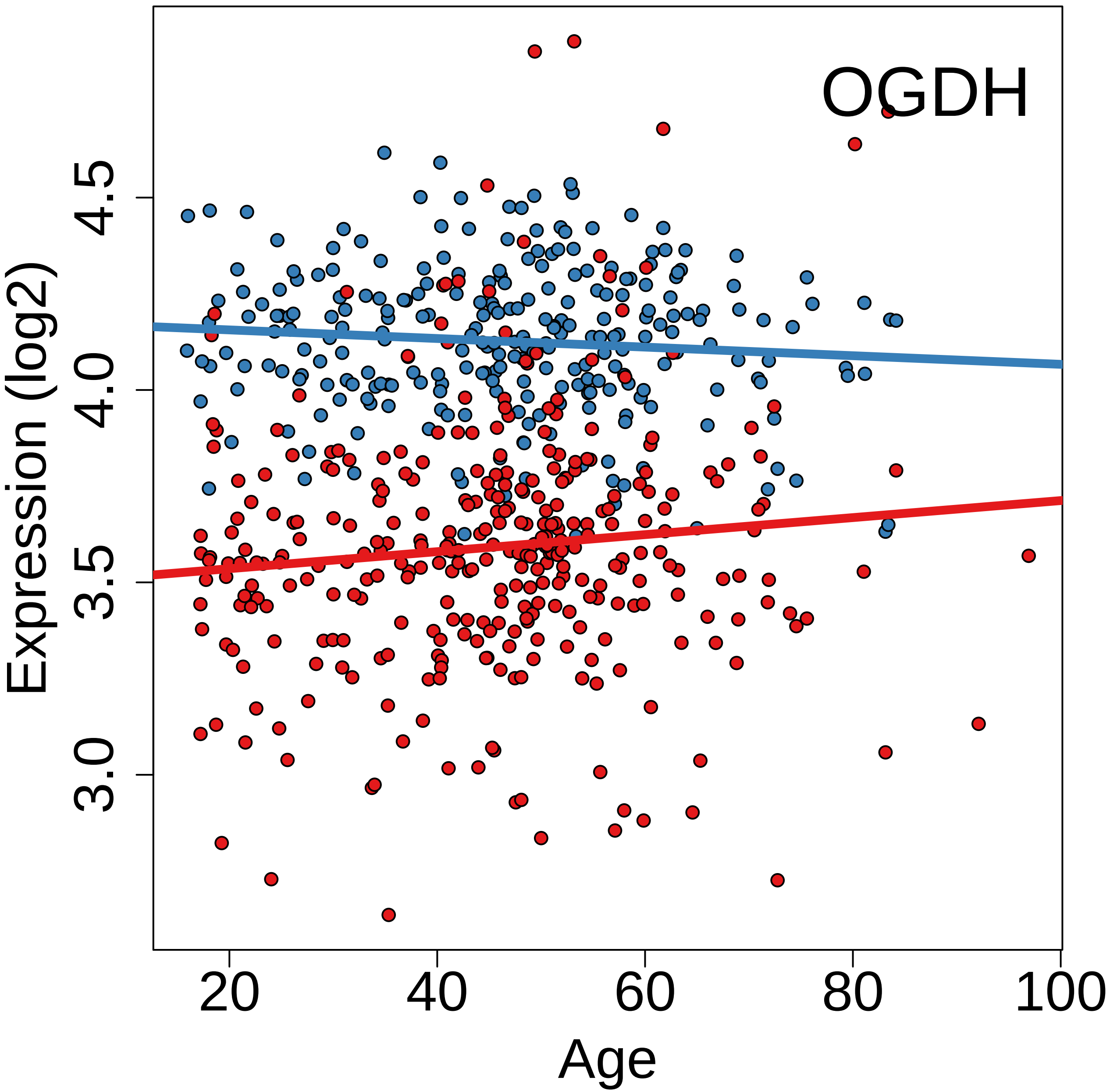
OGDH
Expression (log2)
3.0
3.5
4.0
4.5
20
40
60
80
100
Age

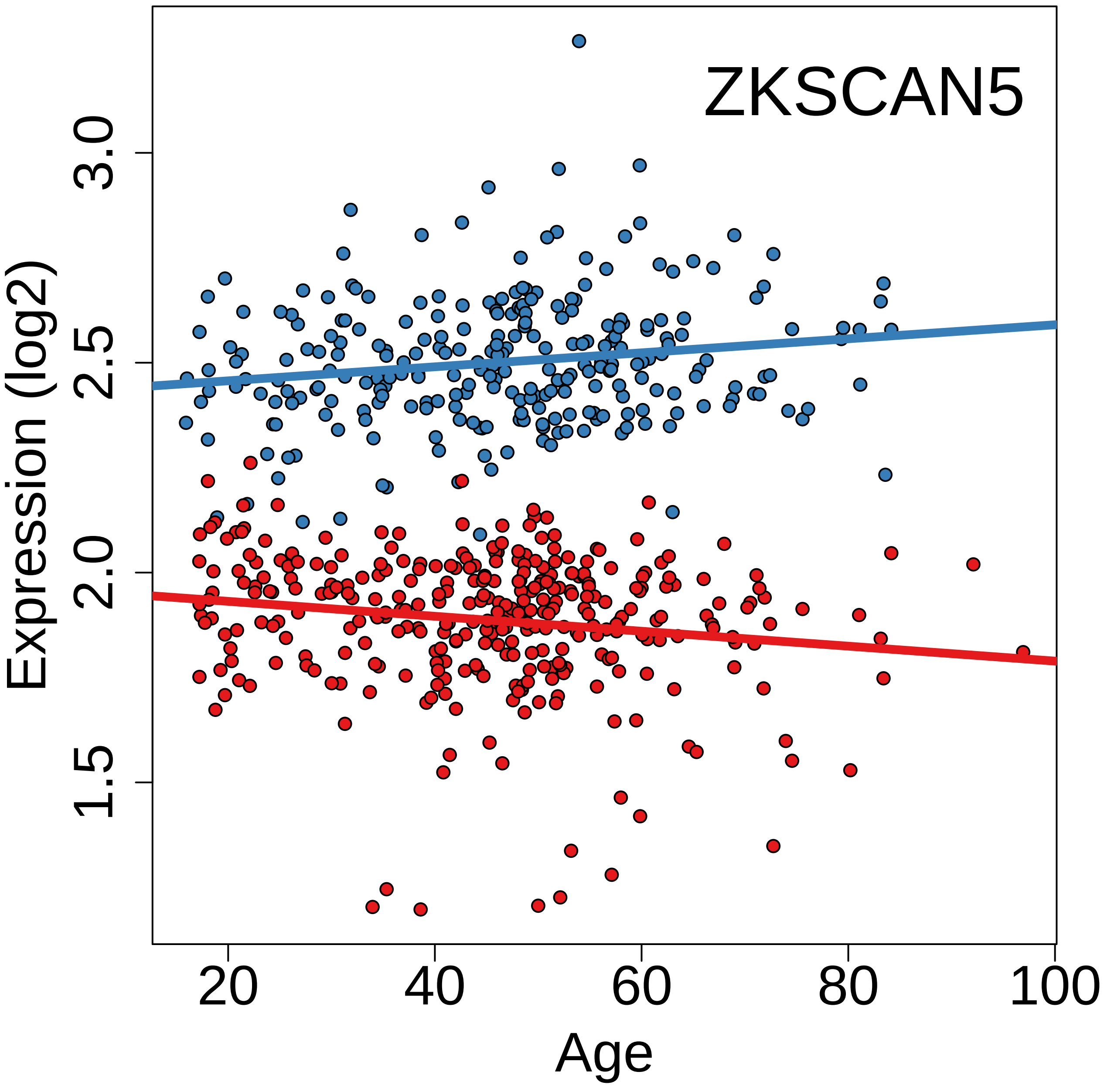
ZKSCAN5
Expression (log2)
1.5
2.0
2.5
3.0
20
40
60
80
100
Age

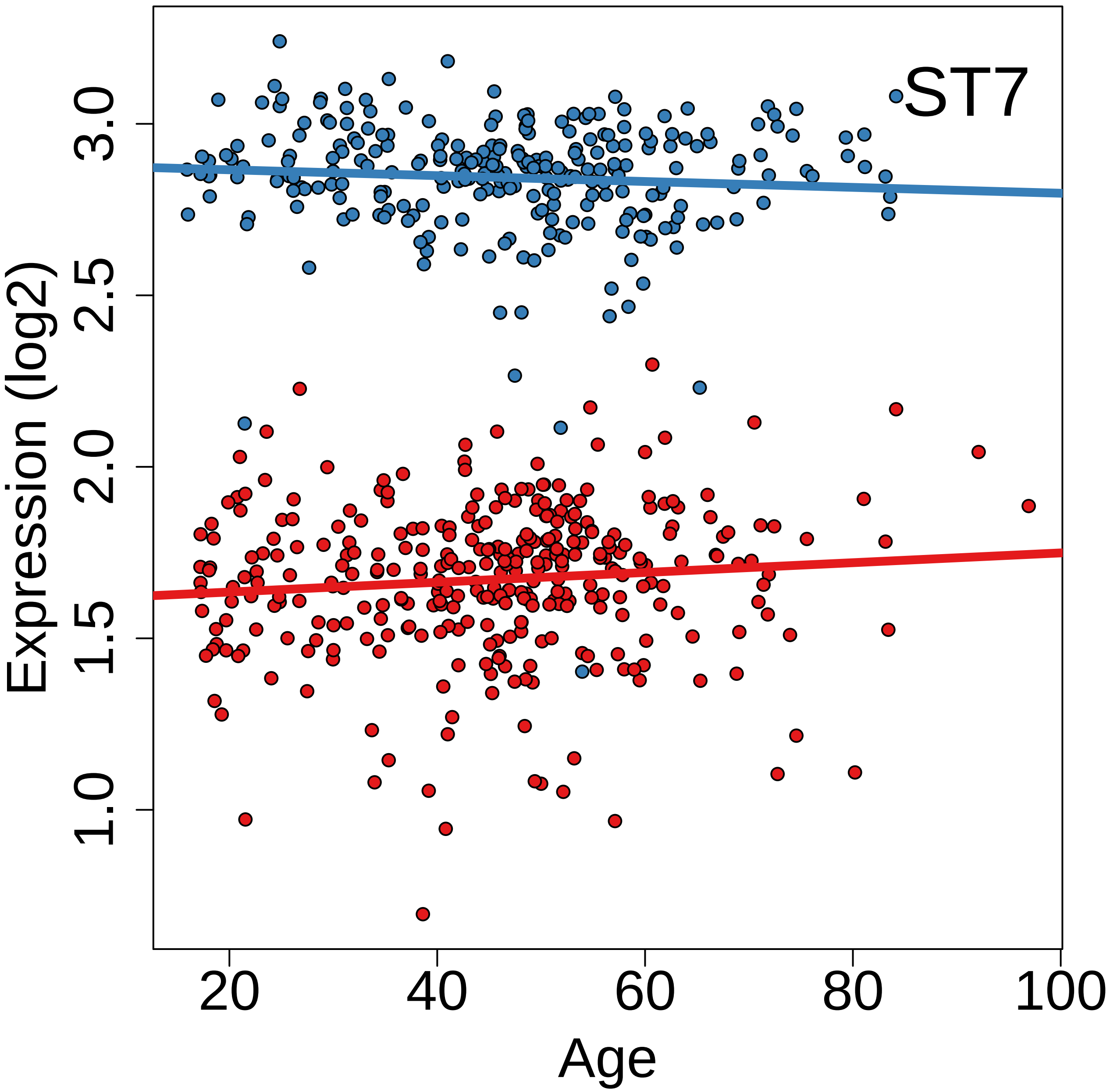
ST7
Expression (log2)
1.0
1.5
2.0
2.5
3.0
20
40
60
80
100
Age

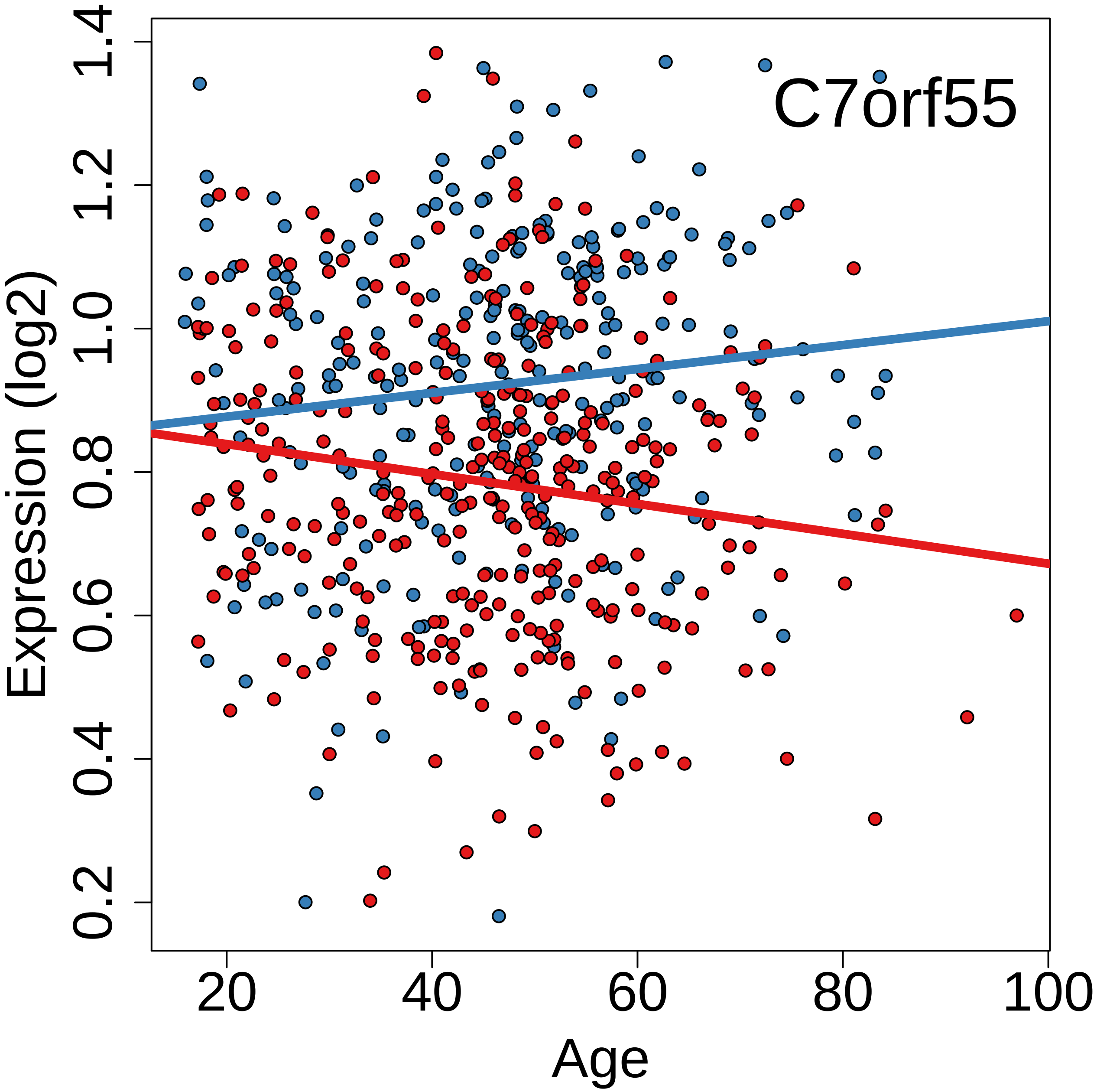
C7orf55
Expression (log2)
Age
0.2
0.4
0.6
0.8
1.0
1.2
1.4
20
40
60
80
100

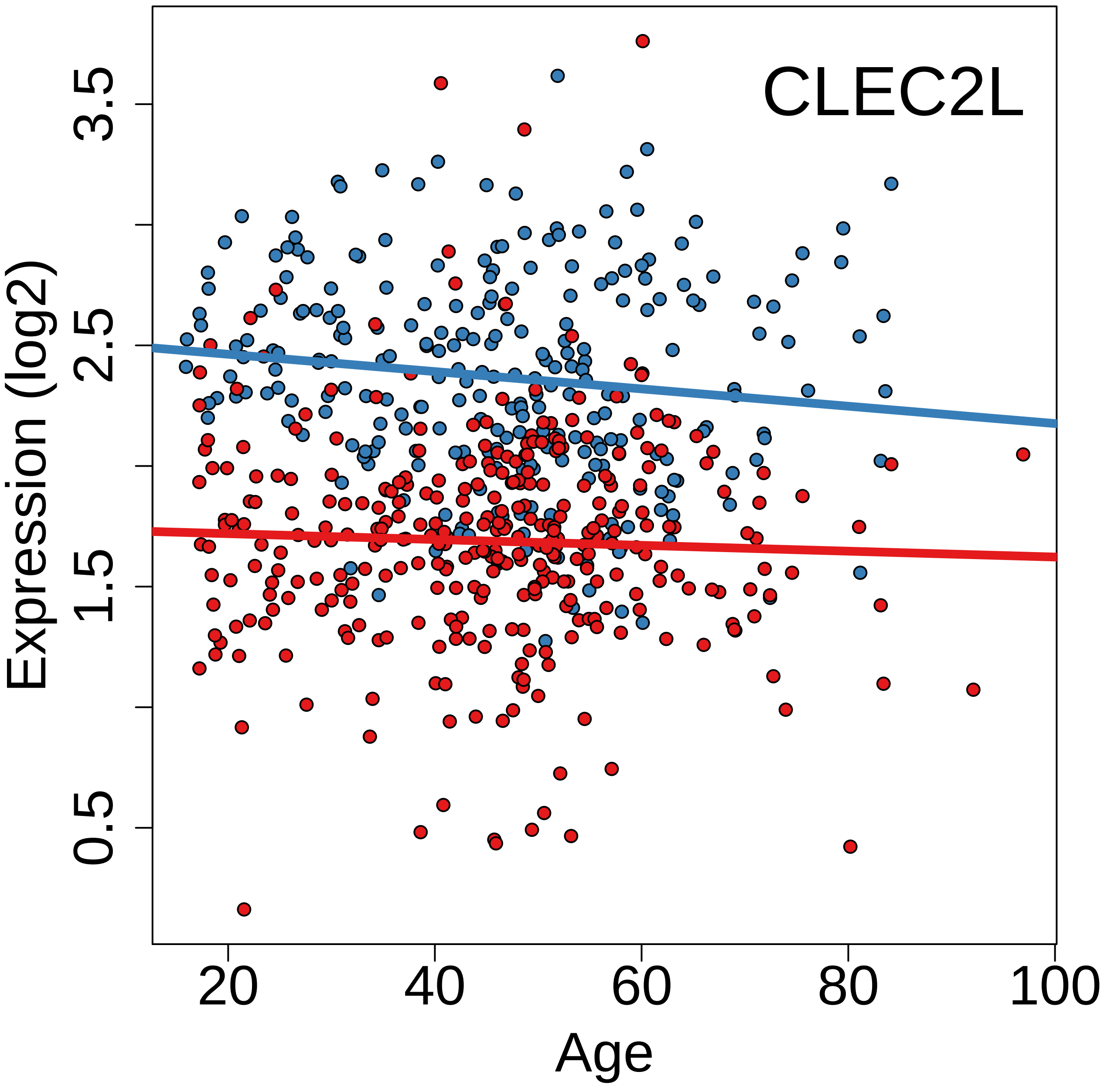
CLEC2L
Expression (log2)
0.5
1.5
2.5
3.5
20
40
60
80
100
Age

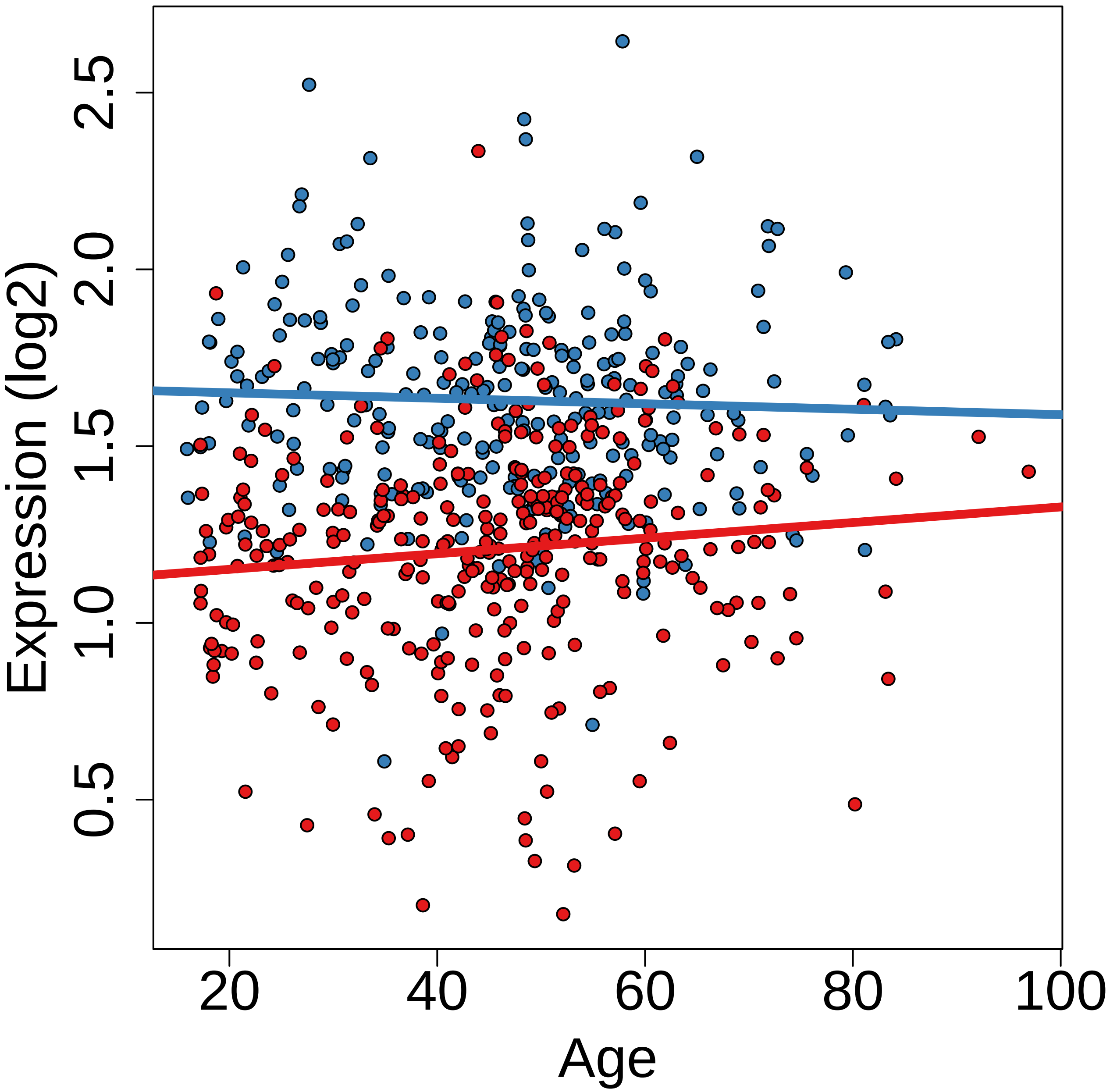
Expression (log2)
2.5
2.0
1.5
1.0
0.5
20
40
60
80
100
Age

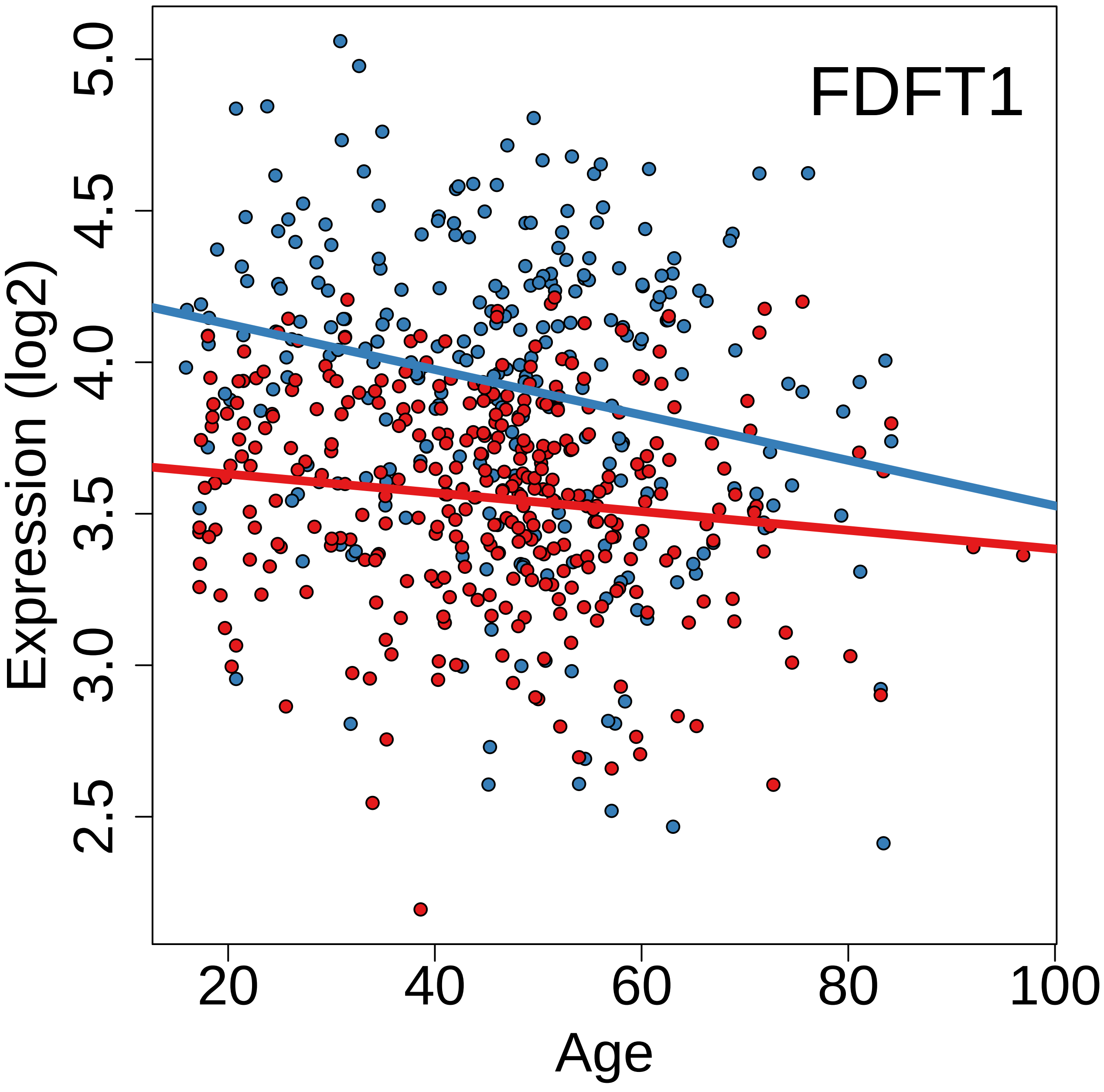
FDFT1
Expression (log2)
Age
2.5
3.0
3.5
4.0
4.5
5.0
20
40
60
80
100

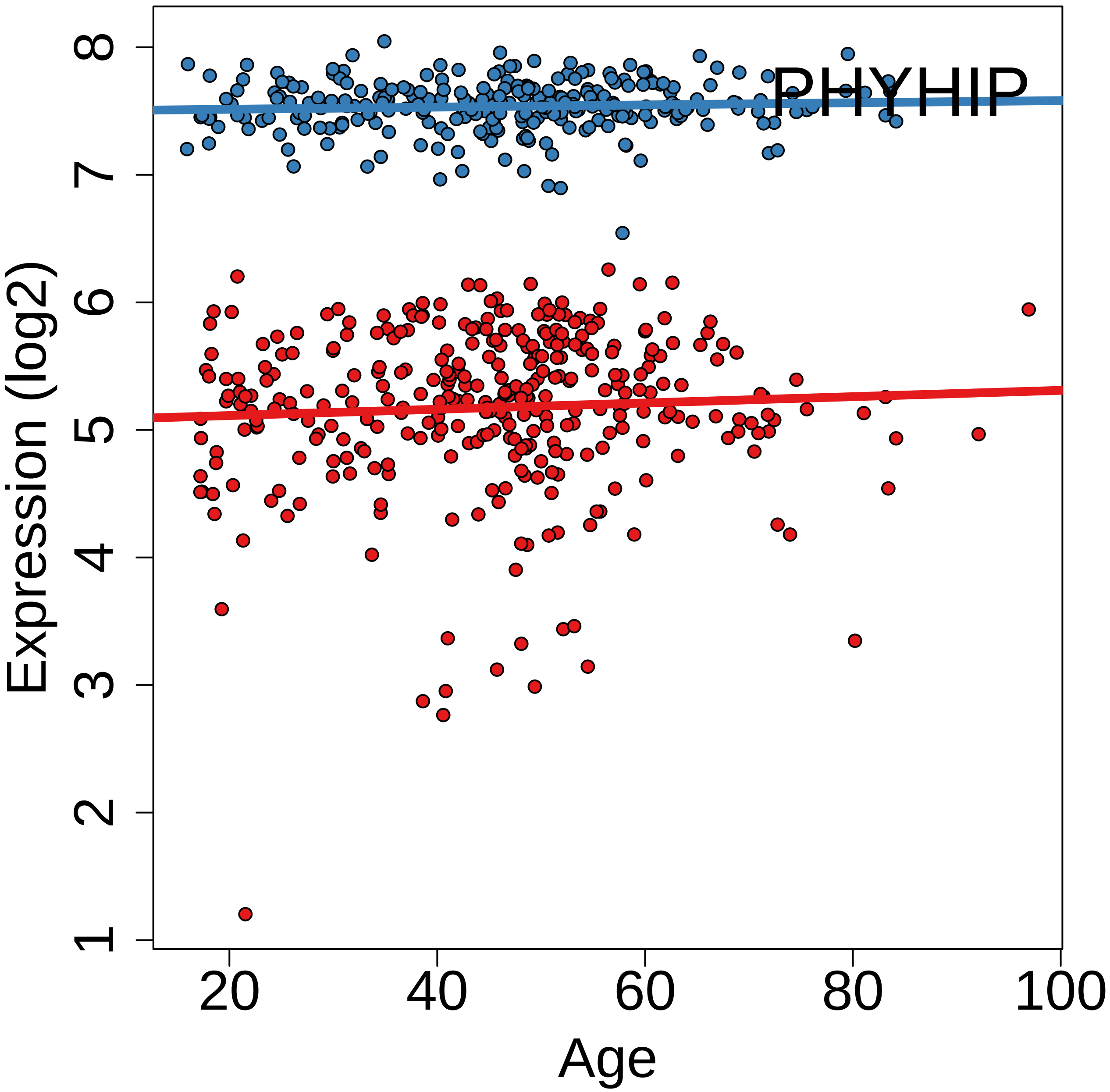
PHYHIP
Expression (log2)
Age
1
2
3
4
5
6
7
8
20
40
60
80
100

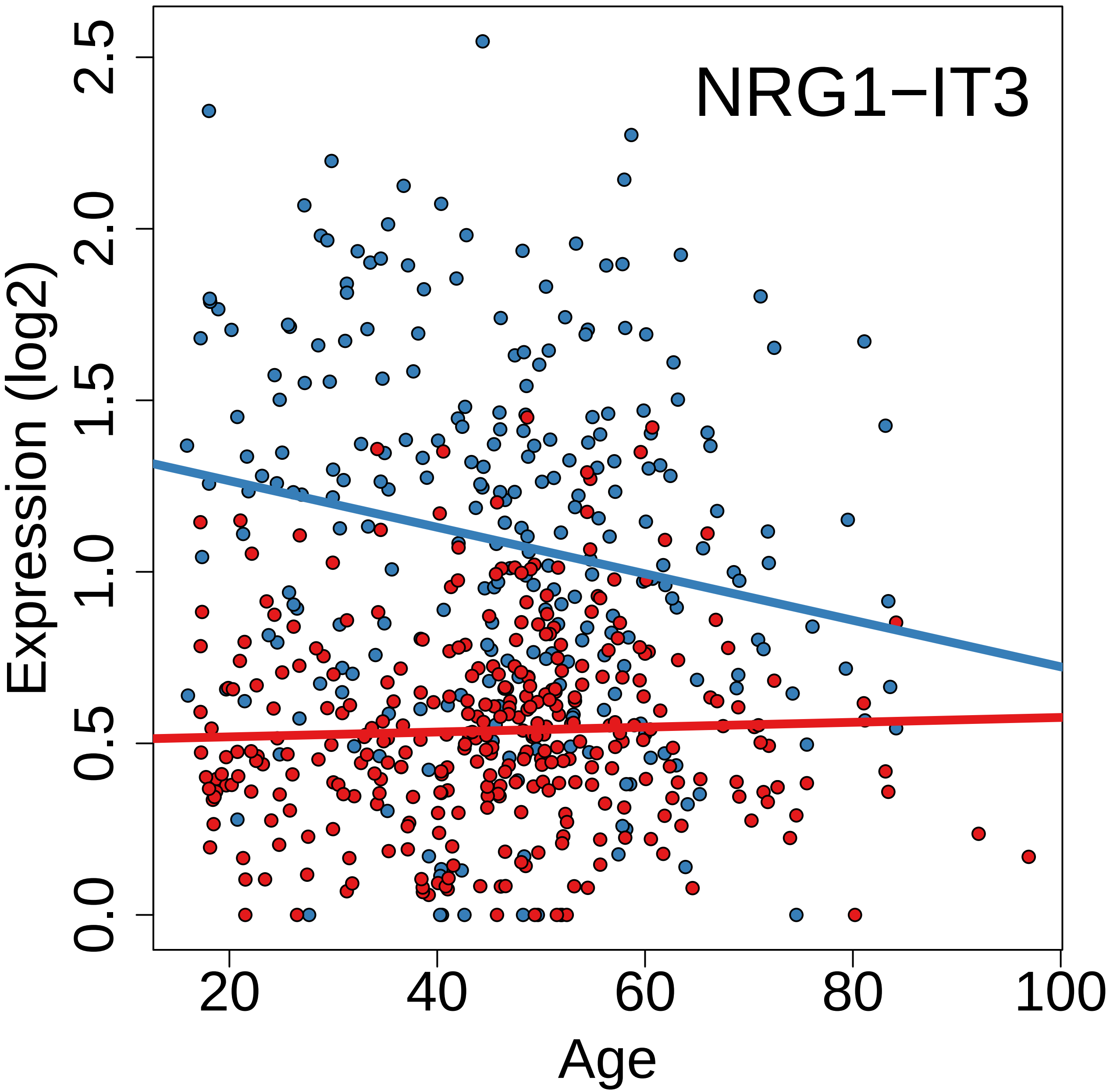
NRG1−IT3
Expression (log2)
0.0
0.5
1.0
1.5
2.0
2.5
20
40
60
80
100
Age

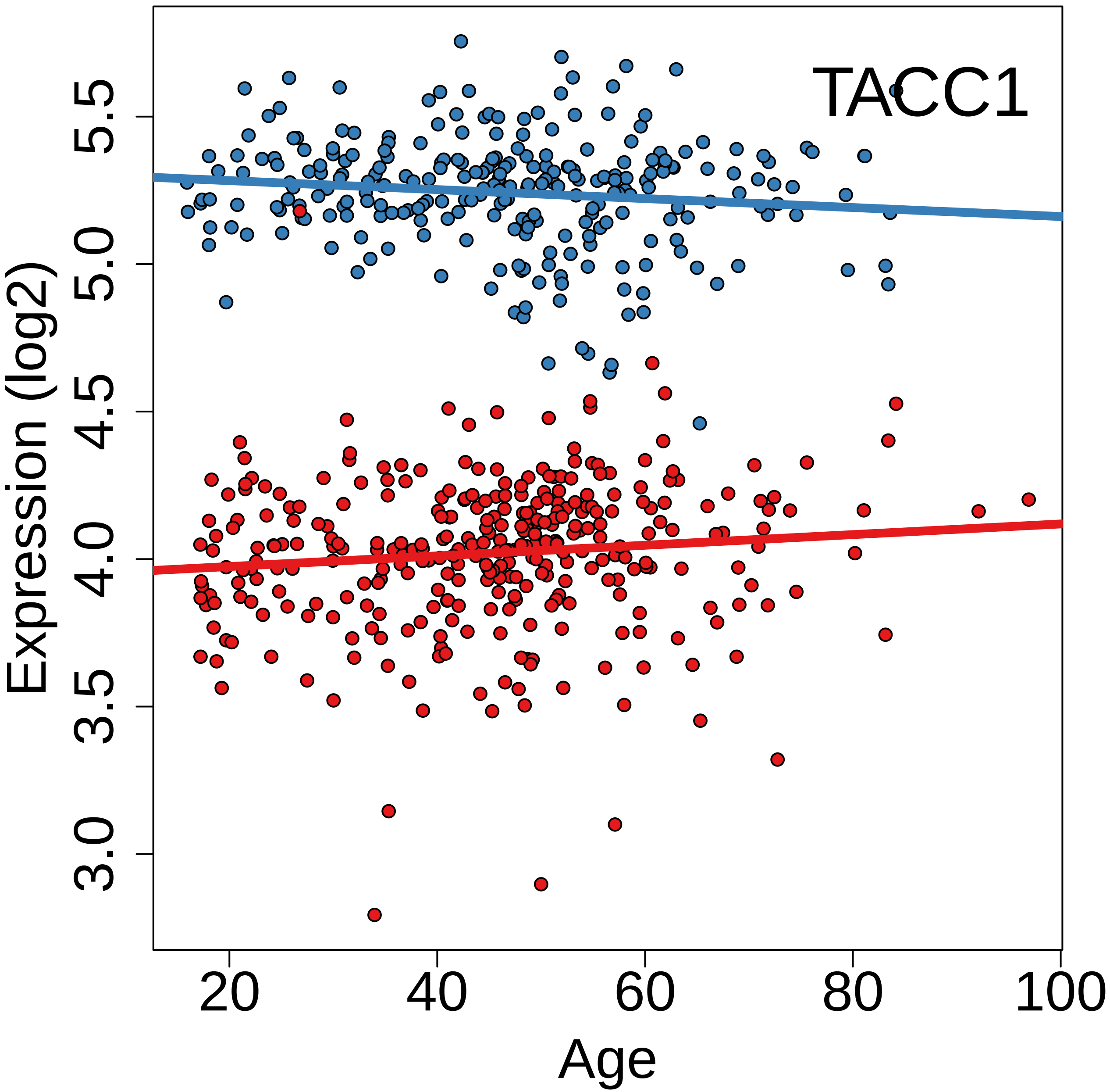
TACC1
Expression (log2)
3.0
3.5
4.0
4.5
5.0
5.5
20
40
60
80
100
Age

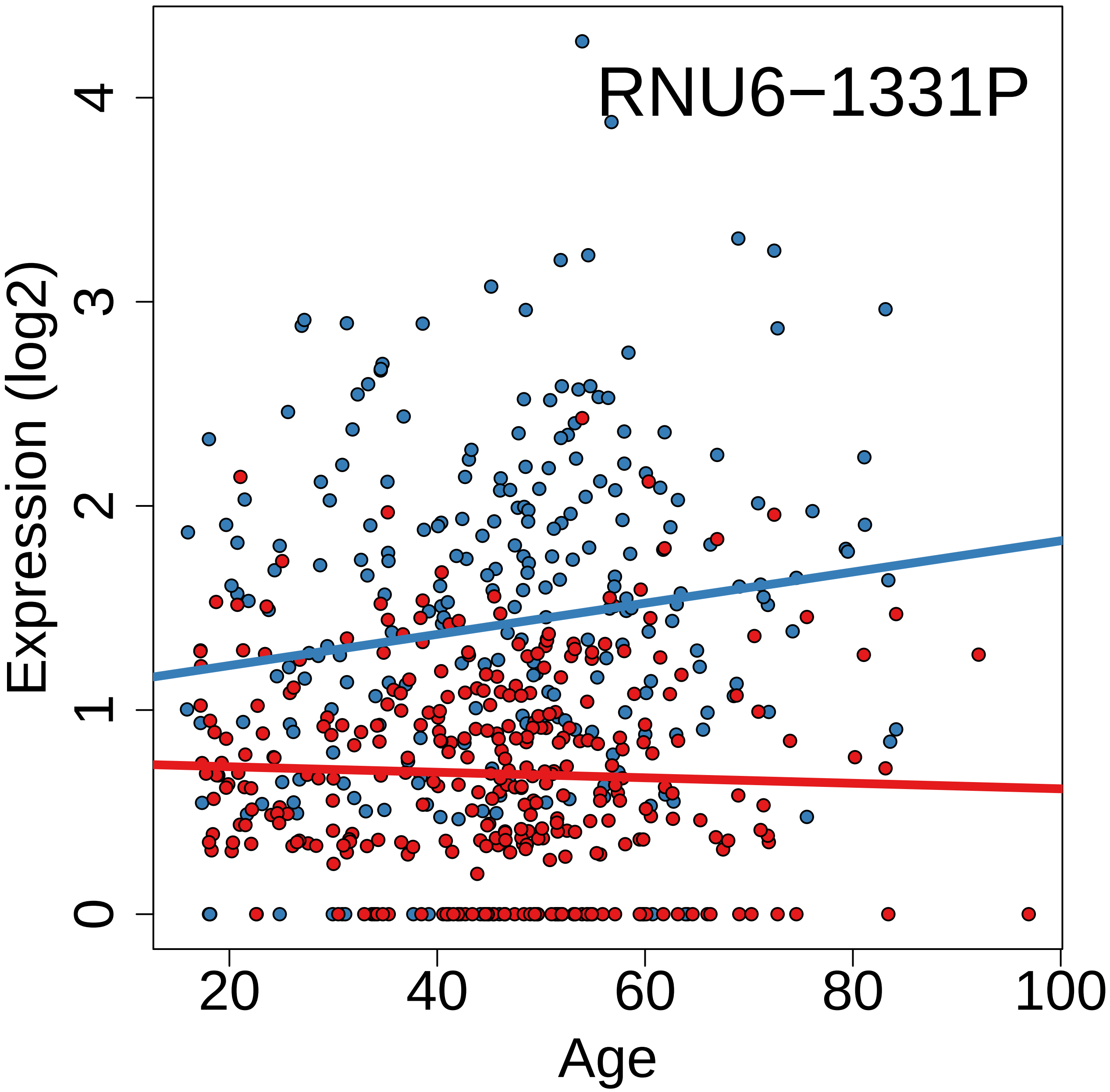
RNU6−1331P
Expression (log2)
0
1
2
3
4
20
40
60
80
100
Age

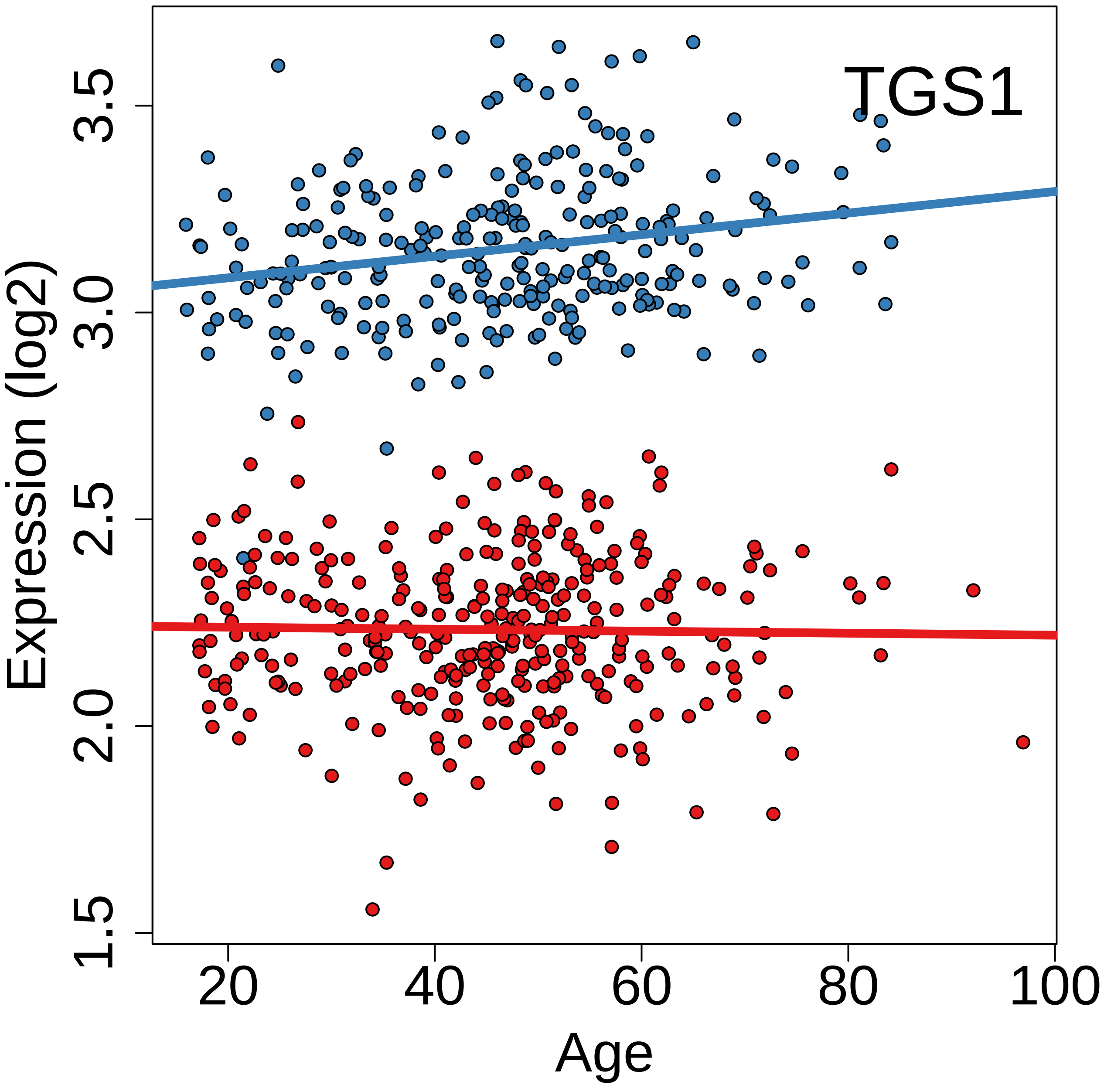

TGS1
Expression (log2)
Age
1.5
2.0
2.5
3.0
3.5
20
40
60
80
100

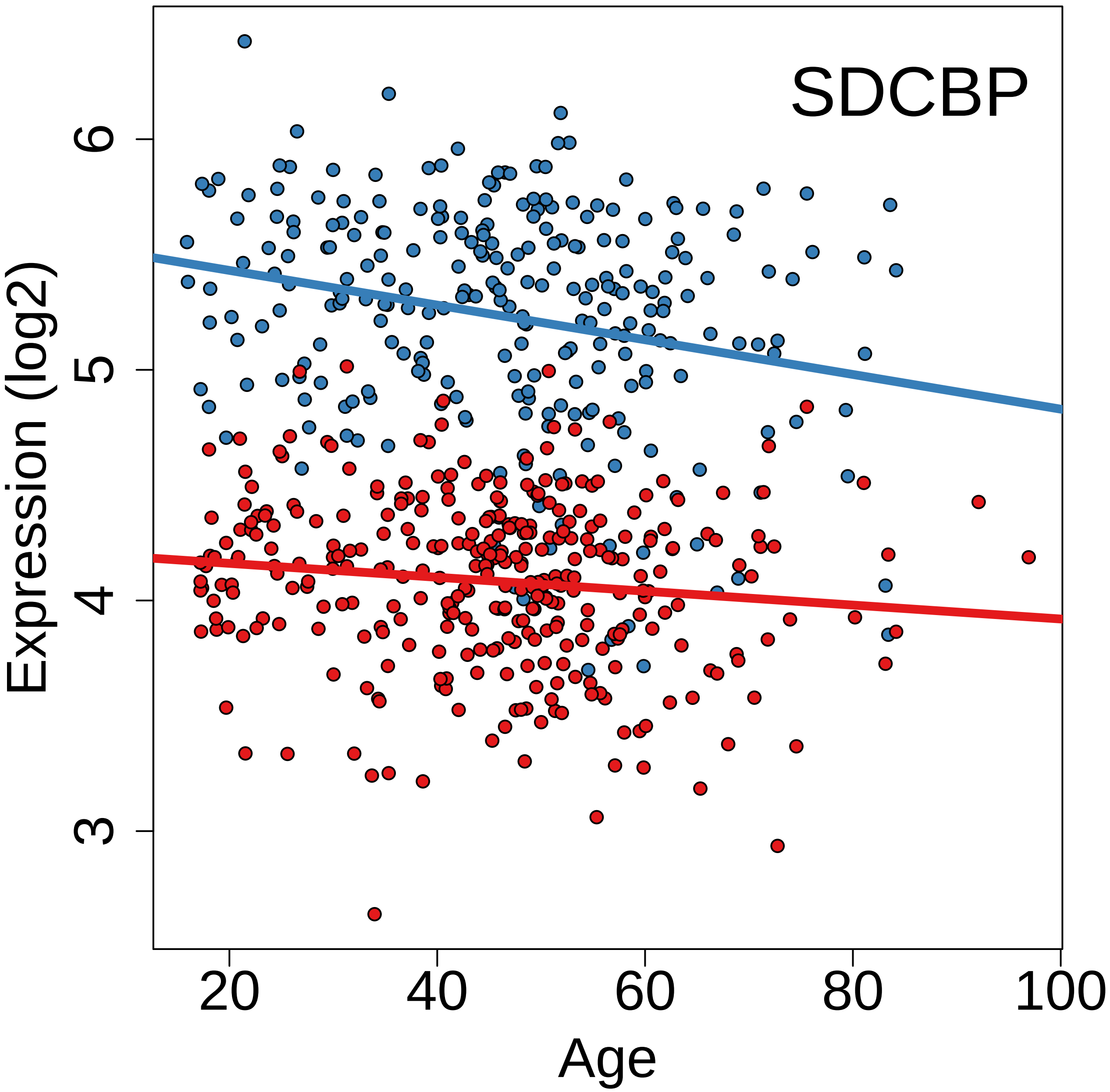

SDCBP
Expression (log2)
3
4
5
6
20
40
60
80
100
Age

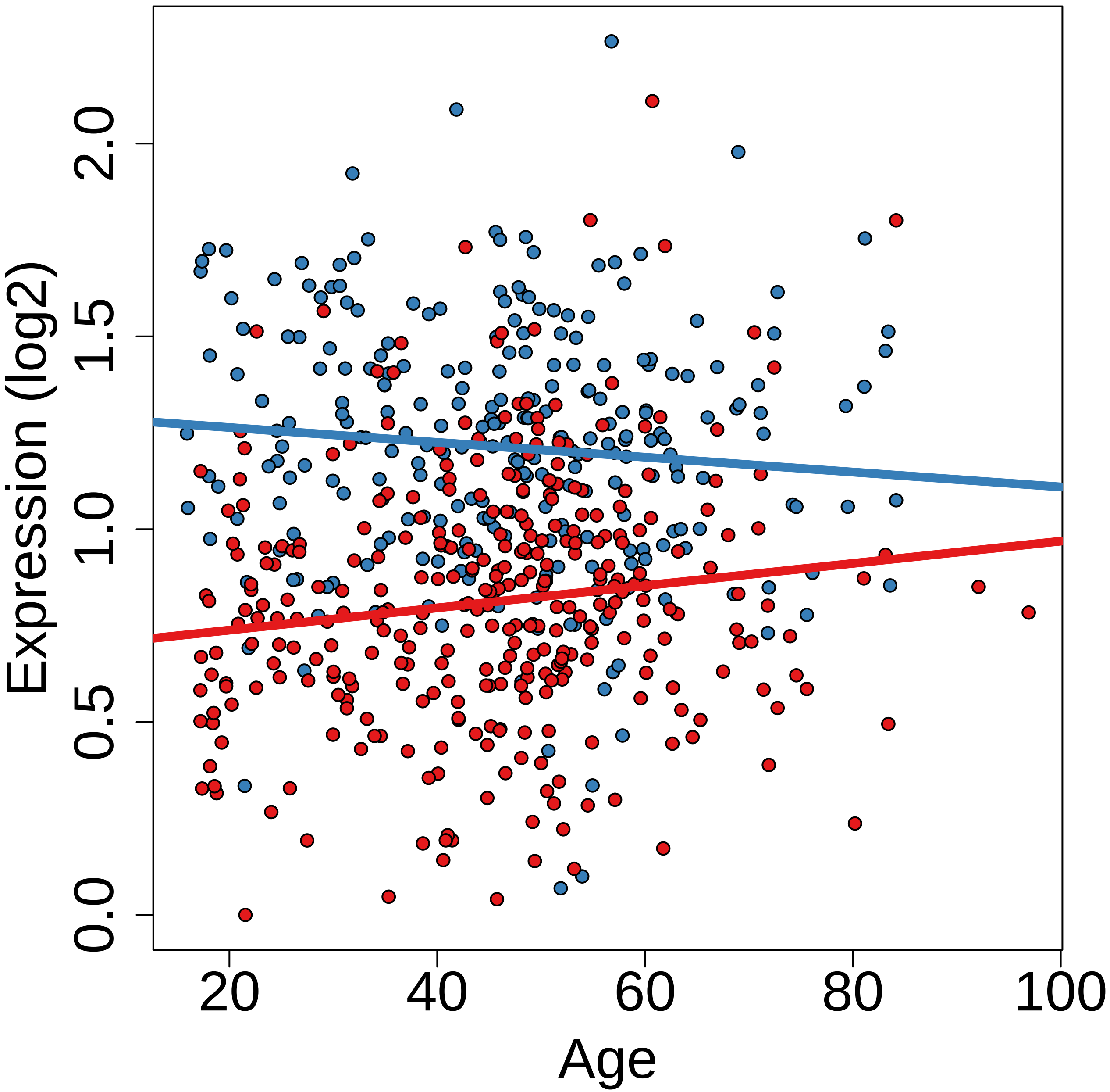

Expression (log2)
0.0
0.5
1.0
1.5
2.0
20
40
60
80
100
Age

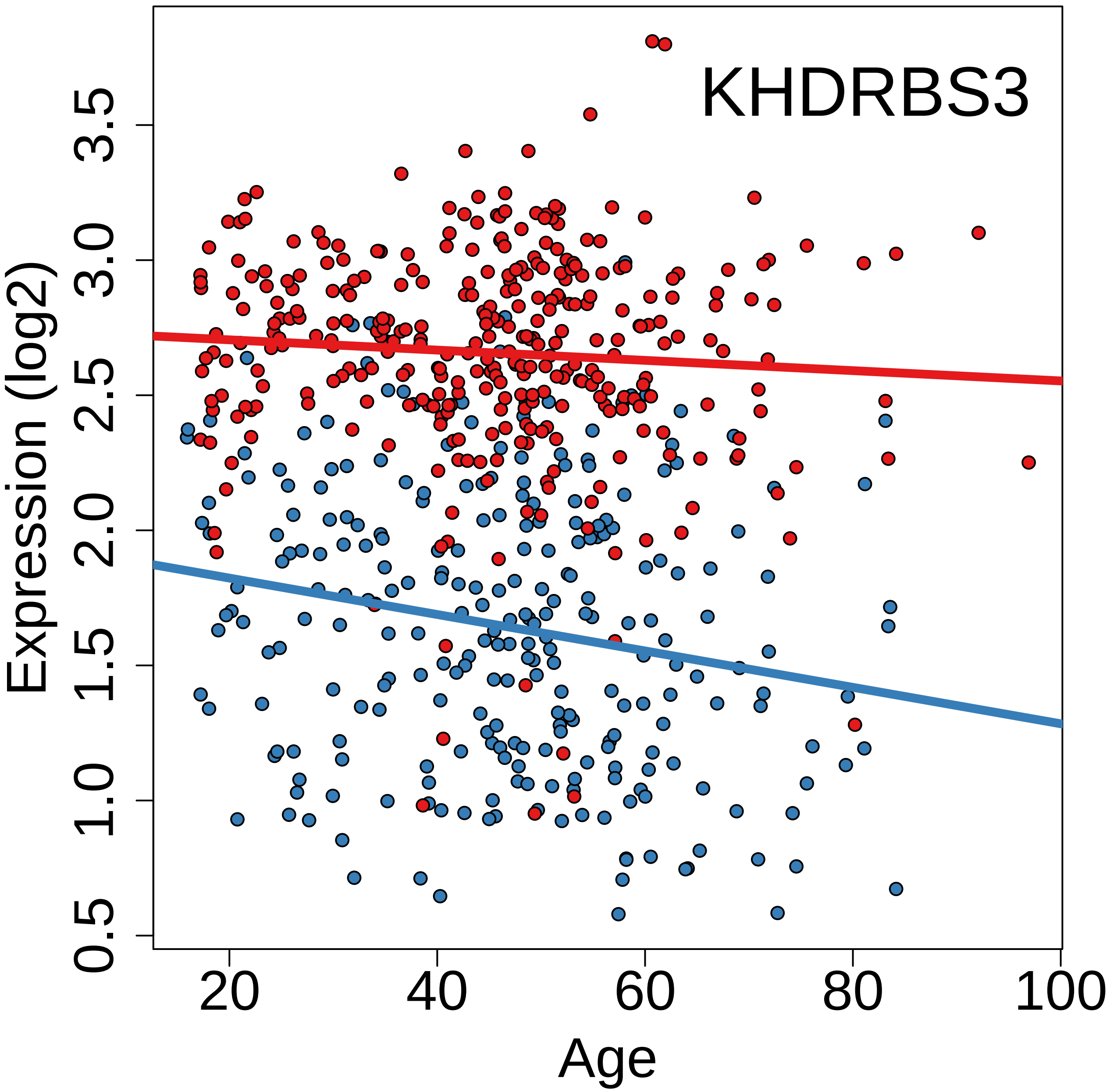
KHDRBS3
Expression (log2)
0.5
1.0
1.5
2.0
2.5
3.0
3.5
20
40
60
80
100
Age

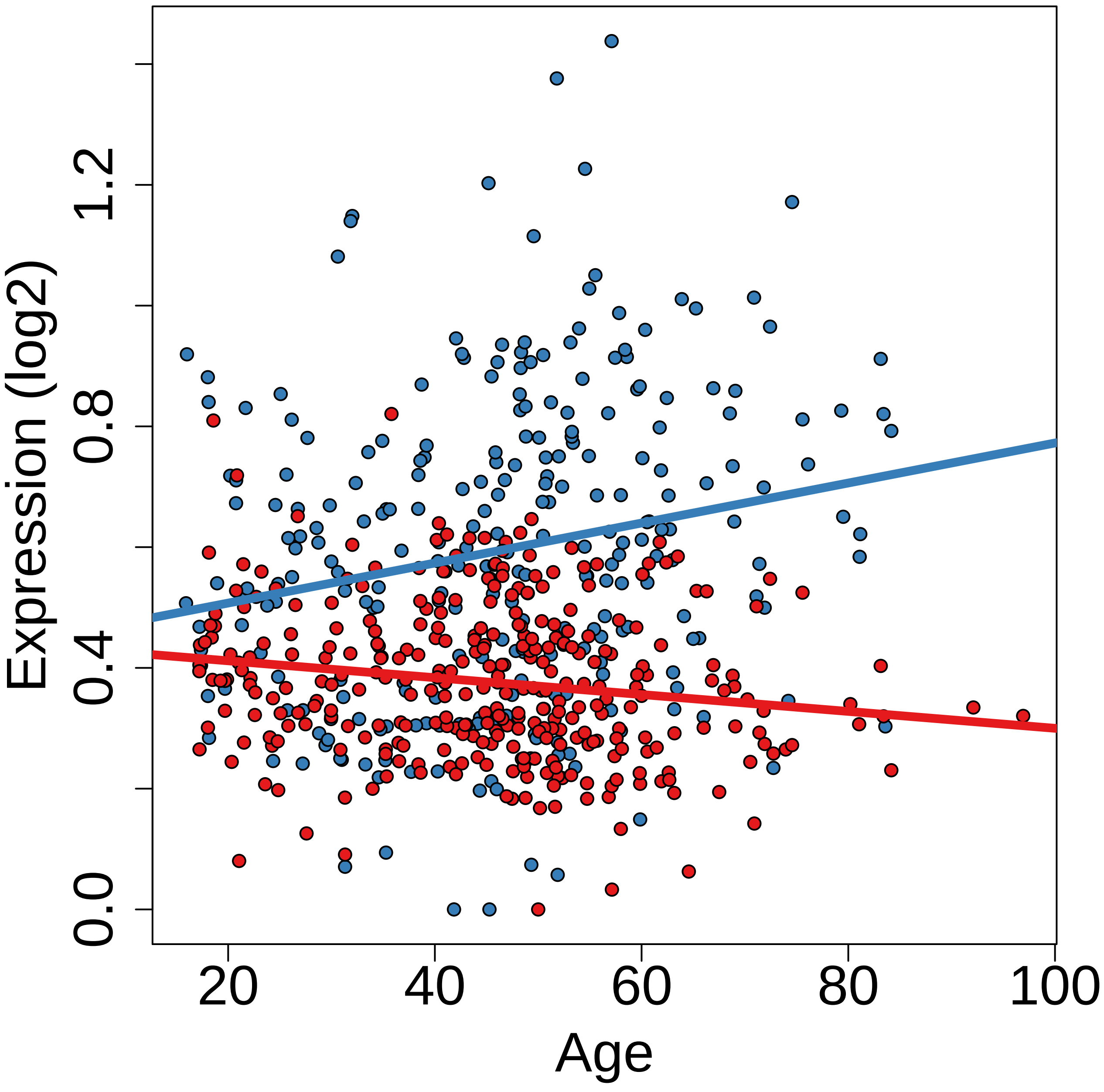
Expression (log2)
0.0
0.4
0.8
1.2
20
40
60
80
100
Age

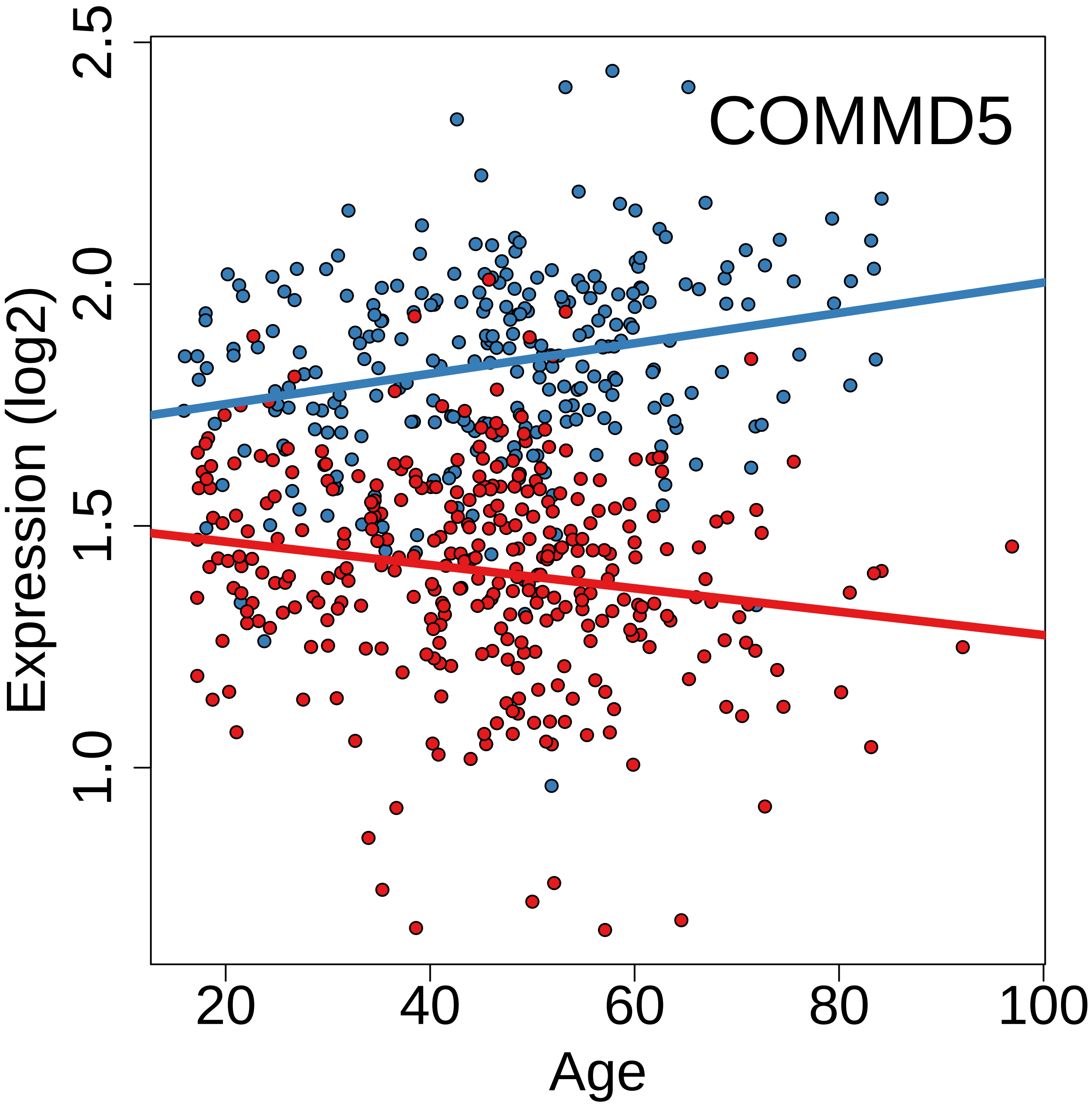
COMMD5
Expression (log2)
2.5
2.0
1.5
1.0
20
40
60
80
100
Age

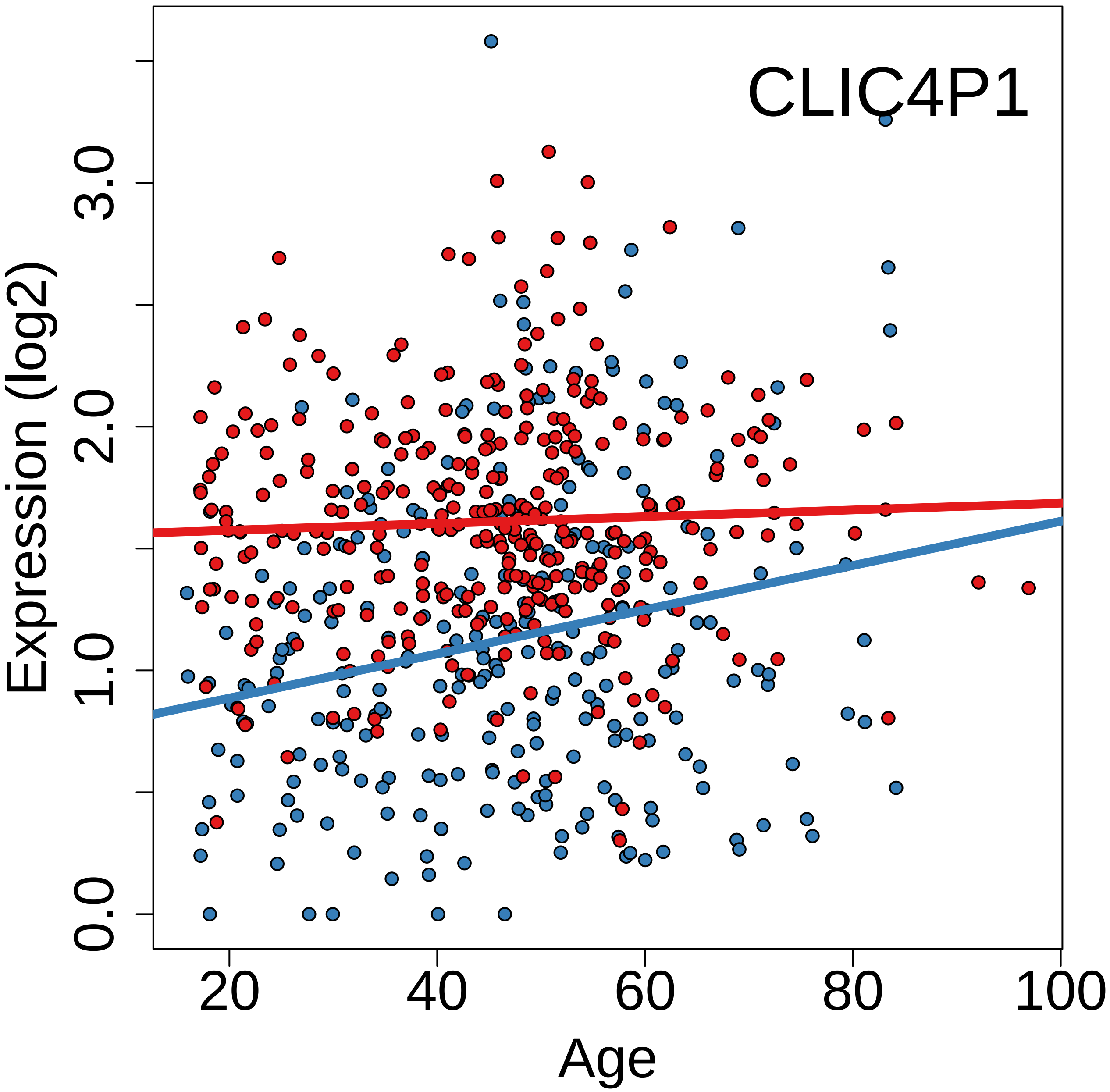

CLIC4P1
Expression (log2)
0.0
1.0
2.0
3.0
20
40
60
80
100
Age

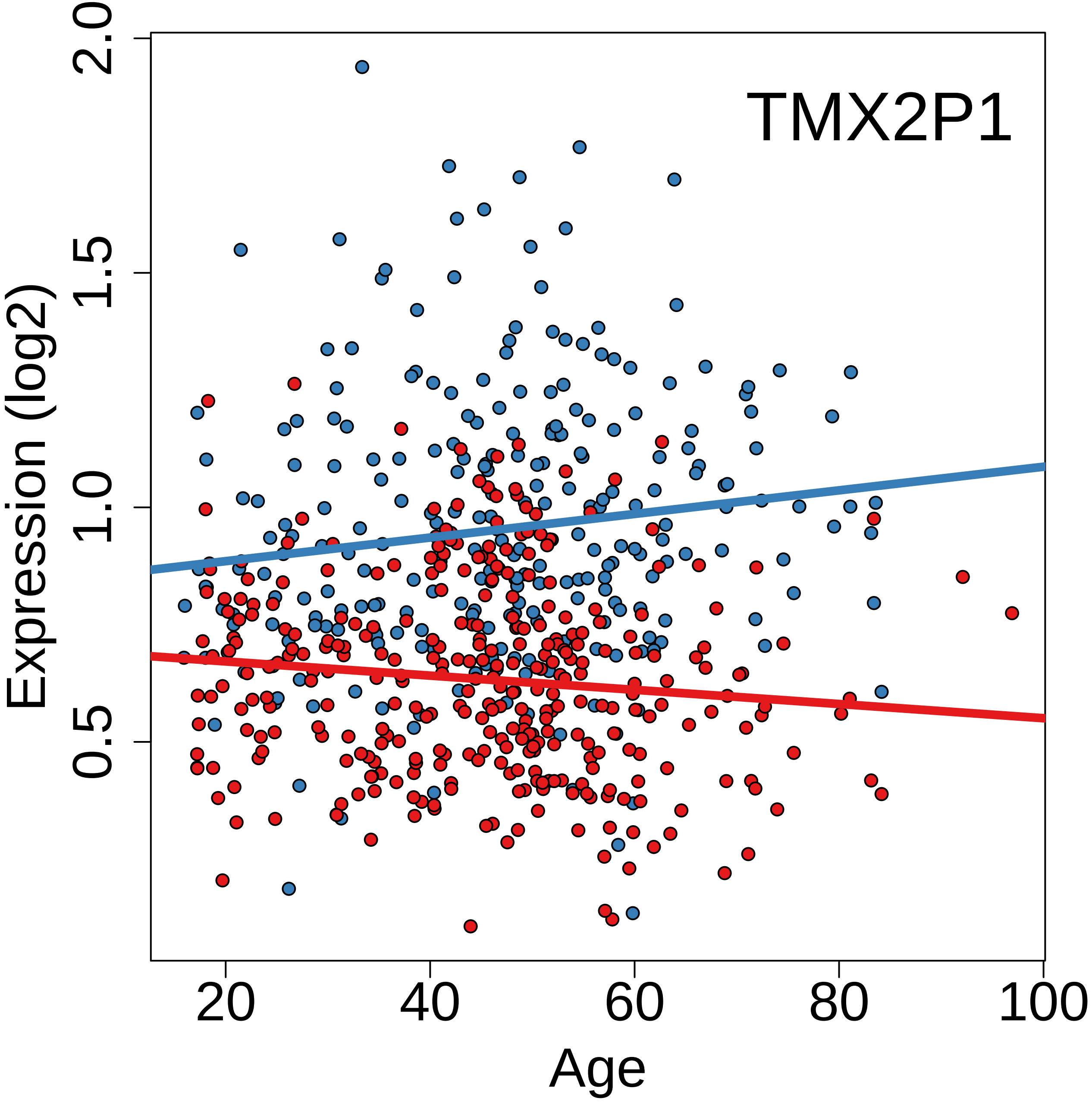
TMX2P1
Expression (log2)
2.0
1.5
1.0
0.5
20
40
60
80
100
Age

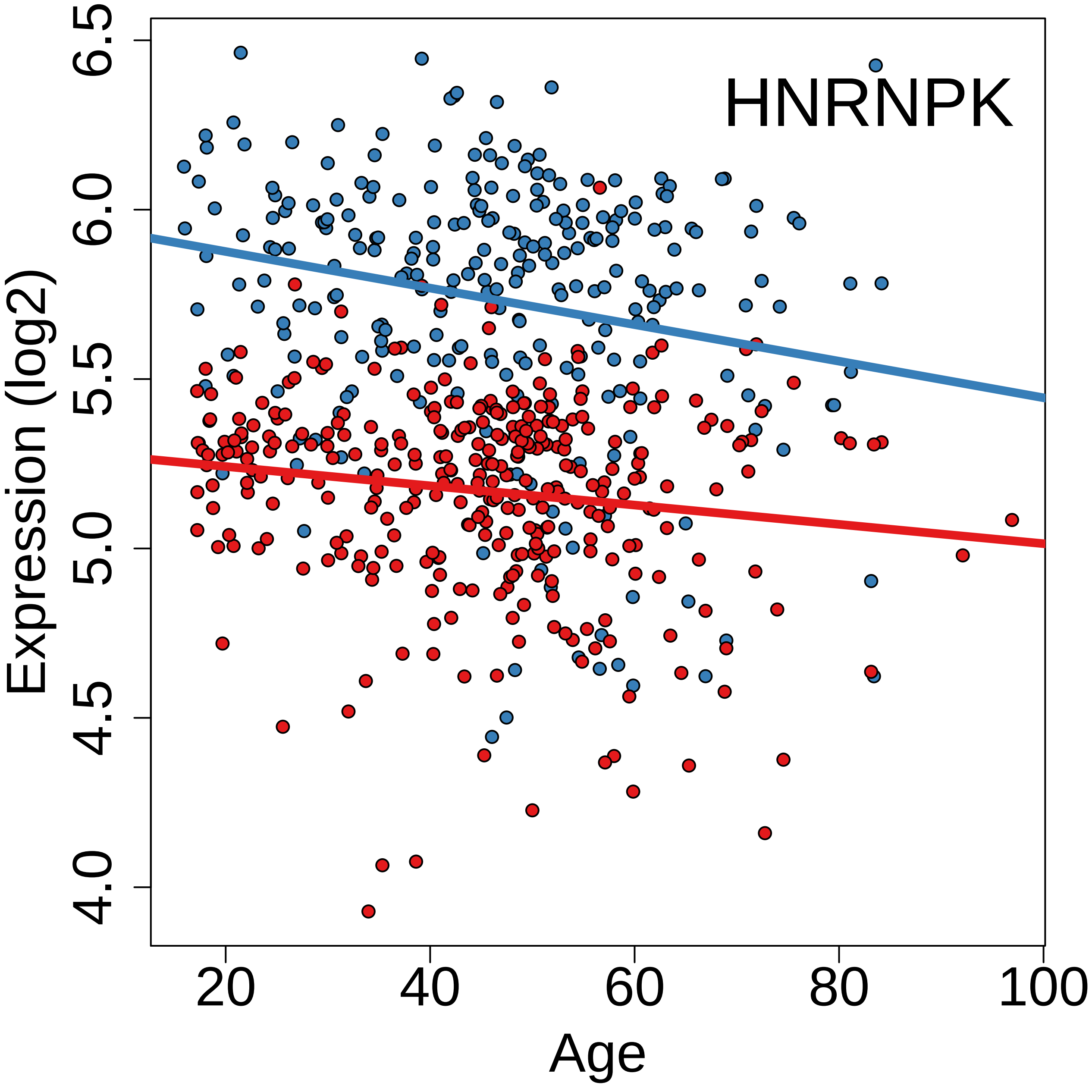
HNRNPK
Expression (log2)
6.5
6.0
5.5
5.0
4.5
4.0
Age
20
40
60
80
100

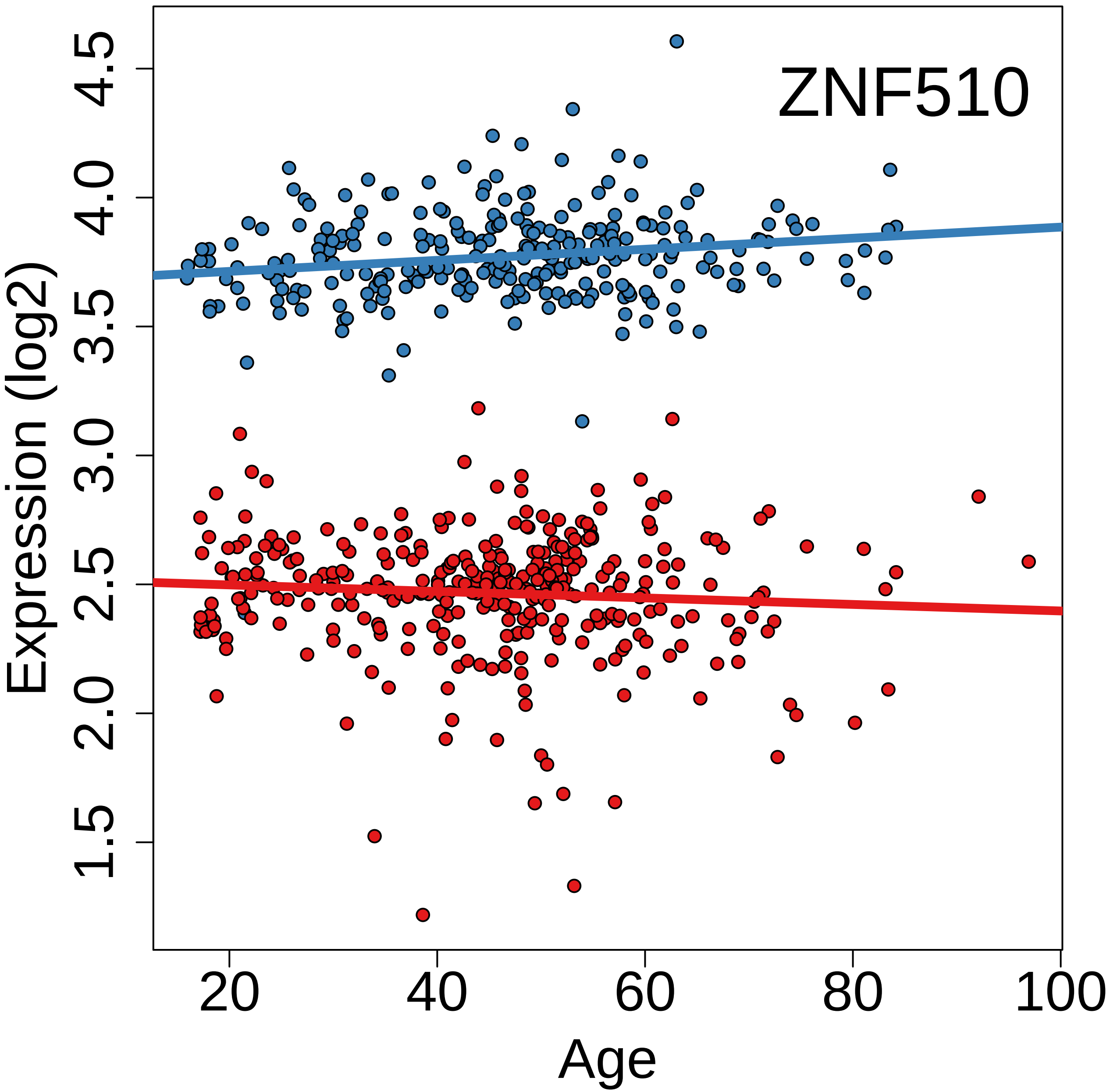
ZNF510
Expression (log2)
1.5
2.0
2.5
3.0
3.5
4.0
4.5
20
40
60
80
100
Age

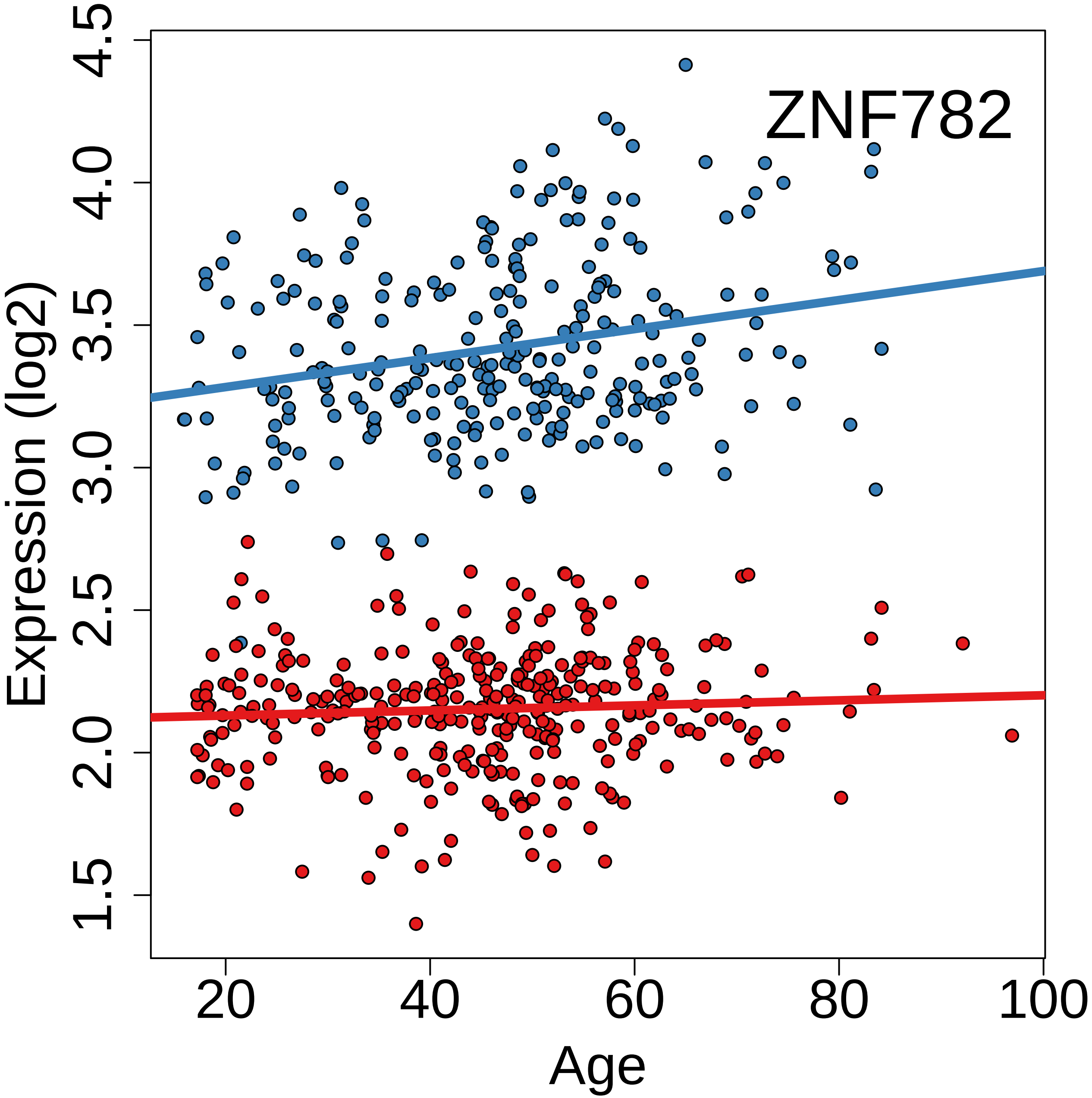
ZNF782
Expression (log2)
1.5
2.0
2.5
3.0
3.5
4.0
4.5
20
40
60
80
100
Age

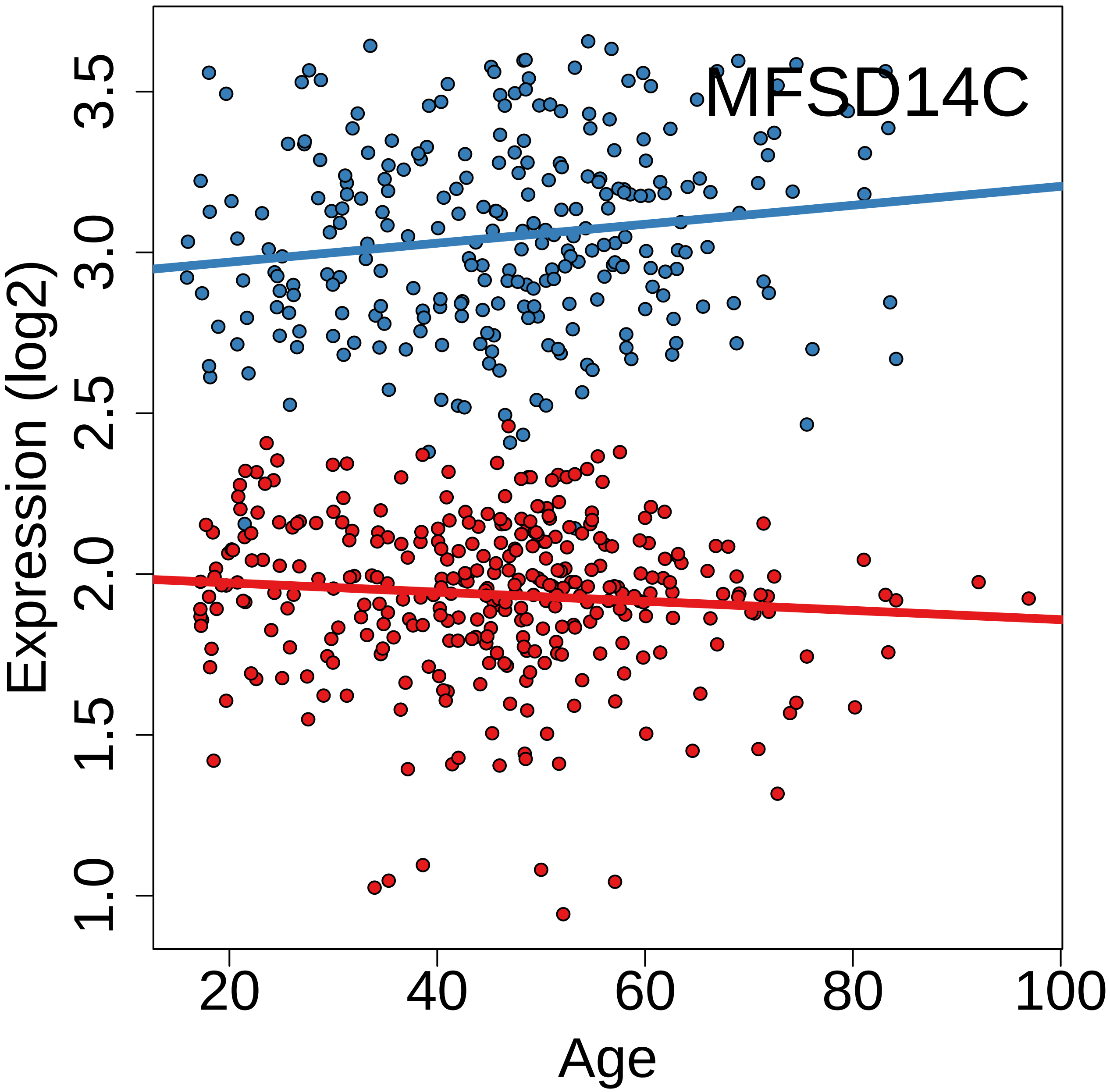
MFSD14C
Expression (log2)
1.0
1.5
2.0
2.5
3.0
3.5
20
40
60
80
100
Age

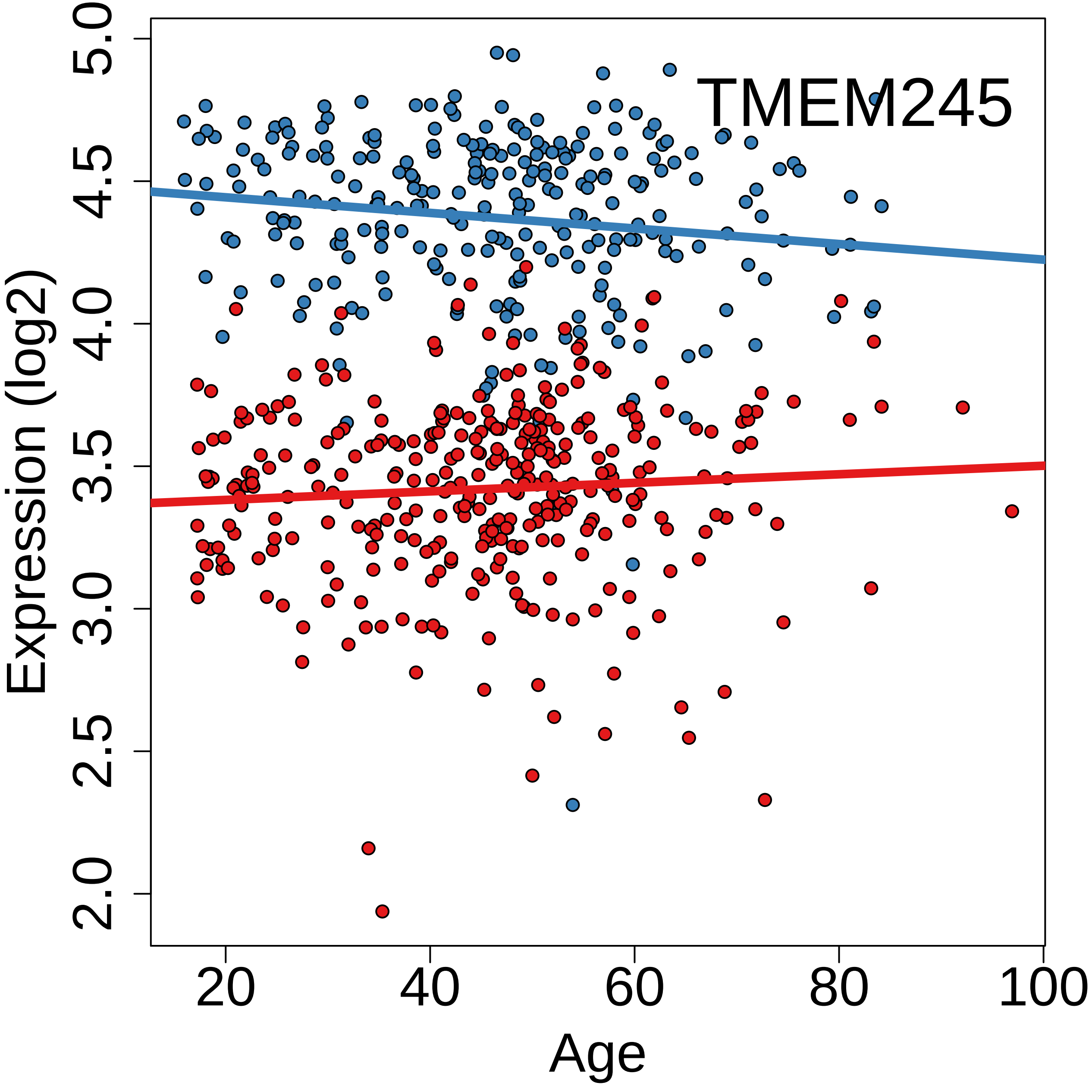
TMEM245
Expression (log2)
5.0
4.5
4.0
3.5
3.0
2.5
2.0
20
40
60
80
100
Age

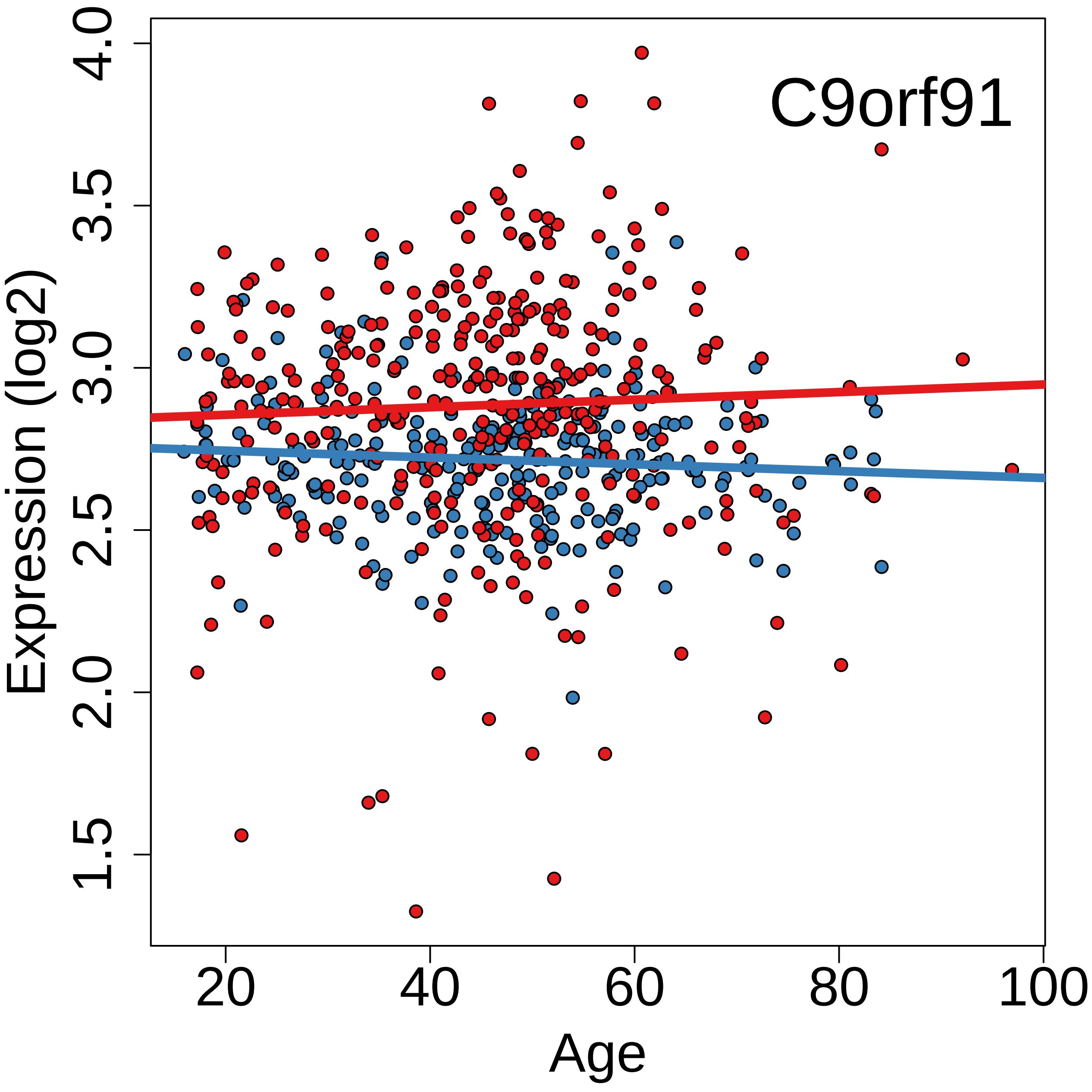
C9orf91
Expression (log2)
4.0
3.5
3.0
2.5
2.0
1.5
20
40
60
80
100
Age

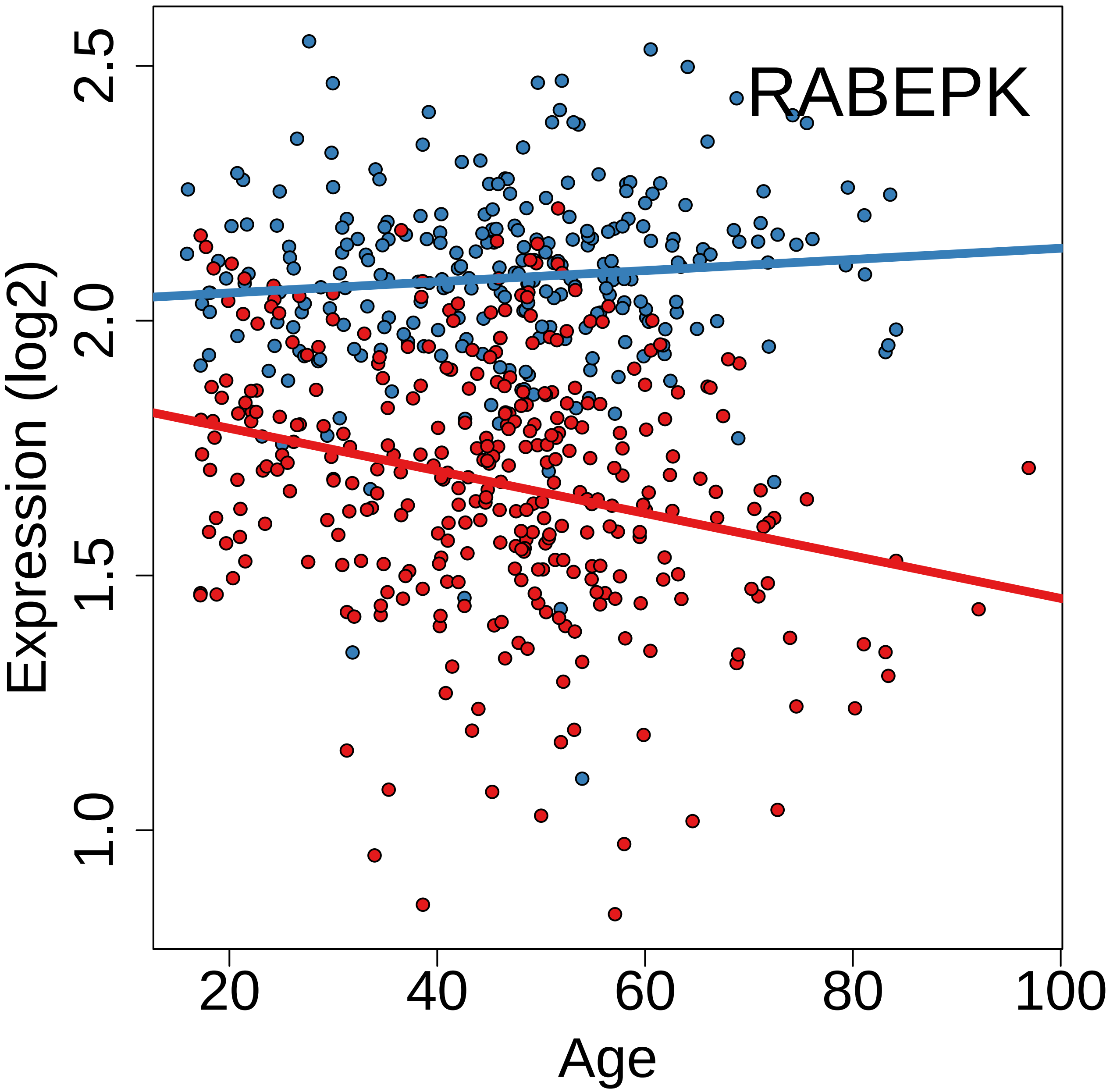
RABEPK
Expression (log2)
1.0
1.5
2.0
2.5
20
40
60
80
100
Age

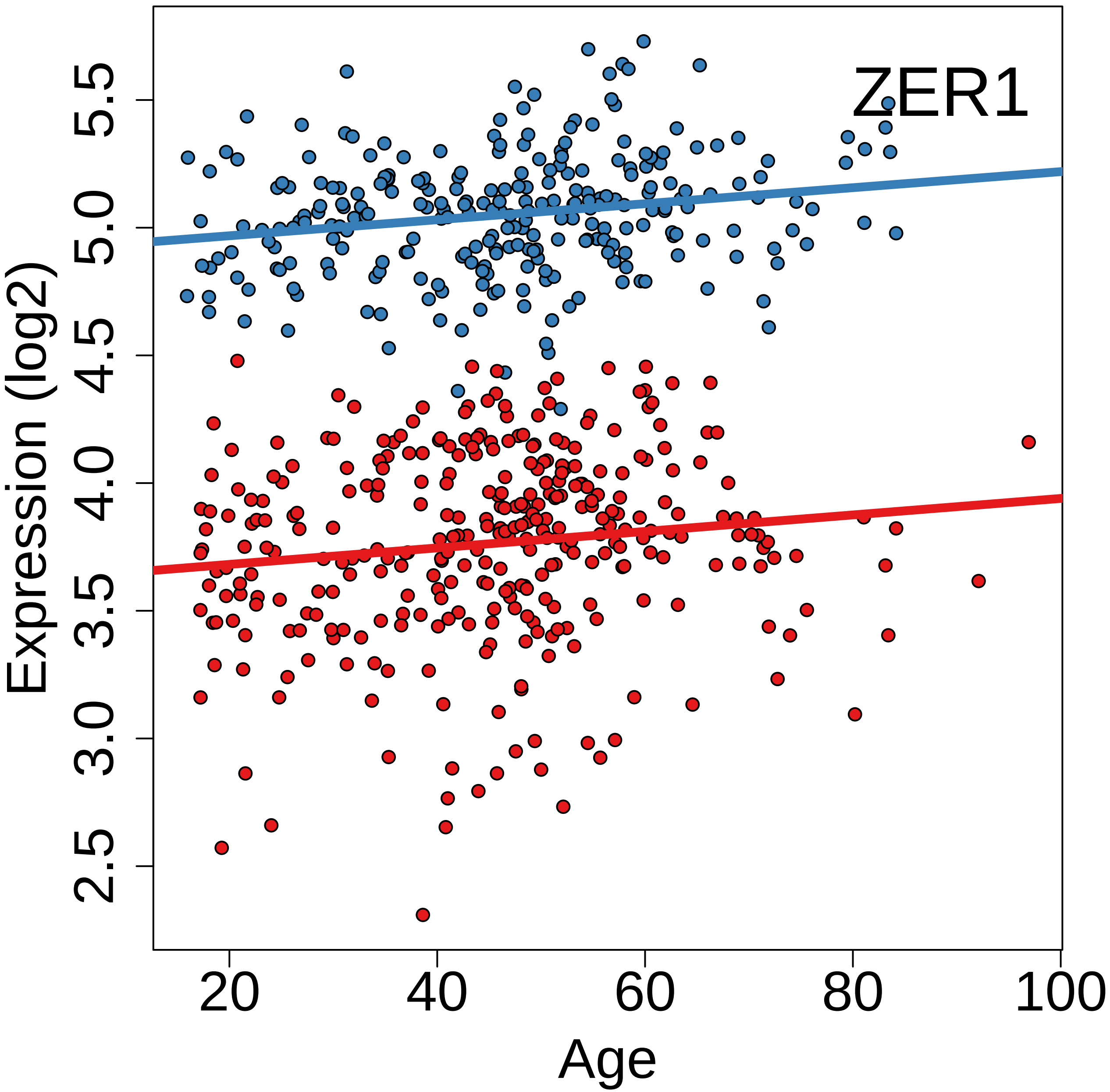
ZER1
Expression (log2)
2.5
3.0
3.5
4.0
4.5
5.0
5.5
20
40
60
80
100
Age

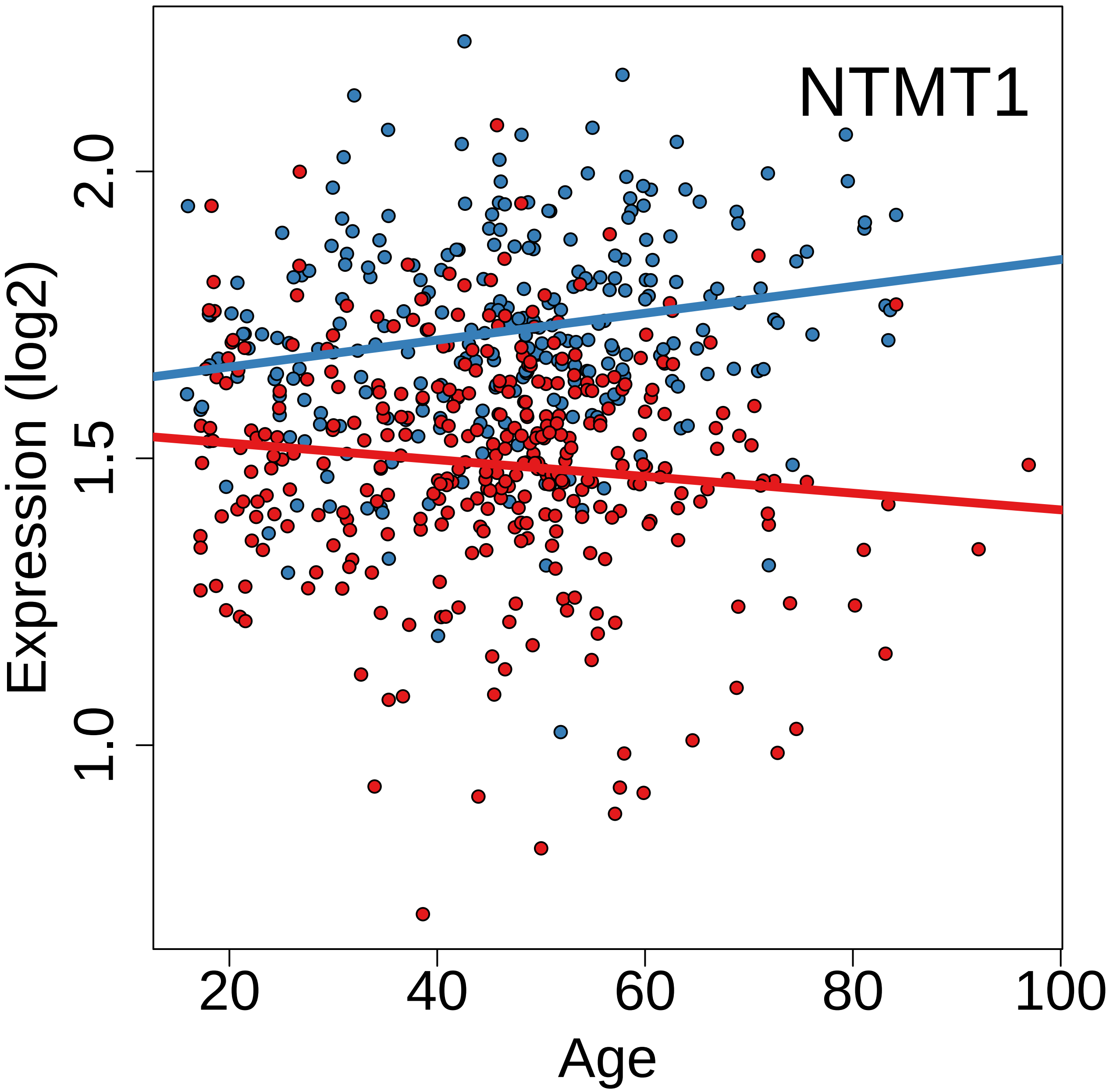
NTMT1
Expression (log2)
2.0
1.5
1.0
20
40
60
80
100
Age

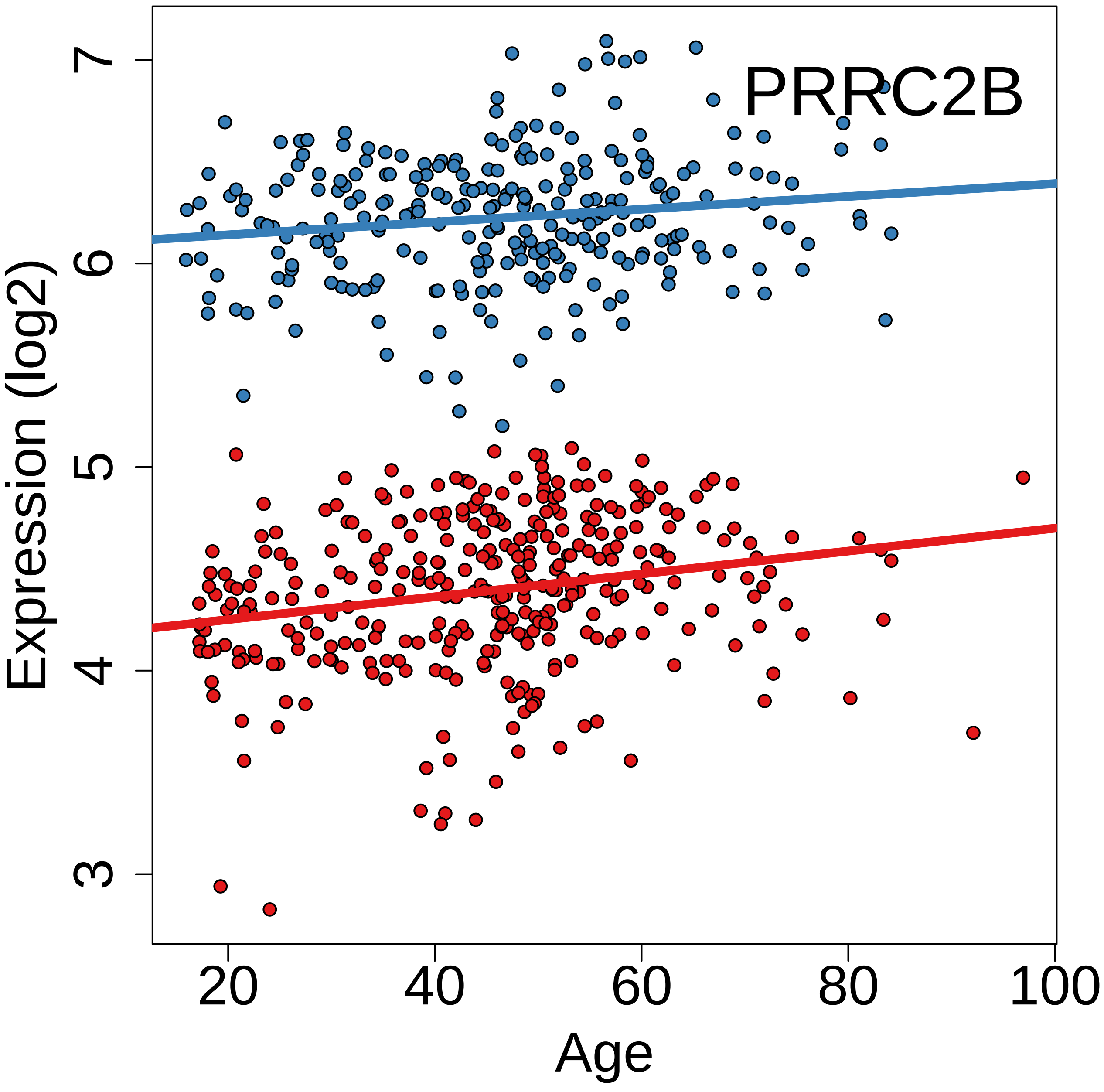
PRRC2B
Expression (log2)
3
4
5
6
7
20
40
60
80
100
Age

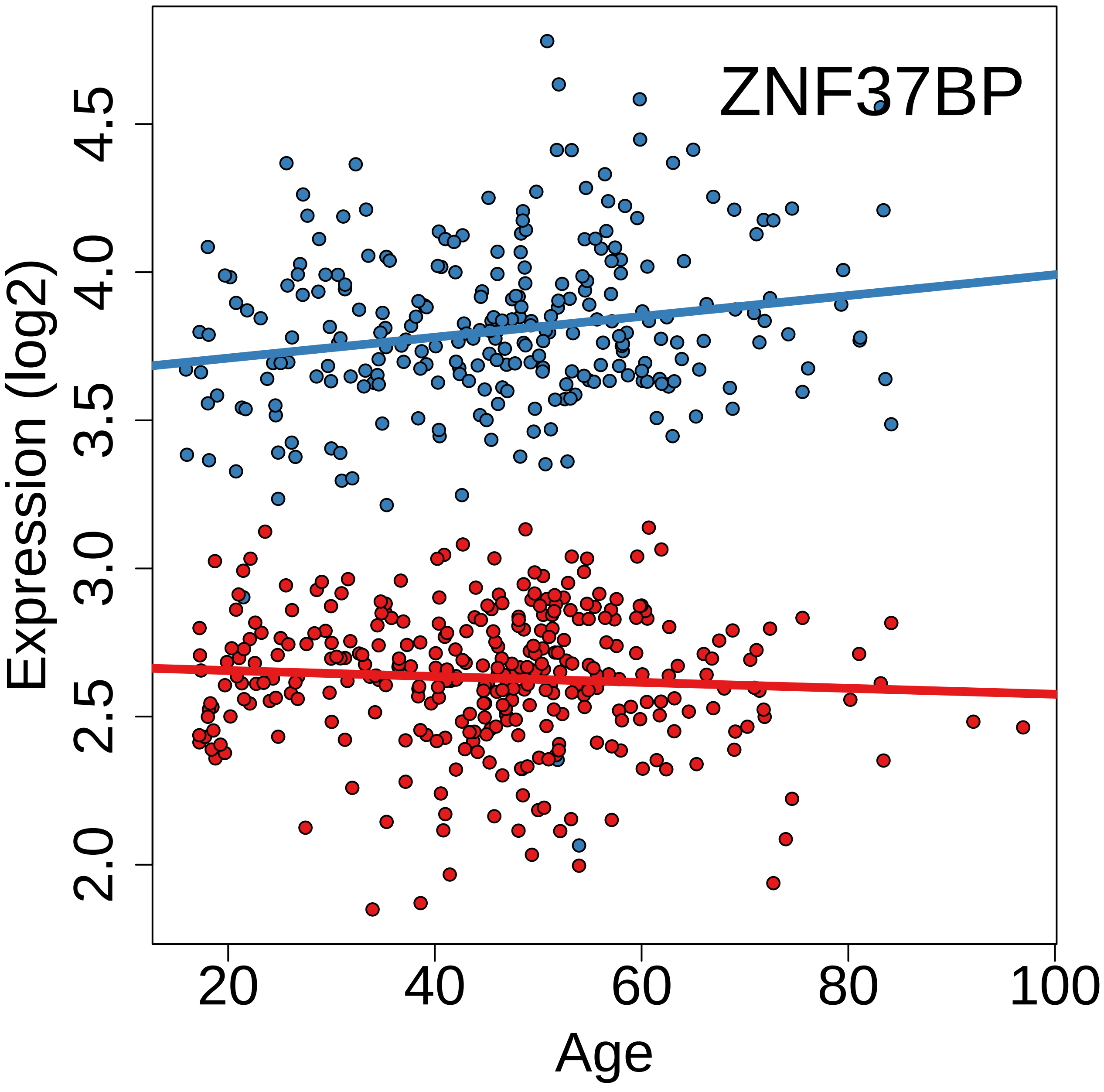
ZNF37BP
Expression (log2)
2.0
2.5
3.0
3.5
4.0
4.5
20
40
60
80
100
Age

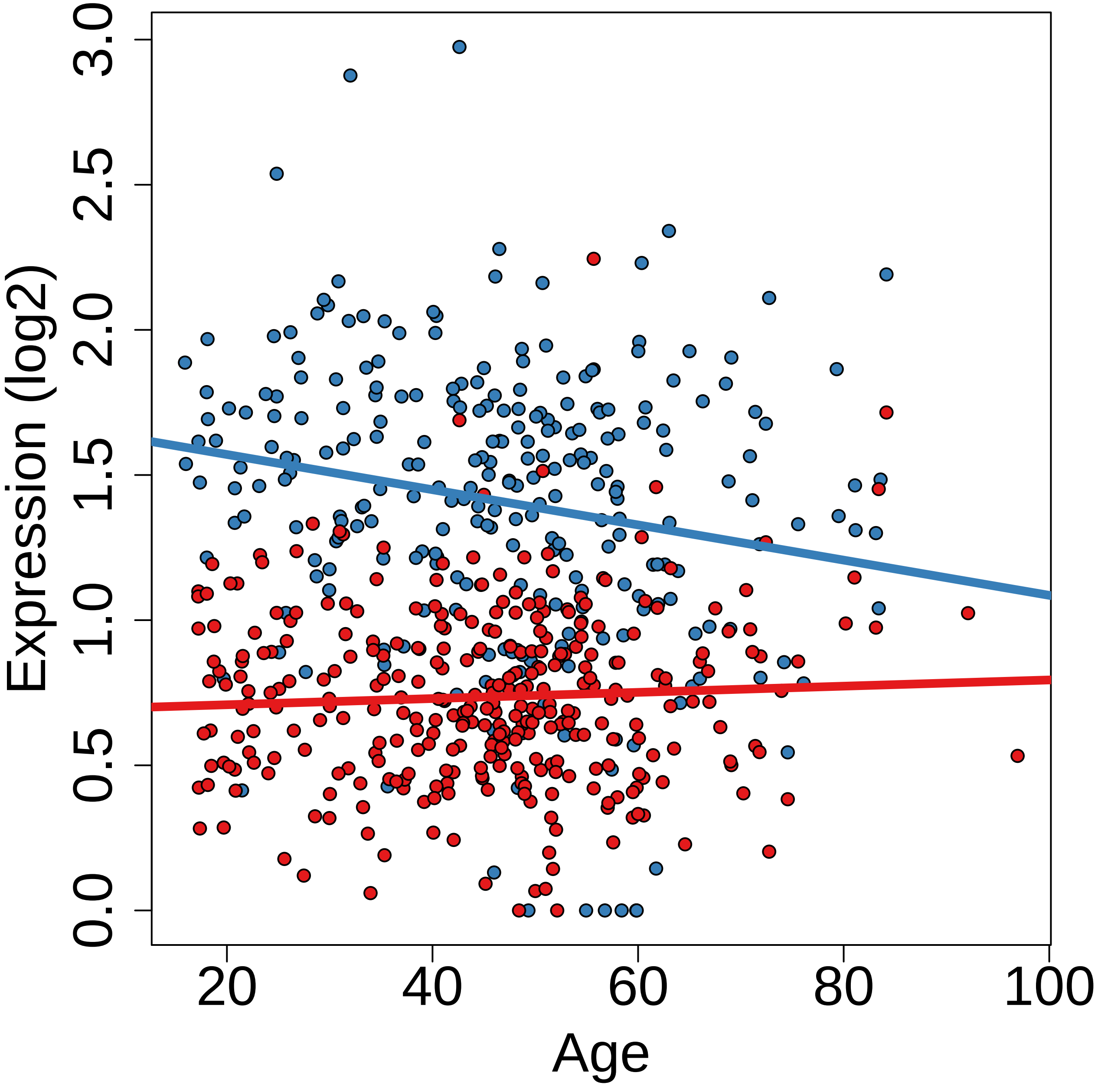

Expression (log2)
0.0
0.5
1.0
1.5
2.0
2.5
3.0
20
40
60
80
100
Age

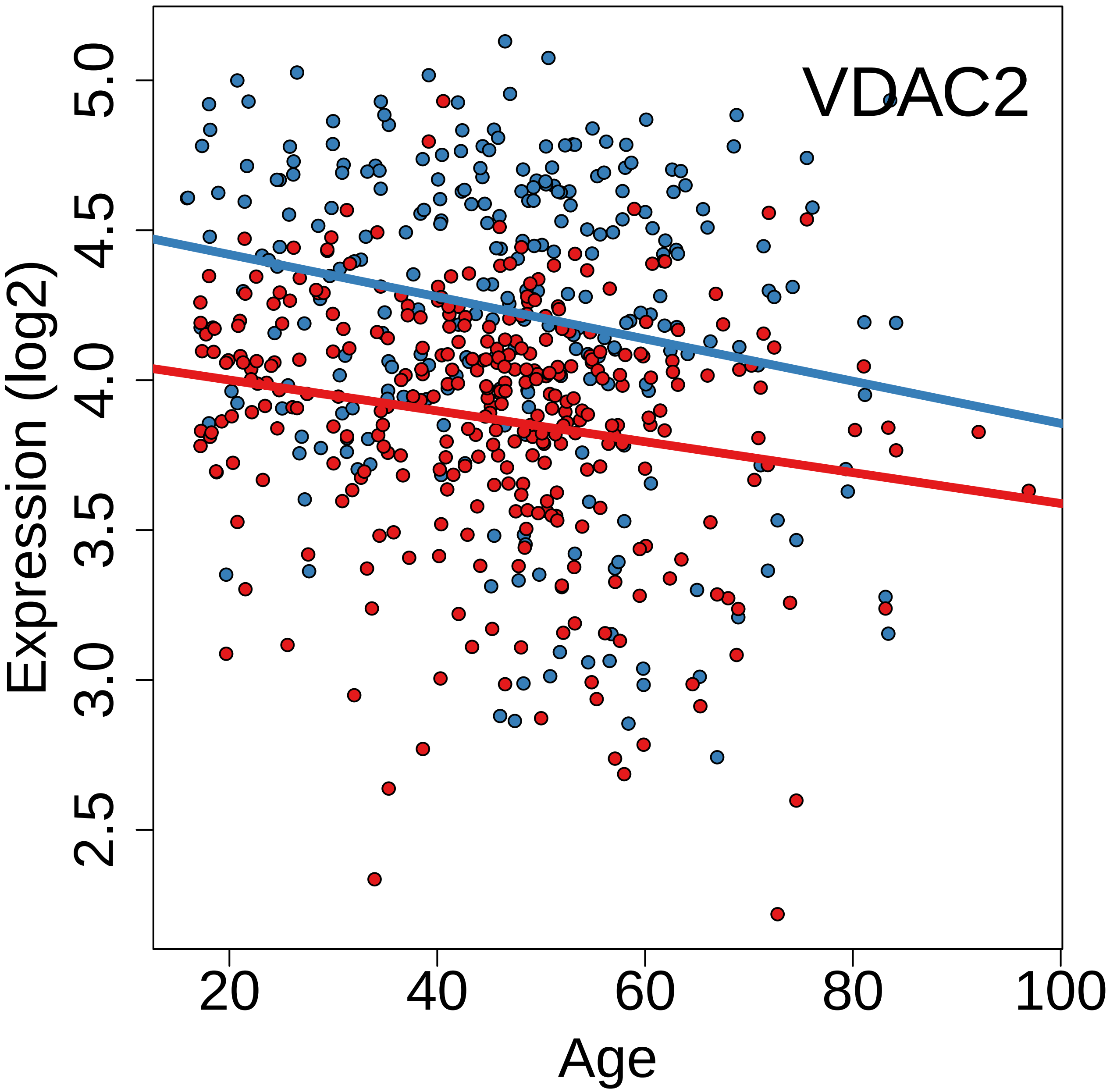
VDAC2
Expression (log2)
5.0
4.5
4.0
3.5
3.0
2.5
20
40
60
80
100
Age

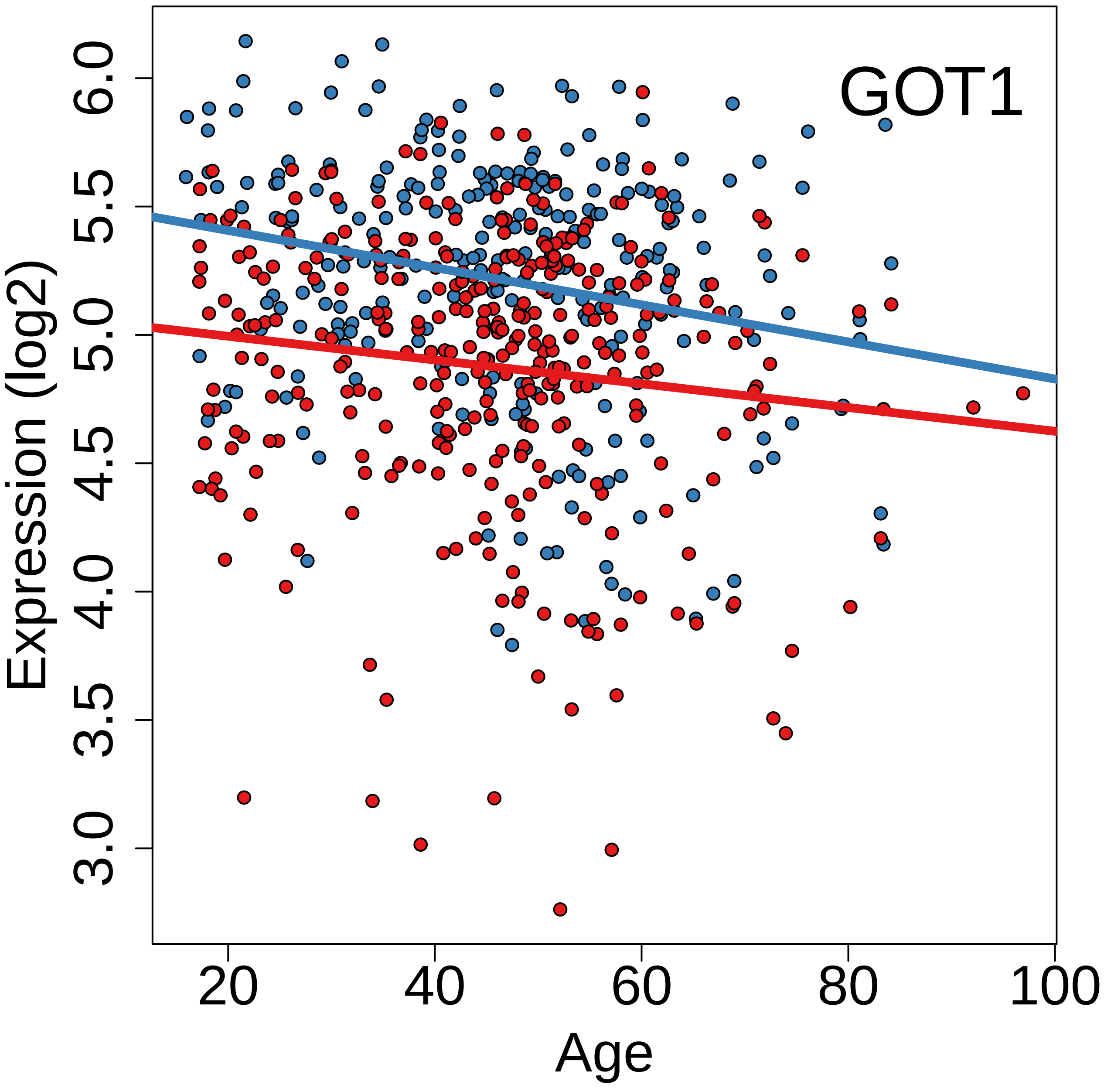
GOT1
Expression (log2)
3.0
3.5
4.0
4.5
5.0
5.5
6.0
20
40
60
80
100
Age

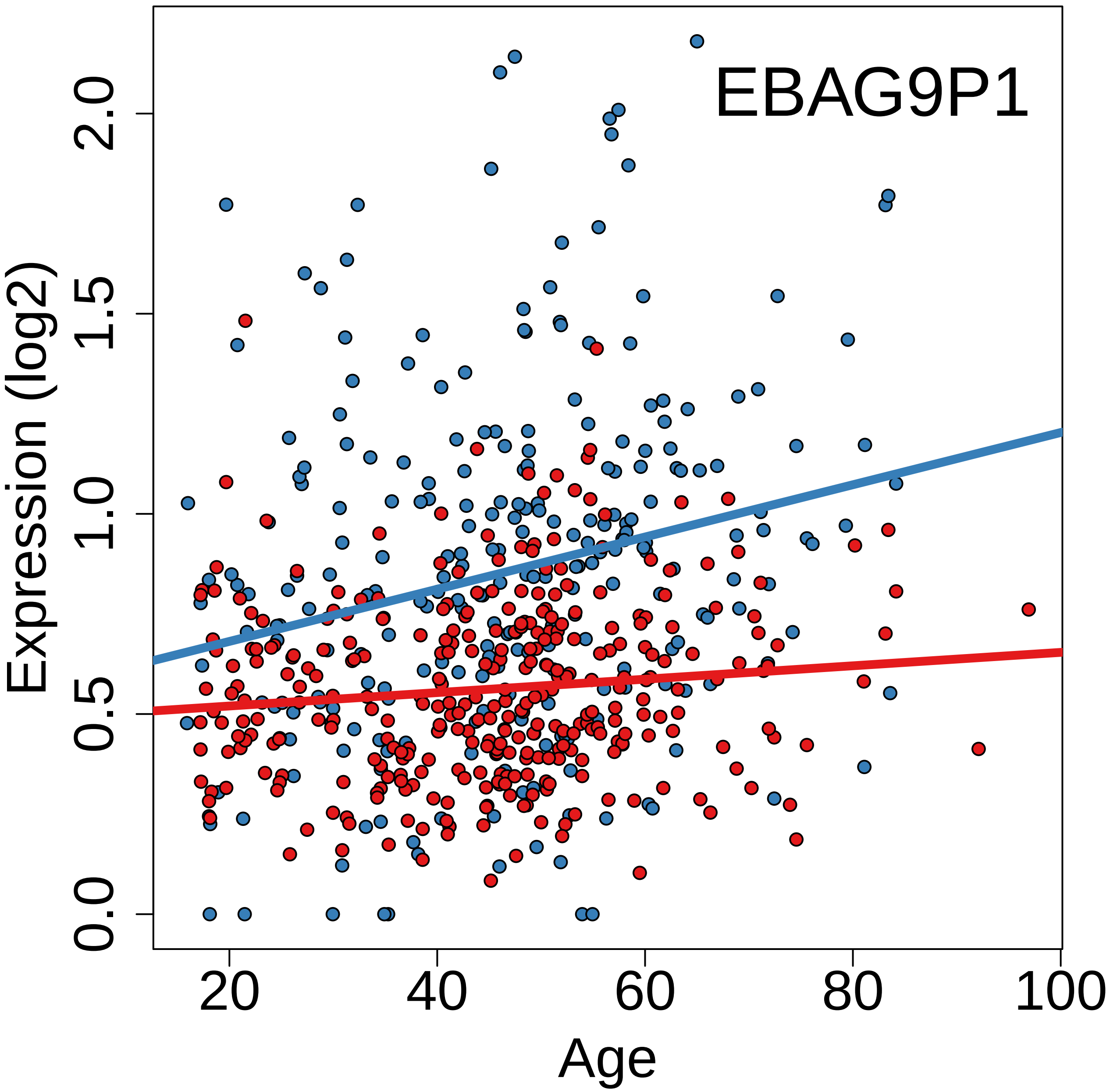
EBAG9P1
Expression (log2)
0.0
0.5
1.0
1.5
2.0
20
40
60
80
100
Age

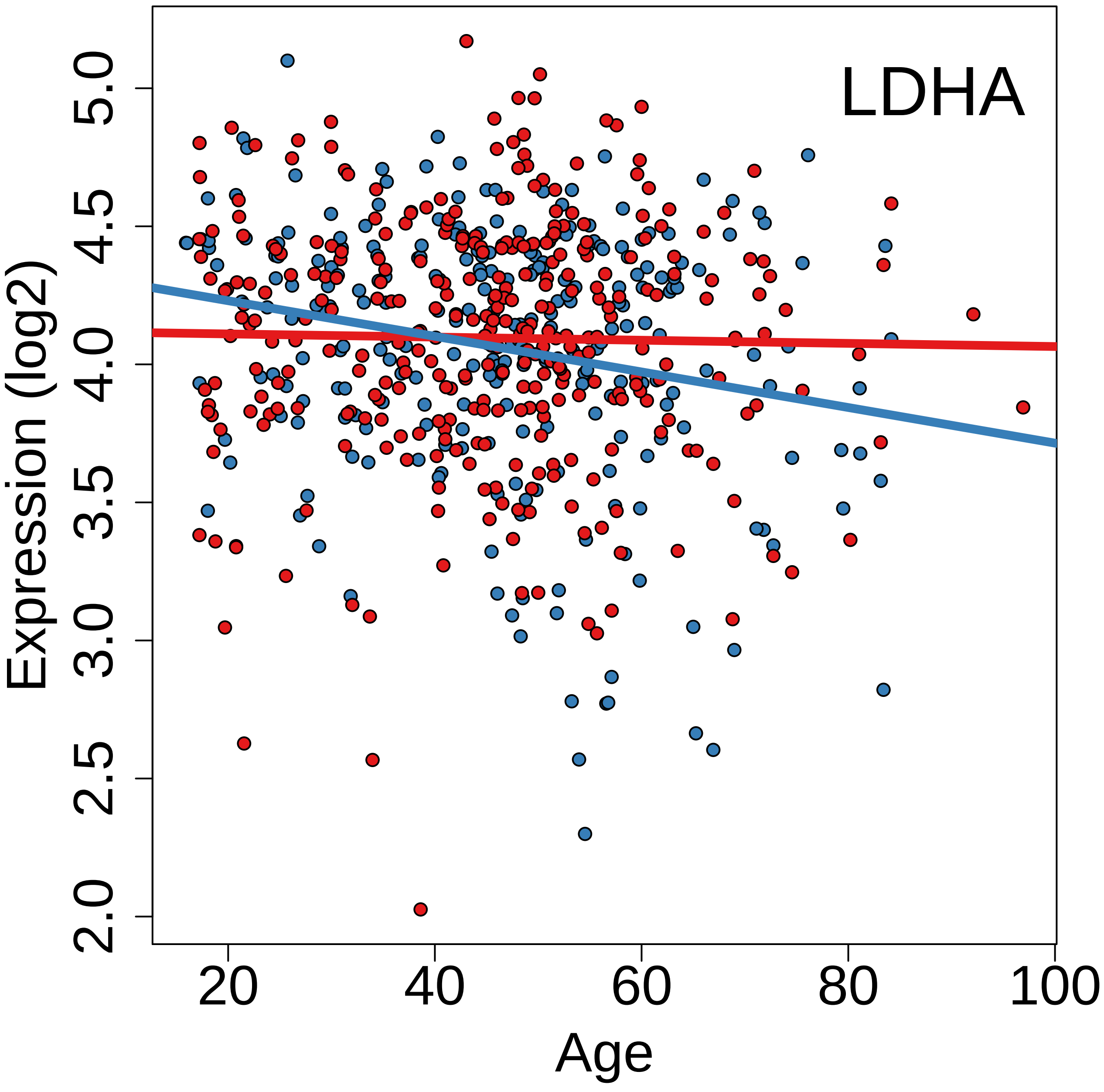
LDHA
Expression (log2)
2.0
2.5
3.0
3.5
4.0
4.5
5.0
20
40
60
80
100
Age

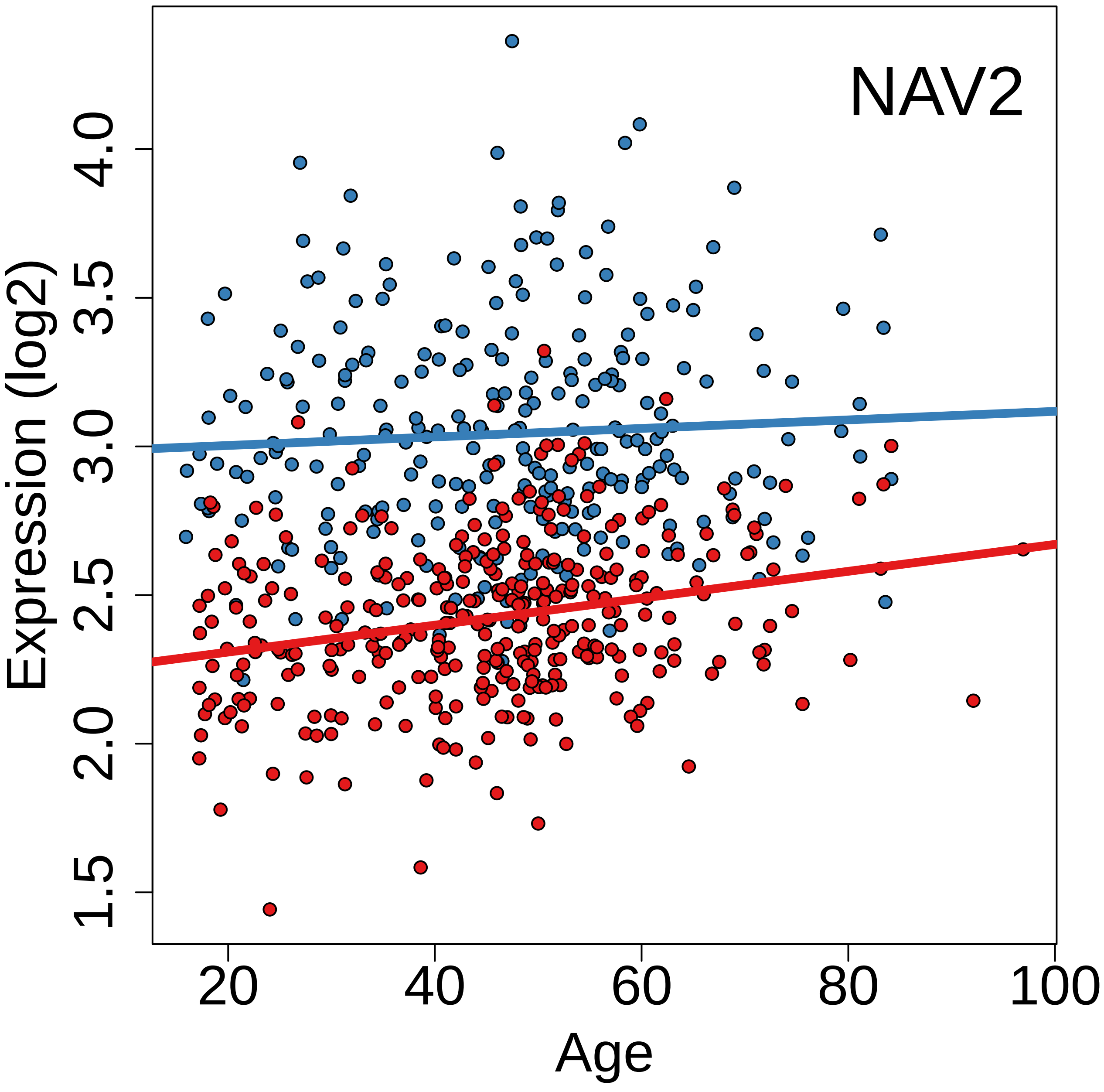

NAV2
Expression (log2)
1.5
2.0
2.5
3.0
3.5
4.0
20
40
60
80
100
Age

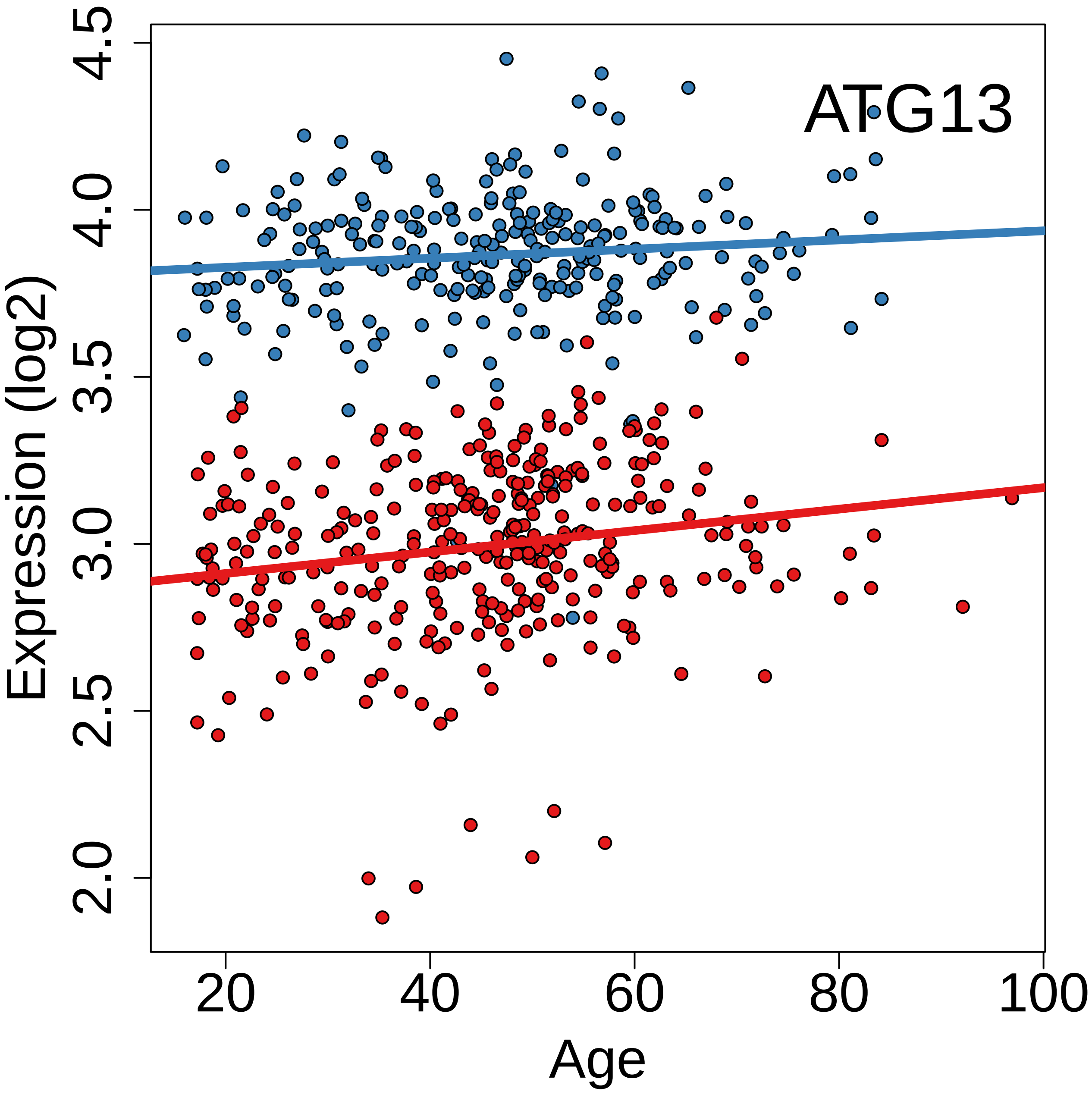
ATG13
Expression (log2)
4.5
4.0
3.5
3.0
2.5
2.0
20
40
60
80
100
Age

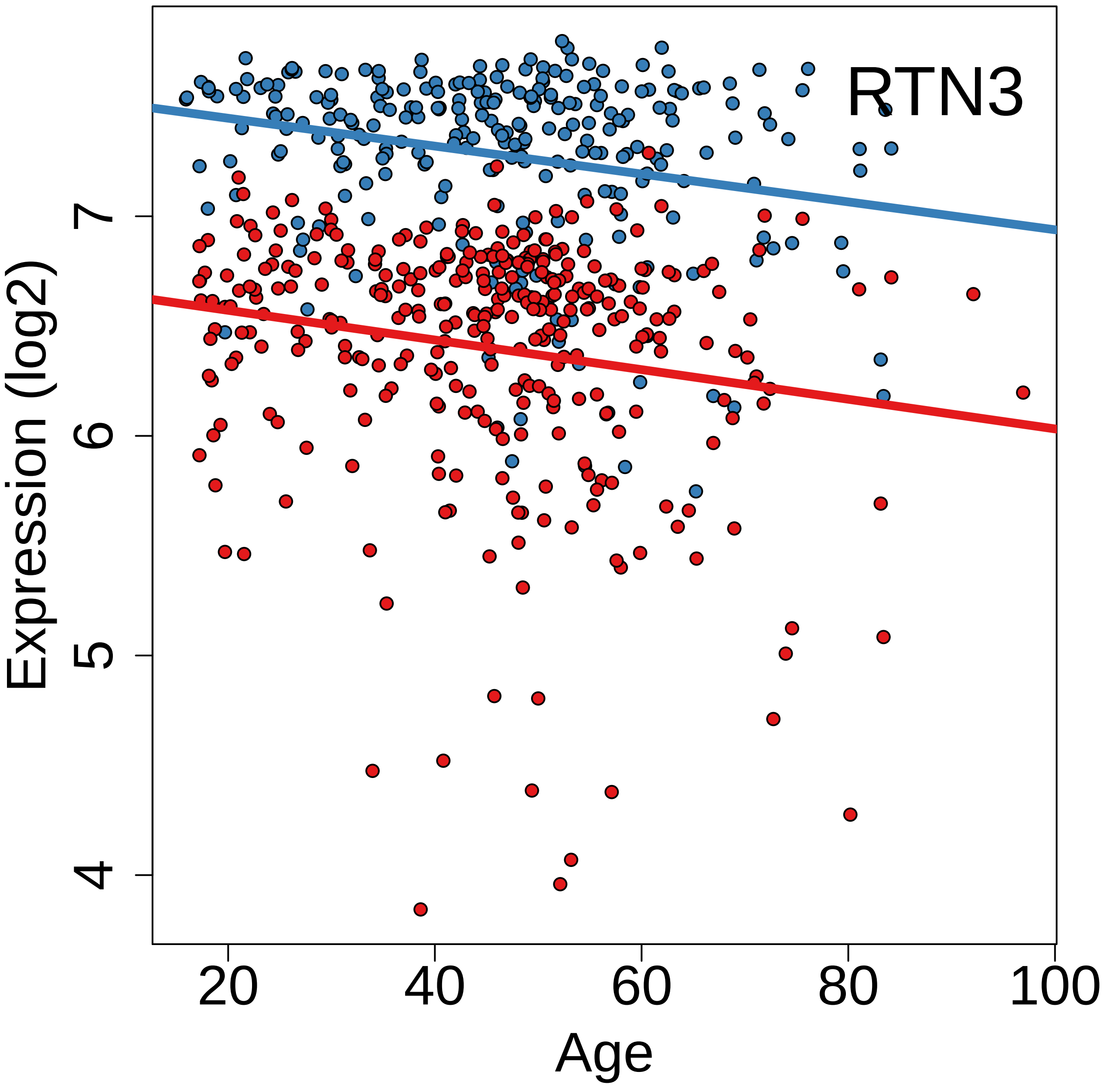

RTN3
Expression (log2)
4
5
6
7
20
40
60
80
100
Age

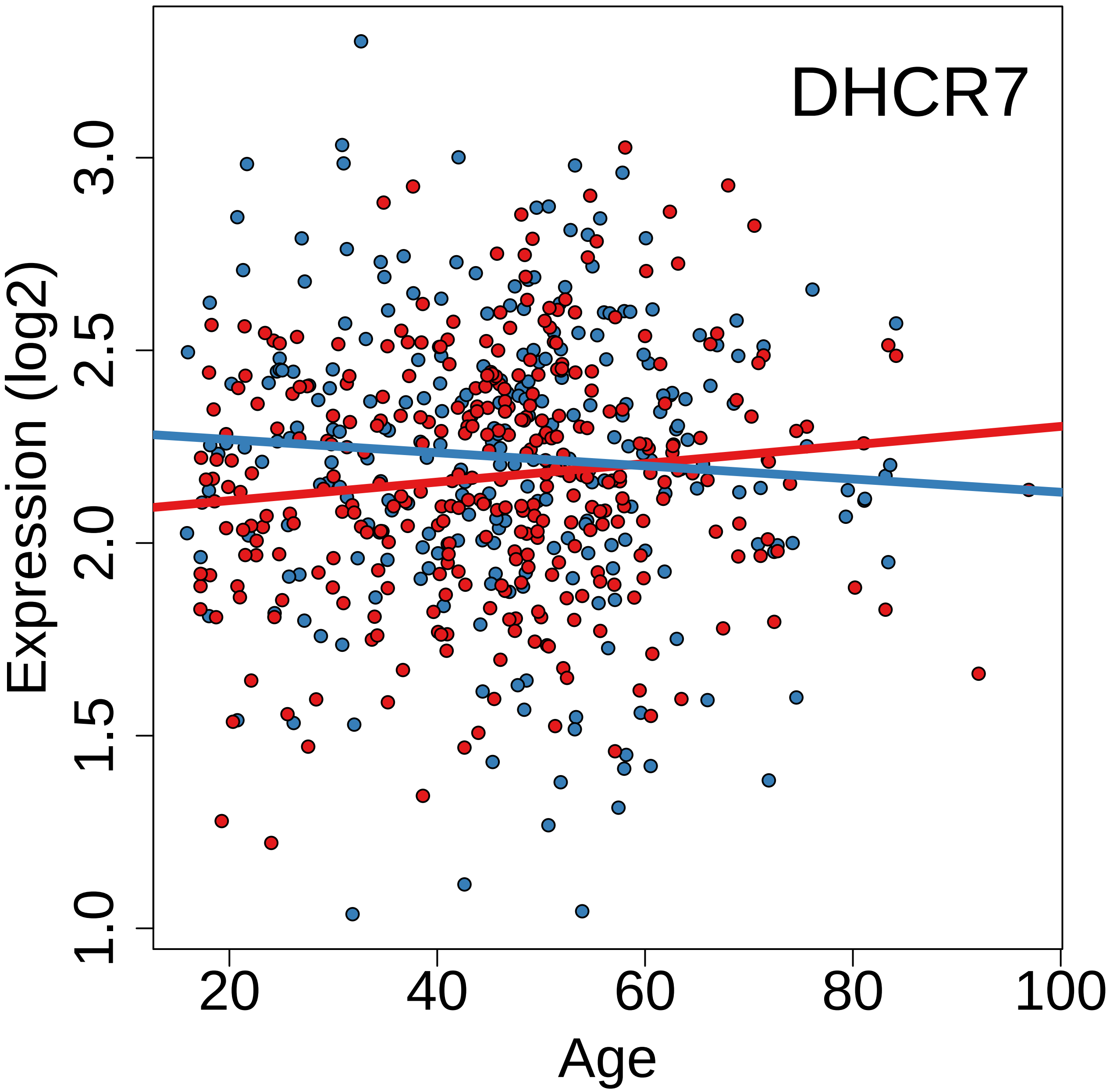

DHCR7
Expression (log2)
1.0
1.5
2.0
2.5
3.0
20
40
60
80
100
Age

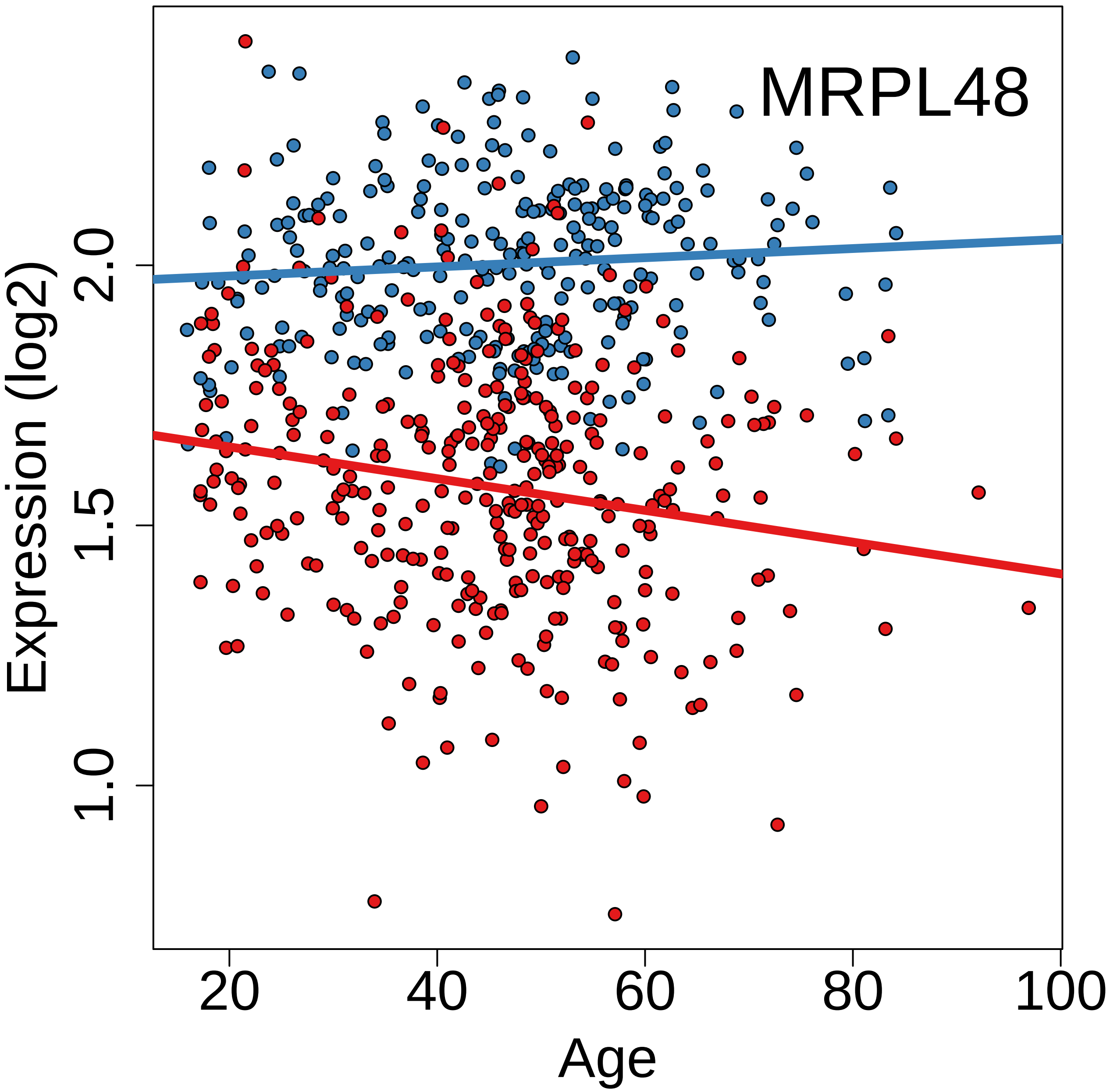
MRPL48
Expression (log2)
2.0
1.5
1.0
20
40
60
80
100
Age

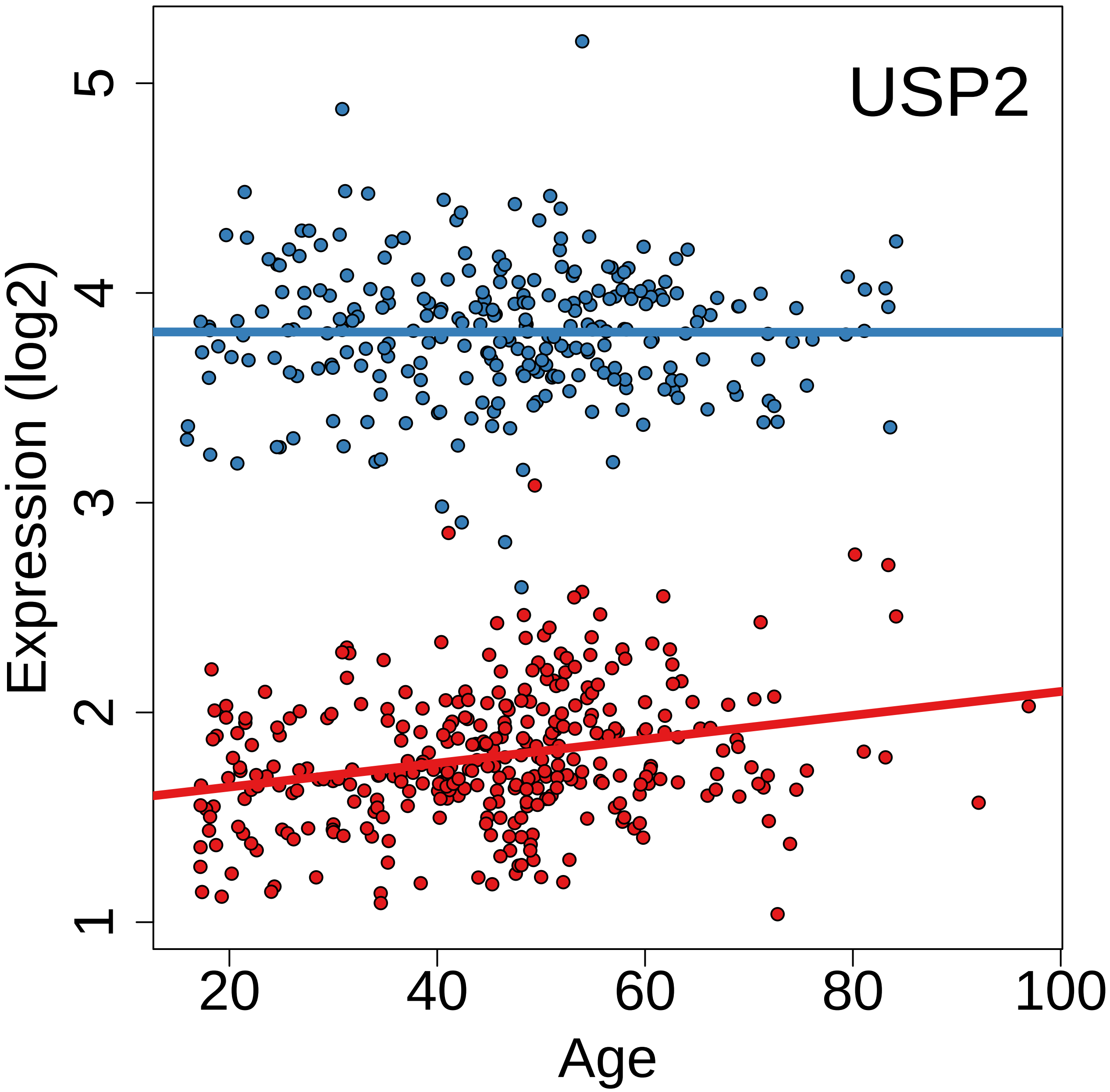
USP2
Expression (log2)
1
2
3
4
5
20
40
60
80
100
Age

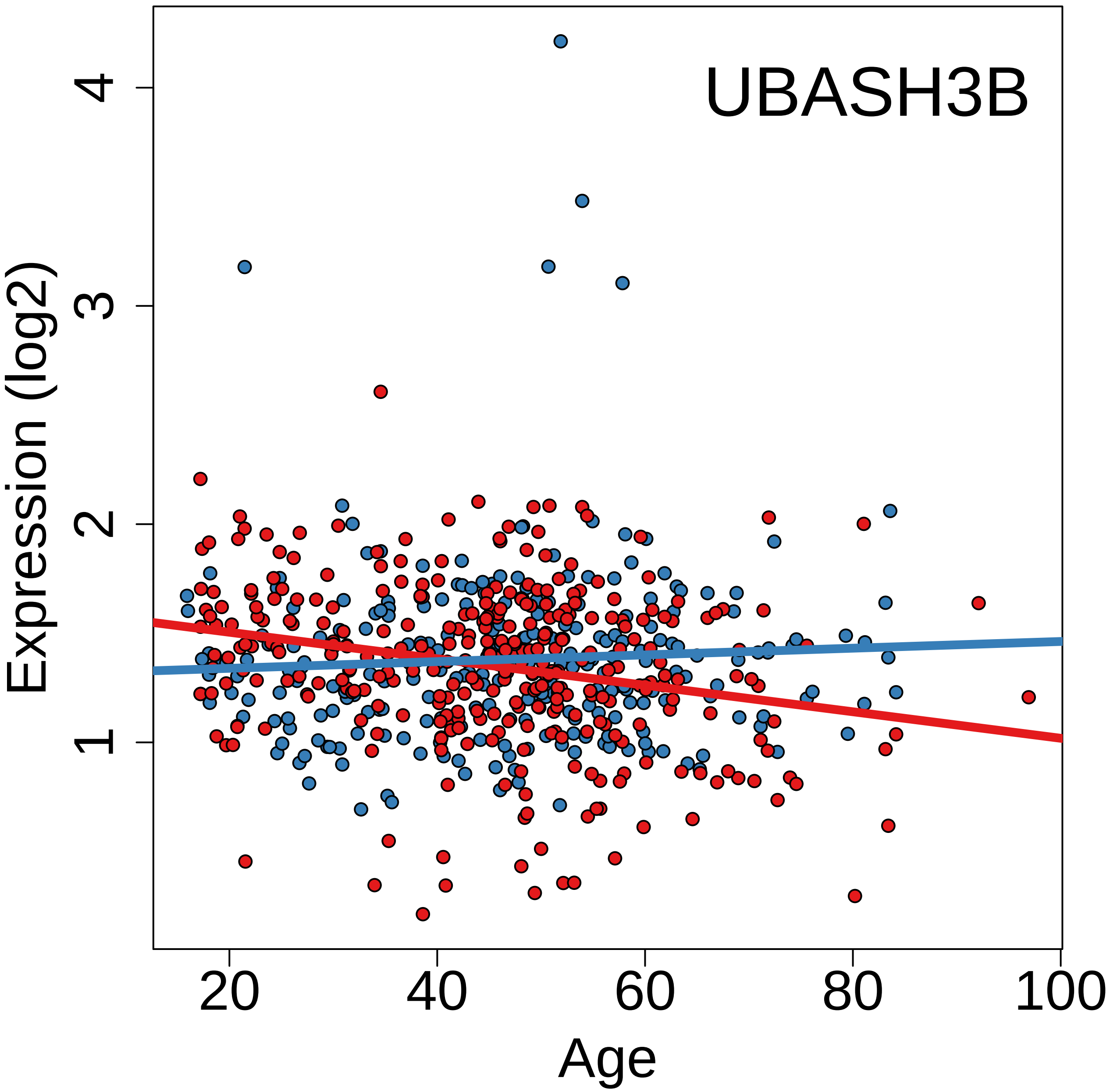
UBASH3B
Expression (log2)
Age
1
2
3
4
20
40
60
80
100

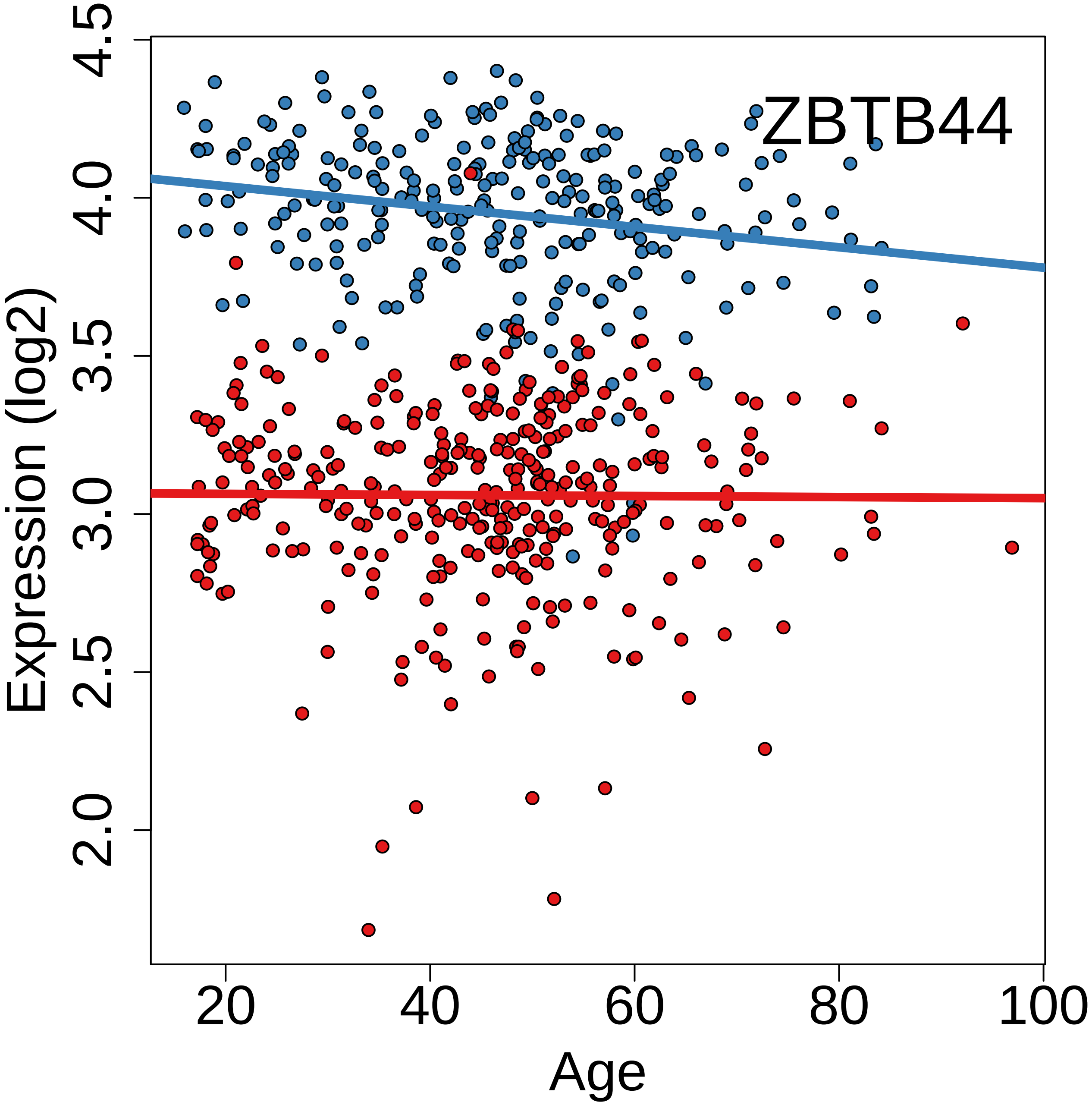
ZBTB44
Expression (log2)
Age
2.0
2.5
3.0
3.5
4.0
4.5
20
40
60
80
100

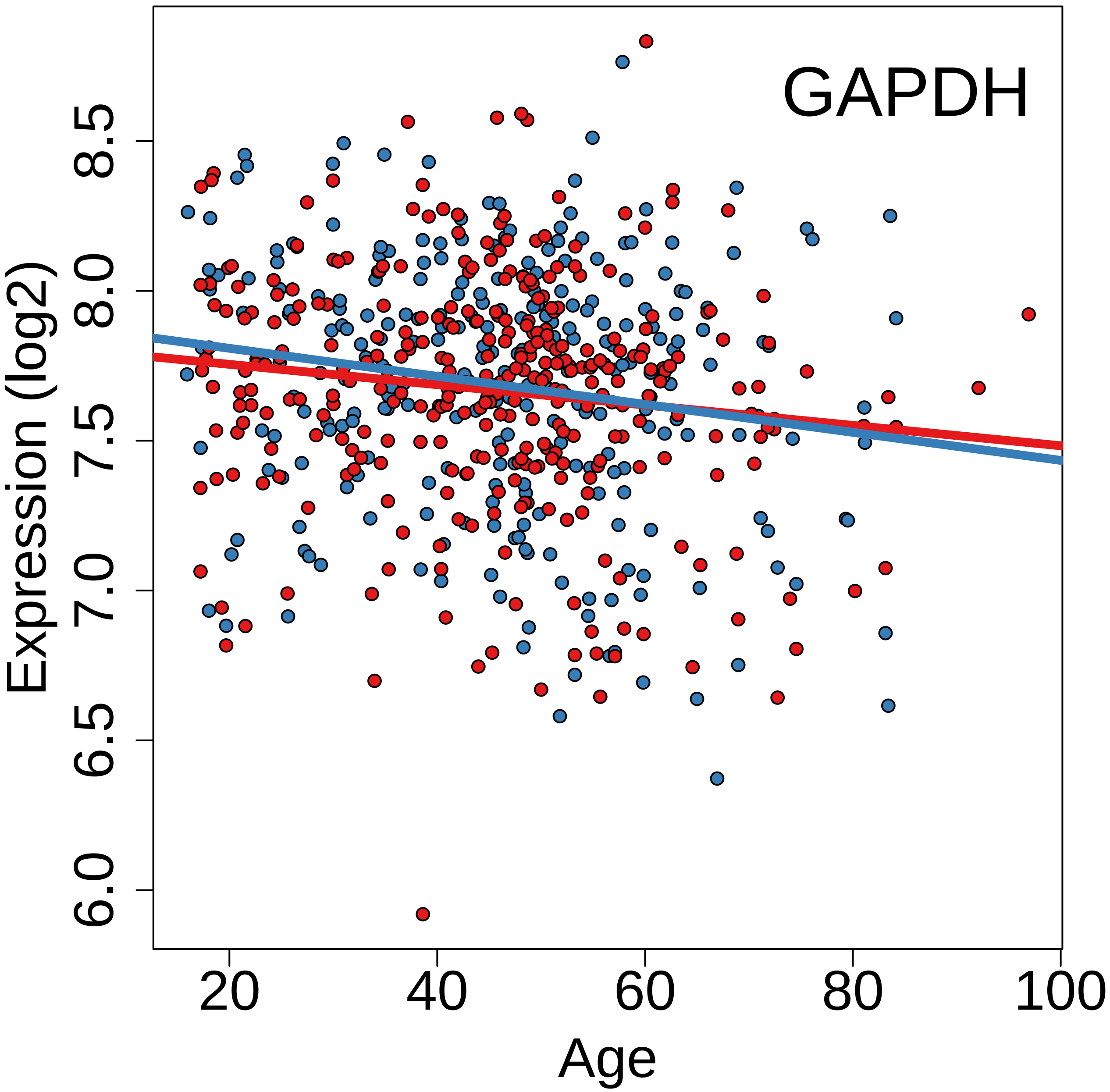

GAPDH
Expression (log2)
Age
6.0
6.5
7.0
7.5
8.0
8.5
20
40
60
80
100

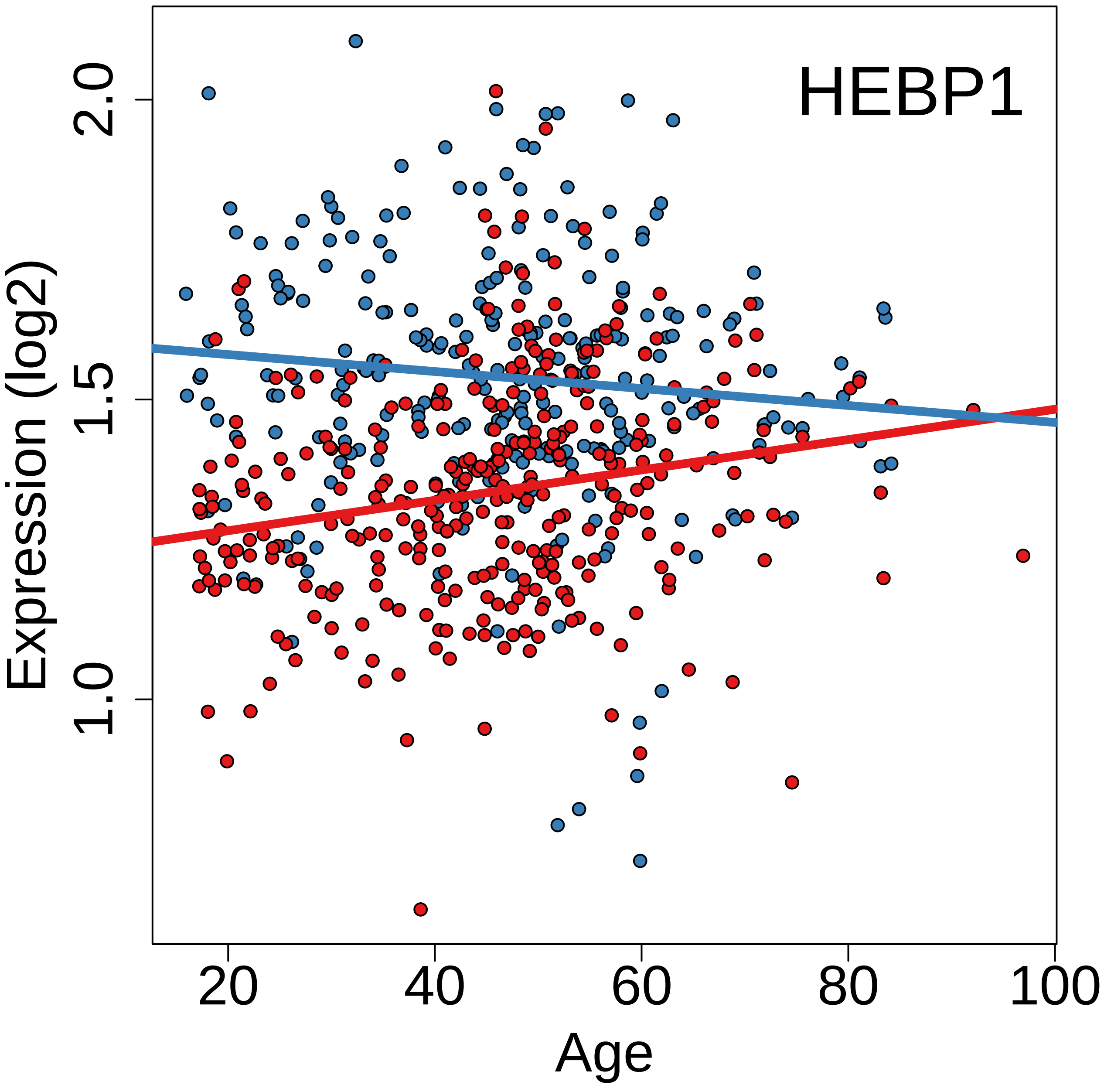
HEBP1
Expression (log2)
1.0
1.5
2.0
20
40
60
80
100
Age

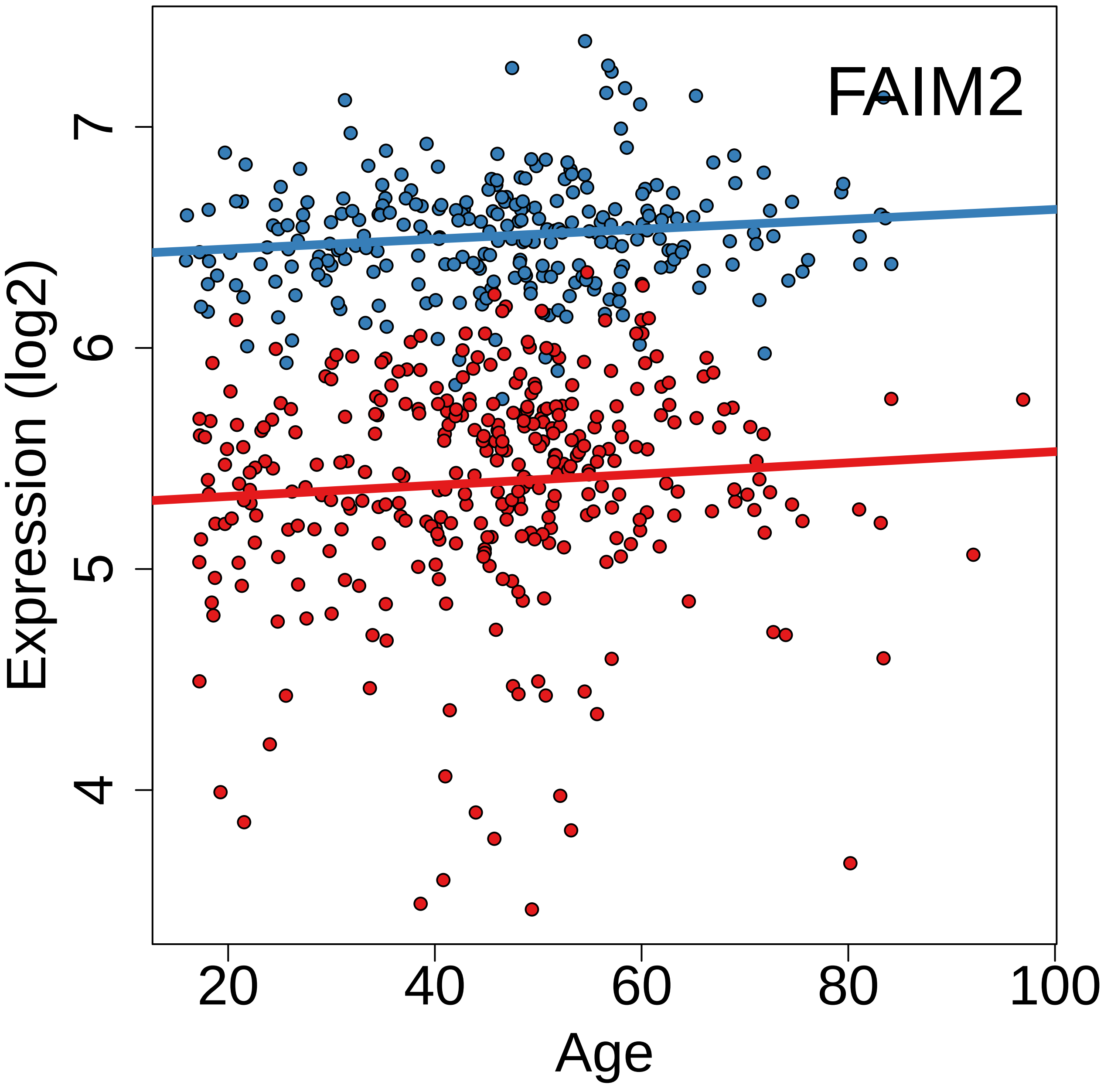
FAIM2
Expression (log2)
Age
4
5
6
7
20
40
60
80
100

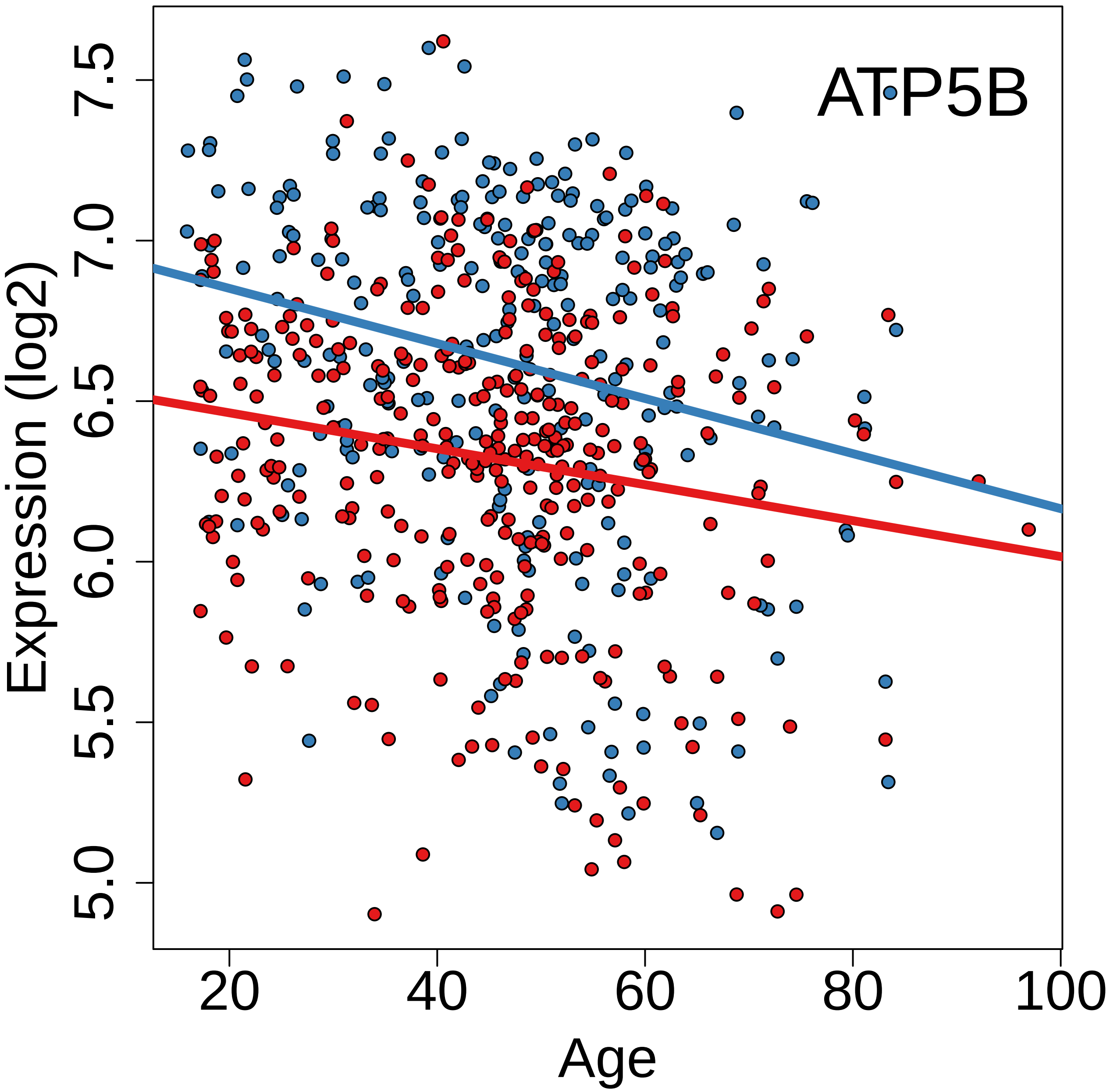
ATP5B
Expression (log2)
5.0
5.5
6.0
6.5
7.0
7.5
20
40
60
80
100
Age

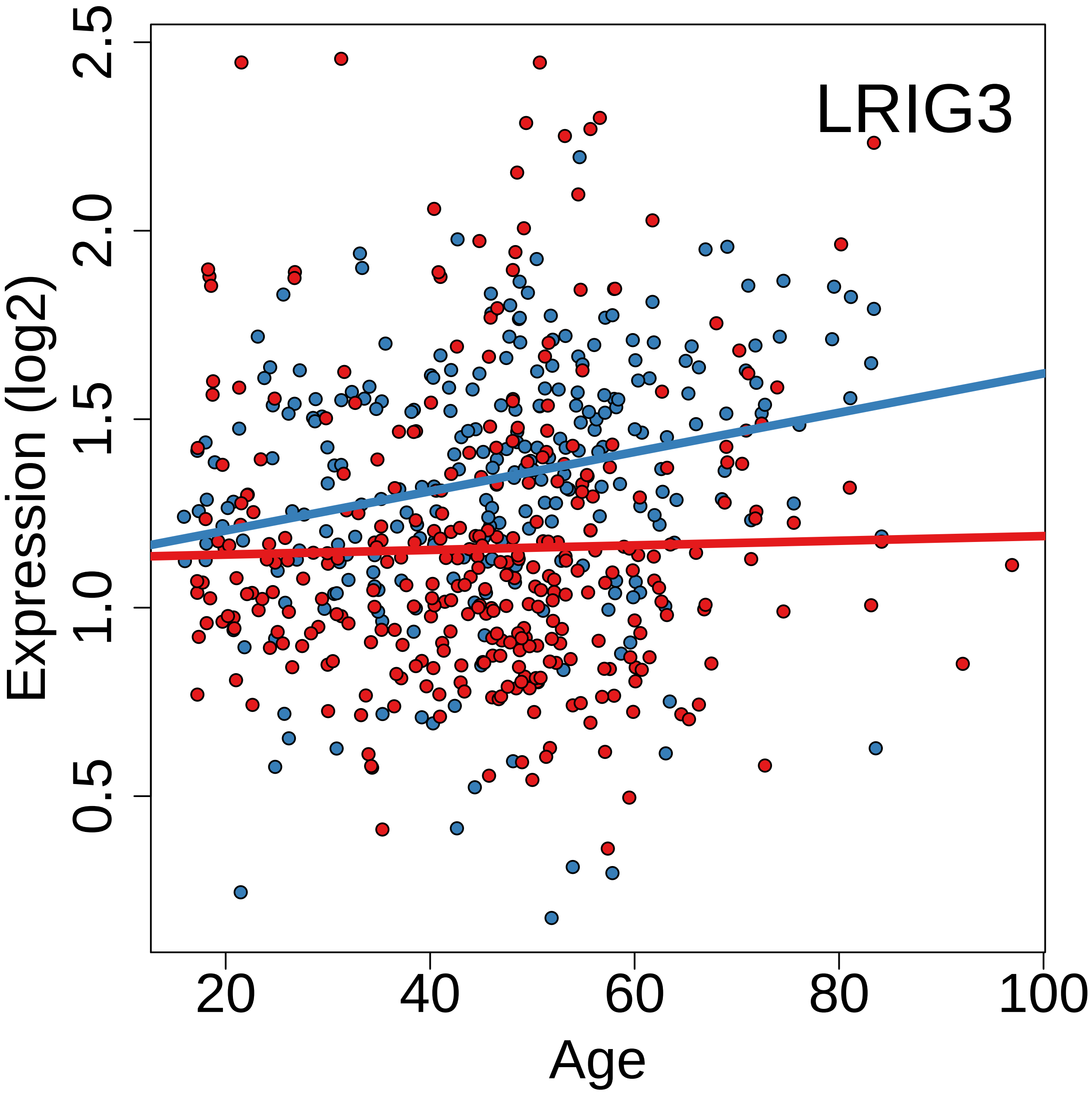
LRIG3
Expression (log2)
Age
0.5
1.0
1.5
2.0
2.5
20
40
60
80
100

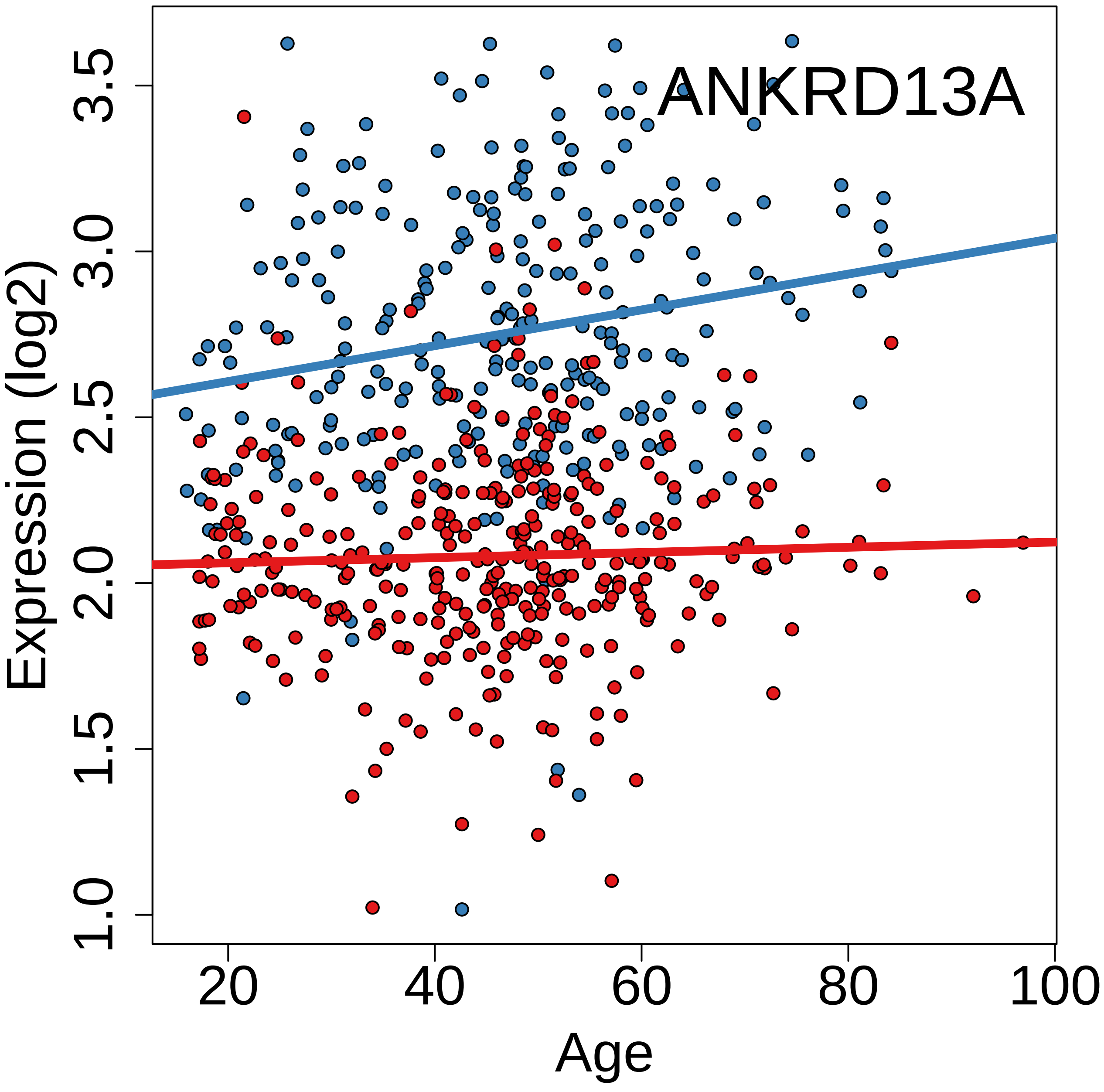
ANKRD13A
Expression (log2)
1.0
1.5
2.0
2.5
3.0
3.5
20
40
60
80
100
Age

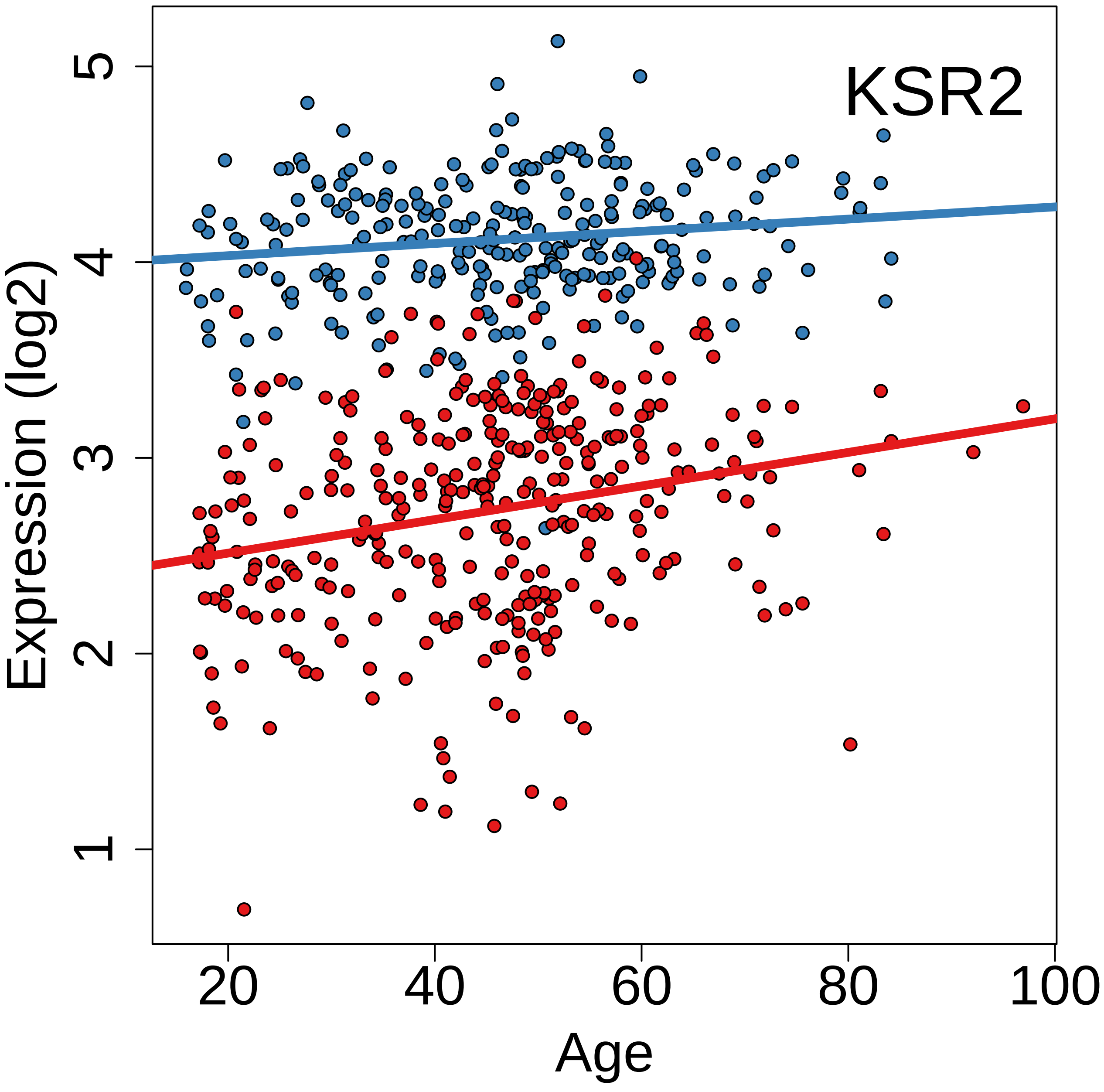
KSR2
Expression (log2)
Age
1
2
3
4
5
20
40
60
80
100

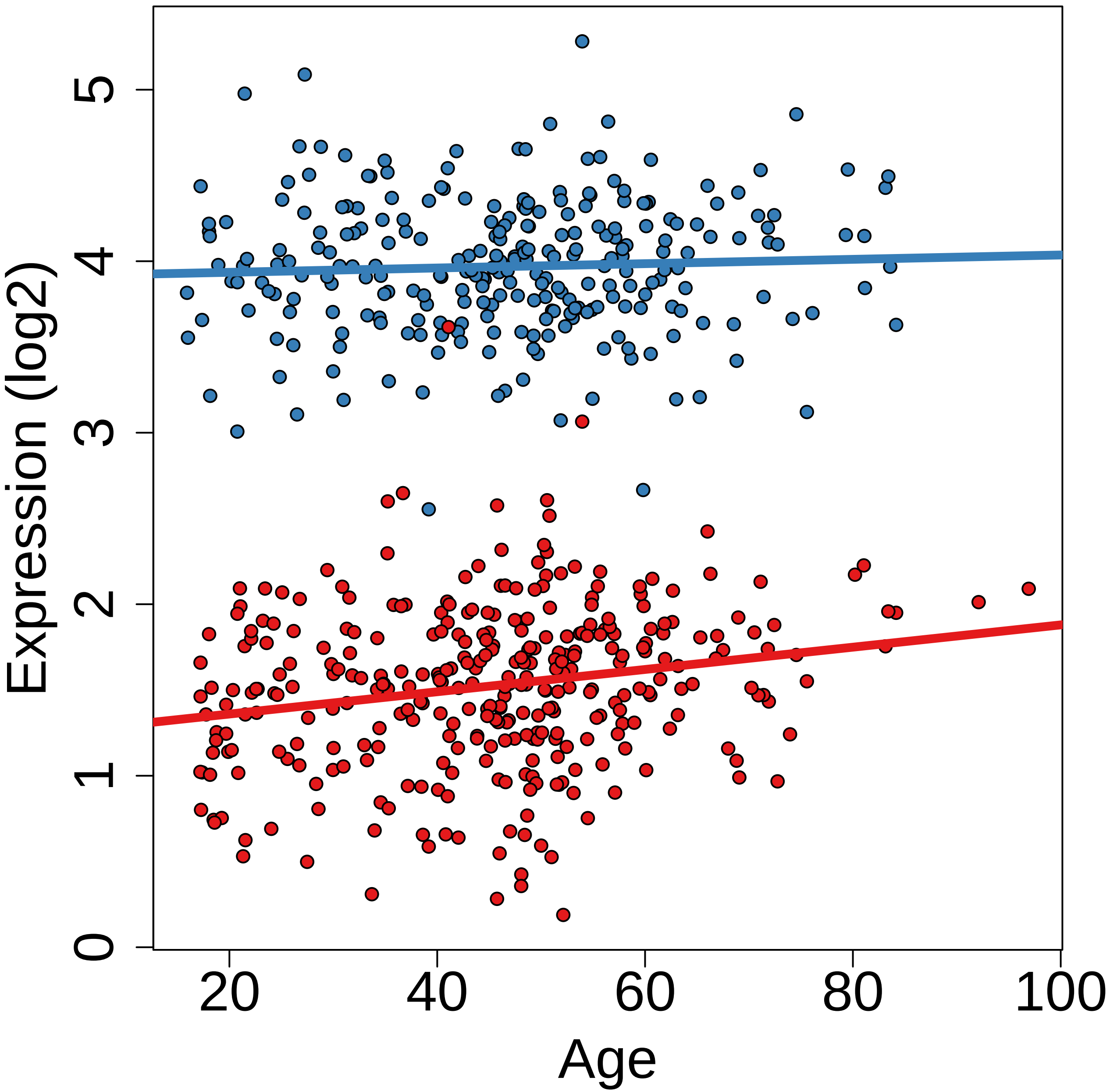
Expression (log2)
0
1
2
3
4
5
20
40
60
80
100
Age

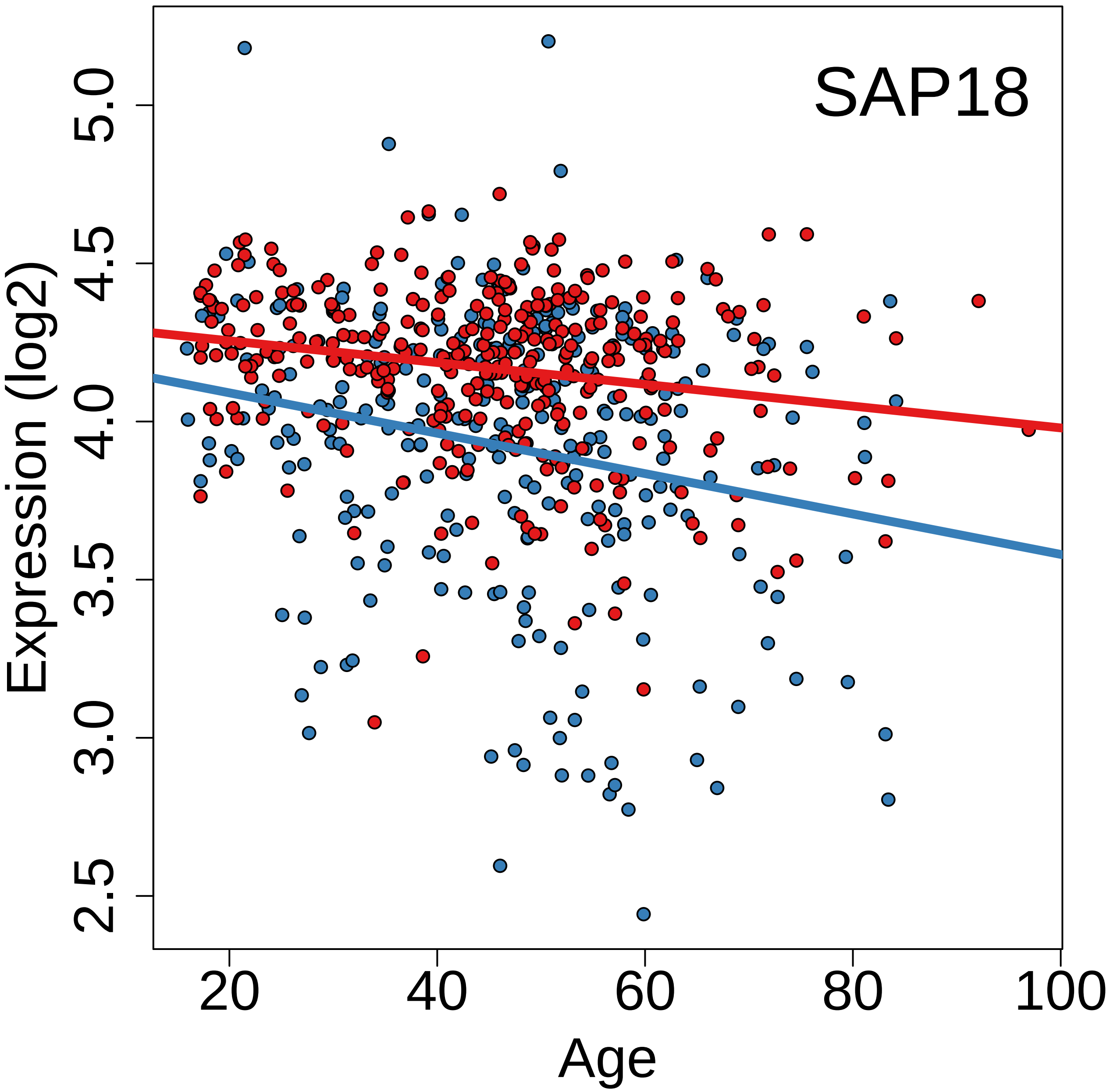
SAP18
Expression (log2)
2.5
3.0
3.5
4.0
4.5
5.0
Age
20
40
60
80
100

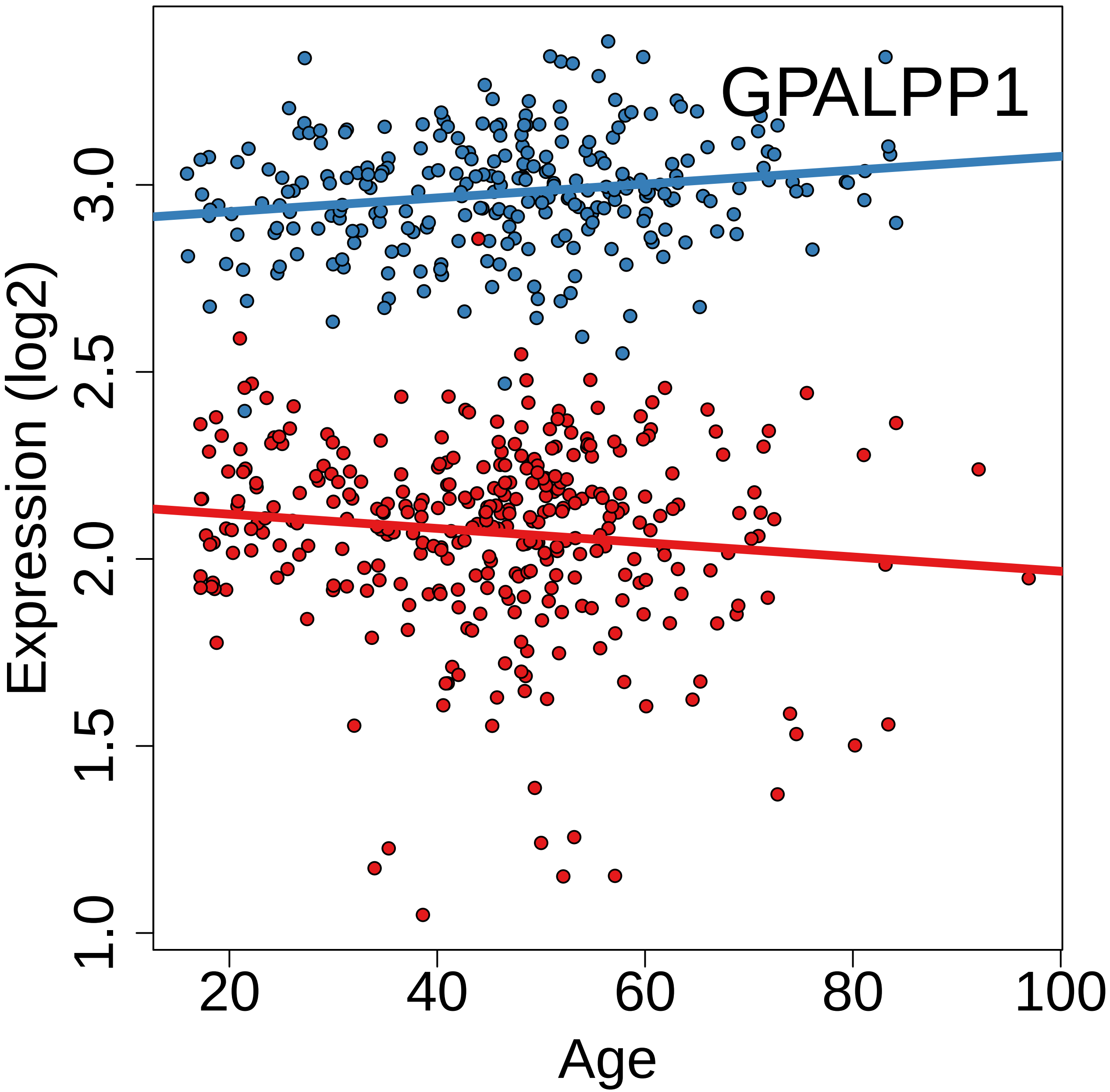

GPALPP1
Expression (log2)
Age
1.0
1.5
2.0
2.5
3.0
20
40
60
80
100

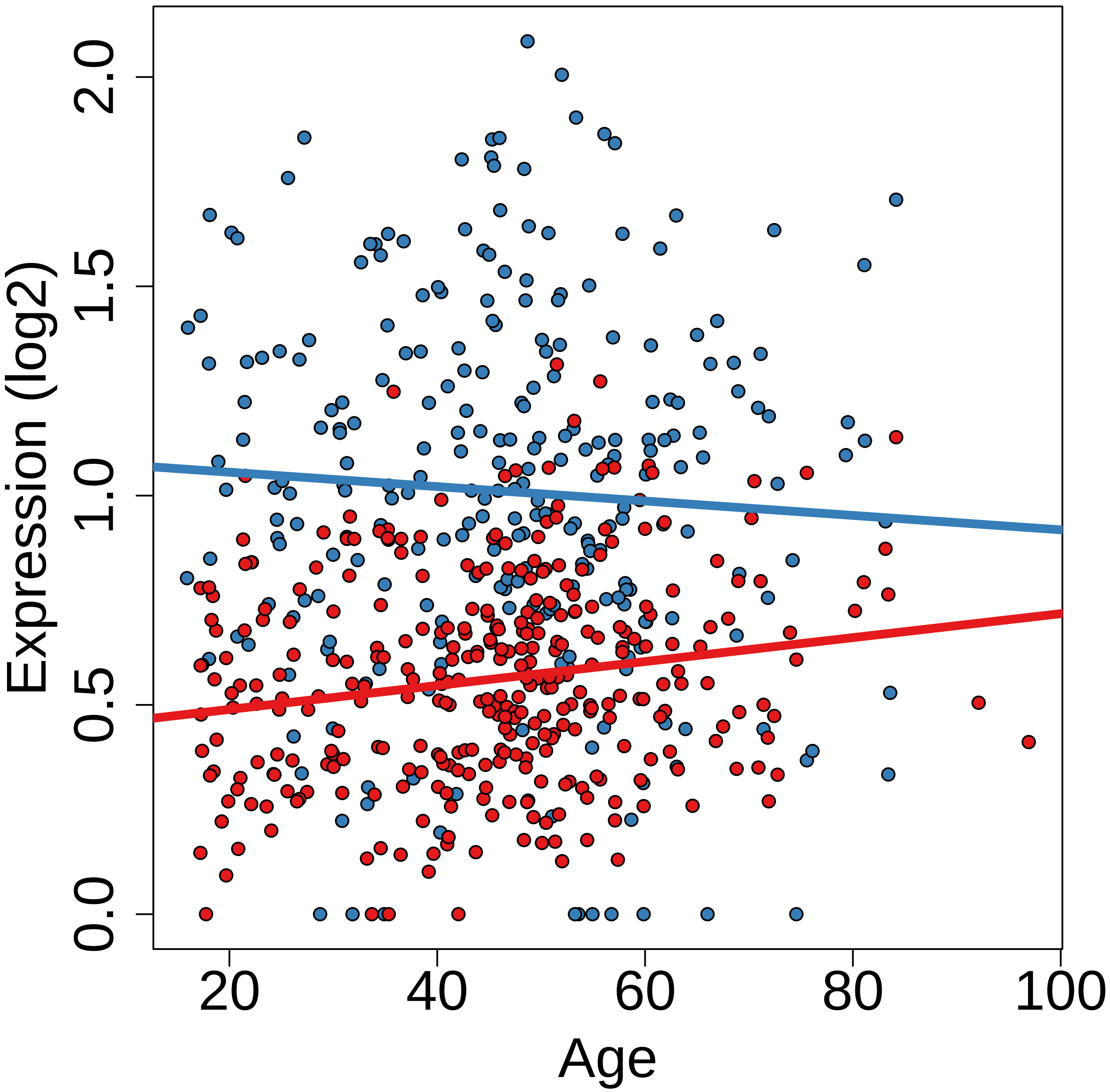

Expression (log2)
0.0
0.5
1.0
1.5
2.0
20
40
60
80
100
Age

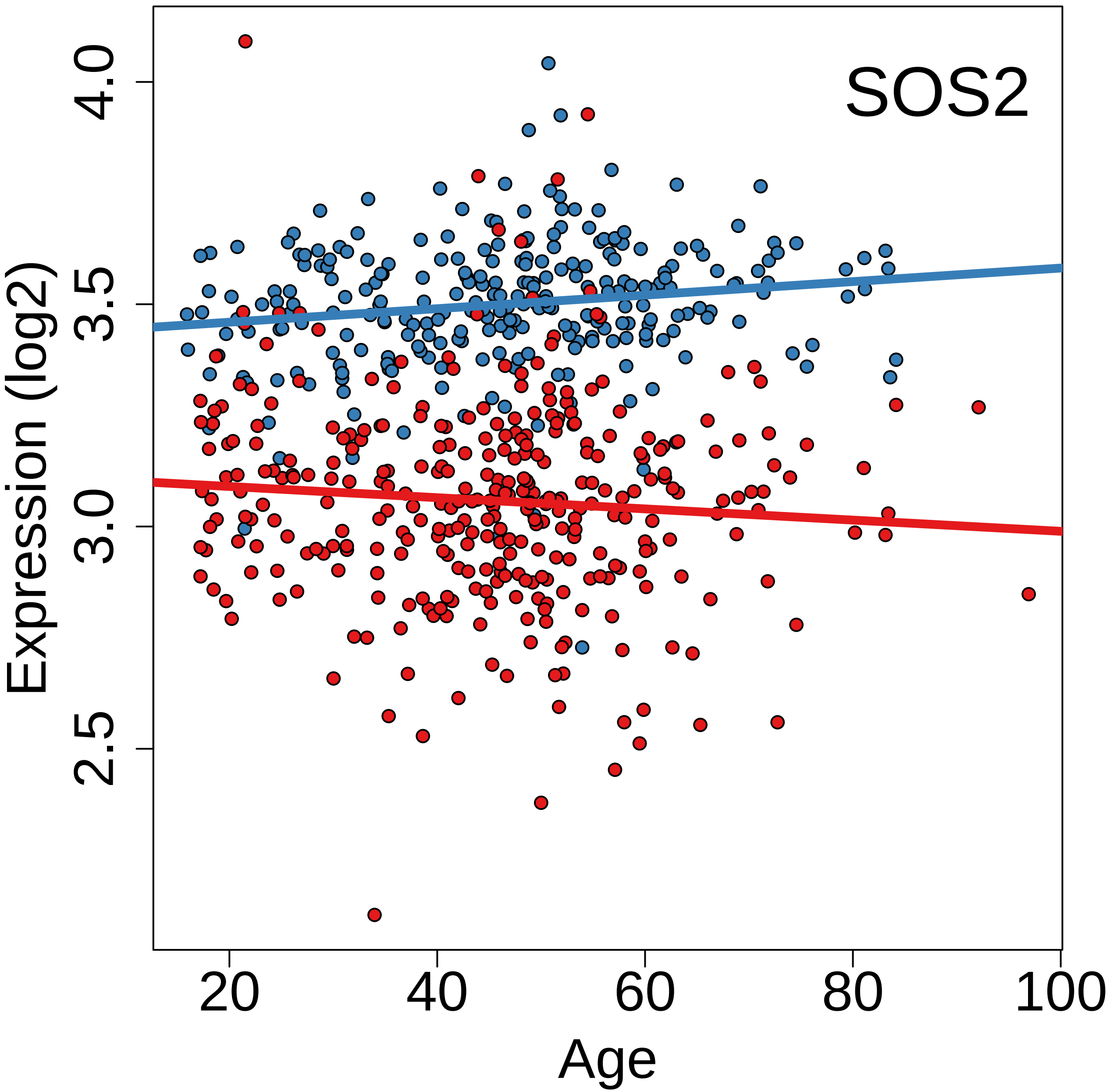

SOS2
Expression (log2)
4.0
3.5
3.0
2.5
20
40
60
80
100
Age

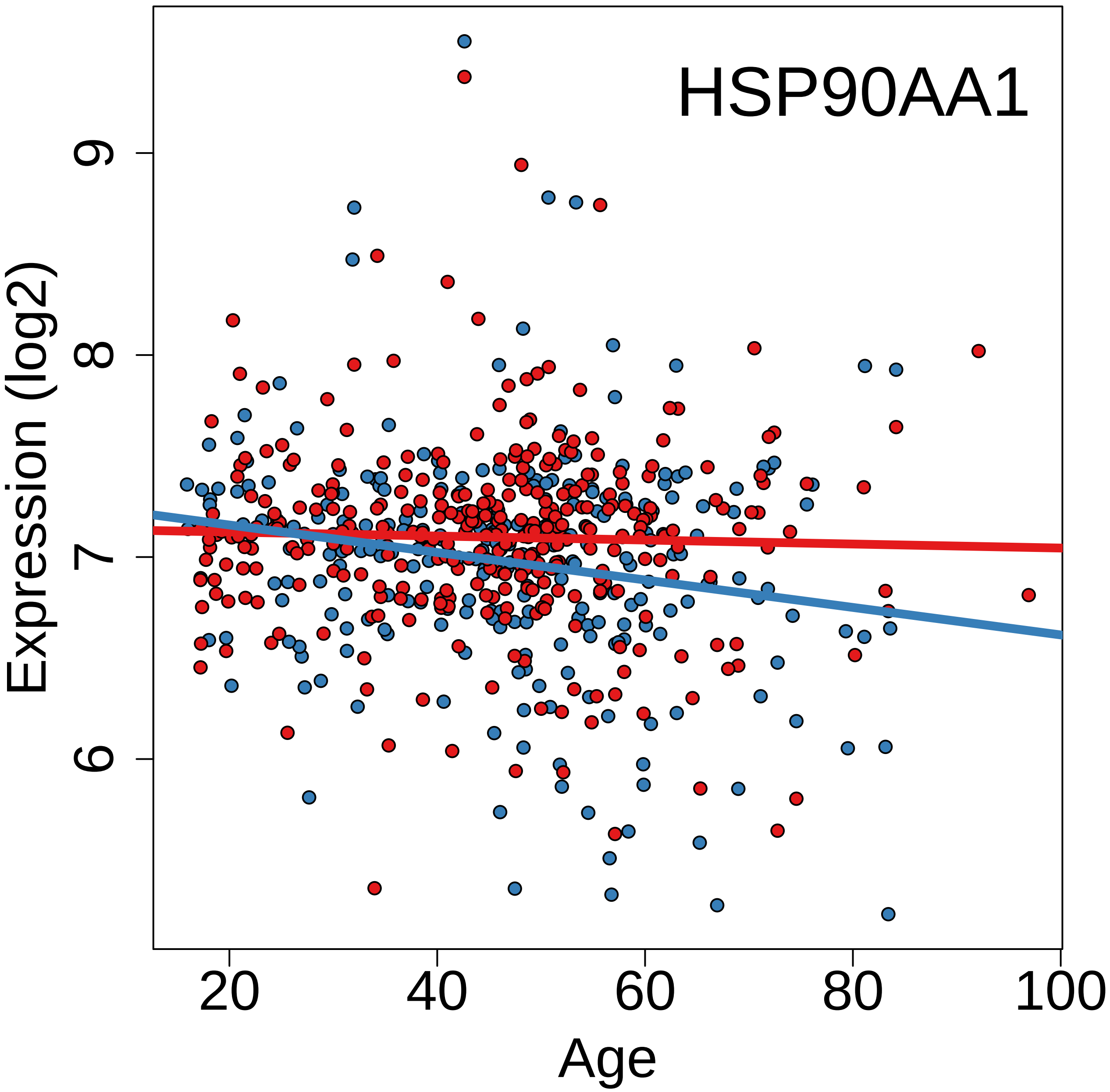
HSP90AA1
Expression (log2)
6
7
8
9
20
40
60
80
100
Age

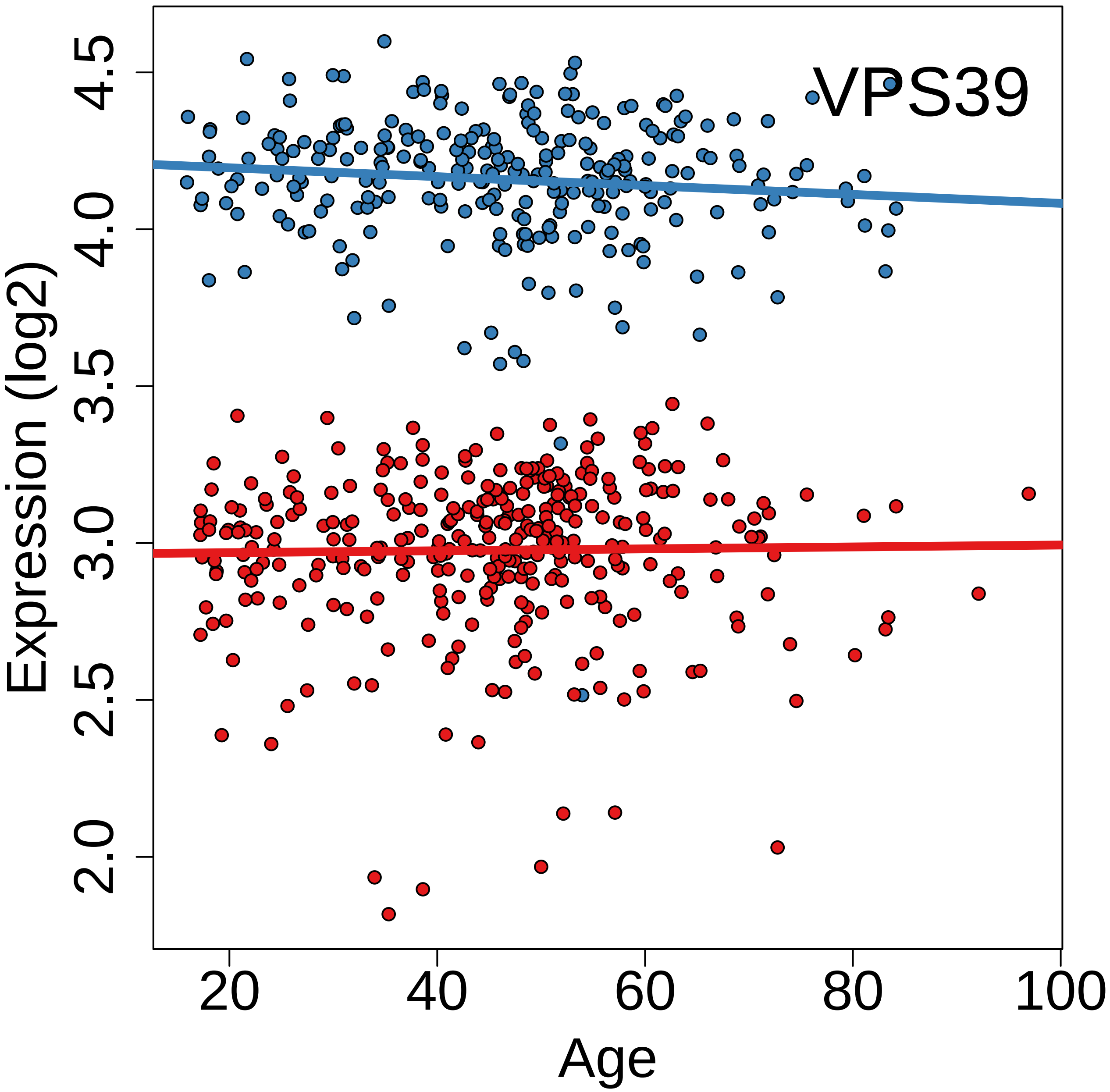
VPS39
Expression (log2)
2.0
2.5
3.0
3.5
4.0
4.5
20
40
60
80
100
Age

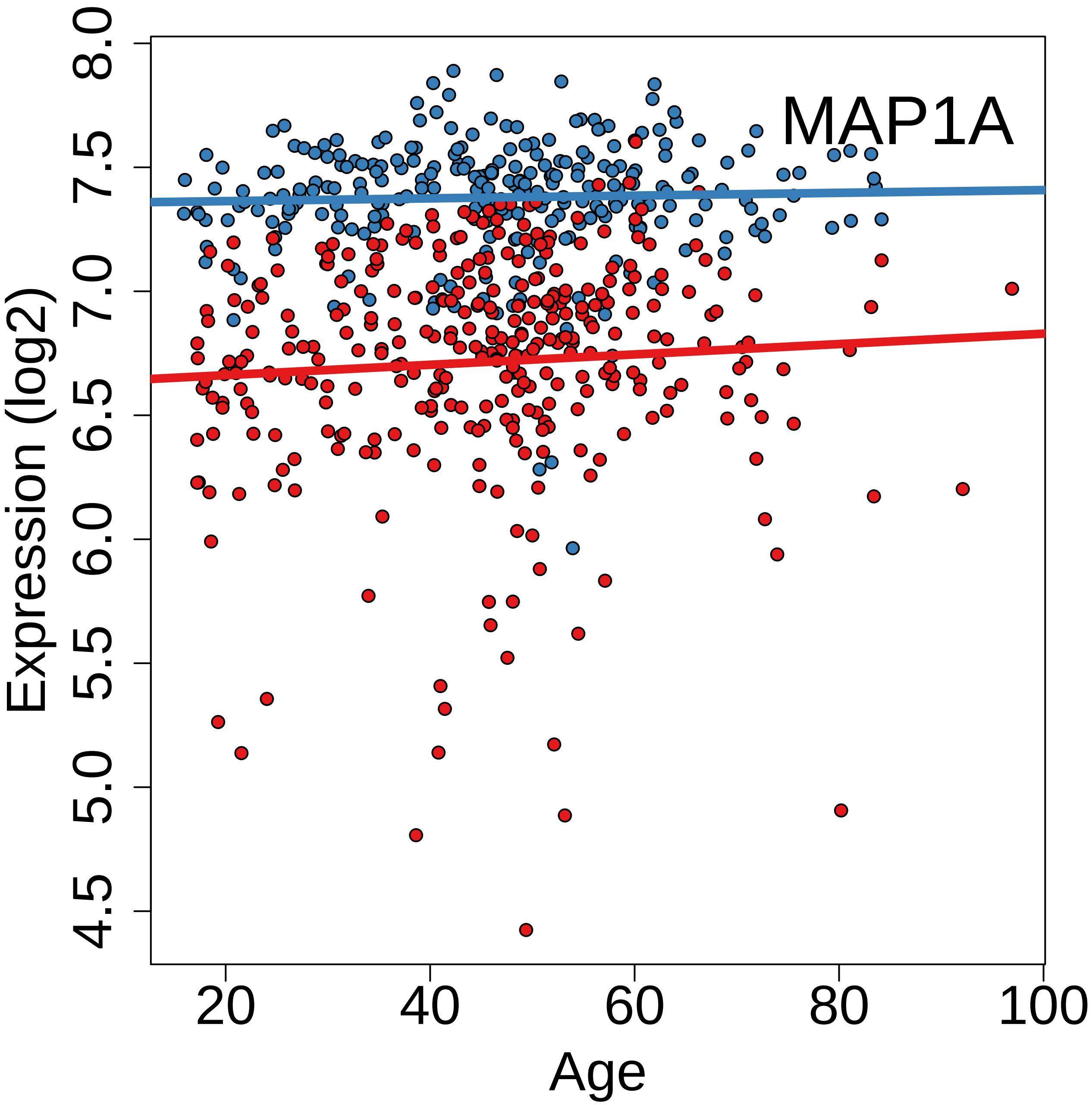
MAP1A
Expression (log2)
Age
8.0
7.5
7.0
6.5
6.0
5.5
5.0
4.5
20
40
60
80
100

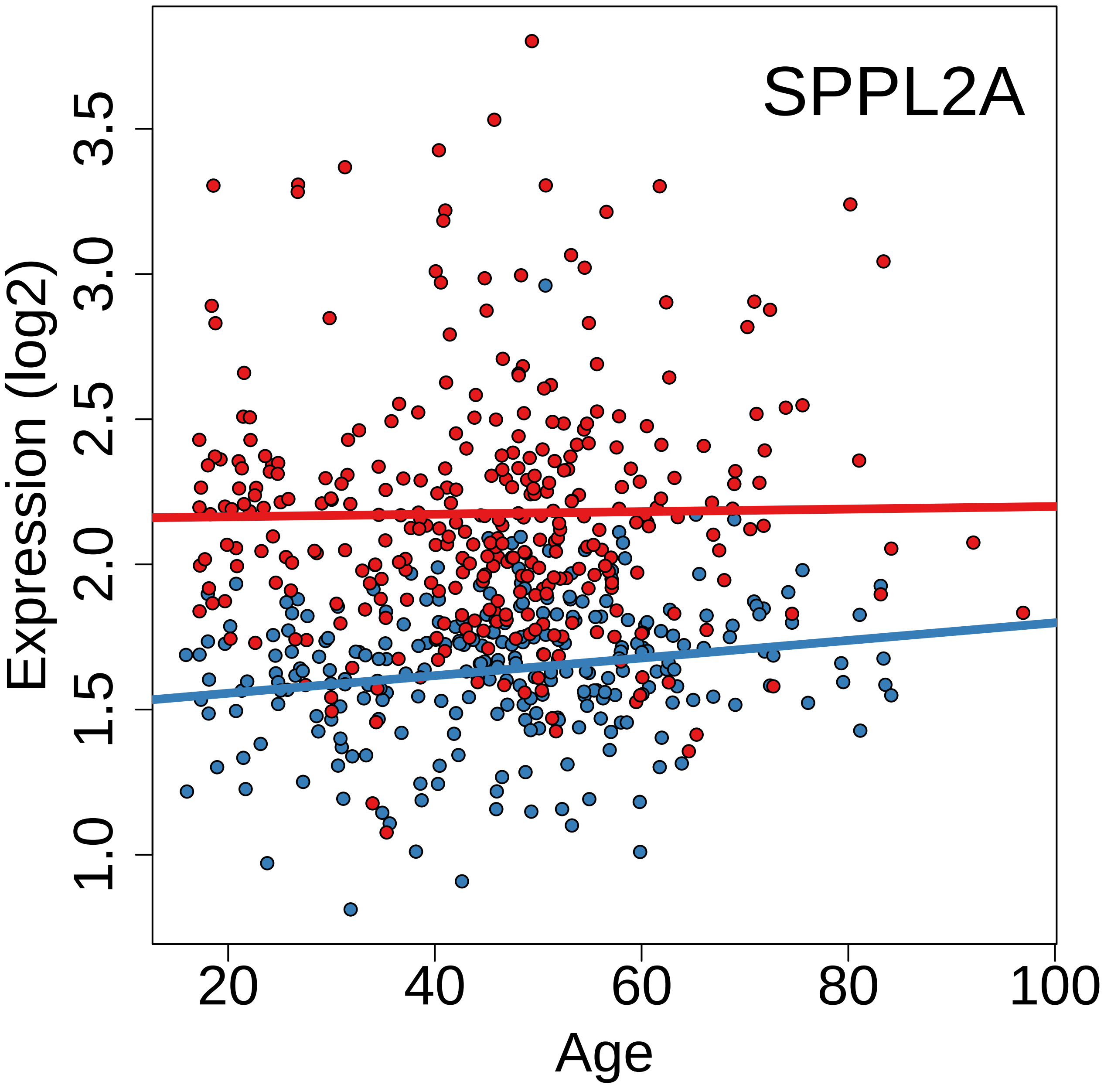

SPPL2A
Expression (log2)
1.0
1.5
2.0
2.5
3.0
3.5
20
40
60
80
100
Age

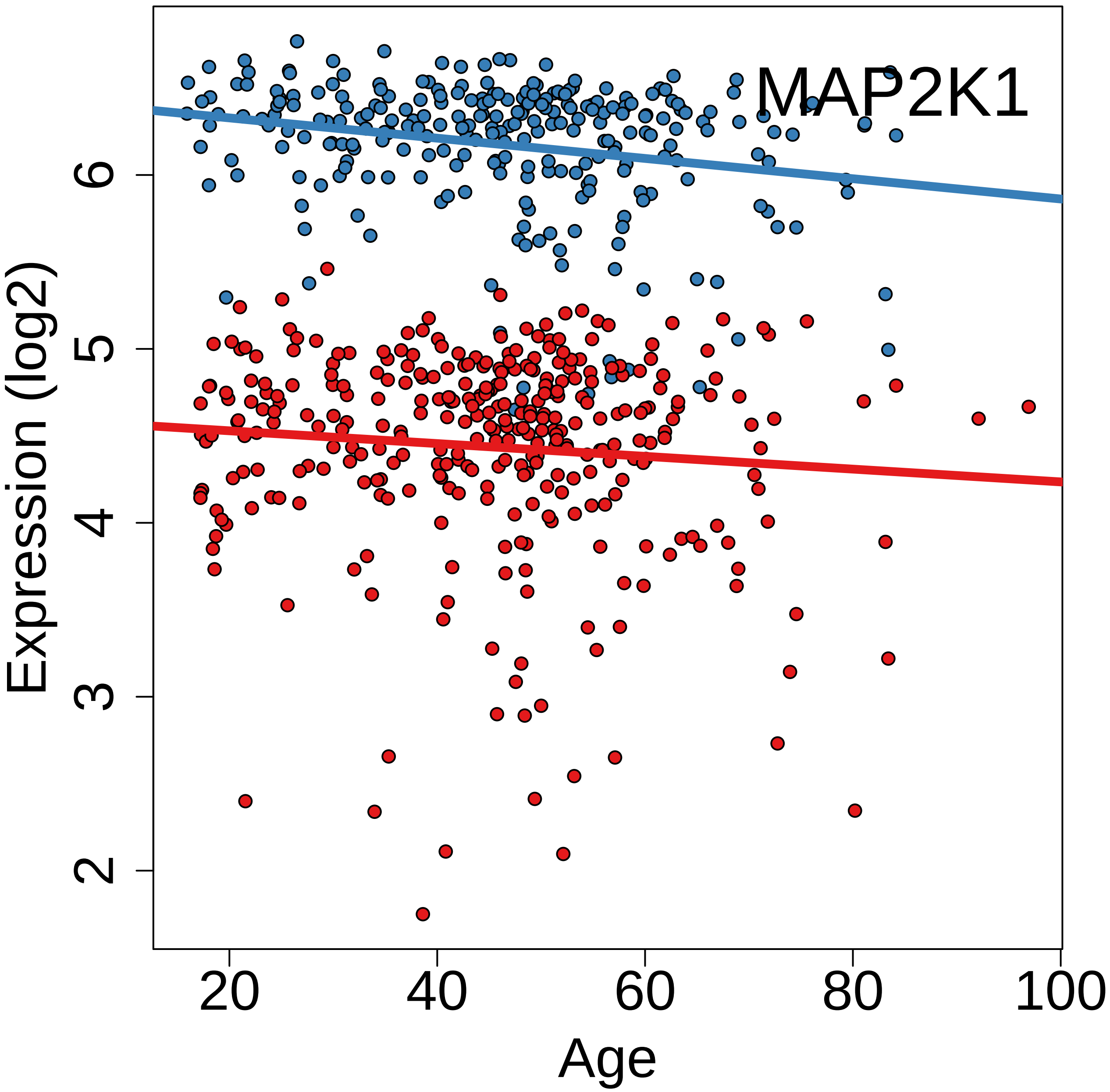
MAP2K1
Expression (log2)
Age
2
3
4
5
6
20
40
60
80
100

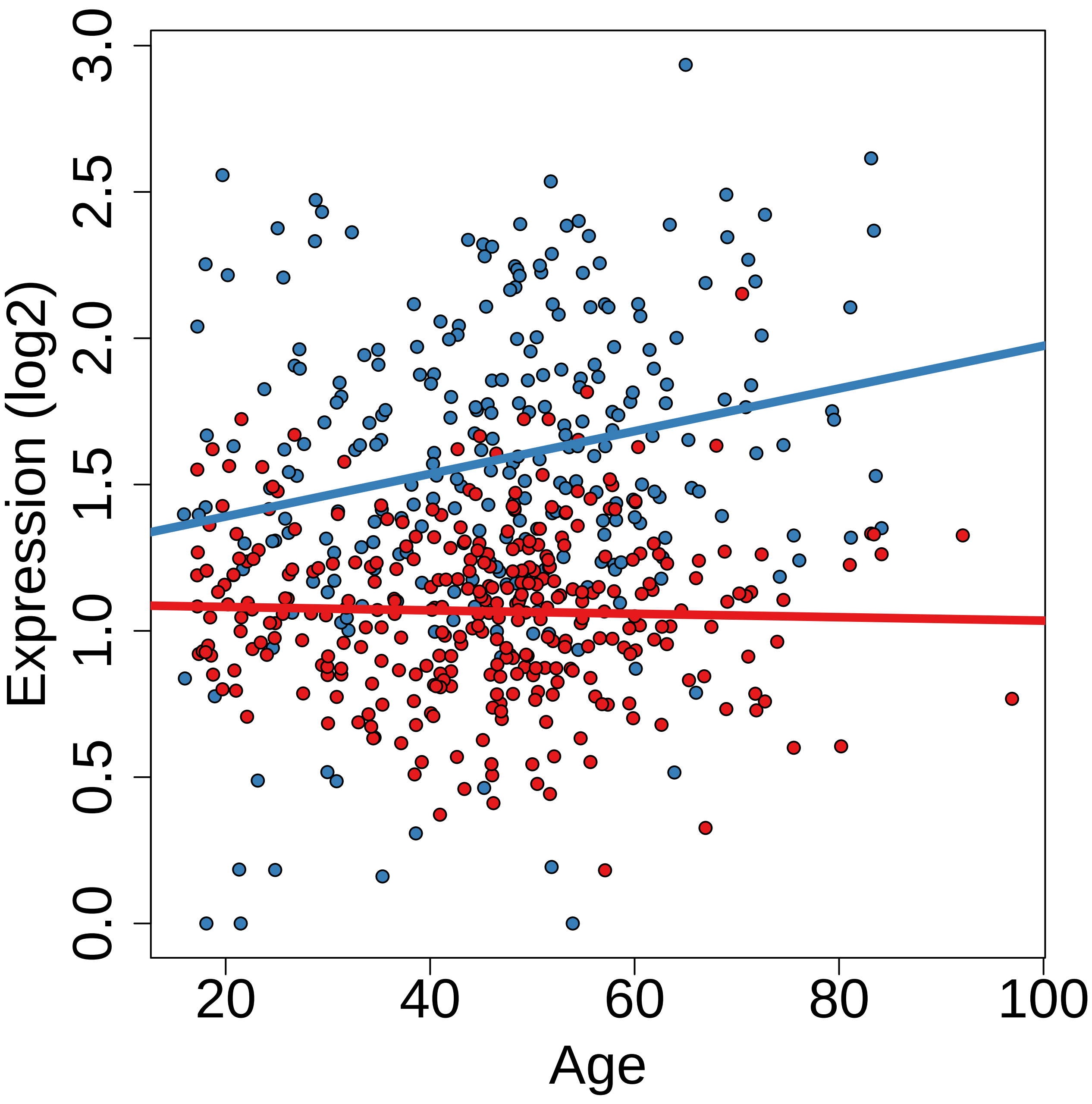

Expression (log2)
0.0
0.5
1.0
1.5
2.0
2.5
3.0
20
40
60
80
100
Age

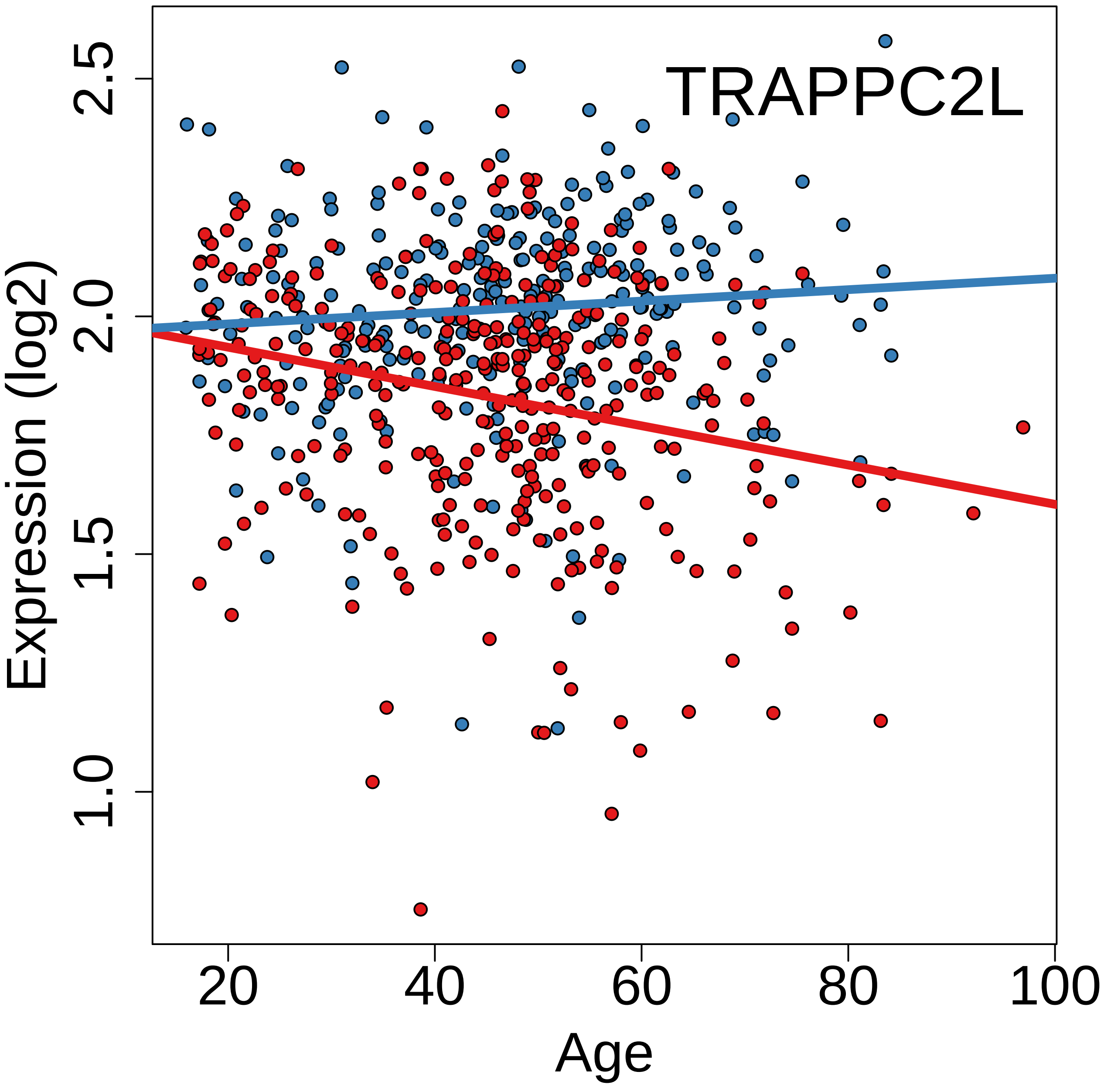

TRAPPC2L
Expression (log2)
2.5
2.0
1.5
1.0
20
40
60
80
100
Age

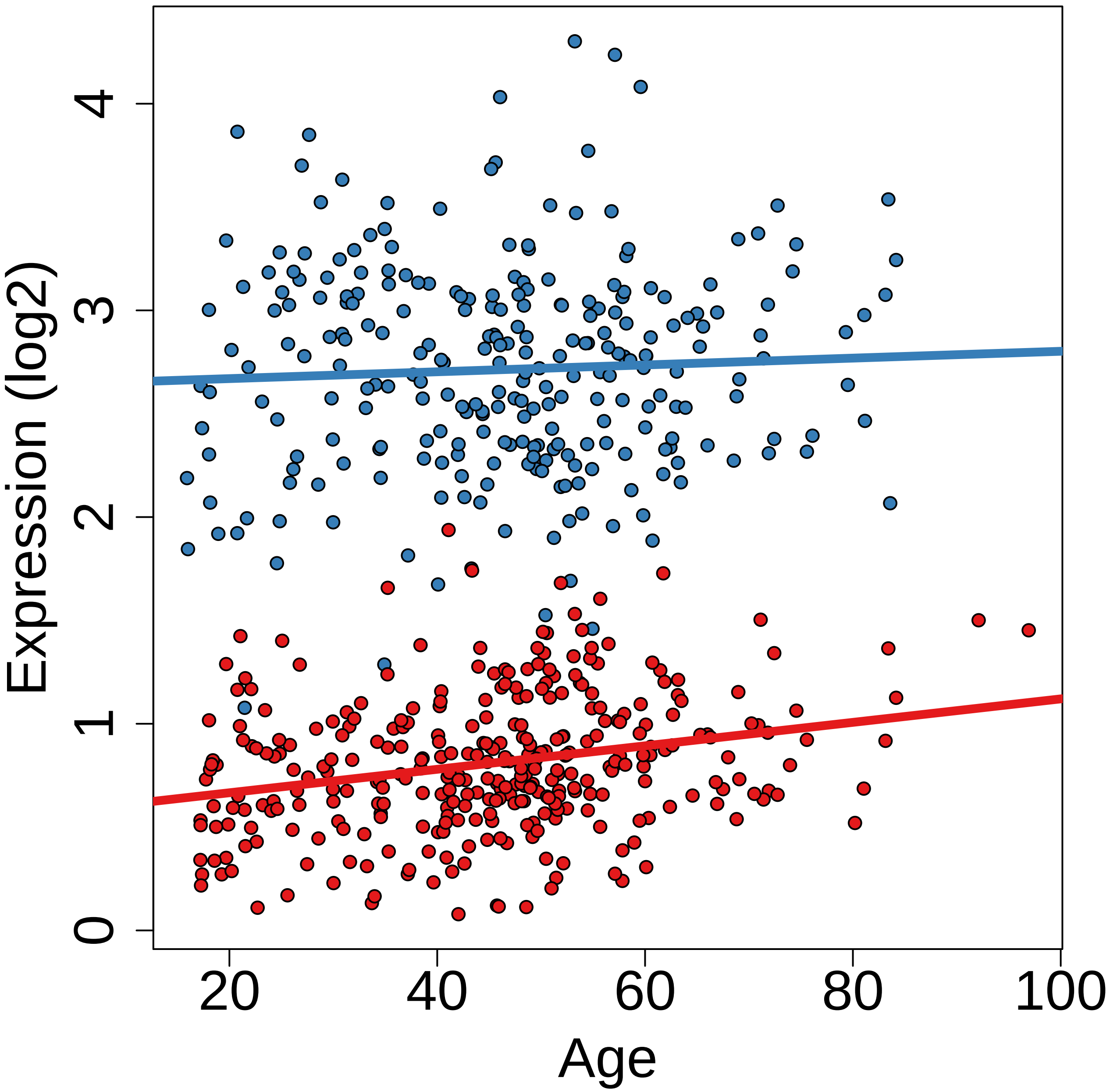
Expression (log2)
0
1
2
3
4
20
40
60
80
100
Age

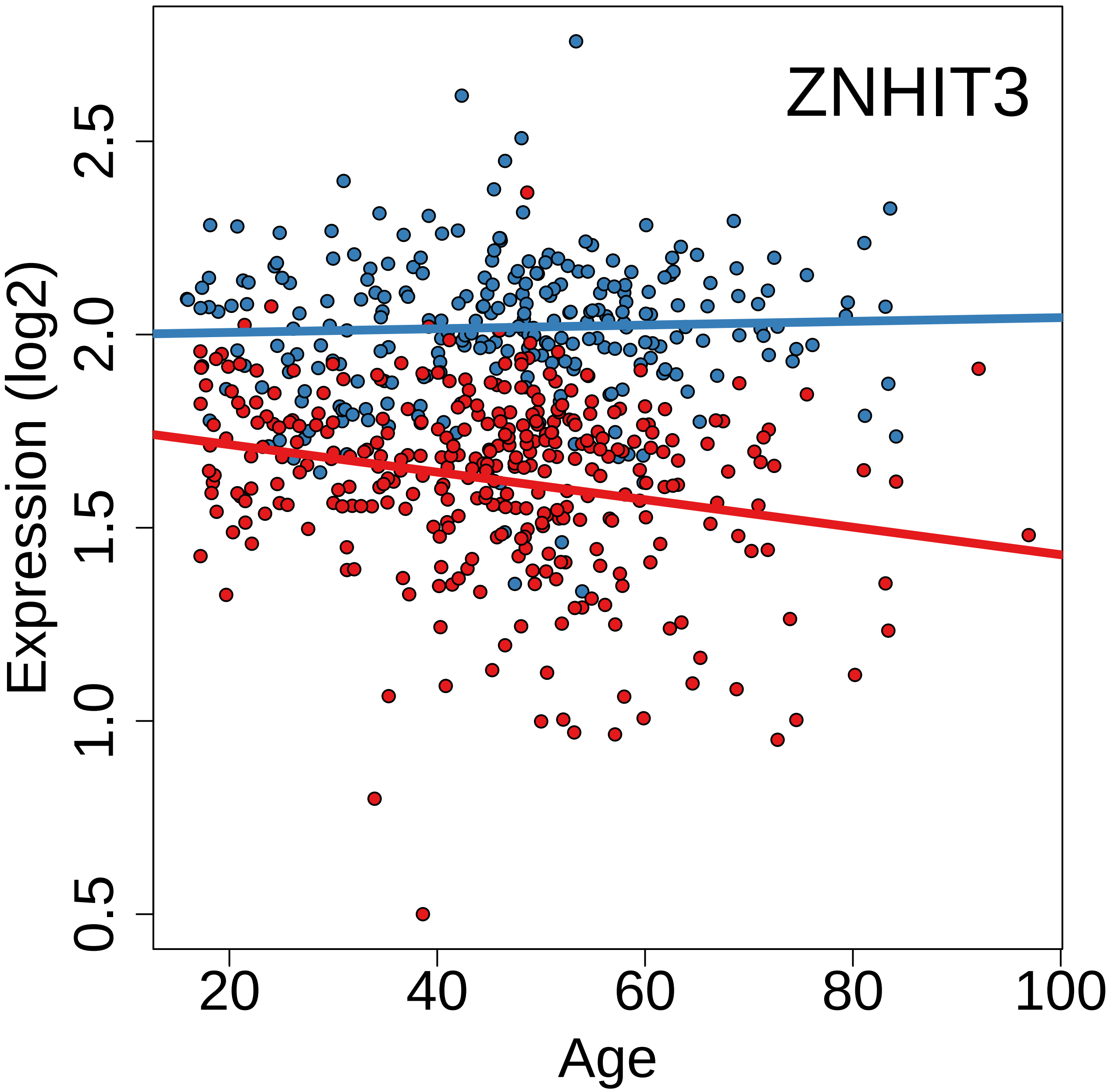

ZNHIT3
Expression (log2)
0.5
1.0
1.5
2.0
2.5
20
40
60
80
100
Age

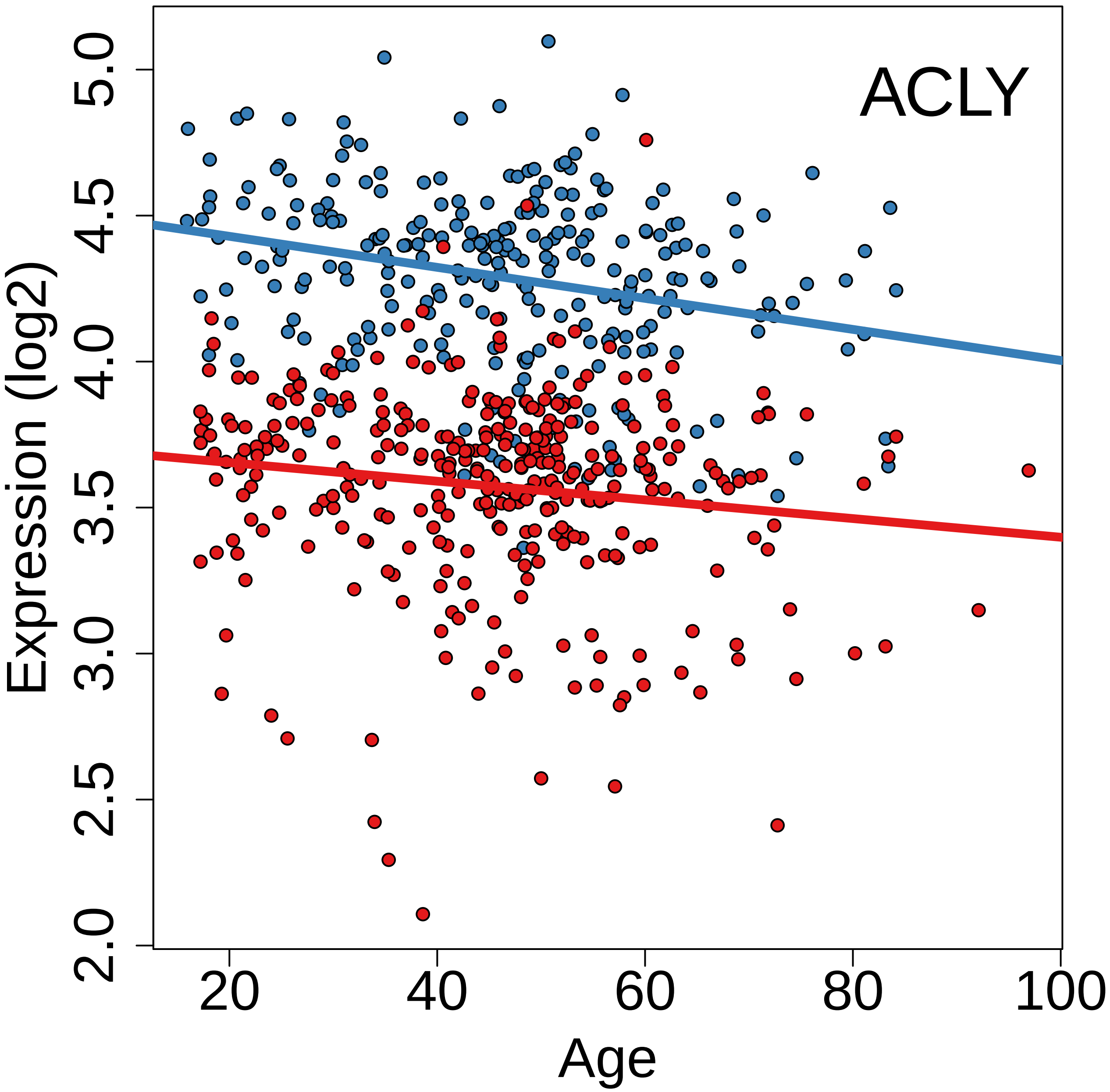
ACLY
Expression (log2)
5.0
4.5
4.0
3.5
3.0
2.5
2.0
20
40
60
80
100
Age

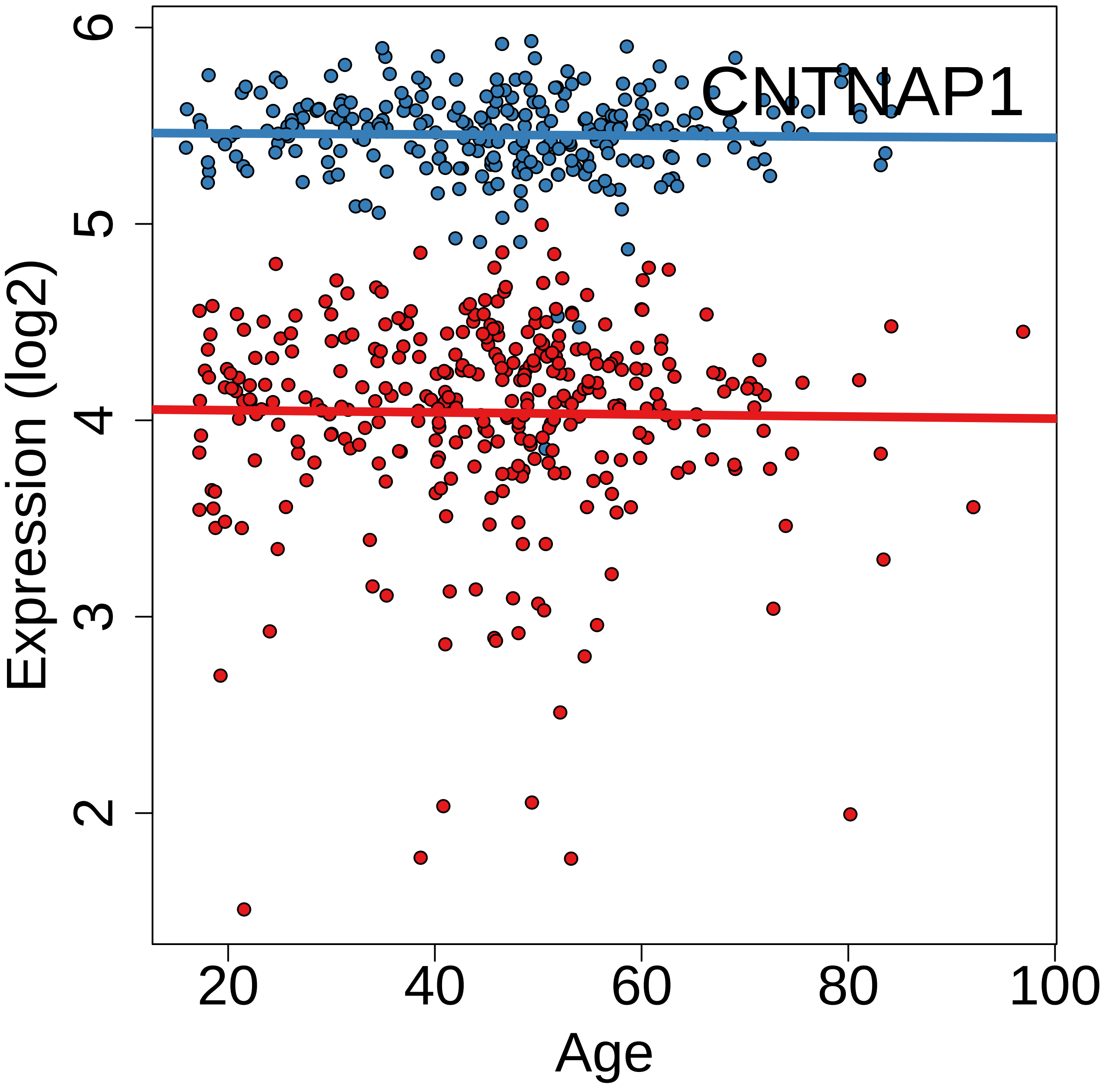
CNTNAP1
Expression (log2)
Age
2
3
4
5
6
20
40
60
80
100

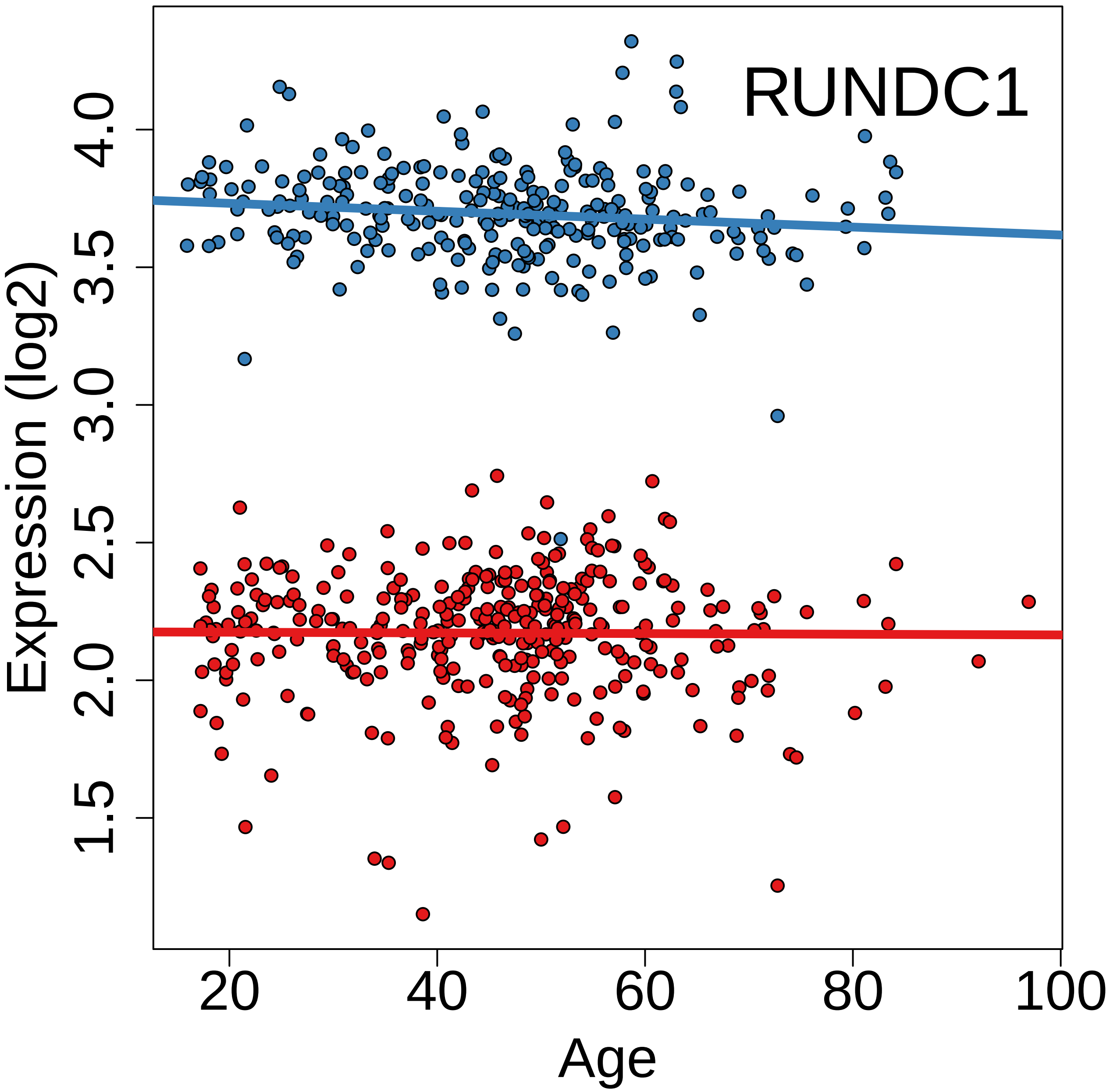

RUNDC1
Expression (log2)
1.5
2.0
2.5
3.0
3.5
4.0
20
40
60
80
100
Age

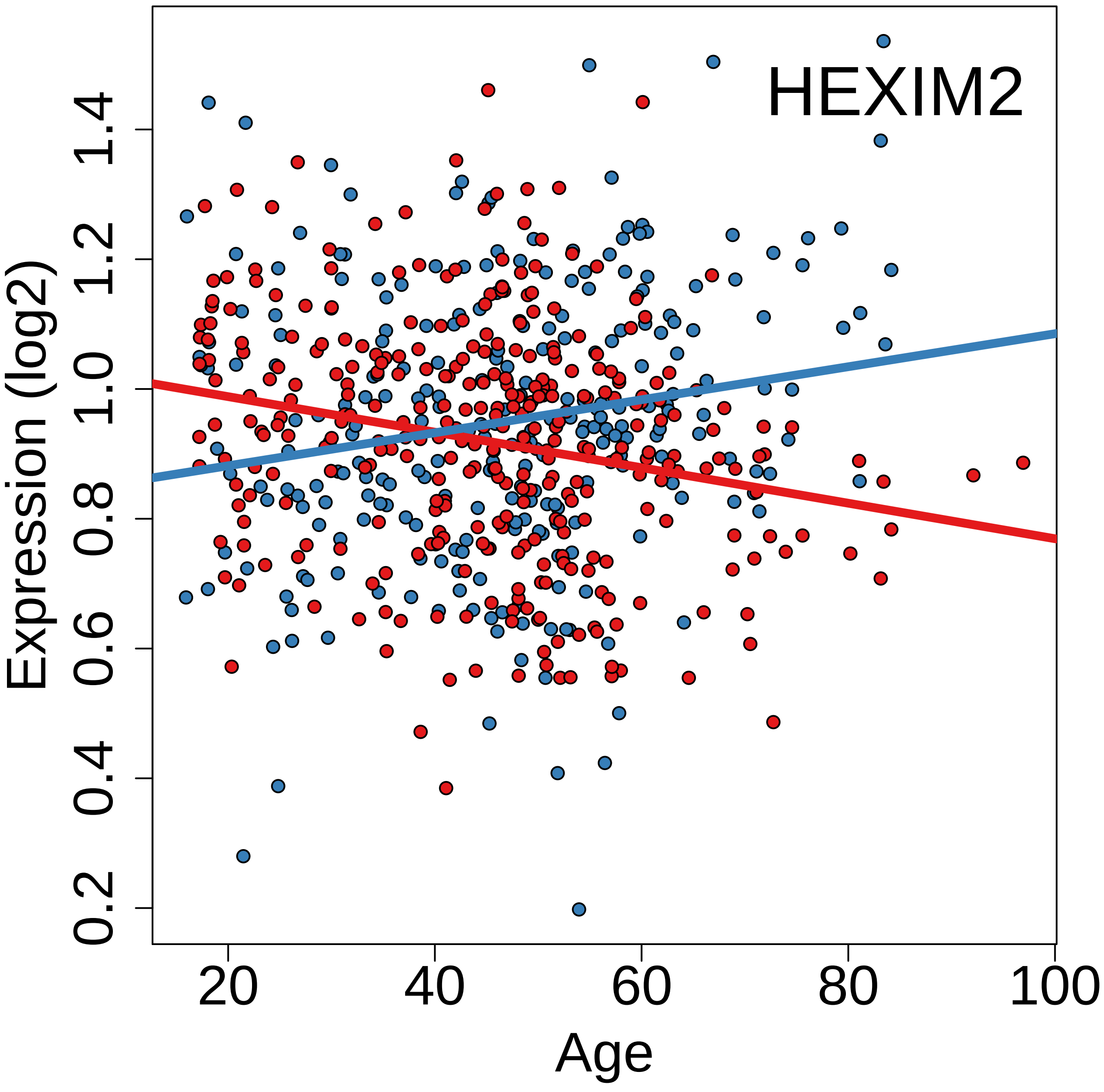
HEXIM2
Expression (log2)
0.2
0.4
0.6
0.8
1.0
1.2
1.4
Age
20
40
60
80
100

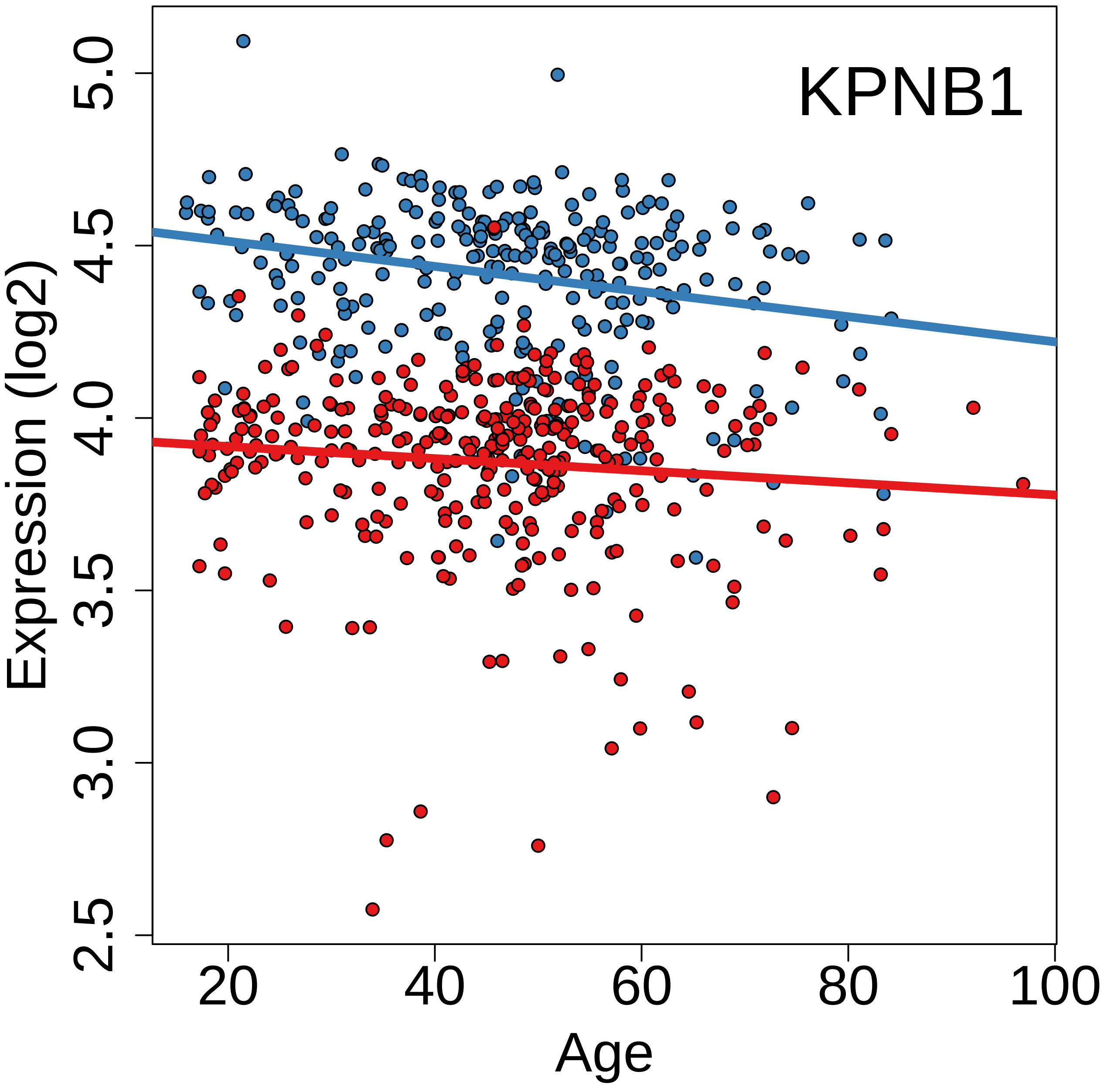
KPNB1
Expression (log2)
5.0
4.5
4.0
3.5
3.0
2.5
20
40
60
80
100
Age

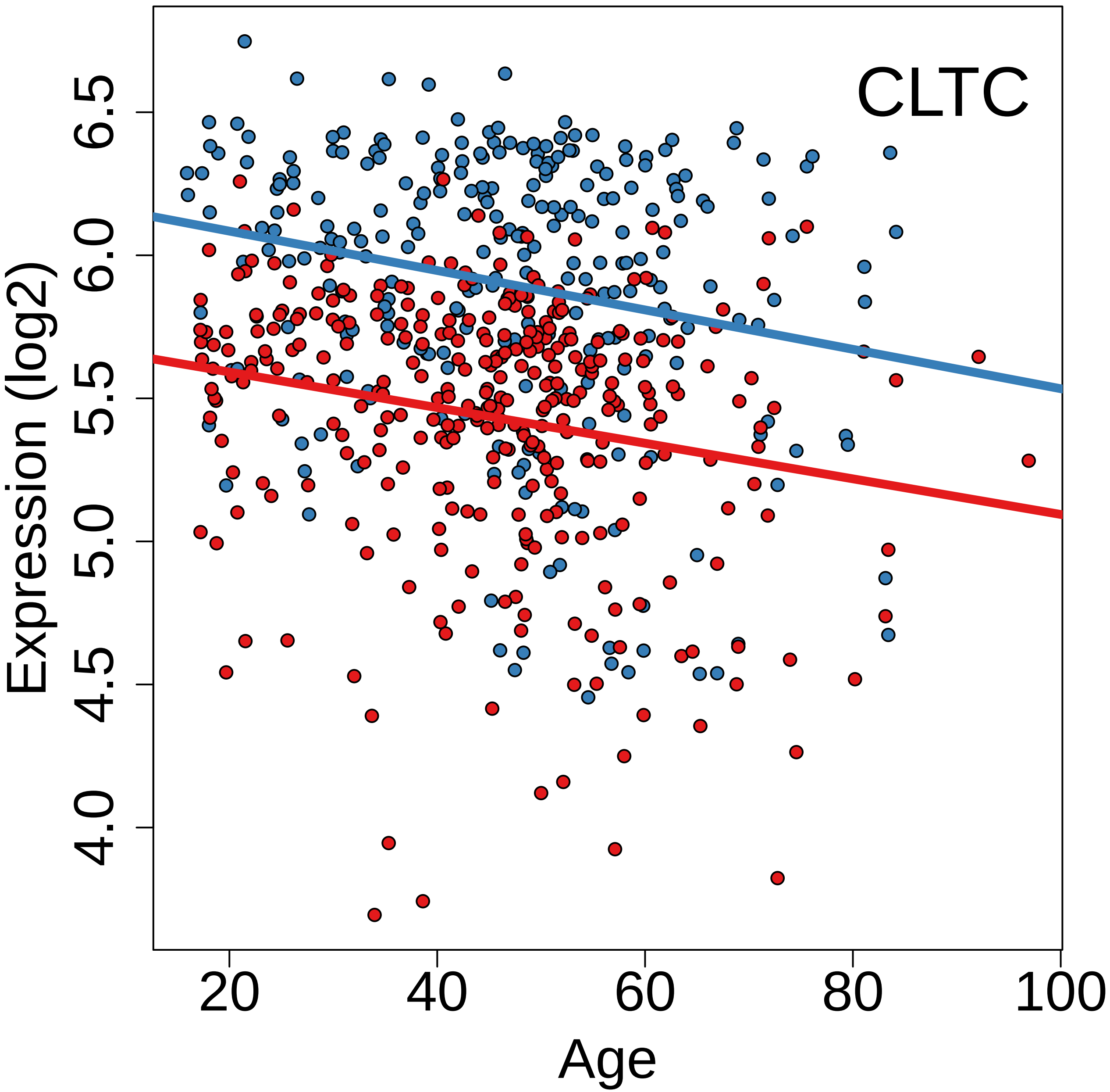
CLTC
Expression (log2)
4.0
4.5
5.0
5.5
6.0
6.5
20
40
60
80
100
Age

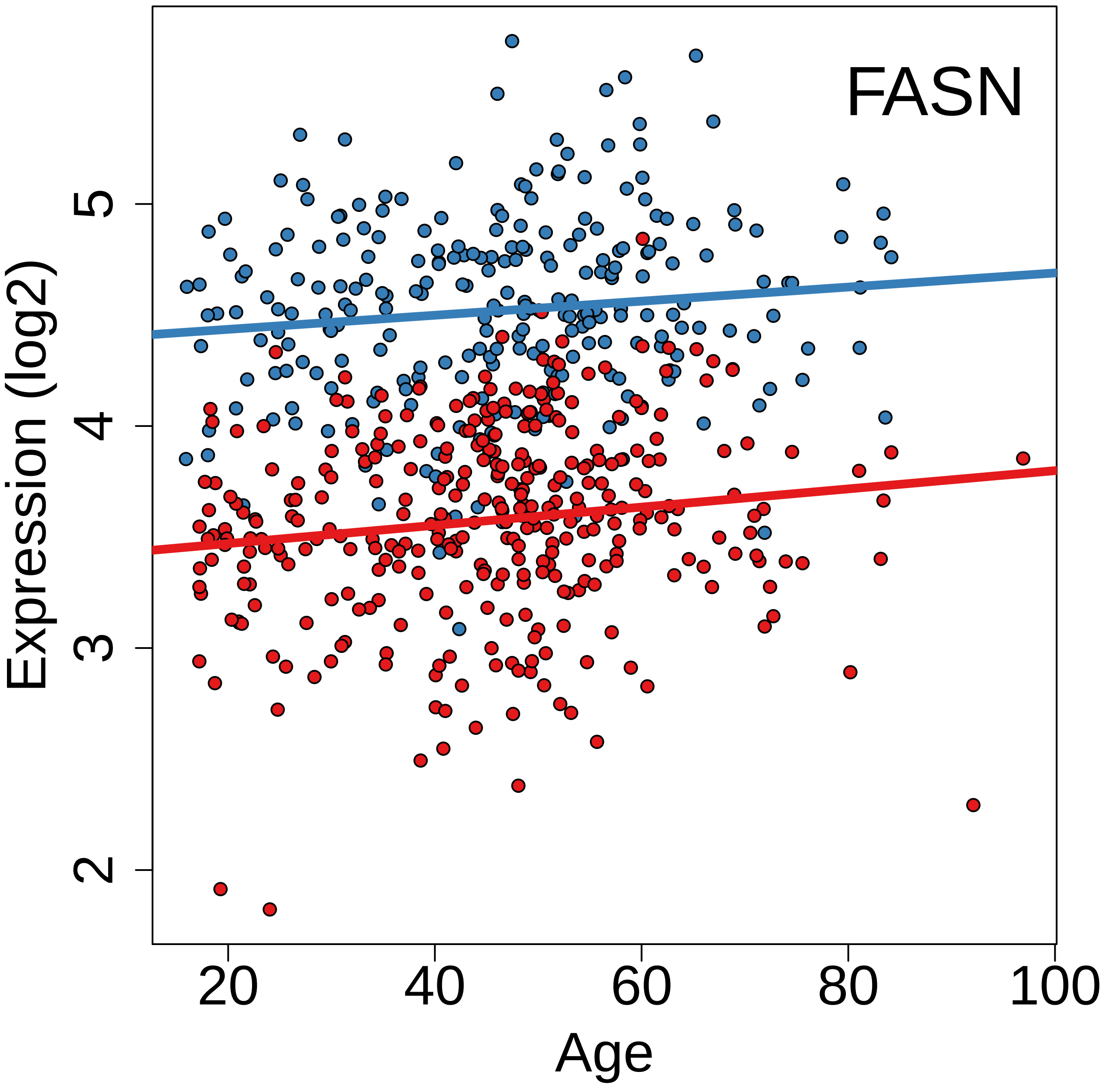
FASN
Expression (log2)
2
3
4
5
20
40
60
80
100
Age

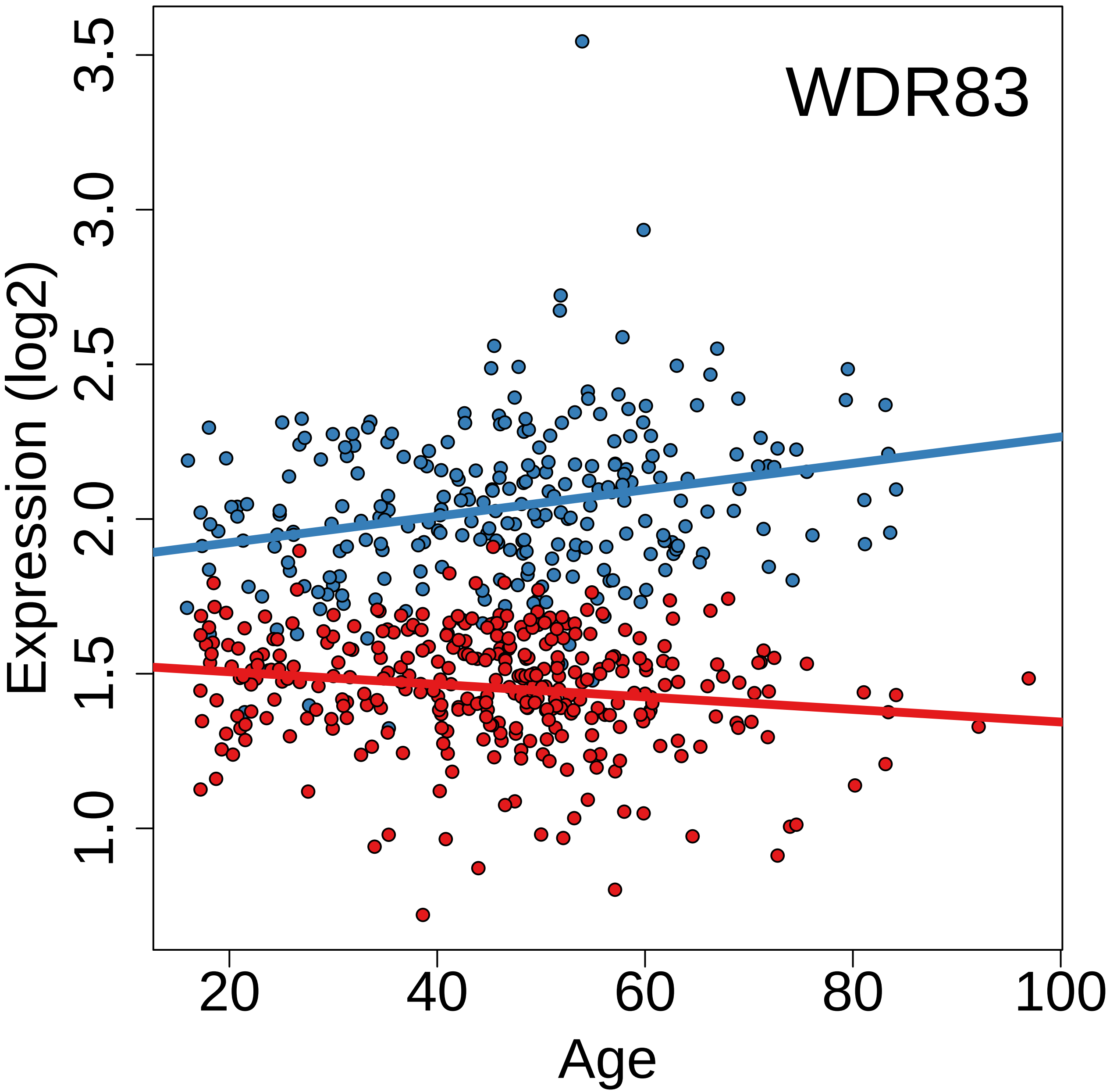
WDR83
Expression (log2)
1.0
1.5
2.0
2.5
3.0
3.5
20
40
60
80
100
Age

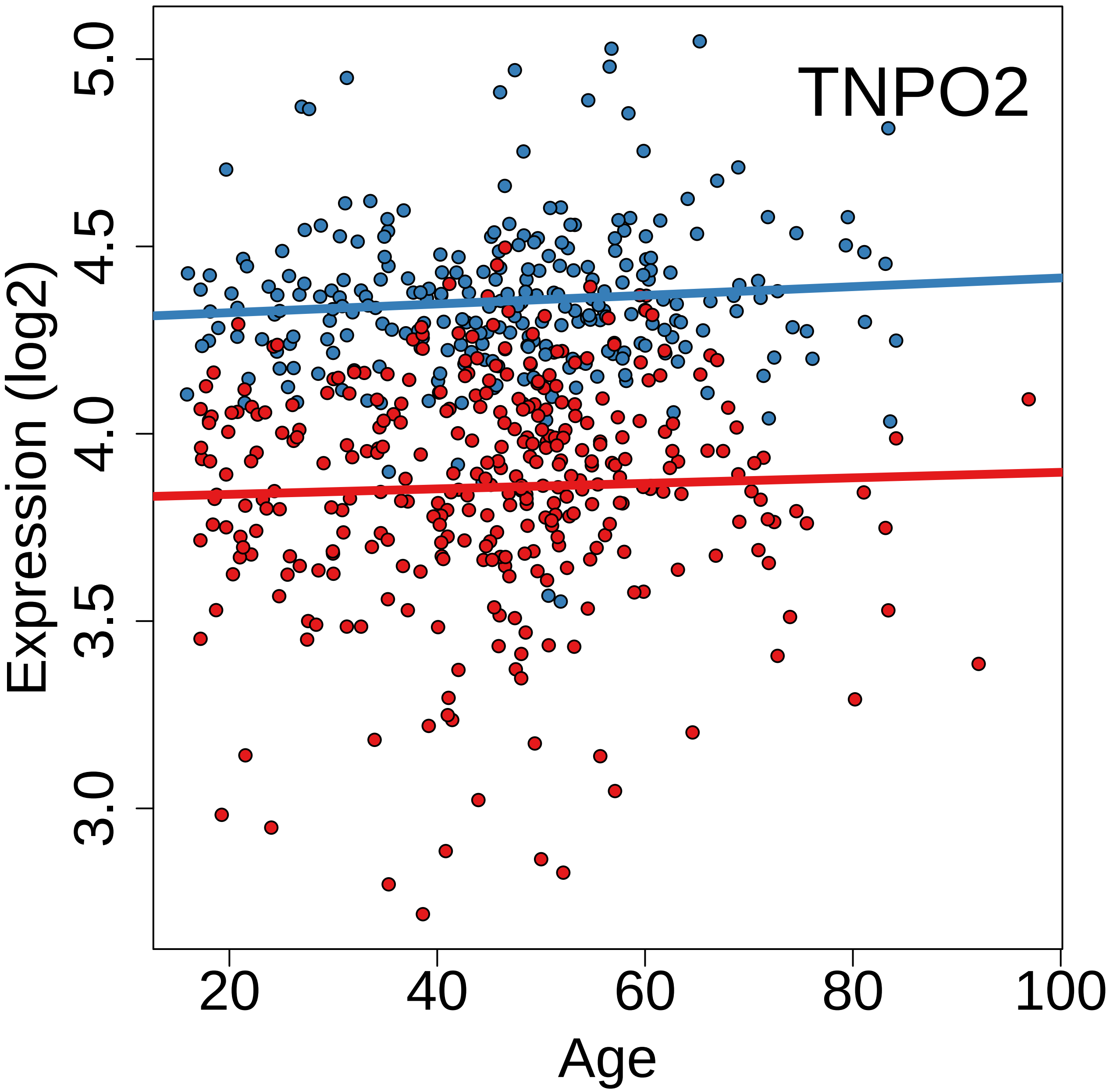
TNPO2
Expression (log2)
3.0
3.5
4.0
4.5
5.0
20
40
60
80
100
Age

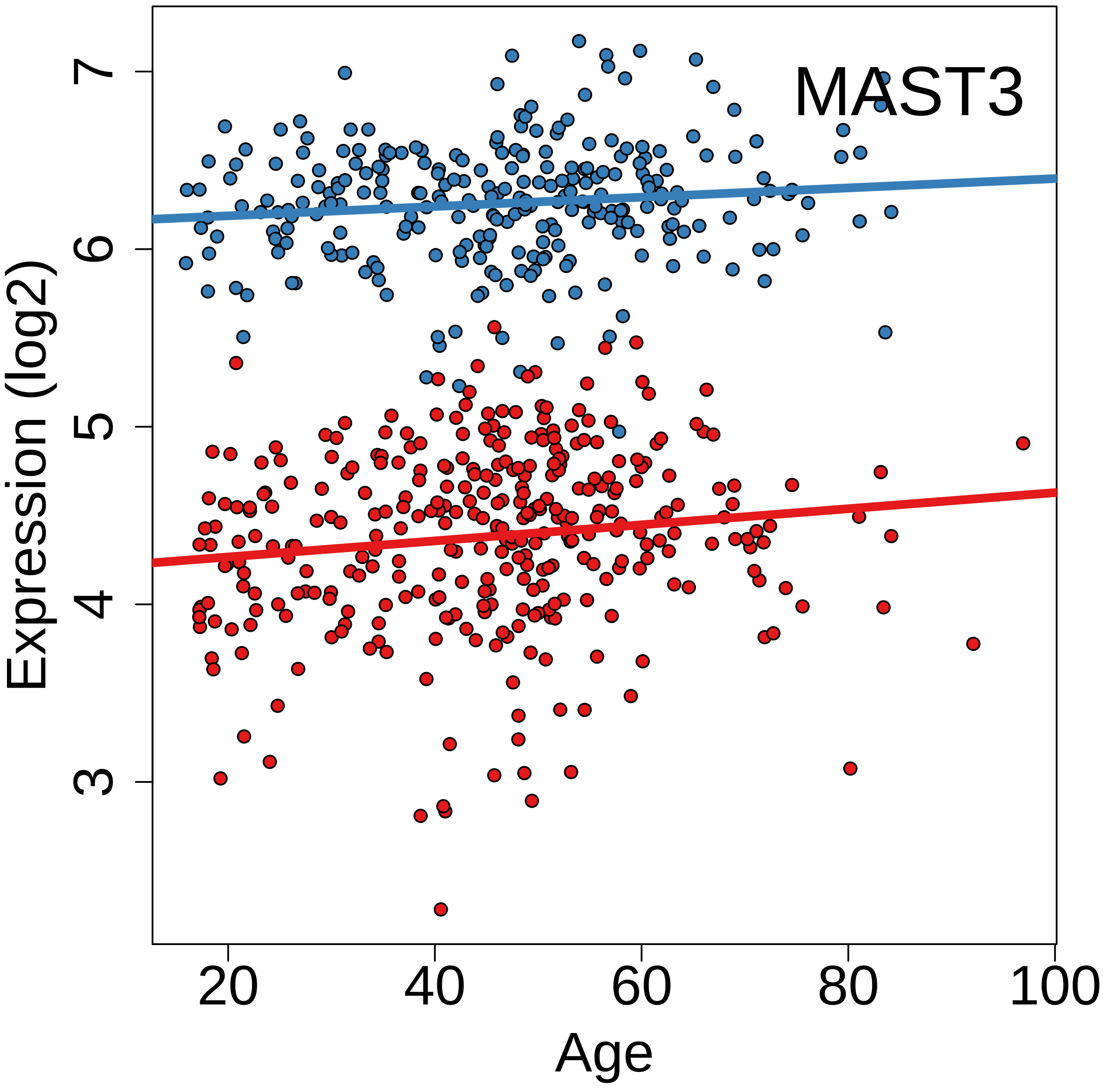
MAST3
Expression (log2)
Age
3
4
5
6
7
20
40
60
80
100

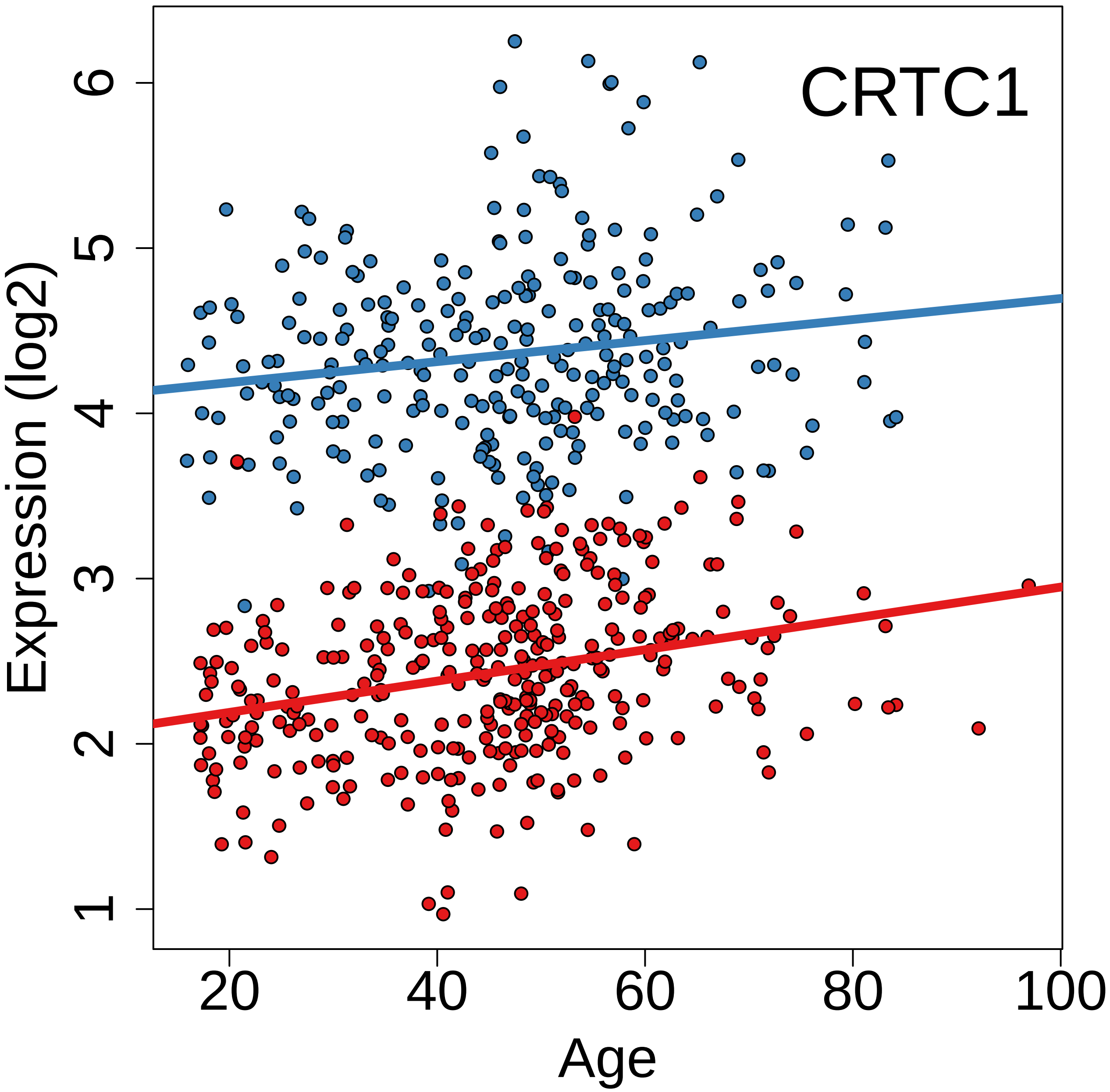
CRTC1
Expression (log2)
Age
1
2
3
4
5
6
20
40
60
80
100

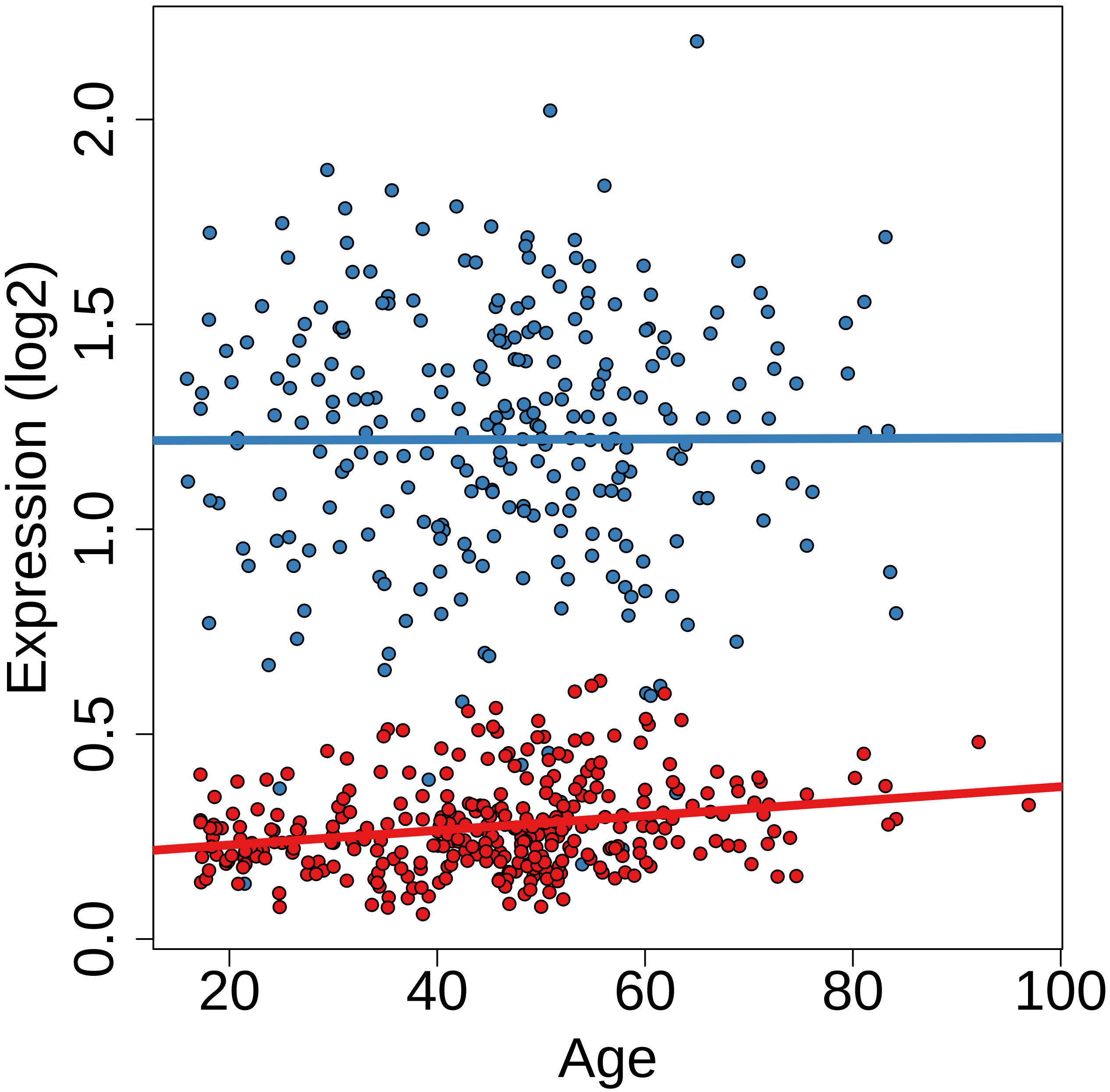

Expression (log2)
0.0
0.5
1.0
1.5
2.0
20
40
60
80
100
Age

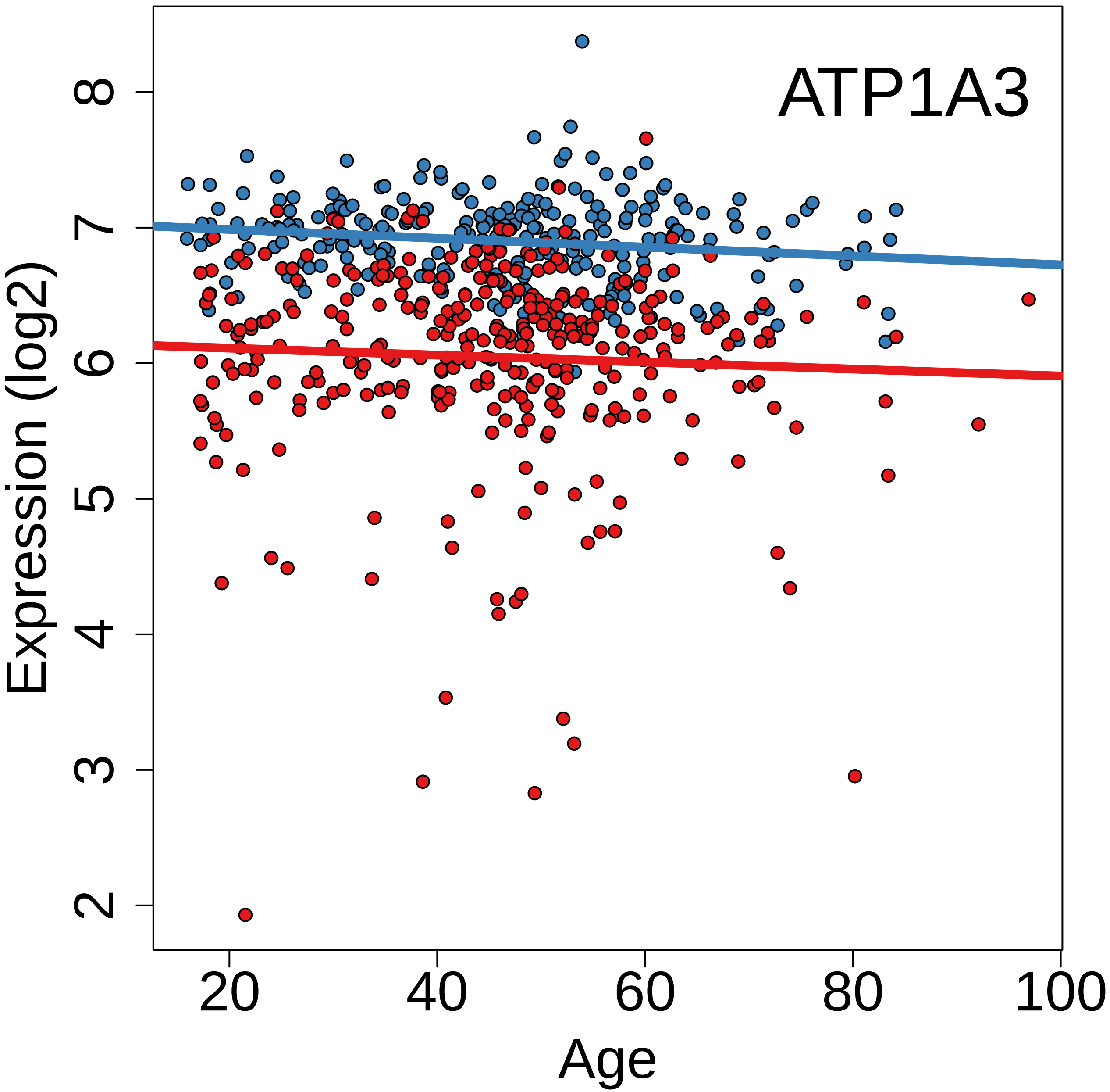
ATP1A3
Expression (log2)
Age
2
3
4
5
6
7
8
20
40
60
80
100

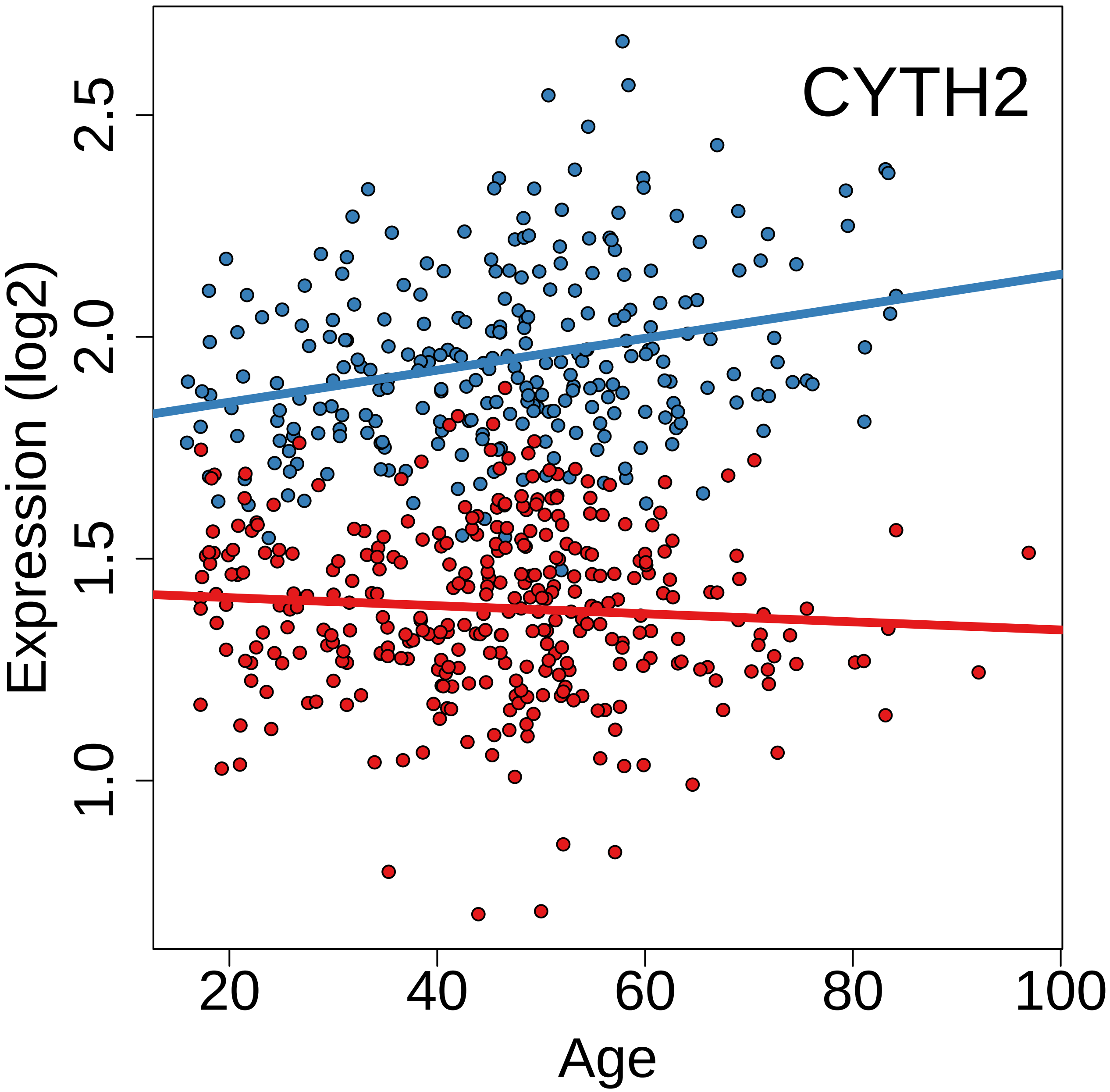
CYTH2
Expression (log2)
2.5
2.0
1.5
1.0
20
40
60
80
100
Age

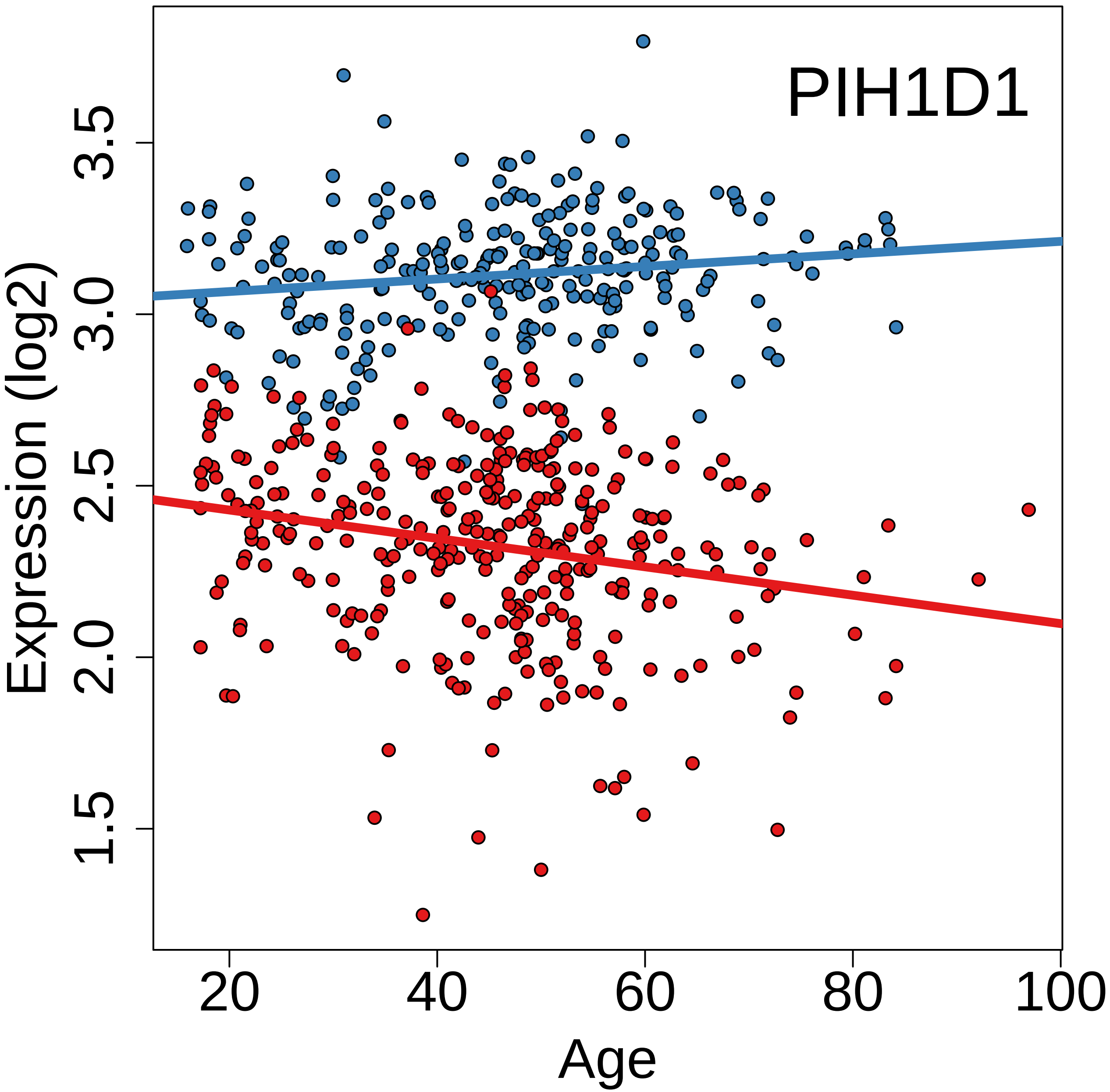
PIH1D1
Expression (log2)
1.5
2.0
2.5
3.0
3.5
20
40
60
80
100
Age

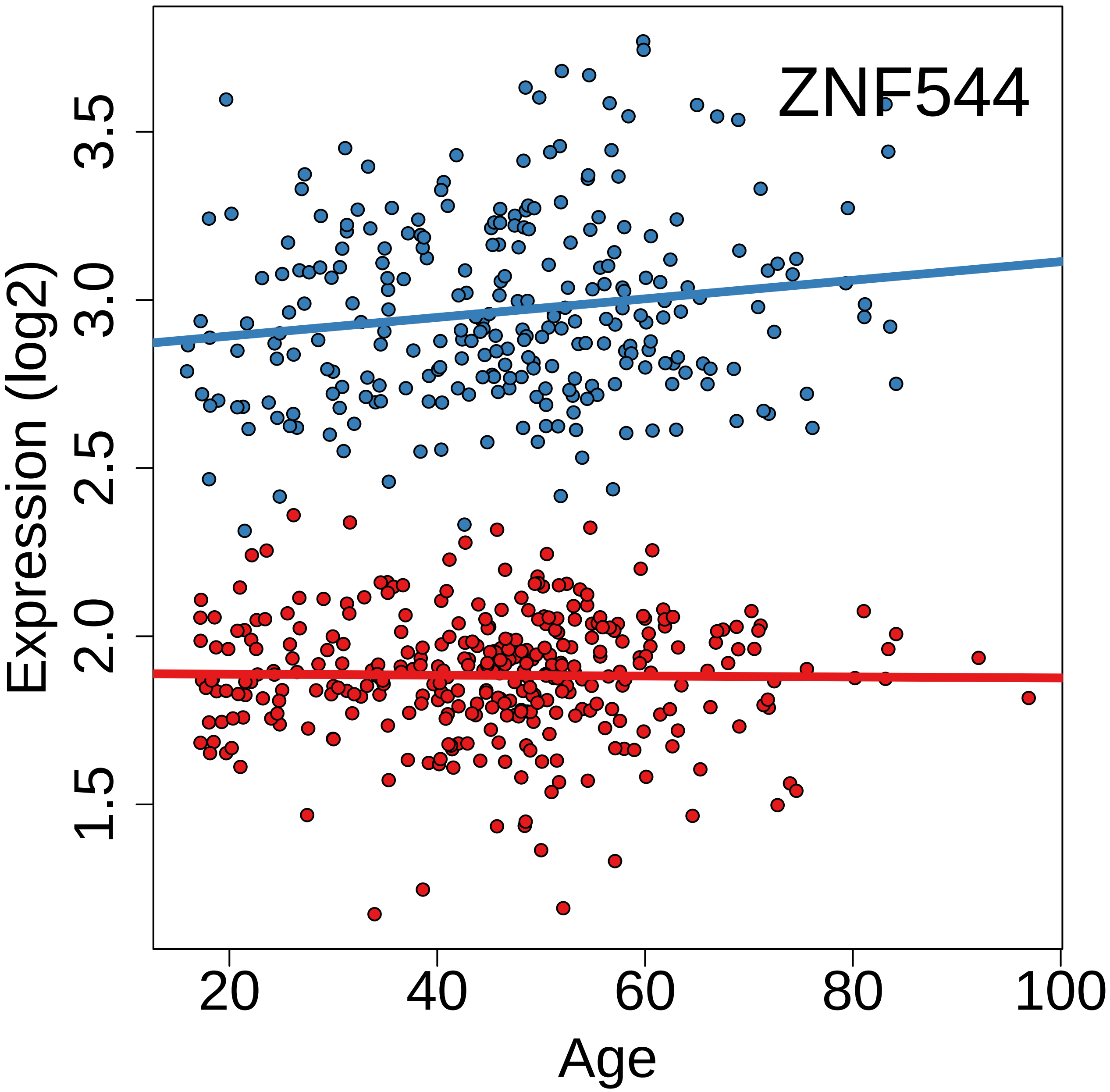
ZNF544
Expression (log2)
1.5
2.0
2.5
3.0
3.5
20
40
60
80
100
Age

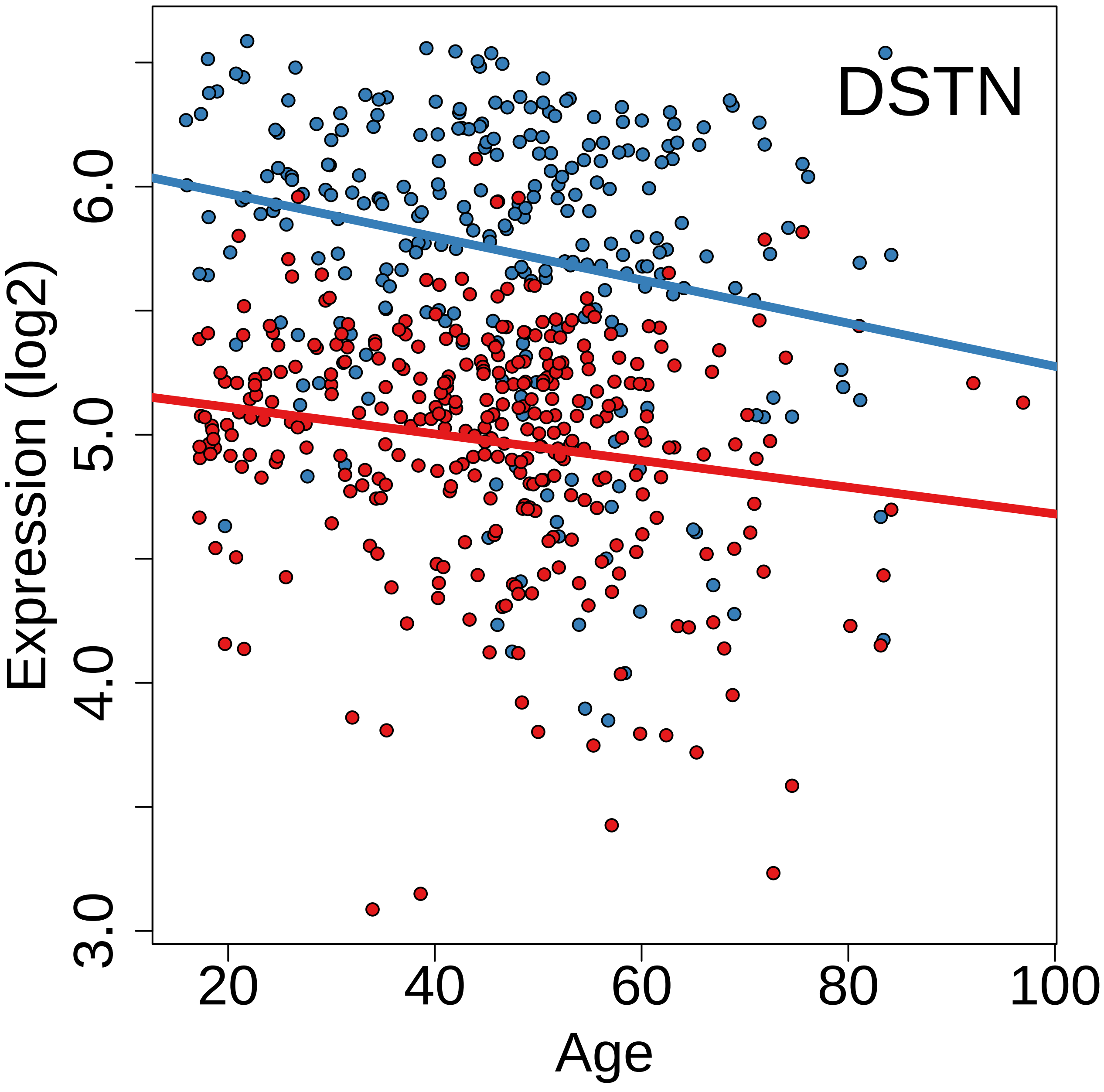
DSTN
Expression (log2)
3.0
4.0
5.0
6.0
20
40
60
80
100
Age

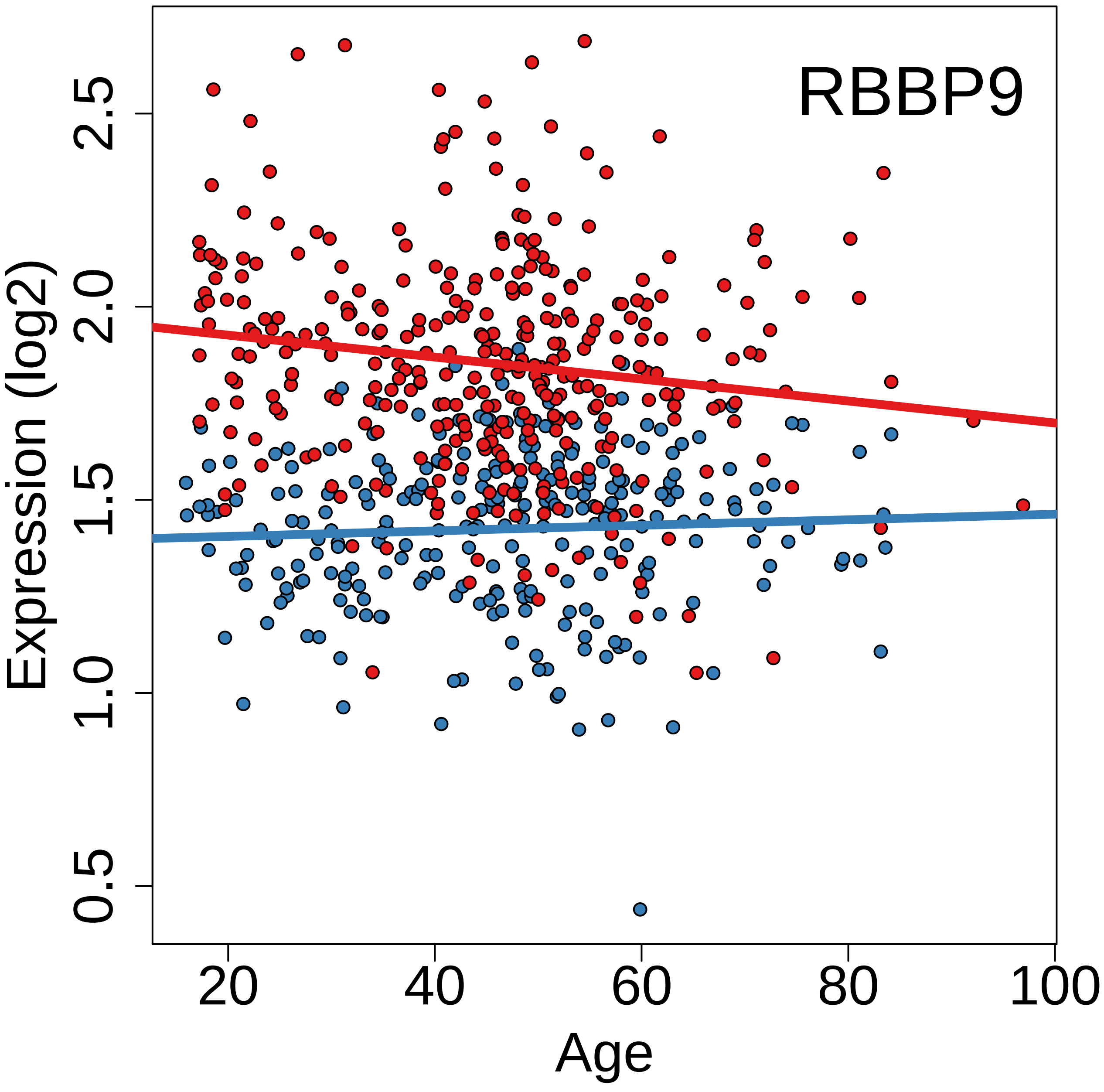
RBBP9
Expression (log2)
0.5
1.0
1.5
2.0
2.5
20
40
60
80
100
Age

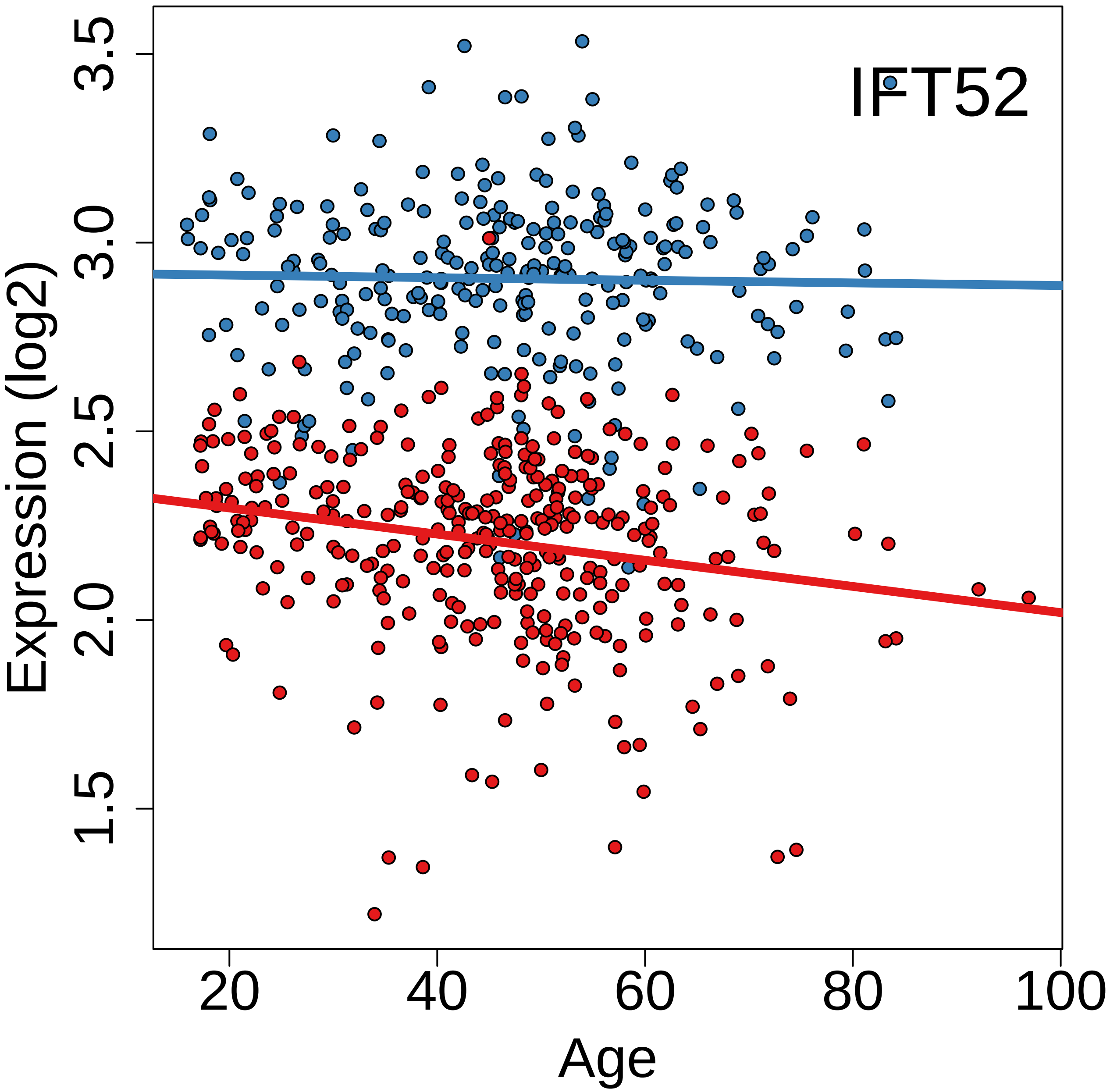

IFT52
Expression (log2)
3.5
3.0
2.5
2.0
1.5
20
40
60
80
100
Age

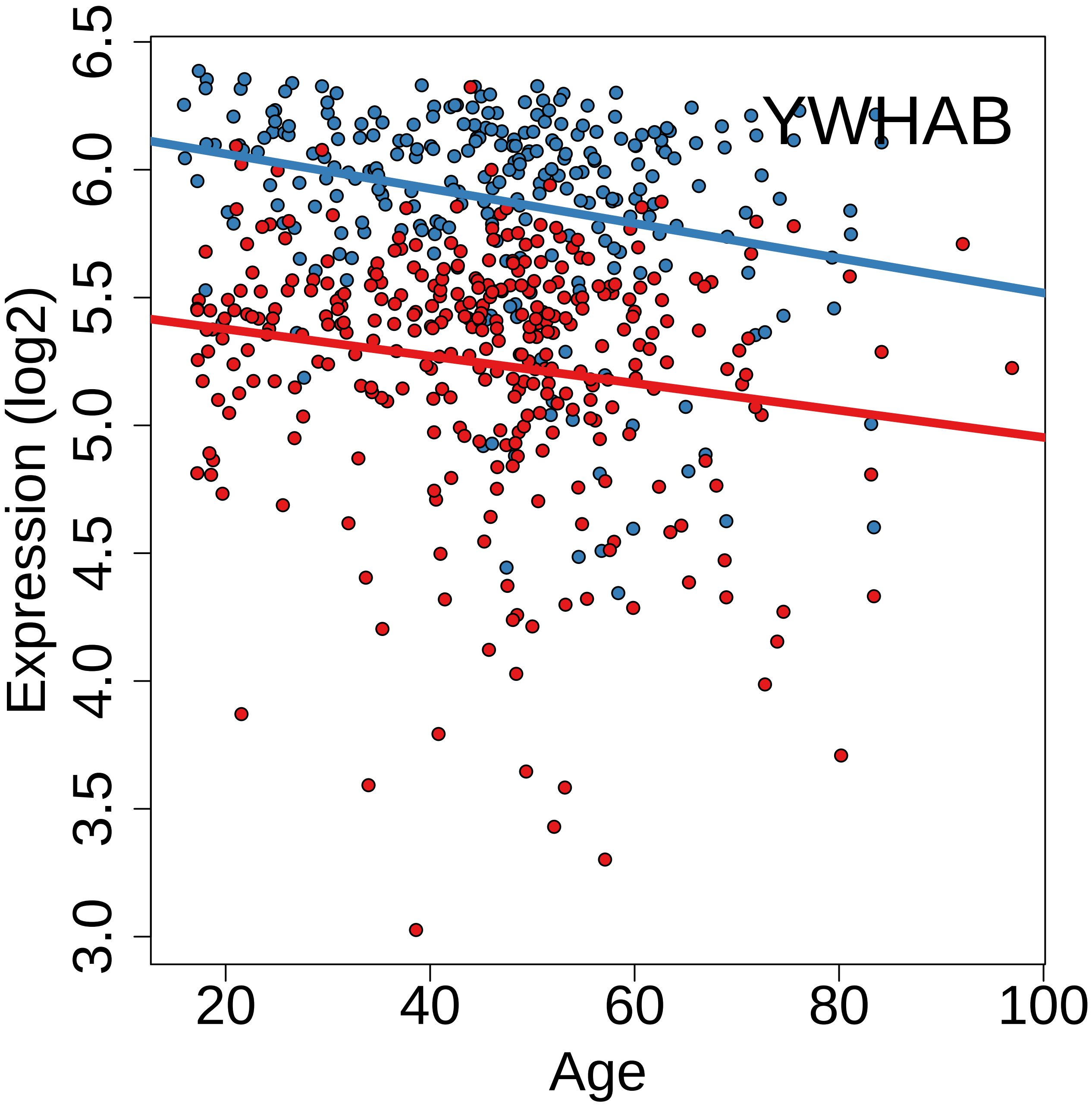

YWHAB
Expression (log2)
Age
3.0
3.5
4.0
4.5
5.0
5.5
6.0
6.5
20
40
60
80
100

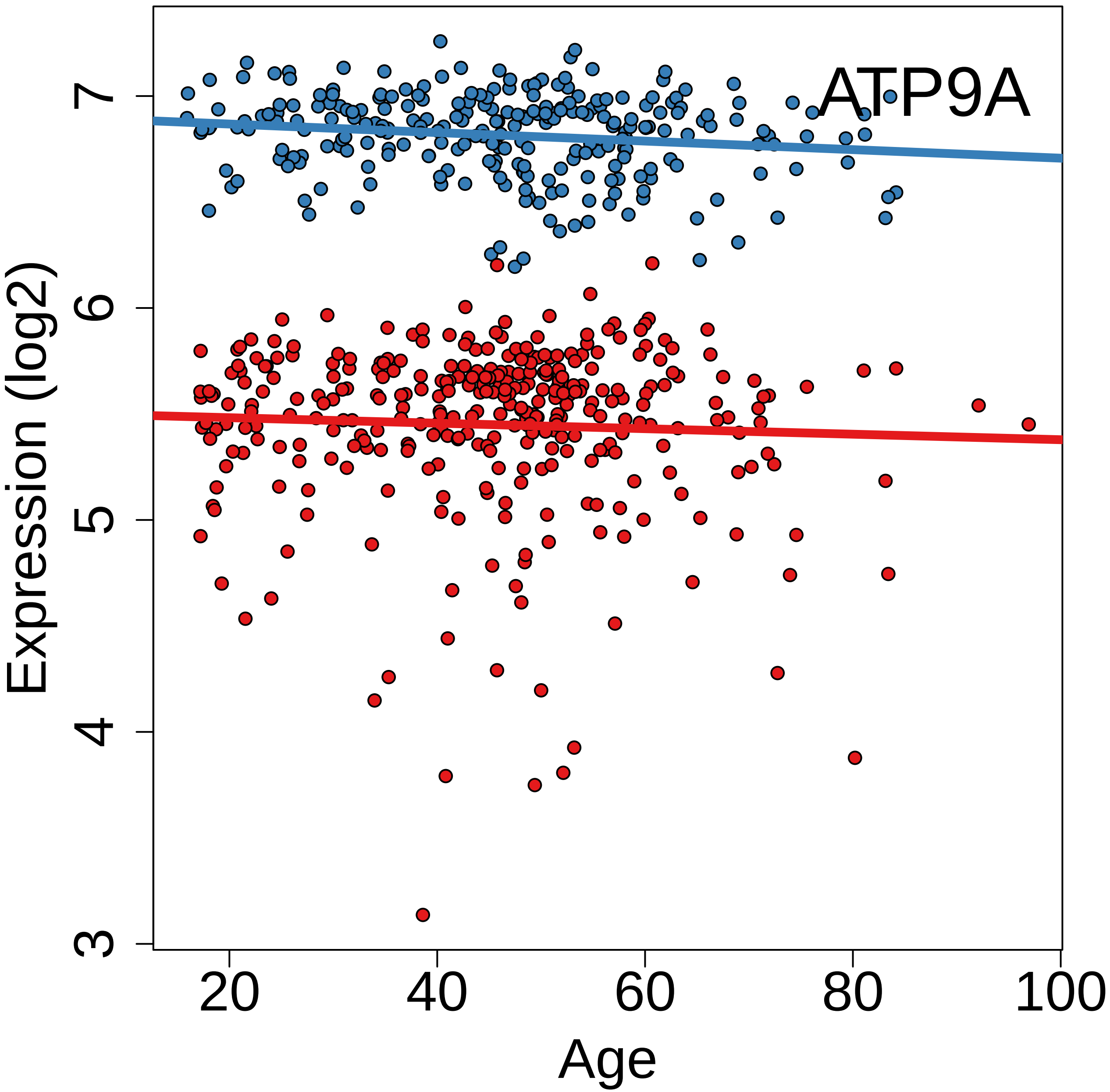
ATP9A
Expression (log2)
3
4
5
6
7
Age
20
40
60
80
100

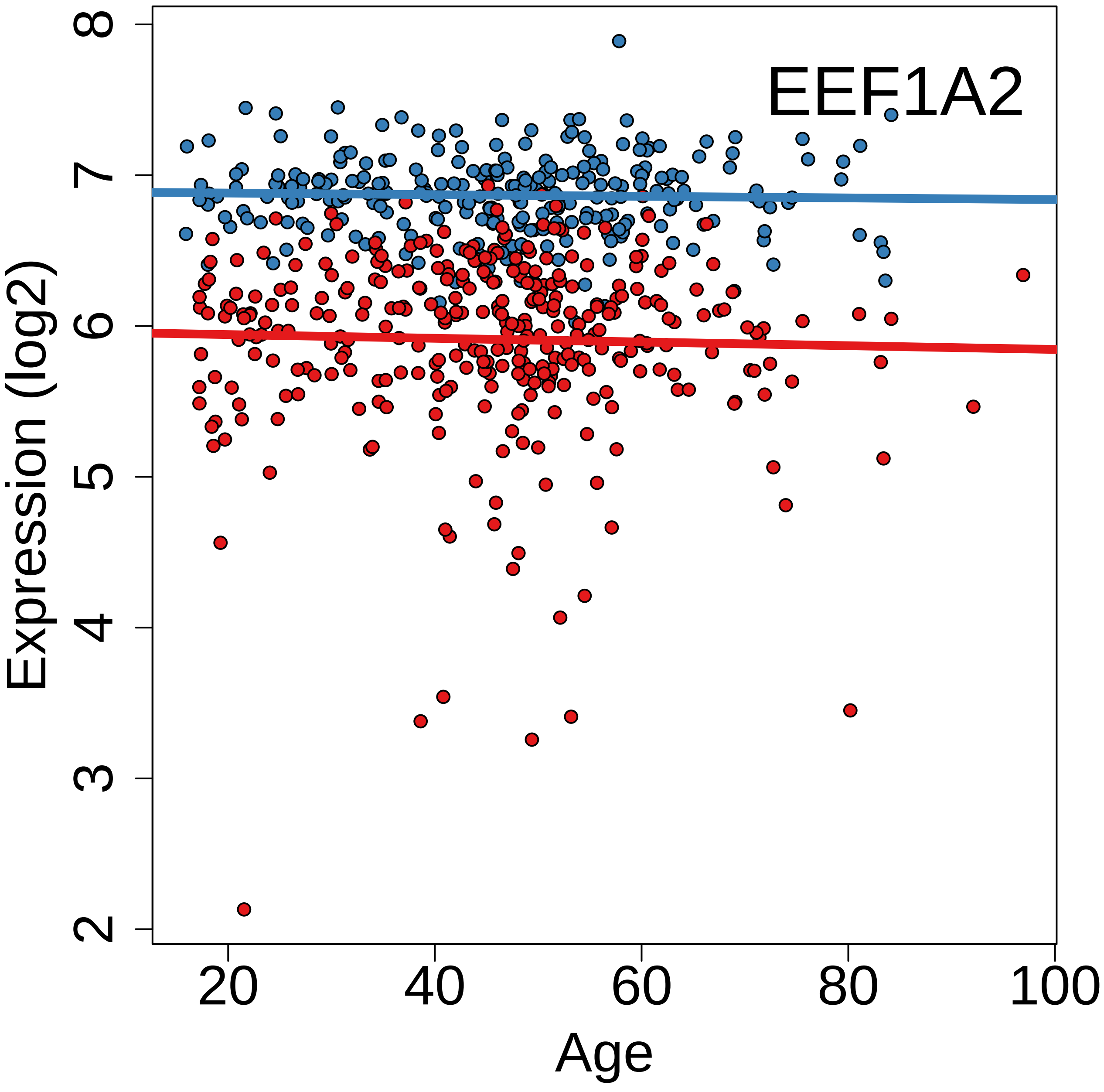
EEF1A2
Expression (log2)
Age
2
3
4
5
6
7
8
20
40
60
80
100

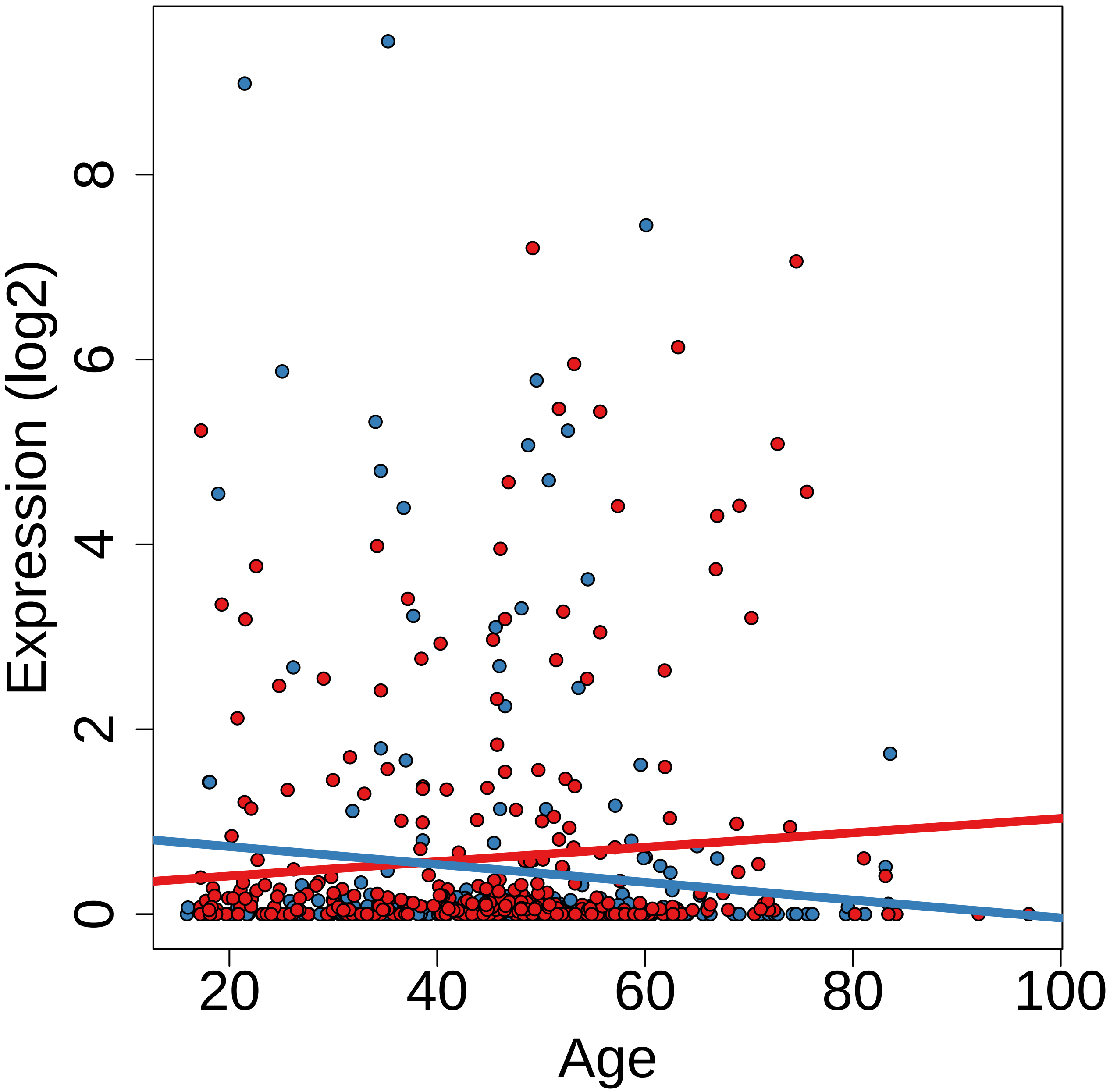

Expression (log2)
0
2
4
6
8
20
40
60
80
100
Age

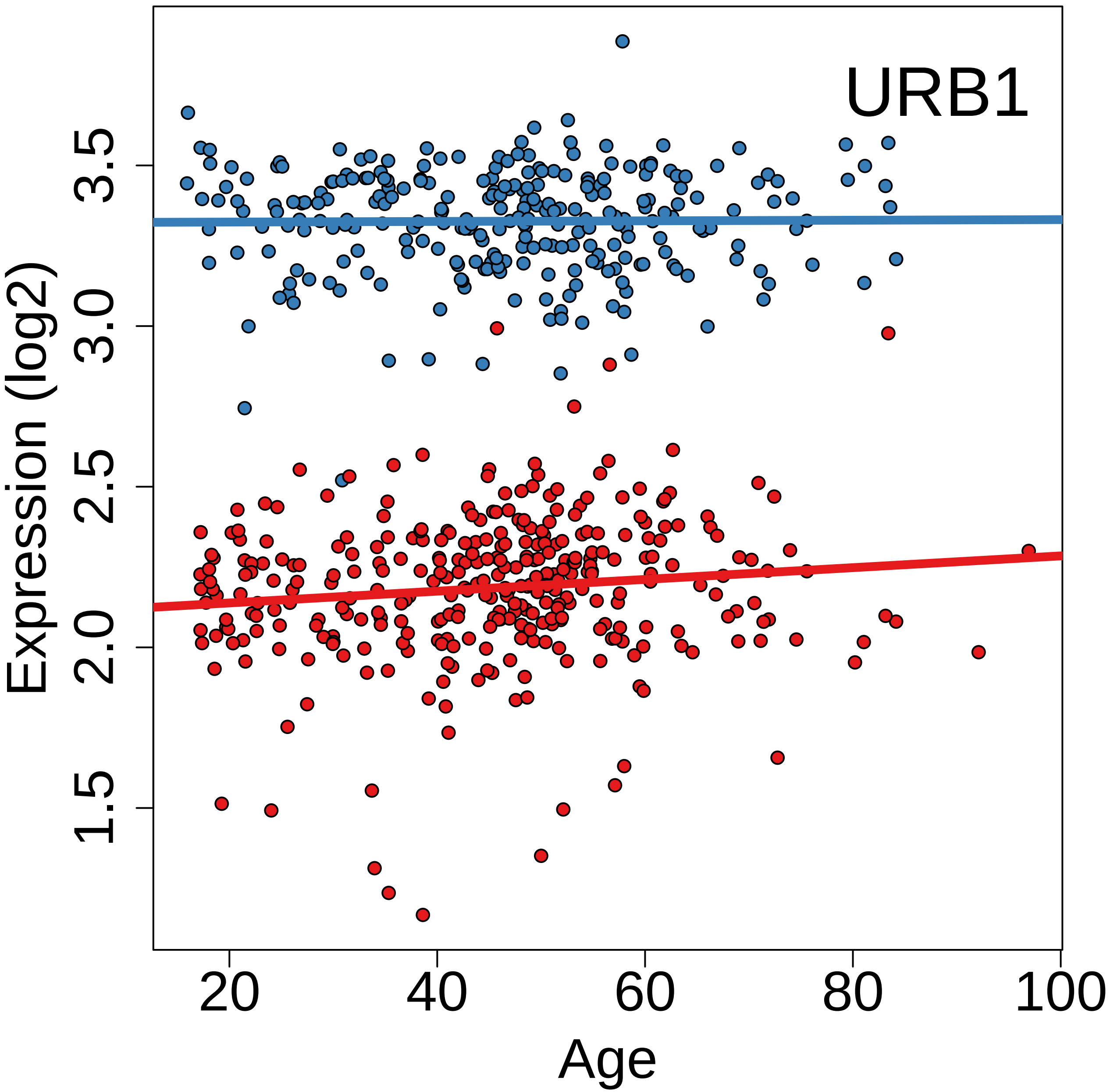

URB1
Expression (log2)
1.5
2.0
2.5
3.0
3.5
20
40
60
80
100
Age

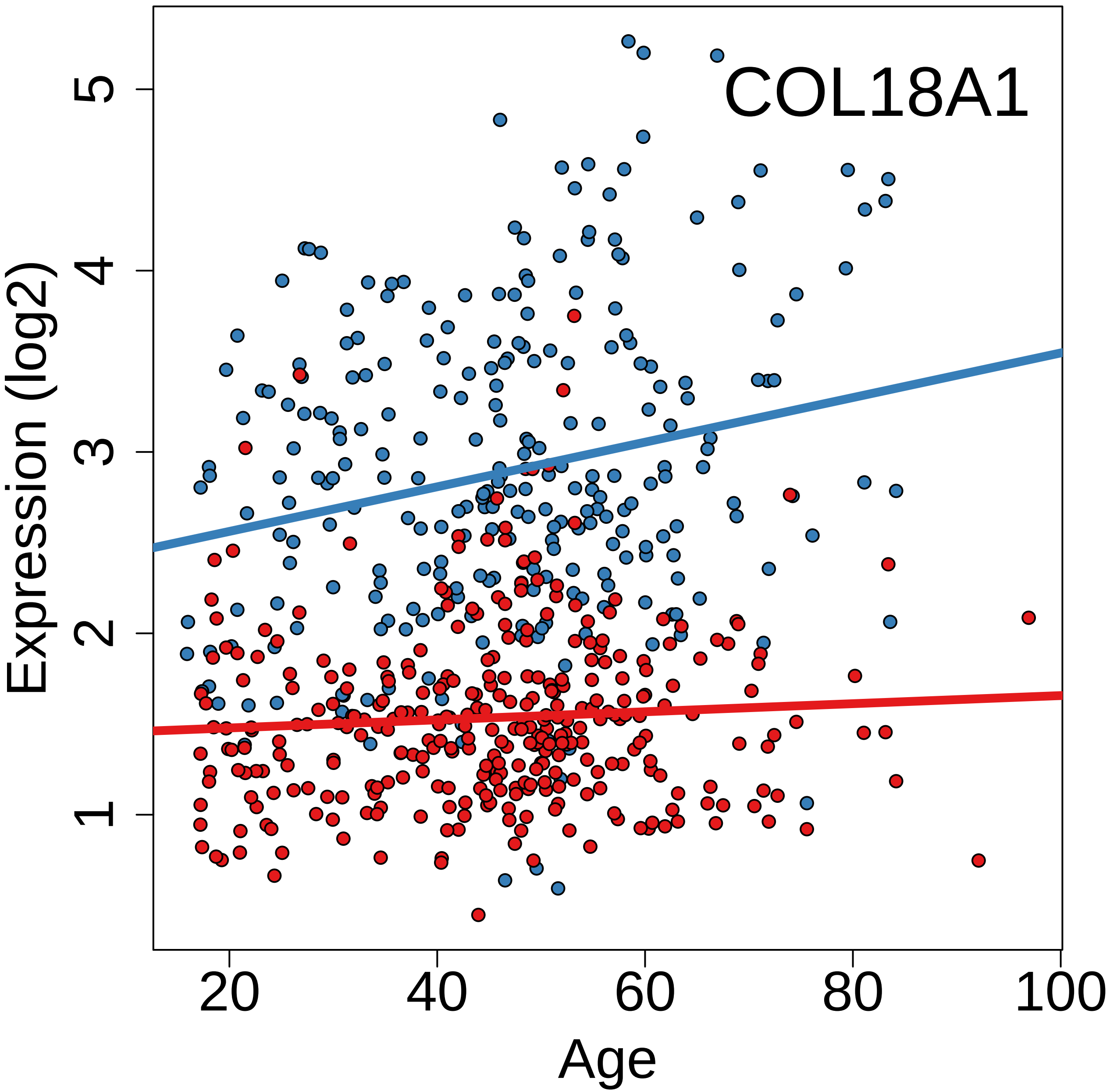
COL18A1
Expression (log2)
1
2
3
4
5
20
40
60
80
100
Age

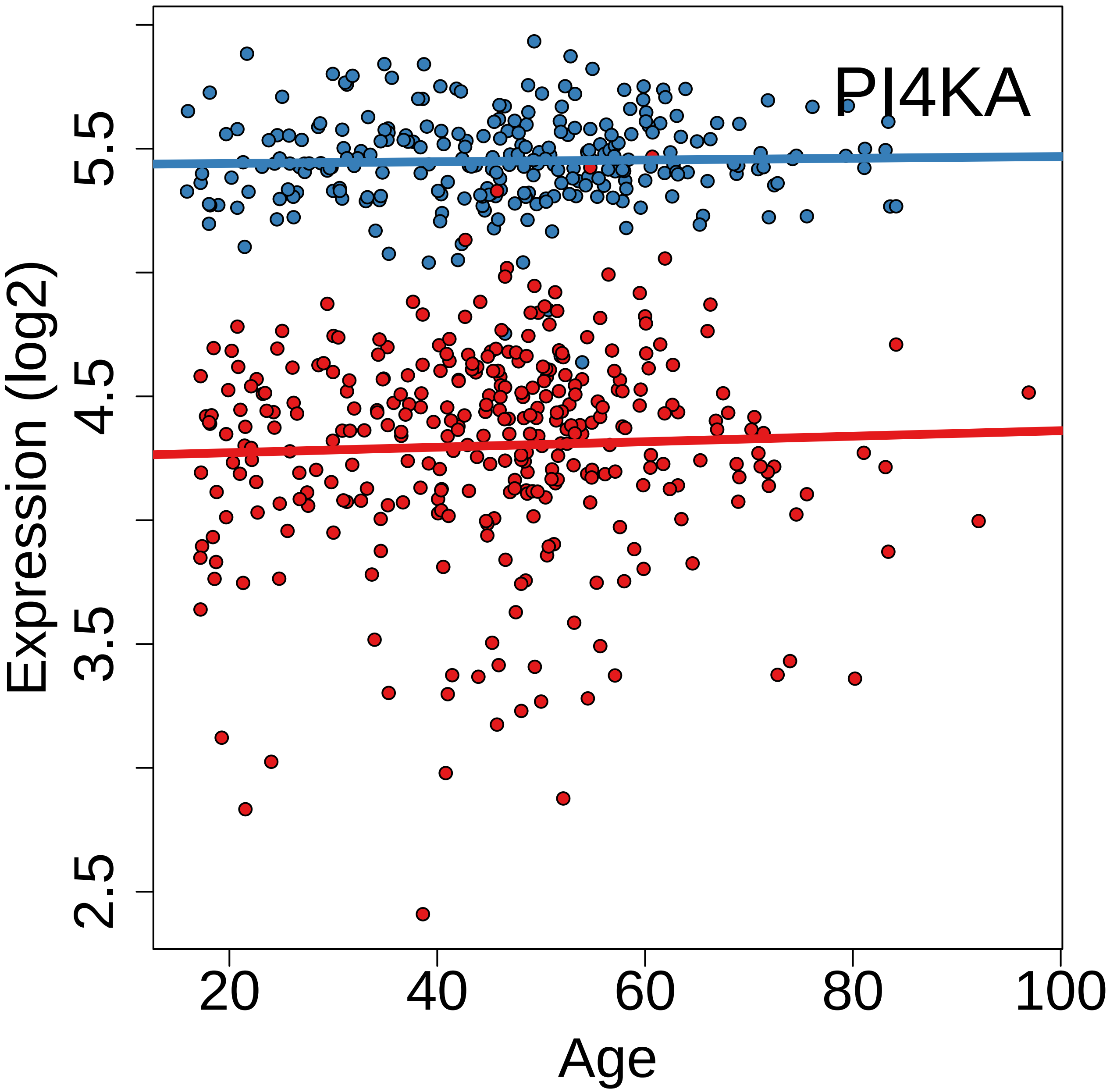
PI4KA
Expression (log2)
2.5
3.5
4.5
5.5
20
40
60
80
100
Age

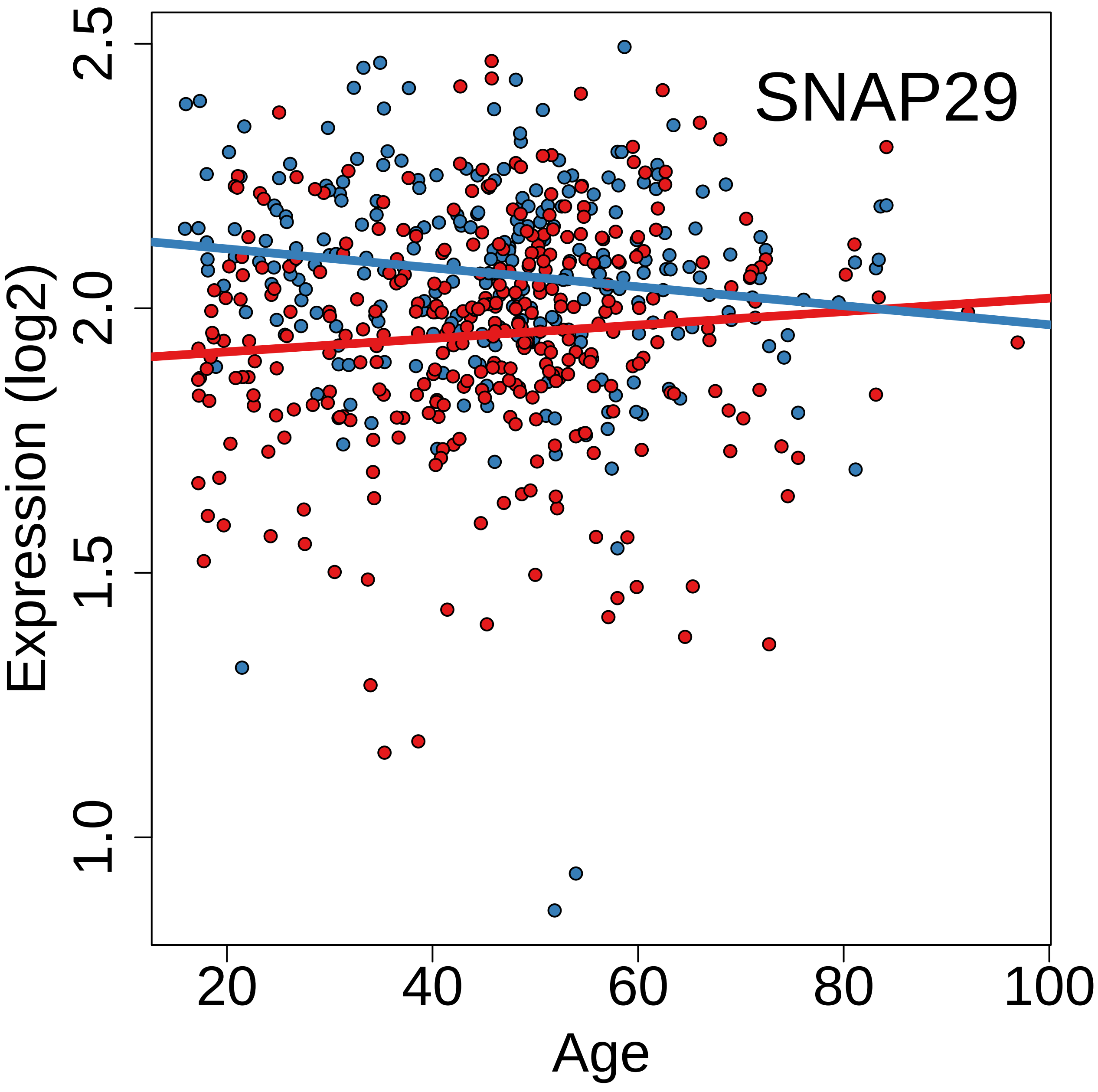

SNAP29
Expression (log2)
1.0
1.5
2.0
2.5
20
40
60
80
100
Age

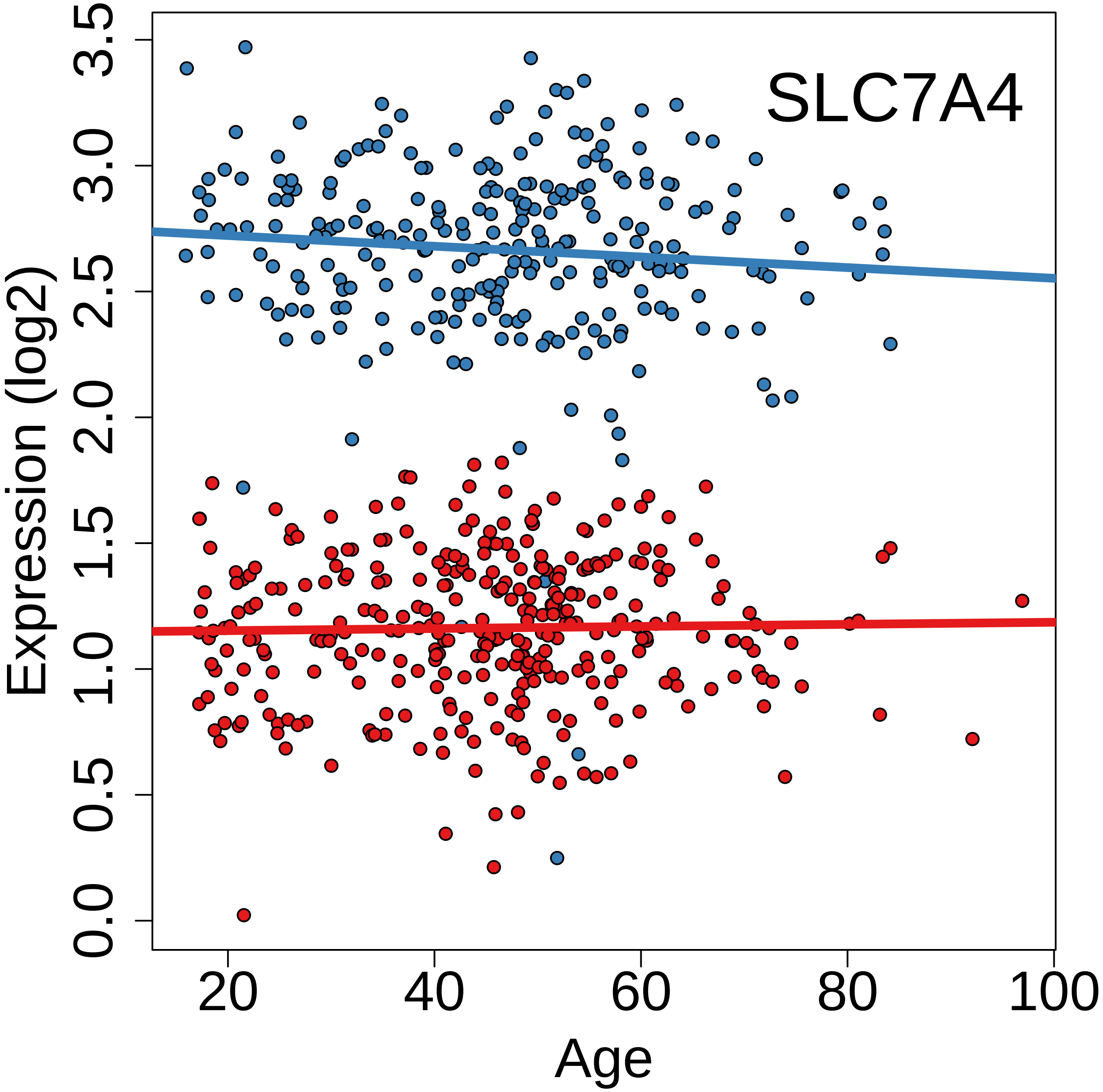

SLC7A4
Expression (log2)
Age
0.0
0.5
1.0
1.5
2.0
2.5
3.0
3.5
20
40
60
80
100

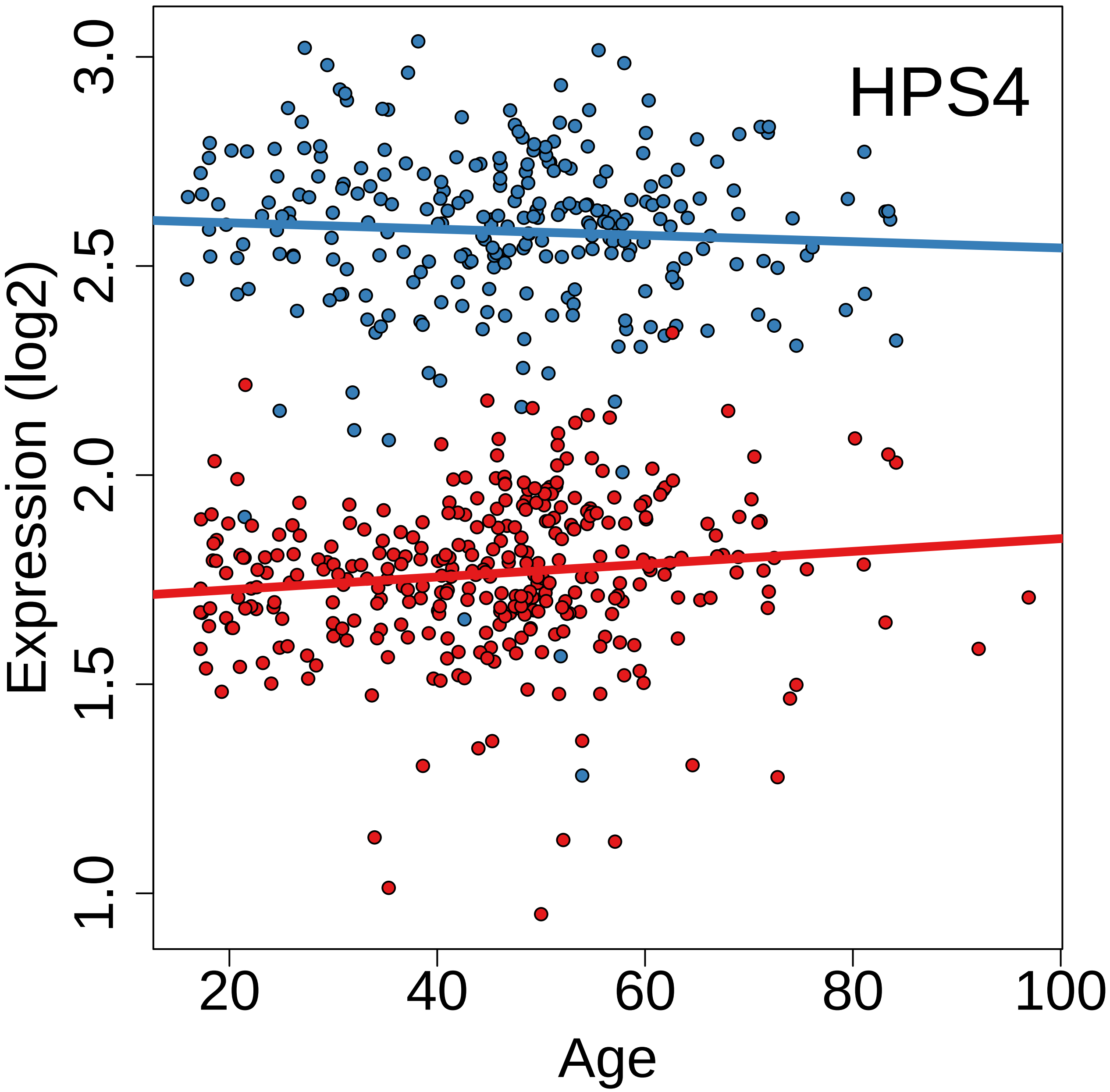
HPS4
Expression (log2)
Age
1.0
1.5
2.0
2.5
3.0
20
40
60
80
100

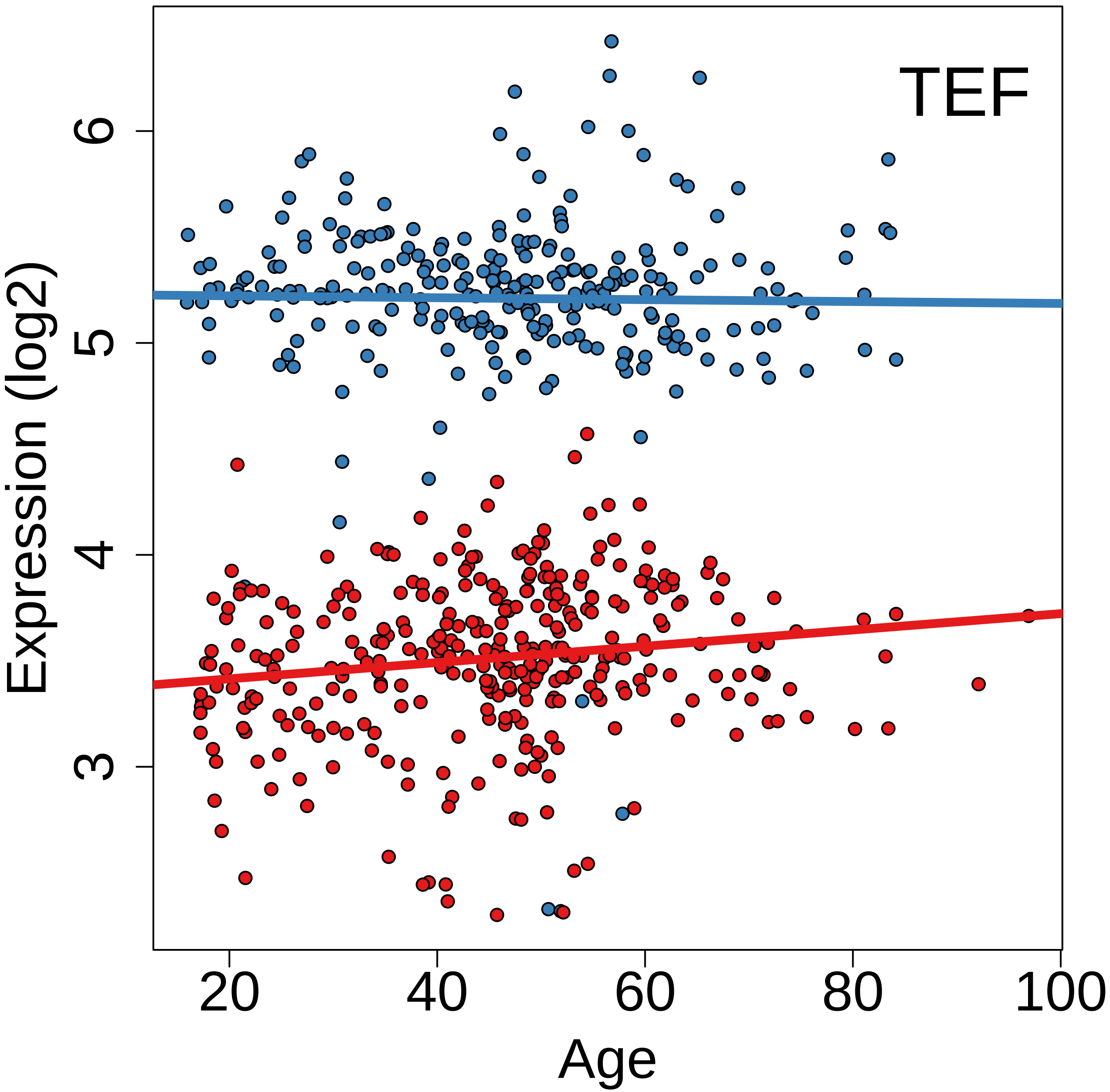
TEF
Expression (log2)
3
4
5
6
20
40
60
80
100
Age

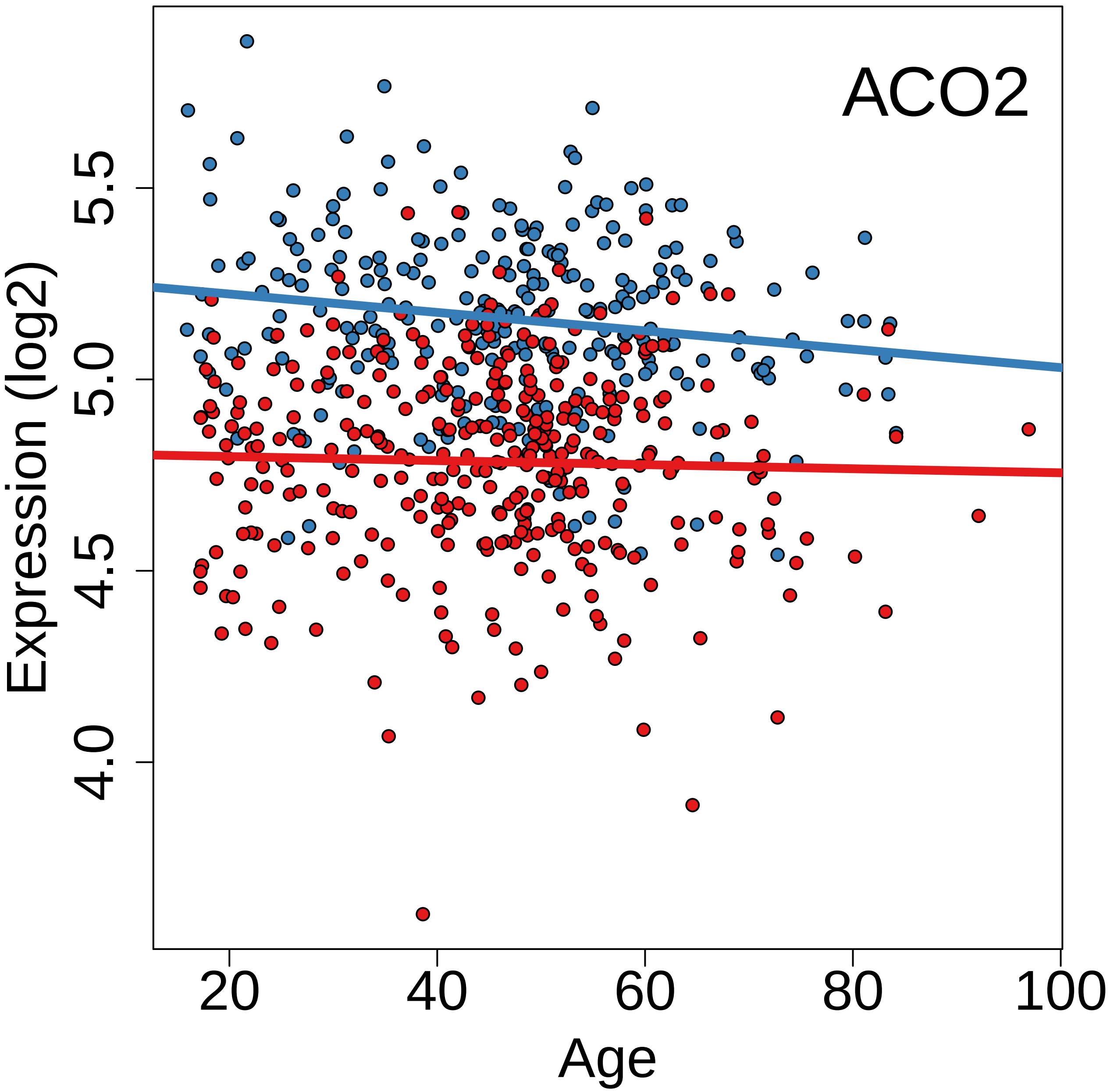
ACO2
Expression (log2)
4.0
4.5
5.0
5.5
20
40
60
80
100
Age

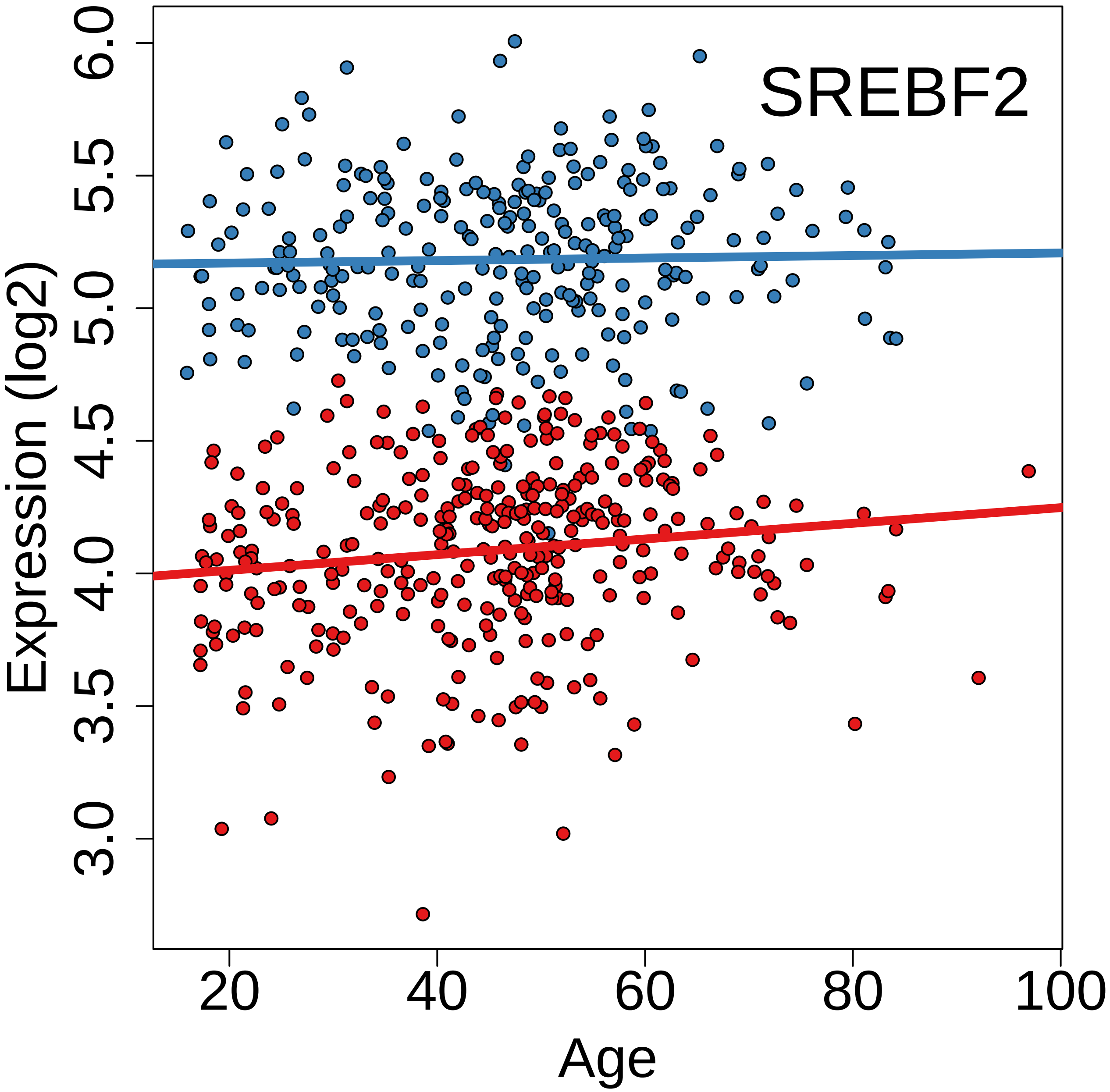
SREBF2
Expression (log2)
Age
6.0
5.5
5.0
4.5
4.0
3.5
3.0
20
40
60
80
100

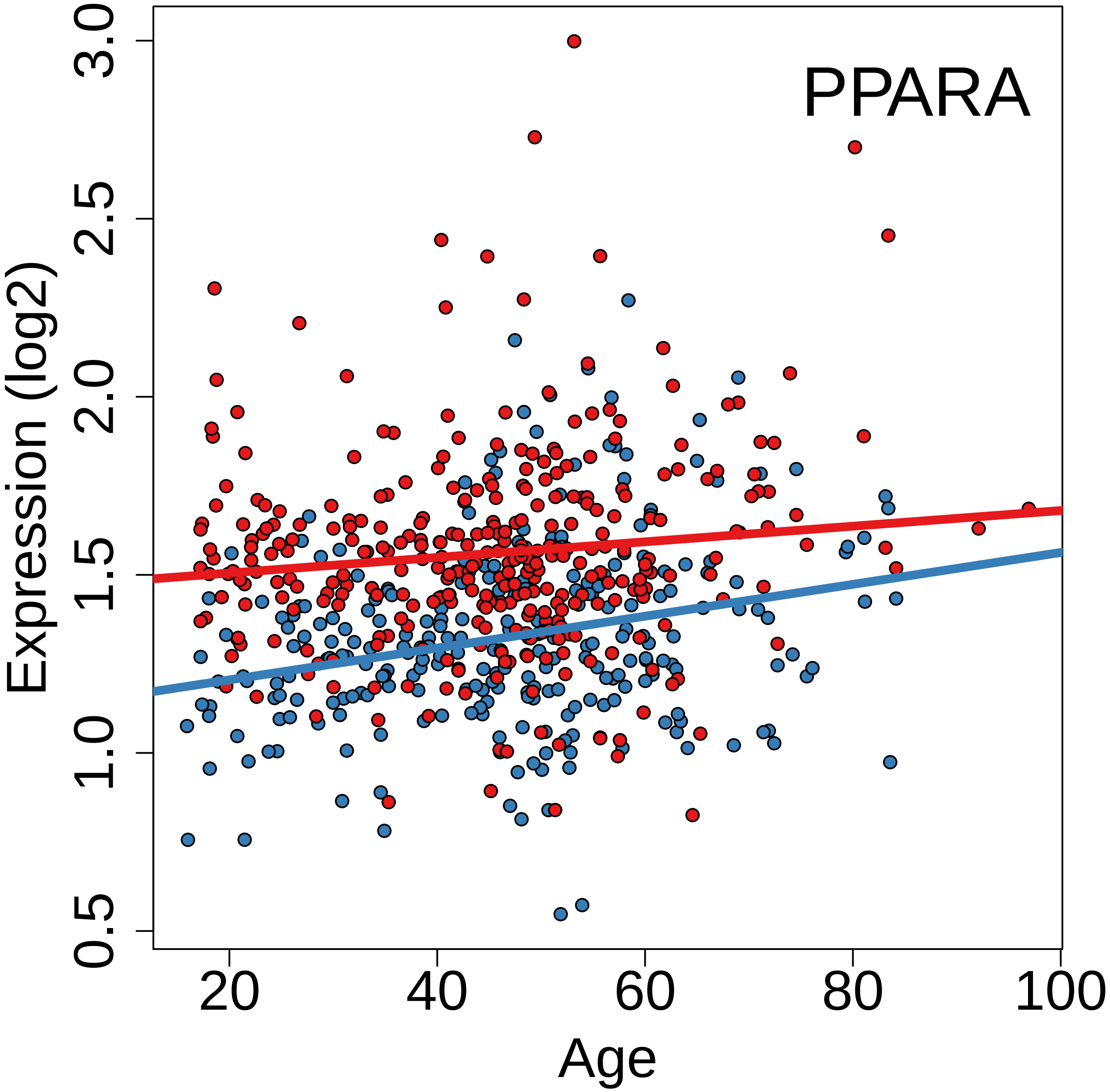
PPARA
Expression (log2)
0.5
1.0
1.5
2.0
2.5
3.0
20
40
60
80
100
Age

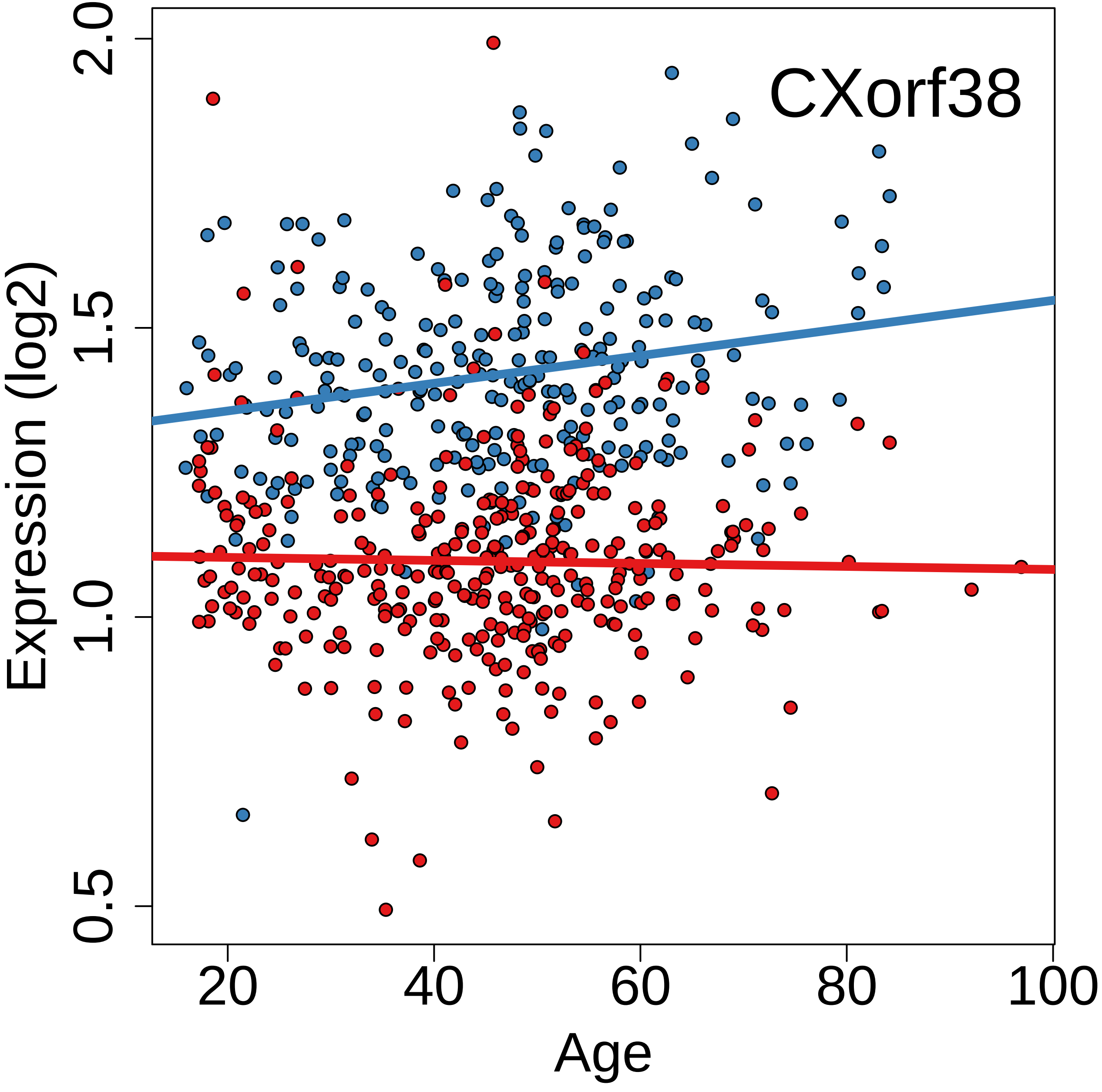
CXorf38
Expression (log2)
2.0
1.5
1.0
0.5
20
40
60
80
100
Age

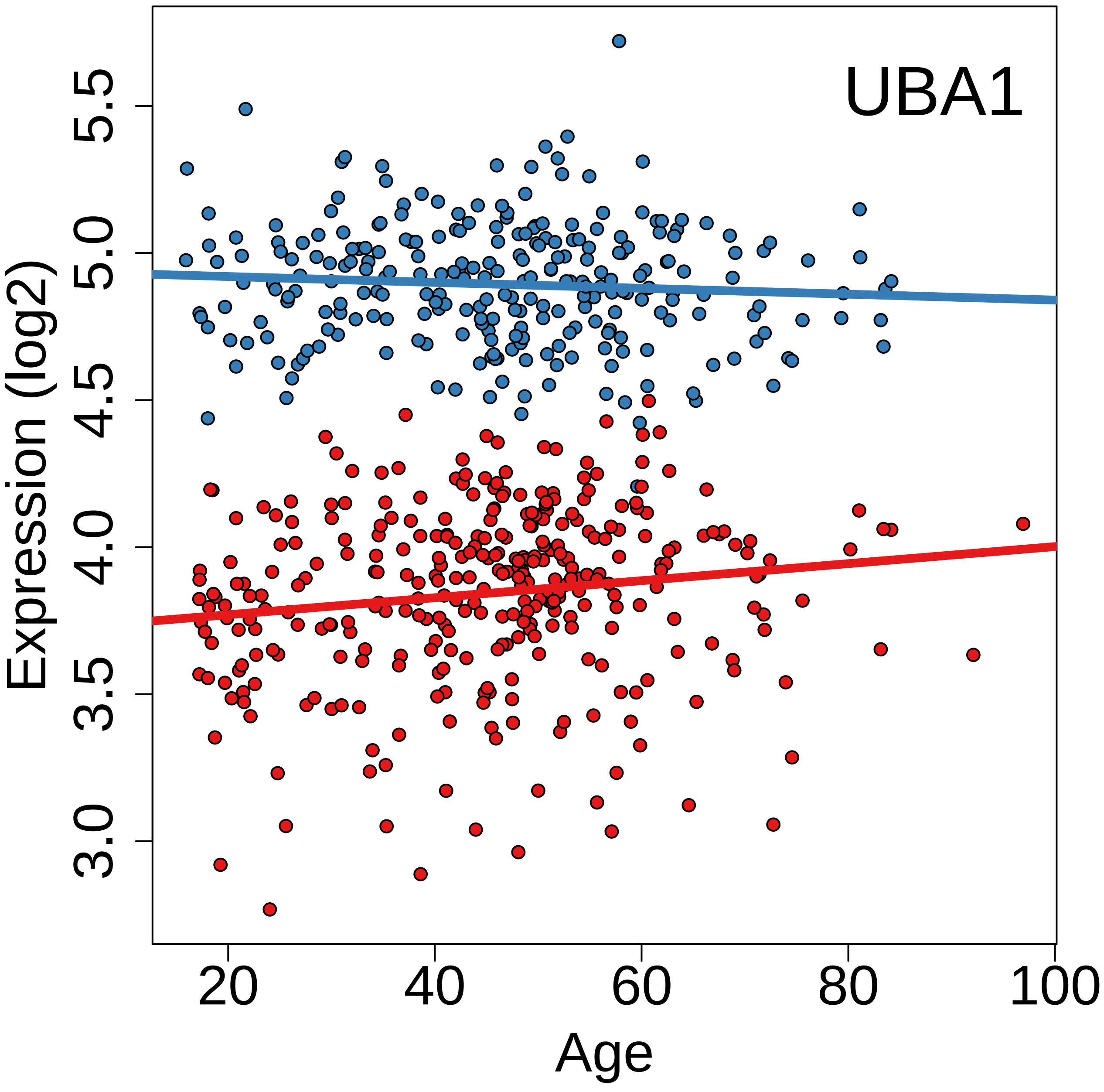

UBA1
Expression (log2)
3.0
3.5
4.0
4.5
5.0
5.5
20
40
60
80
100
Age

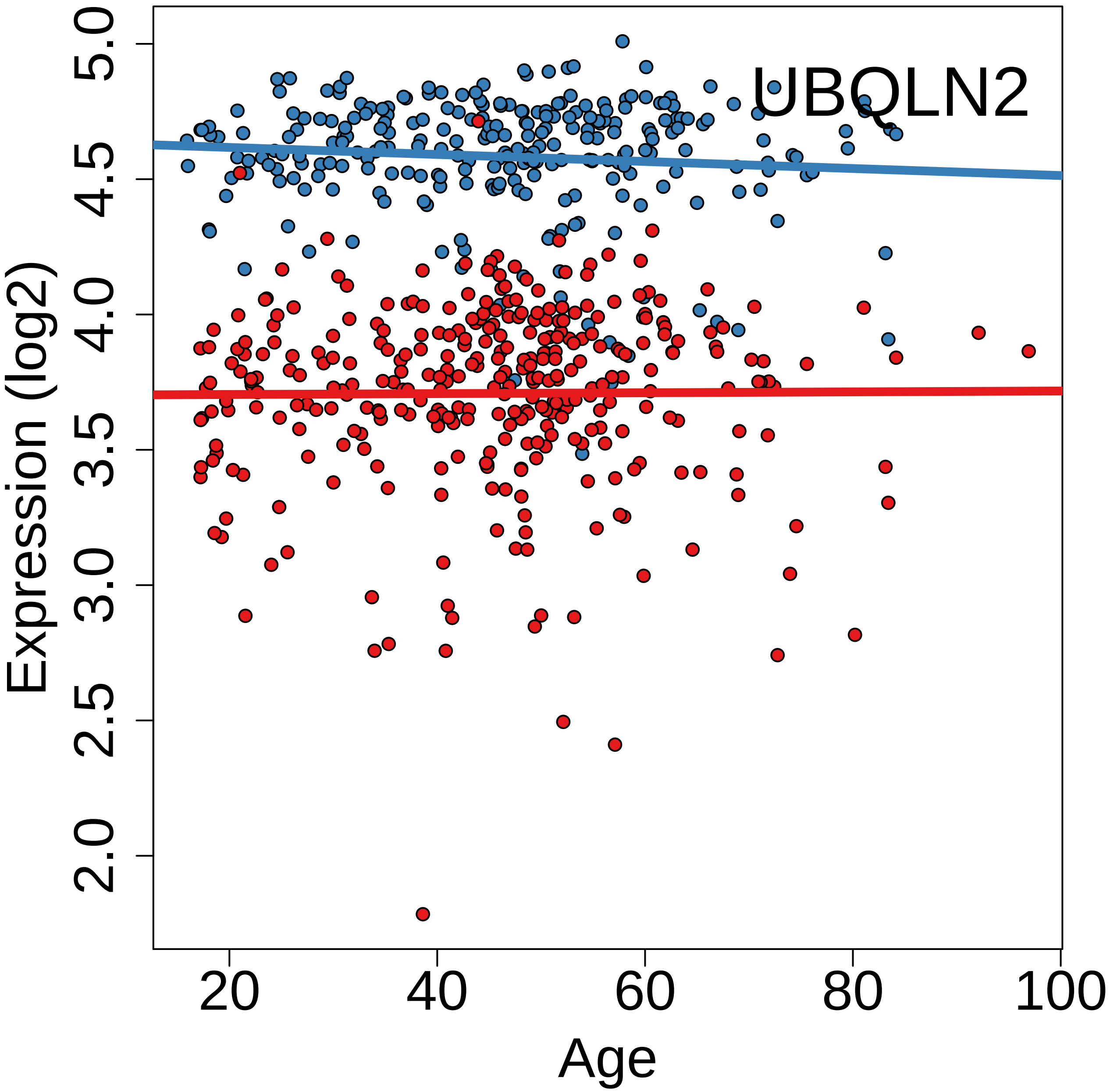
UBQLN2
Expression (log2)
5.0
4.5
4.0
3.5
3.0
2.5
2.0
Age
20
40
60
80
100

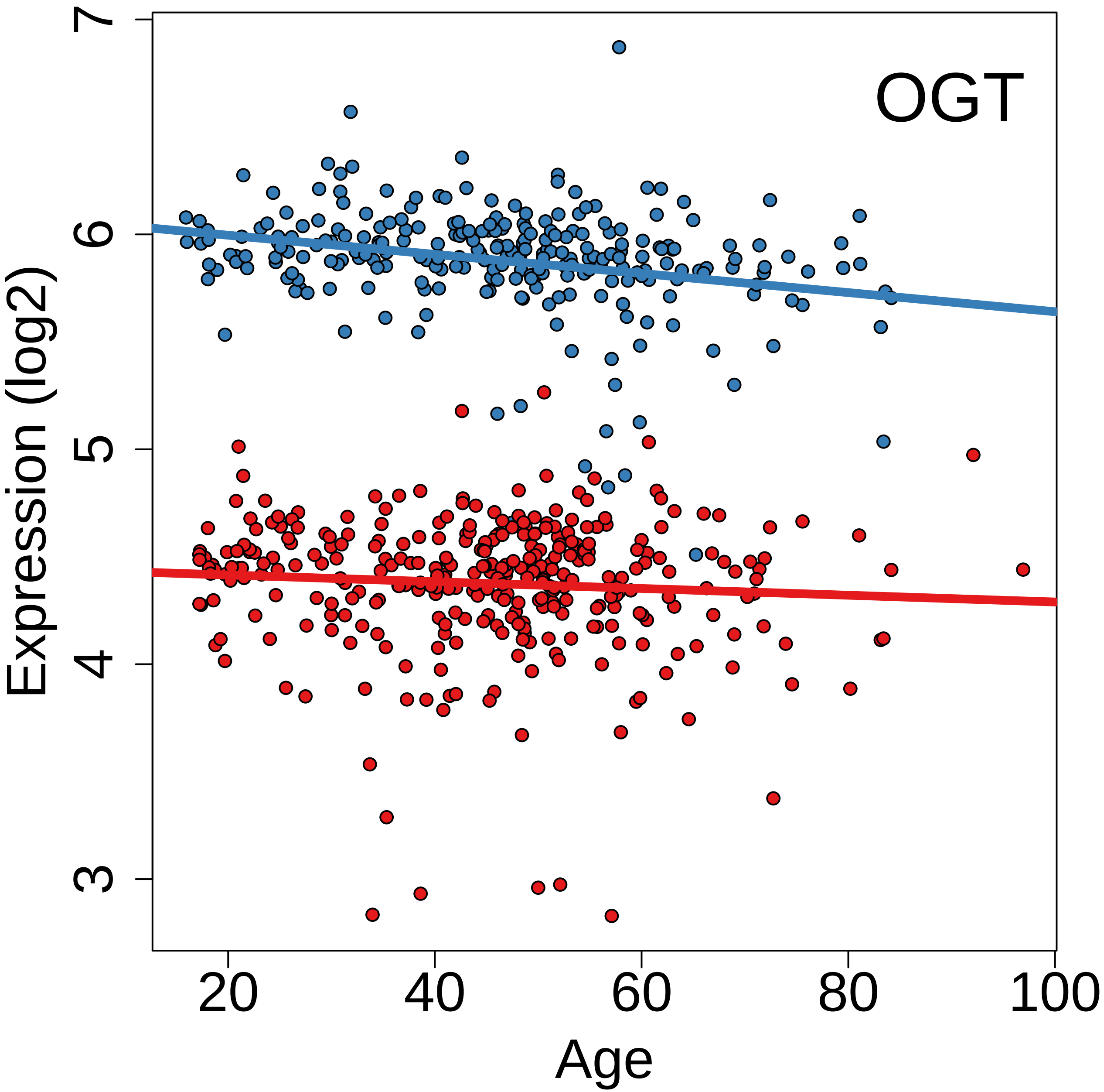
OGT
Expression (log2)
7
6
5
4
3
20
40
60
80
100
Age

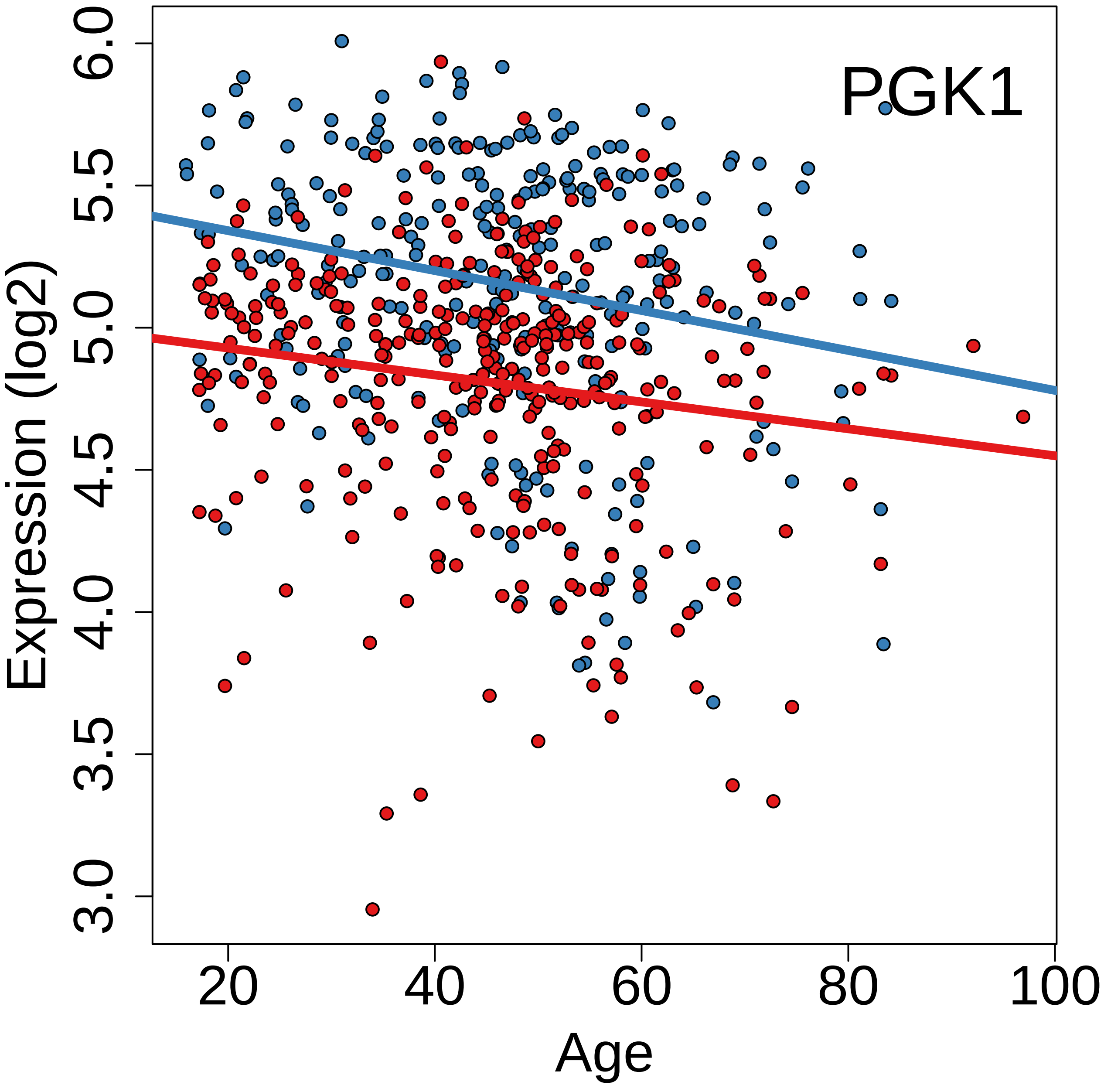

PGK1
Expression (log2)
3.0
3.5
4.0
4.5
5.0
5.5
6.0
20
40
60
80
100
Age

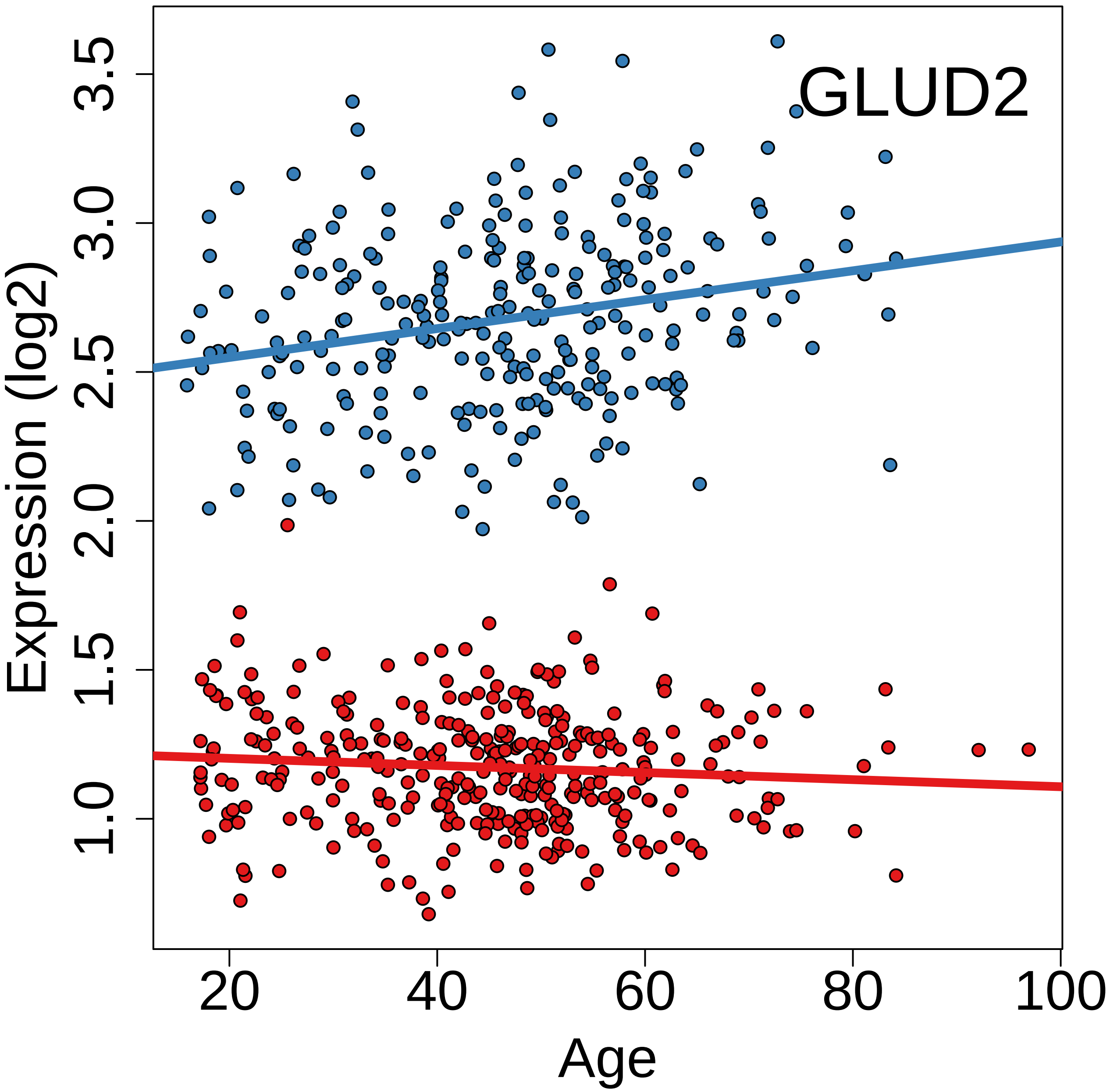

GLUD2
Expression (log2)
1.0
1.5
2.0
2.5
3.0
3.5
20
40
60
80
100
Age

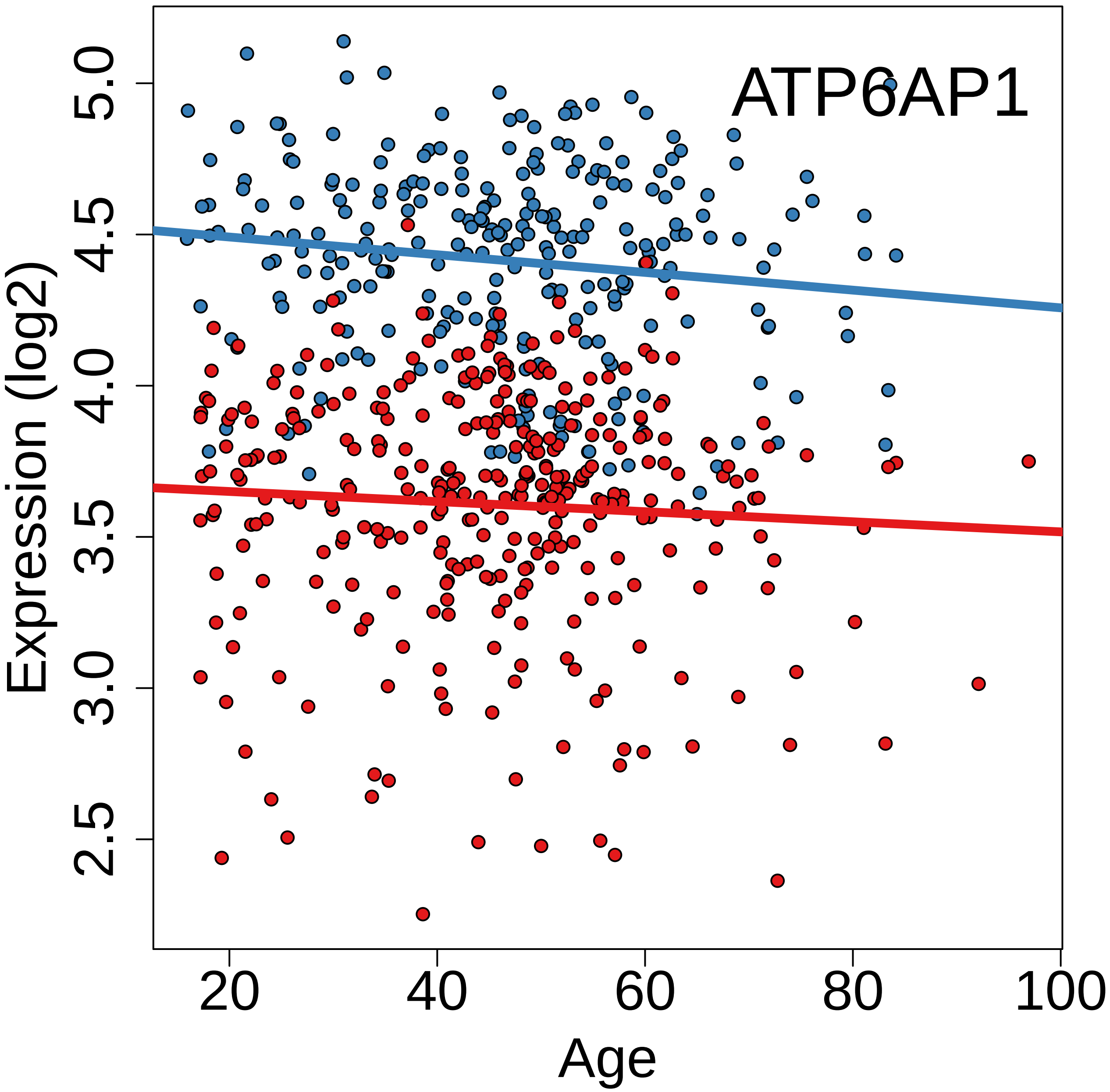
ATP6AP1
Expression (log2)
5.0
4.5
4.0
3.5
3.0
2.5
20
40
60
80
100
Age

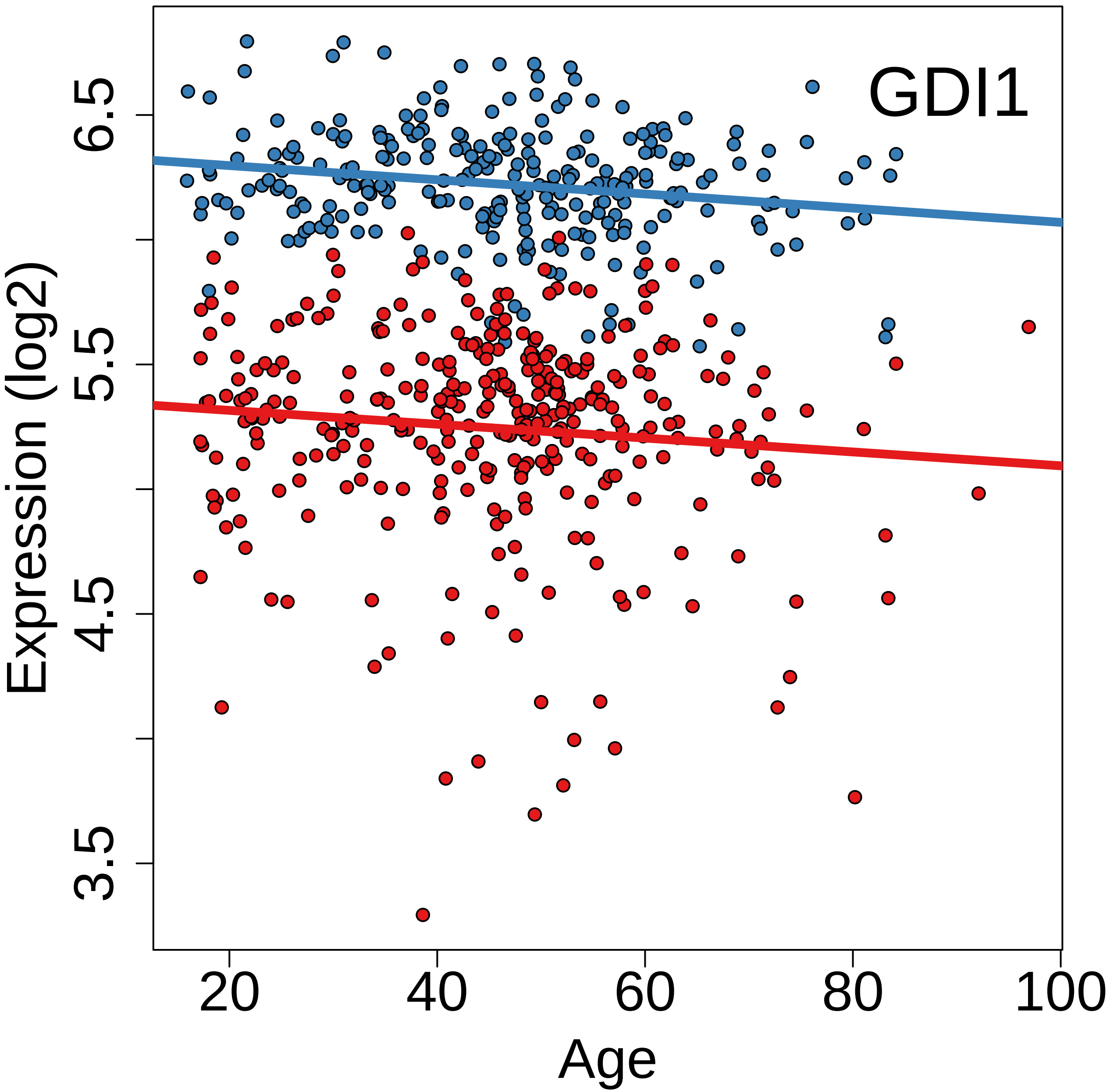
GDI1
Expression (log2)
3.5
4.5
5.5
6.5
20
40
60
80
100
Age

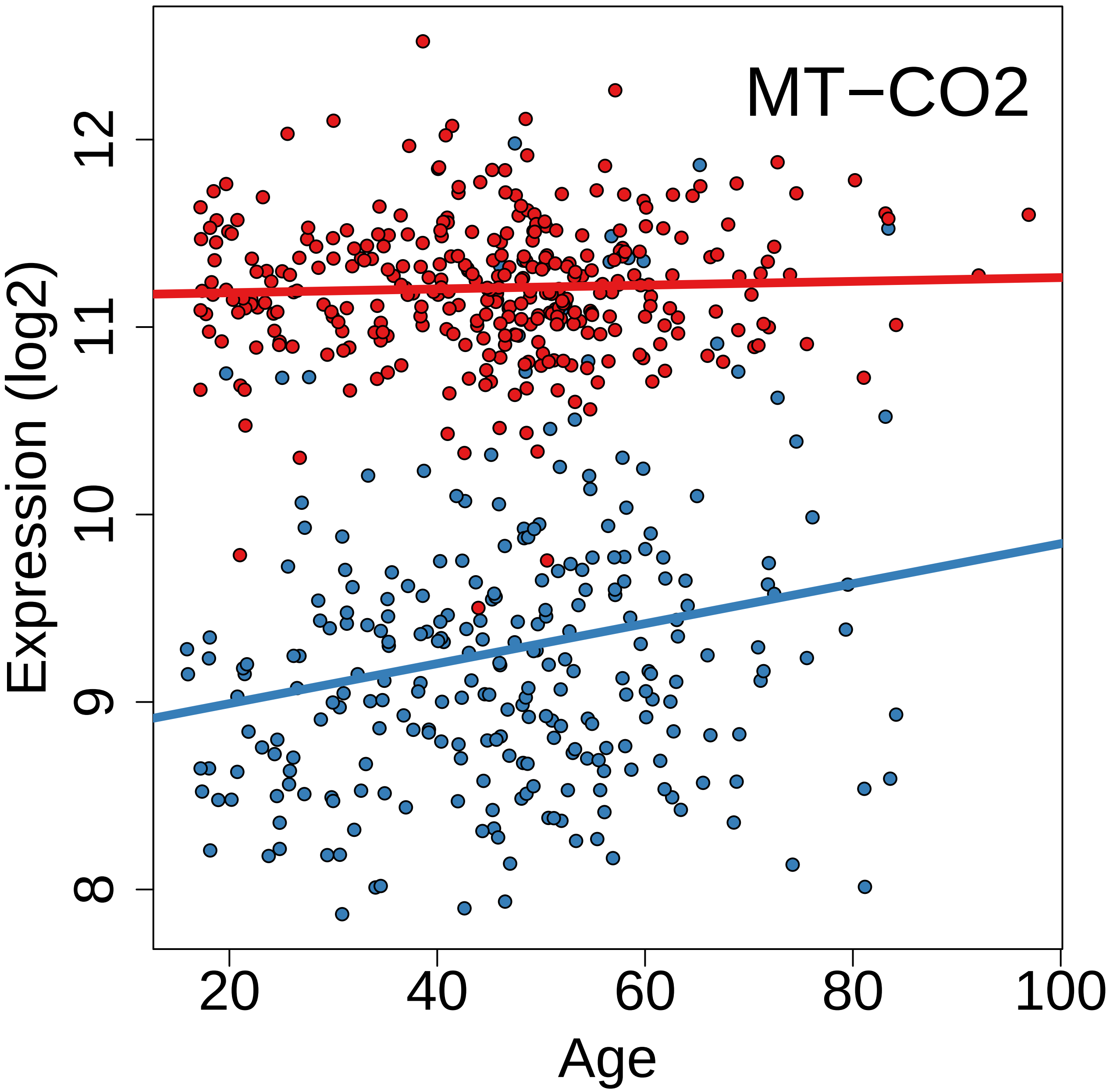
MT−CO2
Expression (log2)
12
11
10
9
8
20
40
60
80
100
Age
