## Supplementary material for "Cell type-specific genetic regulation of expression in the granule cell layer of the human dentate gyrus": suppData3_eQTLexamples.pdf

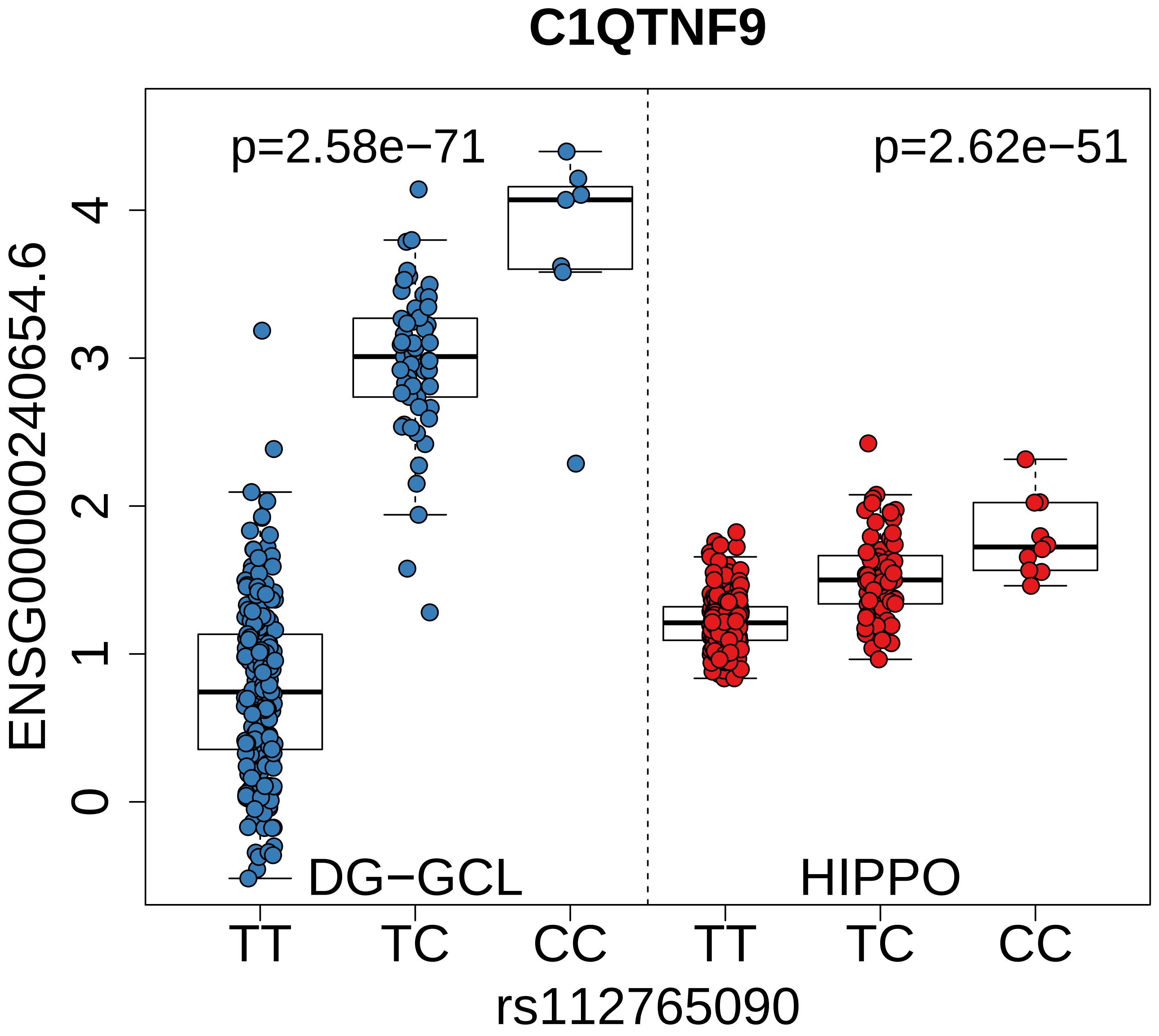
C1QTNF9
p=2.58e−71
p=2.62e−51
ENSG00000240654.6
4
3
2
1
0
DG−GCL
HIPPO
TT
TC
CC
TT
TC
CC
rs112765090

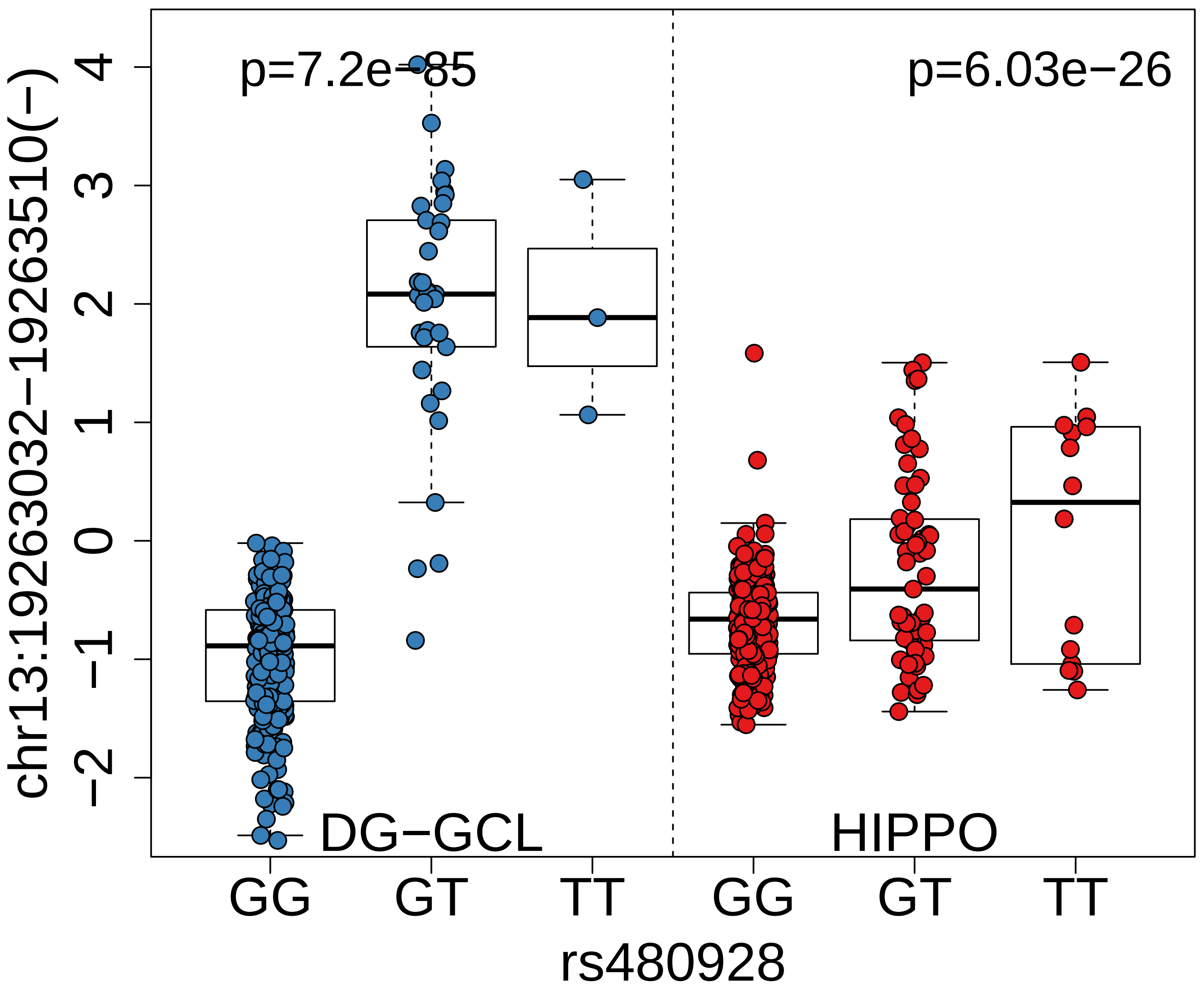
p=7.2e−85
p=6.03e−26
DG−GCL
HIPPO
chr13:19263032−19263510(−)
4
3
2
1
0
−1
−2
GG
GT
TT
GG
GT
TT
rs480928

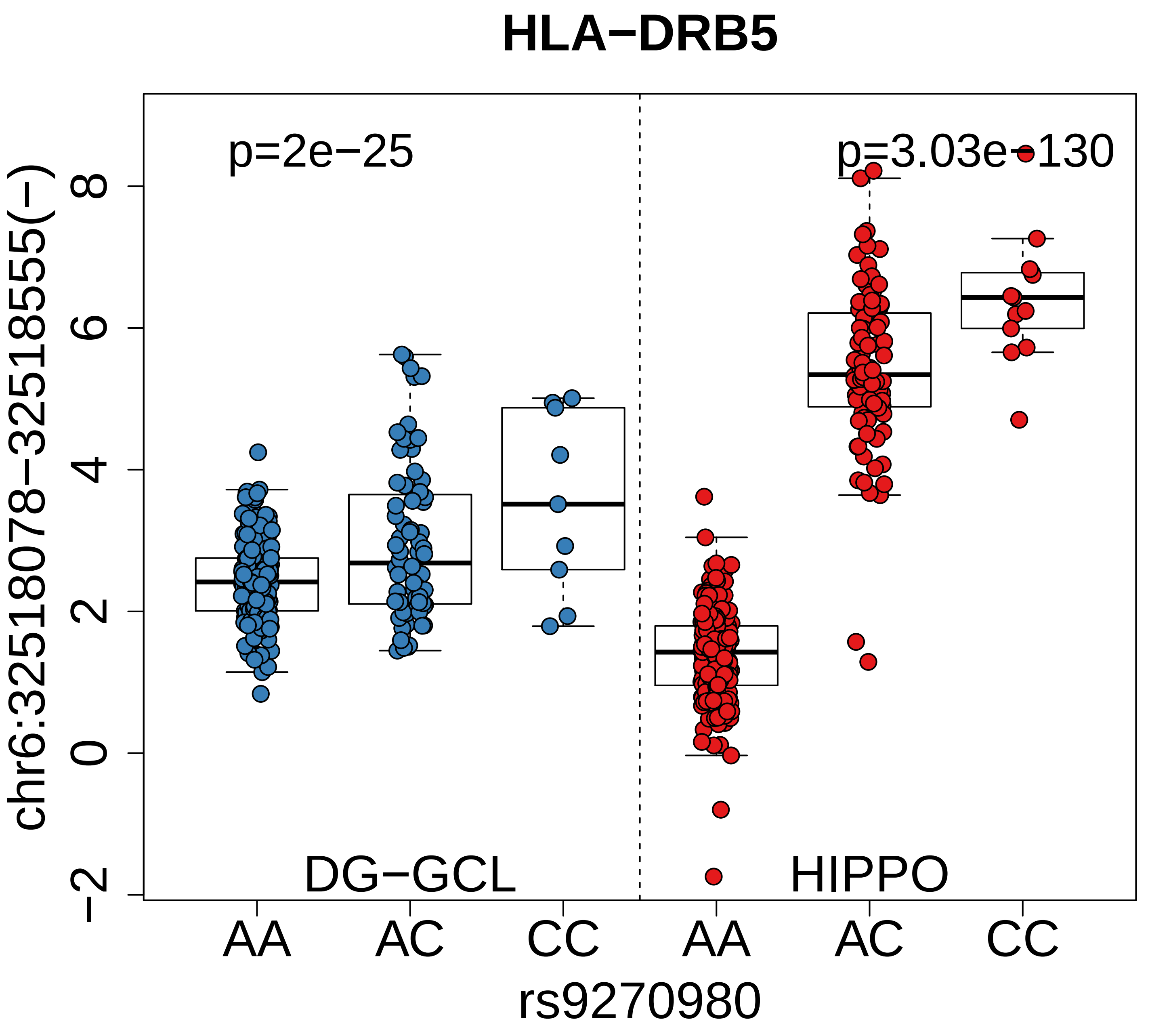

HLA−DRB5
p=2e−25
p=3.03e−130
chr6:32518078−32518555(−)
8
6
4
2
0
−2
DG−GCL
HIPPO
AA
AC
CC
AA
AC
CC
rs9270980

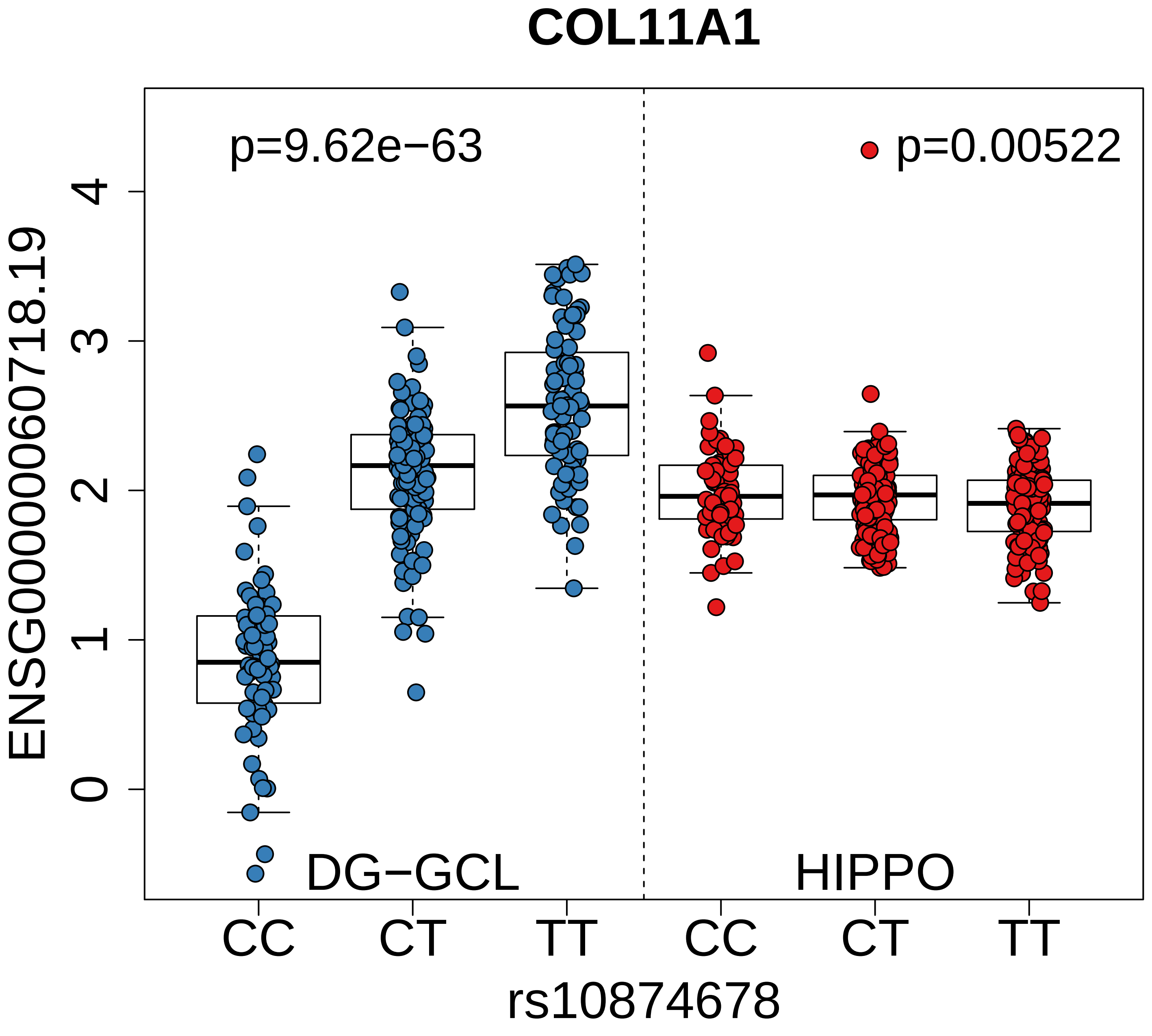

COL11A1
p=9.62e−63
p=0.00522
ENSG00000060718.19
4
3
2
1
0
DG−GCL
HIPPO
CC
CT
TT
CC
CT
TT
rs10874678

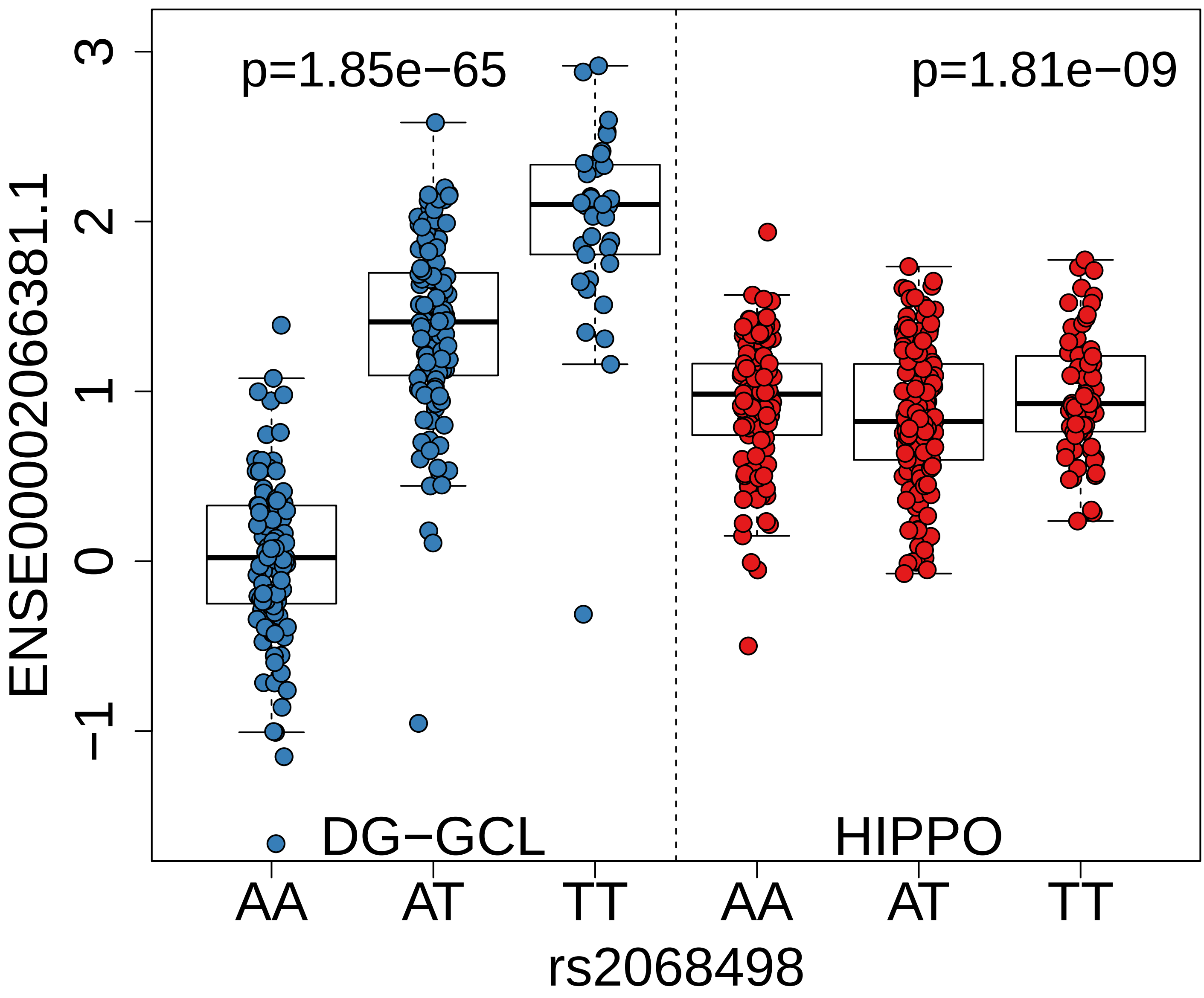
p=1.85e−65
p=1.81e−09
DG−GCL
HIPPO
ENSE00002066381.1
3
2
1
0
−1
AA
AT
TT
AA
AT
TT
rs2068498

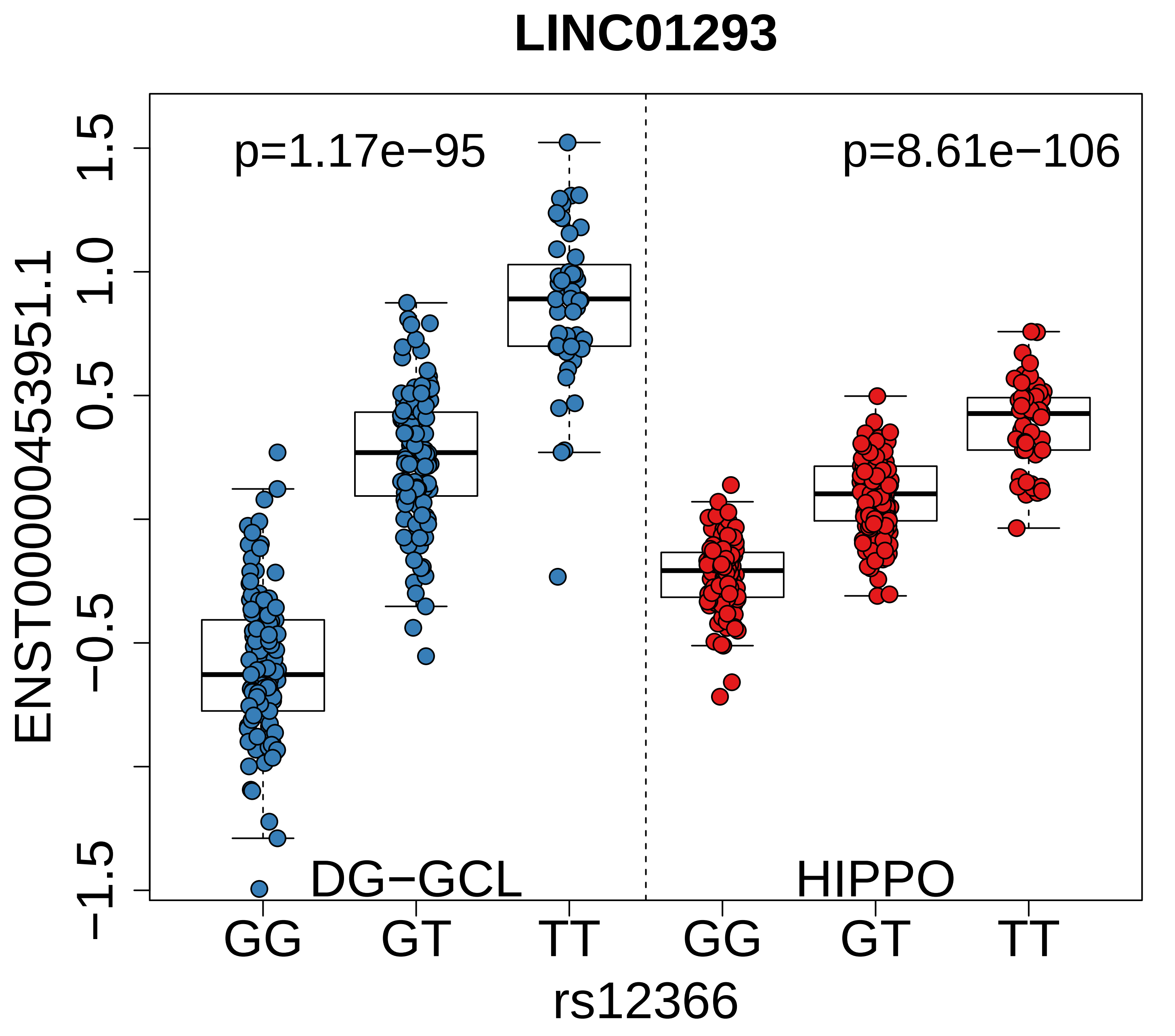
LINC01293
p=1.17e−95
p=8.61e−106
ENST00000453951.1
1.5
1.0
0.5
−0.5
−1.5
DG−GCL
HIPPO
GG
GT
TT
GG
GT
TT
rs12366

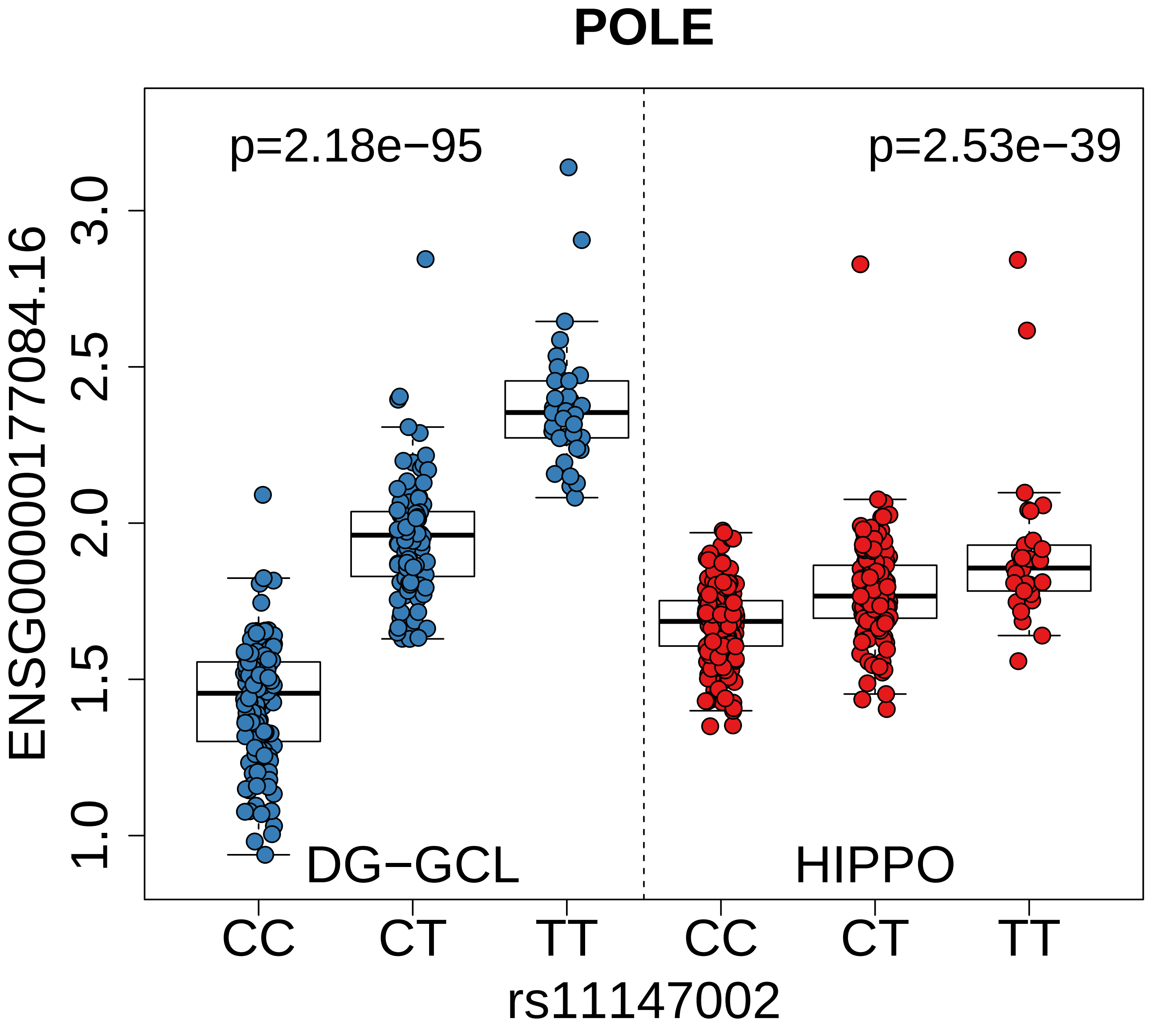
POLE
p=2.18e−95
p=2.53e−39
ENSG00000177084.16
3.0
2.5
2.0
1.5
1.0
DG−GCL
HIPPO
CC
CT
TT
CC
CT
TT
rs11147002

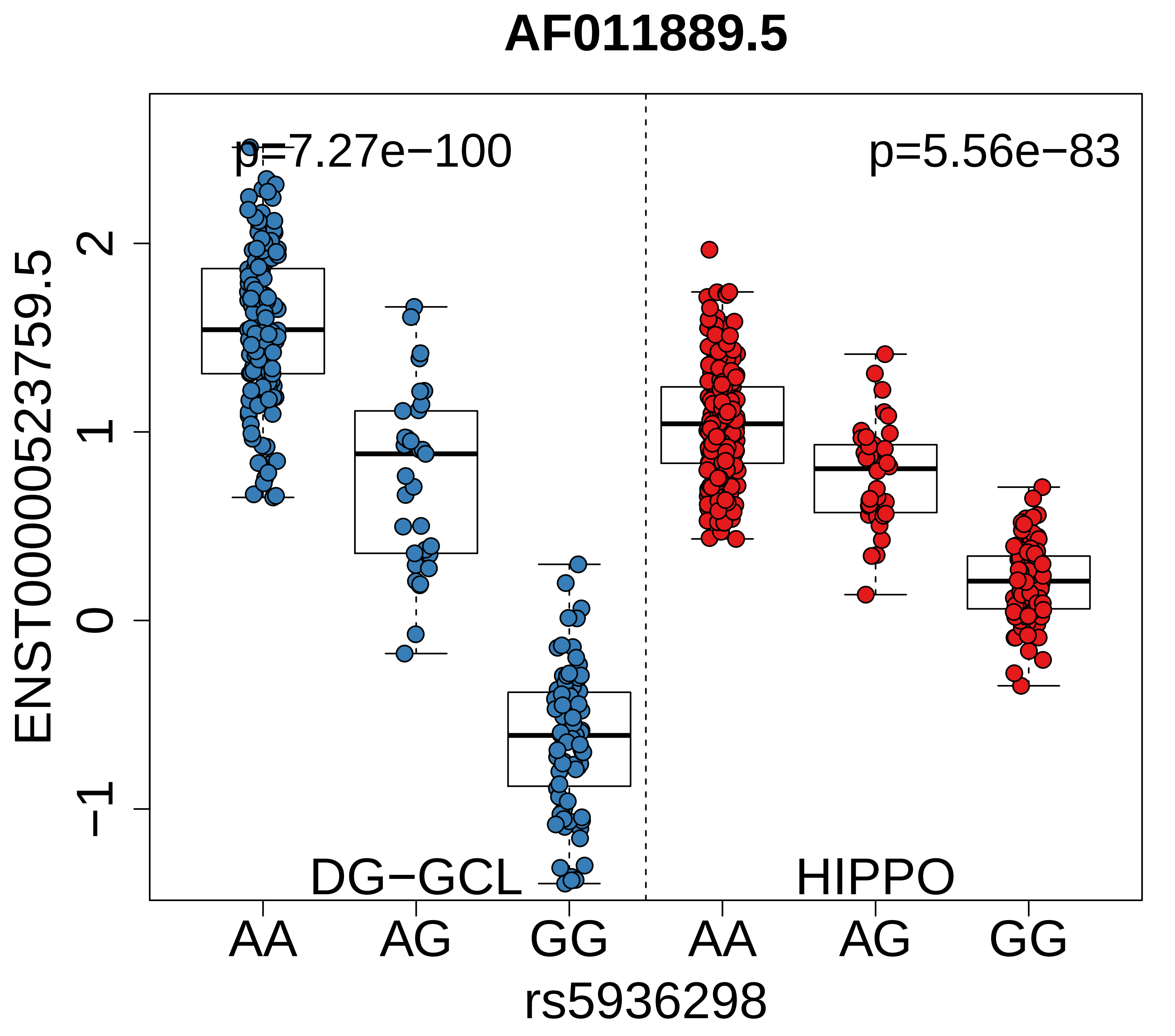

AF011889.5
p=7.27e−100
p=5.56e−83
ENST00000523759.5
2
1
0
−1
DG−GCL
HIPPO
AA
AG
GG
AA
AG
GG
rs5936298

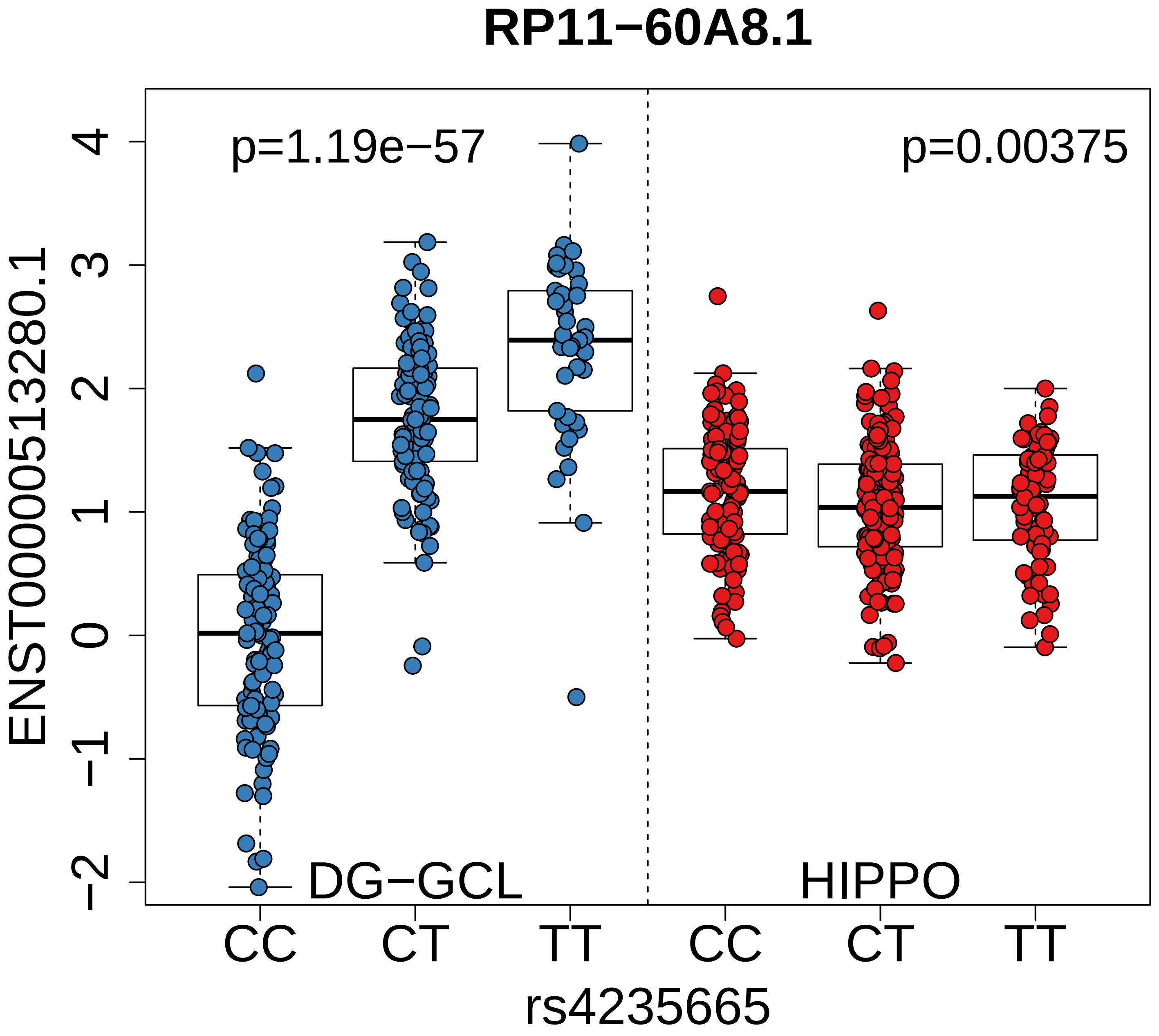

RP11−60A8.1
p=1.19e−57
p=0.00375
ENST00000513280.1
4
3
2
1
0
−1
−2
DG−GCL
HIPPO
CC
CT
TT
CC
CT
TT
rs4235665

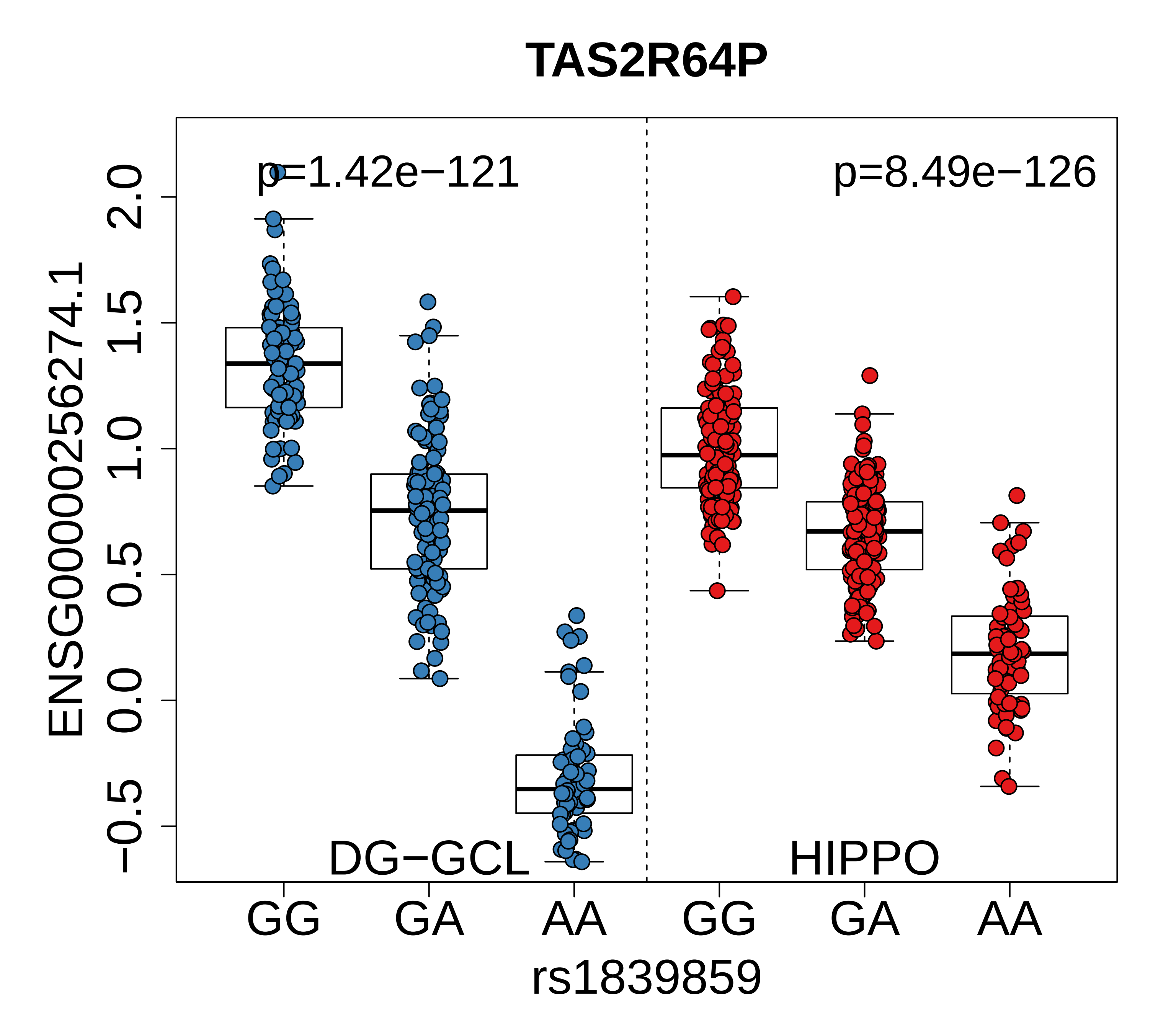

TAS2R64P
p=1.42e−121
p=8.49e−126
ENSG00000256274.1
2.0
1.5
1.0
0.5
0.0
−0.5
DG−GCL
HIPPO
GG
GA
AA
GG
GA
AA
rs1839859

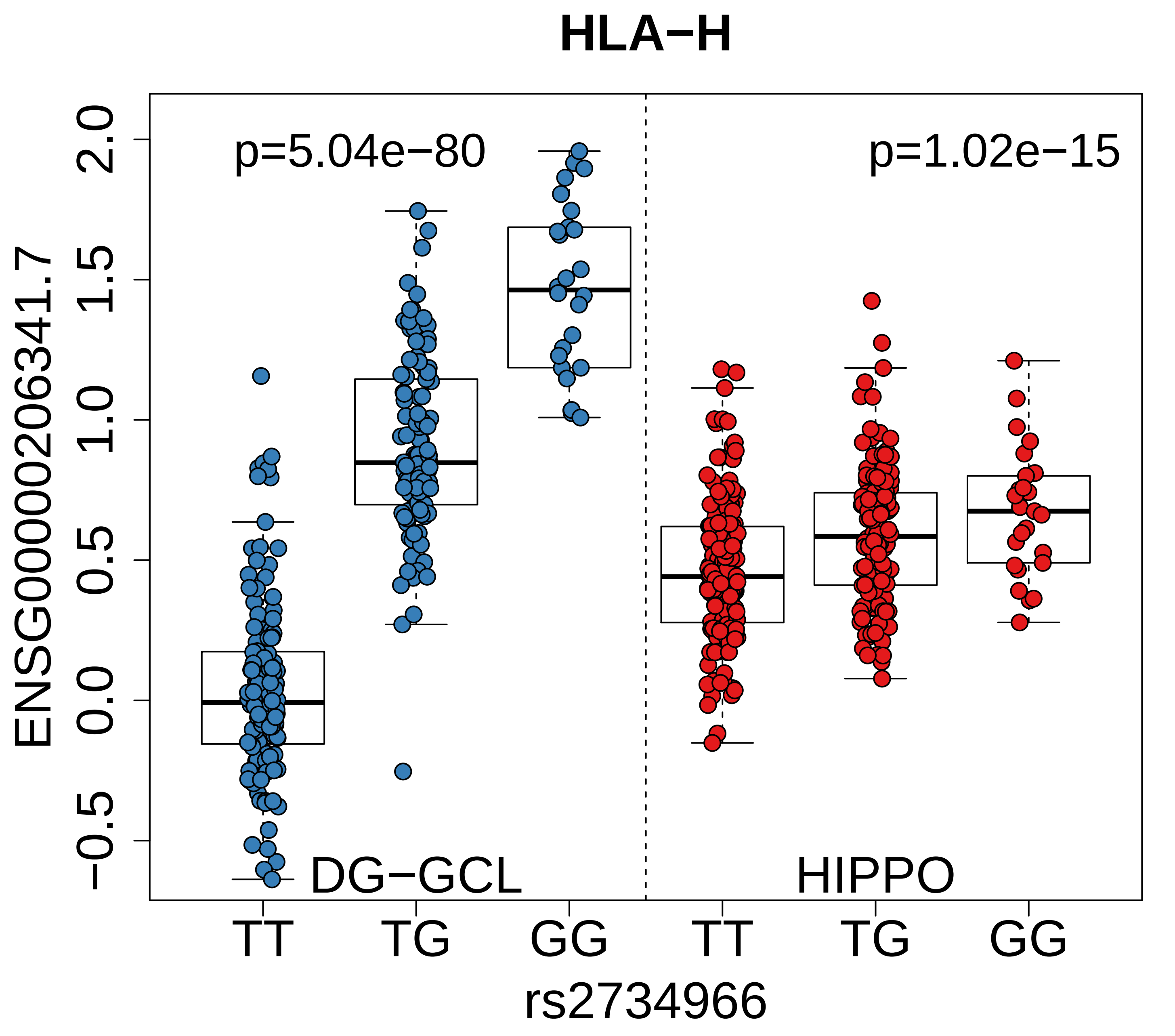
HLA−H
p=5.04e−80
p=1.02e−15
ENSG00000206341.7
2.0
1.5
1.0
0.5
0.0
−0.5
DG−GCL
HIPPO
TT
TG
GG
TT
TG
GG
rs2734966

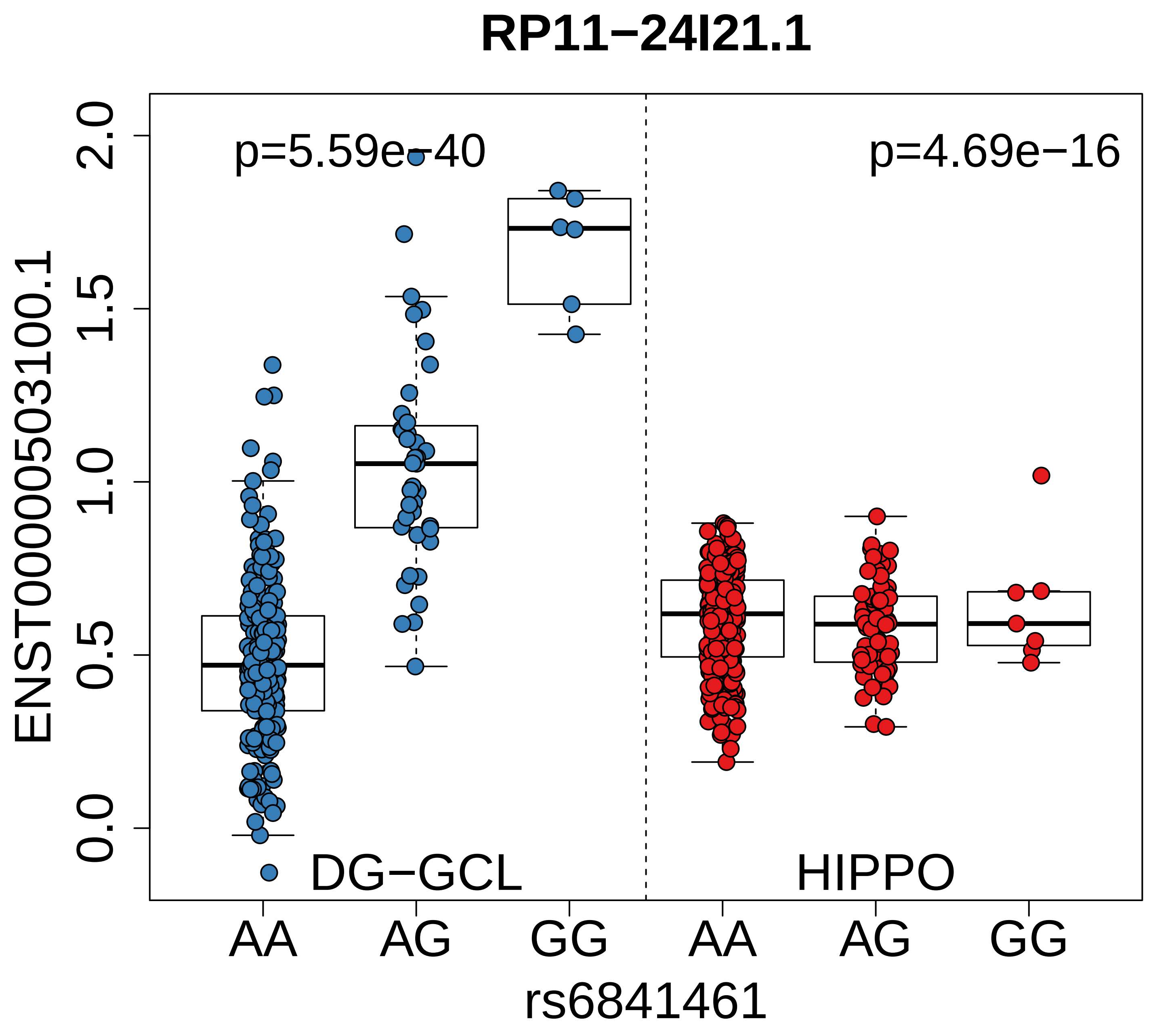
RP11−24I21.1
p=5.59e−40
p=4.69e−16
2.0
1.5
1.0
0.5
0.0
ENST00000503100.1
DG−GCL
HIPPO
AA
AG
GG
AA
AG
GG
rs6841461

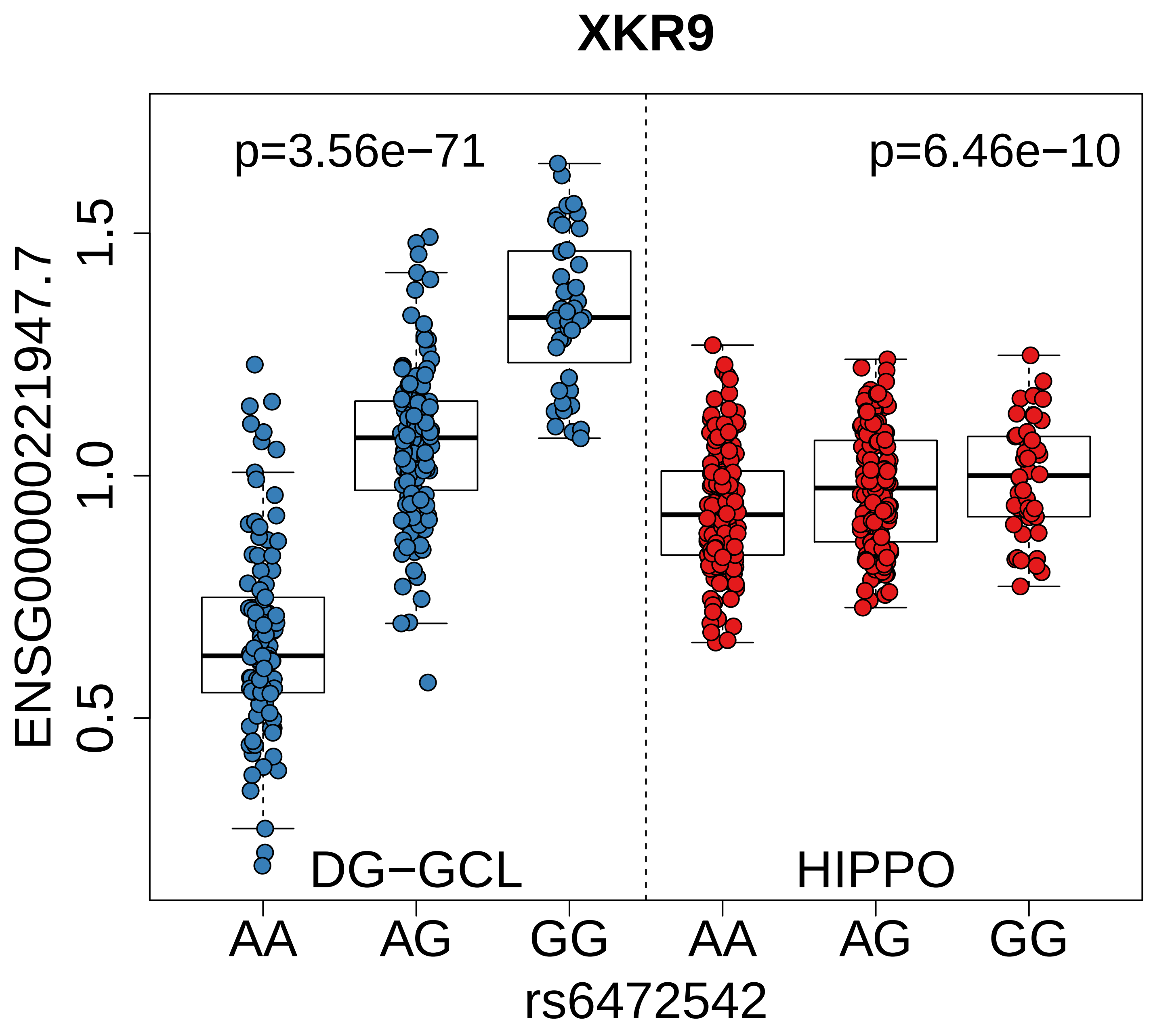

XKR9
p=3.56e−71
p=6.46e−10
ENSG00000221947.7
1.5
1.0
0.5
DG−GCL
HIPPO
AA
AG
GG
AA
AG
GG
rs6472542

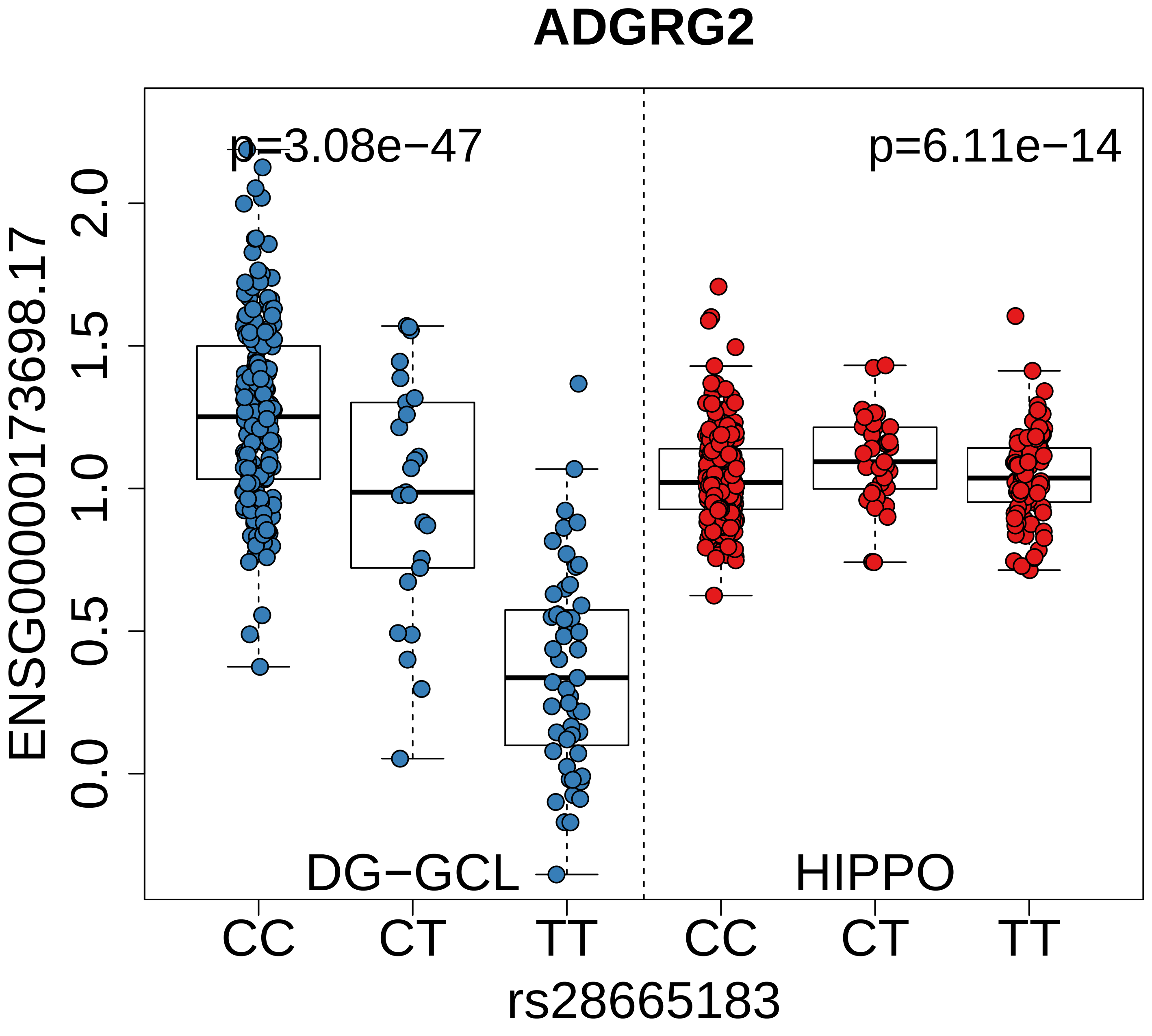

ADGRG2
p=3.08e−47
p=6.11e−14
ENSG00000173698.17
0.0
0.5
1.0
1.5
2.0
DG−GCL
HIPPO
CC
CT
TT
CC
CT
TT
rs28665183

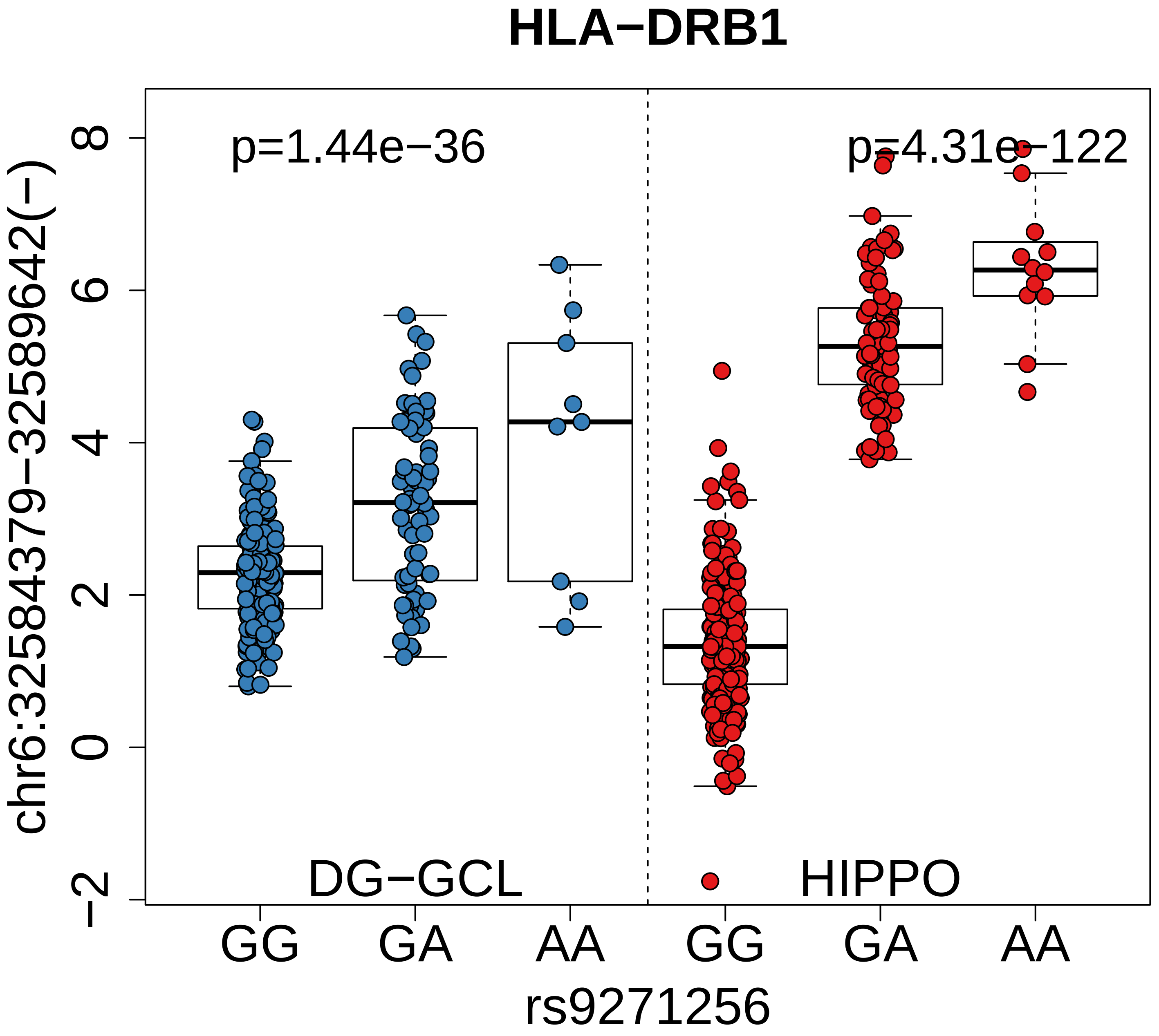

HLA−DRB1
p=1.44e−36
p=4.31e−122
chr6:32584379−32589642(−)
8
6
4
2
0
−2
DG−GCL
HIPPO
GG
GA
AA
GG
GA
AA
rs9271256

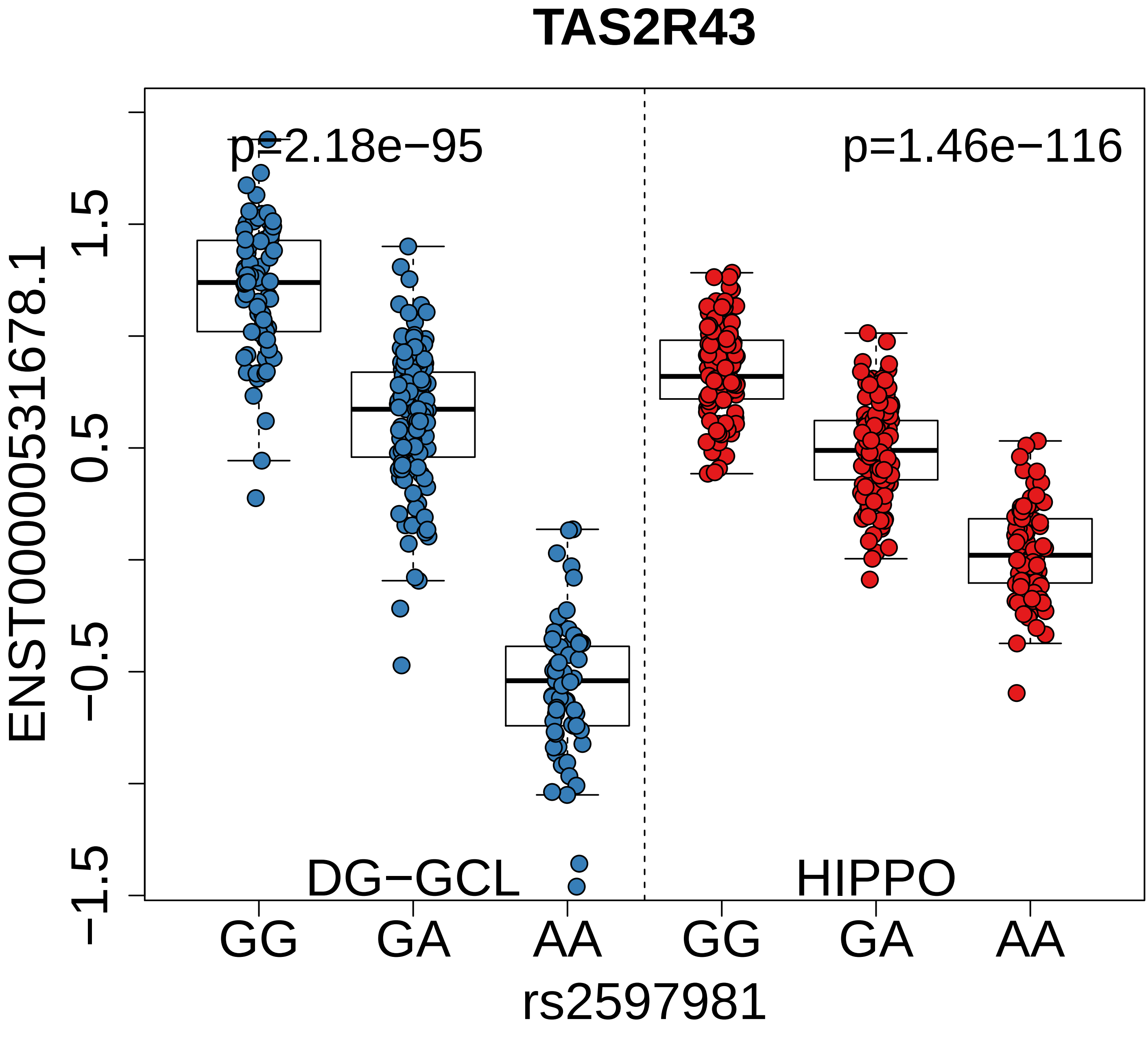
TAS2R43
p=2.18e−95
p=1.46e−116
ENST00000531678.1
1.5
0.5
−0.5
−1.5
DG−GCL
HIPPO
GG
GA
AA
GG
GA
AA
rs2597981

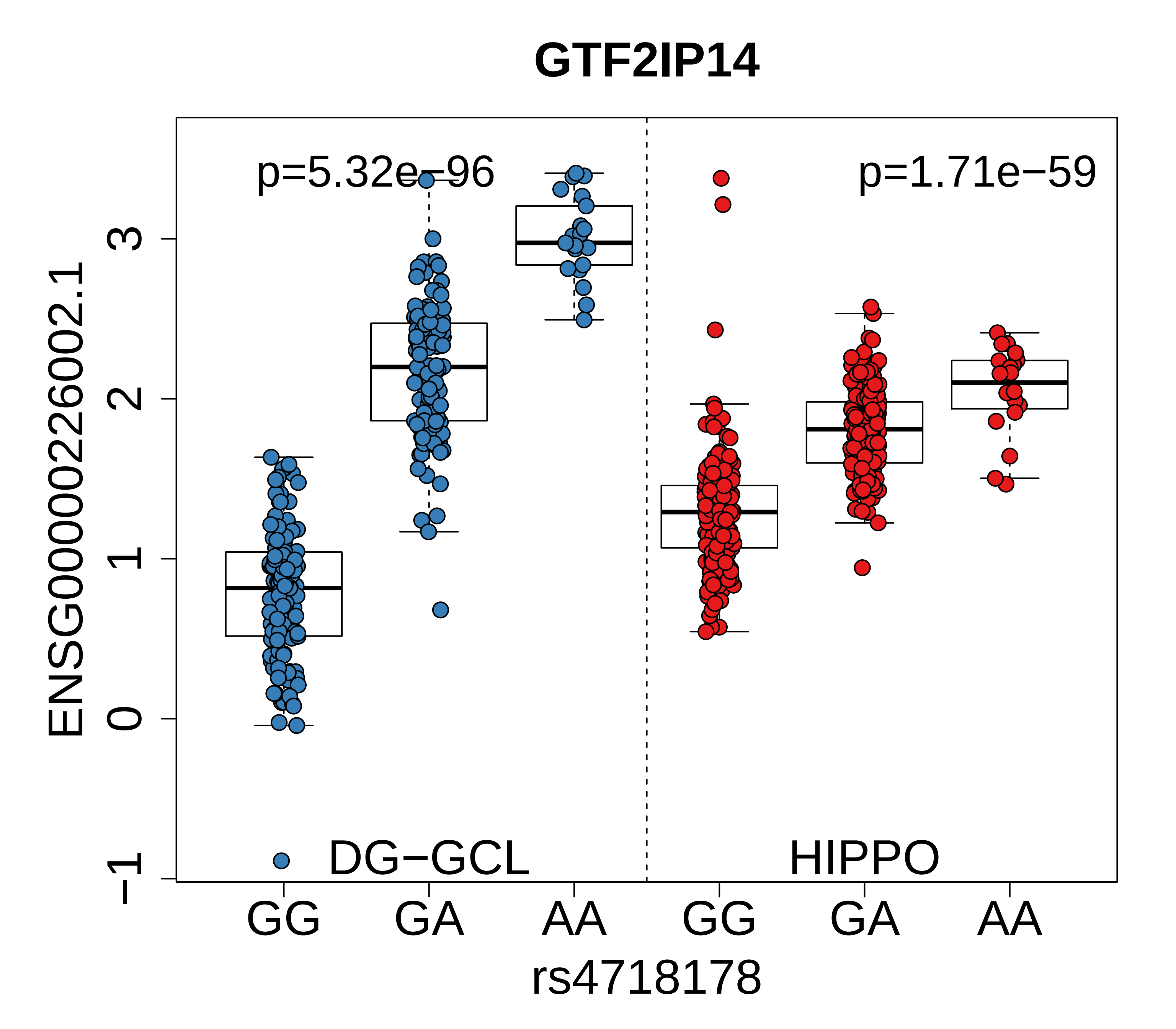

GTF2IP14
p=5.32e−96
p=1.71e−59
ENSG00000226002.1
3
2
1
0
−1
DG−GCL
HIPPO
GG
GA
AA
GG
GA
AA
rs4718178

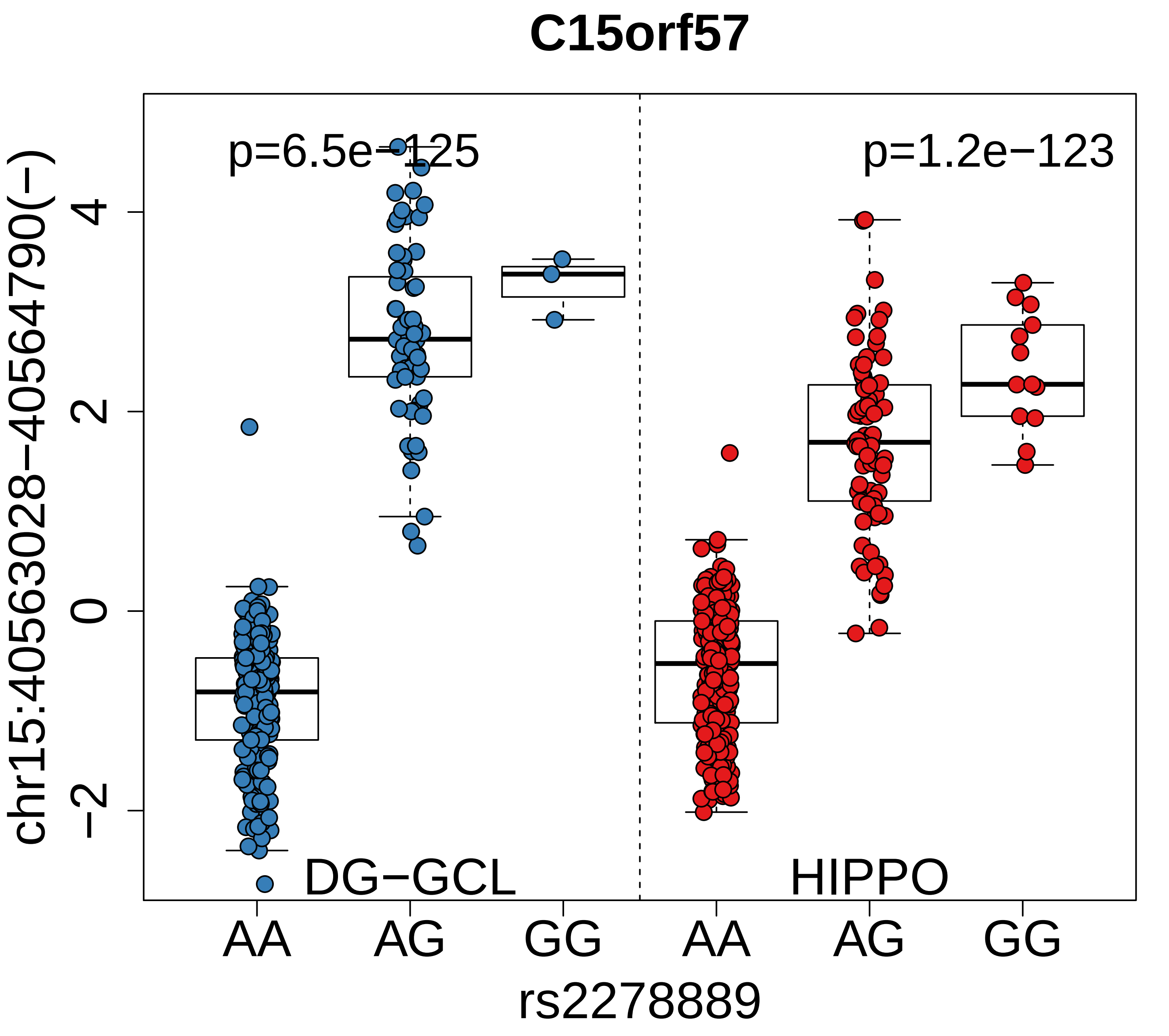
C15orf57
p=6.5e−125
p=1.2e−123
chr15:40563028−40564790(−)
4
2
0
−2
DG−GCL
HIPPO
AA
AG
GG
AA
AG
GG
rs2278889

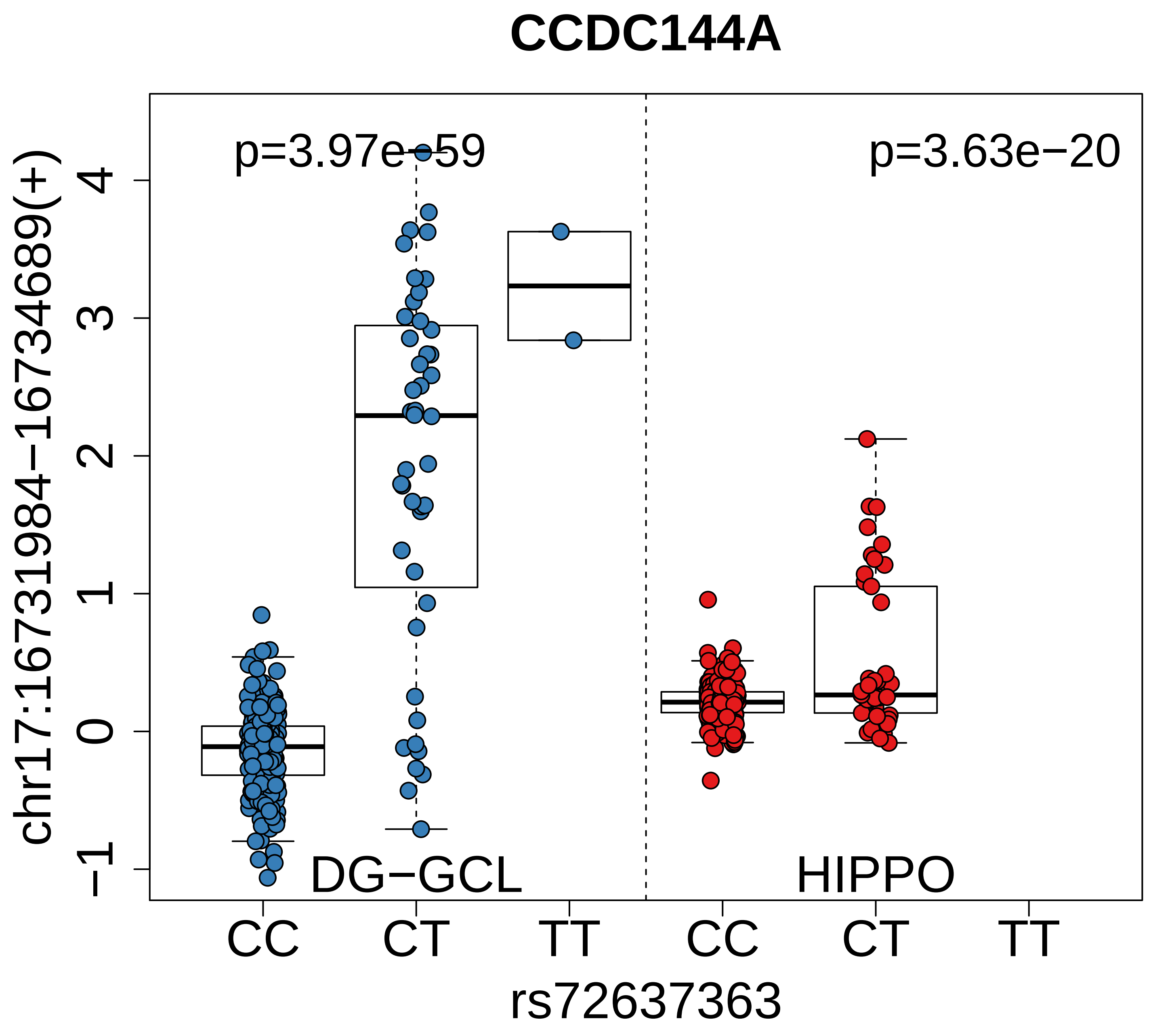

CCDC144A
p=3.97e−59
p=3.63e−20
chr17:16731984−16734689(+)
4
3
2
1
0
−1
DG−GCL
HIPPO
CC
CT
TT
CC
CT
TT
rs72637363

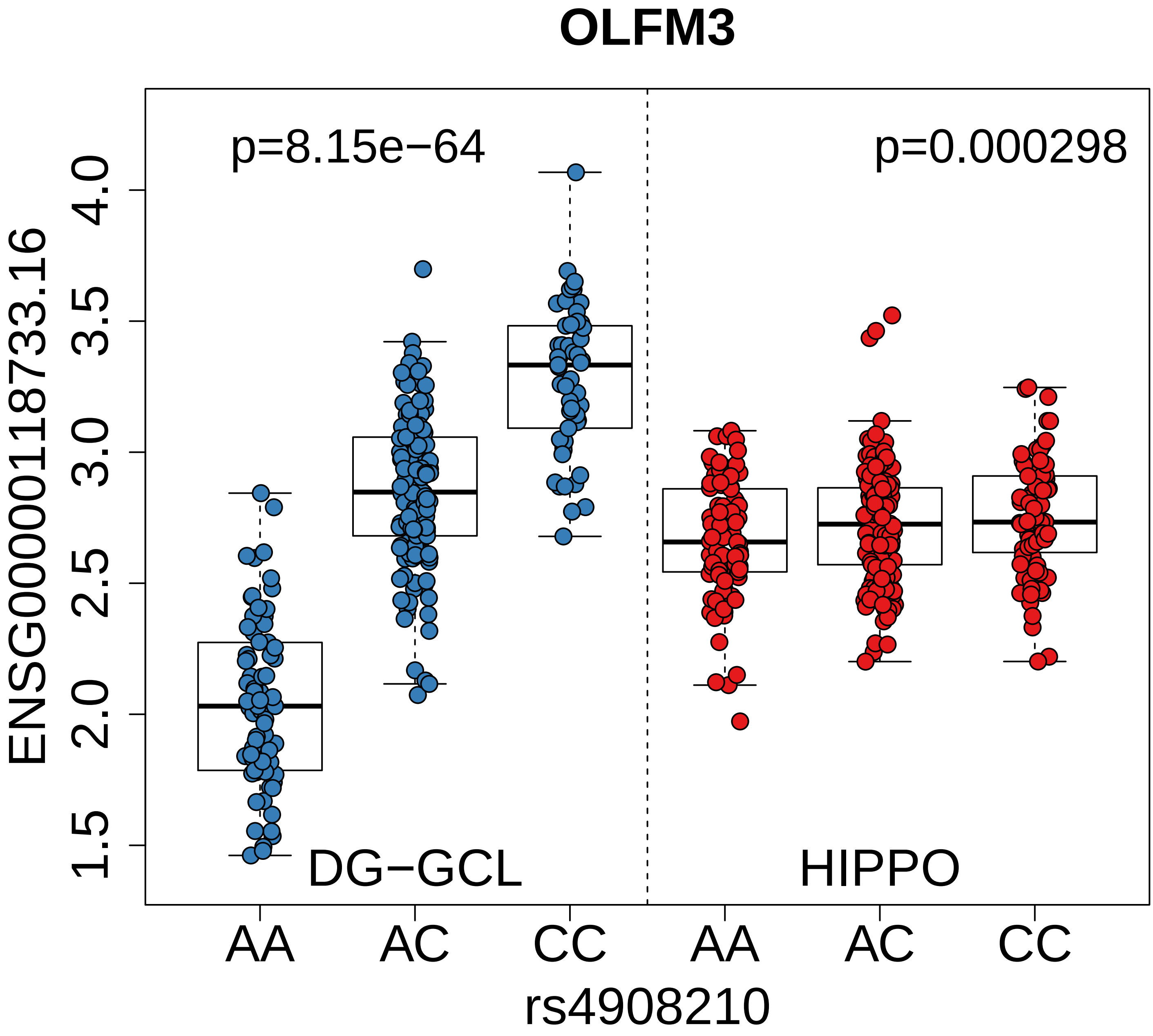

OLFM3
p=8.15e−64
p=0.000298
ENSG00000118733.16
4.0
3.5
3.0
2.5
2.0
1.5
DG−GCL
HIPPO
AA
AC
CC
AA
AC
CC
rs4908210

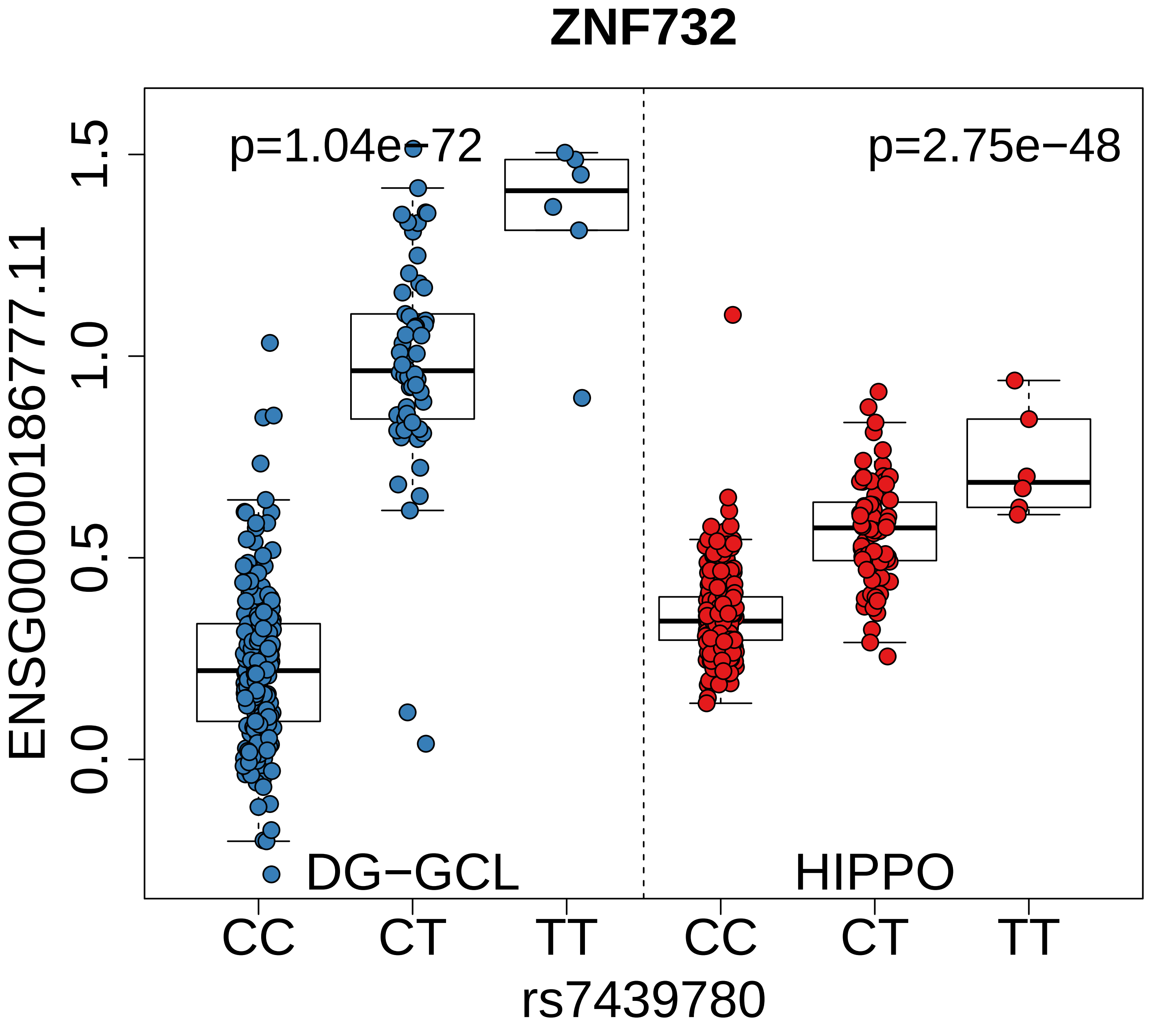

ZNF732
p=1.04e−72
p=2.75e−48
ENSG00000186777.11
1.5
1.0
0.5
0.0
DG−GCL
HIPPO
CC
CT
TT
CC
CT
TT
rs7439780

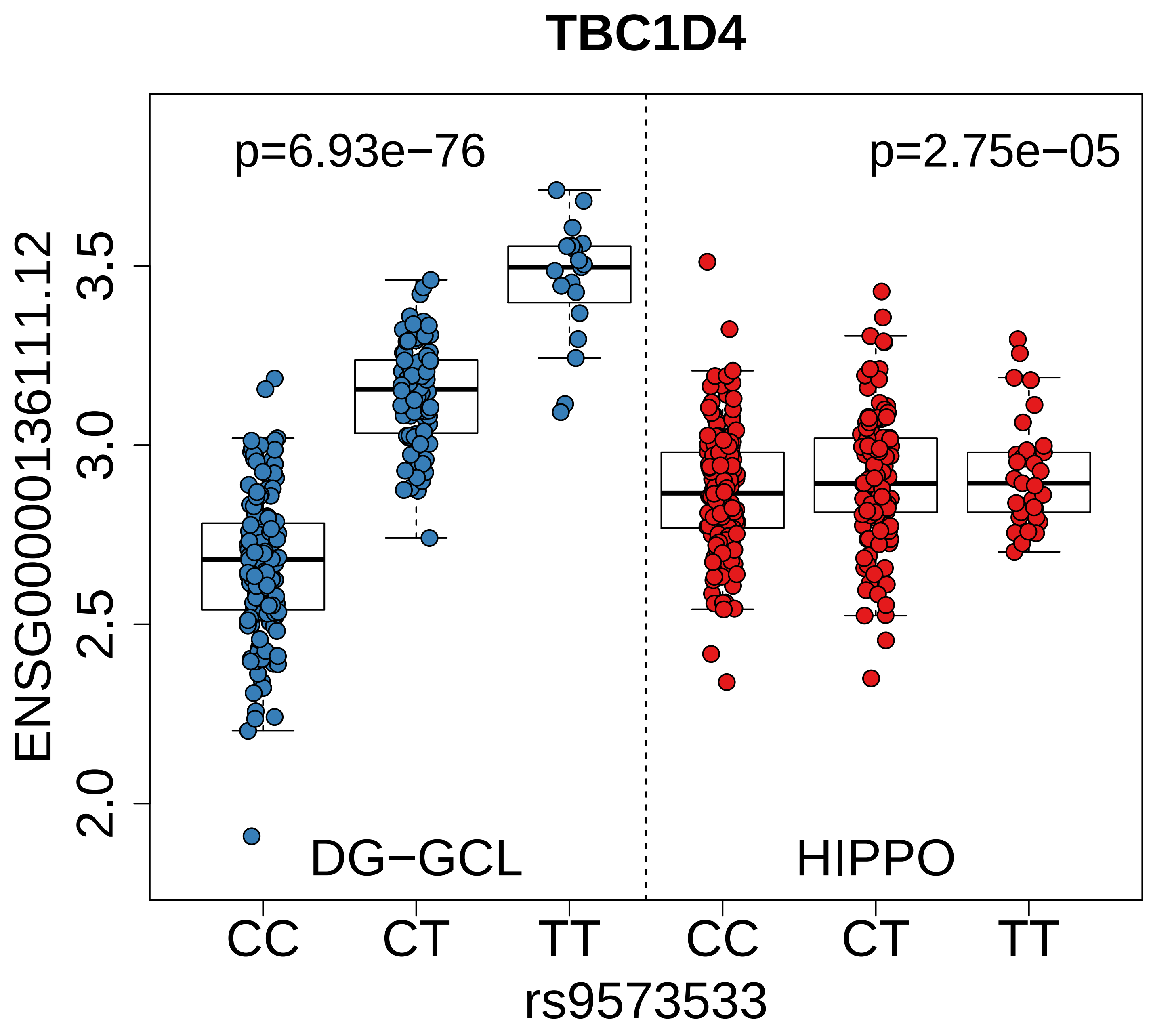

TBC1D4
p=6.93e−76
p=2.75e−05
ENSG00000136111.12
3.5
3.0
2.5
2.0
DG−GCL
HIPPO
CC
CT
TT
CC
CT
TT
rs9573533

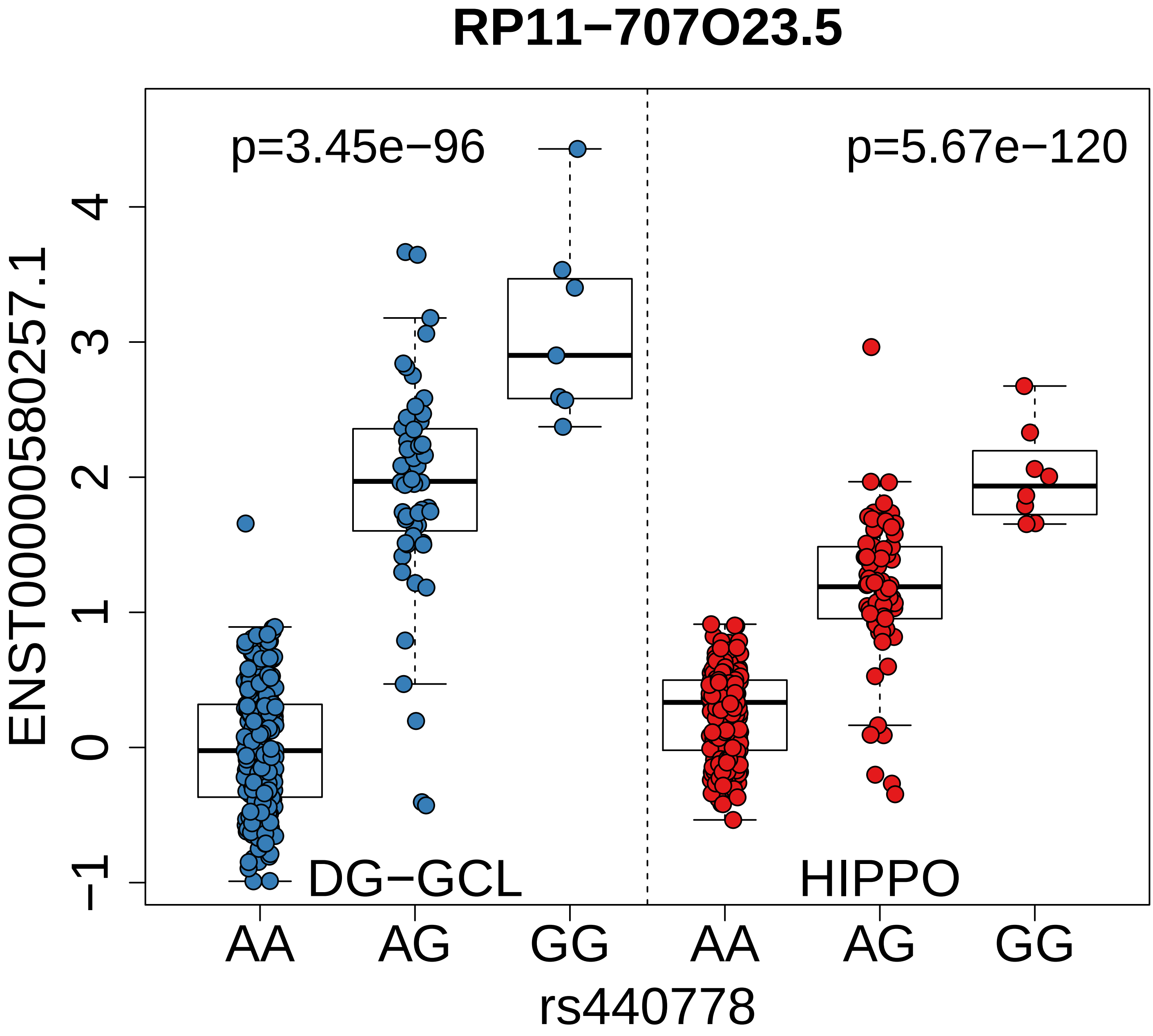
RP11−707O23.5
p=3.45e−96
p=5.67e−120
ENST00000580257.1
4
3
2
1
0
−1
DG−GCL
HIPPO
AA
AG
GG
AA
AG
GG
rs440778

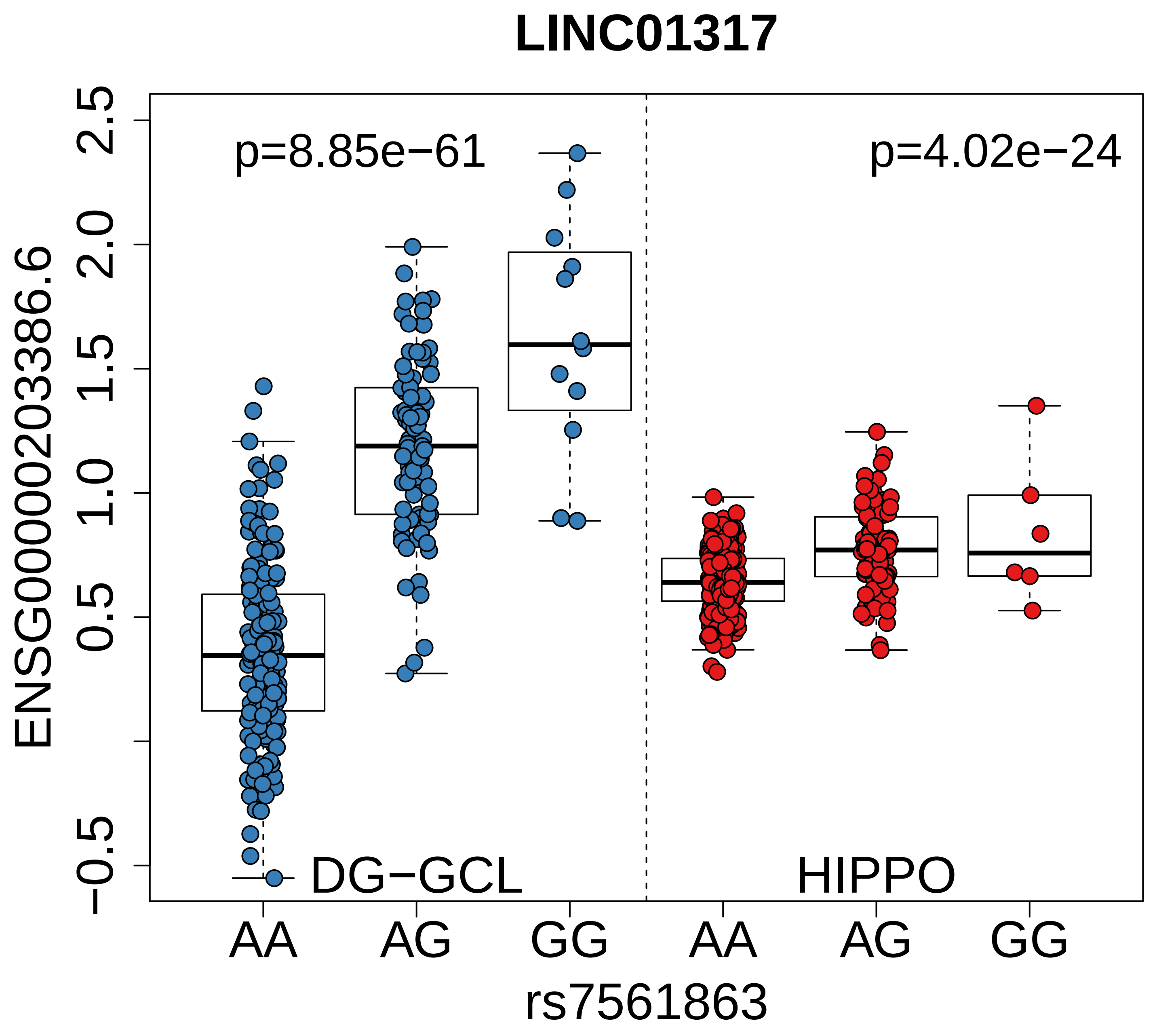

LINC01317
p=8.85e−61
p=4.02e−24
DG−GCL
HIPPO
ENSG00000203386.6
2.5
2.0
1.5
1.0
0.5
−0.5
AA
AG
GG
AA
AG
GG
rs7561863

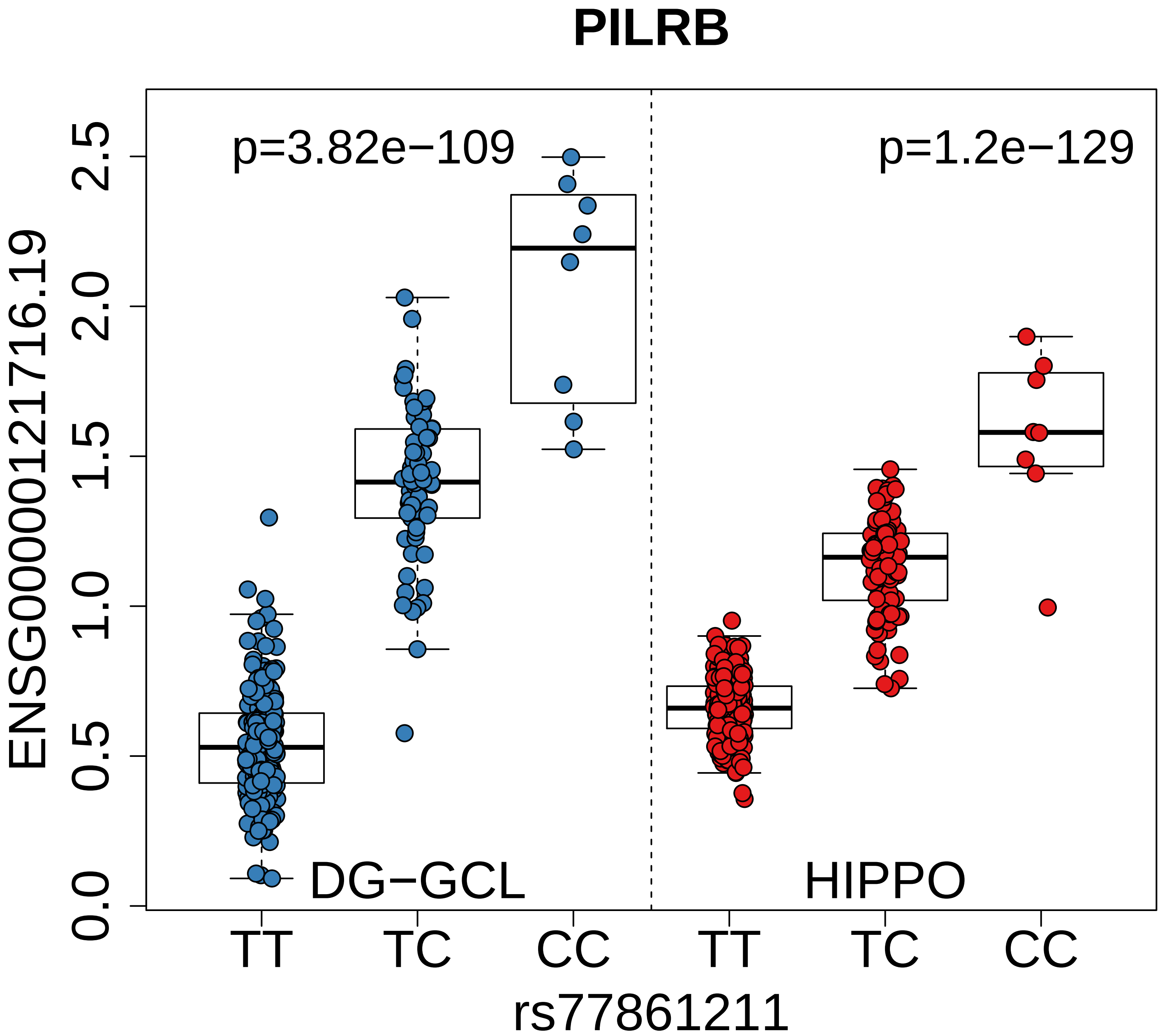

PILRB
p=3.82e−109
p=1.2e−129
ENSG00000121716.19
0.0
0.5
1.0
1.5
2.0
2.5
DG−GCL
HIPPO
TT
TC
CC
TT
TC
CC
rs77861211

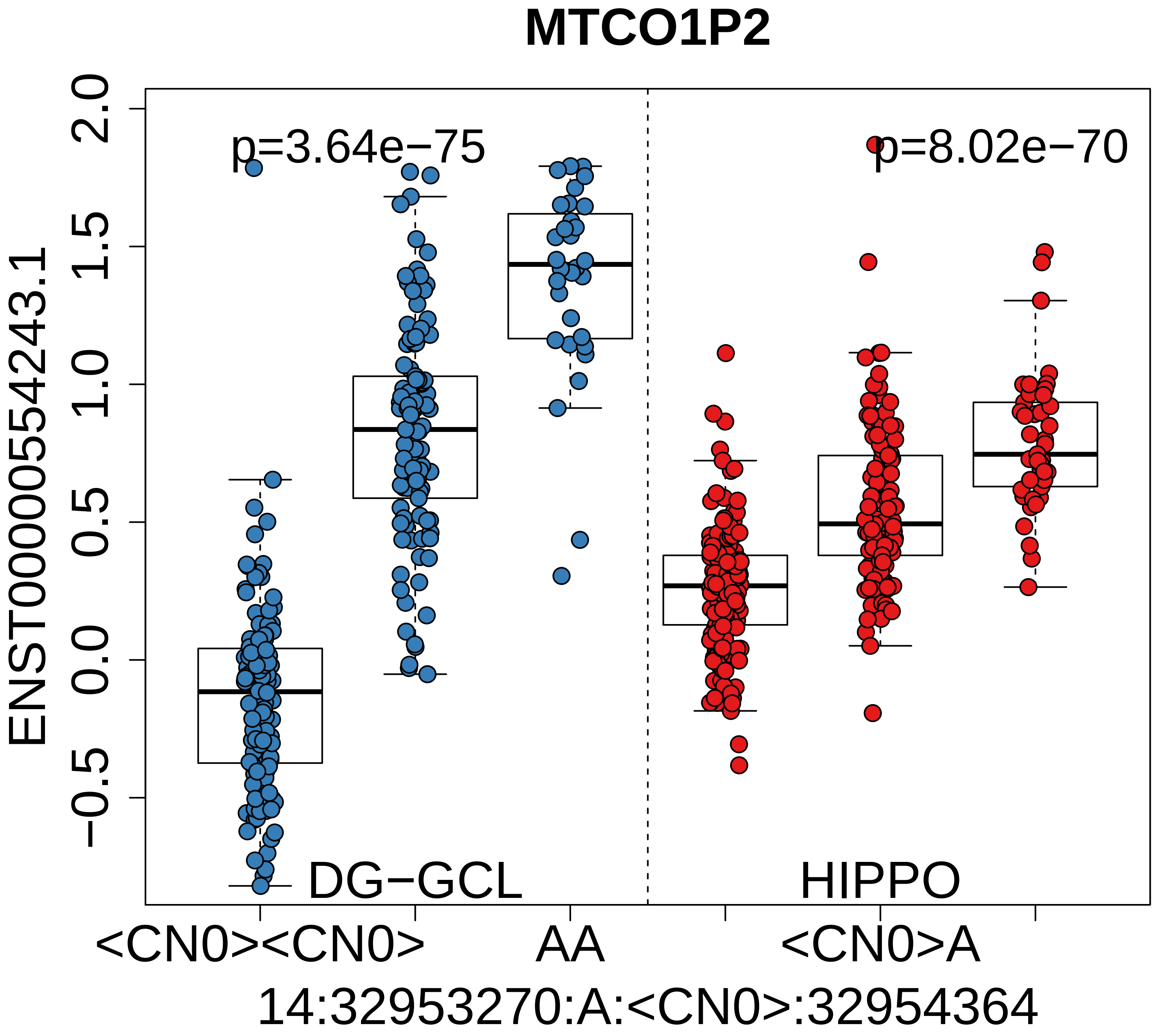

MTCO1P2
p=3.64e−75
p=8.02e−70
DG−GCL
HIPPO
ENST00000554243.1
−0.5
0.0
0.5
1.0
1.5
2.0
<CN0><CN0>
AA
<CN0>A
14:32953270:A:<CN0>:32954364

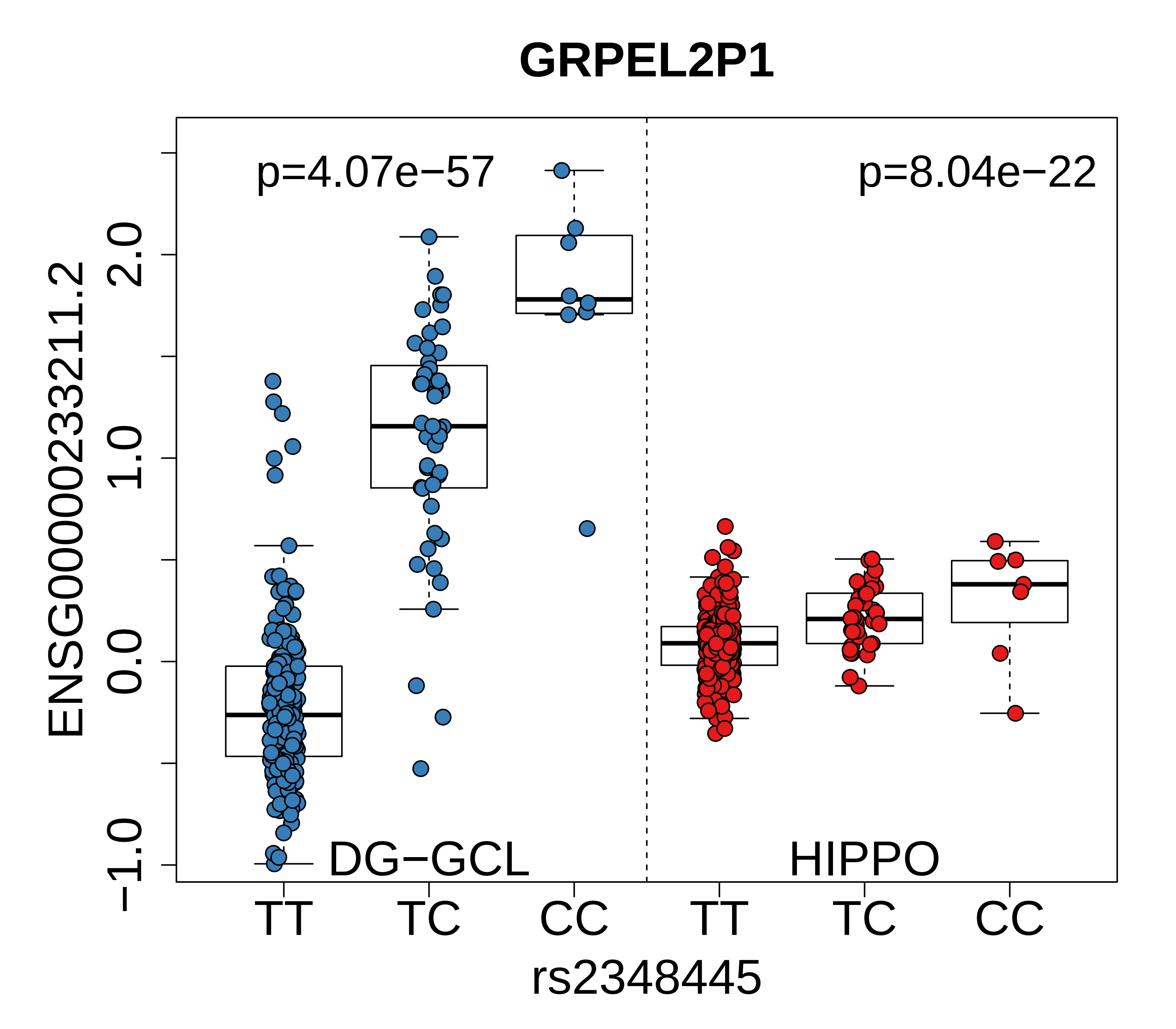
GRPEL2P1
p=4.07e−57
p=8.04e−22
ENSG00000233211.2
2.0
1.0
0.0
−1.0
DG−GCL
HIPPO
TT
TC
CC
TT
TC
CC
rs2348445

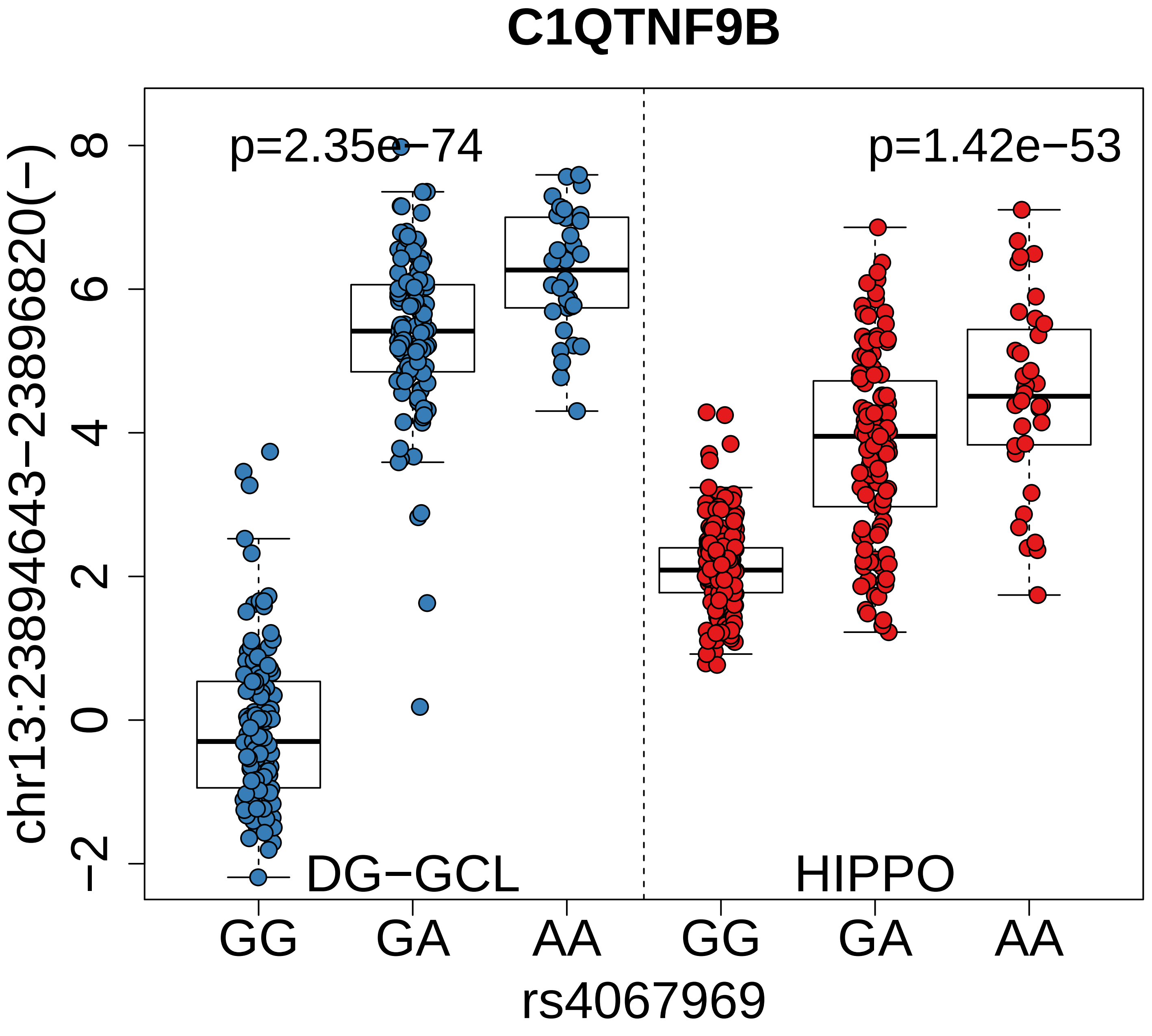
C1QTNF9B
p=2.35e−74
p=1.42e−53
DG−GCL
HIPPO
chr13:23894643−23896820(−)
8
6
4
2
0
−2
GG
GA
AA
GG
GA
AA
rs4067969

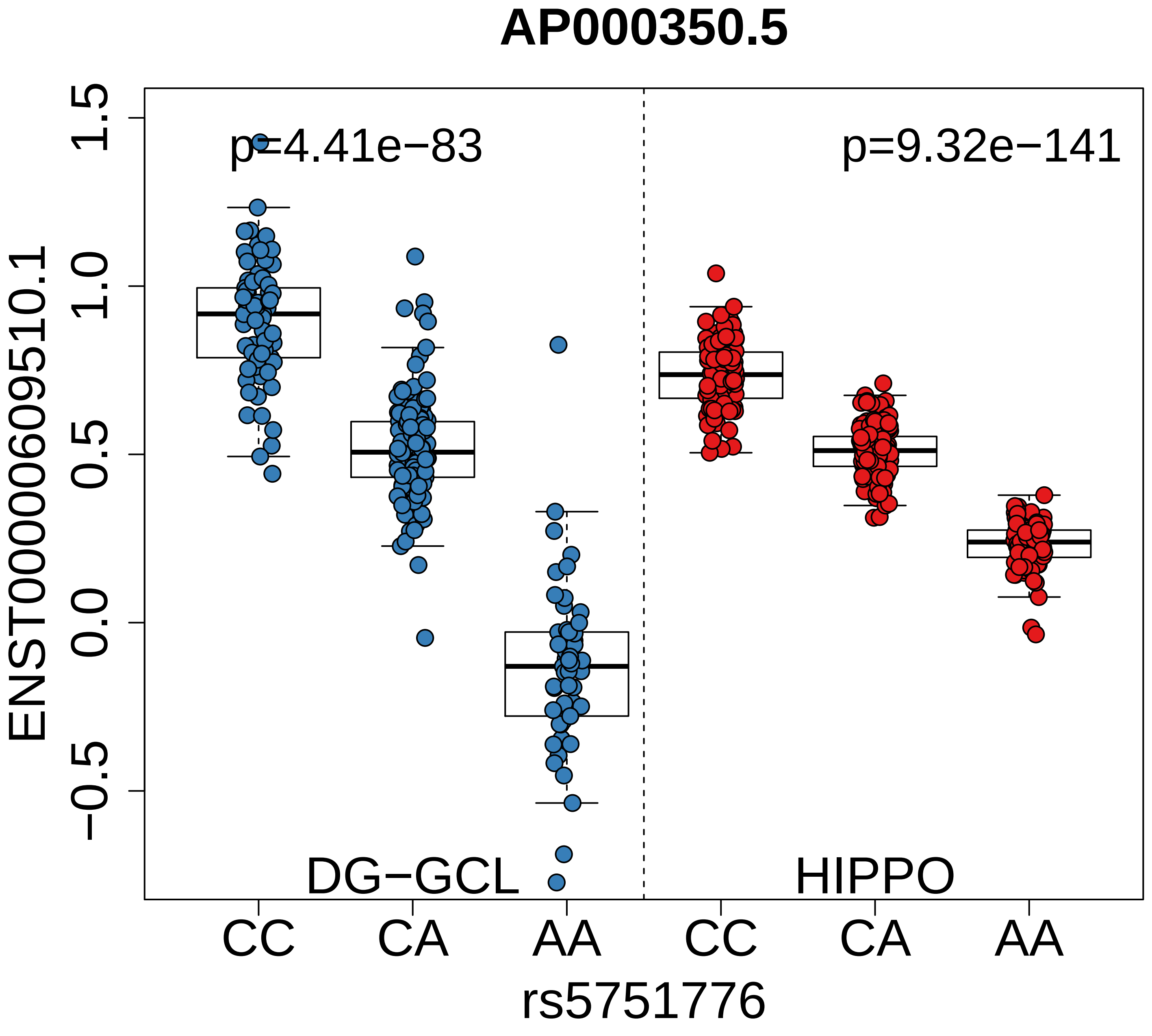

AP000350.5
p=4.41e−83
p=9.32e−141
ENST00000609510.1
1.5
1.0
0.5
0.0
−0.5
DG−GCL
HIPPO
CC
CA
AA
CC
CA
AA
rs5751776

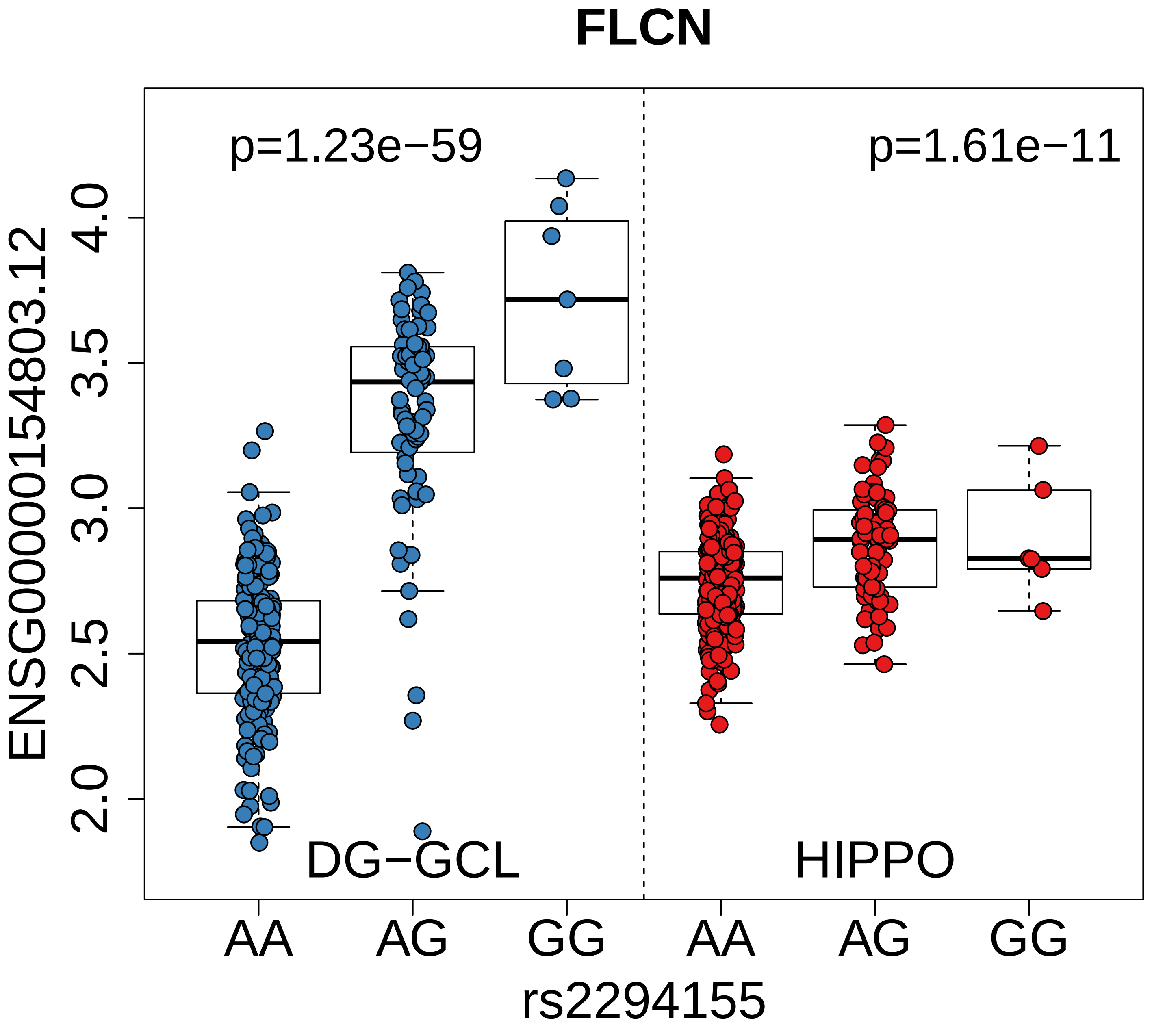
FLCN
p=1.23e−59
p=1.61e−11
ENSG00000154803.12
4.0
3.5
3.0
2.5
2.0
DG−GCL
HIPPO
AA
AG
GG
AA
AG
GG
rs2294155

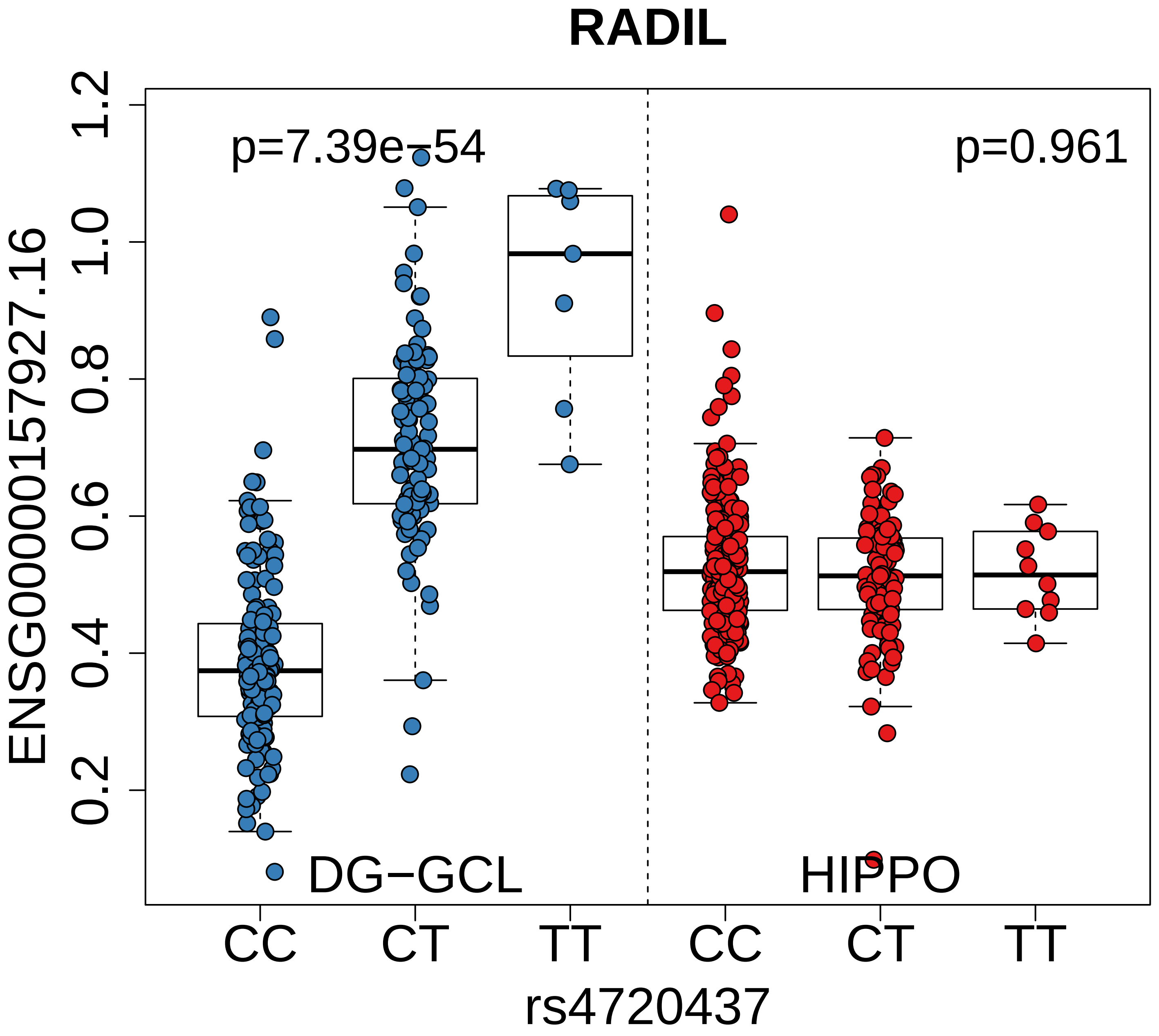

RADIL
p=7.39e−54
p=0.961
ENSG00000157927.16
1.2
1.0
0.8
0.6
0.4
0.2
DG−GCL
HIPPO
CC
CT
TT
CC
CT
TT
rs4720437

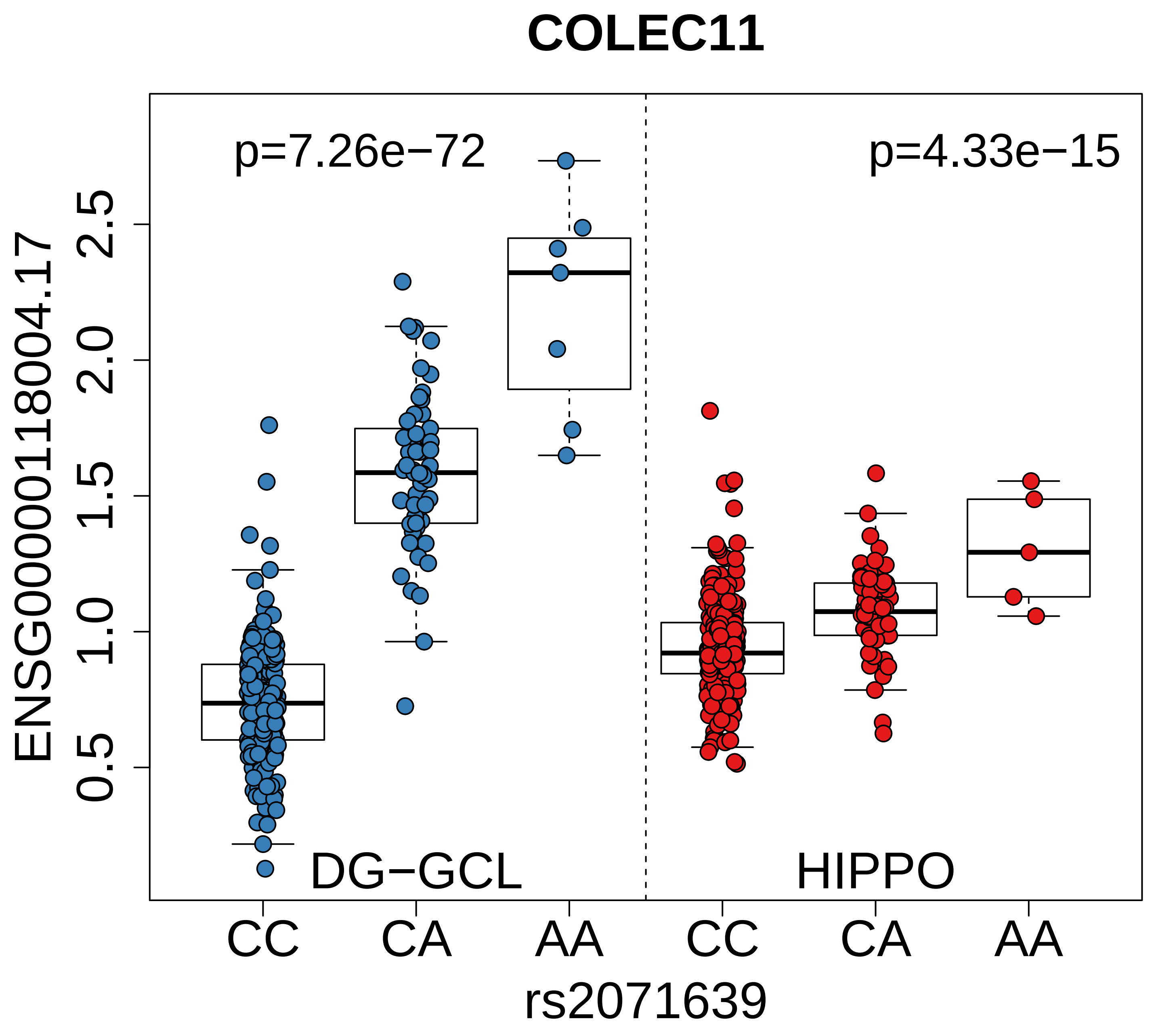

COLEC11
p=7.26e−72
p=4.33e−15
ENSG00000118004.17
0.5
1.0
1.5
2.0
2.5
DG−GCL
HIPPO
CC
CA
AA
CC
CA
AA
rs2071639

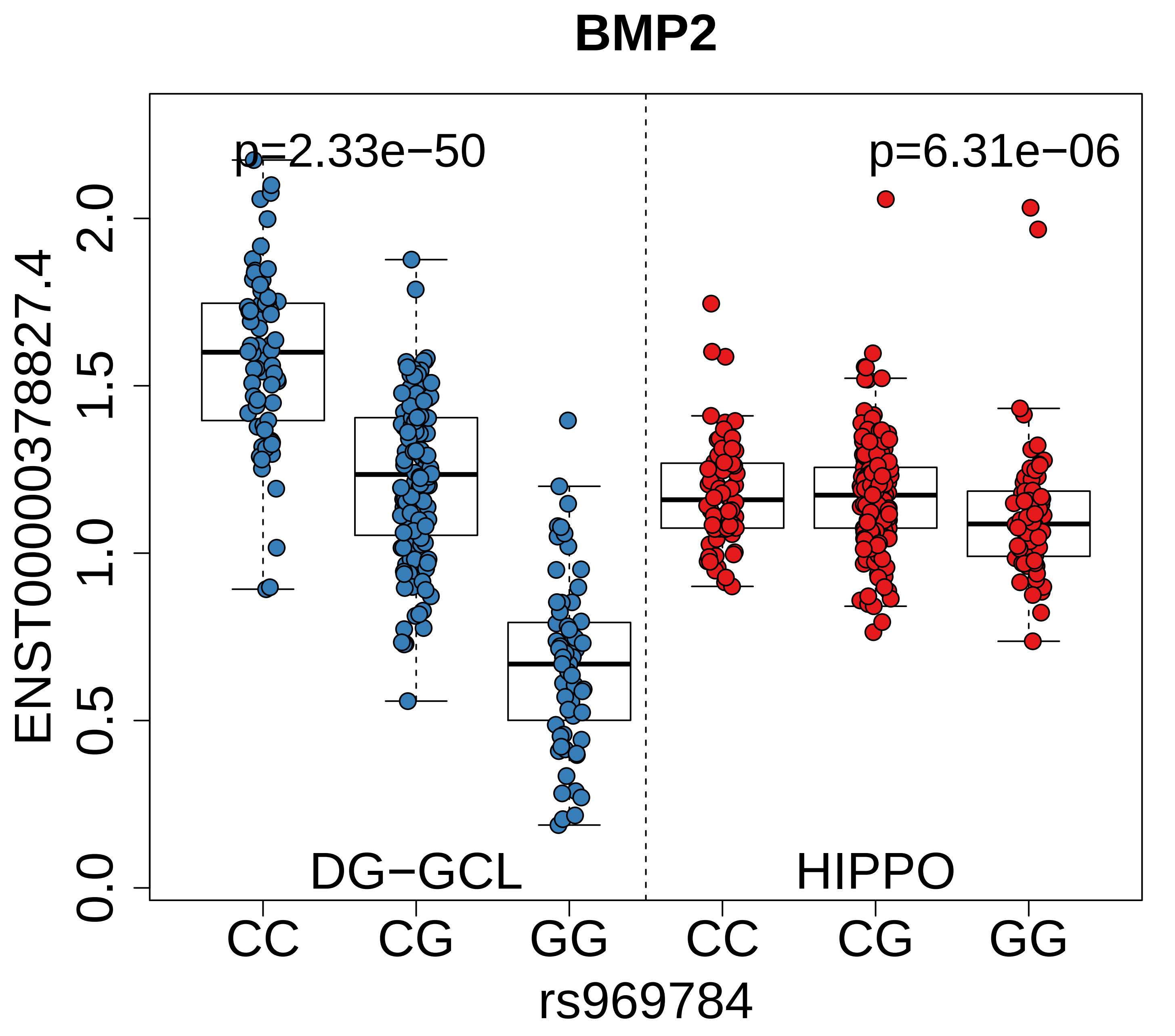
BMP2
p=2.33e−50
p=6.31e−06
ENST00000378827.4
0.0
0.5
1.0
1.5
2.0
DG−GCL
HIPPO
CC
CG
GG
CC
CG
GG
rs969784

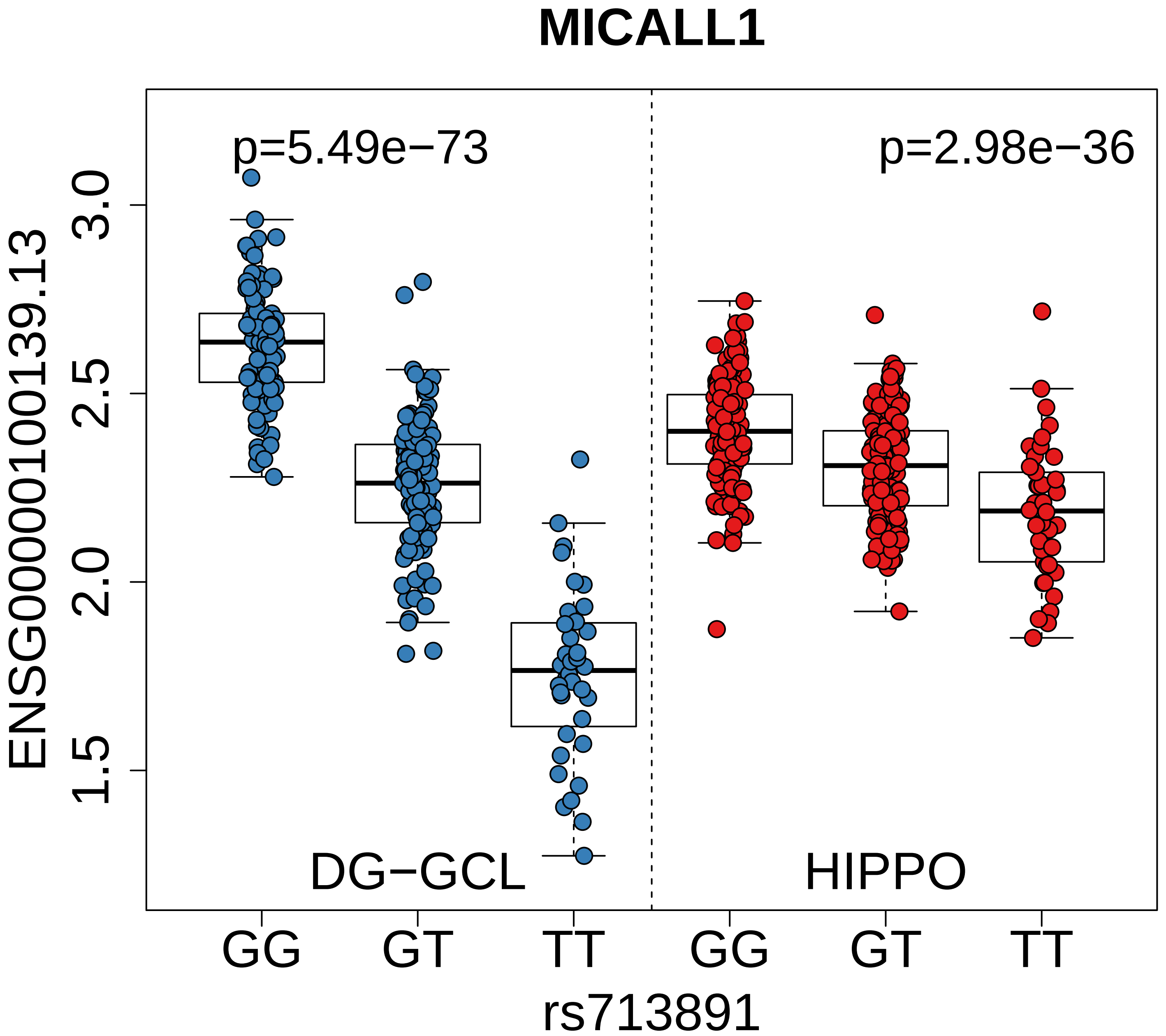
MICALL1
p=5.49e−73
p=2.98e−36
ENSG00000100139.13
3.0
2.5
2.0
1.5
DG−GCL
HIPPO
GG
GT
TT
GG
GT
TT
rs713891

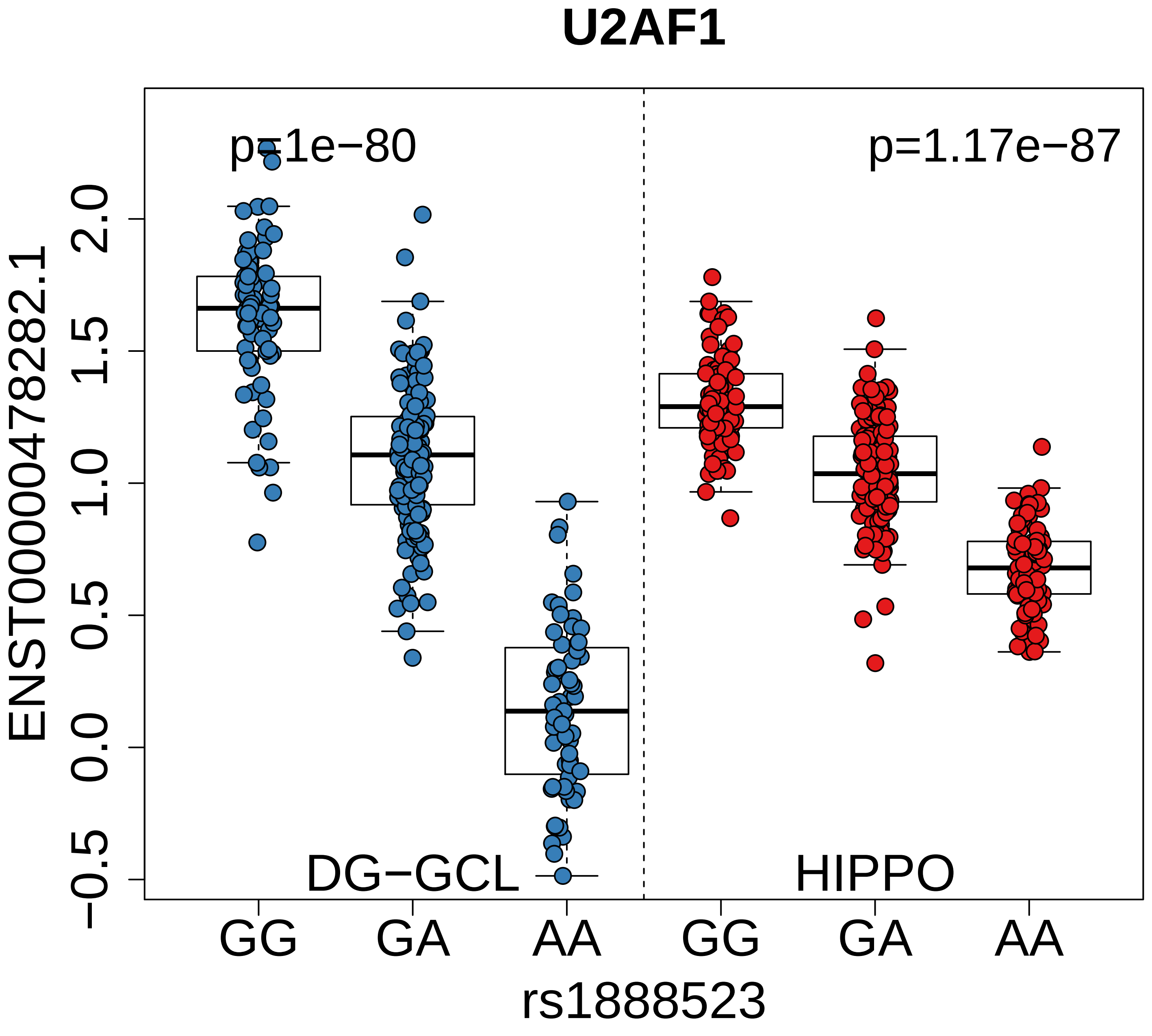

U2AF1
p=1e−80
p=1.17e−87
ENST00000478282.1
2.0
1.5
1.0
0.5
0.0
−0.5
DG−GCL
HIPPO
GG
GA
AA
GG
GA
AA
rs1888523

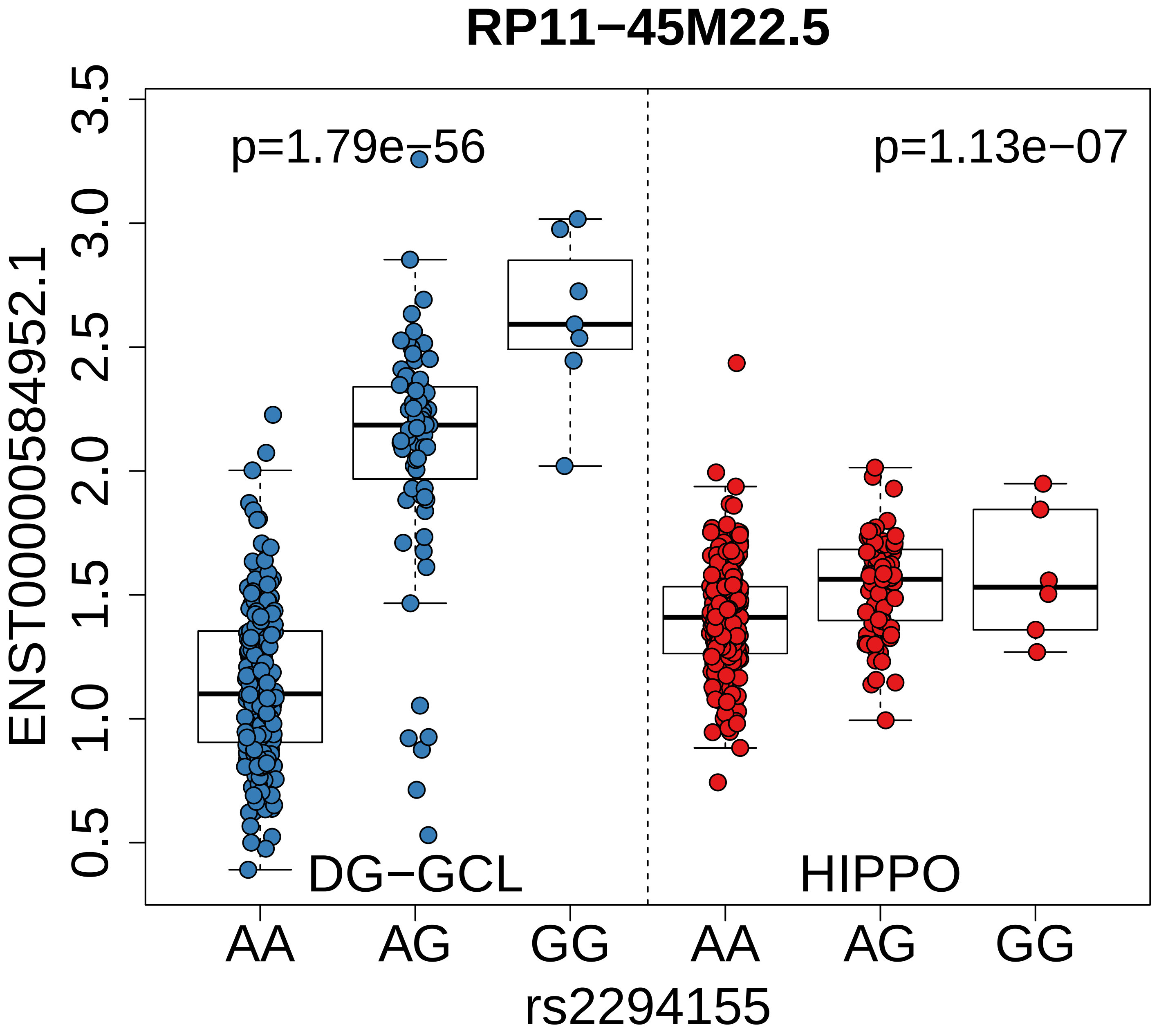
RP11−45M22.5
p=1.79e−56
p=1.13e−07
DG−GCL
HIPPO
ENST00000584952.1
0.5
1.0
1.5
2.0
2.5
3.0
3.5
AA
AG
GG
AA
AG
GG
rs2294155

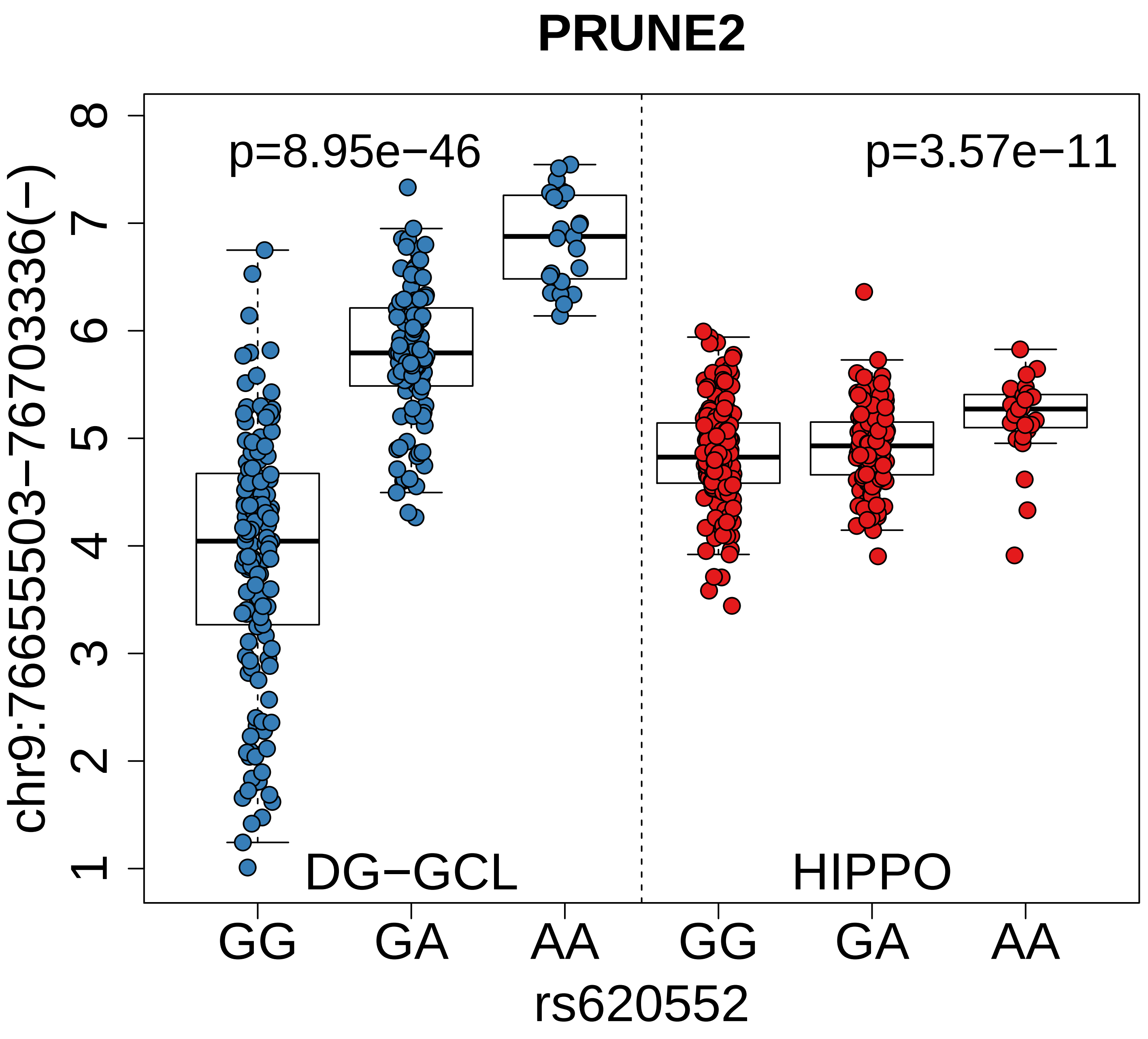
PRUNE2
p=8.95e−46
p=3.57e−11
DG−GCL
HIPPO
chr9:76655503−76703336(−)
1
2
3
4
5
6
7
8
GG
GA
AA
GG
GA
AA
rs620552

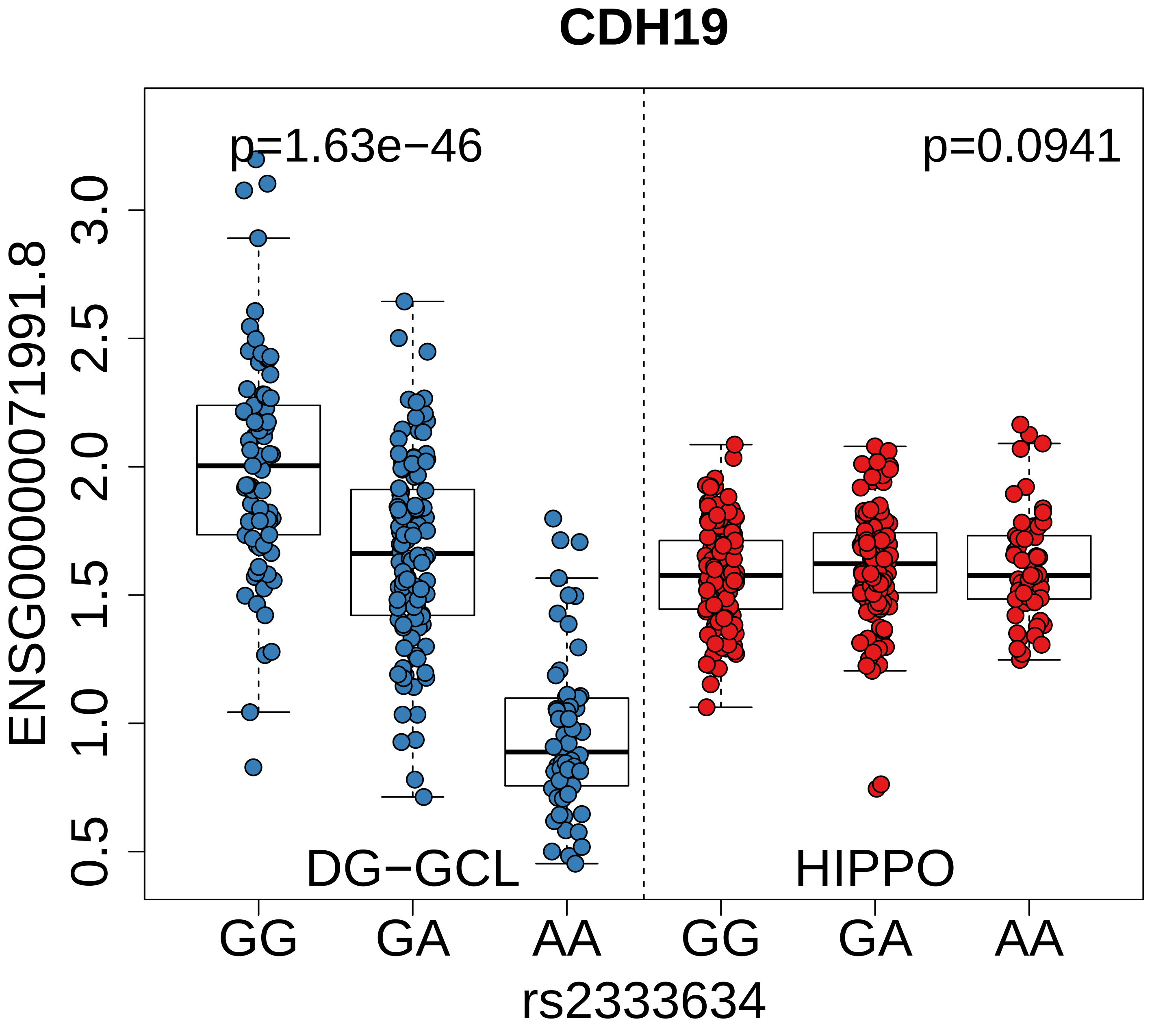
CDH19
p=1.63e−46
p=0.0941
ENSG00000071991.8
3.0
2.5
2.0
1.5
1.0
0.5
DG−GCL
HIPPO
GG
GA
AA
GG
GA
AA
rs2333634

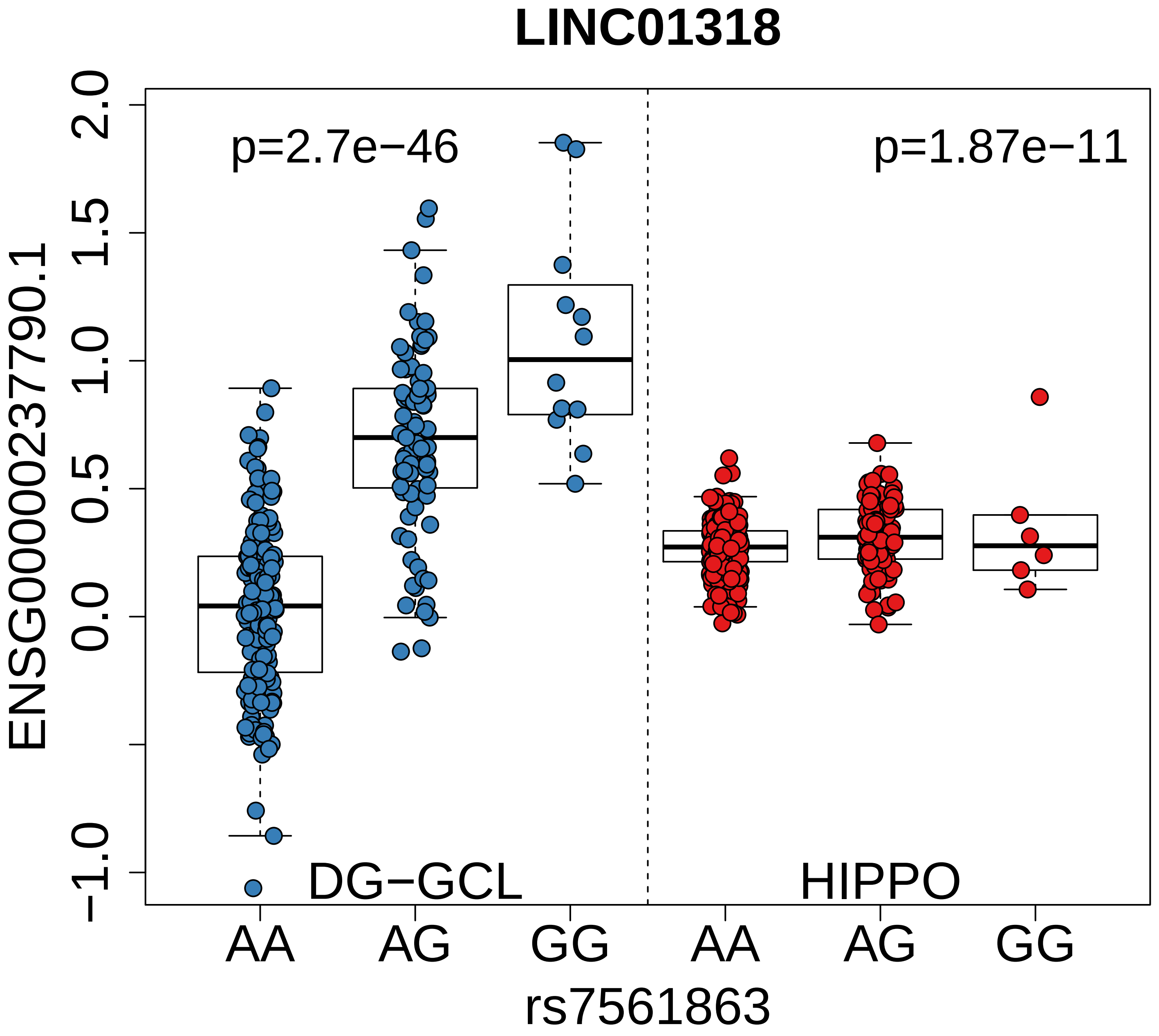
LINC01318
p=2.7e−46
p=1.87e−11
ENSG00000237790.1
2.0
1.5
1.0
0.5
0.0
−1.0
DG−GCL
HIPPO
AA
AG
GG
AA
AG
GG
rs7561863

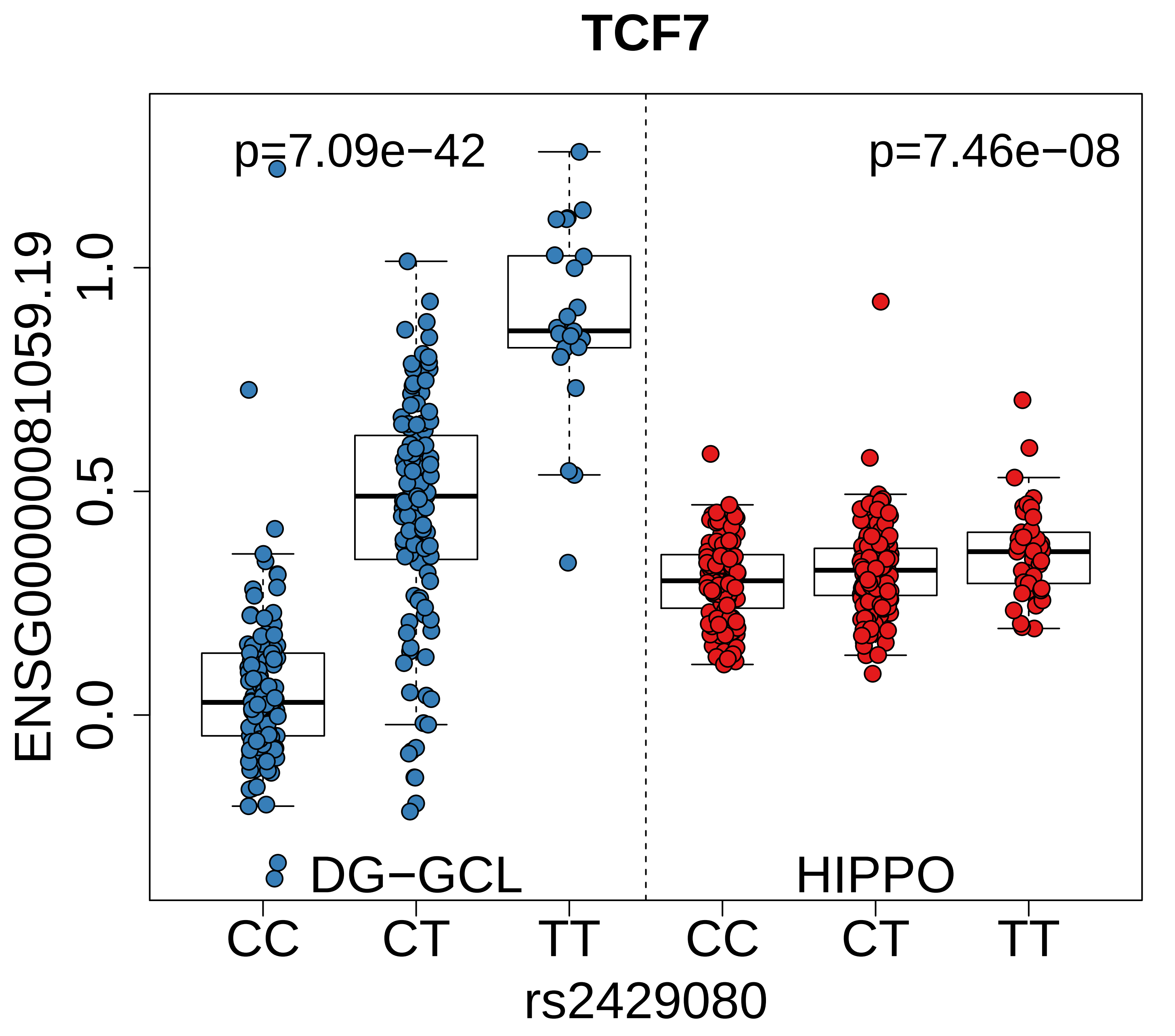
TCF7
p=7.09e−42
p=7.46e−08
ENSG00000081059.19
1.0
0.5
0.0
DG−GCL
HIPPO
CC
CT
TT
CC
CT
TT
rs2429080

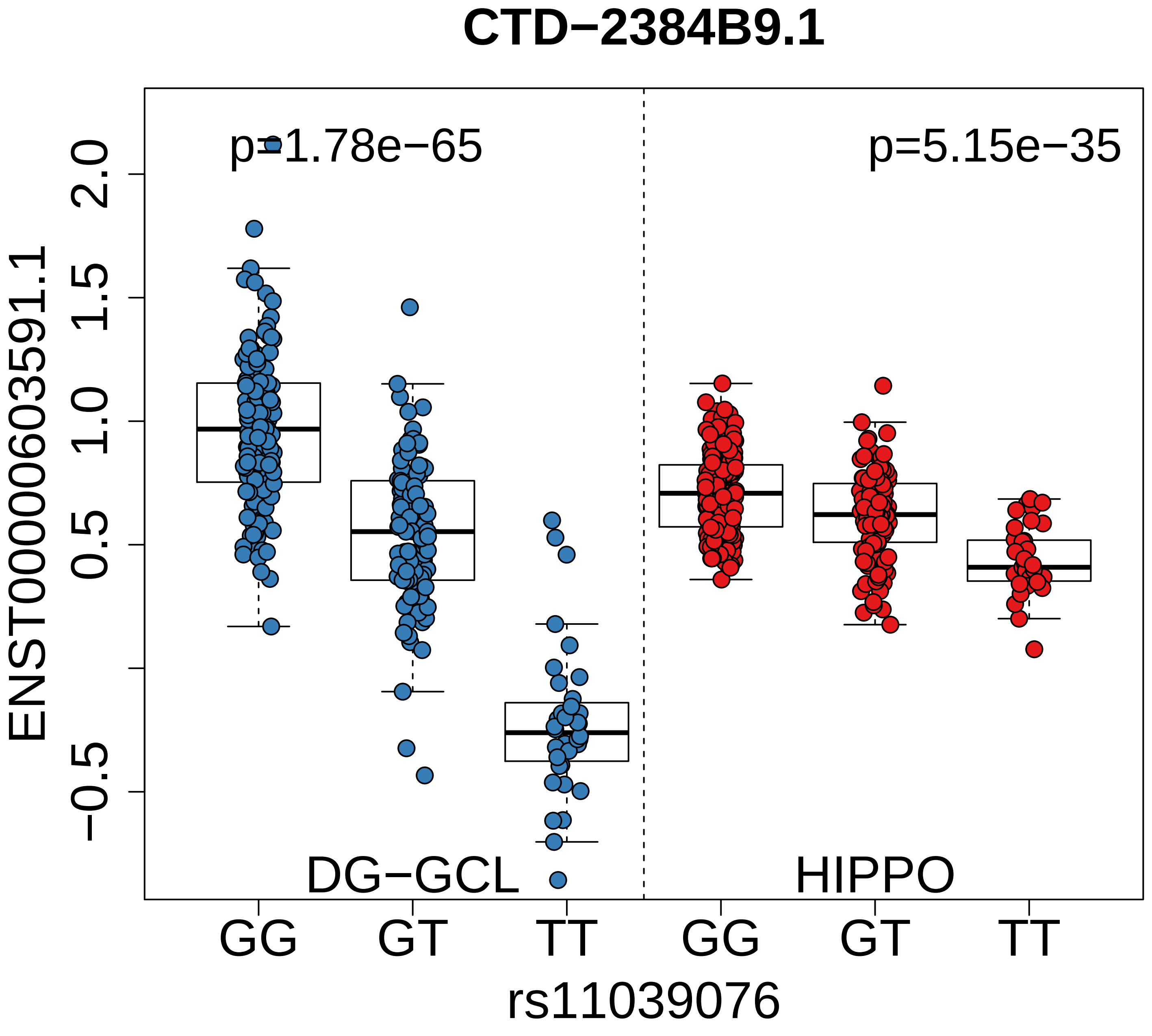
CTD−2384B9.1
p=1.78e−65
p=5.15e−35
ENST00000603591.1
2.0
1.5
1.0
0.5
−0.5
DG−GCL
HIPPO
GG
GT
TT
GG
GT
TT
rs11039076

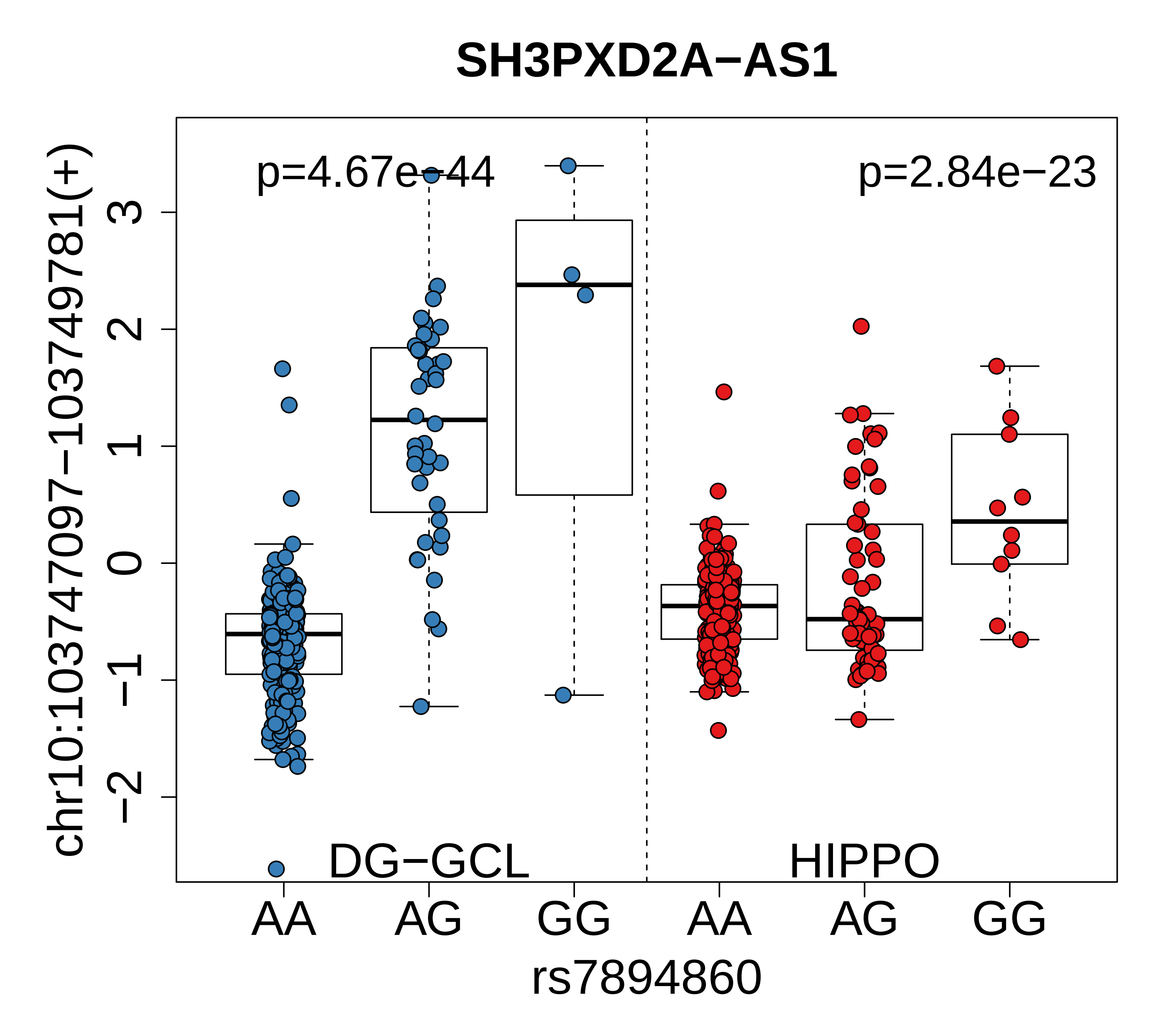

SH3PXD2A−AS1
p=4.67e−44
p=2.84e−23
DG−GCL
HIPPO
chr10:103747097−103749781(+)
3
2
1
0
−1
−2
AA
AG
GG
AA
AG
GG
rs7894860

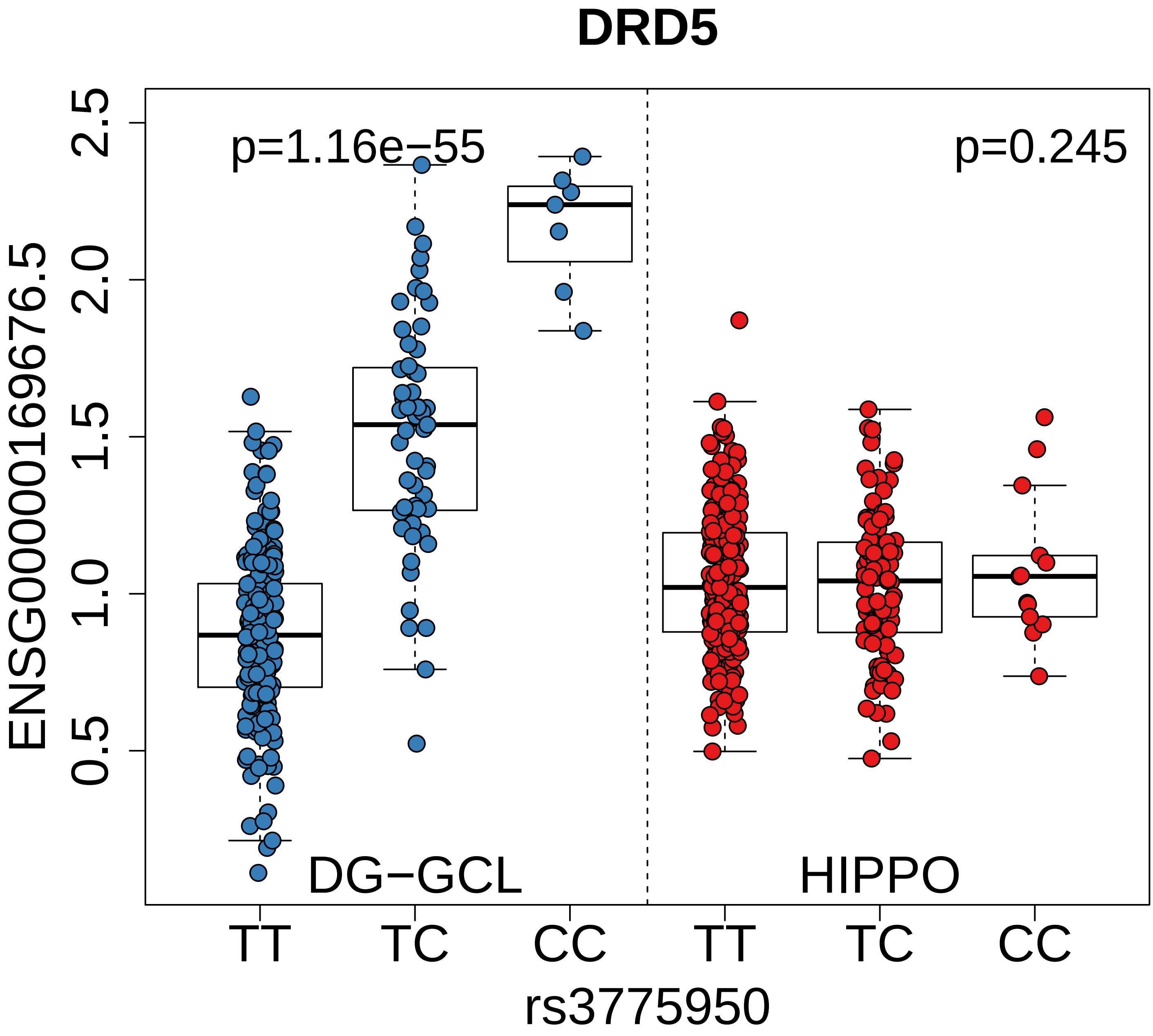
DRD5
p=1.16e−55
p=0.245
ENSG00000169676.5
2.5
2.0
1.5
1.0
0.5
DG−GCL
HIPPO
TT
TC
CC
TT
TC
CC
rs3775950

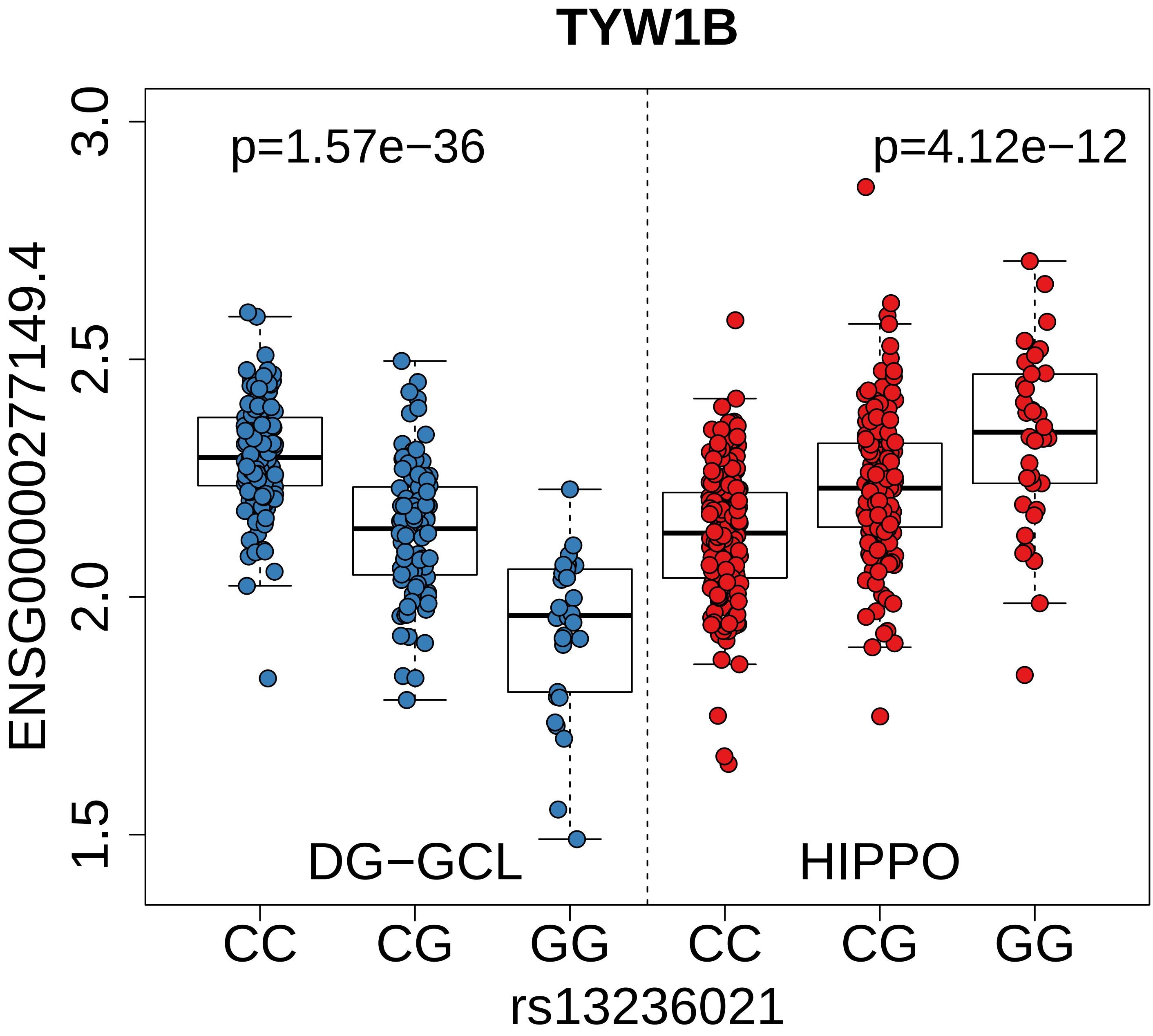
TYW1B
p=1.57e−36
p=4.12e−12
ENSG00000277149.4
3.0
2.5
2.0
1.5
DG−GCL
HIPPO
CC
CG
GG
CC
CG
GG
rs13236021

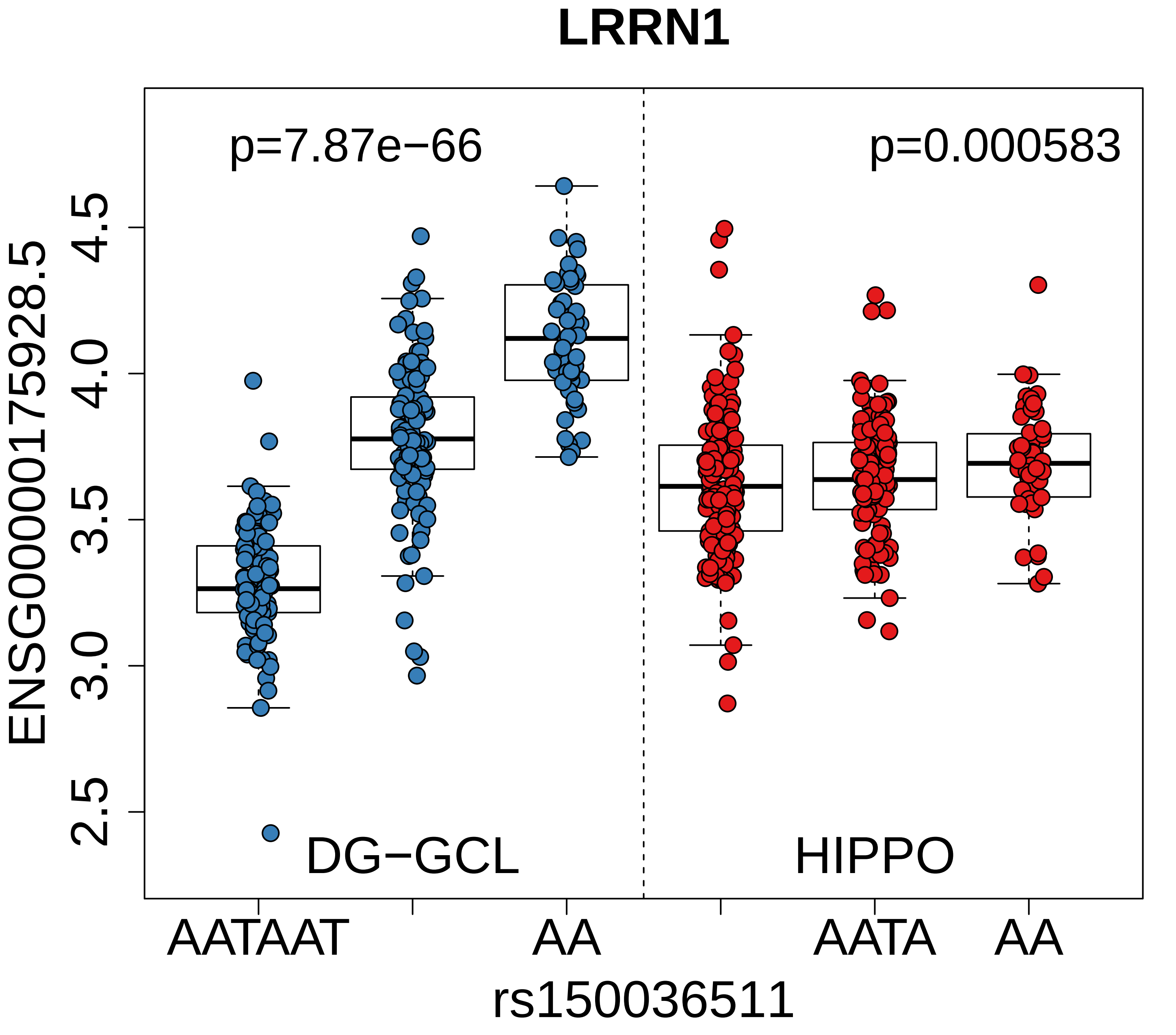

LRRN1
p=7.87e−66
p=0.000583
ENSG00000175928.5
4.5
4.0
3.5
3.0
2.5
DG−GCL
HIPPO
AATAAT
AA
AATA
AA
rs150036511

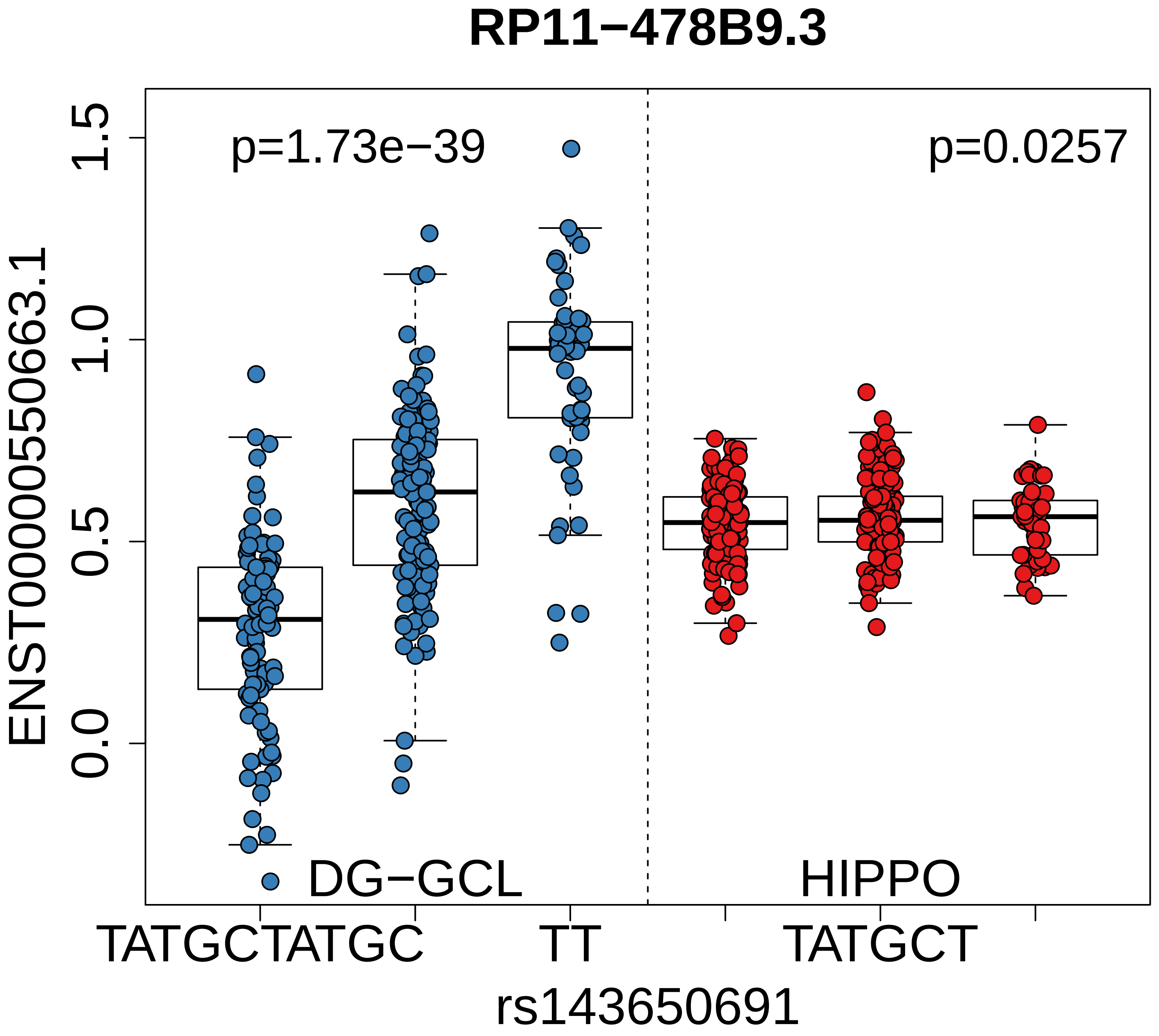

RP11−478B9.3
p=1.73e−39
p=0.0257
ENST00000550663.1
1.5
1.0
0.5
0.0
DG−GCL
HIPPO
TATGCTATGC
TT
TATGCT
rs143650691

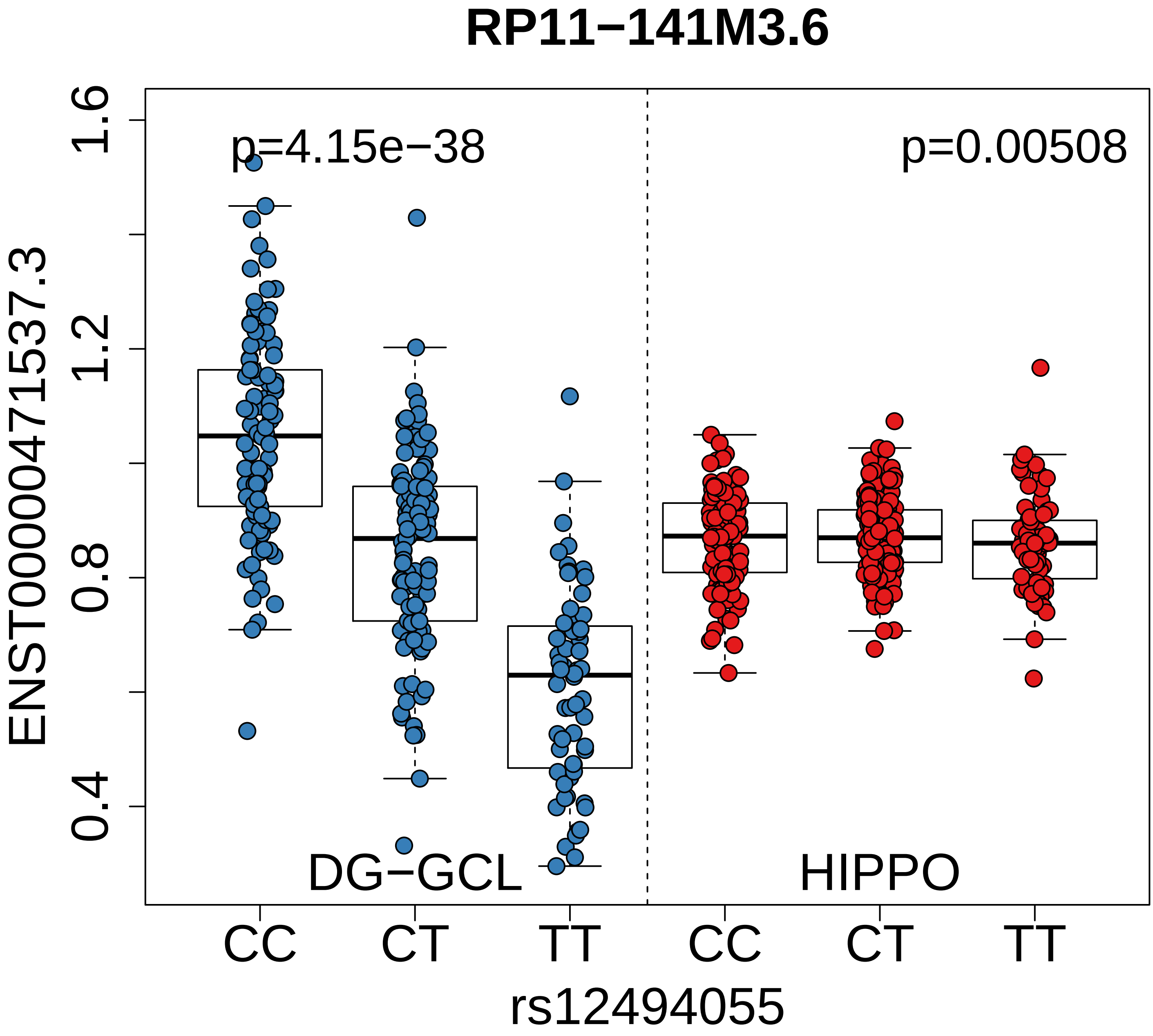

RP11−141M3.6
p=4.15e−38
p=0.00508
1.6
1.2
0.8
0.4
ENST00000471537.3
DG−GCL
HIPPO
CC
CT
TT
CC
CT
TT
rs12494055

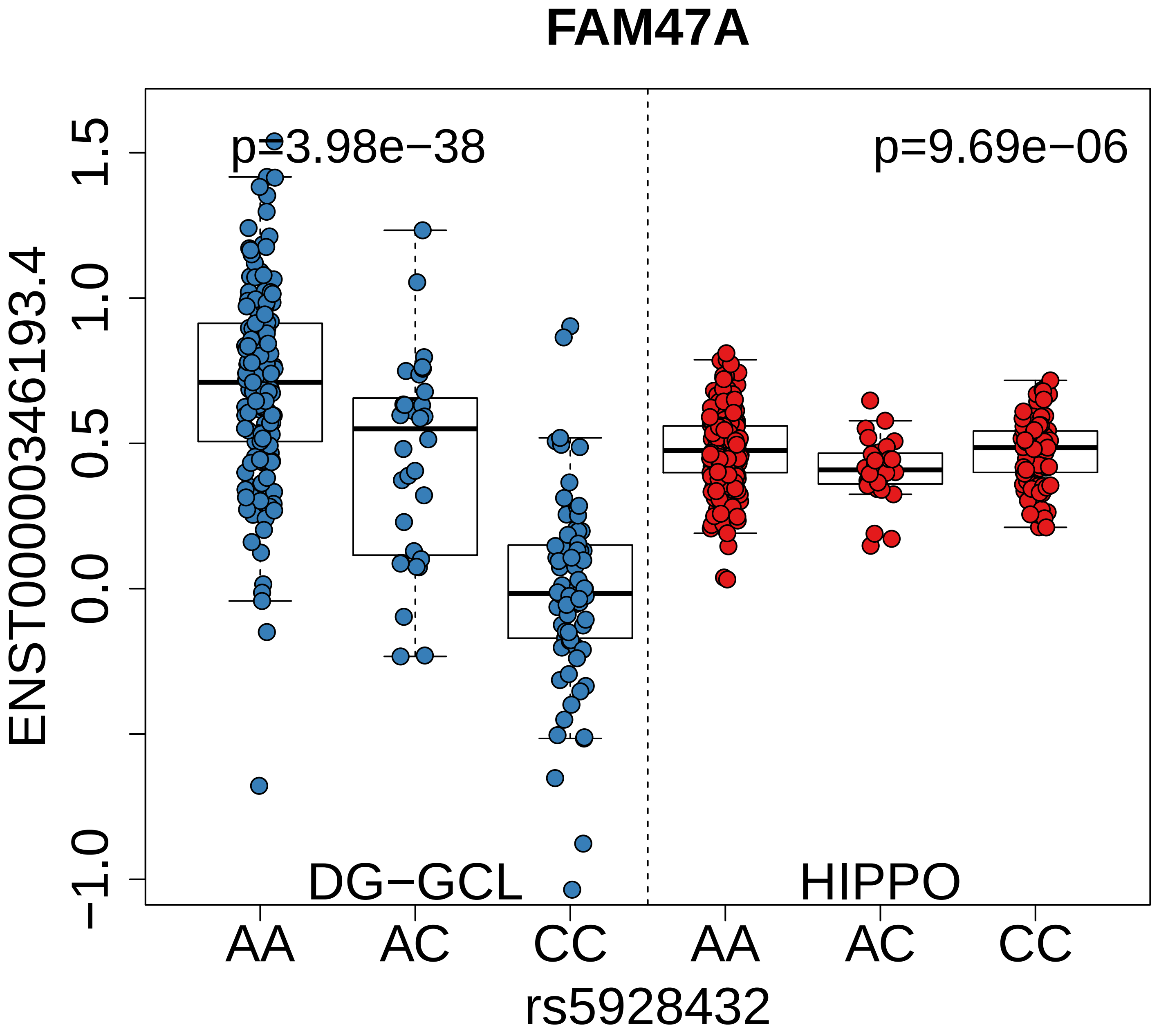
FAM47A
p=3.98e−38
p=9.69e−06
ENST00000346193.4
1.5
1.0
0.5
0.0
−1.0
DG−GCL
HIPPO
AA
AC
CC
AA
AC
CC
rs5928432

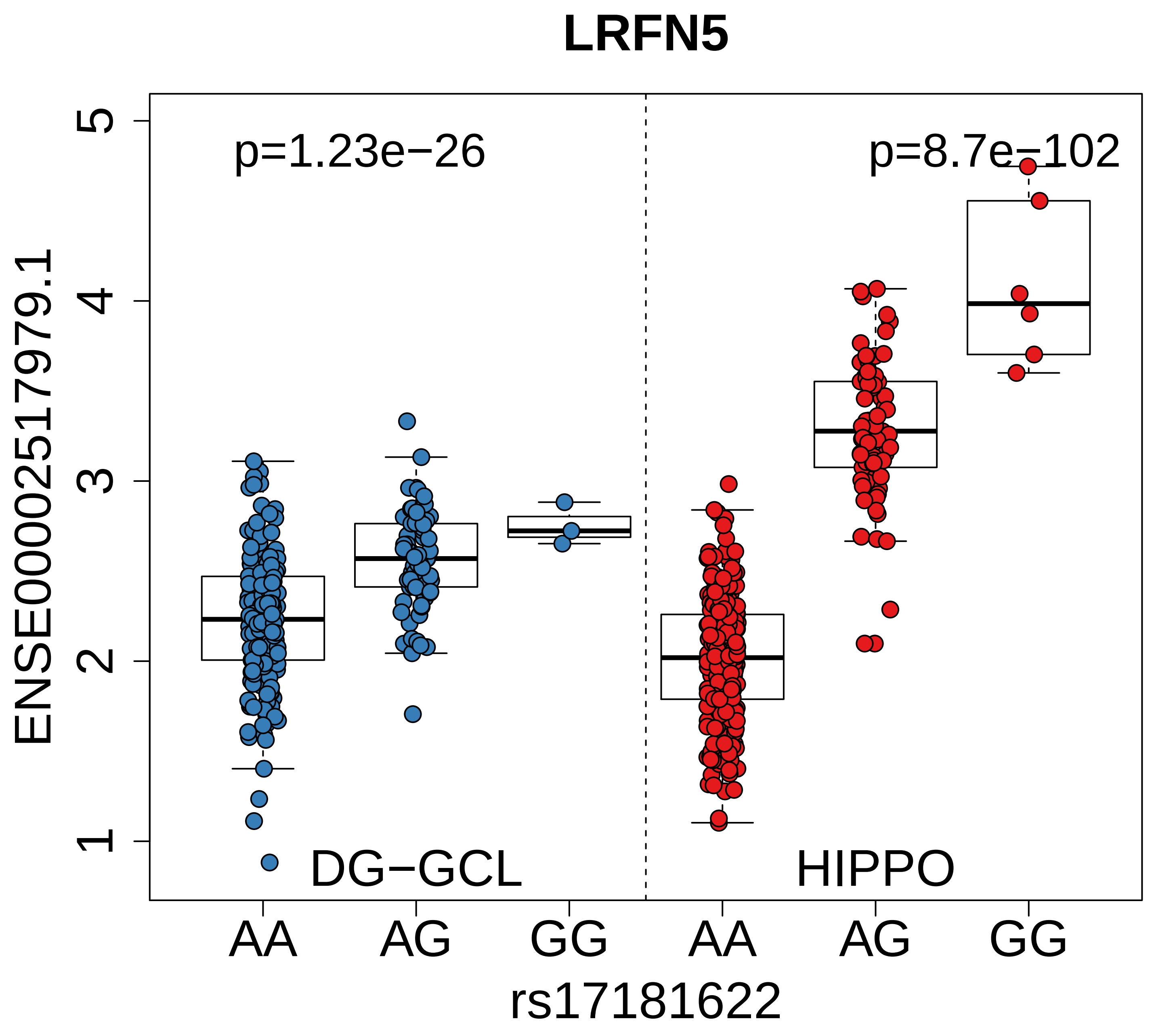
LRFN5
p=1.23e−26
p=8.7e−102
ENSE00002517979.1
1
2
3
4
5
DG−GCL
HIPPO
AA
AG
GG
AA
AG
GG
rs17181622

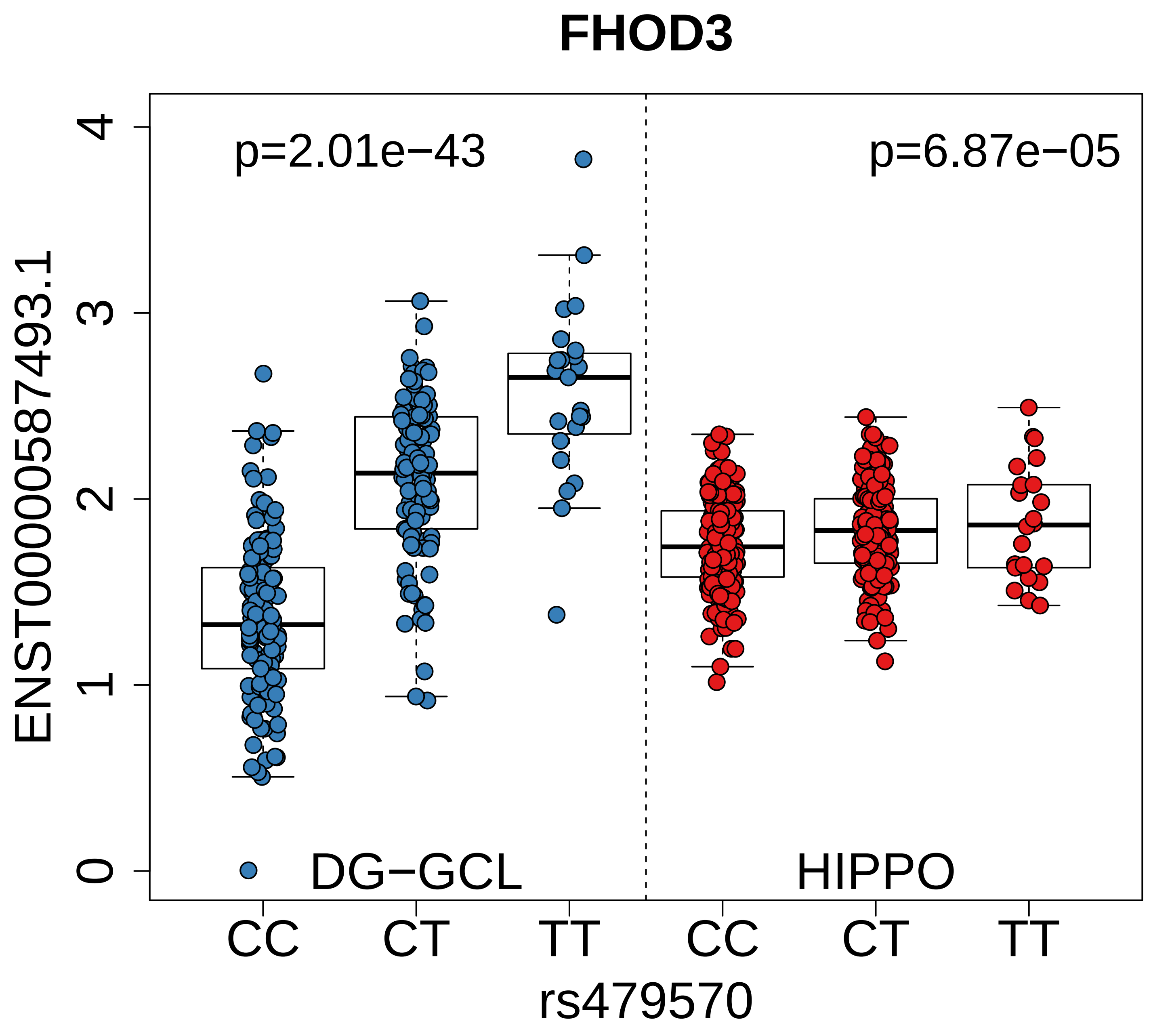

FHOD3
p=2.01e−43
p=6.87e−05
ENST00000587493.1
0
1
2
3
4
DG−GCL
HIPPO
CC
CT
TT
CC
CT
TT
rs479570

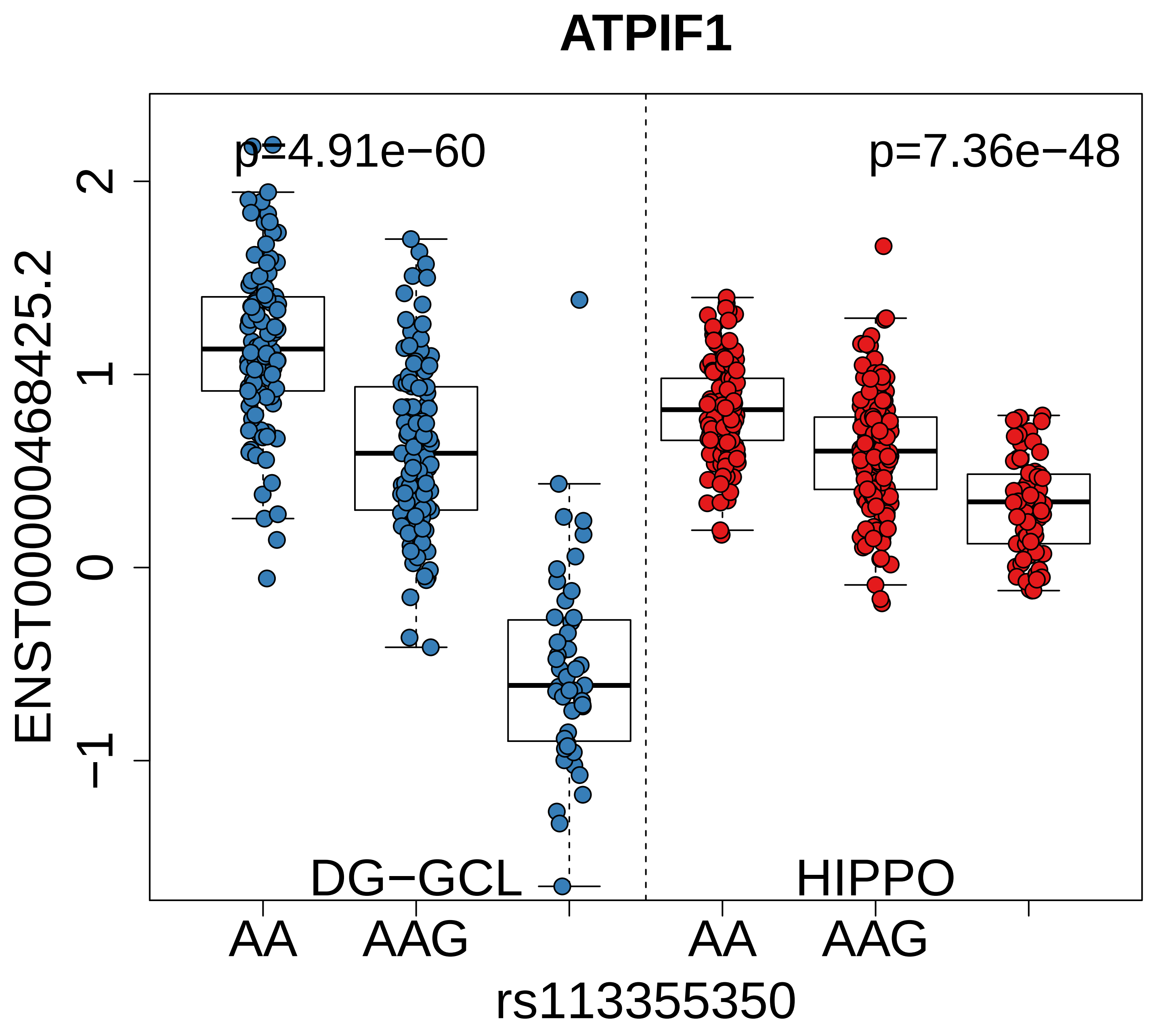
ATPIF1
p=4.91e−60
p=7.36e−48
ENST00000468425.2
2
1
0
−1
DG−GCL
HIPPO
AA
AAG
AA
AAG
rs113355350

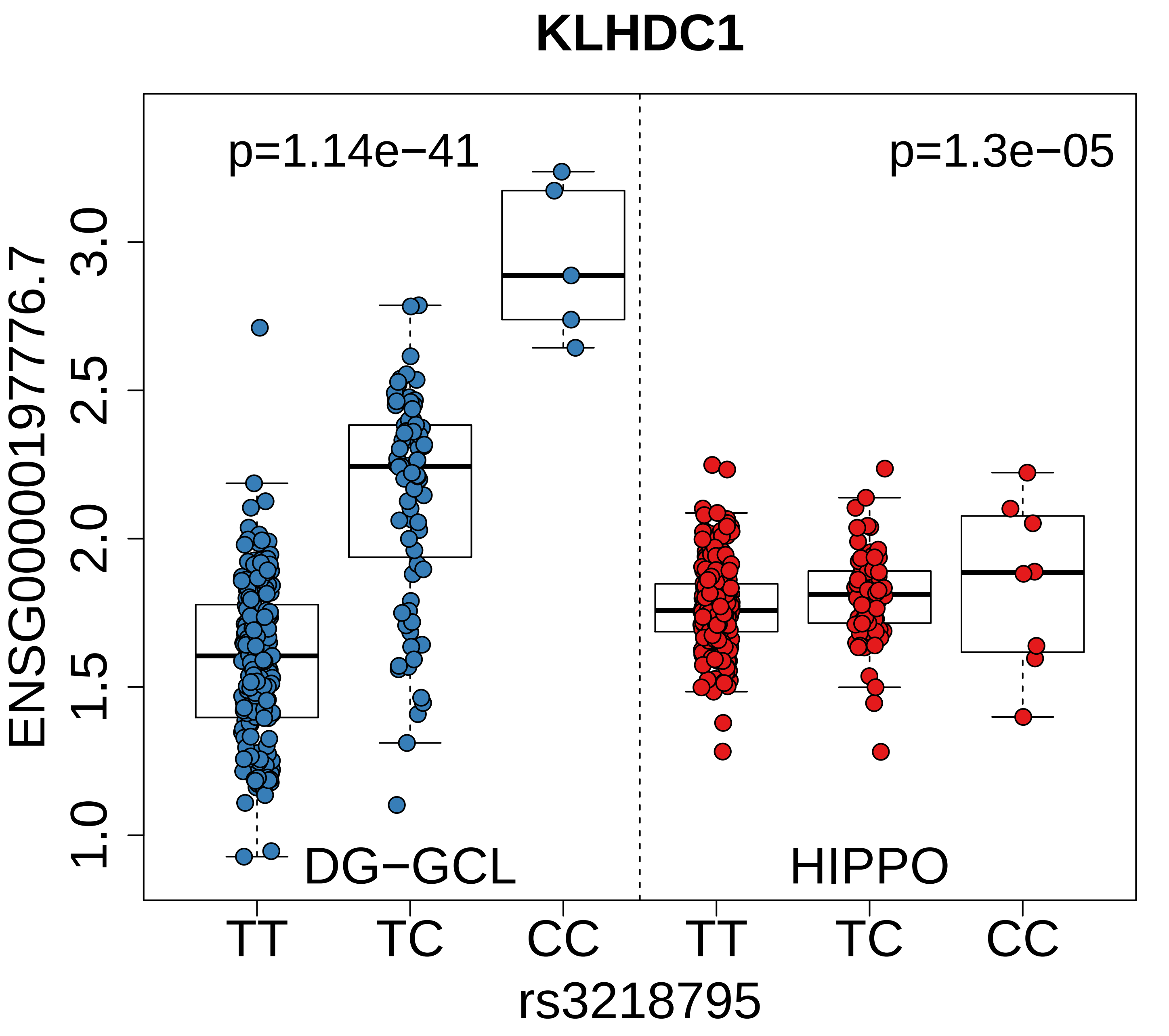
KLHDC1
p=1.14e−41
p=1.3e−05
ENSG00000197776.7
3.0
2.5
2.0
1.5
1.0
DG−GCL
HIPPO
TT
TC
CC
TT
TC
CC
rs3218795

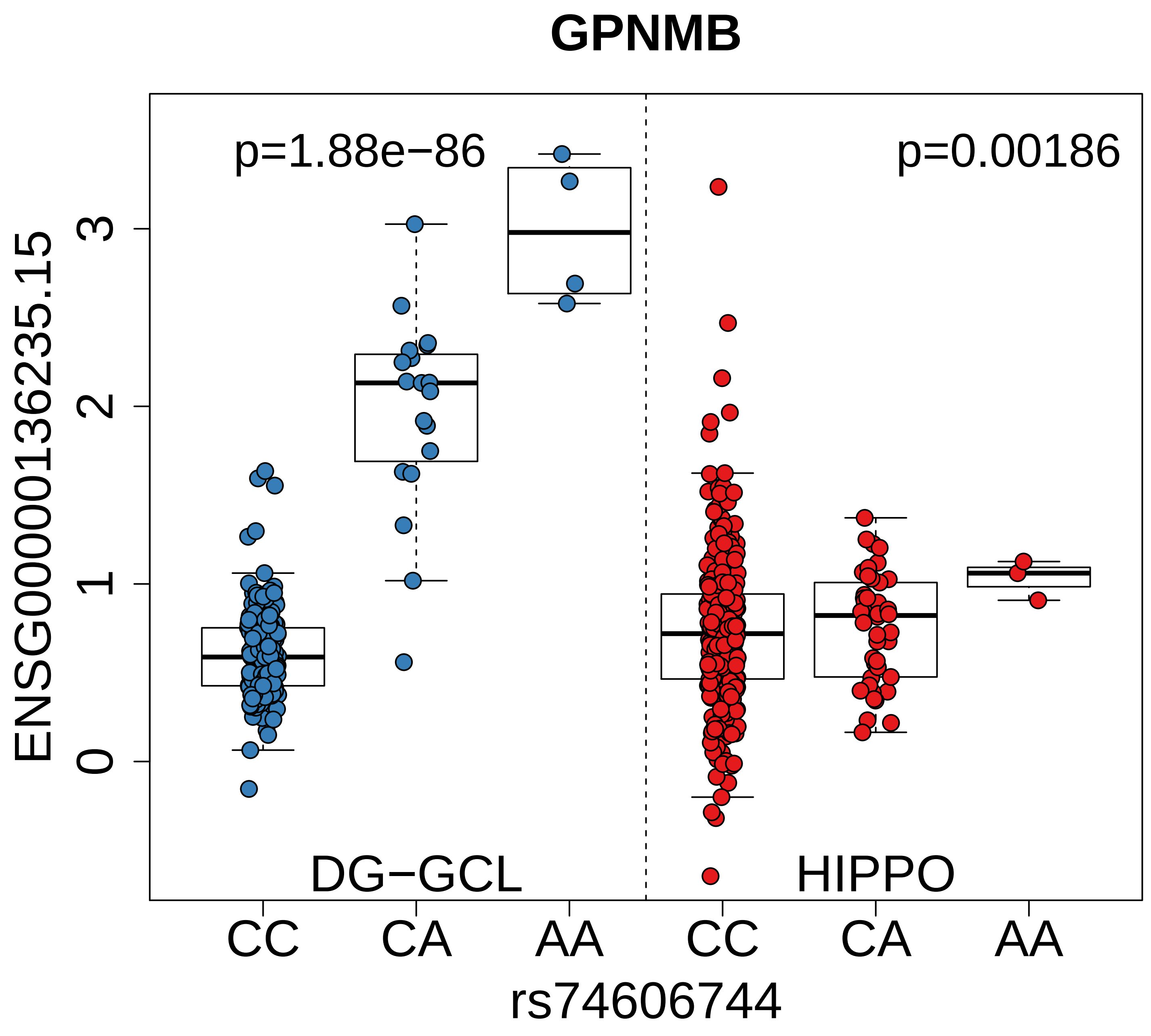

GPNMB
p=1.88e−86
p=0.00186
ENSG00000136235.15
0
1
2
3
DG−GCL
HIPPO
CC
CA
AA
CC
CA
AA
rs74606744

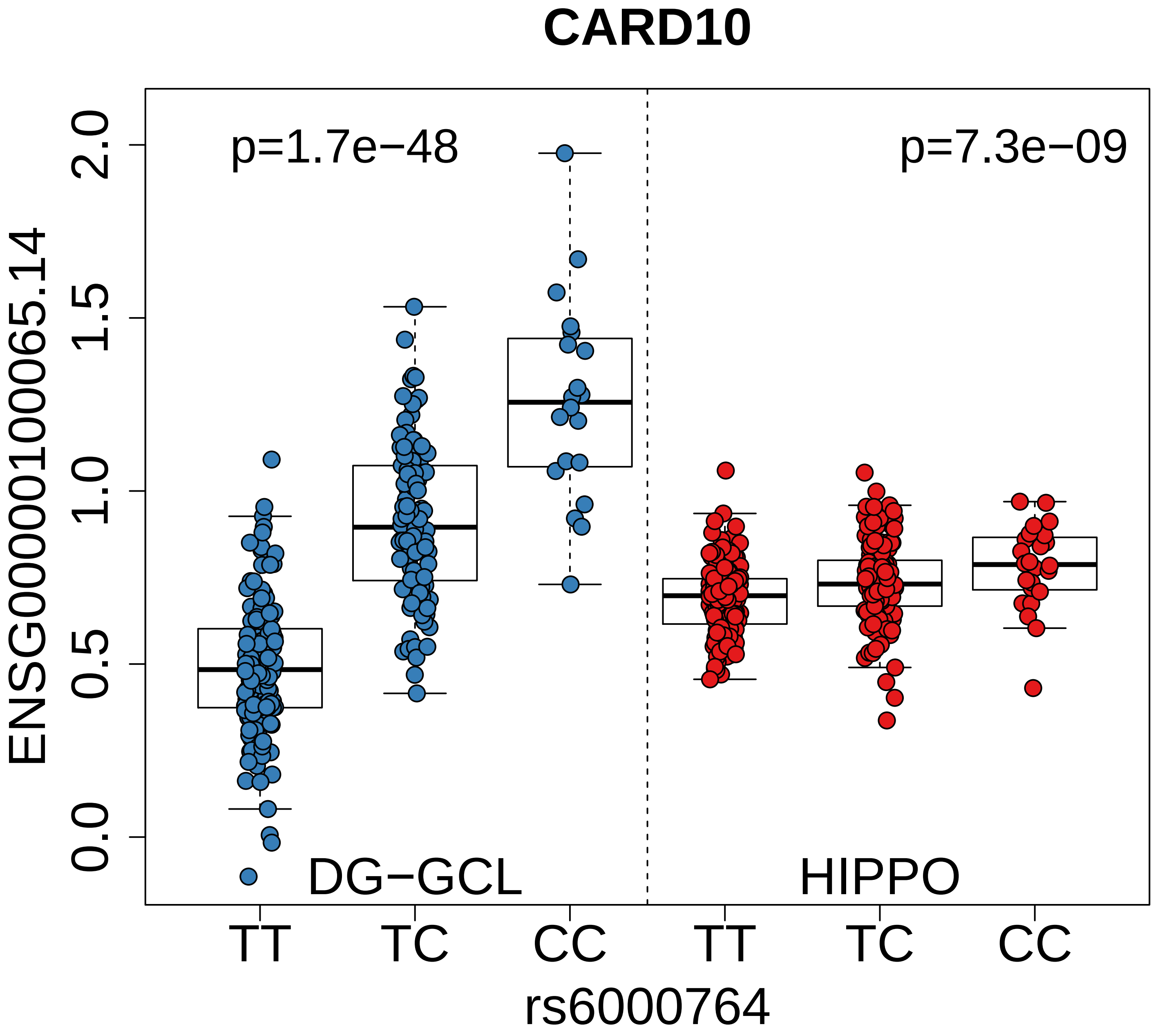
CARD10
p=1.7e−48
p=7.3e−09
ENSG00000100065.14
2.0
1.5
1.0
0.5
0.0
DG−GCL
HIPPO
TT
TC
CC
TT
TC
CC
rs6000764

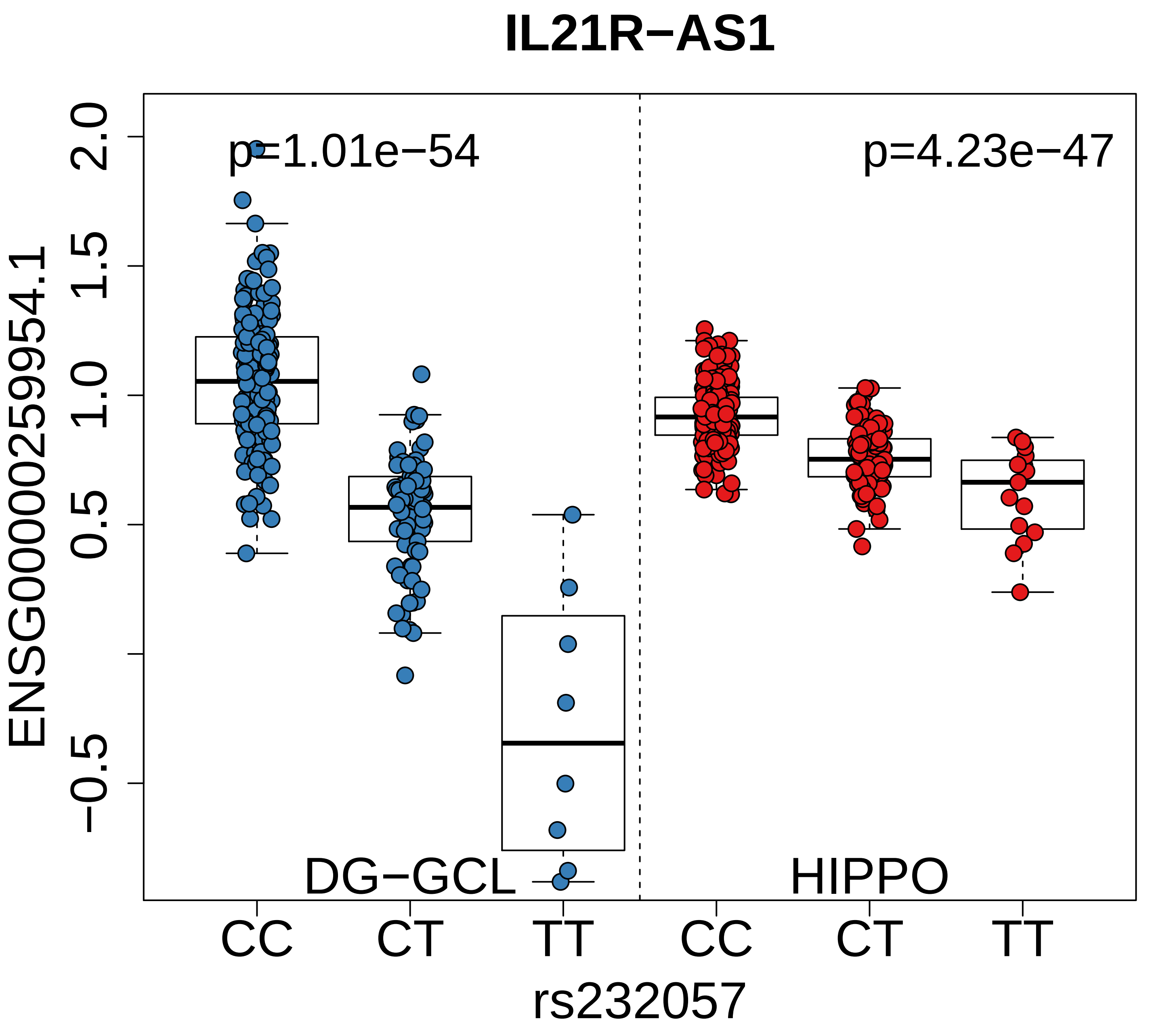
IL21R−AS1
p=1.01e−54
p=4.23e−47
ENSG00000259954.1
2.0
1.5
1.0
0.5
−0.5
DG−GCL
HIPPO
CC
CT
TT
CC
CT
TT
rs232057

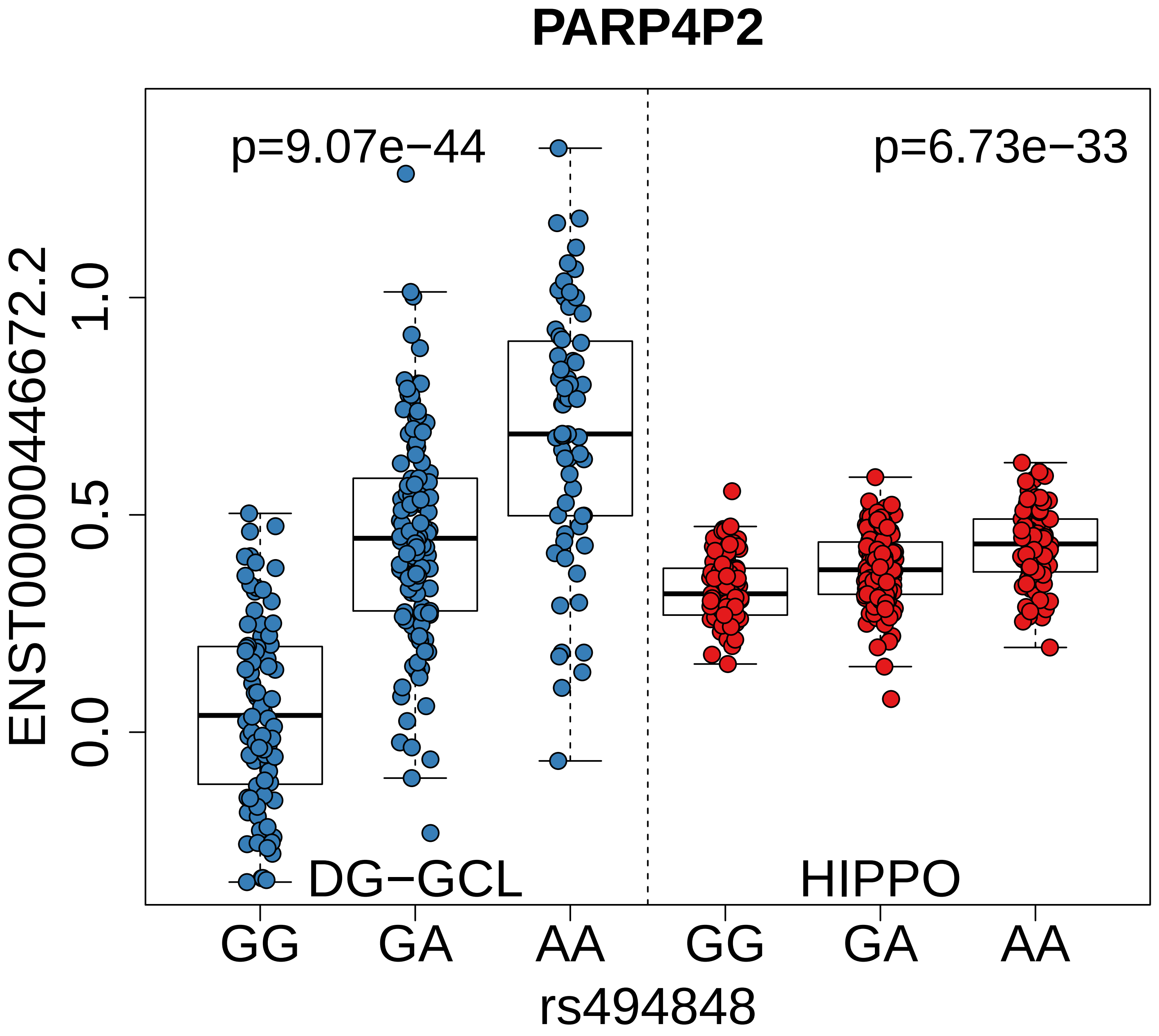
PARP4P2
p=9.07e−44
p=6.73e−33
ENST00000446672.2
1.0
0.5
0.0
DG−GCL
HIPPO
GG
GA
AA
GG
GA
AA
rs494848

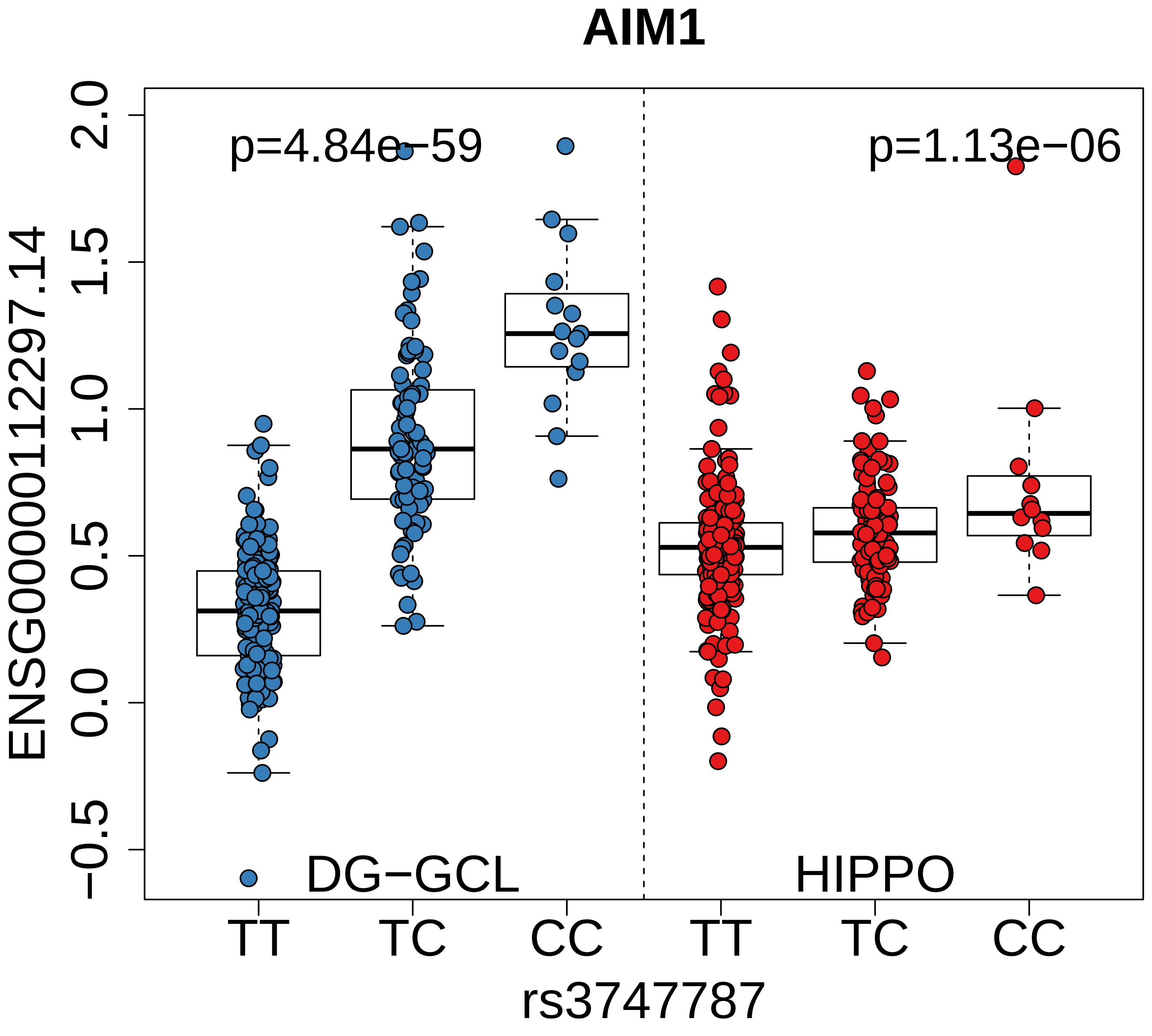
AIM1
p=4.84e−59
p=1.13e−06
ENSG00000112297.14
2.0
1.5
1.0
0.5
0.0
−0.5
DG−GCL
HIPPO
TT
TC
CC
TT
TC
CC
rs3747787

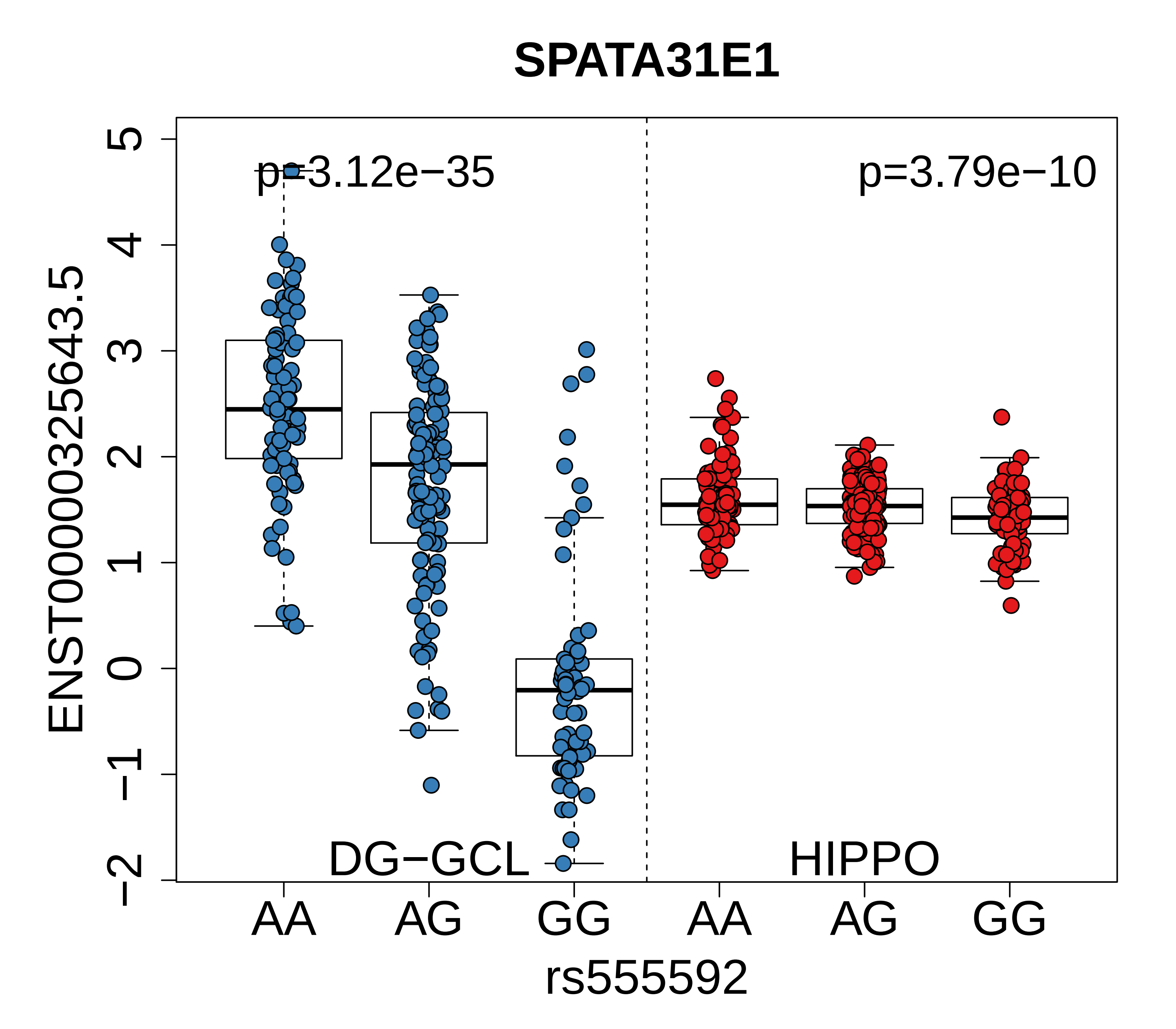

SPATA31E1
p=3.12e−35
p=3.79e−10
ENST00000325643.5
5
4
3
2
1
0
−1
−2
DG−GCL
HIPPO
AA
AG
GG
AA
AG
GG
rs555592

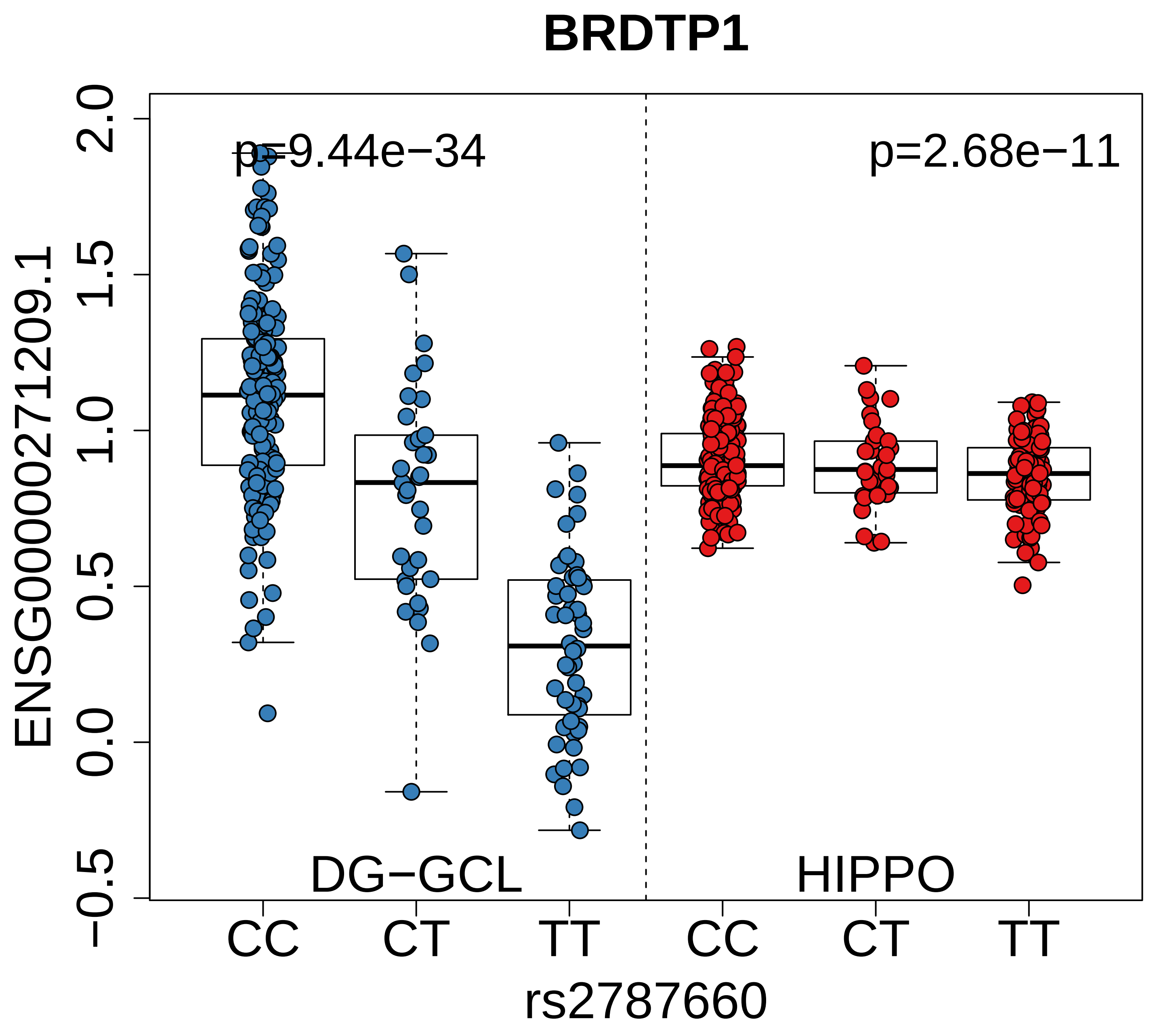

BRDTP1
p=9.44e−34
p=2.68e−11
ENSG00000271209.1
2.0
1.5
1.0
0.5
0.0
−0.5
DG−GCL
HIPPO
CC
CT
TT
CC
CT
TT
rs2787660

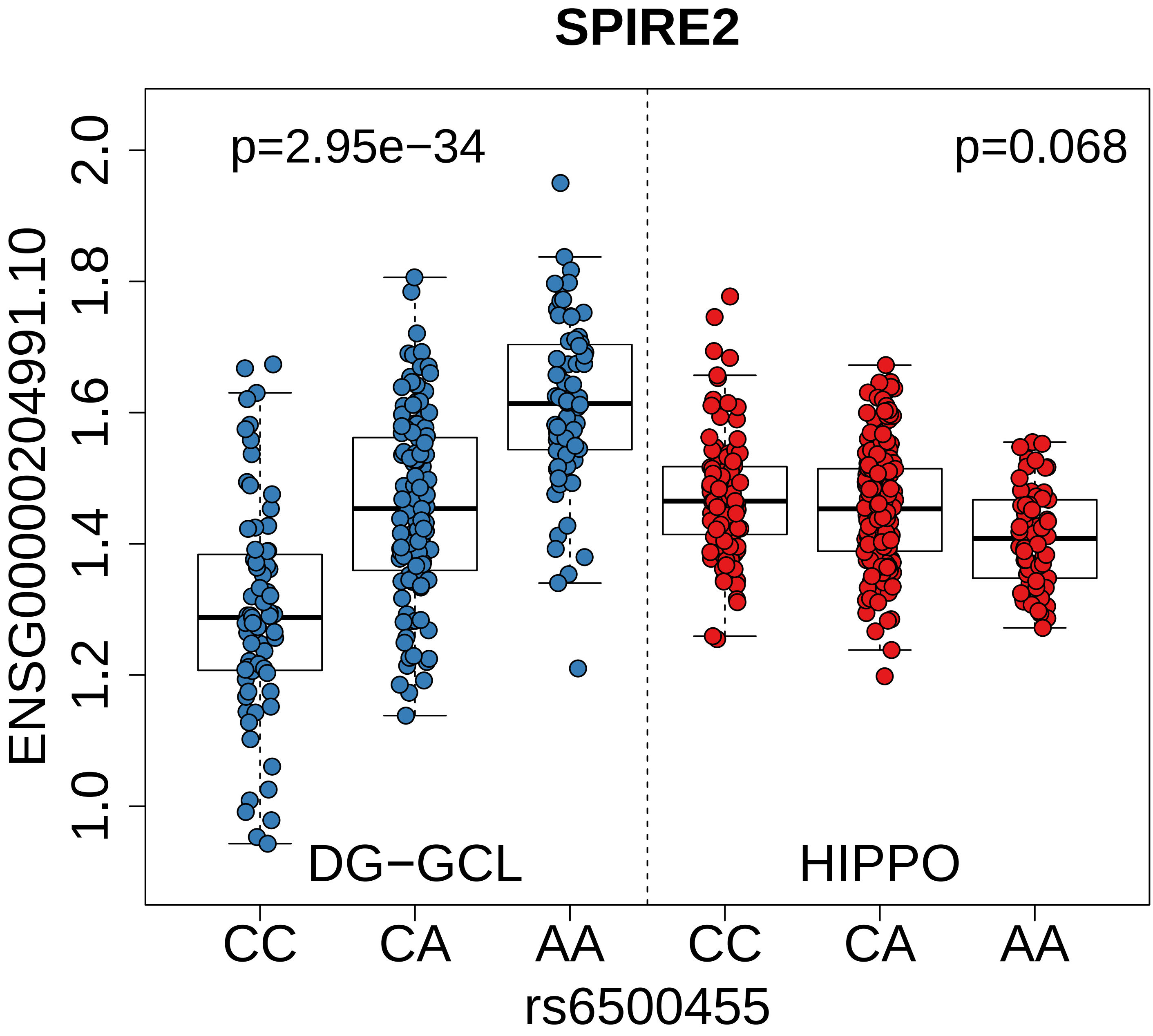
SPIRE2
p=2.95e−34
p=0.068
ENSG00000204991.10
2.0
1.8
1.6
1.4
1.2
1.0
DG−GCL
HIPPO
CC
CA
AA
CC
CA
AA
rs6500455

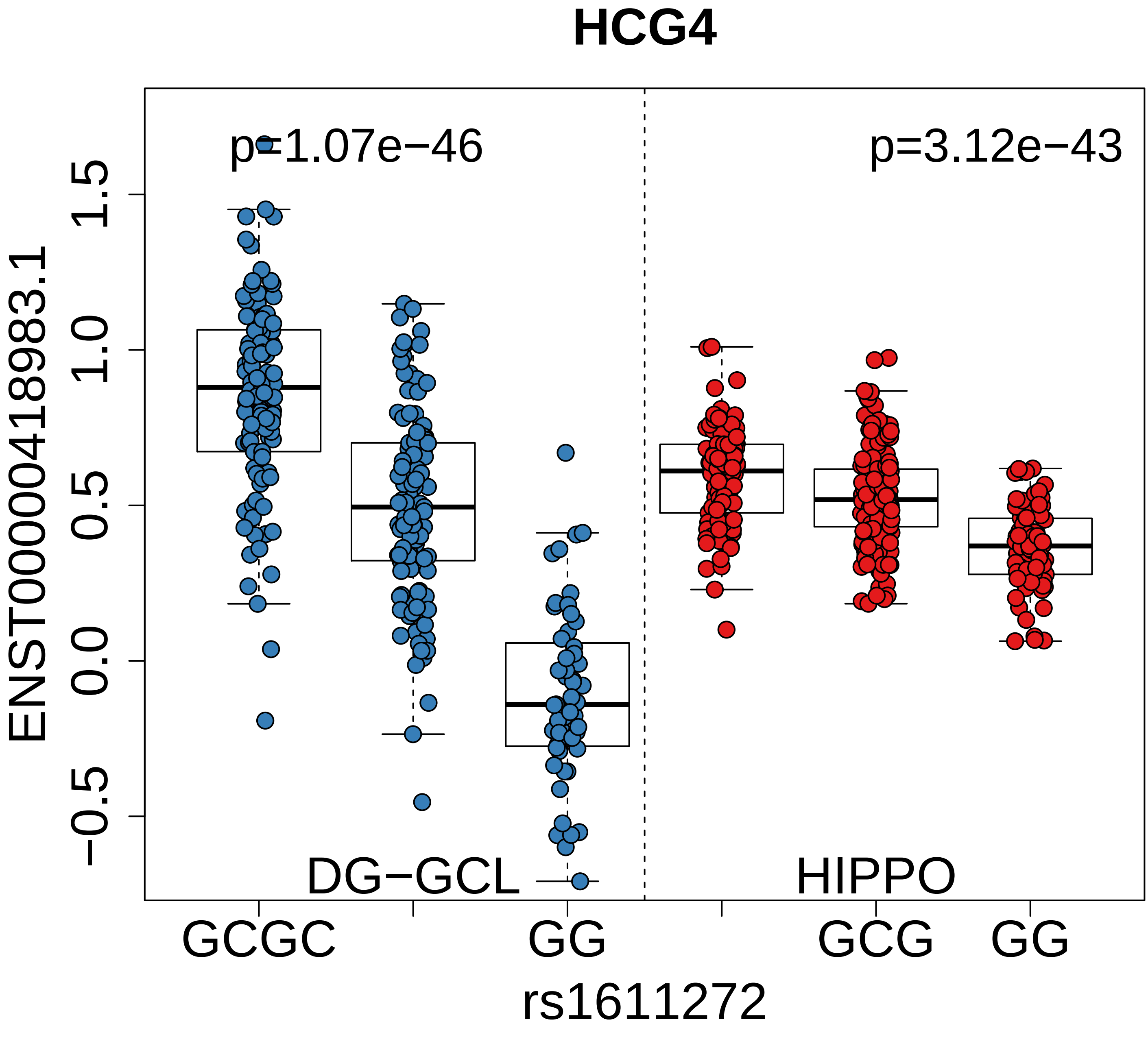
HCG4
p=1.07e−46
p=3.12e−43
ENST00000418983.1
1.5
1.0
0.5
0.0
−0.5
DG−GCL
HIPPO
GCGC
GG
GCG
GG
rs1611272

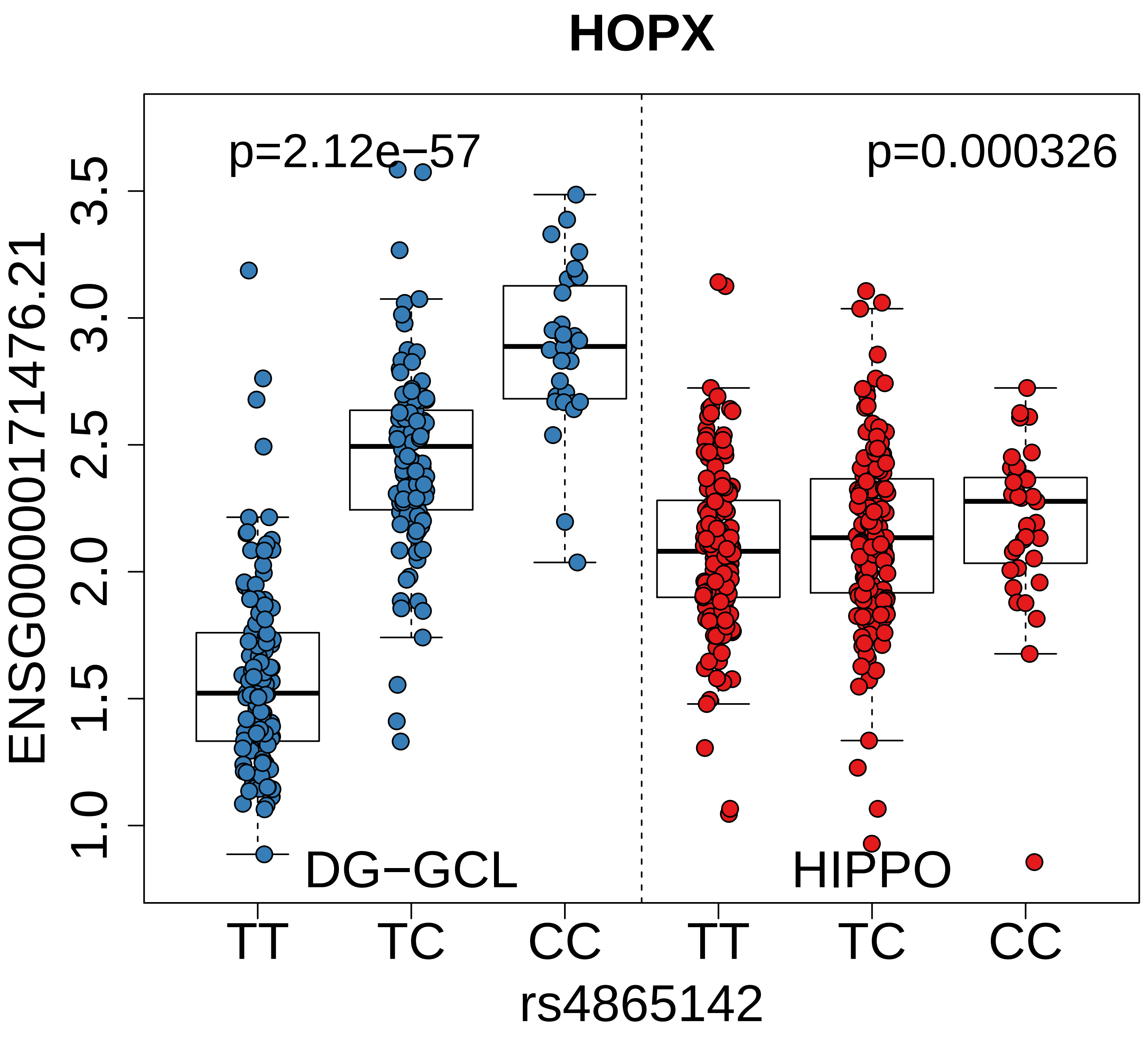
HOPX
p=2.12e−57
p=0.000326
ENSG00000171476.21
3.5
3.0
2.5
2.0
1.5
1.0
DG−GCL
HIPPO
TT
TC
CC
TT
TC
CC
rs4865142

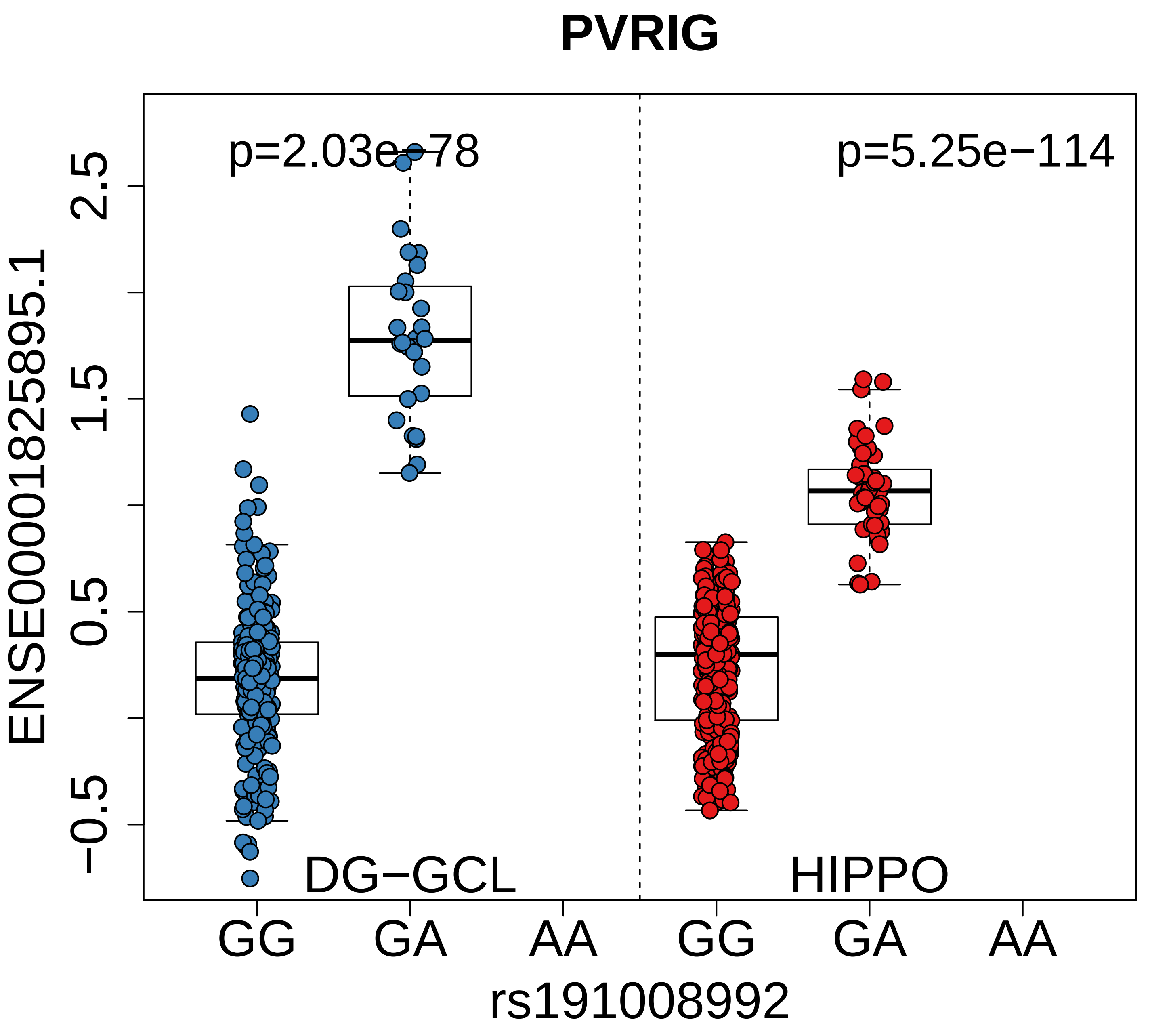

PVRIG
p=2.03e−78
p=5.25e−114
ENSE00001825895.1
2.5
1.5
0.5
−0.5
DG−GCL
HIPPO
GG
GA
AA
GG
GA
AA
rs191008992

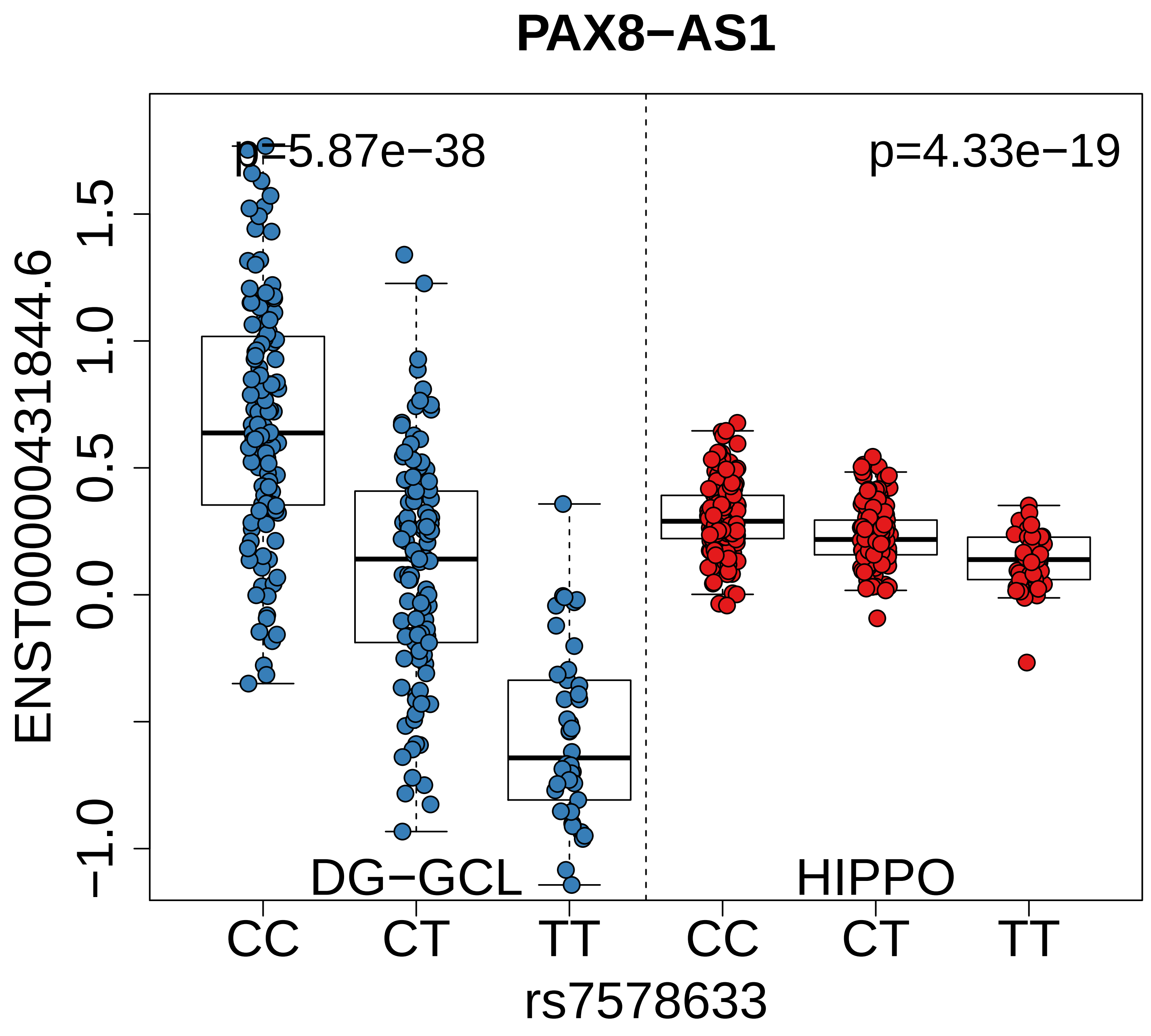

PAX8−AS1
p=5.87e−38
p=4.33e−19
ENST00000431844.6
1.5
1.0
0.5
0.0
−1.0
DG−GCL
HIPPO
CC
CT
TT
CC
CT
TT
rs7578633

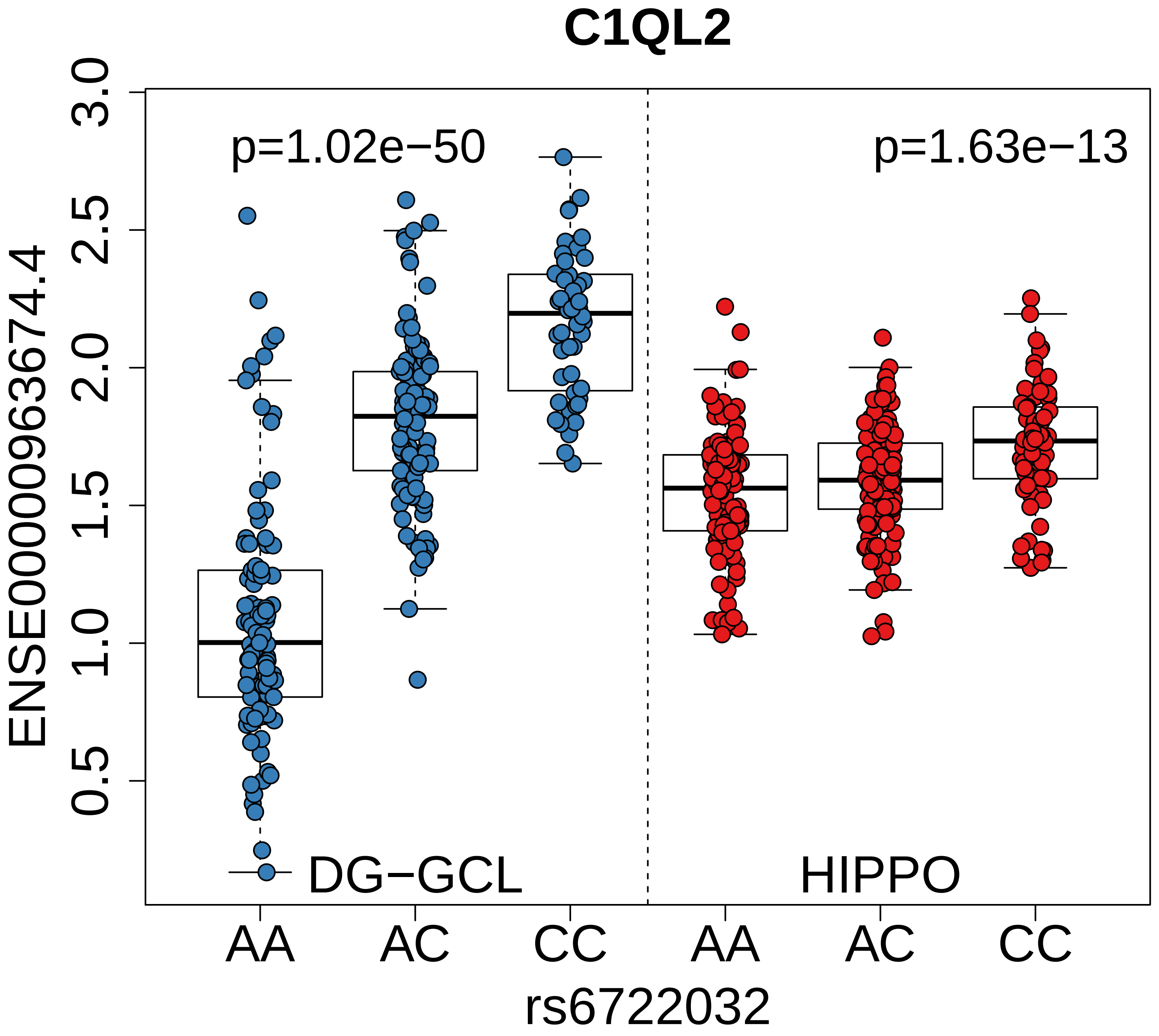
C1QL2
p=1.02e−50
p=1.63e−13
ENSE00000963674.4
3.0
2.5
2.0
1.5
1.0
0.5
DG−GCL
HIPPO
AA
AC
CC
AA
AC
CC
rs6722032

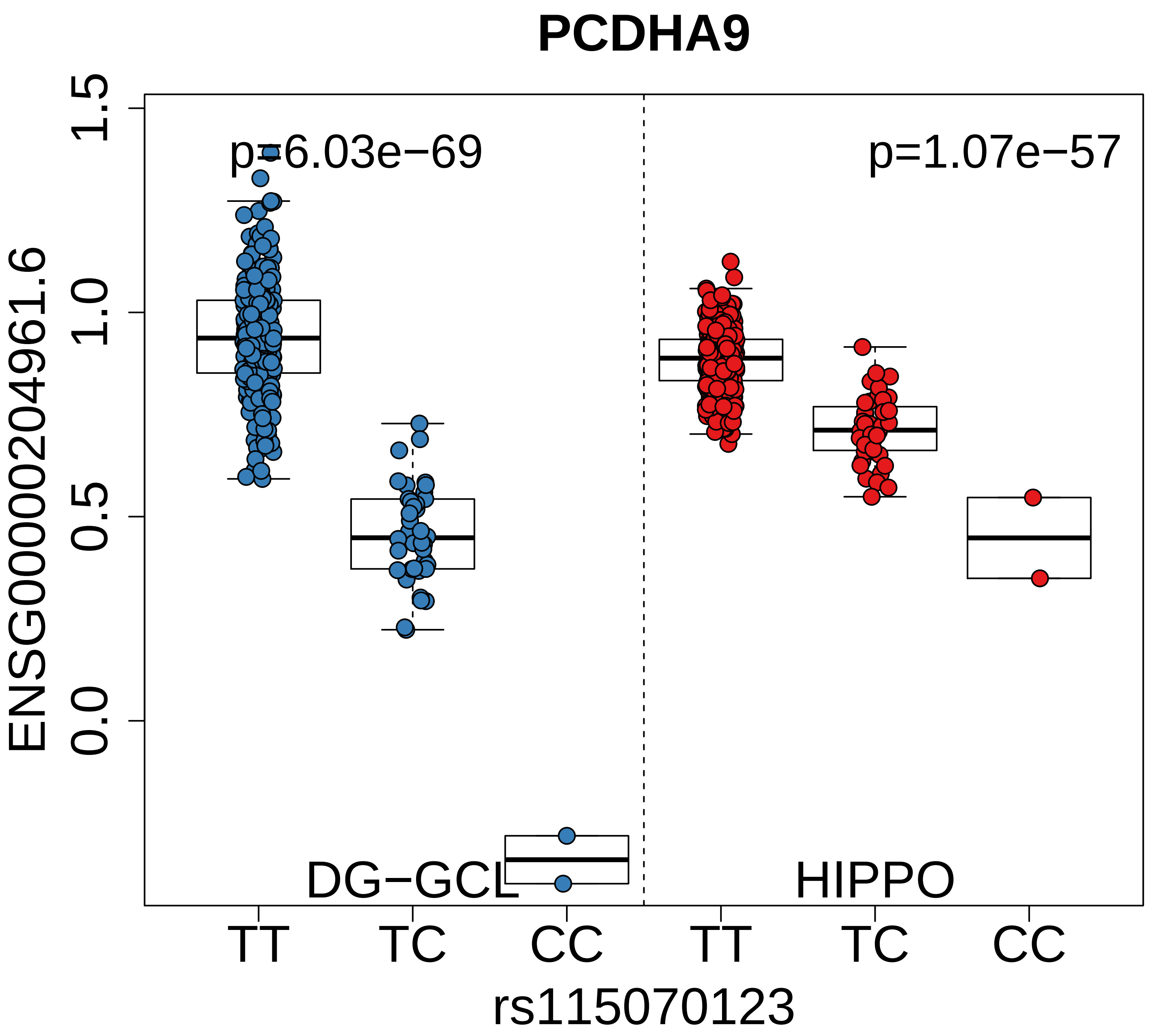

PCDHA9
p=6.03e−69
p=1.07e−57
DG−GCL
HIPPO
ENSG00000204961.6
1.5
1.0
0.5
0.0
TT
TC
CC
TT
TC
CC
rs115070123

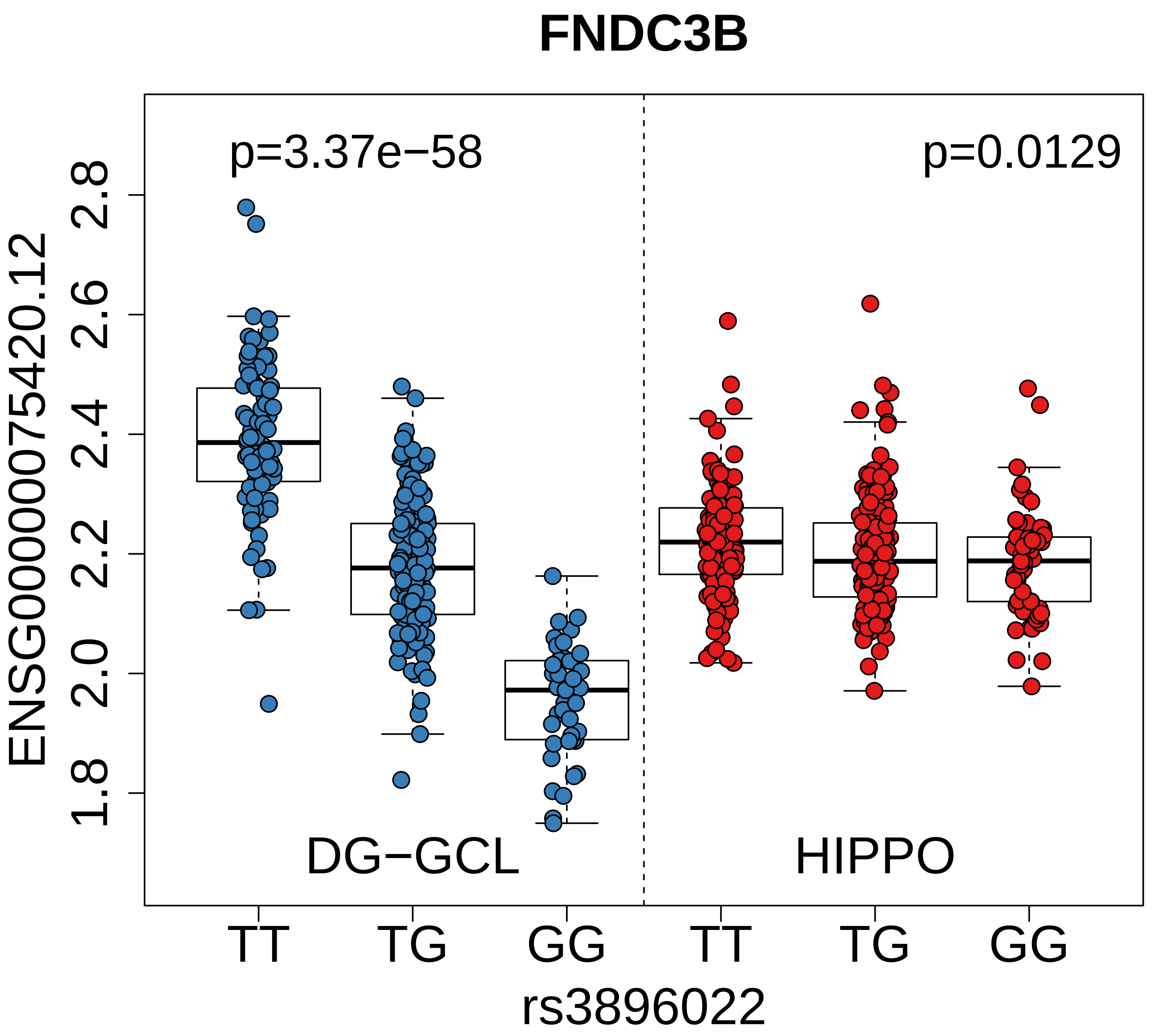

FNDC3B
p=3.37e−58
p=0.0129
ENSG00000075420.12
2.8
2.6
2.4
2.2
2.0
1.8
DG−GCL
HIPPO
TT
TG
GG
TT
TG
GG
rs3896022

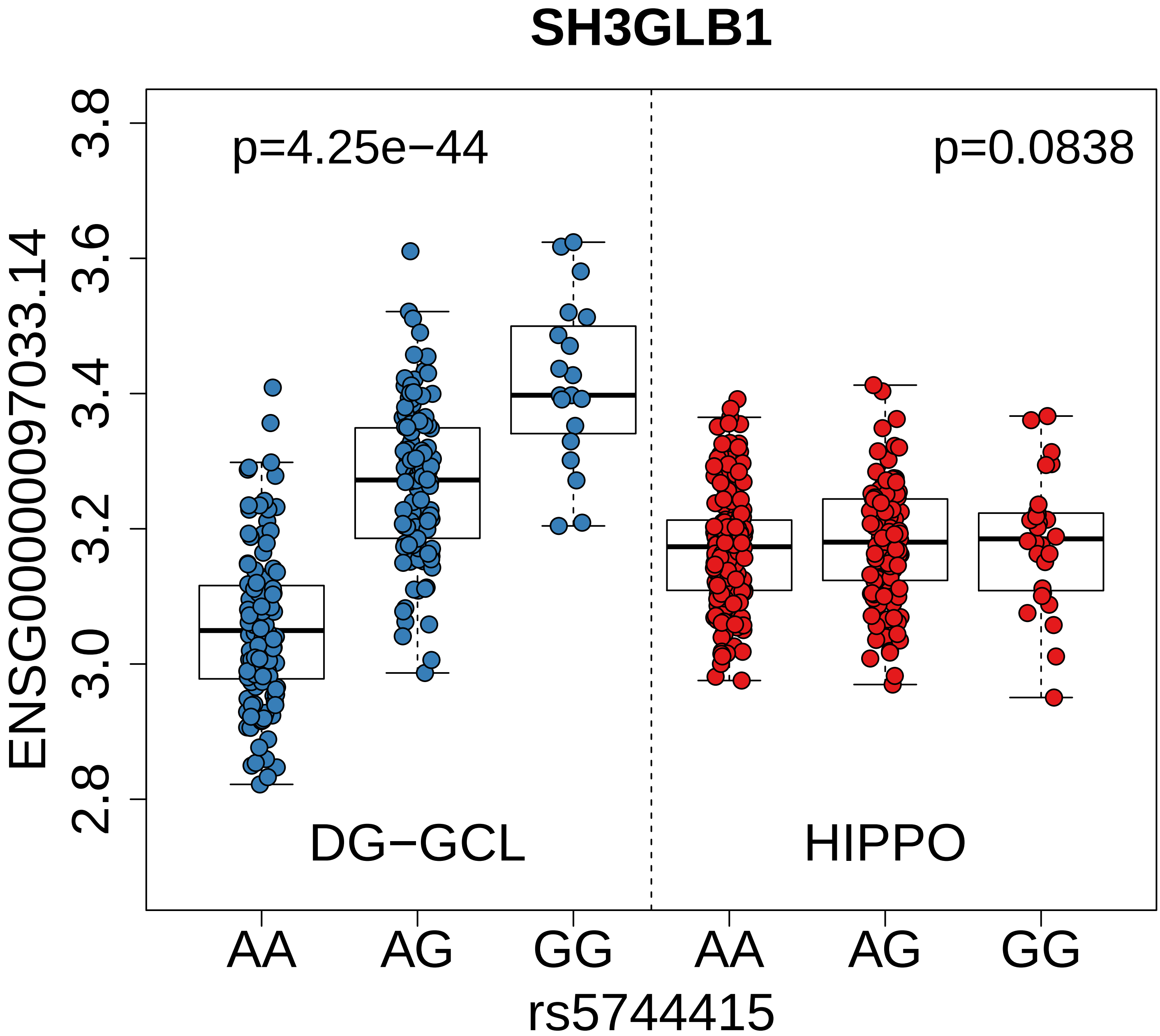
SH3GLB1
p=4.25e−44
p=0.0838
ENSG00000097033.14
3.8
3.6
3.4
3.2
3.0
2.8
DG−GCL
HIPPO
AA
AG
GG
AA
AG
GG
rs5744415

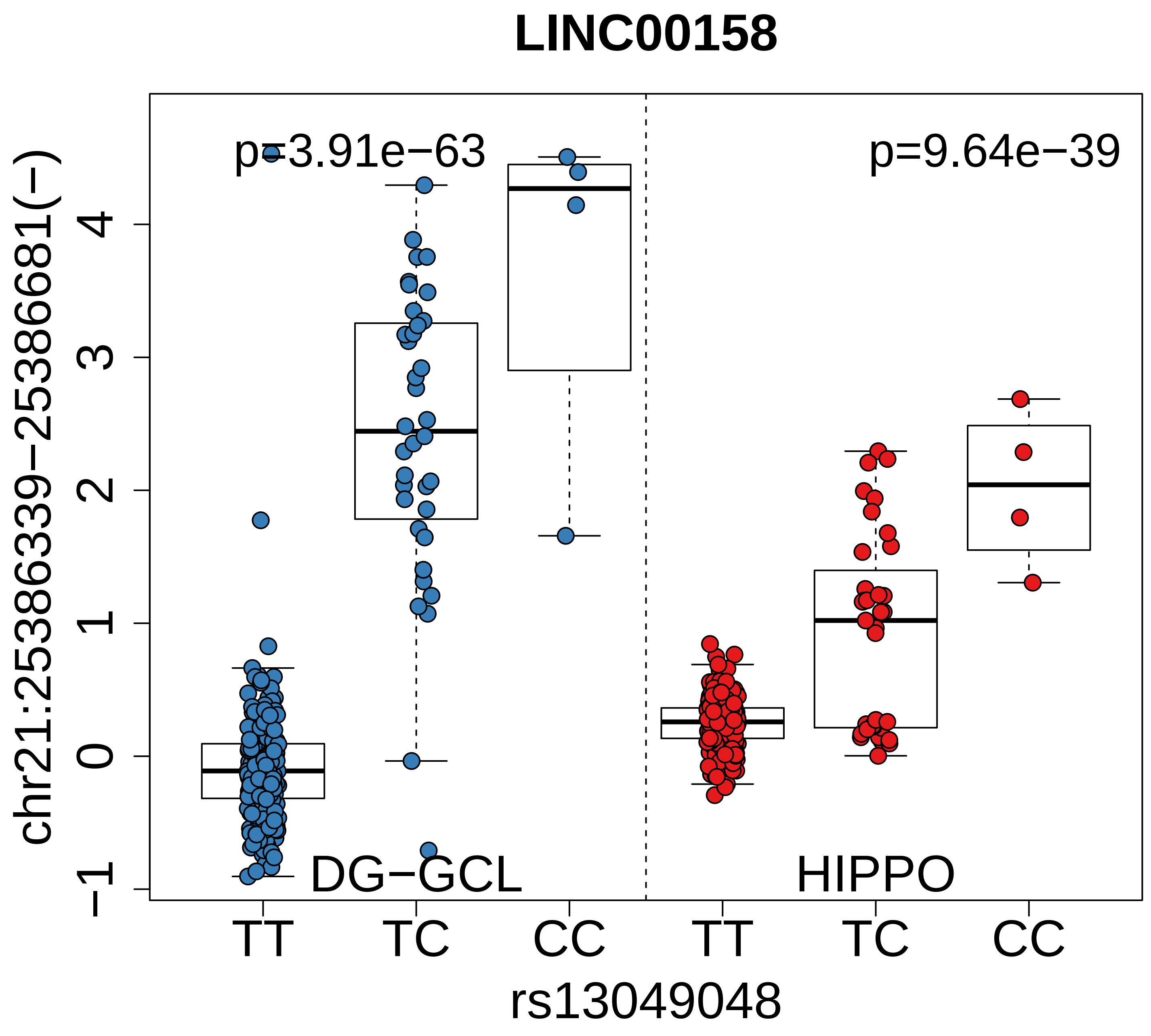

LINC00158
chr21:25386339−25386681(−)
p=3.91e−63
p=9.64e−39
DG−GCL
HIPPO
−1
0
1
2
3
4
TT
TC
CC
TT
TC
CC
rs13049048

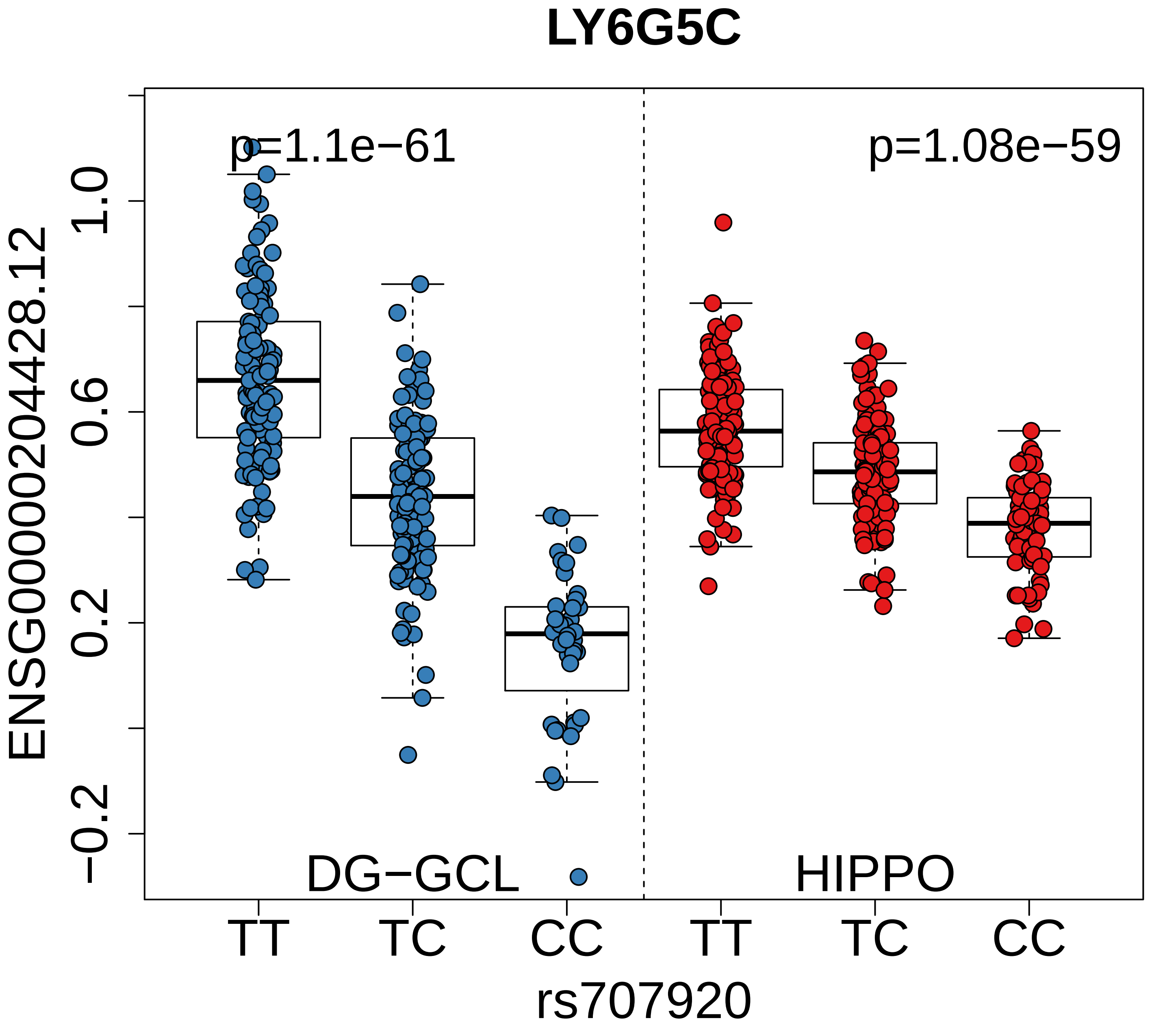

LY6G5C
p=1.1e−61
p=1.08e−59
ENSG00000204428.12
1.0
0.6
0.2
−0.2
DG−GCL
HIPPO
TT
TC
CC
TT
TC
CC
rs707920

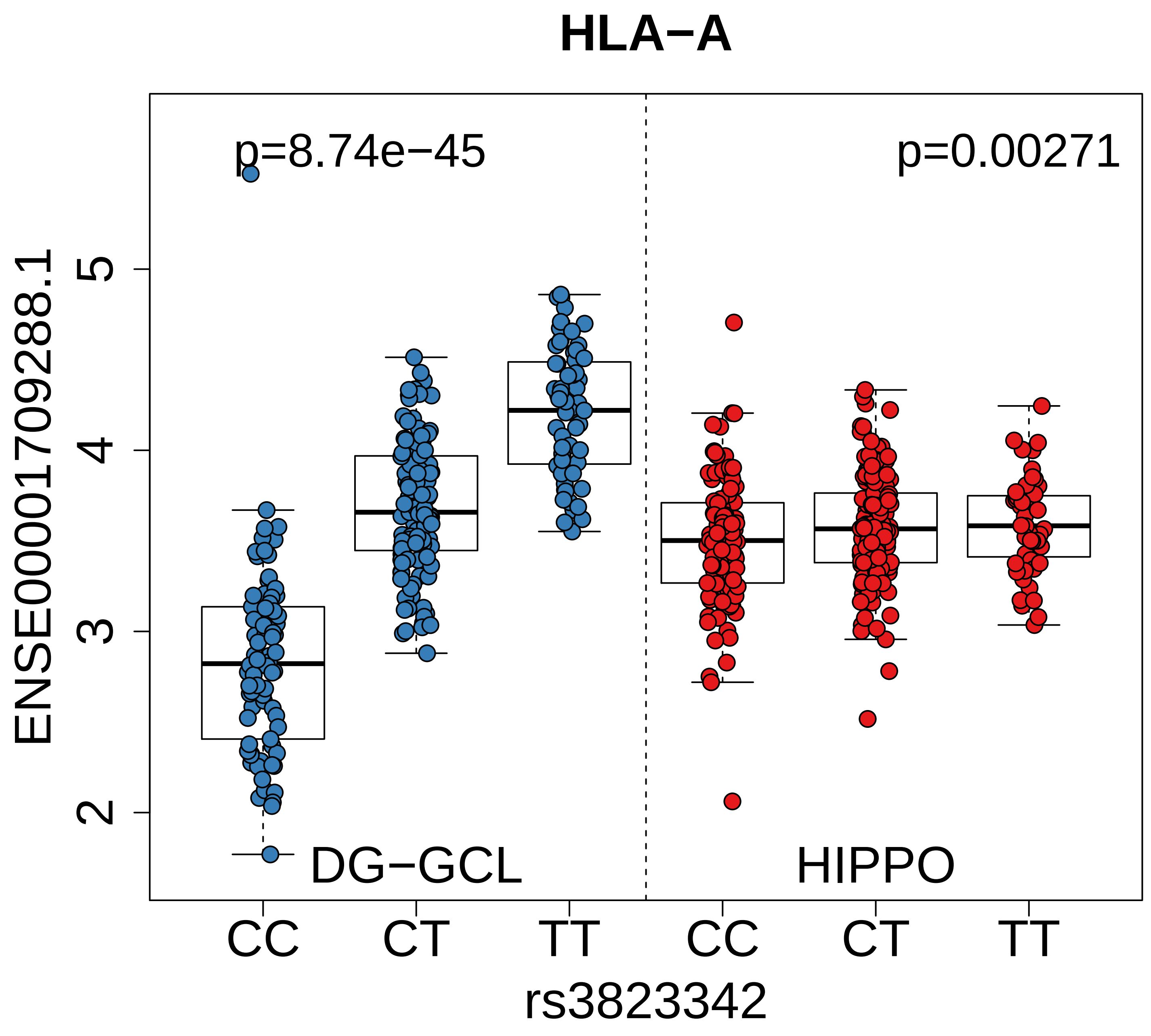
HLA−A
p=8.74e−45
p=0.00271
ENSE00001709288.1
5
4
3
2
DG−GCL
HIPPO
CC
CT
TT
CC
CT
TT
rs3823342

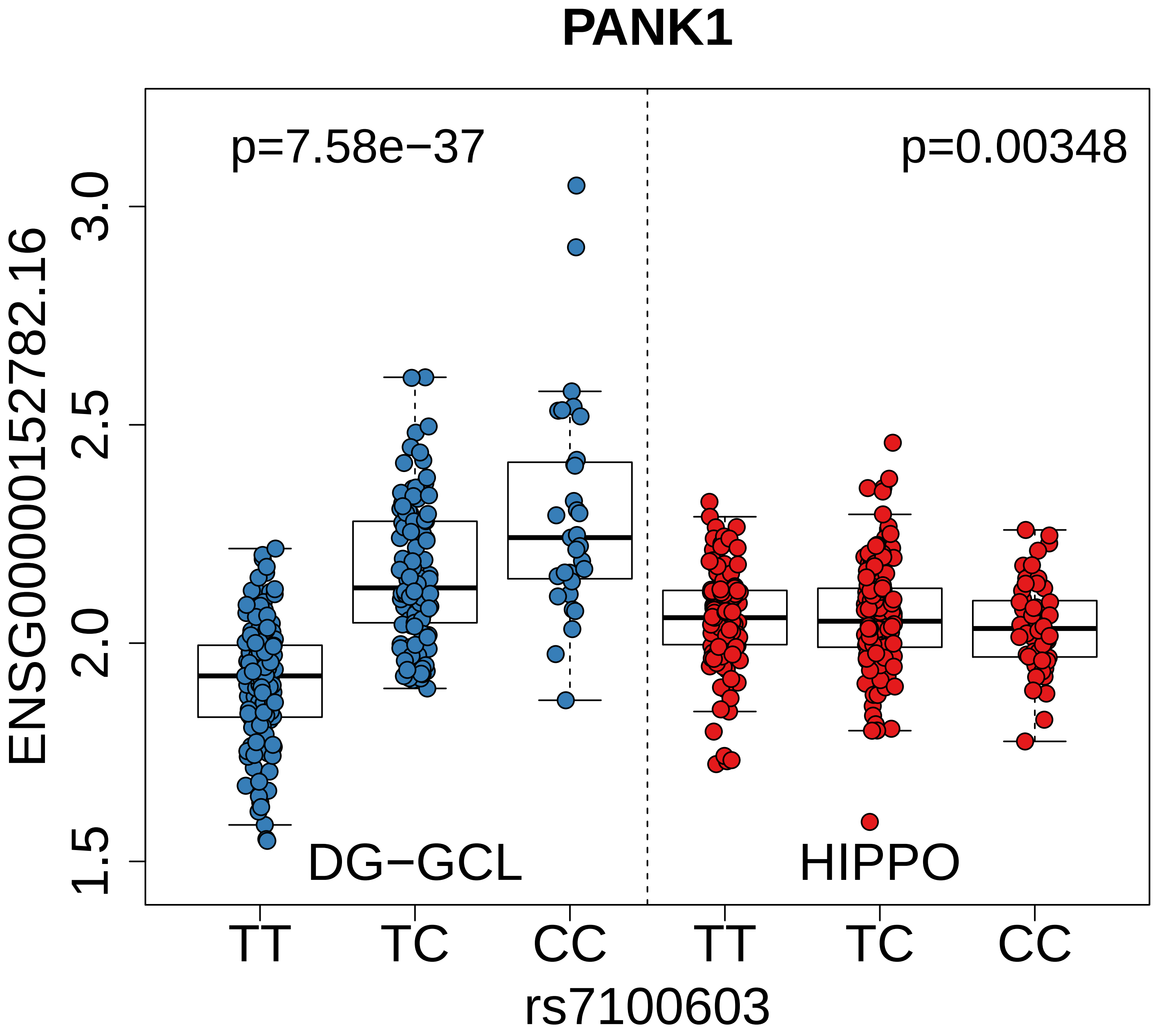
PANK1
p=7.58e−37
p=0.00348
ENSG00000152782.16
3.0
2.5
2.0
1.5
DG−GCL
HIPPO
TT
TC
CC
TT
TC
CC
rs7100603

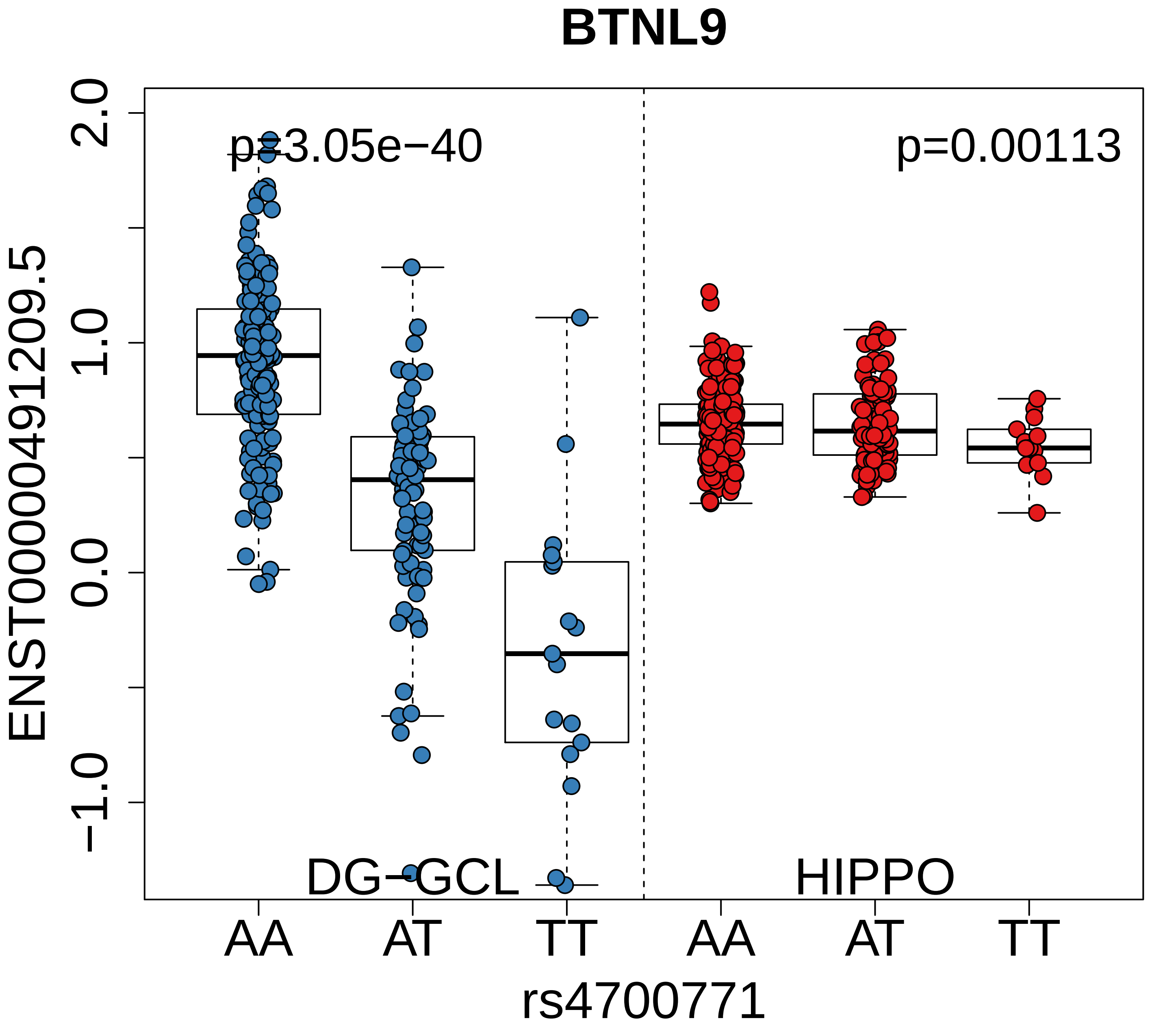
BTNL9
p=3.05e−40
p=0.00113
ENST00000491209.5
2.0
1.0
0.0
−1.0
DG−GCL
HIPPO
AA
AT
TT
AA
AT
TT
rs4700771

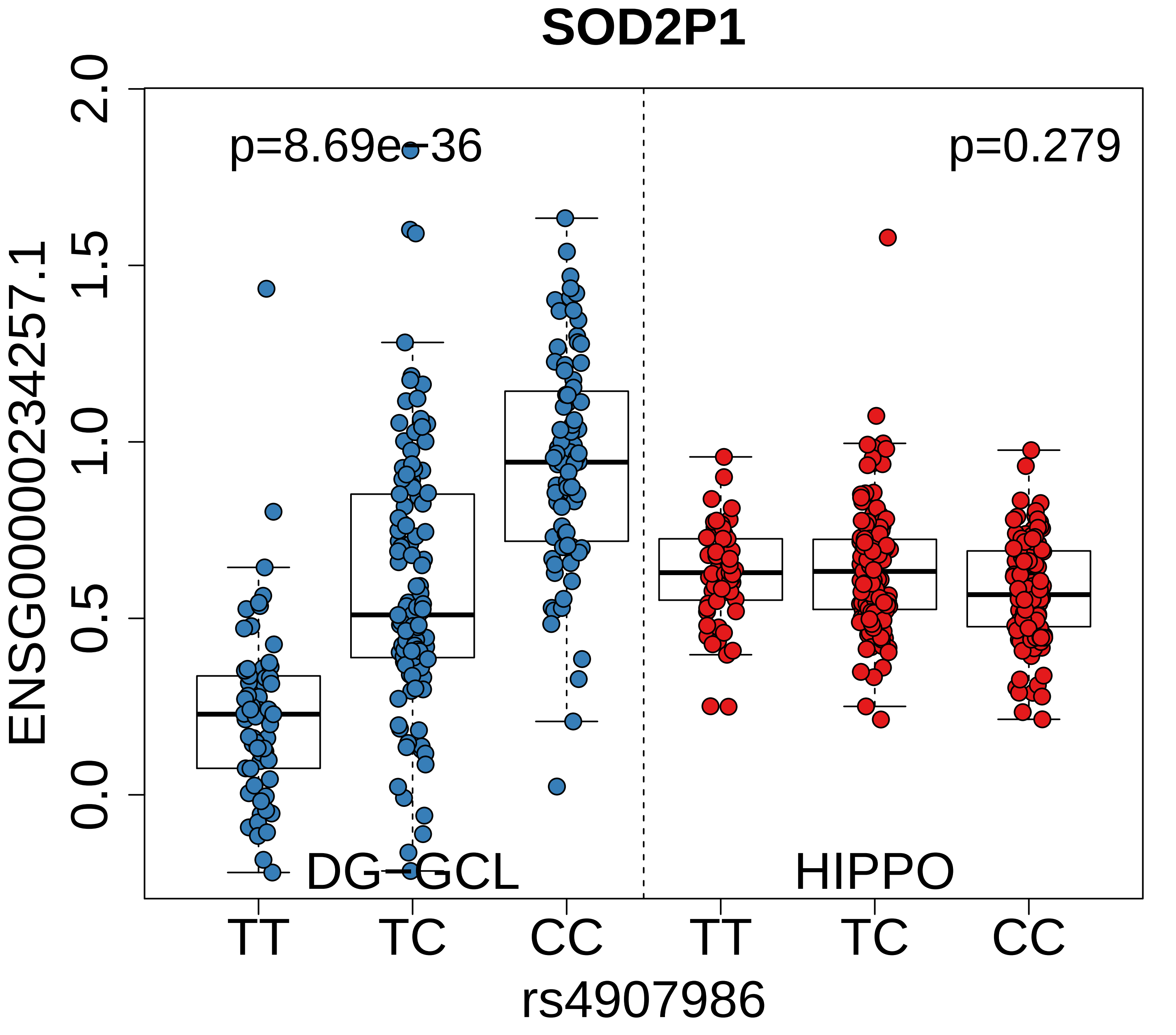
SOD2P1
p=8.69e−36
p=0.279
ENSG00000234257.1
2.0
1.5
1.0
0.5
0.0
DG−GCL
HIPPO
TT
TC
CC
TT
TC
CC
rs4907986

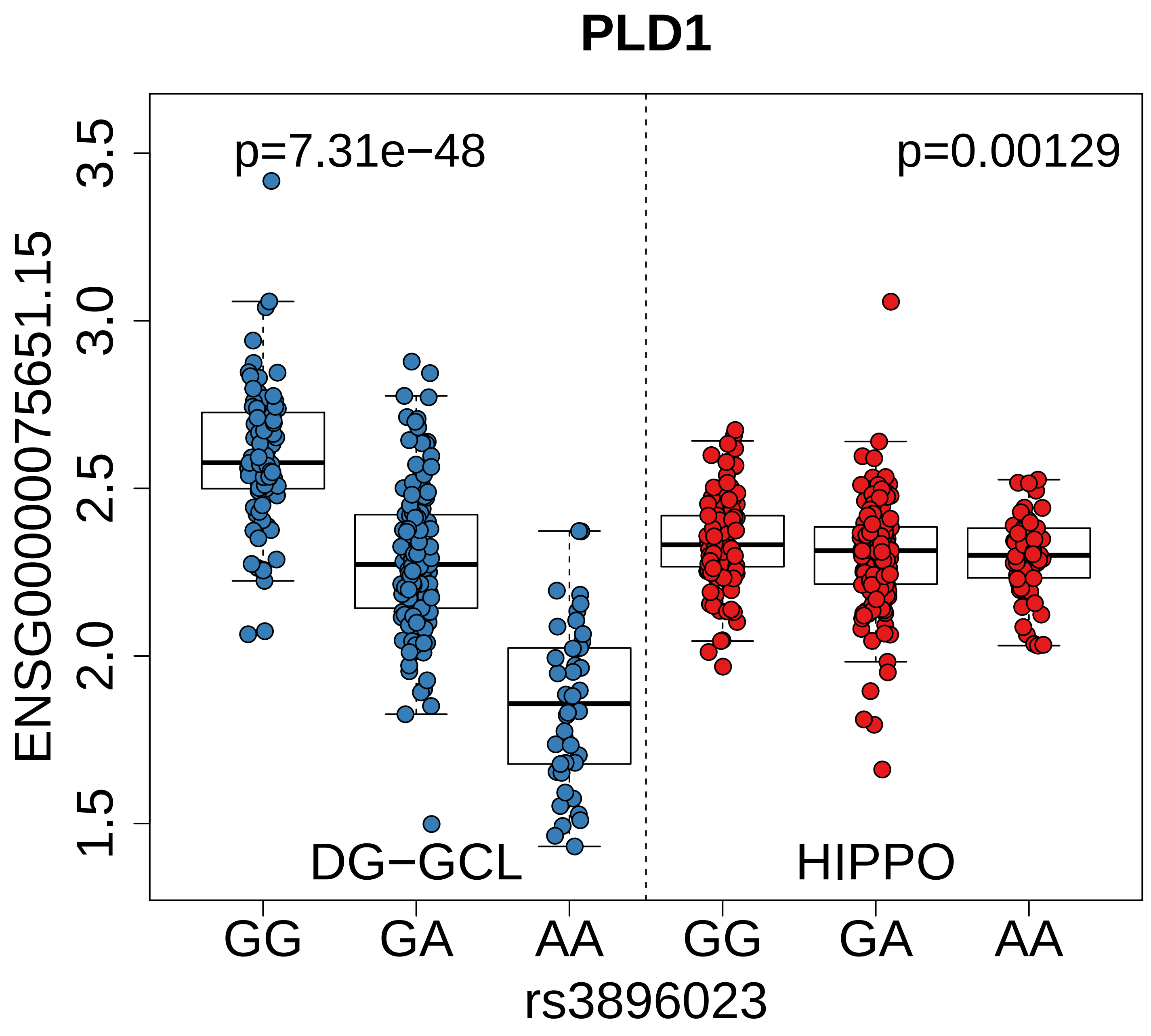

PLD1
p=7.31e−48
p=0.00129
ENSG00000075651.15
3.5
3.0
2.5
2.0
1.5
DG−GCL
HIPPO
GG
GA
AA
GG
GA
AA
rs3896023

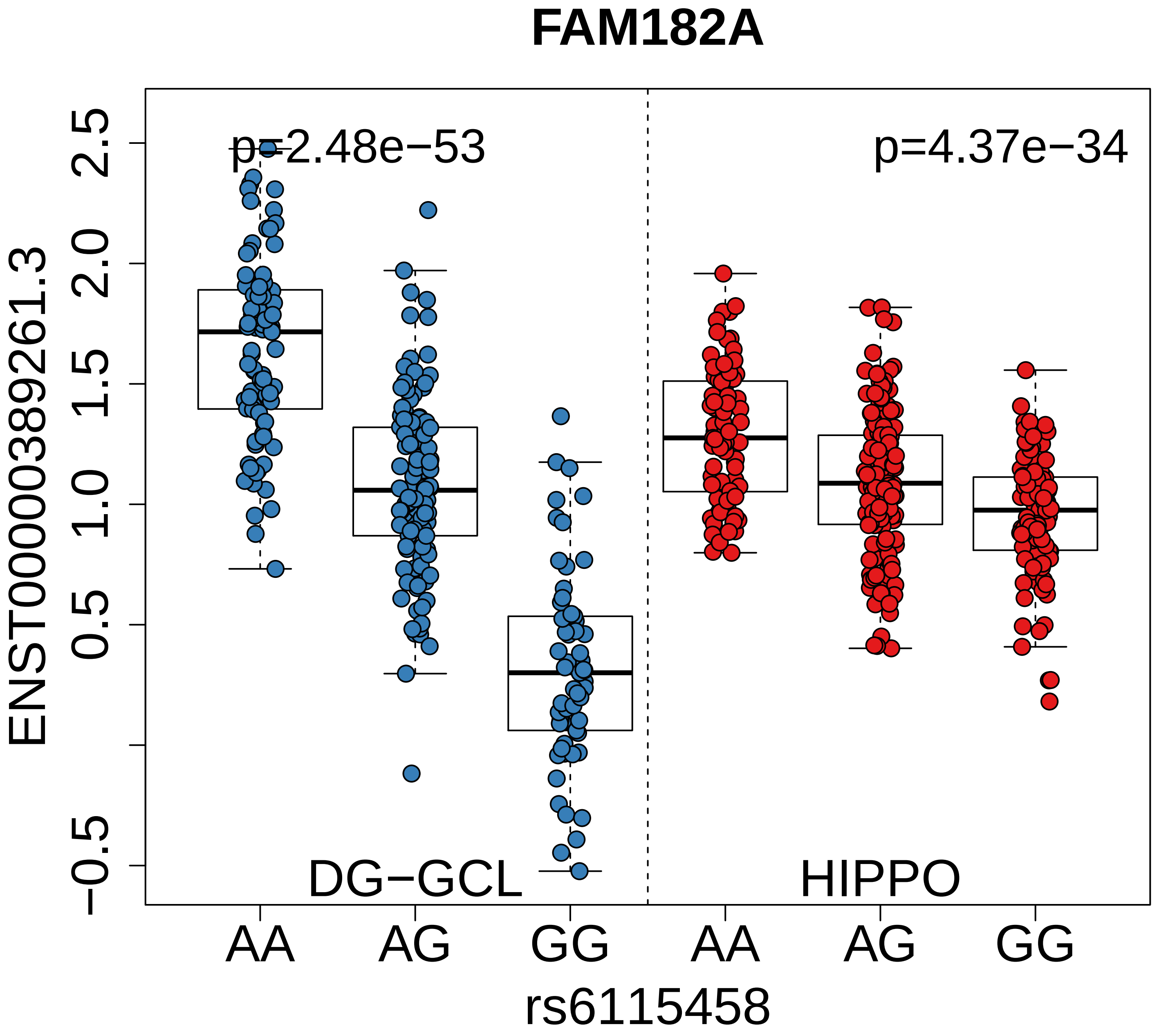
FAM182A
p=2.48e−53
p=4.37e−34
ENST00000389261.3
2.5
2.0
1.5
1.0
0.5
−0.5
DG−GCL
HIPPO
AA
AG
GG
AA
AG
GG
rs6115458

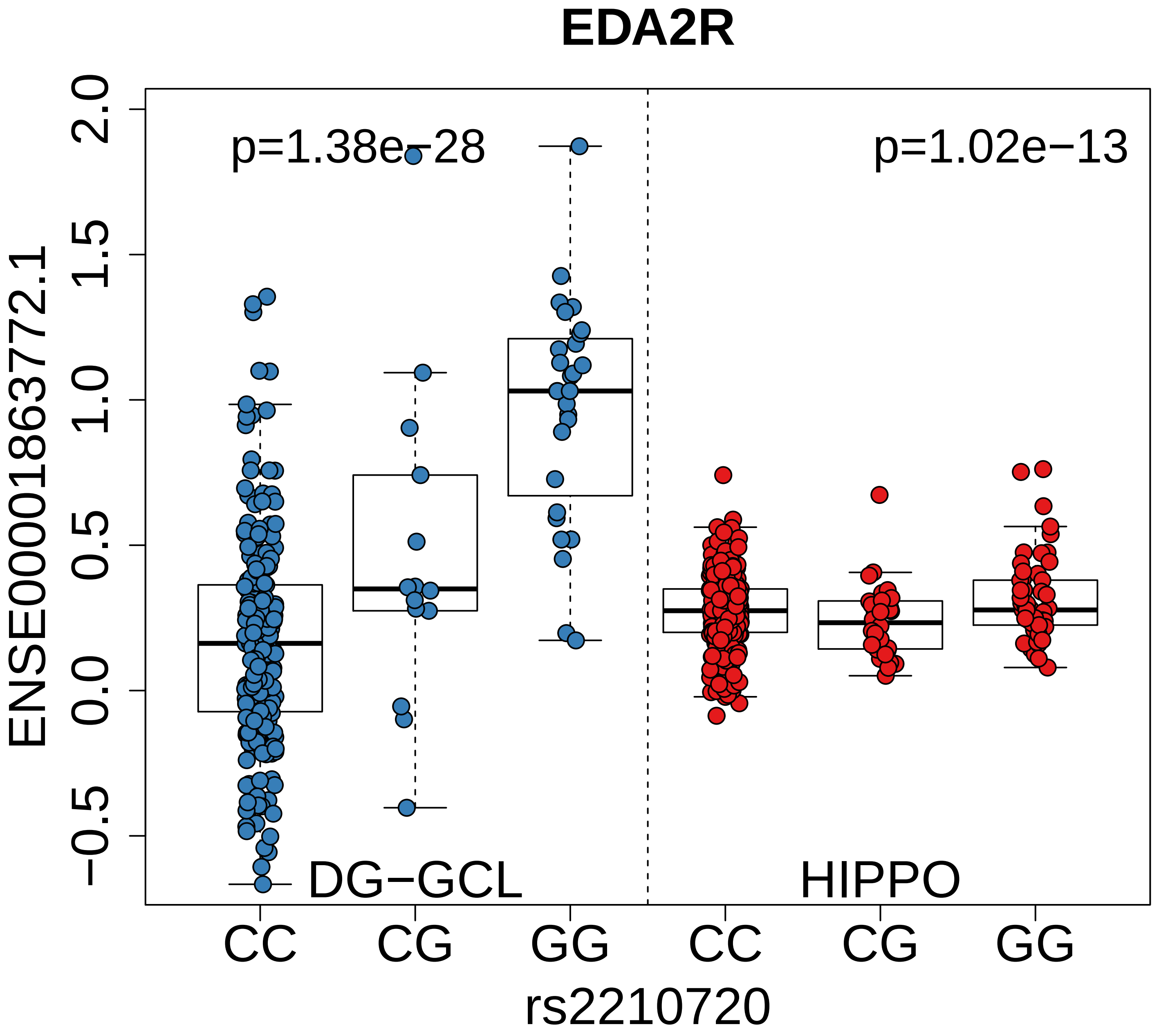
EDA2R
p=1.38e−28
p=1.02e−13
ENSE00001863772.1
2.0
1.5
1.0
0.5
0.0
−0.5
DG−GCL
HIPPO
CC
CG
GG
CC
CG
GG
rs2210720

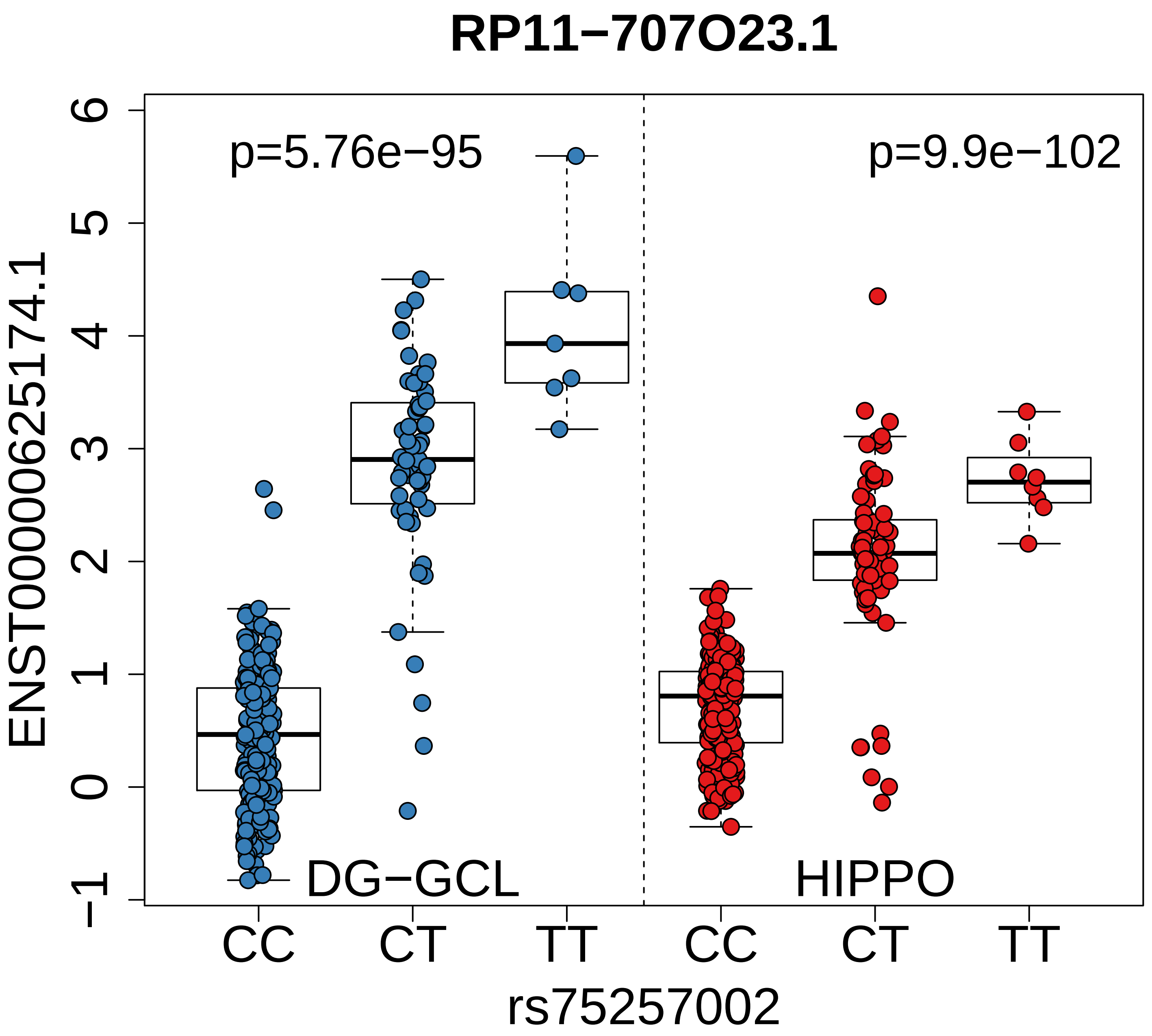

RP11−707O23.1
p=5.76e−95
p=9.9e−102
ENST00000625174.1
6
5
4
3
2
1
0
−1
DG−GCL
HIPPO
CC
CT
TT
CC
CT
TT
rs75257002

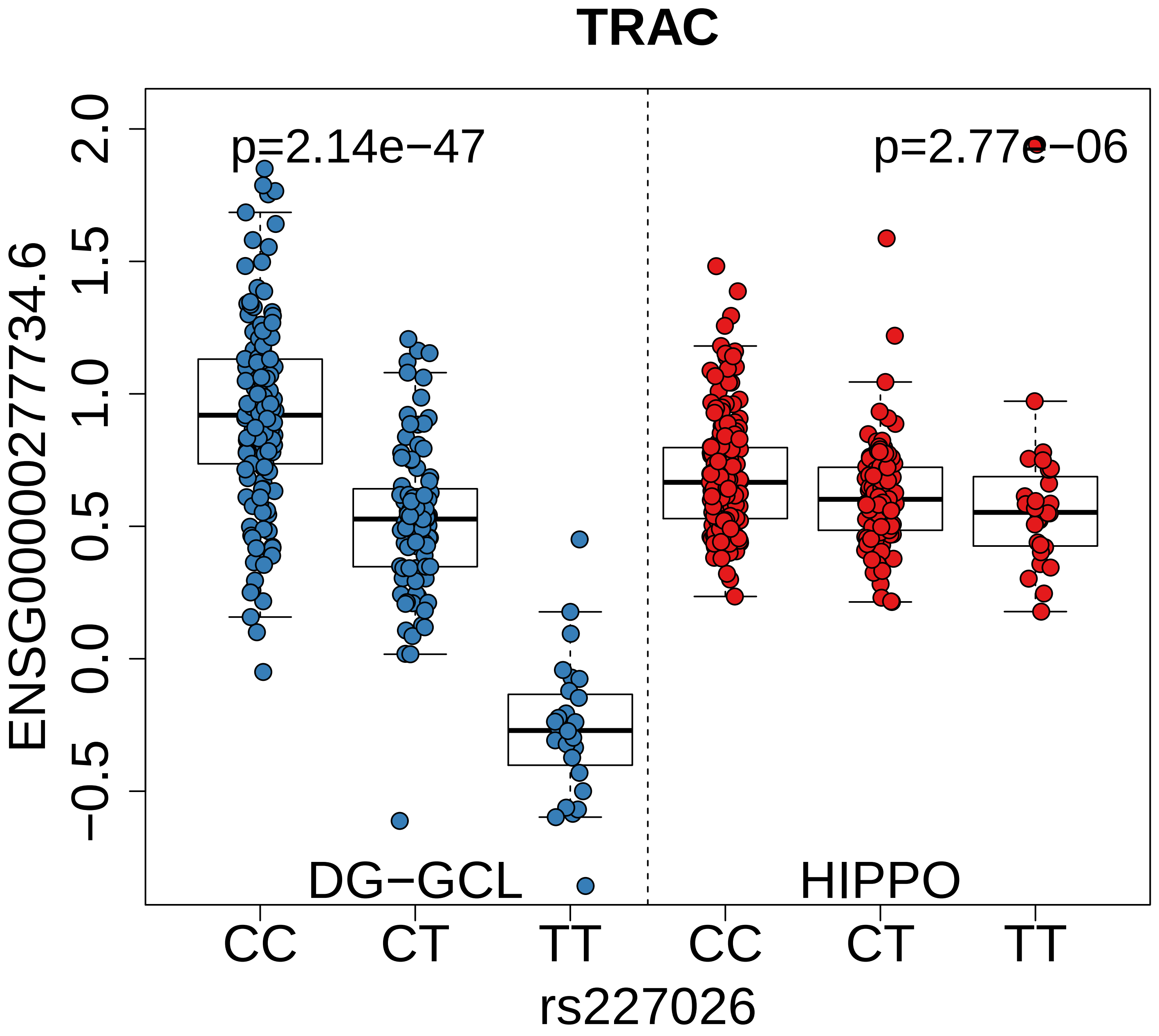
TRAC
p=2.14e−47
p=2.77e−06
ENSG00000277734.6
2.0
1.5
1.0
0.5
0.0
−0.5
DG−GCL
HIPPO
CC
CT
TT
CC
CT
TT
rs227026

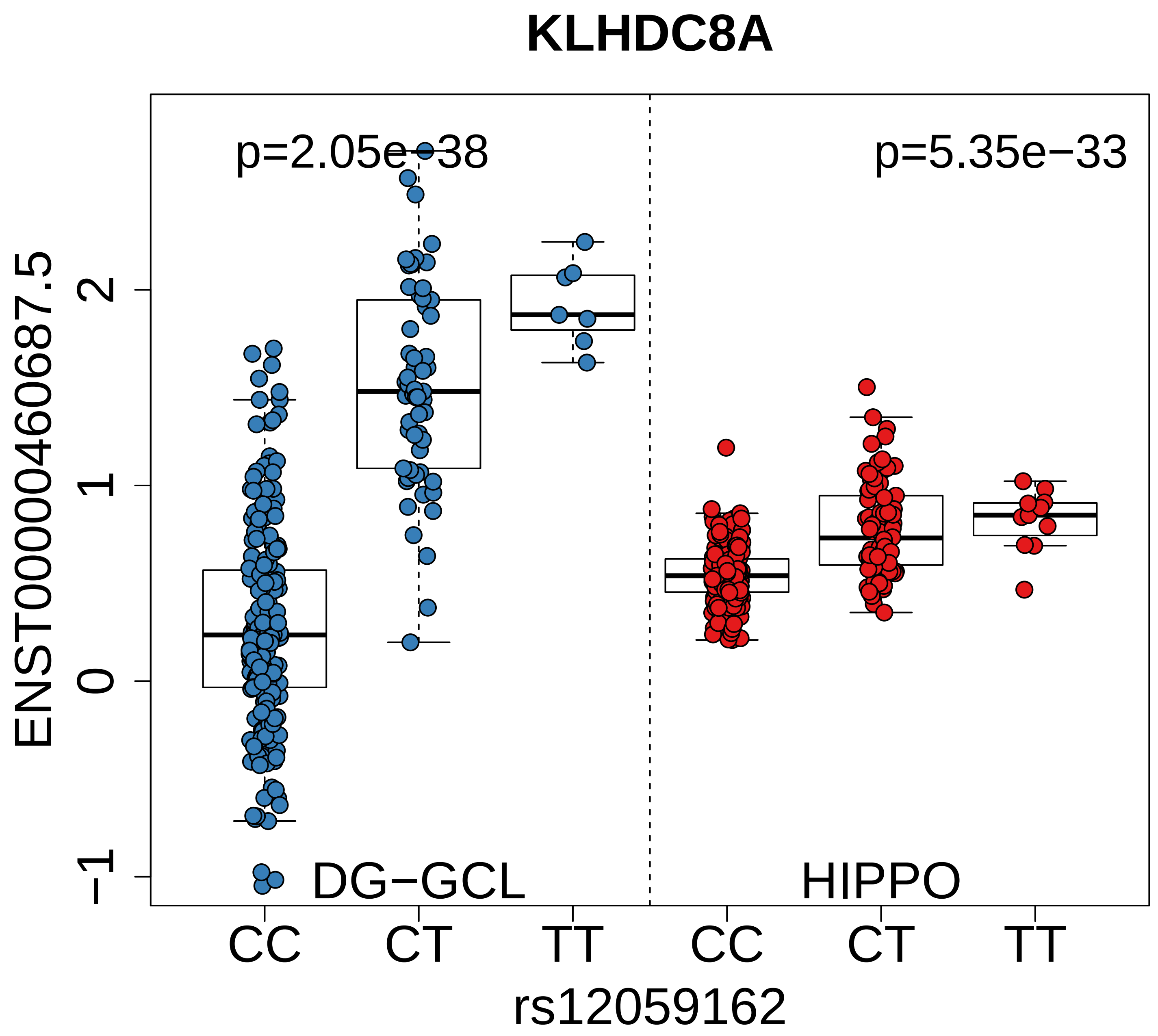

KLHDC8A
p=2.05e−38
p=5.35e−33
ENST00000460687.5
2
1
0
−1
DG−GCL
HIPPO
CC
CT
TT
CC
CT
TT
rs12059162

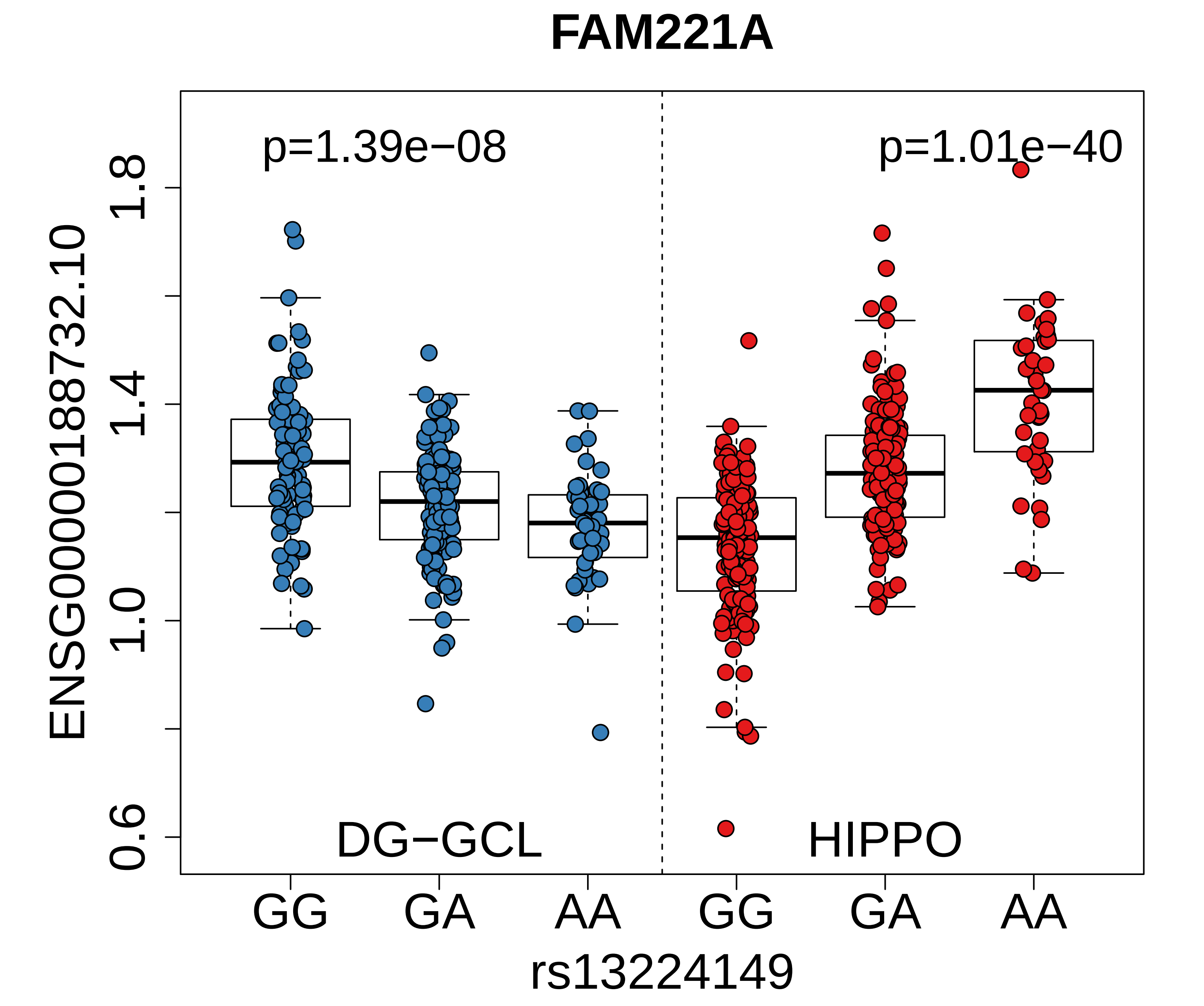
FAM221A
p=1.39e−08
p=1.01e−40
DG−GCL
HIPPO
ENSG00000188732.10
0.6
1.0
1.4
1.8
GG
GA
AA
GG
GA
AA
rs13224149

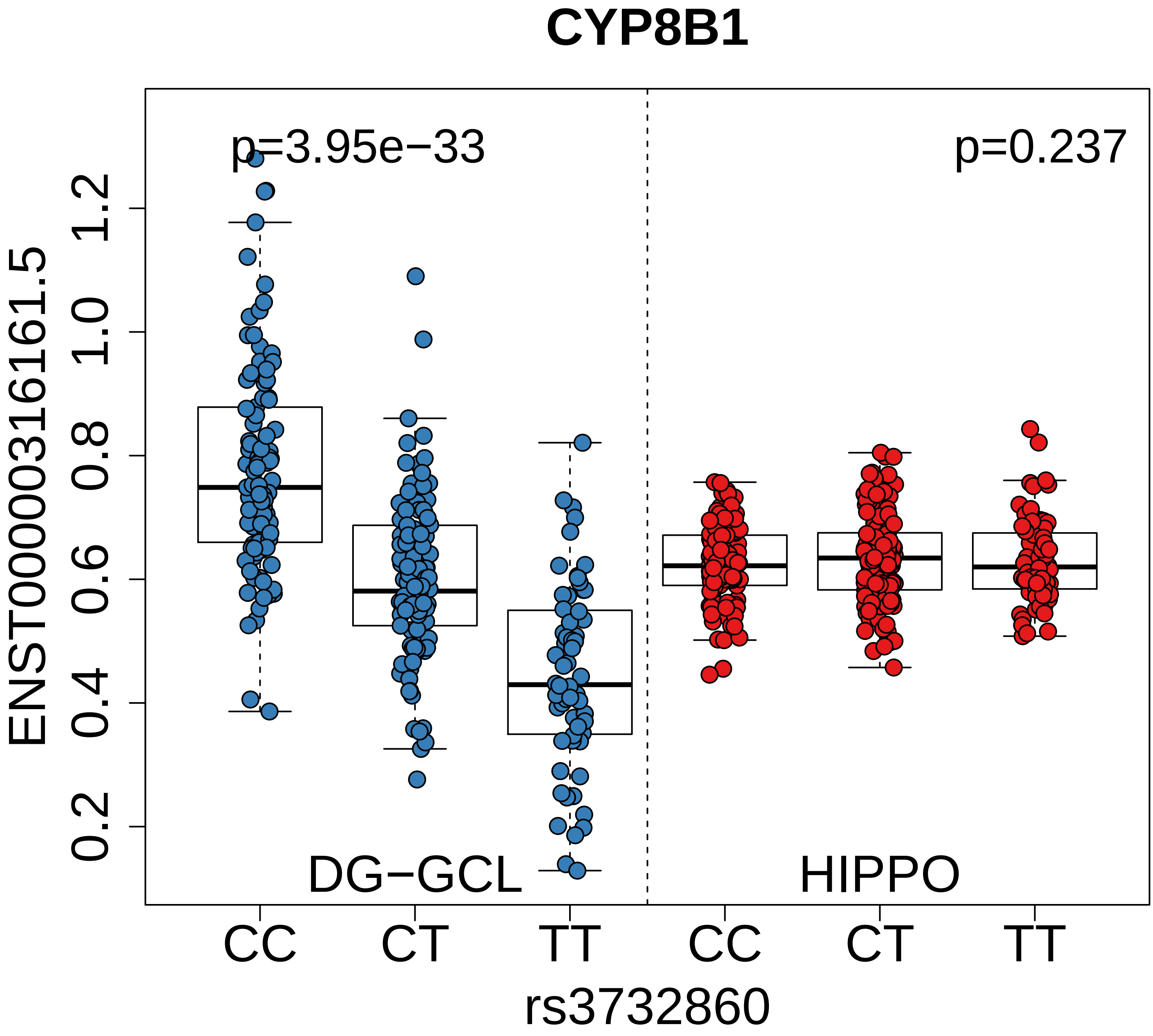

CYP8B1
p=3.95e−33
p=0.237
ENST00000316161.5
1.2
1.0
0.8
0.6
0.4
0.2
DG−GCL
HIPPO
CC
CT
TT
CC
CT
TT
rs3732860

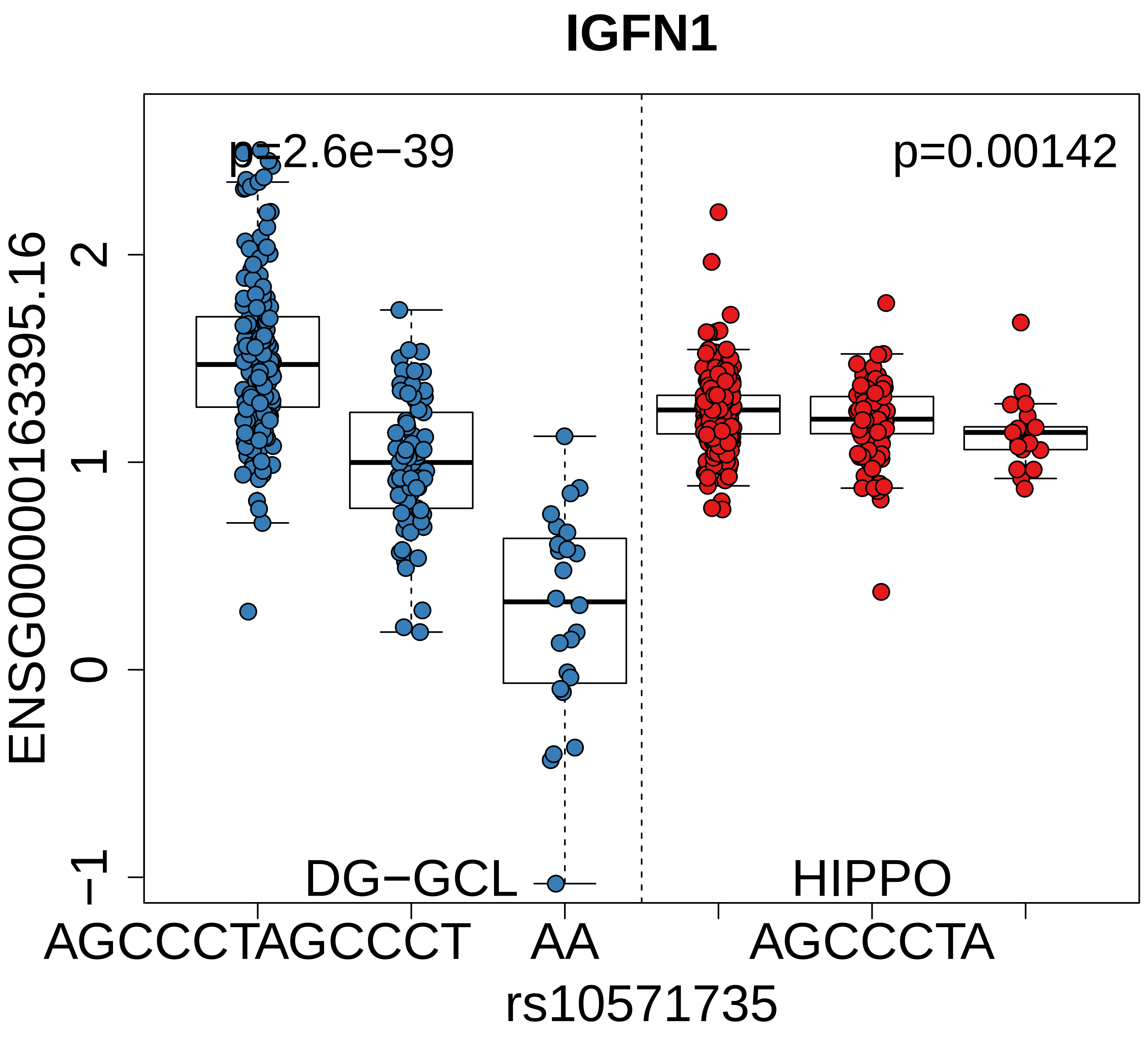

IGFN1
p=2.6e−39
p=0.00142
ENSG00000163395.16
2
1
0
−1
DG−GCL
HIPPO
AGCCCTAGCCCT
AA
AGCCCTA
rs10571735

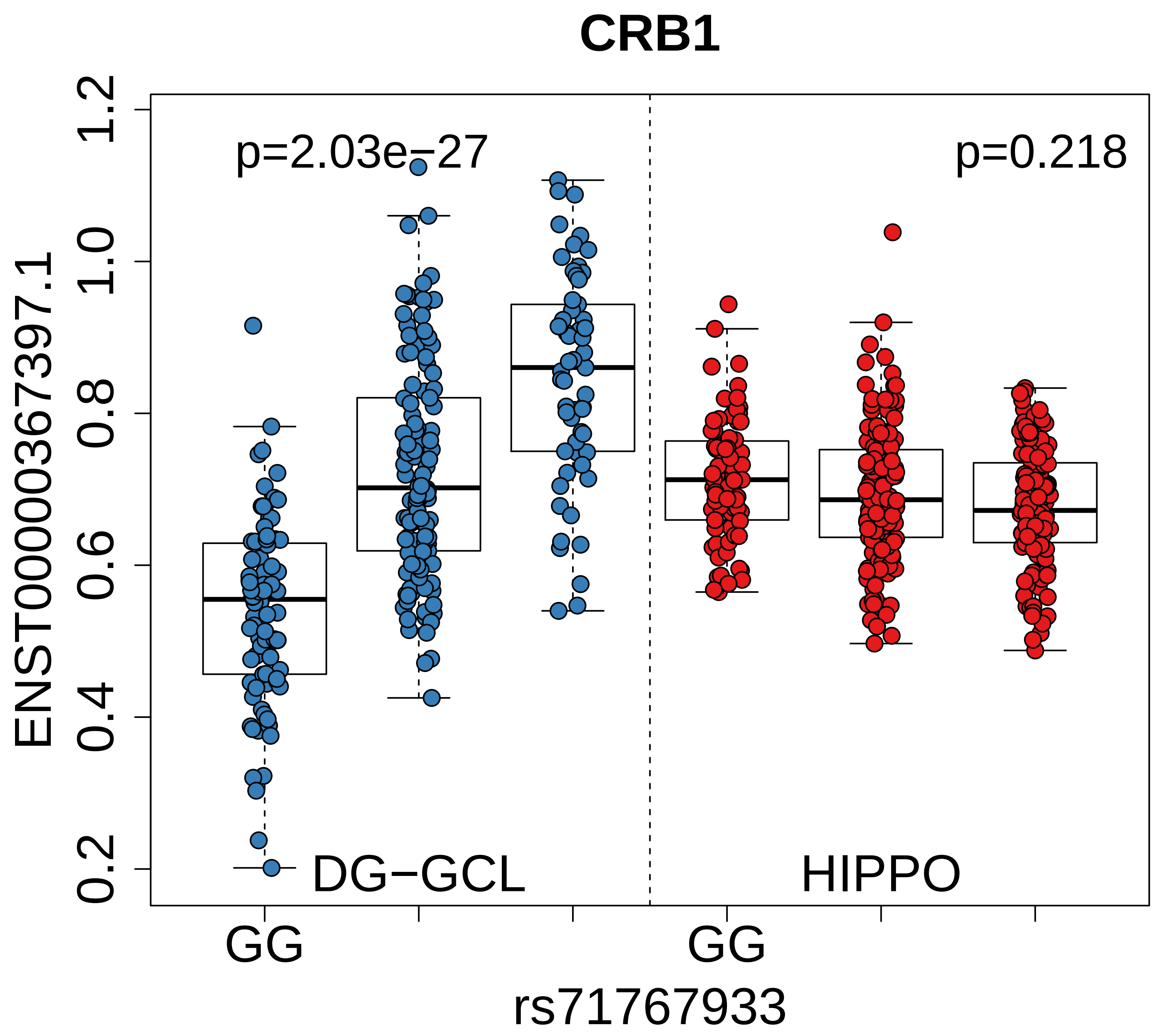
CRB1
p=2.03e−27
p=0.218
ENST00000367397.1
1.2
1.0
0.8
0.6
0.4
0.2
DG−GCL
HIPPO
GG
GG
rs71767933

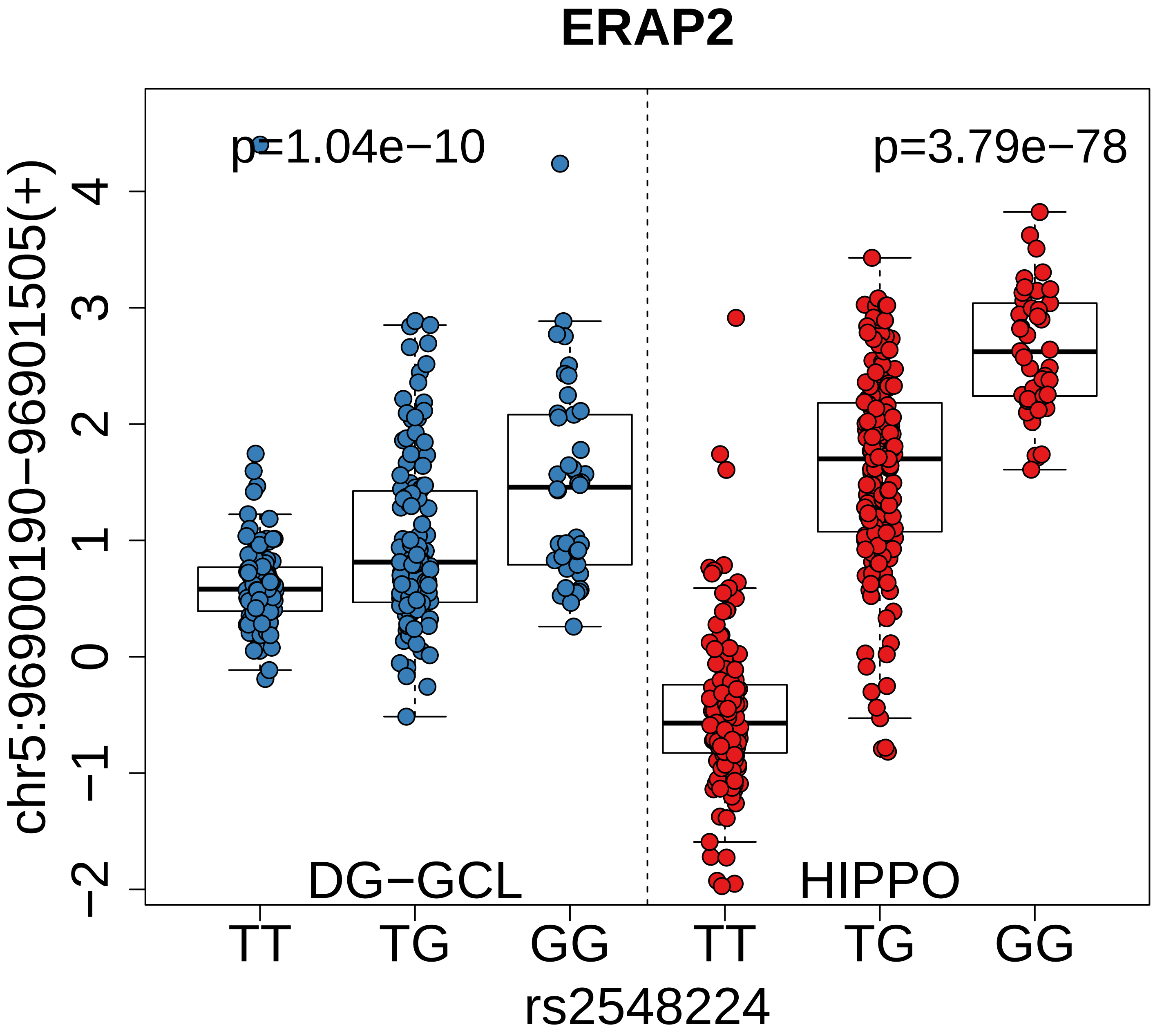

ERAP2
p=1.04e−10
p=3.79e−78
chr5:96900190−96901505(+)
DG−GCL
HIPPO
TT
TG
GG
TT
TG
GG
rs2548224
4
3
2
1
0
−1
−2

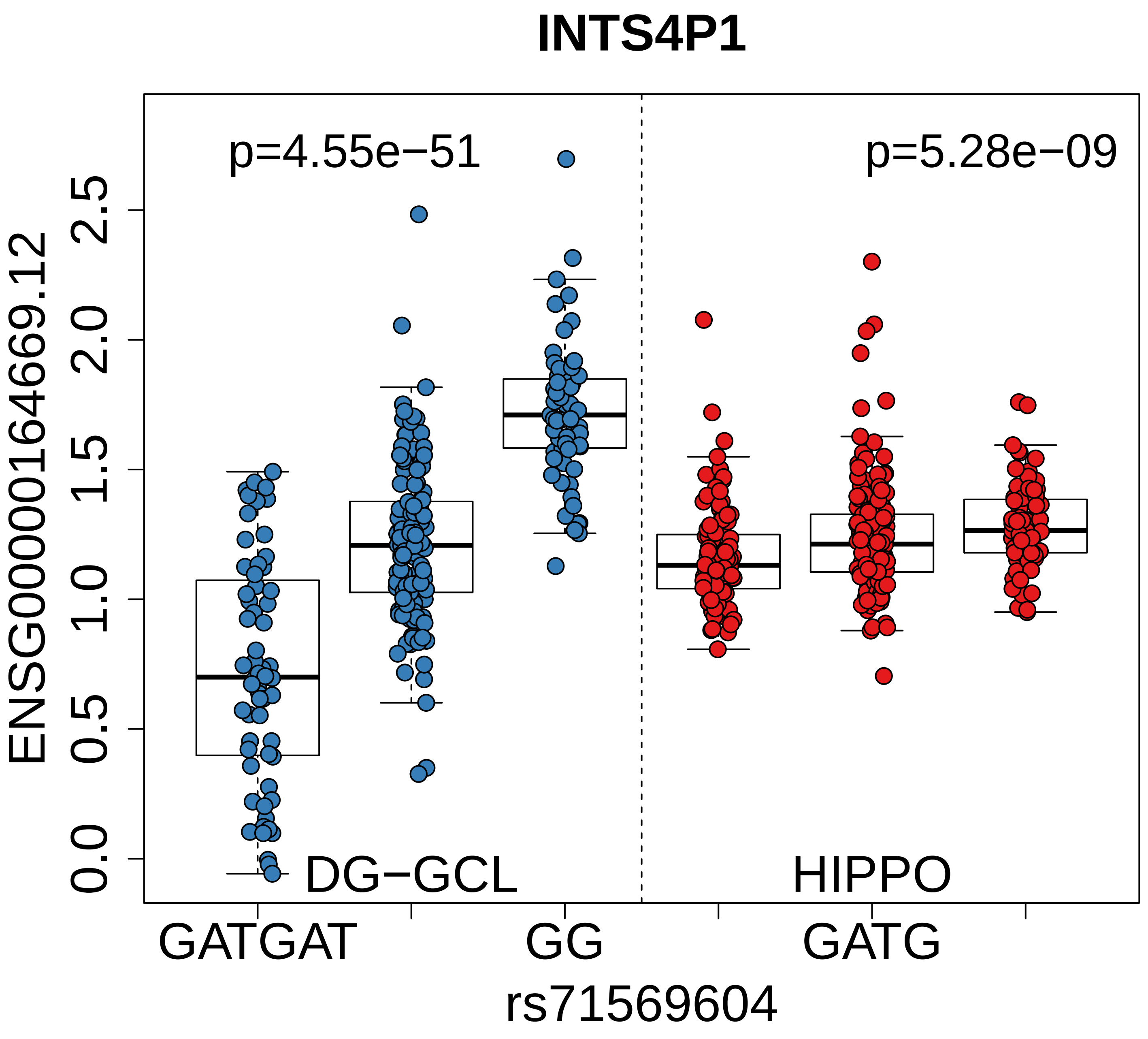

INTS4P1
p=4.55e−51
p=5.28e−09
ENSG00000164669.12
2.5
2.0
1.5
1.0
0.5
0.0
DG−GCL
HIPPO
GATGAT
GG
GATG
rs71569604

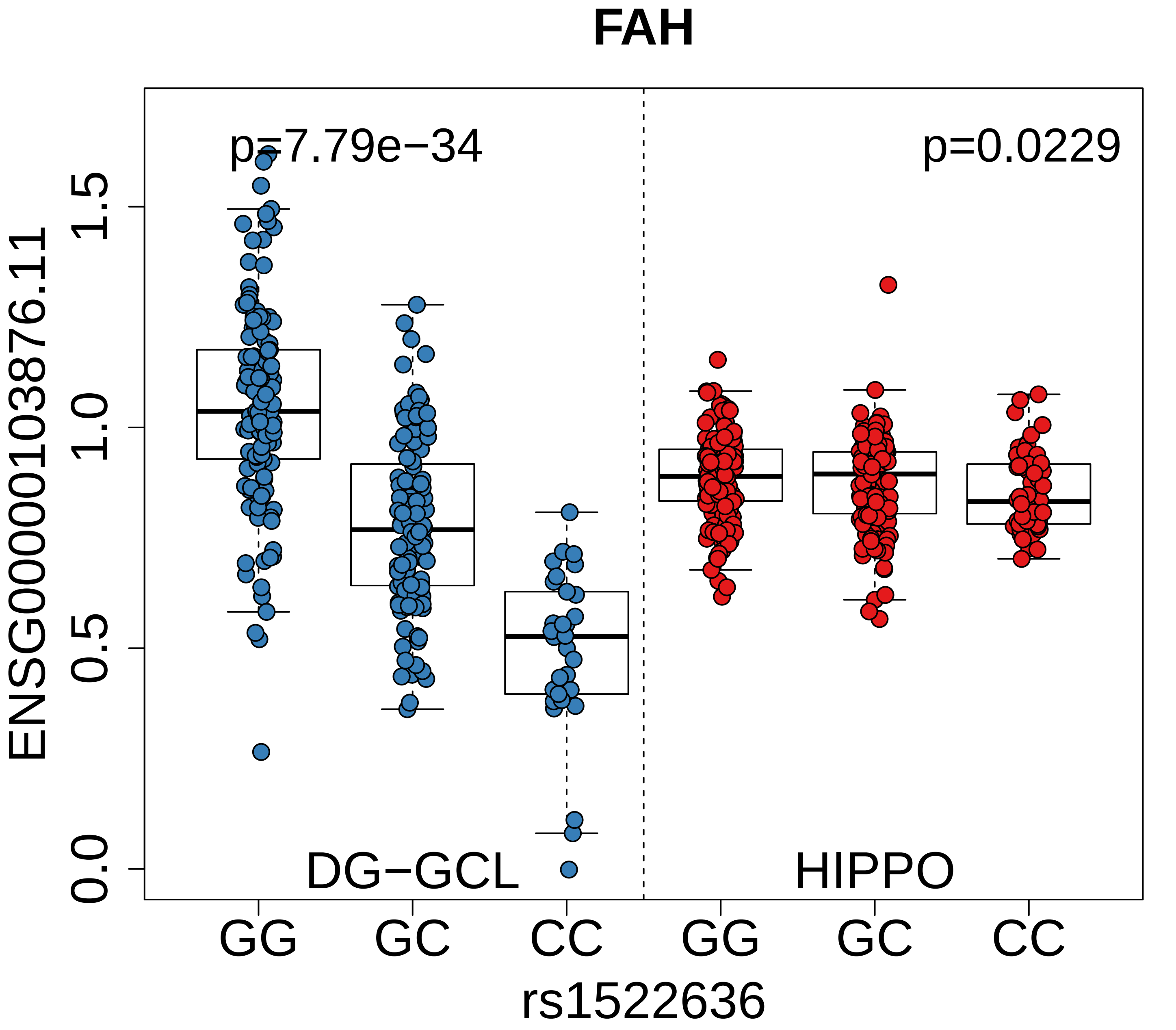

FAH
p=7.79e−34
p=0.0229
ENSG00000103876.11
0.0
0.5
1.0
1.5
DG−GCL
HIPPO
GG
GC
CC
GG
GC
CC
rs1522636

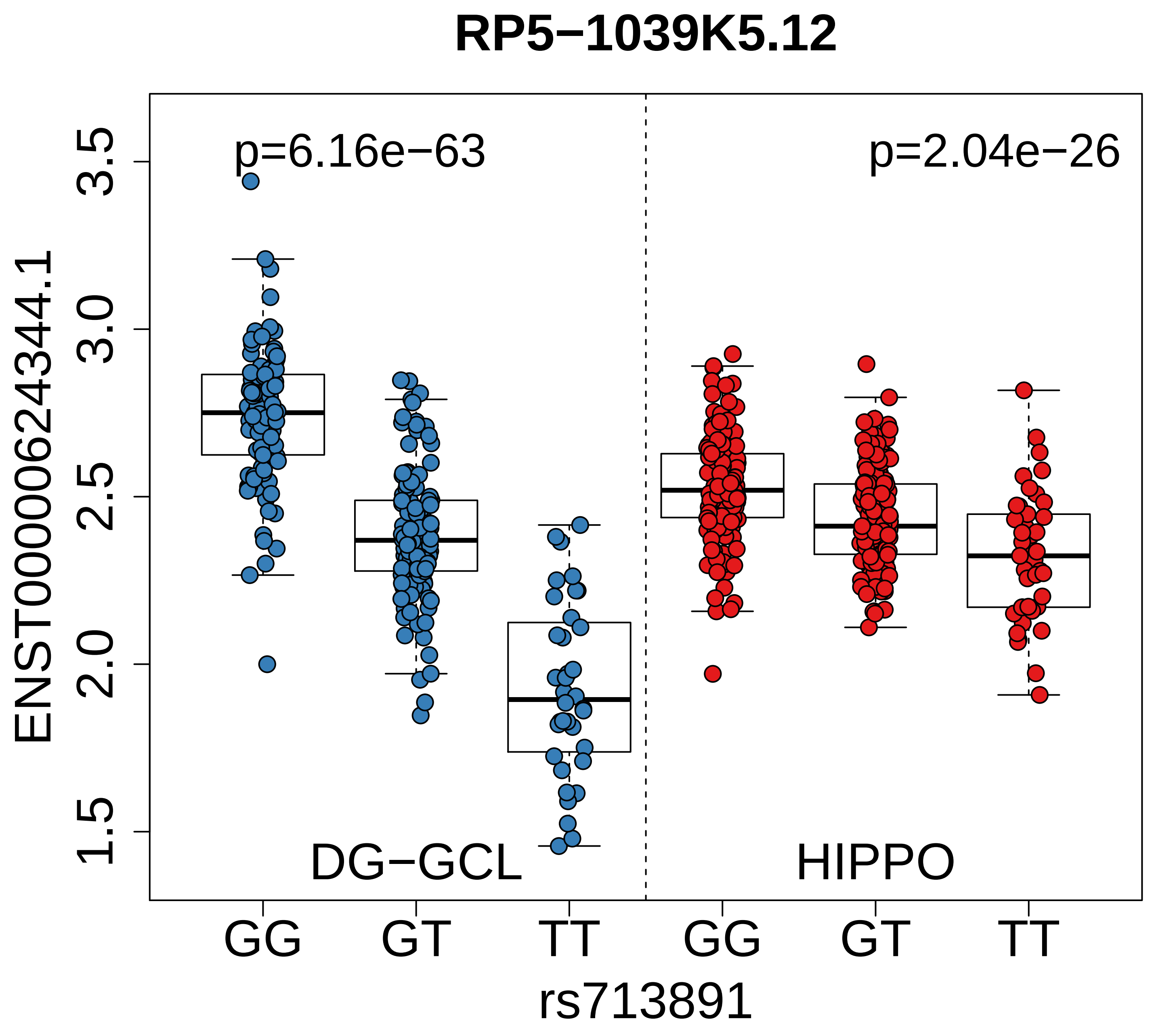
RP5−1039K5.12
p=6.16e−63
p=2.04e−26
ENST00000624344.1
3.5
3.0
2.5
2.0
1.5
DG−GCL
HIPPO
GG
GT
TT
GG
GT
TT
rs713891

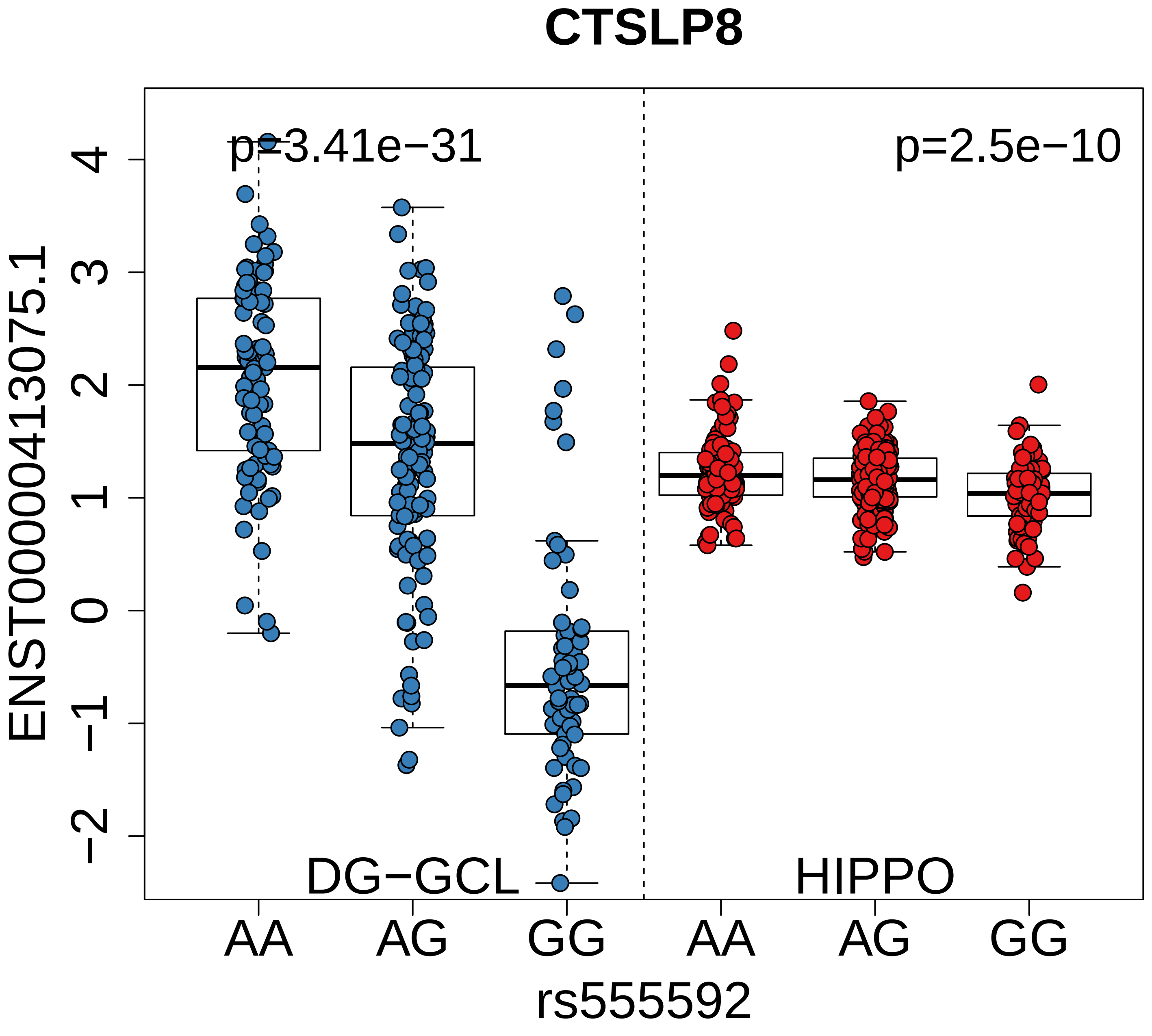
CTSLP8
p=3.41e−31
p=2.5e−10
ENST00000413075.1
4
3
2
1
0
−1
−2
DG−GCL
HIPPO
AA
AG
GG
AA
AG
GG
rs555592

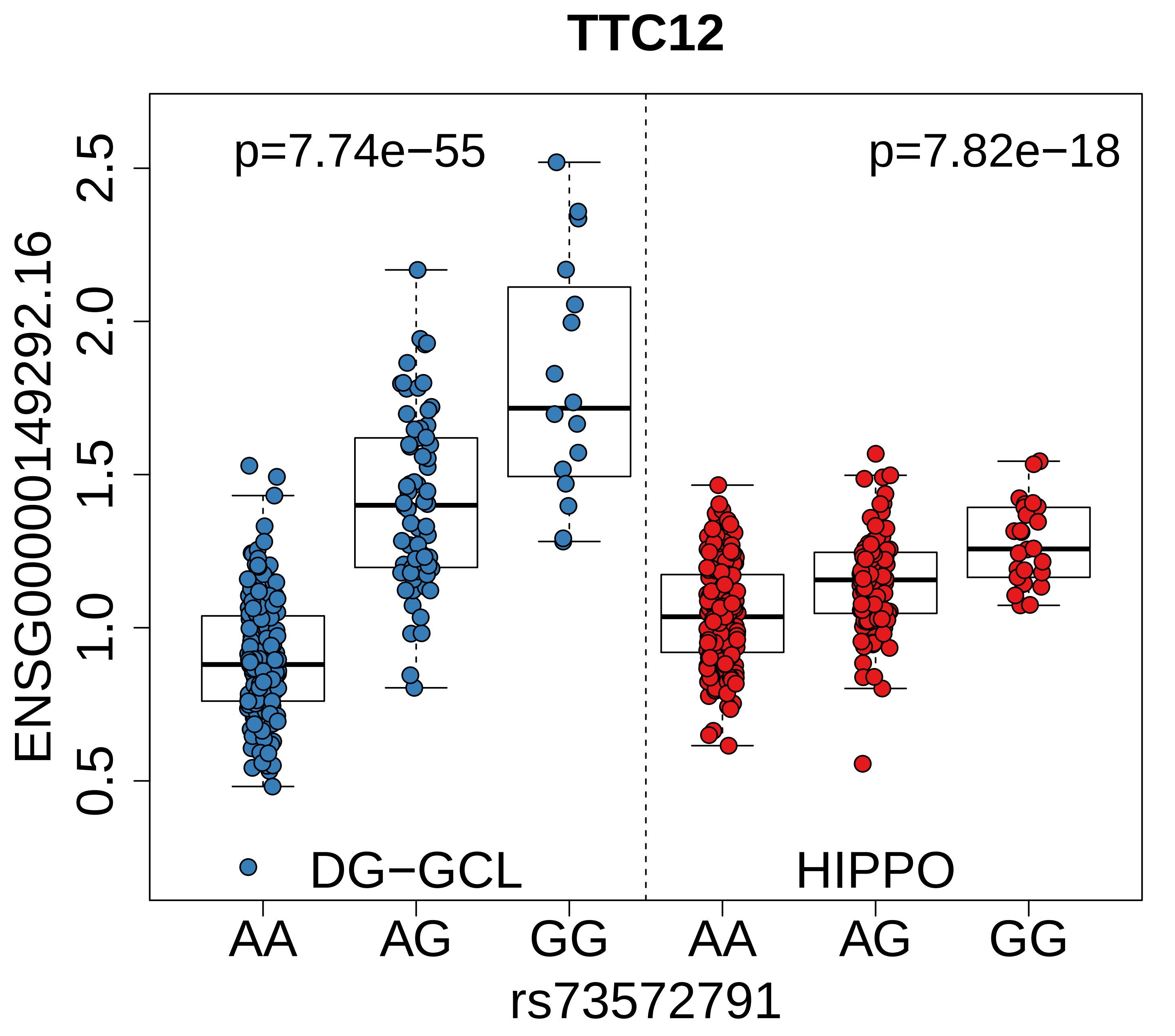

TTC12
p=7.74e−55
p=7.82e−18
ENSG00000149292.16
0.5
1.0
1.5
2.0
2.5
DG−GCL
HIPPO
AA
AG
GG
AA
AG
GG
rs73572791

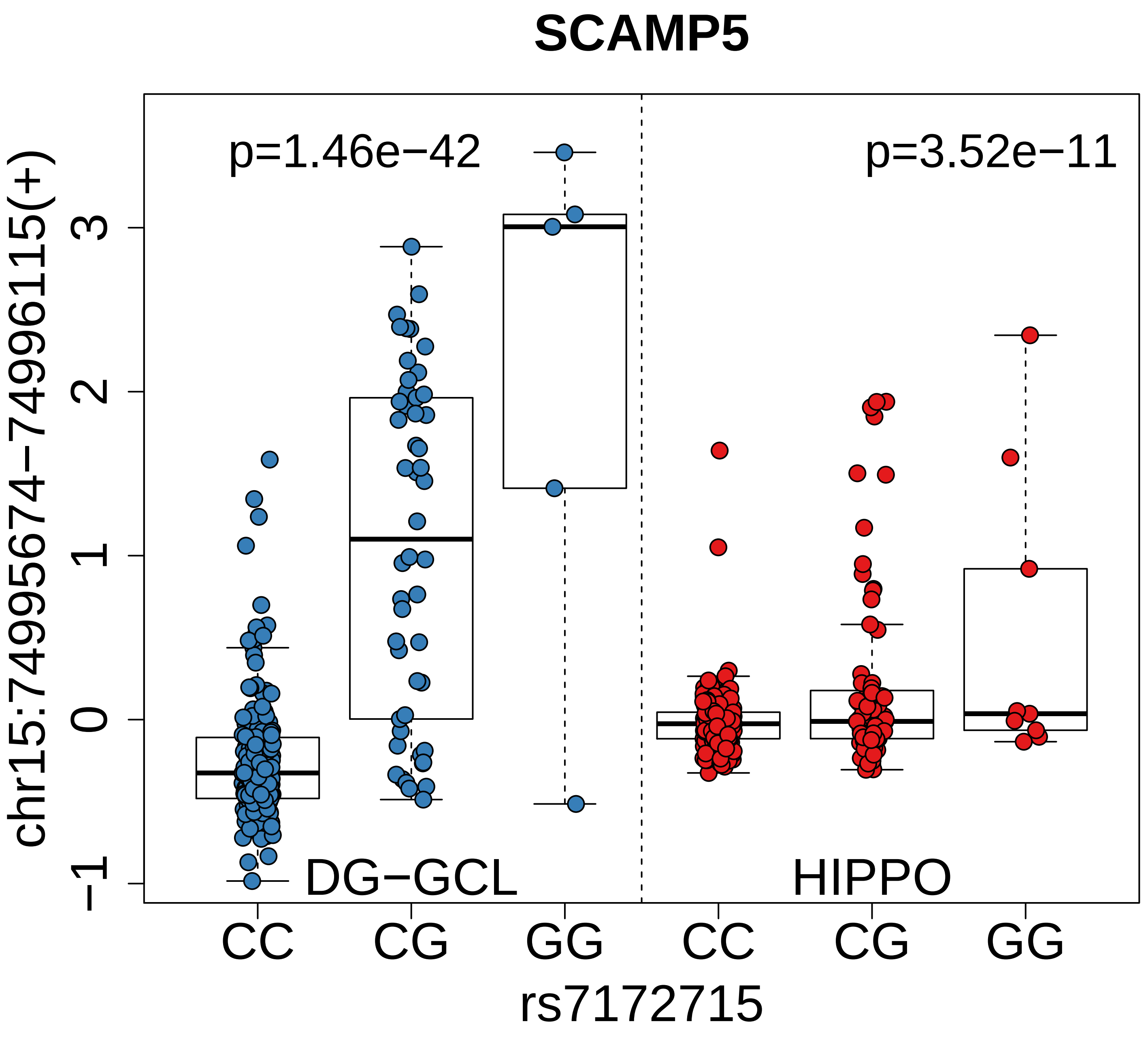

SCAMP5
p=1.46e−42
p=3.52e−11
chr15:74995674−74996115(+)
3
2
1
0
−1
DG−GCL
HIPPO
CC
CG
GG
CC
CG
GG
rs7172715

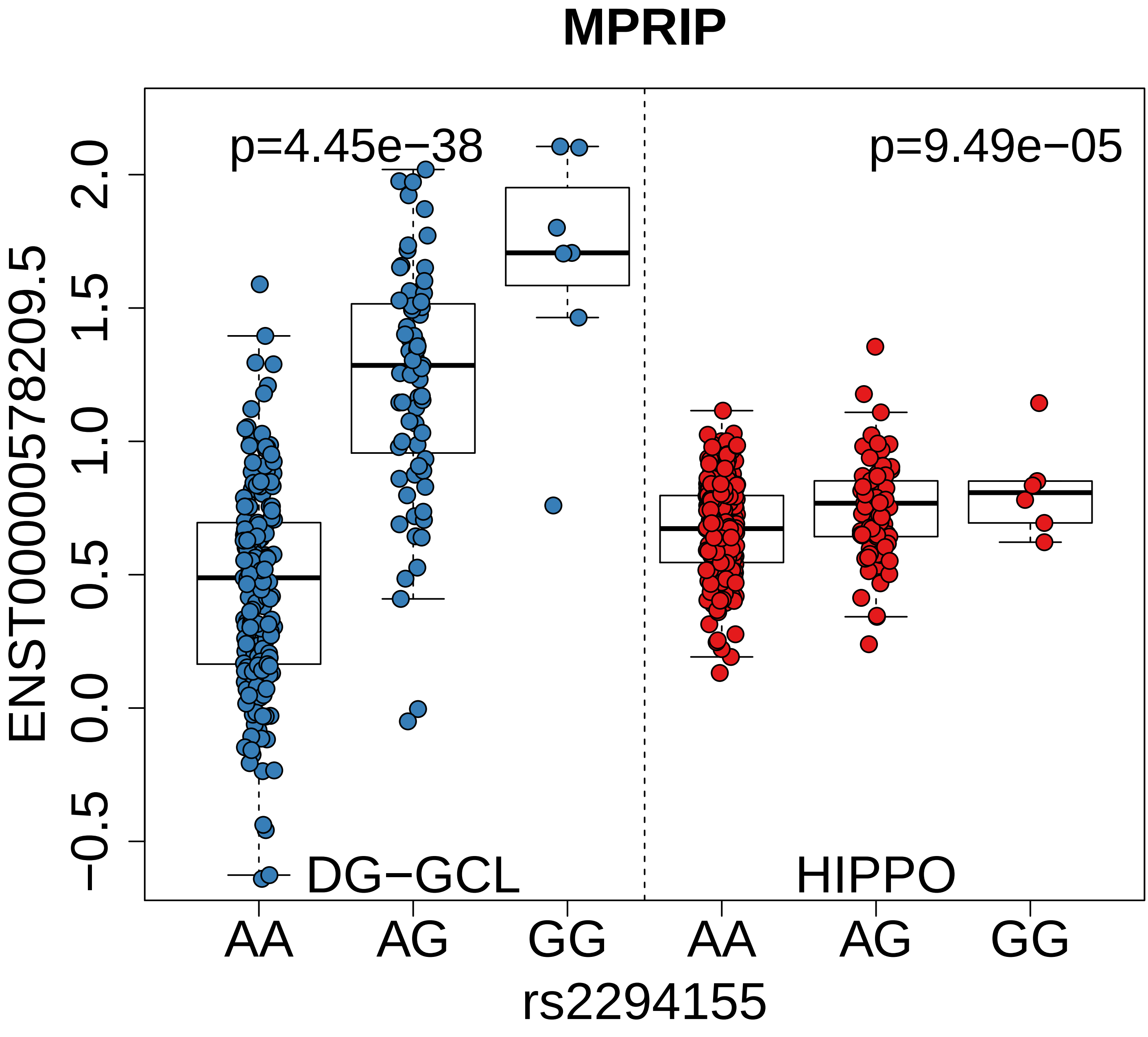
MPRIP
p=4.45e−38
p=9.49e−05
ENST00000578209.5
2.0
1.5
1.0
0.5
0.0
−0.5
DG−GCL
HIPPO
AA
AG
GG
AA
AG
GG
rs2294155

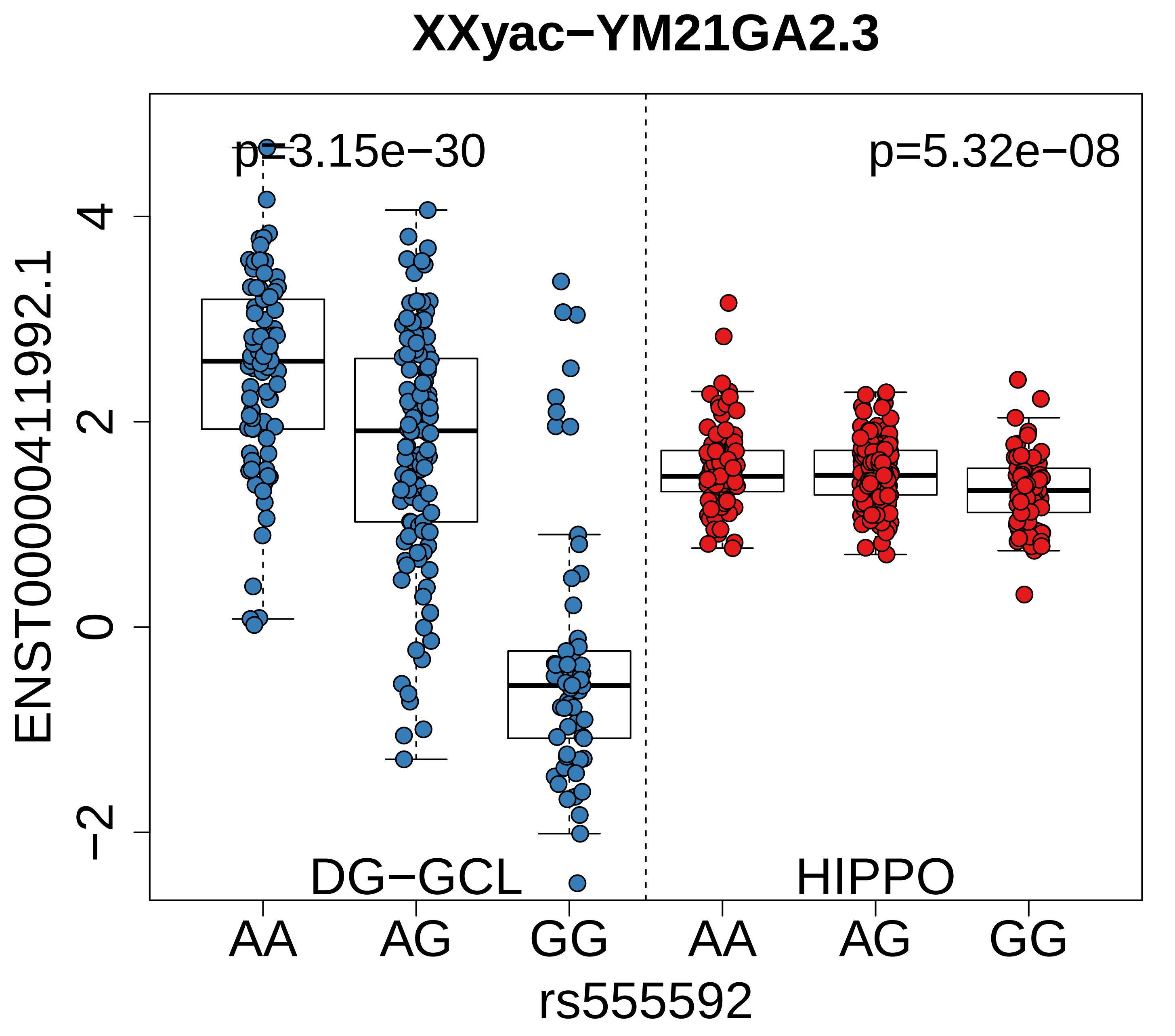

XXyac−YM21GA2.3
p=3.15e−30
p=5.32e−08
ENST00000411992.1
4
2
0
−2
DG−GCL
HIPPO
AA
AG
GG
AA
AG
GG
rs555592

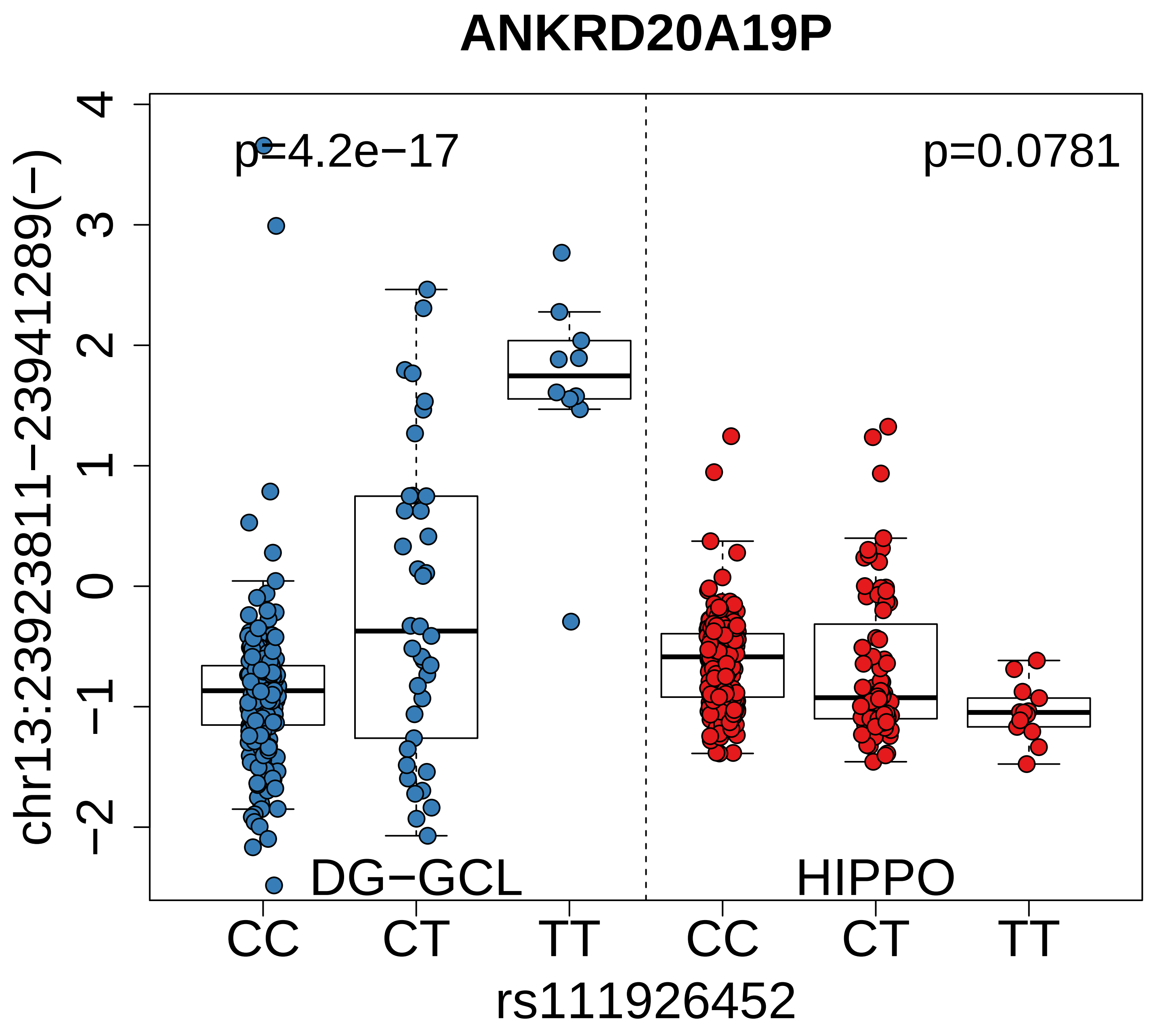
ANKRD20A19P
p=4.2e−17
p=0.0781
DG−GCL
HIPPO
CC
CT
TT
CC
CT
TT
rs111926452
chr13:23923811−23941289(−)
4
3
2
1
0
−1
−2

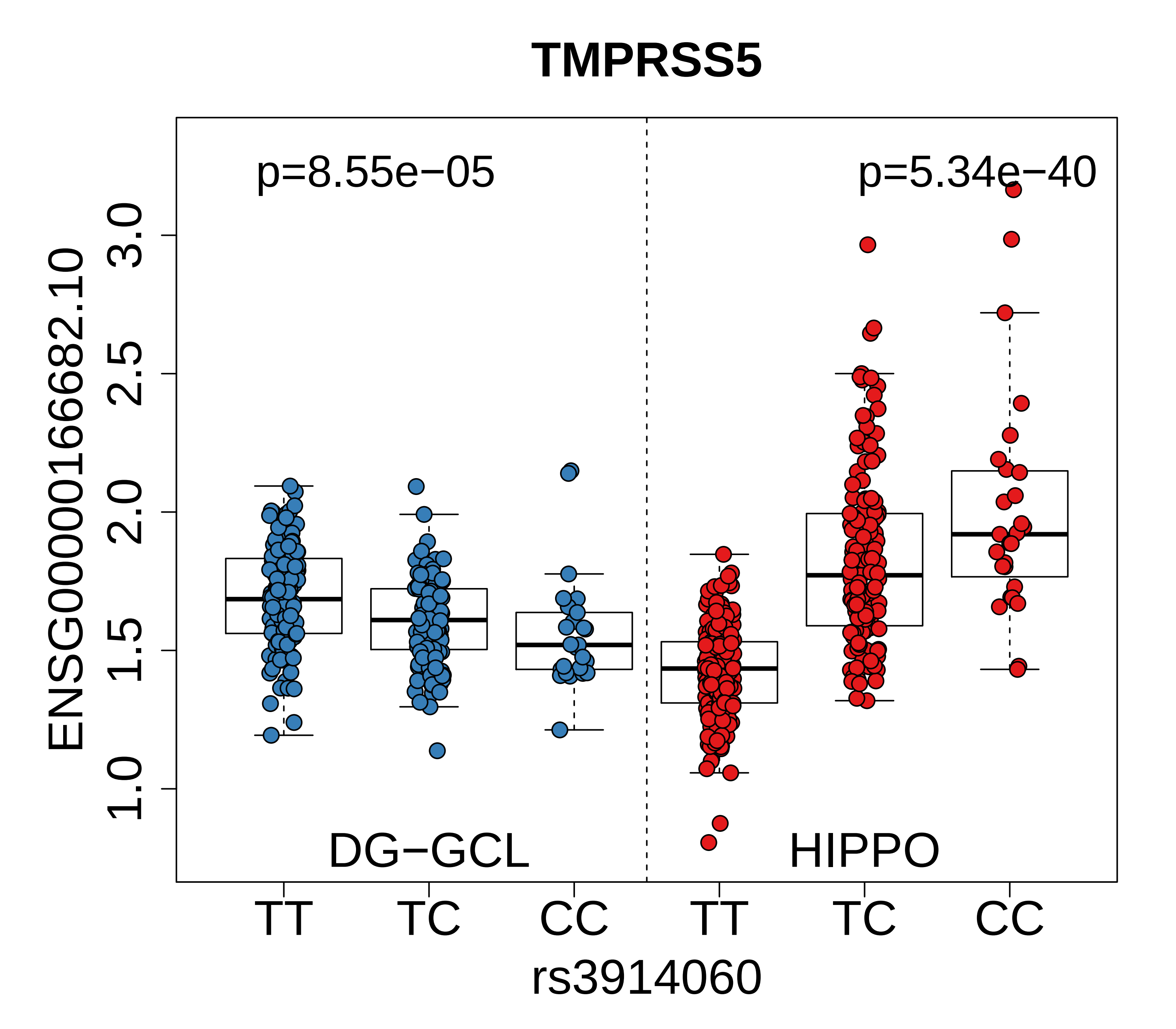
TMPRSS5
p=8.55e−05
p=5.34e−40
ENSG00000166682.10
3.0
2.5
2.0
1.5
1.0
DG−GCL
HIPPO
TT
TC
CC
TT
TC
CC
rs3914060

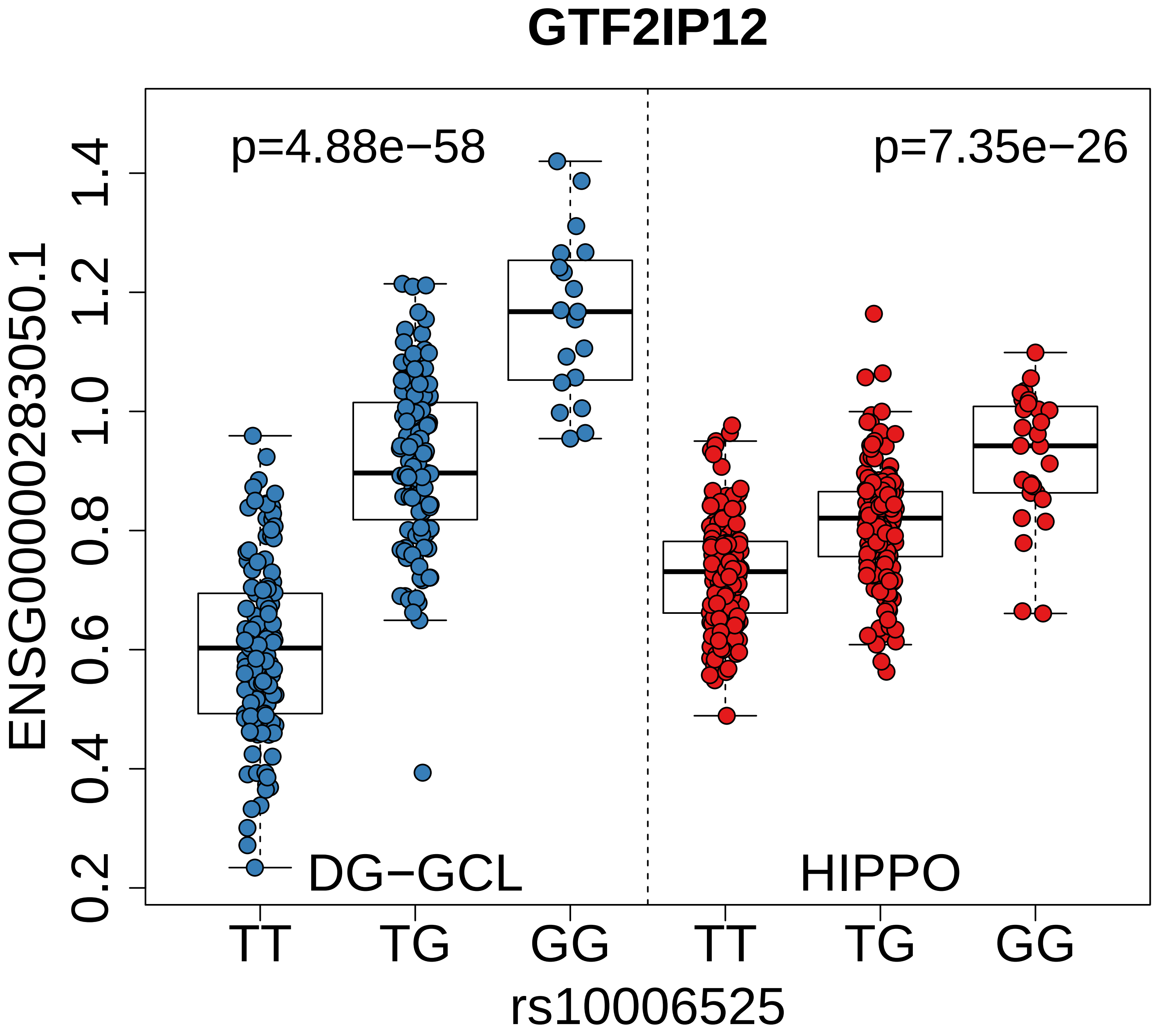
GTF2IP12
p=4.88e−58
p=7.35e−26
ENSG00000283050.1
0.2
0.4
0.6
0.8
1.0
1.2
1.4
DG−GCL
HIPPO
TT
TG
GG
TT
TG
GG
rs10006525

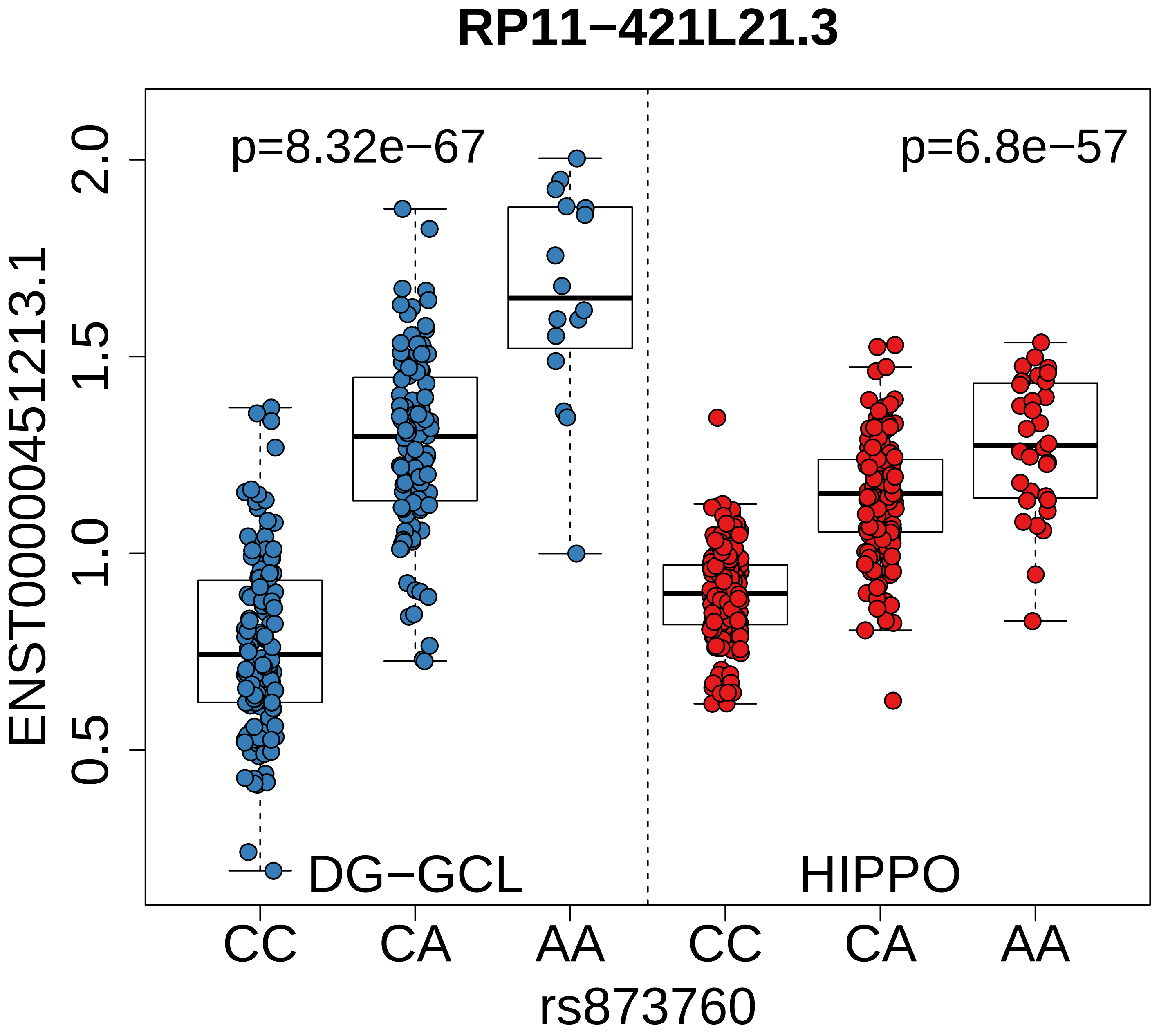

RP11−421L21.3
p=8.32e−67
p=6.8e−57
ENST00000451213.1
2.0
1.5
1.0
0.5
DG−GCL
HIPPO
CC
CA
AA
CC
CA
AA
rs873760

### NEK4P2

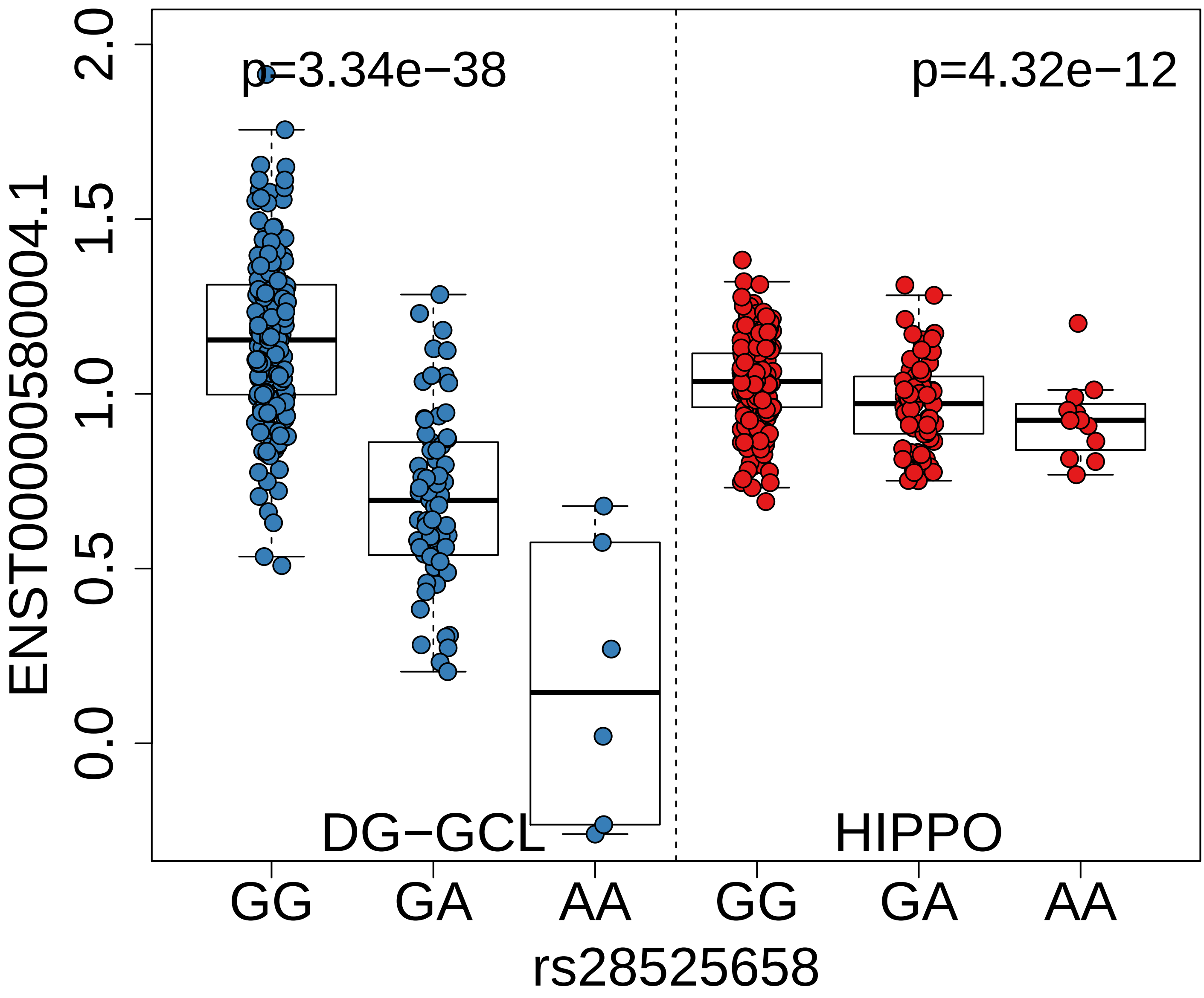

p=3.34e−38
p=4.32e−12
ENST00000580004.1
2.0
1.5
1.0
0.5
0.0
DG−GCL
HIPPO
GG
GA
AA
GG
GA
AA
rs28525658

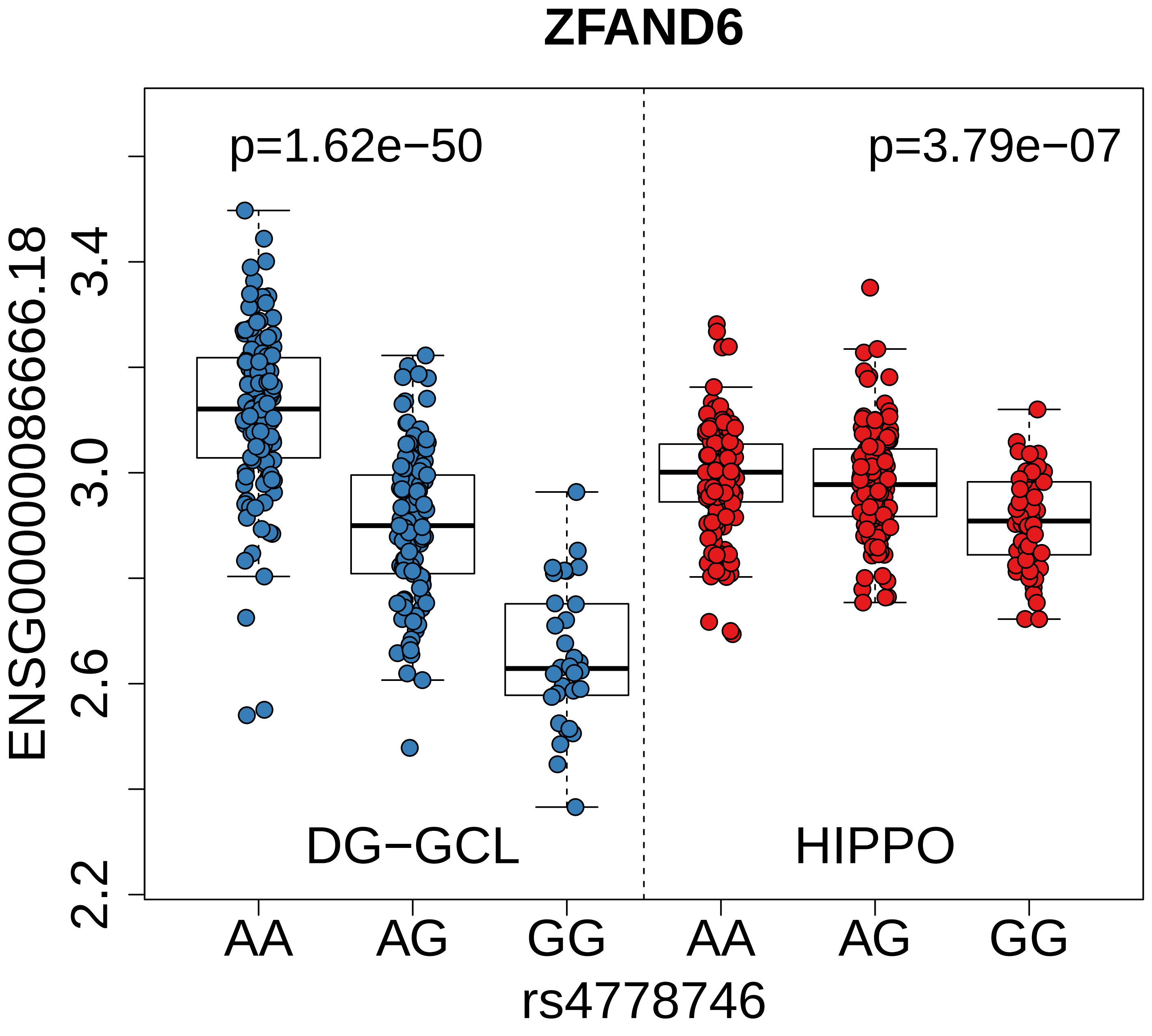

ZFAND6
p=1.62e−50
p=3.79e−07
ENSG00000086666.18
3.4
3.0
2.6
2.2
DG−GCL
HIPPO
AA
AG
GG
AA
AG
GG
rs4778746

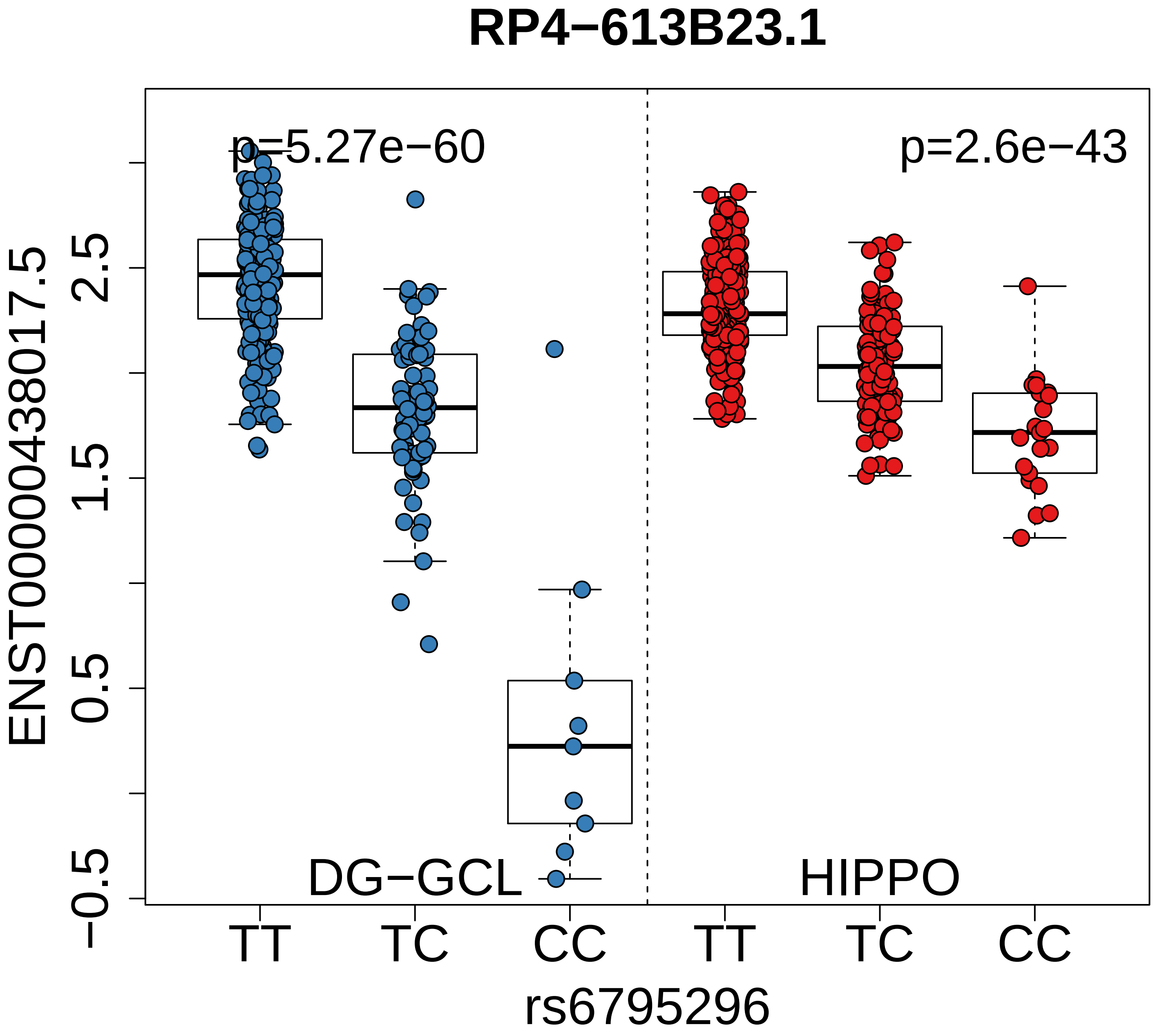

RP4−613B23.1
p=5.27e−60
p=2.6e−43
ENST00000438017.5
2.5
1.5
0.5
−0.5
DG−GCL
HIPPO
TT
TC
CC
TT
TC
CC
rs6795296
