## Supplementary figures and images for "Cell type-specific genetic regulation of expression in the granule cell layer of the human dentate gyrus"

### suppData1_ageExamples.pdf

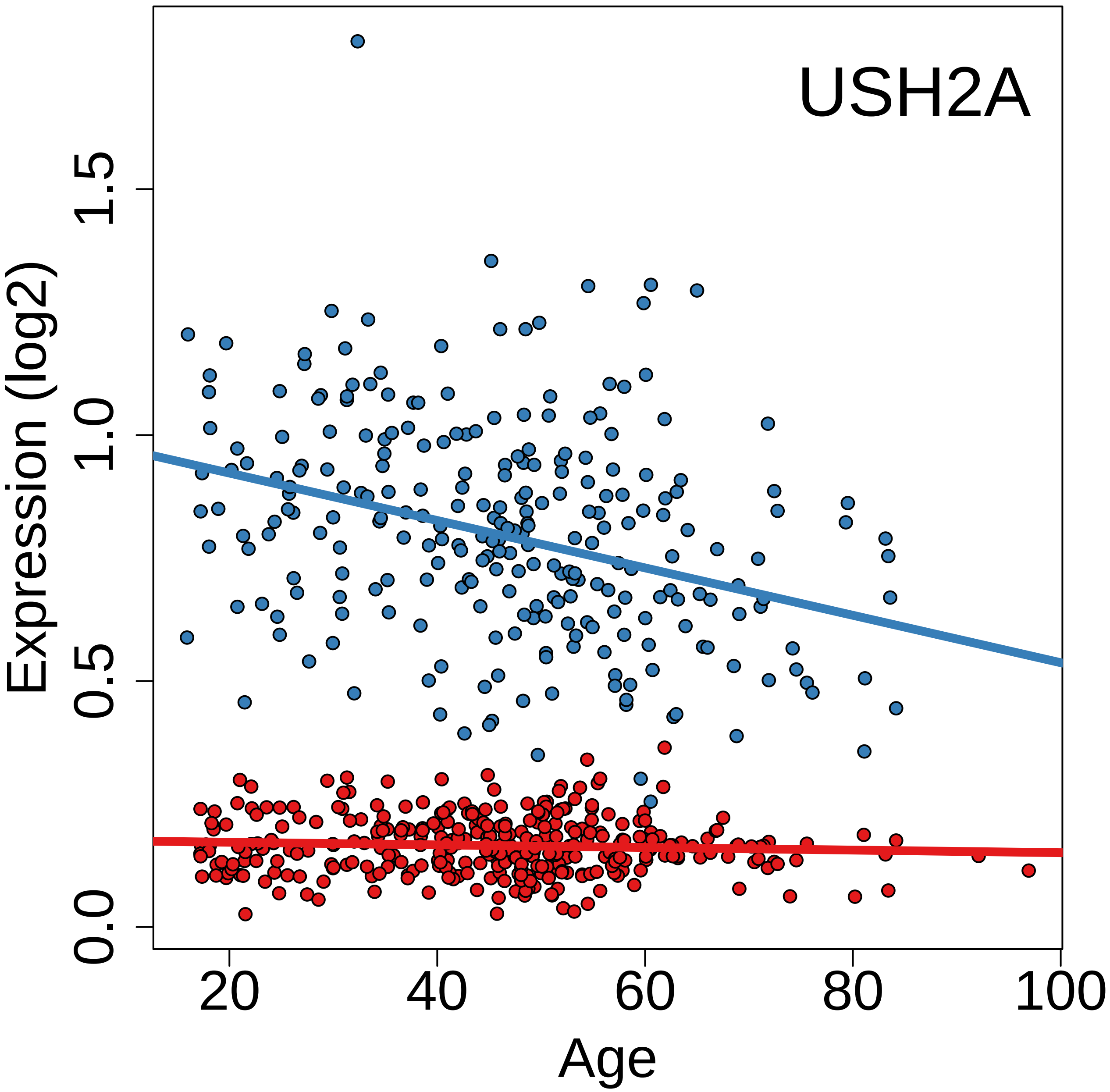

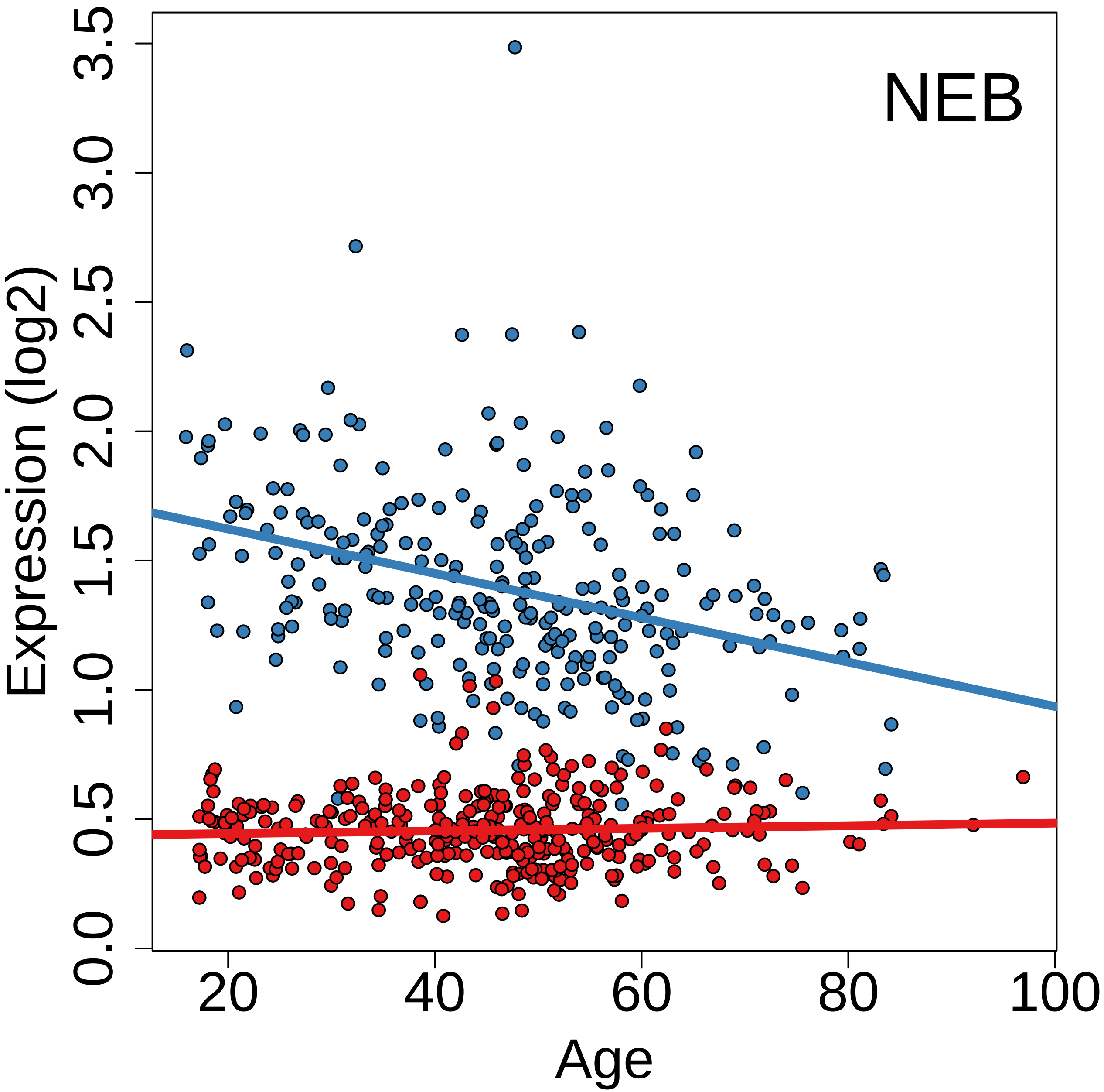

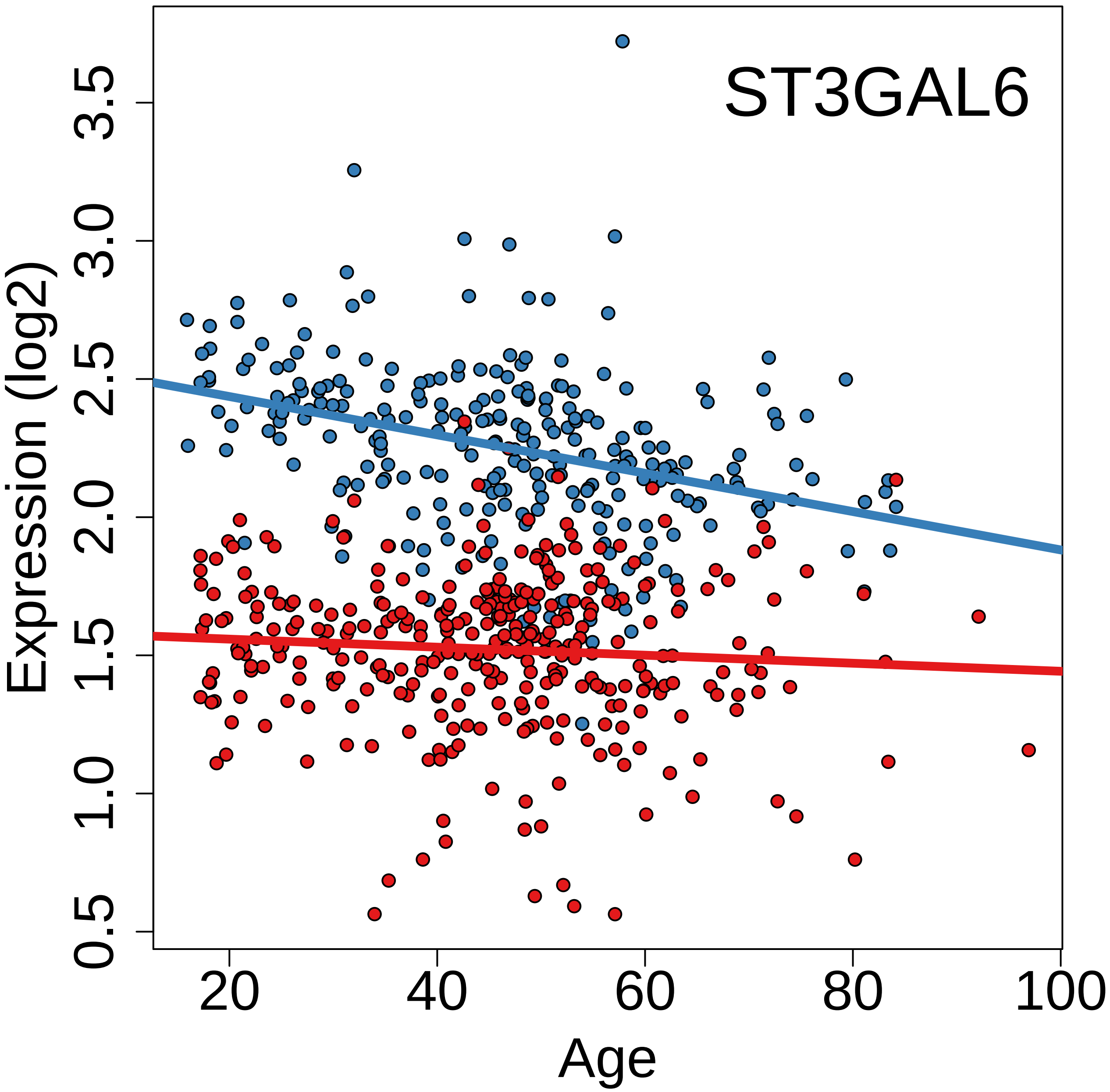

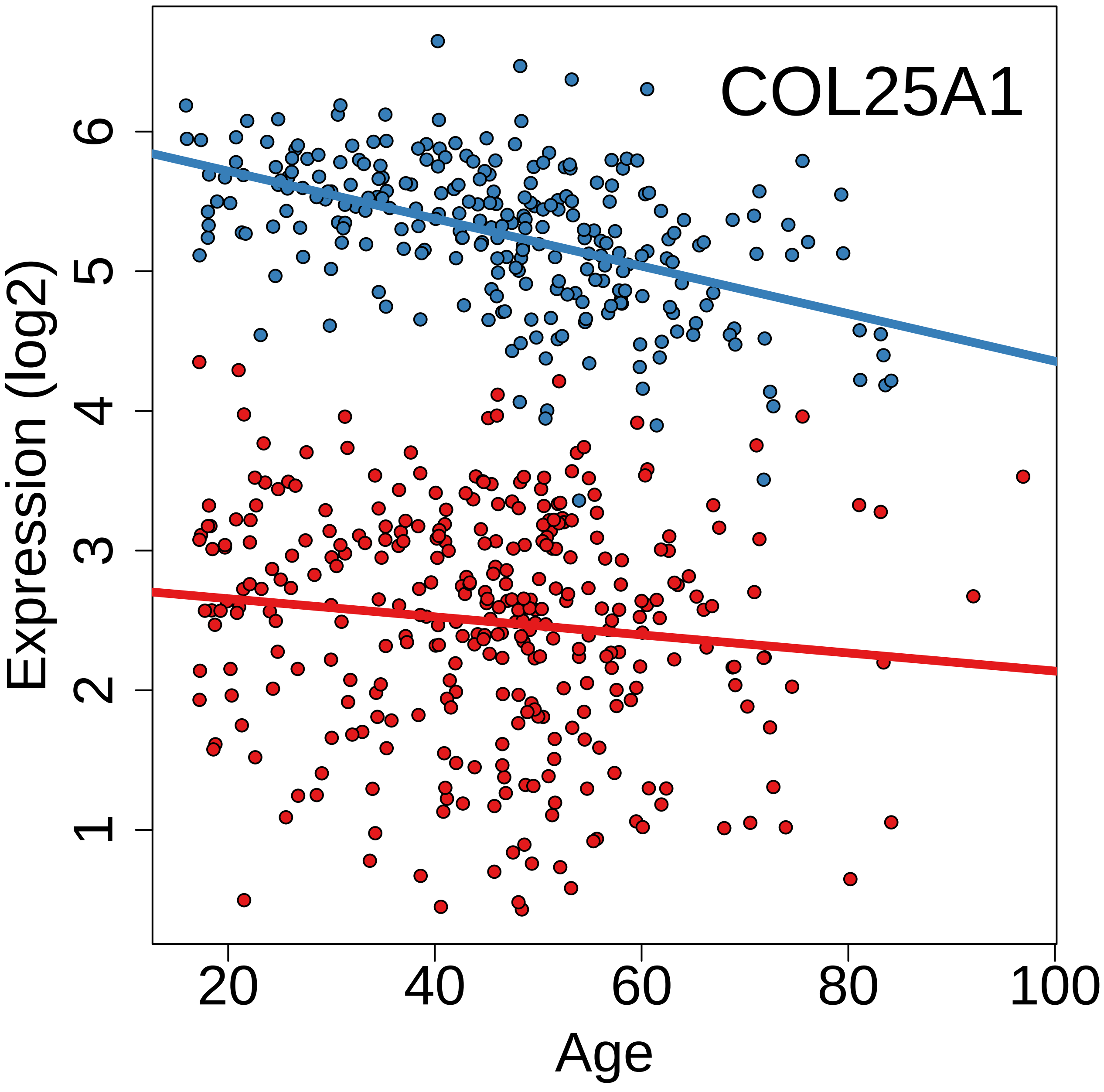

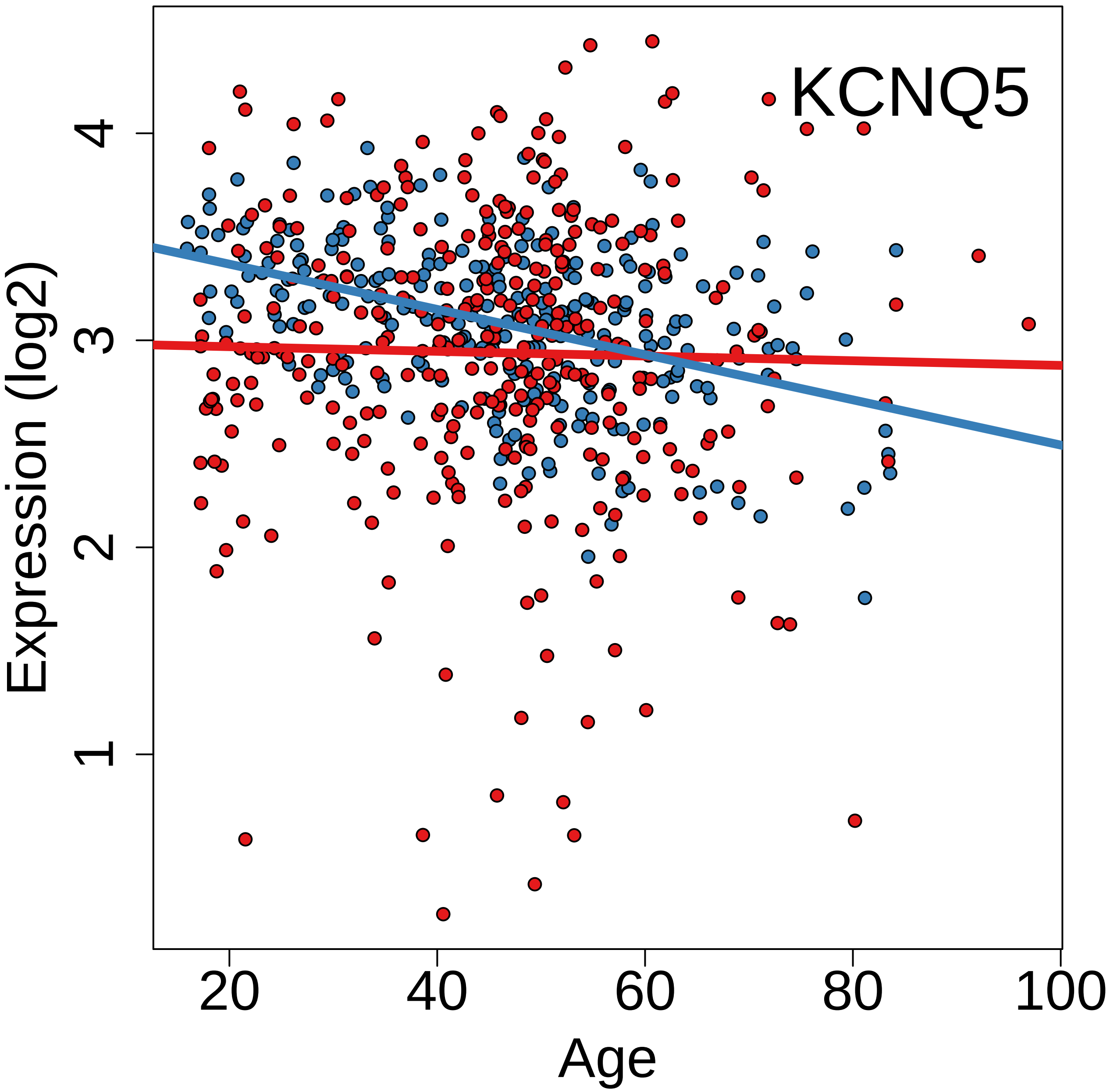

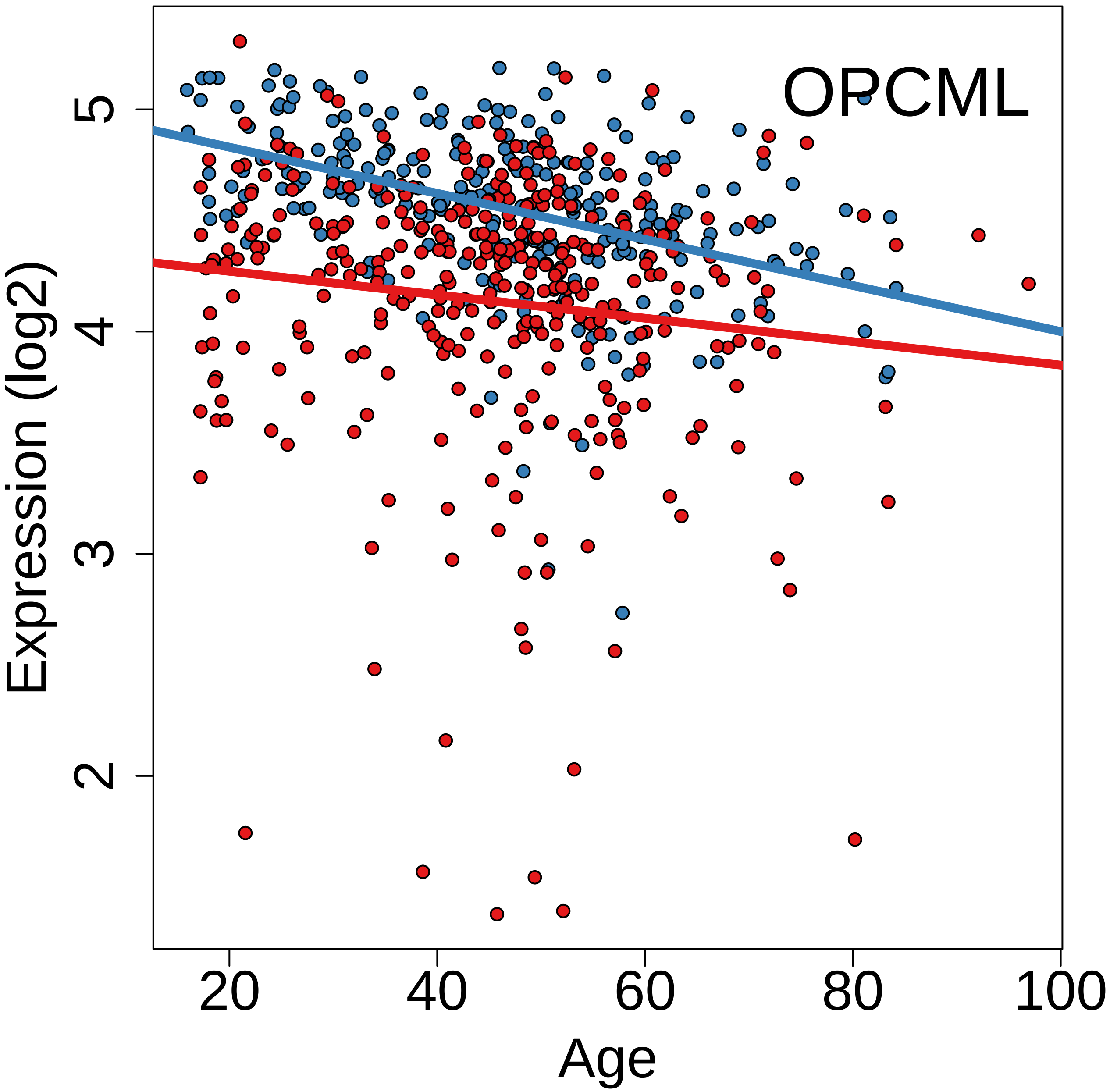

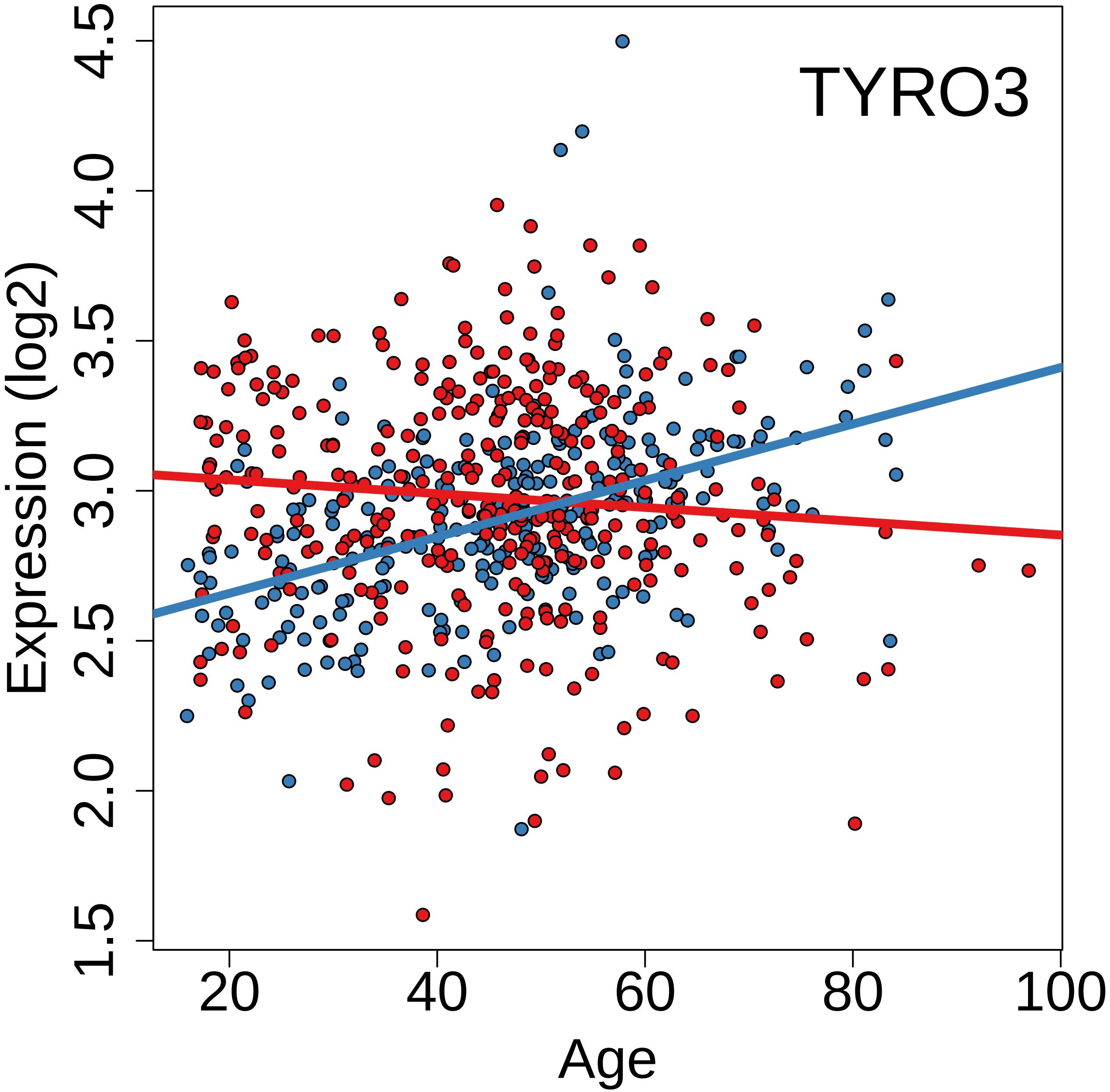

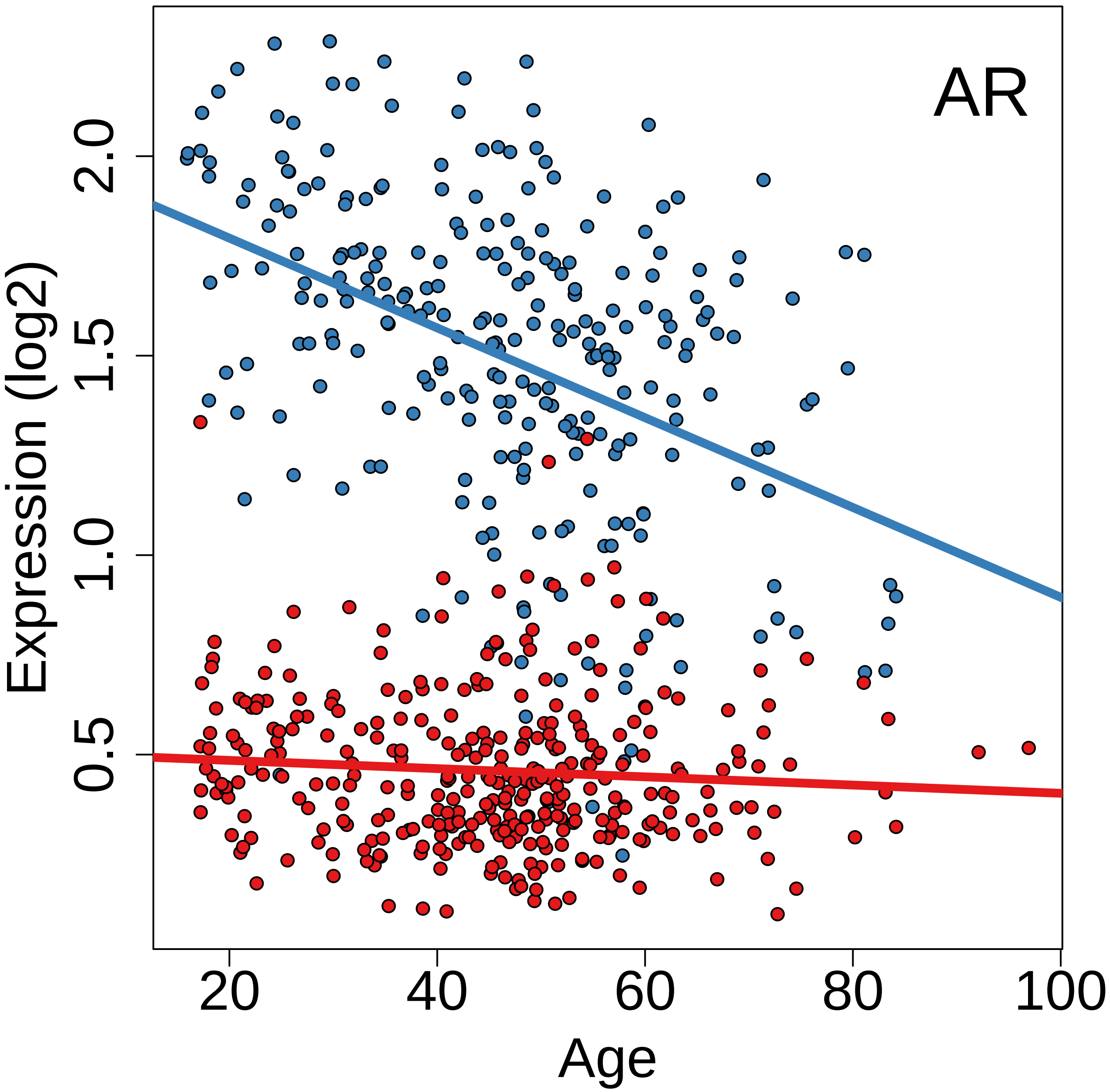

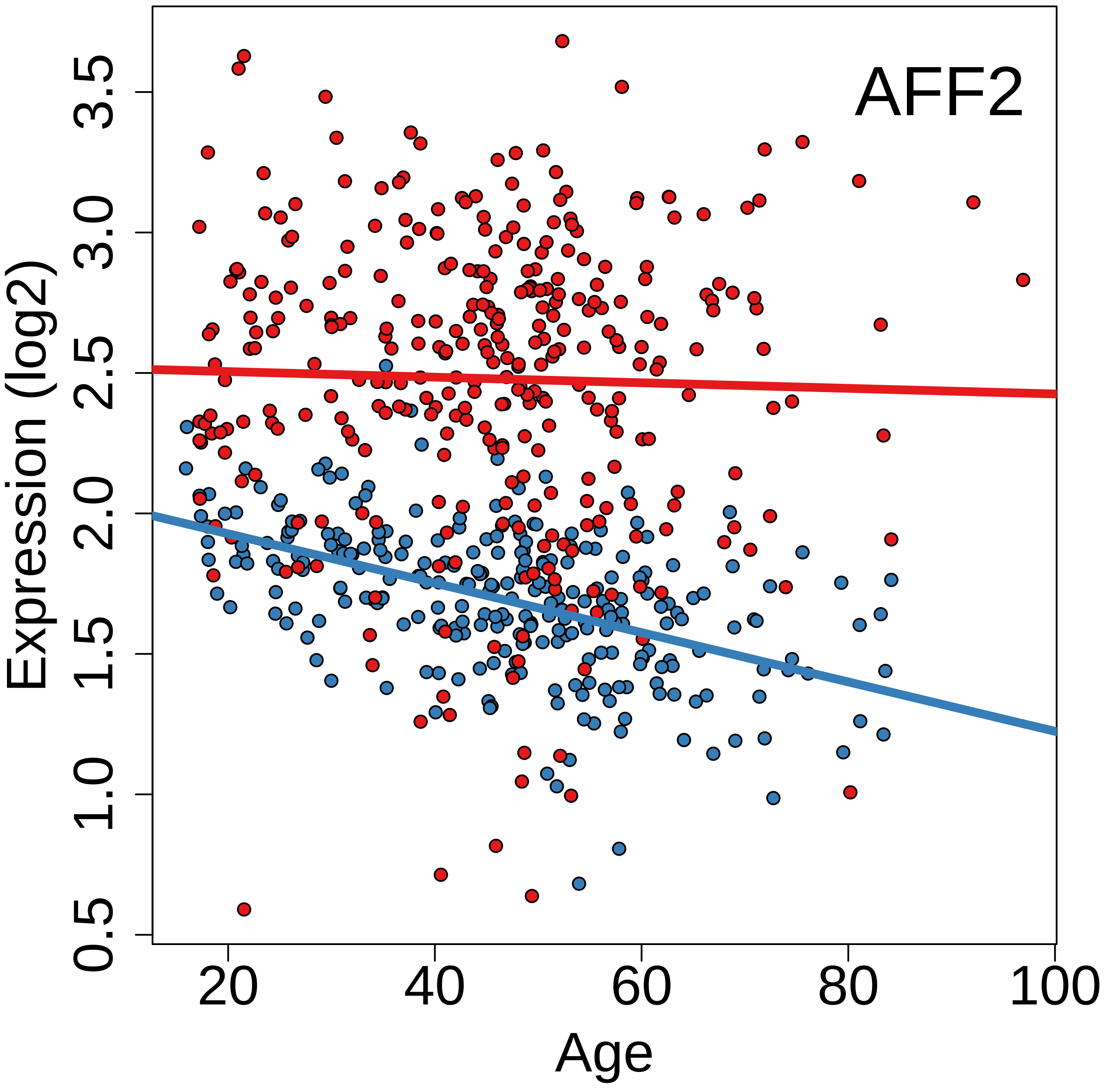
